# Plasma and Liver Lipids are Differentially Regulated After Cannabinoid Treatments in Male and Female Mice

**DOI:** 10.1101/2023.01.30.526249

**Authors:** Clare T Johnson, Taryn Bosquez-Berger, Hannah R Bentz, Gabriel H Dias de Abreu, Ken Mackie, Hui-Chen Lu, Heather B Bradshaw

## Abstract

Cannabinoids (CBs) have sex-dependent behavioral and physiological effects and modulate lipids across the body. To understand how some of these sex differences may be due to differences how both phyto- and endocannabinoids are regulated in the liver and plasma, male and female CD1 mice were administered 10 mg/kg *i.p*. of cannabidiol (CBD), Δ9-tetrahydrocannabinol (THC), or THC+CBD then core blood and liver tissue were collected after 2 hours. Lipids were extracted from both liver and plasma, and samples were screened via HPLC/MS/MS for Δ9-tetrahydrocannabinol (THC), metabolites 11-OH-THC and 11-COOH-THC, cannabidiol (CBD), metabolites 7-OH-CBD and 7-COOH-CBD, ~70 endolipids including the endocannabinoids, *N*-arachidonoyl ethanolamine (Anandamide; AEA) and 2-sn-arachidonoyl glycerol (2-AG). Structural analogs to AEA (*e.g*. lipoamino acids, lipoamines) and 2-acyl glycerols) as well as free fatty acids and prostaglandins were also evaluated. Results show that at 2 hours post injection levels of CBs demonstrate key differences between males and females in both plasma and liver, and that these differences vary when co-administered as opposed to administered alone. Illustrating a link between liver and plasma, directionality of CB differences are similar between the two tissue types as a function of both sex and CB treatment. By contrast, endolipids had very different profiles as a function of sex, CB administration, and tissue type. Importantly, there are baseline differences between male and female mice in endocannabinoids and related lipids, which likely impact how CB administration modulates these endolipids. These data illustrate the complexity of outcomes of CB treatment between males and females on circulating CBs and endolipids and highlight the need to consider these factors when evaluating efficacy of CB drug treatments or usage.

## 1. Introduction

Multiple lines of evidence have established sex differences in cannabinoid (CB) behavioral and molecular pharmacology ^1–5^. Δ9-tetrahydrocannabinol (THC) and cannabidiol (CBD) have different time and dose-response courses for male and female rats on mechanical pain (von Frey test), and temperature ^6^. Sex differences in cognitive function have also been observed; for example, in adult male mice, loss of CB1 receptors is associated with a significant increase in anxiety-like behaviors on the elevated plus maze, but the effect in females is not significant ^7^. By contrast, acute, low doses of inhaled THC (25mg/mL over 30 min, drug formulation provided by NIDA) lead to significant increases in anxiety-like behavior and decreases in locomotion in female adolescent rats, but not males. However, at a high dose of inhaled THC (100mg/mL over 30 min), males demonstrated anxiety-like behaviors whereas females show no significant effects ^8^. CBD also has time- and sex-dependent effects in rats, wherein male Swiss mice show a decrease in immobility on the tail suspension test after an acute injection (3, 10, 30 mg/kg), but females are not affected significantly by the same treatment ^9^. These behavioral differences support a need for deeper understanding of how CB are regulating signaling systems differently in male and female animals.

One explanation for these sex differences may stem from the fact that CBs are metabolized differently in male and female animals and that these metabolites drive different signaling efficacy. Adult female rats have higher peak plasma levels of 11-OH-THC after acute injection with 5 mg/kg of THC compared to males ^1^, and adult female-derived microsomes convert THC to 11-OH-THC at 3 times the rate of male microsomes, while conversion to 11-COOH-THC is similar between sexes ^1^. After a 2.5 mg/kg injection of THC, female rats have higher plasma levels of 11-OH-THC and 11-COOH-THC than males, but males have higher levels of THC ^10^. In humans, after vapor inhalation of 20 or 25 mg THC, levels of the THC metabolite 11-OH-THC show a significantly higher change from baseline in females than males ^11^. This emerging picture of how THC is being metabolized differentially in males and females provides an important framework to understand how THC may be driving differences in sex differences in responses to THC; however, the data on potential sex differences in CBD is much less developed.

Another hypothesis for sex differences in pharmacological responses to CBs is that if males and females have underlying differences in their endogenous CB (eCB) signaling systems which, in turn, are differentially affected by CB pharmacology, then this will drive differences in behavioral/therapeutic outcomes. In support of this hypothesis, evidence shows that differences exist in the distribution of CB receptors, with male rats having significantly more CB1 mRNA in the anterior pituitary than females, though levels in females change across the cycle ^12^. Sex differences also exist in eCB ligand regulation, in that females have significantly more of the eCB ligand, *N*-arachidonoyl ethanolamine (AEA), in the pituitary and hypothalamus than males,^12^ and levels of eCBs and related lipids change across the cycle throughout the CNS ^13^. Removal of the gonads decreases CB1 mRNA in the pituitary of males, as does repeated THC administration ^12^. Support that these different systems are differentially affected by CBs is also seen in that chronic THC administration leads to a significant decrease in CB1 density and sensitization in the prefrontal cortex, hippocampus, cerebellum, and striatum in both males and females, yet females show a greater effect size ^3^. THC has broader effects than signaling through CB1 and has been shown to be active at GPR18 ^14,15^, TRPA1 ^16^, and GPR55 ^17^, and to cause broad-scale changes in the CNS lipidome ^18^. CBD’s direct receptor signaling is less understood; however, it too causes broad-scale changes in the CNS lipidome ^19^.

Endogenous lipids (endolipids-different from exogenous CB lipids and their metabolites) are an important part of the eCB system and include endolipids like AEA, 2-arachidonoyl glycerol (2-AG) and their structural congeners (*see supplemental data for a list of over 60 of these*). We have shown that modulation of the eCB system, such as deletion of eCB biosynthetic enzyme NAPE-PLD ^20^, metabolic enzymes FAAH and MAGL ^21^, deletion of the cannabinoid receptor CB1 ^21^, or administration of THC or CBD results in broad changes in signaling lipids in the brain ^18,19^. Acute, low dose THC administration (3 mg/kg) in female mice results in regio-specific decreases in *N*-acyl ethanolamines (NAEs) ^18^, whereas acute CBD administration (3 mg/kg) in female mice results in regio-specific increases in NAEs the brain ^19^. eCB ligands and related endolipids are not limited to the brain, and from a translational standpoint measuring endolipids in biological specimens that are less invasive to sample, such as plasma, could be particularly informative. Already, we have shown sex differences in endolipids in plasma of patients with gastroparesis ^22^. Likewise, serum collected from cannabis users shows increased palmitic acid, oleic acid, 2-AG, and PEA, but not AEA or OEA compared to non-cannabis users ^23^, which illustrates that plasma endolipids are responsive to changes in systemic CB pharmacology. It will be informative to compare plasma levels of endolipids with CB treatments to other tissues in order to determine how translatable these values are to other parts of the body.

Here, we hypothesize that if underlying sex differences exist in metabolic pathways utilized by CBs, then CBs and their metabolites will be differentially present in liver and plasma after an acute administration of THC, CBD or THC+CBD as a function of genetic sex. We further hypothesize that if there are sex differences in the metabolism and distribution of CBs and their, then plasma and liver endolipids will be significantly and differentially altered by CB treatment as a function of genetic sex.

## 2. Methods

### 2.1 Animals

Adult male and female CD1 mice (6-20 weeks old) were housed with a 12 hr light dark cycle and cared for in accordance with Indiana Care and Use Committee requirements. They were randomly assigned to receive a single i.p. injection of either THC (10 mg/kg; n_male_= 10; n_female_= 10), CBD (10 mg/kg; n_male_=11; n_female_= 9), THC+CBD (10 mg/kg; n_male_=11; n_female_= 11), or vehicle (ethanol; n_male_=8; n_female_= 8). 2 hours after injection, animals were sacrificed via decapitation, livers dissected, and trunk blood collected in heparin-containing tubes on ice that were then centrifuged for 5 min. The upper fraction of plasma was then collected and stored at −80°C until samples were processed. 3mm punches were taken from the medial lobe of the liver closest to the gallbladder to be used for lipid measurement and also stored at −80°C until processing.

### 2.2 Lipid Extraction

On the day of processing, plasma samples were thawed for 5 min at 37° C, vortexed, and between 25-75μL of each sample transferred to a tube for lipid extraction. Methanol was added to each sample for a final volume of 2 mL. 5 μL of 1 μM deuterium labeled anandamide (d8AEA) was used as an internal standard and solutions were vortexed. Samples were then incubated in the dark on ice for 15 min before being centrifuged at 19,000G, 20° C, for 20 min. Supernatants were added to 6 mL of HPLC-grade water to make a 75% organic solution. Lipids were partially purified from this mixture using C-18 solid phase extraction columns (Agilent), eluting with 65%, 75%, and 100% methanol (1.5 mL). On the day of processing liver samples, ~50 vol of methanol were added to each sample in addition to 5 μL of 1 μM d8AEA. Samples were incubated in the dark on ice for 2h, then homogenized via sonication, and centrifuged at 19,000G, 20° C, for 20 min and partially purified as the plasma samples.

### 2.3 Statistical Analysis

20 μL of each sample was assessed for related lipid species up to 6 lipids in one mass spectrometric analysis. Levels of individual lipids were determined based on concentration curves of known standards. Final moles/liter (plasma) and moles/gram (liver) calculations were adjusted based on the average percent recovery of the internal standard and amount of starting material. In plasma samples, 1 male from the THC group and 1 male from the THC+CBD group had concentrations of CBs greater than two standard deviations from the mean and data from all lipid analysis from these subjects were omitted. In liver samples, 1 female from the CBD group, 1 female from the THC+CBD group, 1 male from the CBD group, and 1 male from the THC group were outliers for CBs and excluded from subsequent analysis. Outliers in individual endolipids beyond two standard deviations of the mean were excluded in final analysis (*final sample sizes and all statistical measurements for each lipid analysis group are provided in Supplemental data*). Outlier exclusion was ~3% of all plasma data and 4% of all liver data. Data were analyzed with one-way ANOVAs with Fisher’s Least Significant Differences post-hoc (veh compared to CB treatment by sex) and independent t-tests (males versus female). Significance was set p ≤ 0.05, and trending significance as 0.05 < p ≤ 0.10 ^19,21^.

## 3. Results

### 3.1 Plasma and liver levels of cannabinoids and CB metabolites are sex and treatment dependent

Levels of CBD and THC were detectable in all plasma and liver samples in predicted groups (*i.e*. CBD in CBD or T+C treatment, THC in THC or T+C treatment) (Figures 1Ai-ii and 1i-iii). Likewise, the THC metabolites, (±)-11-Hydroxy-Δ^9^-THC (11-OH THC) and (+)-11-Nor-Δ^9^-THC-9-carboxylic acid (11-COOH THC) and the CBD metabolites, 7-hydroxy cannabidiol (7-OH-CBD) and 7-carboxy cannabidiol (7-COOH-CBD) were detected in all predicted samples with the exception that 7-COOH-CBD was not detected in plasma from males in the CBD alone treatment group, though one male in that treatment group had detectable levels in the liver (Fig. 1Aiii-vi, Fig. 1Eiii-vi). Stock solutions used to generate injections were used to generate standard curves for mass spectrometric analysis and THC and CBD were equimolar in the injection solutions. This information allowed for direct comparisons of the molarity of THC and CBD in plasma and liver at the 2-hour time point post injection. Therefore, if THC and CBD were transported and metabolized at the same rate in each treatment group, they would have equivalent amounts recovered across treatment groups.

**Figure 1.**
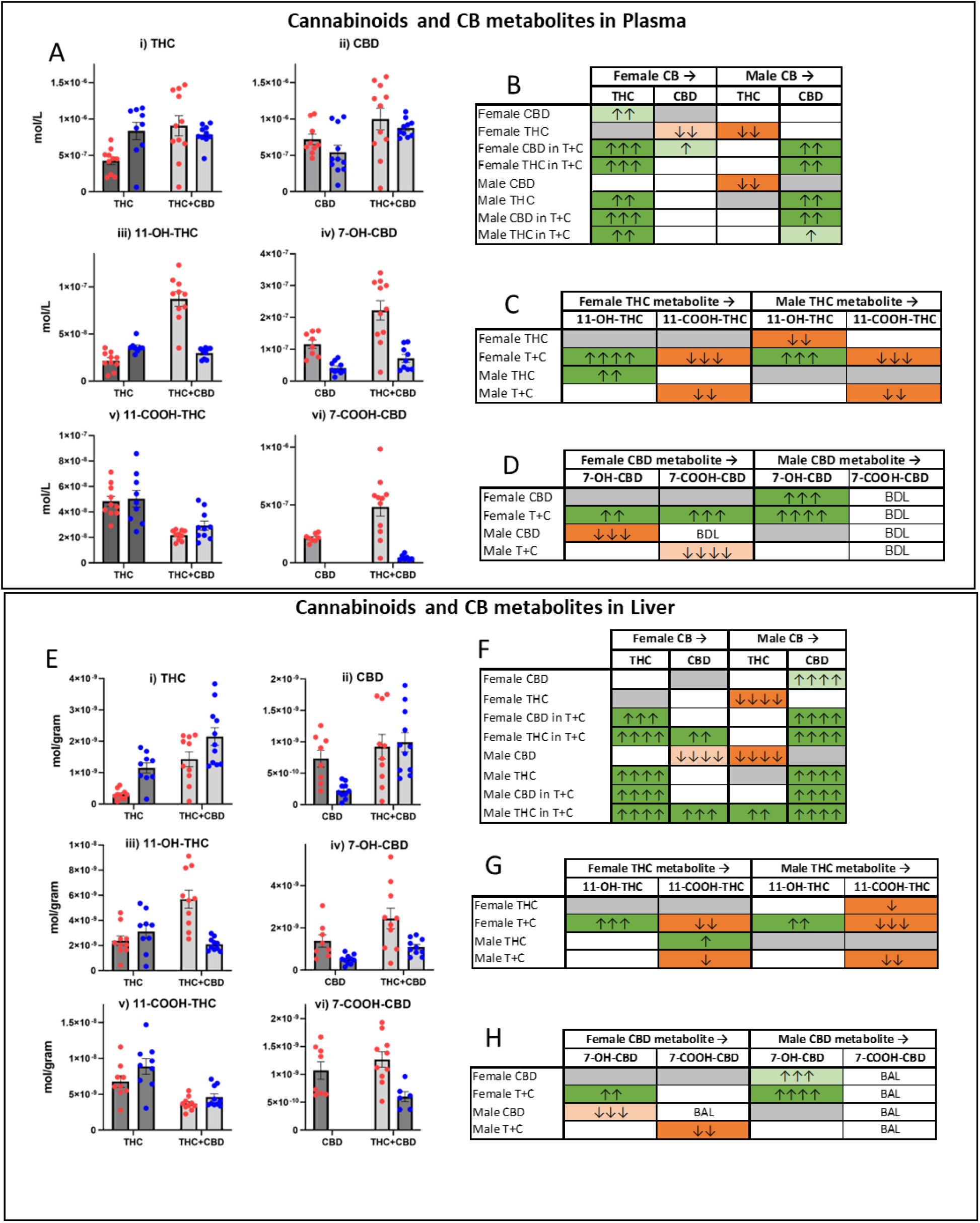
Levels of CBD, THC, and their metabolites in the plasma and liver of male and female mice 2hr post-i.p. injection of 10mg/kg CBD, THC, or THC+CBD. **A)** Mean ±SEM concentrations (mol/L) for each cannabinoid (CB) measured in plasma: i) THC, ii) CBD, iii) 11-hydroxy Δ^9^-tetrahydrocannabinol (11-OH-THC), iv) 7-hydroxy cannabidiol (7-OH-CBD), v) 11-nor-9-carboxy-Δ^9^-tetrahydrocannabinol (11-COOH-THC), vi) 7-carboxy cannabidiol (7-COOH-CBD). Individual measurements from each female (red) and male (blue) animal are represented by a corresponding dot. **B)** Heatmaps illustrating significant differences between the plasma levels of CBs measured in each CB treatment group. Data are represented as comparisons of levels of female THC alone treatment group compared to all other treatment groups (far left column); then female CBD alone treatment group compared to all other treatment groups (second to the left column). Data represented as comparisons of levels of male THC alone treatment group compared to all other treatment groups (third to the left column); then male CBD alone treatment group compared to all other treatment groups (far right column). **C)** Heatmaps illustrating differences in plasma THC metabolite levels in the THC-containing treatment groups relative to levels in the THC alone treatment group. **D)** Heatmaps illustrating differences in CBD metabolite levels in CB treatment group plasma relative to levels in the CBD alone treatment groups. **E)** Mean ±SEM concentrations (mol/g) for each cannabinoid (CB) measured in liver: i) THC, ii) CBD, iii) 11-hydroxy Δ^9^-tetrahydrocannabinol (11-OH-THC), iv) 7-hydroxy cannabidiol (7-OH-CBD), v) 11-nor-9-carboxy-Δ^9^-tetrahydrocannabinol (11-COOH-THC), vi) 7-carboxy cannabidiol (7-COOH-CBD). Individual measurements from each female (red) and male (blue) animal are represented by a corresponding dot. **F)** Heatmaps illustrating significant differences between the liver levels of CBs measured for each CB treatment group. Data are represented as comparisons of levels of female THC alone treatment group compared to all other treatment groups (far left column); then female CBD alone treatment group compared to all other treatment groups (second to the left column). Data represented as comparisons of levels of male THC alone treatment group compared to all other treatment groups (third to the left column); then male CBD alone treatment group compared to all other treatment groups (far right column). **G)** Heatmaps illustrating differences in liver THC metabolite levels in the THC-containing treatment groups relative to levels in the THC alone treatment group. **H)** Heatmaps illustrating differences in CBD metabolite levels in the liver of CB treatment groups relative to liver levels in the CBD alone treatment groups. Dark shades depict significant differences (p < 0.05), light shades depict trending differences (0.05 < p < 0.10). Green indicates significant increases, orange indicates significant decreases, arrows indicate effect size. *See Methods for description of heatmap analytics. See supplemental data for all statistical analyses*.

Figures 1Ai-ii, 1B, 1Ei-ii, and 1F show that in both the plasma and liver, females who received THC alone had significantly lower levels of THC than all other groups except for the level of CBD in the male CBD alone group (plasma) or both the male and female CBD alone group (liver). Similarly for both plasma and liver, males who received CBD alone had significantly lower CBD levels than all other groups except for THC levels in females who received THC alone (liver) or females in the THC alone and CBD alone groups (plasma). Overall, levels in CBD females tended to be higher than in CBD males.

CB metabolites demonstrated even greater variability among groups. Levels of 11-OH-THC (Fig. 1Aiii, C, 1Eiii, G) were significantly higher in females in the T+C group than all other groups in both plasma and liver; however, female plasma levels of 11-OH-THC in the THC group were significantly lower than THC males. 7-OH-CBD levels (Fig. 1iv, D) in T+C females were, likewise, significantly higher than all other groups and levels in CBD females were higher than in CBD males. Liver 7-OH-CBD levels (Fig. 1Eiv, H) in T+C females were, likewise, significantly higher than all other groups. Plasma 11-COOH-THC levels (Fig. 1v, C) showed no sex differences within treatment groups; however, levels in the T+C groups were significantly lower than THC alone. Liver levels of 11-COOH-THC (Fig. 1Ev, G) were lowest in the male T+C group, though males in the THC alone group had significantly higher levels than females in the THC alone group. In both plasma and liver, levels of 7-COOH CBD (Fig. 1Avi, D, 1Evi, H) showed the greatest sex differences, wherein levels were either below detectable levels (plasma) or below analytical levels (liver) in CBD males and significantly lower in T+C males.

### 3.2 Directional changes in overall levels of endogenous plasma and liver lipids are sex and CB treatment dependent

Of the 65 endolipids evaluated in plasma (*see supplemental data Table 1 for complete list and Fig. 2.1 for heatmap analysis*), 44 were present at levels high enough for detection and analysis in female and male plasma samples (*see supplemental data Figs. 1.2, 1.3*). Additional endolipids in females (16 in females, 17 in males) were detected in only a few plasma samples, so these were not statistically evaluated and are denoted as BAL (*see Supplemental Figs. 1.2, 1.3*). 6 endolipids were not present at detectable limits in female samples and are denoted at BDL, while 5 were not present at detectable limits in male samples. Likewise, of the 86 endolipids evaluated in the liver, 74 were present at levels high enough for detection and analysis in all treatment groups in females and 71 in males; 11 were present but BAL in males, 10 were present but BAL in females; 2 were BDL in females, and 4 were BDL in males. Figure 2 illustrates a summary of the overall sex difference analyses in which male endolipid levels in either plasma or liver are compared to female levels in the same tissue (females acting as the “baseline”; Female → Male), endolipid levels in each CB treatment are compared to vehicle (Veh→ CB), endolipid levels in CB treatments (THC or THC:CBD; T+C) are compared to levels in CBD alone treated animals (CBD → CB), or levels in T+C are compared to THC alone treated animals.

**Figure 2.**
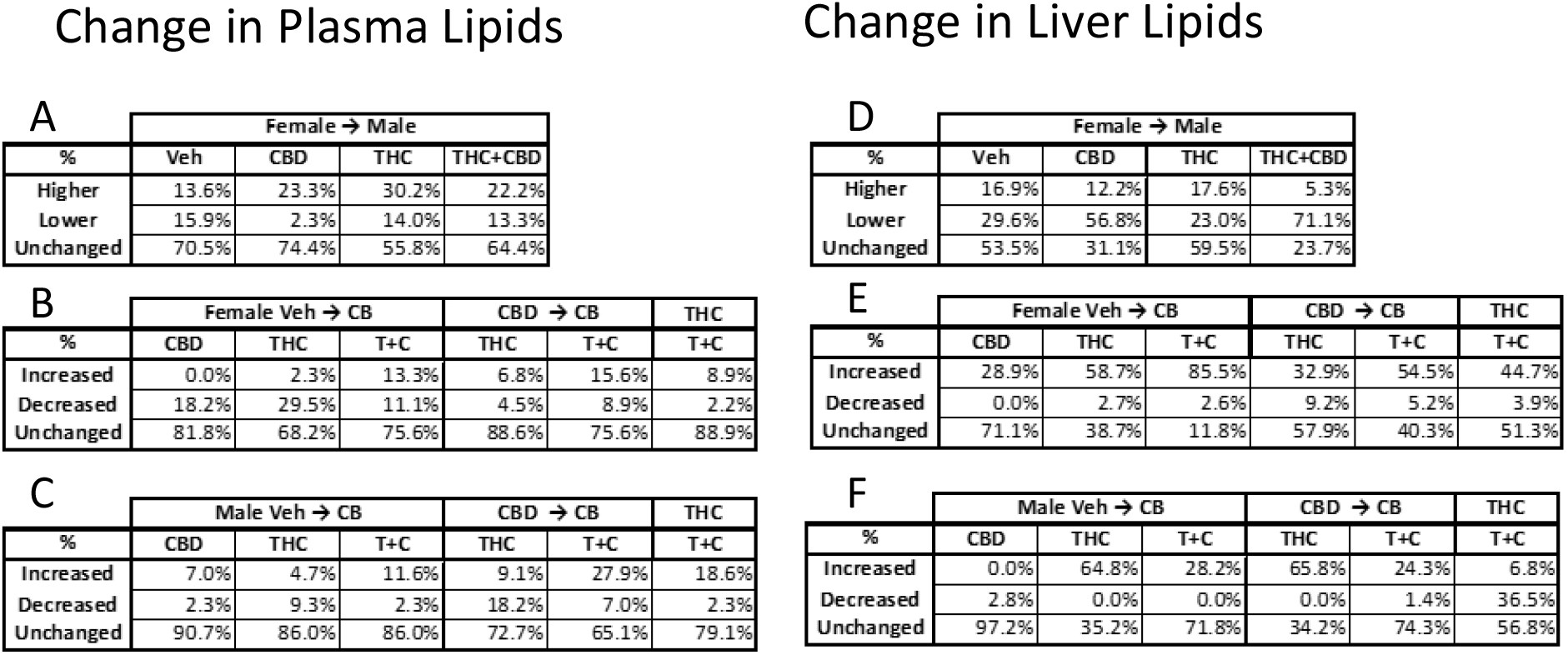
Overall directional changes in endogenous plasma and liver lipids 2hr post-i.p. injection of 10mg/kg CBD, THC, or THC+CBD. Percentages calculated from the total number of endogenous plasma lipids detected that were either higher, lower, or had no change relative to the compared treatment group (*see Methods and Supplemental Figure 1 for analytical details*). **A)** Percent changes in plasma lipid levels from females (→) to males across treatment groups. **B)** Percent changes in lipid levels from the female vehicle → to each female CB treatment group; from female CBD → to female THC and T+C; and from female THC → to female T+C. **C)** Percent changes in plasma lipid levels from the male vehicle → to the male CB treatment groups; from male CBD → to male THC and T+C; and from male THC → to T+C. **D)** Percent changes in liver lipid levels from females (→) to males across treatment groups. **E)** Percent changes in liver lipid levels from the female vehicle → to each female CB treatment group; from female CBD → to female THC and T+C; and from female THC → to female T+C. **F)** Percent changes in liver lipid levels from the male vehicle → to the male CB treatment groups; from male CBD → to male THC and T+C; and from male THC → to T+C.

Figure 2A shows that in the plasma of vehicle-treated animals 15.9% of endolipids were significantly lower in males compared to females, 13.6% were significantly higher, and 70.5% were not significantly different. In the liver of vehicle treated animals (Fig. 2D) 29.6% were significantly lower in males compared to females, 16.9% were significantly higher and 53.5% were not significantly different between males and females. This same analysis is also shown for male CB treatment groups (10 mg/kg CBD, THC, or T+C) relative to females. In plasma, each CB treatment increased the percentage of endolipids that were *significantly higher* in males compared to females, with THC causing an additional 15% more endolipids affected (Fig. 2A). A unique feature of CBD treatment in plasma was that the percentage of lipids that were lower in males compared to females was reduced to only 2%, meaning that those lipids were either decreased in the female or increased in the male, putting them in the same range. Data show that this was a differential effect and did not occur all in one direction (see supplemental data for all heatmaps). In the liver each CB treatment resulted in a reduction in the number of endolipids that were significantly higher in males than females (Fig. 2D) with only 5.3% of liver endolipids higher in males than females after T+C treatment.

Figures 2B and 2E report the percentage of endolipids that changed in female plasma (Fig. 2B) or female liver (Fig. 2E) from three levels of analysis: 1) Each CB treatment compared to vehicle (Veh → CB); 2) CB treatment (THC or T+C) compared to CBD (CBD → CB); or 3) T+C treatment compared to THC. Likewise, these comparisons were made for male plasma (Fig. 2C) and male liver (Fig. 2F). *Overall, female plasma and liver endolipids were more affected by CB treatment than males*. In both sexes and tissues, CBD had the least effect on endolipids at this 2-hour time point compared to Veh, but the direction of changes that did occur differed between tissue and sex. In females, CBD caused no increases in plasma endolipids (0%) and no decreases in liver endolipids (0%) but caused decreases in 18.2% of plasma endolipids and increases in 28.9% of liver endolipids. In males, the opposite relationship appeared wherein CBD caused decreases in only 2.3% of plasma endolipids and no increases in liver endolipids (0%), while causing increases in 7% of plasma endolipids and decreases in 2.8% of liver endolipids. In the male liver, THC alone had the greatest effect on endolipids, but the female liver was most affected by T+C.

Comparisons within CB treatment groups illustrate that male plasma endolipids were more affected between CB treatments than females. Compared to levels measured after CBD treatment, male plasma endolipids were changed by 27% compared to THC treatment and 34% compared to T+C. Female levels only changed by 11% and 24% respectively. Comparing plasma endolipids after THC to T+C, males changed 21% and females 11%. Female and male liver endolipids were more similarly affected between sexes, and more affected than plasma endolipids. Compared to levels measured after CBD treatment, male liver endolipids were changed by 65.8% compared to THC treatment and 24.3% compared to T+C. Female levels changed by 32.9% and 54.5% respectively. Comparing liver endolipids after THC to T+C, males changed 43% and females 49%. In both sexes and tissue types, endolipids that changed with THC or T+C relative to CBD or T+C relative to THC were predominantly increased.

### 3.3 Liver levels of *N*-acyl ethanolamines are more variable with sex and CB treatment than plasma

Levels of *N*-acyl ethanolamines showed greater sex-dependence and responsiveness to CB treatment in the liver (Figures 3Ei-vi) than plasma (Figures 3Ai-vi). After vehicle injection, males had higher liver levels of *N-*palmitoyl ethanolamine (PEA), *N*-stearoyl ethanolamine (SEA), and *N*-oleoyl ethanolamine (OEA), and lower levels of *N*-docosahexaenoyl ethanolamine (DEA) than females, but *N*-linoleoyl ethanolamine (LEA) and AEA levels did not differ (Fig. 3E, 3F). In contrast with the liver, only a subset of NAEs in the plasma, PEA and SEA, were higher in males relative to females after vehicle treatment (Fig. 3Aii, 2Aiii, 2B).

**Figure 3.**
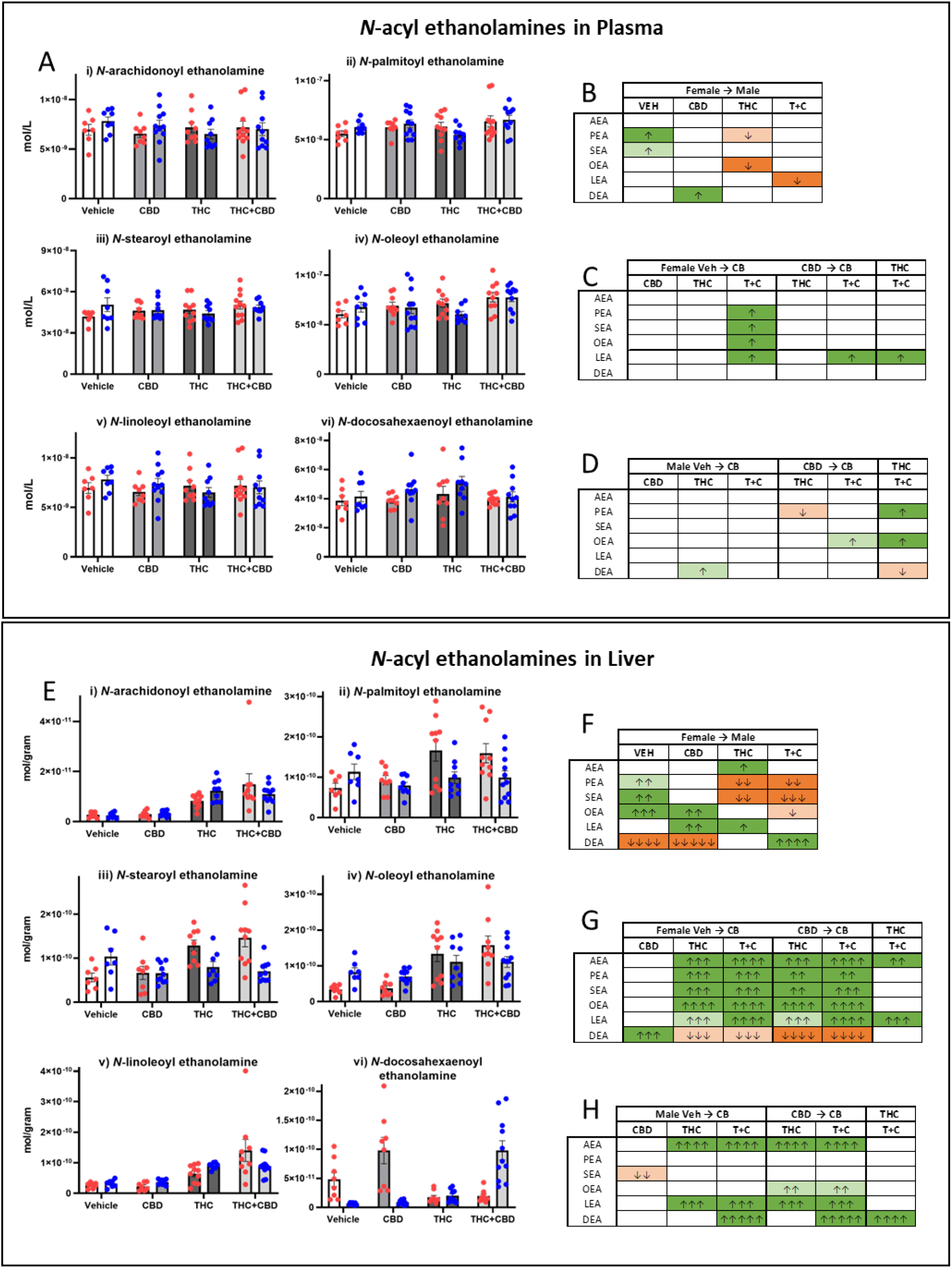
Levels of plasma and liver *N*-acyl ethanolamines 2hr after (10mg/kg) CBD, THC, or THC+CBD administration to male and female mice. **A)** Plasma concentrations (mol/L; means ±SEM) of each *N-*acyl ethanolamine (NAE): i) *N*-arachidonoyl ethanolamine (AEA), ii) *N*-palmitoyl ethanolamine (PEA), iii) *N*-stearoyl ethanolamine (SEA), iv) *N*-oleoyl ethanolamine (OEA), v) *N*-linoleoyl ethanolamine (LEA), and vi) *N*-docosahexaenoyl ethanolamine (DEA). Individual measurements from each female (red) and male (blue) animal are represented by a corresponding dot. **B)** Heatmaps illustrating significant differences in plasma NAE levels in males compared to (→) females across all treatment groups. **C)** Heatmaps illustrating significant differences in plasma NAE levels from female vehicle →to female CB treatment groups; from female CBD → to female THC and T+C treatment groups; and from female THC → to female T+C treatment group. **D)** Heatmaps illustrating significant differences in plasma NAE levels from male vehicle → to male CB treatment groups; from male CBD → to male THC and T+C treatment groups; and from male THC → to male T+C treatment group. **E)** Liver concentrations (mol/g; means ±SEM) of each *N-*acyl ethanolamine (NAE): i) *N*-arachidonoyl ethanolamine (AEA), ii) *N*-palmitoyl ethanolamine (PEA), iii) *N*-stearoyl ethanolamine (SEA), iv) *N*-oleoyl ethanolamine (OEA), v) *N*-linoleoyl ethanolamine (LEA), and vi) *N*-docosahexaenoyl ethanolamine (DEA). Individual measurements from each female (red) and male (blue) animal are represented by a corresponding dot. **F)** Heatmaps illustrating significant differences in liver NAE levels in males compared to (→) females across all treatment groups. **G)** Heatmaps illustrating significant differences in liver NAE levels from female vehicle →to female CB treatment groups; from female CBD → to female THC and T+C treatment groups; and from female THC → to female T+C treatment group. **H)** Heatmaps illustrating significant differences in liver NAE levels from male vehicle → to male CB treatment groups; from male CBD → to male THC and T+C treatment groups; and from male THC → to male T+C treatment group. Dark shades depict significant differences (p < 0.05), light shades depict trending differences (0.05 < p < 0.10). Green indicates significant increases, orange indicates significant decreases, arrows indicate effect size. *See Methods for description of heatmap analytics. See supplemental data for all statistical analyses*.

CB treatment tended to attenuate the sex differences in liver NAEs with vehicle treatment, though to varying degrees. After CBD treatment, males continued to show higher liver levels of OEA relative to females, though the fold-difference between males and females was smaller than what was measured after vehicle. DEA levels also remained lower in males relative to females, but the fold difference was slightly greater. Males also had significantly higher LEA levels compared to females but significant differences in SEA and PEA were abolished. Likewise, in plasma, CBD treatment abolished the sex differences in PEA and SEA, while also significantly increasing DEA in males relative to females.

THC treatment resulted in sex differences in liver NAEs in the same direction as what was seen with CBD treatment, but at a greater magnitude. In the liver, THC-treated males had significantly lower levels of PEA and SEA, and higher levels of LEA and AEA, but no differences in OEA and DEA compared to females. The sex-differences in plasma NAEs after THC treatment most closely resemble those seen in the liver after THC treatment (Fig. 3B). However, the differences in plasma exhibited smaller fold-changes than liver NAEs. In the liver, combination T+C treatment resulted in a near complete reversal of sex differences seen after vehicle treatment. Males had significantly lower levels of PEA, SEA, and OEA than females and higher levels of DEA, but LEA and AEA remained unaffected. Plasma did not show the same effect of T+C treatment, although males no longer had significantly higher levels of NAEs than females, instead having lower levels of LEA (Fig. 3Av, 3B).

The sex differences in NAEs after CB treatment in the liver are largely the result of increases in NAEs in female liver after CB treatment, rather than decreases in male liver NAEs. Assessment of differences within sex showed that females were highly affected by THC treatment, whether it was administered alone or in combination with CBD (Fig 3G). All NAEs except DEA were significantly higher in the liver of THC and THC+CBD treated females compared to both vehicle and CBD treated females. In female plasma, NAEs (PEA, SEA, OEA, LEA) were significantly higher than vehicle only in the T+C group. Levels of NAEs in CBD and THC treated female plasma looked similar to levels in vehicle treated females. NAEs in male livers were less broadly affected by THC treatment, with the only differences relative to vehicle being higher levels of LEA and AEA after THC treatment, and LEA, AEA, and DEA after THC+CBD treatment. NAEs in both plasma and liver from both male and female CBD treated animals did not vary significantly from vehicle.

### 3.4 Plasma and liver levels of 2-acyl glycerols are highly variable by sex and THC or THC+CBD treatment

Plasma 2-acyl glycerols were more dynamic than plasma NAEs and differed by both CB treatment and sex (Fig. 4A-D). 2-arachidonoyl glycerol (2-AG) and 2-oleoyl glycerol (2-OG) levels were significantly higher in males than females in vehicle controls, however, only levels of 2-AG remained higher after CBD treatment. Similar to plasma, with vehicle injection, male liver had significantly higher levels of 2-AG and 2-OG, as well as 2-LG than females (Fig. 4E-F). However, unlike plasma, all of these compounds remained elevated in male liver after CBD treatment along with levels of 2-PG. THC and T+C treatment resulted in the most tissue-dependent sex differences in 2-acyl glycerols. In plasma, levels of 2-OG and 2-palmitoyl glycerol (2-PG) were significantly lower in males than females with THC treatment, but levels of 2-OG and 2-LG were significantly higher in males than females with T+C treatment. In contrast, the only sex differences after THC and T+C treatment in the liver were elevated levels of 2-LG and decreased levels of 2-AG respectively (Fig. 4F).

**Figure 4.**
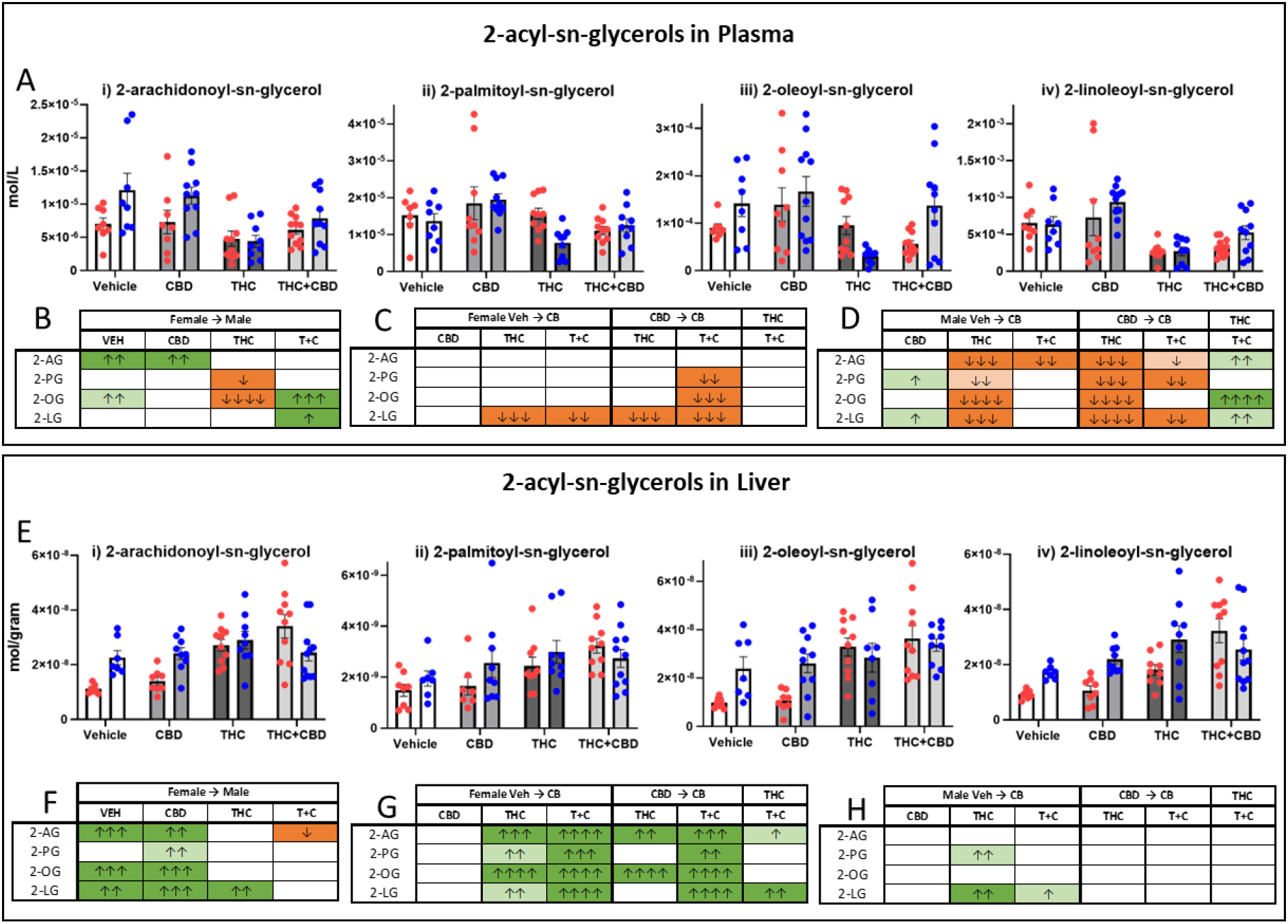
Levels of plasma and liver 2-acyl glycerols 2hr after (10mg/kg) CBD, THC, or THC+CBD administration to male and female mice. **A)** Plasma concentrations (mol/L; means ±SEM) of each 2-acyl glycerol: i) 2-arachidonoyl glycerol (2-AG), ii) 2-palmitoyl glycerol (2-PG), iii) 2-oleoyl glycerol (2-OG), iv) 2-linoleoyl glycerol (2-OG). Individual measurements from each female (red) and male (blue) animal are represented by a corresponding dot. **B)** Heatmaps illustrating significant differences in plasma 2-acyl glycerol levels in males compared to (→) females across all treatment groups. **C)** Heatmaps illustrating significant differences in plasma 2-acyl glycerol levels from female vehicle → to female CB treatment groups; from female CBD → to female THC and T+C treatment groups; and from female THC → to female T+C treatment group. **D)** Heatmaps illustrating significant differences in plasma 2-acyl glycerol levels from male vehicle → to male CB treatment groups; male CBD → male THC and T+C treatment groups; and male THC → male T+C treatment group. **E)** Liver concentrations (mol/g; means ±SEM) of each 2-acyl glycerol: i) 2-arachidonoyl glycerol (2-AG), ii) 2-palmitoyl glycerol (2-PG), iii) 2-oleoyl glycerol (2-OG), iv) 2-linoleoyl glycerol (2-OG). Individual measurements from each female (red) and male (blue) animal are represented by a corresponding dot. **F)** Heatmaps illustrating significant differences in 2-acyl glycerol levels in male liver compared to (→) female across all treatment groups. **G)** Heatmaps illustrating significant differences in liver 2-acyl glycerol levels from female vehicle → to female CB treatment groups; from female CBD → to female THC and T+C treatment groups; and from female THC → to female T+C treatment group. **H)** Heatmaps illustrating significant differences in liver 2-acyl glycerol levels from male vehicle → to male CB treatment groups; male CBD → male THC and T+C treatment groups; and male THC → male T+C

Comparing 2-acyl glycerol levels after CB treatment to vehicle in female plasma, only 2-LG levels were significantly lower in the THC and T+C groups (Fig. 4C). Conversely, levels of all 2-acyl glycerols were higher in the THC and T+C groups compared to vehicle in female liver (Fig. 4G). Comparisons within CB treatment in female plasma (Fig. 4C) showed that levels of 2-PG, 2-OG, and 2-LG were significantly lower in the T+C group compared to the CBD group, with no significant differences between THC and T+C. In contrast, all 2-acyl glycerols were significantly higher in the T+C group compared to the CBD group in female liver, but only 2-AG and 2-LG were higher in the T+C group compared to THC.

Comparisons of 2-acyl glycerol levels after CB treatment to vehicle were more dynamic in male plasma than female (Fig. 4D)-all 2-acyl glycerol species significantly decreased with THC treatment, 2-PG and 2-LG significantly increased with CBD, and 2-AG decreased with T+C. In male liver, the only differences in 2-acyl glycerols after CB treatment compared to vehicle were increased levels of 2-PG and 2-LG after THC treatment and increased levels of 2-LG after T+C treatment. Again, male plasma lipid levels were more dynamic than female (Fig. 4D) when comparing between CB treatment groups with all 2-acyl glycerol species significantly lower in the THC treatment compared to CBD, and all but 2-OG being significantly lower in the T+C group compared to CBD. However, the fold-changes were smaller in the combination group. This is reflected in the comparison in the T+C group to THC alone showing that lipid levels were significantly higher. No differences existed in 2-acyl glycerol levels between CB treatments in the male liver.

### 3.5 Levels of Free Fatty Acids in plasma and liver differ as a function of sex and CB treatment

In plasma there were more overall sex differences in free fatty acids (FFAs) than as a result of CB treatments (Fig. 5Ai-vi; B); conversely, in the liver there were more overall differences in FFAs in response to CB treatment (Figures 5Ei-v, 5F). Plasma levels of arachidonic acid (ARA) were higher in males with CBD than females with CBD, but significantly lower with THC and T+C treatment compared to females. However, plasma levels of linoleic acid (LIN) were significantly lower in males than females with vehicle. Plasma eicosapentaenoic acid (EIC) levels were significantly higher in males than females in vehicle, CBD, and T+C groups. Similarly, liver levels of EIC were higher in males than females across all treatment groups. In liver tissue from the vehicle group, males had higher levels of all FFAs except LIN, which did not significantly differ from females. Males in the CBD group also tended to have higher liver levels of LIN and ARA than females. The only compounds that were lower in male liver than female were oleic acid (OLE) in the THC alone treatment group, and LIN in the T+C treatment group.

**Figure 5.**
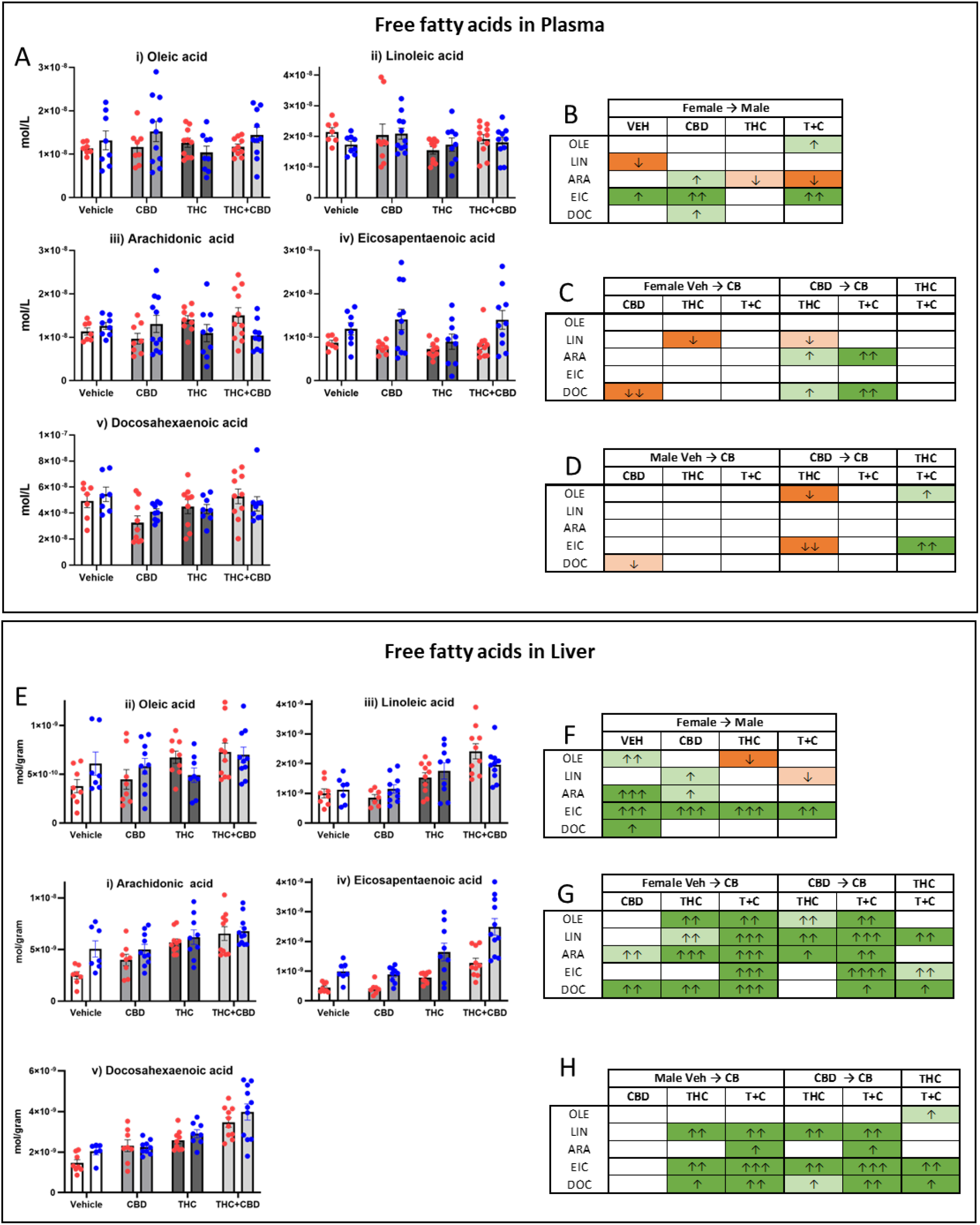
Levels of plasma and liver free fatty acids 2hr after (10mg/kg) CBD, THC, or THC+CBD administration to male and female mice. **A)** Plasma concentrations (mol/L; means ±SEM) of each free fatty acid: i) oleic acid (OLE), ii) linoleic acid (LIN), iii) arachidonic acid (ARA), iv) eicosapentaenoic acid (EIC), v) docosahexaenoic acid (DOC). Individual measurements from each female (red) and male (blue) animal are represented by a corresponding dot. **B)** Heatmaps illustrating significant differences in plasma free fatty acids in males compared to (→) females across treatment groups. **C)** Heatmaps illustrating significant differences in plasma free fatty acid levels from female vehicle → to female CB treatment groups; from female CBD → to female THC and T+C treatment groups; and from female THC → to female T+C. **D)** Heatmaps illustrating significant differences in plasma free fatty acid levels from male vehicle → to male CB treatment groups; from male CBD → to male THC and T+C treatment groups; and from male THC → male T+C. **E)** Liver concentrations (mol/g; means ±SEM) of each free fatty acid: i) oleic acid (OLE), ii) linoleic acid (LIN), iii) arachidonic acid (ARA), iv) eicosapentaenoic acid (EIC), v) docosahexaenoic acid (DOC). Individual measurements from each female (red) and male (blue) animal are represented by a corresponding dot. **F)** Heatmaps illustrating significant differences in liver free fatty acids in males compared to (→) females across treatment groups. **G)** Heatmaps illustrating significant differences in liver free fatty acid levels from female vehicle → to female CB treatment groups; from female CBD → to female THC and T+C treatment groups; and from female THC → to female T+C. **H)** Heatmaps illustrating significant differences in liver free fatty acid levels from male vehicle → to male CB treatment groups; from male CBD → to male THC and T+C treatment groups; and from male THC → male T+C. Dark shades depict significant differences (p < 0.05), light shades depict trending differences (0.05 < p < 0.10). Green indicates significant increases, orange indicates significant decreases, arrows indicate effect size. *See Methods for description of heatmap analytics. See supplemental data for all statistical analyses*.

In male plasma the only significant difference in FFAs after CB treatment compared to vehicle was a lower level of DOC after CBD. Female plasma also showed significantly lower levels of DOC after CBD treatment compared to vehicle. In comparing CBD treatment to THC or T+C in female plasma, levels of ARA and DOC were significantly higher in both THC-containing treatments while the level of LIN was significantly lower in THC only. In male plasma, levels of EIC and OLE were significantly lower in THC compared to CBD. There were no differences in FFAs between THC and T+C treatment in female plasma; however, levels of EIC and OLE were higher in T+C compared to THC in males. Similar to plasma, neither male nor female liver FFAs were highly affected by CBD treatment compared to vehicle. Instead, most changes were seen in the T+C treatment group, and all significant differences were increases compared to baseline (Figures 5G, 5H). Females were more affected by THC treatment than males, with all compounds higher than vehicle (Fig 5G). In males, the key significant differences after THC treatment were higher levels of LIN, EIC, and DOC compared to vehicle (Fig 5H).

### 3.6 Changes in levels of lipoamines like *N*-acyl phenylalanines are sex, tissue, and CB treatment dependent

There was little to no concurrency in the levels of *N*-acyl phenylalanines (NAPhe) between plasma and liver as a function of sex or CB treatment. Sex differences in plasma levels had a mosaic pattern with levels being lower in males with vehicle treatment, but males having higher overall levels with THC treatment (Figures 6Ai-vi, 6B). Conversely, levels of NAPhe were significantly lower in males across all groups in the liver with CBD and T+C groups showing the most pronounced differences (Figures 6Ei-vi, 6F).

**Figure 6.**
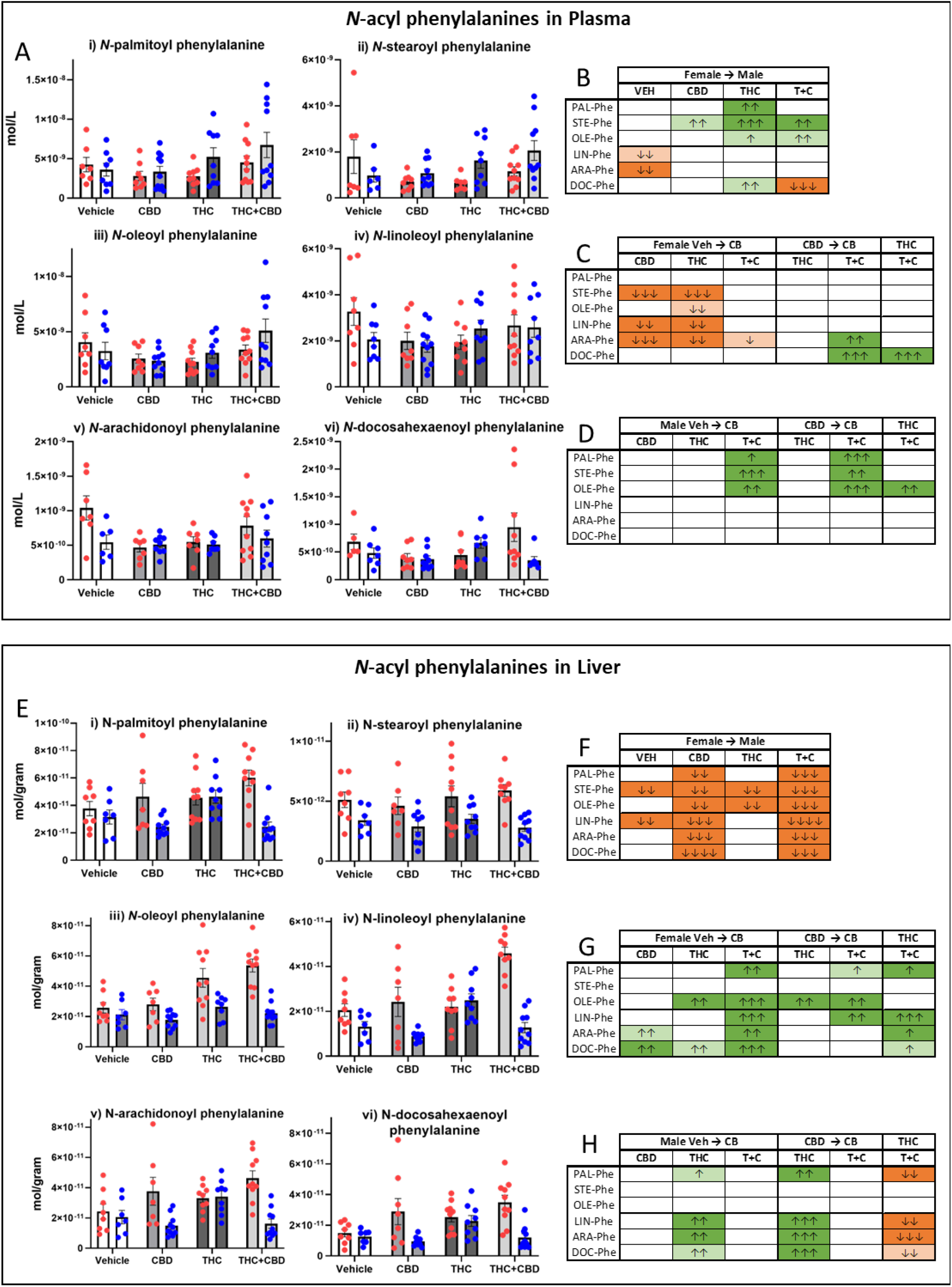
Levels of plasma and liver *N*-acyl phenylalanines 2hr after (10mg/kg) CBD, THC, or THC+CBD administration to male and female mice. **A)** Plasma concentrations (mol/L; means ±SEM) of each *N-*acyl phenylalanine: i) *N*-palmitoyl phenylalanine (PAL-Phe), ii) *N*-stearoyl phenylalanine (STE-Phe), iii) *N*-oleoyl phenylalanine (OLE-Phe), iv) *N*-linoleoyl phenylalanine (LIN-Phe), v) *N*-arachidonoyl phenylalanine (ARA-Phe), vi) *N*-docosahexaenoyl phenylalanine (DOC-Phe). Individual measurements from each female (red) and male (blue) animal are represented by a corresponding dot. **B)** Heatmaps illustrating significant differences in plasma *N*-acyl phenylalanine levels in males compared to (→) females across treatment groups. **C)** Heatmaps illustrating significant differences in plasma *N*-acyl phenylalanine levels from female vehicle → to female CB treatment groups; from female CBD → to female THC and T+C treatment groups; and from female THC → to female T+C. **D)** Heatmaps illustrating significant differences in plasma *N*-acyl phenylalanine levels from male vehicle → to male CB treatment groups; from male CBD → to male THC and T+C treatment groups; and from male THC → to male T+C. **E)** Liver concentrations (mol/g; means ±SEM) of each *N*-acyl phenylalanine: i) *N*-palmitoyl phenylalanine (PAL-Phe), ii) *N*-stearoyl phenylalanine (STE-Phe), iii) *N*-oleoyl phenylalanine (OLE-Phe), iv) *N*-linoleoyl phenylalanine (LIN-Phe), v) *N*-arachidonoyl phenylalanine (ARA-Phe), vi) *N*-docosahexaenoyl phenylalanine (DOC-Phe). Individual measurements from each female (red) and male (blue) animal are represented by a corresponding dot. **F)** Heatmaps illustrating significant differences in liver *N*-acyl phenylalanine levels in males compared to (→) females across treatment groups. **G)** Heatmaps illustrating significant differences in liver *N*-acyl phenylalanine levels from female vehicle → to female CB treatment groups; from female CBD → to female THC and T+C treatment groups; and from female THC → to female T+C. **H)** Heatmaps illustrating significant differences in liver *N*-acyl phenylalanine levels from male vehicle → to male CB treatment groups; from male CBD → to male THC and T+C treatment groups; and from male THC → to male T+C. Dark shades depict significant differences (p < 0.05), light shades depict trending differences (0.05 < p < 0.10). Green indicates significant increases, orange indicates significant decreases, arrows indicate effect size. *See Methods for description of heatmap analytics. See supplemental data for all statistical analyses*.

Effects of CB treatment on female plasma showed that both CBD and THC treatment caused multiple decreases and only minimal effects with T+C, whereas there were no differences in males compared to vehicle with CBD and THC but significant increases with T+C. When comparing CBD treatment to THC and T+C, plasma NAPhe levels were significantly increased in the T+C group in both males and females, though the specific lipids were different. Supplemental data figures 1.4 and 1.5 show a very similar pattern for *N*-acyl tyrosines and *N*-acyl valines respectively with levels decreasing in female plasma with CB treatment and increasing in male plasma. In females, liver levels of *N*-docosahexaenoyl phenylalanine were increased by all CB treatments (Fig 6G). The combination T+C treatment also led to increased levels of *N*-acyl phenylalanines except *N*-stearoyl phenylalanine. This is in contrast with male T+C animals, whose levels of liver *N*-acyl phenylalanines did not differ significantly from vehicle. Instead, the only treatment that impacted levels in males was THC alone, which had significantly higher levels than vehicle and CBD animals (Fig 6H).

## 4. Discussion

Previous studies measuring CBs in plasma, blood, and serum have demonstrated that their concentration varies depending on dose, route of administration, and time post-treatment ^1,8,24^. In rats, several groups have reported that THC levels remain similar between males and females ^1,4,8,24^, though female adolescent rats tended to have higher concentrations after injection (5 mg/kg, peak concentrations occurring 30 min post-injection for females and 15 min for males)^24^ and inhalation (25 mg/mL over 30 min, male and female peak concentrations both occurring 5 min post-inhalation session)^8^, and adult female rats tended to have higher concentrations after injection (5 mg/kg, both male and female peak concentrations occurring at 1 h) ^1^. In humans, females have significantly higher plasma peak concentrations than males when administered THC (5 mg) orally while in a fasted state ^25^. However, it has also been found that adult male rats given an injection of THC (2.5 mg/kg) have significantly higher THC levels than females (from 15 min post-injection to 240 min post-injection) ^10^. This finding is consistent with our results in that males had significantly higher THC levels than females at 2 hours post-injection even though the dose here was 10 mg/kg.

Sex differences in plasma CBD have been less extensively evaluated. In one human study where CBD was administered orally to men and women (1500-6000 mg), the maximum blood concentration increased at a rate that is less than 1:1 with dose, and the peak concentration of CBD occurred 4-5 h post-administration and CBD metabolites at 3.5-5 h post-administration regardless of dose. However, data from both sexes was combined for the publication analysis, which meant that we could not compare any potential sex differences here ^26^. Comparison of a single oral dose of CBD (115mg/kg) given to male and female rats shows that females reach a peak plasma concentration after 7:45 hrs, while males reach a peak concentration at 8:25 hrs ^27^. While accumulation of CBD in tissue was not evaluated after acute administration, after 28 days of chronic treatment, males and females did not significantly differ in CBD levels in adipose or muscle tissue, but females had significantly higher levels of CBD than males in liver ^27^. A more comprehensive evaluation of plasma THC and CBD over time and at multiple doses in both males and females would help determine which variables account for these differences in THC accumulation across studies and clarify the pharmacokinetics of CBD in both sexes.

Our results show that the combination of THC+CBD results in higher levels of both THC and CBD in females and higher levels of CBD in males than when each CB is administered alone. Pooled analysis of data from men and women given THC+CBD (1600 μg) via *i.v*. injection show the peak plasma THC concentration is 30 ηg/mL at 5 min, peak CBD concentration is 22 ηg/mL at 7 min, peak 11-OH-THC occurs at 8 min, and peak 11-COOH-THC occurs at 53 min post-injection ^28^. Analysis of males and females separately shows sex differences in a THC-dominant cannabis concentrate with less than 1% CBD. Men that inhaled these 70-90% THC cannabis concentrates had significantly higher levels of THC and CBD in their plasma than females immediately post-consumption, but levels of 11-OH-THC did not differ, and the differences were no longer significant 1h after ingestion ^29^. This supports the need for analysis of male and female samples separately to evaluate sex-dependent effects that might be informative, and a similarly rigorous evaluation of the time-course of THC+CBD at multiple doses.

Overall evaluation of CB metabolites has shown that females accumulate higher levels of 11-OH-THC than males ^1,8,10,24^. Here, 2-hours post-treatment, females had higher 11-OH-THC levels only when THC was delivered in combination with CBD. These differences illustrate the rapidity in which CB metabolism takes place and that the metabolism is consistently sex dependent. Both CBD and THC are metabolized by cytochrome P450 enzymes such as CYP2J2, through which they also prevent AEA degradation (THC to a greater extent than CBD) ^30^. CBD, THC, and their metabolites 11-OH-THC, 11-COOH-THC, 7-OH-CBD, and 7-COOH-CBD also act as CYP inhibitors ^31^. However, male and female mice have different CYP mRNA expression profiles ^32^, and liver expression changes across the estrous cycle and in response to sex steroids ^33^. Therefore, sex differences in CYPs might explain the differences in metabolism seen here. Indeed, metabolism of THC to 11-OH-THC in microsomal preparations derived from females occurs at 3 times the rate of the metabolism in cells derived from males ^1^.

Here, we reported that male and female mice have different concentrations of many endolipids prior to and after CB administration. Baseline sex differences in lipid signaling molecules have been documented in humans, including differences in 2-acyl glycerols ^34^ and AEA which modulates naturally across the menstrual cycle ^35^, as well as across the estrous cycle in the CNS in rodents ^13^. Male and female rats also show differences in liver triacylglycerols, phosphatidylcholines, and phosphatidylethanolamines in response to dietary manipulations ^36,37^. Therefore, we can hypothesize that differences in hormonal environment likely drive sex differences in both baseline and treatment-induced plasma and liver lipids. It is beyond the scope of this report to speculate on the signaling outcomes of the dynamics of the differences in endolipids described here; however, we have recent reports that delve into some of the known signaling pathways of these understudied endolipids that would be a starting point to understanding their relevance. A few highlights of that research show that these endolipids act as agonists and antagonists at TRPV1-4 receptors ^38^ and agonists at GPR119 ^39^. Investigation of cannabinoid exposure on endolipids shows that changes do occur in the liver. For instance, primary hepatocytes from male mice show that THC leads to dose- and time-dependent increases in AEA, 2-AG, 2-OG, and 2-PG, but not PEA, OEA, DHEA, or EPEA ^40^. Interestingly, activation of hepatic stellate cells without pharmacologic intervention but through culture results in increased NAEs, 2-acyl glycerols, and some FFAs ^41^. Selective knockdown of liver NAPE-PLD in mice results in increased lipid droplet size in the liver, lower levels of 1/2-AG, 2-OG, 1/2-LG, OEA and LEA, but not AEA or PEA, higher levels of ARA, and changes in oxysterols and bile acids ^42^. That the changes in these plasma and liver endolipids are both CB and sex dependent provides an important reminder that the regulation of lipid signaling molecules by drugs and genetic sex is multifactorial and the focus on only a few signaling lipids within the cannabinoid field (*e.g*. Anandamide and 2-AG) greatly limits our understanding of these systems.

## Notes

### Competing Interest Statement

The authors have declared no competing interest.

