## Supplemental Data for "Plasma and Liver Lipids are Differentially Regulated After Cannabinoid Treatments in Male and Female Mice"

### Supplemental Figures

**Supplemental Table 6.2 Descriptive statistics for *N*-acyl phenylalanines (mol/g) in liver collected from male and female adult mice 2h after receiving an i.p. injection of 10mg/kg CBD, THC, or THC+CBD** .. 101

**Supplemental Table 7.1 Descriptive statistics for *N*-acyl tyrosines (mol/L) in plasma collected from male and female adult mice 2h after receiving an i.p. injection of 10mg/kg CBD, THC, or THC+CBD** .. 123

**Supplemental Table 10.1 Descriptive statistics for *N*-acyl glycines (mol/L) in plasma collected from male and female adult mice 2h after receiving an i.p. injection of 10mg/kg CBD, THC, or THC+CBD** .. 182

**Supplemental Table 11.1 Descriptive statistics for *N*-acyl leucines (mol/L) in plasma collected from male and female adult mice 2h after receiving an i.p. injection of 10mg/kg CBD, THC, or THC+CBD** .. 203

**Supplemental Table 12.1 Descriptive statistics for *N*-acyl prolines (mol/L) in plasma collected from male and female adult mice 2h after receiving an i.p. injection of 10mg/kg CBD, THC, or THC+CBD** .. 223

**Supplemental Table 14.1 Descriptive statistics for *N*-acyl taurines (mol/L) in plasma collected from male and female adult mice 2h after receiving an i.p. injection of 10mg/kg CBD, THC, or THC+CBD** .. 251

**Supplemental Table 16.1 Descriptive statistics for *N*-acyl methionines (mol/g) in liver collected from male and female adult mice 2h after receiving an i.p. injection of 10mg/kg CBD, THC, or THC+CBD** .. 275

**Supplemental Table 17.1 Descriptive statistics for *N*-acyl tryptophans (mol/L) in liver collected from male and female adult mice 2h after receiving an i.p. injection of 10mg/kg CBD, THC, or THC+CBD** .. 288

Supplemental Table 1. Lipids in HPLC-MS/MS Screening Library

| *N*-acyl alanine | $[M–H]^-$ | Fragment |
|---|---|---|
| *N*-palmitoyl alanine | 327 | 88.09 |
| *N*-stearoyl alanine | 355 | 88.09 |
| *N*-oleoyl alanine | 353 | 88.09 |
| *N*-linoleoyl alanine | 351 | 88.09 |
| *N*-arachidonoyl alanine | 375 | 88.09 |
| *N*-docosahexaenoyl alanine | 399 | 88.09 |
| *N*-acyl ethanolamine | $[M+H]^+$ | Fragment |
| *N*-palmitoyl ethanolamine | 300 | 62.1 |
| *N*-stearoyl ethanolamine | 328 | 62.1 |
| *N*-oleoyl ethanolamine | 326 | 62.1 |
| *N*-linoleoyl ethanolamine | 324 | 62.1 |
| *N*-arachidonoyl ethanolamine | 348 | 62.1 |
| *N*-docosahexaenoyl ethanolamine | 373 | 62.1 |
| N-acyl GABA | $[M+H]^+$ | Fragment |
| N-palmitoyl GABA | 341 | 102.1 |
| N-stearoyl GABA | 369 | 102.1 |
| N-oleoyl GABA | 367 | 102.1 |
| N-linoleoyl GABA | 365 | 102.1 |
| N-arachidonoyl GABA | 389 | 102.1 |
| N-docosahexaenoyl GABA | 413 | 102.1 |
| *N*-acyl glycine | $[M–H]^-$ | Fragment |
| *N*-palmitoyl glycine | 312 | 74.2 |
| *N*-stearoyl glycine | 340 | 74.2 |
| *N*-oleoyl glycine | 338 | 74.2 |
| *N*-linoleoyl glycine | 336 | 74.2 |
| *N*-arachidonoyl glycine | 360 | 74.2 |
| *N*-docosahexaenoyl glycine | 384 | 74.2 |
| *N*-acyl leucine | $[M–H]^-$ | Fragment |
| *N*-palmitoyl leucine | 369 | 130.1 |
| *N*-stearoyl leucine | 397 | 130.1 |
| *N*-oleoyl leucine | 395 | 130.1 |
| *N*-linoleoyl leucine | 393 | 130.1 |
| *N*-docosahexaenoyl leucine | 441 | 130.1 |
| *N*-acyl methionine | $[M–H]^-$ | Fragment |
| *N*-palmitoyl methionine | 387 | 148.2 |
| *N*-stearoyl methionine | 415 | 148.2 |
| *N*-oleoyl methionine | 413 | 148.2 |
| *N*-linoleoyl methionine | 411 | 148.2 |
| *N*-arachidonoyl methionine | 435 | 148.2 |
| *N*-docosahexaenoyl methionine | 459 | 148.2 |
| *N*-acyl phenylalanine | $[M–H]^-$ | Fragment |
| *N*-palmitoyl phenylalanine | 403 | 164.1 |
| *N*-stearoyl phenylalanine | 431 | 164.1 |
| *N*-oleoyl phenylalanine | 429 | 164.1 |

| *N*-acyl proline | $[M–H]^-$ | Fragment |
|---|---|---|
| *N*-palmitoyl proline | 353 | 114.12 |
| *N*-stearoyl proline | 381 | 114.12 |
| *N*-oleoyl proline | 378 | 114.12 |
| *N*-linoleoyl proline | 377 | 114.12 |
| *N*-arachidonoyl proline | 401 | 114.12 |
| *N*-docosahexaenoyl proline | 425 | 114.12 |
| *N*-acyl serine | $[M–H]^-$ | Fragment |
| *N*-palmitoyl serine | 342 | 74 |
| *N*-stearoyl serine | 370 | 74 |
| *N*-oleoyl serine | 368 | 74 |
| *N*-linoleoyl serine | 366 | 74 |
| *N*-arachidonoyl serine | 390 | 74 |
| *N*-docosahexaenoyl serine | 414 | 74 |
| *N*-acyl taurine | $[M–H]^-$ | Fragment |
| *N*-palmitoyl taurine | 363 | 124 |
| *N*-stearoyl taurine | 391 | 124 |
| *N*-oleoyl taurine | 389 | 124 |
| *N*-arachidonoyl taurine | 411 | 124 |
| N-acyl tryptophan | $[M–H]^-$ | Fragment |
| N-palmitoyl tryptophan | 442 | 203.1 |
| N-stearoyl tryptophan | 470 | 203.1 |
| N-oleoyl tryptophan | 468 | 203.1 |
| N-linoleoyl tryptophan | 466 | 203.1 |
| N-arachidonoyl tryptophan | 490 | 203.1 |
| N-docosahexaenoyl tryptophan | 514 | 203.1 |
| *N*-acyl tyrosine | $[M–H]^-$ | Fragment |
| *N*-palmitoyl tyrosine | 419 | 180.18 |
| *N*-stearoyl tyrosine | 447 | 180.18 |
| *N*-oleoyl tyrosine | 445 | 180.18 |
| *N*-linoleoyl tyrosine | 443 | 180.18 |
| *N*-arachidonoyl tyrosine | 466 | 180.18 |
| *N*-docosahexaenoyl tyrosine | 491 | 180.18 |
| *N*-acyl valine | $[M–H]^-$ | Fragment |
| *N*-palmitoyl valine | 354 | 116.31 |
| *N*-stearoyl valine | 383 | 116.14 |
| *N*-oleoyl valine | 381 | 116.14 |
| *N*-linoleoyl valine | 379 | 116.14 |
| *N*-docosahexaenoyl valine | 427 | 116.14 |
| Free Fatty Acids | $[M–H]^-$ | Fragment |
| Oleic acid | 282 | 263 |
| Linoleic acid | 280 | 261 |
| Arachidonic acid | 304 | 285 |
| Docosahexaenoic acid | 328 | 283 |
| Eicosapentaenoic acid | 302 | 257 |

| | | |
|---|---|---|
| *N*-linoleoyl phenylalanine | 427 | 164.1 |
| *N*-arachidonoyl phenylalanine | 451 | 164.1 |
| *N*-docosahexaenoyl phenylalanine | 475 | 164.1 |
| Phytocannabinoids | $[M+H]^+$ | Fragment |
| CBD | 315 | 191 |
| THC | 315 | 123.2 |
| Phytocannabinoid Metabolites | $[M–H]^-$ | Fragment |
| 7-OH-CBD | 330 | 173 |
| 11-OH-THC | 330 | 173 |
| 7-COOH-CBD | 344 | 191 |
| 11-COOH-THC | 344 | 191 |

| 2-acyl glycerol | $[M+H]^+$ | Fragment |
|---|---|---|
| 2-acyl glycerol | $[M+H]^+$ | Fragment |
| 2-arachidonoyl glycerol | 379 | 287.5 |
| 2-linoleoyl glycerol | 356 | 245 |
| 2-oleoyl glycerol | 358 | 265.2 |
| 2-palmitoyl glycerol | 332 | 239.5 |
| Prostaglandins | $[M+H]^+$ | Fragment |
| PGE2 | 351 | 315 |
| PGF2α | 353 | 309.2 |
| 6-keto-PGF1α | 369 | 206.9 |

Lipids in the HPLC/MS/MS screening library. Masses of parent and ion fragments for each lipid in the HPLC/MS/MS screening library. All compounds are run in negative ionization mode ($[M – H]^-$ parent ion) except *N*-acyl ethanolamine, 2-acyl glycerol, and phytocannabinoid methods, which are run in positive ionization mode ($[M + H]^+$ parent ion).

Supplemental Figure 1.1. Heatmaps of significant differences between lipids measured in plasma from male mice relative to levels measured in plasma from female mice 2h after a 10mg/kg injection of CBD, THC, or THC+CBD.

| | VEH | CBD | THC | THC+CBD |
|---|---|---|---|---|
| ***N-acyl alanine*** | | | | |
| *N-palmitoyl alanine* | | | | |
| *N-stearoyl alanine* | | | ↓↓ | ↓ |
| *N-oleoyl alanine* | ↑ | | | |
| *N-linoleoyl alanine* | | | | |
| *N-arachidonoyl alanine* | BAL | BDL | BDL | BAL |
| *N-docosahexaenoyl alanine* | BDL | BAL | BDL | BDL |
| ***N-acyl ethanolamine*** | | | | |
| *N-arachidonoyl ethanolamine* | | | | |
| *N-palmitoyl ethanolamine* | ↑ | | ↓ | |
| *N-stearoyl ethanolamine* | ↑ | | | |
| *N-oleoyl ethanolamine* | | | ↓ | |
| *N-linoleoyl ethanolamine* | | | | ↓ |
| *N-docosahexaenoyl ethanolamine* | | ↑ | | |
| ***N-acyl glycine*** | | | | |
| *N-palmitoyl glycine* | ↓ | ↑ | | ↓ |
| *N-stearoyl glycine* | | ↑ | | |
| *N-oleoyl glycine* | | | | |
| *N-linoleoyl glycine* | | | ↑↑ | |
| *N-arachidonoyl glycine* | BDL | BDL | BAL | BDL |
| *N-docosahexaenoyl leucine* | BDL | BDL | BDL | BDL |
| ***N-acyl leucine*** | | | | |
| *N-palmitoyl leucine* | | | ↑↑ | |
| *N-stearoyl leucine* | | | ↑↑ | |
| *N-oleoyl leucine* | | | | ↑↑ |
| *N-linoleoyl leucine* | | | ↑↑ | |
| *N-docosahexaenoyl leucine* | | ↑↑ | | |
| ***N-acyl phenylalanine*** | | | | |
| *N-palmitoyl phenylalanine* | | | ↑↑ | |
| *N-stearoyl phenylalanine* | | ↑↑ | ↑↑↑ | ↑↑ |
| *N-oleoyl phenylalanine* | | | ↑ | ↑↑ |
| *N-linoleoyl phenylalanine* | ↓↓ | | | |
| *N-arachidonoyl phenylalanine* | ↓↓ | | | |
| *N-docosahexaenoyl phenylalanine* | | | ↑↑ | ↓↓↓ |
| ***N-acyl proline*** | | | | |
| *N-palmitoyl proline* | BAL | BAL | BAL | BAL |
| *N-stearoyl proline* | BAL | BAL | BAL | BAL |
| *N-oleoyl proline* | BDL | BDL | BAL | BDL |
| *N-linoleoyl proline* | BDL | BAL | BDL | BDL |
| *N-arachidonoyl proline* | BAL | BDL | BDL | BDL |
| *N-docosahexaenoyl proline* | BDL | BAL | BDL | BAL |

| | VEH | CBD | THC | THC+CBD |
|---|---|---|---|---|
| ***N-acyl serine*** | | | | |
| *N-palmitoyl serine* | | ↑↑ | | ↑↑↑ |
| *N-stearoyl serine* | BAL | BAL | BAL | |
| *N-oleoyl serine* | BAL | BAL | BAL | BAL |
| *N-linoleoyl serine* | BDL | BAL | BAL | BDL |
| *N-arachidonoyl serine* | BDL | BAL | BAL | BDL |
| *N-docosahexaenoyl serine* | BDL | BDL | BDL | BDL |
| ***N-acyl taurine*** | | | | |
| *N-palmitoyl taurine* | BAL | BAL | BAL | BAL |
| *N-stearoyl taurine* | BAL | BAL | BAL | BAL |
| *N-oleoyl taurine* | BAL | BAL | BAL | ↓ |
| *N-arachidonoyl taurine* | BDL | BAL | BDL | BDL |
| ***N-acyl tyrosine*** | | | | |
| *N-palmitoyl tyrosine* | | | ↑↑ | ↑↑ |
| *N-stearoyl tyrosine* | ↓↓↓ | | | |
| *N-oleoyl tyrosine* | | | ↑↑↑ | ↑↑↑ |
| *N-linoleoyl tyrosine* | | | ↑↑↑ | |
| *N-arachidonoyl tyrosine* | BAL | BAL | BAL | BAL |
| *N-docosahexaenoyl tyrosine* | | BAL | BAL | BAL |
| ***N-acyl valine*** | | | | |
| *N-palmitoyl valine* | ↓↓ | ↓ | ↑↑ | |
| *N-stearoyl valine* | | | | |
| *N-oleoyl valine* | | | | |
| *N-linoleoyl valine* | ↓↓ | | ↑↑↑ | |
| *N-docosahexaenoyl valine* | BAL | BAL | BAL | BAL |
| **2-acyl-sn-glycerol** | | | | |
| 2-arachidonoyl glycerol | ↑↑ | ↑↑ | | |
| 2-palmitoyl glycerol | | | ↓ | |
| 2-oleoyl glycerol | ↑↑ | | ↓↓↓↓ | ↑↑↑ |
| 2-linoleoyl glycerol | | | | ↑ |
| **Free Fatty Acids** | | | | |
| Oleic acid | | | | ↑ |
| Linoleic acid | ↓ | | | |
| Arachidonic acid | | ↑ | ↓ | ↓ |
| Eicosapentaenoic acid | ↑ | ↑↑ | | ↑↑ |
| Docosahexaenoic acid | | ↑ | | |

Significance ($p<0.05$) is depicted by dark shades of color, while trending towards significance ($0.05 < p < 0.10$) is depicted by light shades of color. Direction of difference, male relative to female, is depicted by color and arrow direction. When male concentrations are higher than female, the cell is green with up arrows. When male concentrations are lower than female, the cell is orange with down arrows. Fold change is represented by the number of arrows. 1 arrow represents a 1-1.49-fold change, 2 arrows represent a 1.5-1.99-fold difference, 3 arrows represent a 2-2.99-fold difference, 4 arrows represent a 3-9.99-fold difference, and 5 arrows represents a 10-fold or greater difference in males relative to females.

Supplemental Figure 1.2. Heatmaps of significant differences between lipids measured in liver from male mice relative to levels measured in liver from female mice 2h after a 10mg/kg injection of CBD, THC, or THC+CBD.

| | VEH | CBD | THC | THC+CBD |
|---|---|---|---|---|
| ***N-acyl alanine*** | | | | |
| *N-palmitoyl alanine* | | | | ↓ |
| *N-stearoyl alanine* | | ↓↓ | ↓ | ↓ |
| *N-oleoyl alanine* | | | ↓↓ | ↓ |
| *N-linoleoyl alanine* | ↓ | ↓ | ↑ | ↓↓ |
| *N-arachidonoyl alanine* | | ↓ | | ↓ |
| *N-docosahexaenoyl alanine* | | ↓↓ | | ↓ |
| ***N-acyl ethanolamine*** | | | | |
| *N-arachidonoyl ethanolamine* | | | ↑ | |
| *N-palmitoyl ethanolamine* | ↑↑ | | ↓↓ | ↓↓ |
| *N-stearoyl ethanolamine* | ↑↑ | | ↓↓ | ↓↓↓ |
| *N-oleoyl ethanolamine* | ↑↑↑ | ↑↑ | | ↓ |
| *N-linoleoyl ethanolamine* | | ↑↑ | ↑ | |
| *N-docosahexaenoyl ethanolamine* | ↓↓↓↓ | ↓↓↓↓↓ | | ↑↑↑↑ |
| ***N-acyl GABA*** | | | | |
| *N-palmitoyl GABA* | | ↓↓↓↓ | ↑↑ | ↓↓↓ |
| *N-stearoyl GABA* | ↑↑↑ | | | |
| *N-oleoyl GABA* | ↓↓ | ↓↓↓↓ | ↓↓ | ↓↓↓ |
| *N-linoleoyl GABA* | BAL | ↓↓↓↓ | ↑↑ | ↓↓ |
| *N-arachidonoyl GABA* | BAL | BAL | | ↓↓ |
| *N-docosahexaenoyl GABA* | BAL | | | ↓↓ |
| ***N-acyl glycine*** | | | | |
| *N-palmitoyl glycine* | | ↓ | ↑ | ↓↓ |
| *N-stearoyl glycine* | | | | ↓↓↓ |
| *N-oleoyl glycine* | | | ↓ | |
| *N-linoleoyl glycine* | | | ↑ | |
| *N-arachidonoyl glycine* | | | ↑ | |
| *N-docosahexaenoyl leucine* | | ↓↓ | ↑ | |
| ***N-acyl leucine*** | | | | |
| *N-palmitoyl leucine* | | ↓↓↓ | | ↓↓↓ |
| *N-stearoyl leucine* | | ↓↓ | | ↓↓↓ |
| *N-oleoyl leucine* | | ↓↓ | ↓ | ↓↓↓ |
| *N-linoleoyl leucine* | ↓↓ | ↓↓↓ | | ↓↓↓ |
| *N-docosahexaenoyl leucine* | | ↓↓↓↓ | | ↓↓↓ |
| ***N-acyl methionine*** | | | | |
| *N-palmitoyl methionine* | ↓↓ | ↓↓ | | ↓↓↓↓ |
| *N-stearoyl methionine* | | ↓↓↓↓ | | ↓↓↓↓ |
| *N-oleoyl methionine* | ↓ | ↓ | ↓↓ | ↓↓↓ |
| *N-linoleoyl methionine* | ↓↓ | ↓↓↓ | | ↓↓↓↓ |
| *N-arachidonoyl methionine* | | ↓↓ | | ↓↓↓ |
| *N-docosahexaenoyl methionine* | ↓↓ | ↓↓↓ | | ↓↓ |
| ***N- acyl phenylalanine*** | | | | |
| *N-palmitoyl phenylalanine* | | ↓↓ | | ↓↓↓ |
| *N-stearoyl phenylalanine* | ↓↓ | ↓↓ | ↓↓ | ↓↓↓ |
| *N-oleoyl phenylalanine* | | ↓↓ | ↓↓ | ↓↓↓ |
| *N-linoleoyl phenylalanine* | ↓↓ | ↓↓↓ | | ↓↓↓↓ |
| *N-arachidonoyl phenylalanine* | | ↓↓↓ | | ↓↓↓ |
| *N-docosahexaenoyl phenylalanine* | | ↓↓↓↓ | | ↓↓↓ |
| ***N- acyl proline*** | | | | |
| *N-palmitoyl proline* | ↓↓↓ | ↓↓ | ↑↑ | ↑↑ |
| *N-stearoyl proline* | BAL | BAL | BAL | BAL |
| *N-oleoyl proline* | BAL | BAL | BAL | BAL |
| *N-linoleoyl proline* | BAL | BAL | BDL | BAL |
| *N-arachidonoyl proline* | BAL | BAL | BDL | BAL |
| *N-docosahexaenoyl proline* | BDL | BAL | BAL | BDL |

| | VEH | CBD | THC | THC+CBD |
|---|---|---|---|---|
| ***N-acyl serine*** | | | | |
| *N-palmitoyl serine* | | | | ↓↓ |
| *N-stearoyl serine* | | | | ↓↓ |
| *N-oleoyl serine* | ↓↓↓ | | ↓↓ | ↓↓↓ |
| *N-linoleoyl serine* | ↓ | | | ↓↓ |
| *N-arachidonoyl serine* | | ↓↓ | | ↓↓↓ |
| *N-docosahexaenoyl serine* | | ↓↓↓↓ | | ↓↓↓ |
| ***N-acyl taurine*** | | | | |
| *N-palmitoyl taurine* | ↓ | | | |
| *N-stearoyl taurine* | ↓↓ | | ↓↓↓ | ↓ |
| *N-oleoyl taurine* | | | ↓↓↓ | ↑ |
| *N-arachidonoyl taurine* | ↓↓ | | | |
| ***N-acyl tryptophan*** | | | | |
| *N-palmitoyl tryptophan* | BAL | BAL | BAL | BAL |
| *N-stearoyl tryptophan* | BAL | BAL | BAL | BAL |
| *N-oleoyl tryptophan* | BDL | BAL | BDL | BAL |
| *N-linoleoyl tryptophan* | BAL | | BAL | ↓↓ |
| *N-arachidonoyl tryptophan* | BDL | BDL | BDL | BDL |
| *N-docosahexaenoyl tryptophan* | BDL | BDL | BDL | BAL |
| ***N-acyl tyrosine*** | | | | |
| *N-palmitoyl tyrosine* | ↓↓ | ↓↓↓ | | ↓↓↓ |
| *N-stearoyl tyrosine* | ↓↓↓ | ↓↓↓↓ | | ↓↓↓ |
| *N-oleoyl tyrosine* | | ↓↓ | ↓↓ | ↓↓↓ |
| *N-linoleoyl tyrosine* | ↓↓ | ↓↓↓ | | ↓↓↓↓ |
| *N-arachidonoyl tyrosine* | ↓↓ | ↓↓↓ | | ↓↓↓ |
| *N-docosahexaenoyl tyrosine* | | ↓↓↓ | | ↓↓↓ |
| ***N-acyl valine*** | | | | |
| *N-palmitoyl valine* | | ↓↓↓ | ↑ | ↓↓↓ |
| *N-stearoyl valine* | | ↓↓↓↓ | | ↓↓ |
| *N-oleoyl valine* | | ↓↓ | ↓ | ↓↓↓ |
| *N-linoleoyl valine* | ↓↓ | ↓↓↓↓ | | ↓↓↓↓ |
| *N-docosahexaenoyl valine* | ↑↑ | ↓↓↓↓ | | ↓↓↓ |
| **2-acyl-sn-glycerol** | | | | |
| 2-arachidonoyl glycerol | ↑↑↑ | ↑↑ | | ↓ |
| 2-palmitoyl glycerol | | ↑↑ | | |
| 2-oleoyl glycerol | ↑↑↑ | ↑↑↑ | | |
| 2-linoleoyl glycerol | ↑↑ | ↑↑↑ | ↑↑ | |
| **Free Fatty Acids** | | | | |
| Oleic acid | ↑↑ | | ↓ | |
| Linoleic acid | | ↑ | | ↓ |
| Arachidonic acid | ↑↑↑ | ↑ | | |
| Eicosapentaenoic acid | ↑↑↑ | ↑↑↑ | ↑↑↑ | ↑↑ |
| Docosahexaenoic acid | ↑ | | | |
| **Prostaglandins** | | | | |
| $PGE_2$ | | ↓ | ↓ | |
| $PGE_{1\alpha}$ | | | | |
| 6-keto-$PGF_{1\alpha}$ | BAL | BAL | BAL | |

Significance ($p<0.05$) is depicted by dark shades of color, while trending towards significance ($0.05 < p < 0.10$) is depicted by light shades of color. Direction of difference, male relative to female, is depicted by color and arrow direction. When male concentrations are higher than female, the cell is green with up arrows. When male concentrations are lower than female, the cell is orange with down arrows. Fold change is represented by the number of arrows. 1 arrow represents a 1-1.49-fold change, 2 arrows represent a 1.5-1.99-fold difference, 3 arrows represent a 2-2.99-fold difference, 4 arrows represent a 3-9.99-fold difference, and 5 arrows represents a 10-fold or greater difference in males relative to females.

Supplemental Figure 1.3. Heatmaps showing differences between lipids measured in the plasma of female mice 2h after a 10mg/kg injection of CBD, THC, or THC+CBD relative to levels measured in the plasma of female mice 2h after injection of vehicle.

| | Veh → CB | | | CBD → CB | | THC→ |
|---|---|---|---|---|---|---|
| | CBD | THC | THC+CBD | THC | THC + CBD | THC + CBD |
| ***N-acyl alanine*** | | | | | | |
| *N-palmitoyl alanine* | | | | | | |
| *N-stearoyl alanine* | | | | | | |
| *N-oleoyl alanine* | | | | | | |
| *N-linoleoyl alanine* | | | | | ↓ | |
| *N-arachidonoyl alanine* | BAL | BAL | BAL | BDL | BAL | BAL |
| *N-docosahexaenoyl alanine* | BDL | BDL | BDL | BDL | BDL | BDL |
| ***N-acyl ethanolamine*** | | | | | | |
| *N-arachidonoyl ethanolamine* | | | | | | |
| *N-palmitoyl ethanolamine* | | | ↑ | | | |
| *N-stearoyl ethanolamine* | | | ↑ | | | |
| *N-oleoyl ethanolamine* | | | ↑ | | | |
| *N-linoleoyl ethanolamine* | | | ↑ | | ↑ | ↑ |
| *N-docosahexaenoyl ethanolamine* | | | | | | |
| ***N-acyl glycine*** | | | | | | |
| *N-palmitoyl glycine* | | | | | ↑ | |
| *N-stearoyl glycine* | | ↑ | | ↑↑ | | ↓↓ |
| *N-oleoyl glycine* | | | | | | |
| *N-linoleoyl glycine* | | ↓↓ | | | | ↑↑ |
| *N-arachidonoyl glycine* | BDL | BDL | BDL | BDL | BDL | BDL |
| *N-docosahexaenoyl glycine* | BDL | BDL | BDL | BDL | BDL | BDL |
| ***N-acyl leucine*** | | | | | | |
| *N-palmitoyl leucine* | | | | | | |
| *N-stearoyl leucine* | | | | | | |
| *N-oleoyl leucine* | | | | | | |
| *N-linoleoyl leucine* | | | | | | |
| *N-docosahexaenoyl leucine* | | | ↑↑↑ | | ↑↑↑ | |
| ***N-acyl phenylalanine*** | | | | | | |
| *N-palmitoyl phenylalanine* | | | | | | |
| *N-stearoyl phenylalanine* | ↓↓↓ | ↓↓↓ | | | | |
| *N-oleoyl phenylalanine* | | ↓↓ | | | | |
| *N-linoleoyl phenylalanine* | ↓↓ | ↓↓ | | | | |
| *N-arachidonoyl phenylalanine* | ↓↓↓ | ↓↓ | ↓ | | ↑↑ | |
| *N-docosahexaenoyl phenylalanine* | | | | | ↑↑↑ | ↑↑↑ |
| ***N-acyl proline*** | | | | | | |
| *N-palmitoyl proline* | BAL | BAL | BAL | BAL | BAL | BAL |
| *N-stearoyl proline* | BAL | BAL | BDL | BAL | BDL | BAL |
| *N-oleoyl proline* | BDL | BDL | BDL | BDL | BDL | BDL |
| *N-linoleoyl proline* | BAL | BDL | BDL | BAL | BAL | BDL |
| *N-arachidonoyl proline* | BAL | BAL | BAL | BDL | BDL | BDL |
| *N-docosahexaenoyl proline* | BAL | BDL | BDL | BAL | BAL | BDL |

| | Veh → CB | | | CBD → CB | | THC→ |
|---|---|---|---|---|---|---|
| | CBD | THC | THC+CBD | THC | THC + CBD | THC + CBD |
| ***N-acyl serine*** | | | | | | |
| *N-palmitoyl serine* | | | ↓↓ | | | |
| *N-stearoyl serine* | BAL | | | BAL | BAL | |
| *N-oleoyl serine* | BAL | BAL | BDL | BAL | BAL | BAL |
| *N-linoleoyl serine* | BAL | BAL | BDL | BAL | BAL | BAL |
| *N-arachidonoyl serine* | BDL | BDL | BDL | BDL | BDL | BDL |
| *N-docosahexaenoyl serine* | BDL | BDL | BDL | BDL | BDL | BDL |
| ***N-acyl taurine*** | | | | | | |
| *N-palmitoyl taurine* | BAL | BAL | BAL | BAL | BAL | BAL |
| *N-stearoyl taurine* | BAL | BAL | BAL | BAL | BAL | BAL |
| *N-oleoyl taurine* | BAL | BAL | BAL | | | |
| *N-arachidonoyl taurine* | BAL | BDL | BDL | BAL | BAL | BDL |
| ***N-acyl tyrosine*** | | | | | | |
| *N-palmitoyl tyrosine* | | ↓↓ | | | | |
| *N-stearoyl tyrosine* | ↓↓ | | ↓↓ | | | |
| *N-oleoyl tyrosine* | | ↓↓↓↓ | ↓↓↓ | | | |
| *N-linoleoyl tyrosine* | | ↓↓↓ | | | | |
| *N-arachidonoyl tyrosine* | BAL | BAL | BAL | BAL | BAL | BAL |
| *N-docosahexaenoyl tyrosine* | | BAL | | BAL | | BAL |
| ***N-acyl valine*** | | | | | | |
| *N-palmitoyl valine* | ↓↓ | ↓↓↓ | | | | ↑↑ |
| *N- stearoyl valine* | ↓↓ | ↓↓ | | | | |
| *N-oleoyl valine* | | | | | | |
| *N-linoleoyl valine* | ↓↓ | ↓↓↓ | | | | |
| *N-docosahexaenoyl valine* | BAL | BAL | BAL | BAL | BAL | BAL |
| **2-acyl-sn-glycerol** | | | | | | |
| 2-arachidonoyl-sn-glycerol | | | | | | |
| 2-palmitoyl-sn-glycerol | | | | | ↓↓ | |
| 2-oleoyl-sn-glycerol | | | | | ↓↓↓ | |
| 2-linoleoyl-sn-glycerol | | ↓↓↓ | ↓↓ | ↓↓↓ | ↓↓↓ | |
| **Free Fatty Acids** | | | | | | |
| Oleic acid | | | | | | |
| Linoleic acid | | ↓ | | ↓ | | |
| Arachidonic acid | | | | ↑ | ↑↑ | |
| Eicosapentaenoic acid | | | | | | |
| Docosahexaenoic acid | ↓↓ | | | ↑ | ↑↑ | |

Significance ($p<0.05$) is depicted by dark shades of color, while trending towards significance ($0.05 < p < 0.10$) is depicted by light shades of color. Direction of difference, female CB relative to female vehicle, is depicted by color and arrow direction. When mean CB treatment group concentrations are higher than mean vehicle group concentrations, the cell is green with up arrows. When mean CB group concentrations are lower than mean vehicle concentrations, the cell is orange with down arrows. Fold-change is represented by the number of arrows. 1 arrow represents a 1-1.49-fold change, 2 arrows represent a 1.5-1.99-fold difference, 3 arrows represent a 2-2.99-fold difference, 4 arrows represent a 3-9.99-fold difference, and 5 arrows represents a 10-fold or greater difference in CB relative to vehicle.

Supplemental Figure 1.4. Heatmaps showing differences between lipids measured in the liver of female mice 2h after a 10mg/kg injection of CBD, THC, or THC+CBD relative to levels measured in the liver of female mice 2h after injection of vehicle.

| | Veh → CB | | | CBD → CB | | THC→ |
|---|---|---|---|---|---|---|
| | CBD | THC | THC+CBD | THC | THC + CBD | THC + CBD |
| ***N-acyl alanine*** | | | | | | |
| *N-palmitoyl alanine* | | ↑ | ↑ | | | |
| *N-stearoyl alanine* | ↑↑ | | | ↓ | ↓ | |
| *N-oleoyl alanine* | | ↑↑ | ↑↑ | ↑↑ | ↑ | |
| *N-linoleoyl alanine* | | ↑ | ↑↑ | | ↑↑ | ↑ |
| *N-arachidonoyl alanine* | | ↑ | ↑ | ↑ | | |
| *N-docosahexaenoyl alanine* | ↑↑ | ↑↑ | ↑↑ | | | |
| ***N-acyl ethanolamine*** | | | | | | |
| *N-arachidonoyl ethanolamine* | | ↑↑↑ | ↑↑↑↑ | ↑↑↑ | ↑↑↑↑ | ↑↑ |
| *N-palmitoyl ethanolamine* | | ↑↑↑ | ↑↑↑ | ↑↑ | ↑↑ | |
| *N-stearoyl ethanolamine* | | ↑↑↑ | ↑↑↑ | ↑↑ | ↑↑↑ | |
| *N-oleoyl ethanolamine* | | ↑↑↑↑ | ↑↑↑↑ | ↑↑↑↑ | ↑↑↑↑ | |
| *N-linoleoyl ethanolamine* | | ↑↑↑ | ↑↑↑↑ | ↑↑↑ | ↑↑↑↑ | ↑↑↑ |
| *N-docosahexaenoyl ethanolamine* | ↑↑↑ | ↓↓↓ | ↓↓↓ | ↓↓↓↓ | ↓↓↓↓ | |
| ***N-acyl glycine*** | | | | | | |
| *N-palmitoyl glycine* | | | ↑↑ | | ↑ | ↑ |
| *N-stearoyl glycine* | ↑↑ | ↑↑ | ↑↑ | | | |
| *N-oleoyl glycine* | | ↑↑ | ↑↑↑ | ↑↑↑ | ↑↑↑ | |
| *N-linoleoyl glycine* | | ↑↑ | ↑↑↑ | | ↑↑ | ↑↑ |
| *N-arachidonoyl glycine* | | ↑↑ | ↑↑↑ | ↑ | ↑↑ | |
| *N-docosahexaenoyl glycine* | ↑↑ | ↑↑ | ↑↑↑ | | ↑ | |
| ***N-acyl GABA*** | | | | | | |
| *N-palmitoyl GABA* | ↑↑↑ | | ↑↑↑ | ↓↓ | | ↑↑ |
| *N-stearoyl GABA* | | | | | | |
| *N-oleoyl GABA* | ↑↑ | ↑↑ | ↑↑↑ | | | |
| *N-linoleoyl GABA* | ↑↑↑↑ | | ↑↑↑ | ↓↓↓↓ | | ↑↑↑ |
| *N-arachidonoyl GABA* | | | ↑↑ | | | |
| *N-docosahexaenoyl GABA* | | ↑↑↑ | ↑↑↑ | | | |
| ***N-acyl leucine*** | | | | | | |
| *N-palmitoyl leucine* | | | ↑↑ | | | ↑ |
| *N-stearoyl leucine* | | | ↑↑↑ | | ↑↑ | ↑↑ |
| *N-oleoyl leucine* | | ↑↑ | ↑↑↑ | ↑↑ | ↑↑ | ↑ |
| *N-linoleoyl leucine* | ↑↑ | | ↑↑↑ | | | ↑↑ |
| *N-docosahexaenoyl leucine* | | | ↑↑↑ | | | ↑ |
| ***N-acyl methionine*** | | | | | | |
| *N-palmitoyl methionine* | | | ↑↑ | | ↑↑ | ↑ |
| *N-stearoyl methionine* | ↑↑↑ | | ↑↑ | ↓↓ | | ↑↑ |
| *N-oleoyl methionine* | | ↑↑ | ↑↑ | ↑↑ | ↑↑ | |
| *N-linoleoyl methionine* | | | ↑↑↑ | | ↑↑ | ↑↑↑ |
| *N-arachidonoyl methionine* | | | ↑↑ | | | |
| *N-docosahexaenoyl methionine* | | | ↑↑↑ | | ↑↑ | ↑↑ |
| ***N*- acyl phenylalanine** | | | | | | |
| *N-palmitoyl phenylalanine* | | | ↑↑ | | ↑ | ↑ |
| *N-stearoyl phenylalanine* | | | | | | |
| *N-oleoyl phenylalanine* | | ↑↑ | ↑↑↑ | ↑↑ | ↑↑ | |
| *N-linoleoyl phenylalanine* | | | ↑↑↑ | | ↑↑ | ↑↑↑ |
| *N-arachidonoyl phenylalanine* | ↑↑ | | ↑↑ | | | ↑ |
| *N-docosahexaenoyl phenylalanine* | ↑↑ | ↑↑ | ↑↑↑ | | | ↑ |
| ***N*- acyl proline** | | | | | | |
| *N-palmitoyl proline* | | ↓↓ | ↓↓↓ | ↓↓ | ↓↓↓ | |
| *N-stearoyl proline* | BAL | BAL | BAL | BAL | BAL | BAL |
| *N-oleoyl proline* | ↑↑ | BAL | | BAL | | BAL |
| *N-linoleoyl proline* | BAL | BDL | BAL | BDL | BAL | BAL |
| *N-arachidonoyl proline* | BAL | BDL | BAL | BDL | BAL | BAL |
| *N-docosahexaenoyl proline* | BAL | BDL | BDL | BDL | BDL | BDL |

| | Veh → CB | | | CBD → CB | | THC→ |
|---|---|---|---|---|---|---|
| | CBD | THC | THC+CBD | THC | THC + CBD | THC + CBD |
| ***N-acyl serine*** | | | | | | |
| *N-palmitoyl serine* | | ↑↑ | ↑↑↑ | | | |
| *N-stearoyl serine* | | | | | | |
| *N-oleoyl serine* | | ↑↑↑ | ↑↑↑ | ↑↑↑ | ↑↑ | |
| *N-linoleoyl serine* | | | ↑↑↑ | | ↑↑ | ↑↑ |
| *N-arachidonoyl serine* | ↑↑↑ | ↑↑↑ | ↑↑↑↑ | | | |
| *N-docosahexaenoyl serine* | ↑↑↑ | | ↑↑↑ | | | ↑↑↑ |
| ***N-acyl taurine*** | | | | | | |
| *N-palmitoyl taurine* | | ↑ | | ↑↑ | | |
| *N-stearoyl taurine* | | ↑↑ | ↑↑↑ | ↑↑ | ↑↑↑ | |
| *N-oleoyl taurine* | | ↑↑↑ | ↑↑ | ↑↑↑↑ | ↑↑↑ | ↓ |
| *N-arachidonoyl taurine* | | ↑↑↑ | ↑↑↑ | ↑↑↑ | ↑↑↑ | |
| ***N-acyl tryptophan*** | | | | | | |
| *N-palmitoyl tryptophan* | BAL | BAL | BAL | BAL | BAL | BAL |
| *N-stearoyl tryptophan* | BAL | BAL | BAL | BDL | BAL | BAL |
| *N-oleoyl tryptophan* | BAL | BDL | BAL | BAL | BAL | BAL |
| *N-linoleoyl tryptophan* | BAL | BAL | BAL | | | |
| *N-arachidonoyl tryptophan* | BDL | BDL | BDL | BDL | BDL | BDL |
| *N-docosahexaenoyl tryptophan* | BDL | BDL | BAL | BDL | BAL | BAL |
| ***N-acyl tyrosine*** | | | | | | |
| *N-palmitoyl tyrosine* | | | ↑ | | ↑ | ↑ |
| *N-stearoyl tyrosine* | ↑↑↑ | | ↑↑ | ↓↓ | | ↑↑ |
| *N-oleoyl tyrosine* | | ↑↑↑ | ↑↑↑ | ↑↑ | ↑↑ | |
| *N-linoleoyl tyrosine* | | | ↑↑↑ | | ↑↑ | ↑↑ |
| *N-arachidonoyl tyrosine* | | ↑ | ↑↑ | | | |
| *N-docosahexaenoyl tyrosine* | | ↑ | ↑↑ | | ↑↑ | ↑ |
| ***N-acyl valine*** | | | | | | |
| *N-palmitoyl valine* | | | ↑↑ | | | ↑↑ |
| *N- stearoyl valine* | ↑↑↑ | | | | | |
| *N-oleoyl valine* | | ↑↑ | ↑↑↑ | ↑↑ | ↑↑↑ | ↑ |
| *N-linoleoyl valine* | | | ↑↑↑ | | ↑ | ↑↑↑ |
| *N-docosahexaenoyl valine* | ↑↑↑ | ↑↑↑↑ | ↑↑↑↑ | | ↑ | |
| **2-acyl-sn-glycerol** | | | | | | |
| 2-arachidonoyl-sn-glycerol | | ↑↑↑ | ↑↑↑↑ | ↑↑ | ↑↑↑ | ↑ |
| 2-palmitoyl-sn-glycerol | | ↑↑ | ↑↑↑ | | ↑↑ | |
| 2-oleoyl-sn-glycerol | | ↑↑↑↑ | ↑↑↑↑ | ↑↑↑↑ | ↑↑↑↑ | |
| 2-linoleoyl-sn-glycerol | | ↑↑ | ↑↑↑↑ | | ↑↑↑↑ | ↑↑ |
| **Free Fatty Acids** | | | | | | |
| Oleic acid | | ↑↑ | ↑↑ | ↑↑ | ↑↑ | |
| Linoleic acid | | ↑↑ | ↑↑↑ | ↑↑ | ↑↑↑ | ↑↑ |
| Arachidonic acid | ↑↑ | ↑↑↑ | ↑↑↑ | ↑ | ↑↑ | |
| Eicosapentaenoic acid | | | ↑↑↑ | | ↑↑↑↑ | ↑↑ |
| Docosahexaenoic acid | ↑↑ | ↑↑ | ↑↑↑ | | ↑ | ↑ |
| **Prostaglandins** | | | | | | |
| $PGE_2$ | ↑↑ | ↑↑↑ | ↑↑ | ↑ | | ↓ |
| $PGE_{1\alpha}$ | ↑↑ | ↑↑ | | | ↓ | ↓ |
| 6-keto-$PGF_{1\alpha}$ | | ↑↑↑ | | | | |

Significance (p<0.05) is depicted by dark shades of color, while trending towards significance (0.05 <p < 0.10) is depicted by light shades of color. Direction of difference, female CB relative to female vehicle, is depicted by color and arrow direction. When mean CB treatment group concentrations are higher than mean vehicle group concentrations, the cell is green with up arrows. When mean CB group concentrations are lower than mean vehicle concentrations, the cell is orange with down arrows. Fold-change is represented by the number of arrows. 1 arrow represents a 1-1.49-fold change, 2 arrows represent a 1.5-1.99-fold difference, 3 arrows represent a 2-2.99-fold difference, 4 arrows represent a 3-9.99-fold difference, and 5 arrows represents a 10-fold or greater difference in CB relative to vehicle.

Supplemental Figure 1.5. Heatmaps of differences between lipids measured in the plasma of male mice 2h after a 10mg/kg injection of CBD, THC, or THC+CBD relative to levels measured in the plasma of male mice 2h after injection of vehicle.

| | Veh → CB | | | CBD → CB | | THC→ |
|---|---|---|---|---|---|---|
| | CBD | THC | THC+CBD | THC | THC + CBD | THC + CBD |
| ***N-acyl alanine*** | | | | | | |
| *N-palmitoyl alanine* | | | | | | |
| *N-stearoyl alanine* | | | | | | |
| *N-oleoyl alanine* | | | | | | |
| *N-linoleoyl alanine* | | | | | | |
| *N-arachidonoyl alanine* | BDL | BDL | BDL | BDL | BDL | BDL |
| *N-docosahexaenoyl alanine* | BAL | BDL | BDL | BAL | BAL | BDL |
| ***N-acyl ethanolamine*** | | | | | | |
| *N-arachidonoyl ethanolamine* | | | | | | |
| *N-palmitoyl ethanolamine* | | | | ↓ | | ↑ |
| *N-stearoyl ethanolamine* | | | | | | |
| *N-oleoyl ethanolamine* | | | | | ↑ | ↑ |
| *N-linoleoyl ethanolamine* | | | | | | |
| *N-docosahexaenoyl ethanolamine* | | ↑ | | | | ↓ |
| ***N-acyl glycine*** | | | | | | |
| *N-palmitoyl glycine* | ↑ | | | | | |
| *N-stearoyl glycine* | | | | | | |
| *N-oleoyl glycine* | | | | | | |
| *N-linoleoyl glycine* | | ↑↑ | | ↑ | | |
| *N-arachidonoyl glycine* | BDL | BAL | BDL | BAL | BDL | BAL |
| *N-docosahexaenoyl glycine* | BDL | BDL | BDL | BDL | BDL | BDL |
| ***N-acyl leucine*** | | | | | | |
| *N-palmitoyl leucine* | | | ↑↑ | | ↑↑ | |
| *N-stearoyl leucine* | | | | | ↑↑ | |
| *N-oleoyl leucine* | | | | | ↑↑ | |
| *N-linoleoyl leucine* | | | | | | |
| *N-docosahexaenoyl leucine* | | | | | | |
| ***N-acyl phenylalanine*** | | | | | | |
| *N-palmitoyl phenylalanine* | | | ↑↑ | | ↑↑↑ | |
| *N-stearoyl phenylalanine* | | | ↑↑↑ | | ↑↑ | |
| *N-oleoyl phenylalanine* | | | ↑↑ | | ↑↑↑ | ↑↑ |
| *N-linoleoyl phenylalanine* | | | | | | |
| *N-arachidonoyl phenylalanine* | | | | | | |
| *N-docosahexaenoyl phenylalanine* | | | | | | |
| ***N-acyl proline*** | | | | | | |
| *N-palmitoyl proline* | BAL | BAL | BAL | | BAL | BAL |
| *N-stearoyl proline* | BAL | BAL | BAL | BAL | BAL | BAL |
| *N-oleoyl proline* | BDL | BAL | BDL | BAL | BDL | BAL |
| *N-linoleoyl proline* | BAL | BDL | BDL | BAL | BAL | BDL |
| *N-arachidonoyl proline* | BDL | BDL | BDL | BDL | BDL | BDL |
| *N-docosahexaenoyl proline* | BDL | BDL | BDL | BDL | BAL | BAL |

| | Veh → CB | | | CBD → CB | | THC→ |
|---|---|---|---|---|---|---|
| | CBD | THC | THC+CBD | THC | THC + CBD | THC + CBD |
| ***N-acyl serine*** | | | | | | |
| *N-palmitoyl serine* | | | | ↓↓ | | |
| *N-stearoyl serine* | BAL | BAL | BAL | BAL | BAL | BAL |
| *N-oleoyl serine* | BAL | BAL | BAL | BAL | BAL | BAL |
| *N-linoleoyl serine* | BDL | BAL | BDL | BAL | BDL | BAL |
| *N-arachidonoyl serine* | BAL | BAL | BDL | BAL | BDL | BAL |
| *N-docosahexaenoyl serine* | BDL | BDL | BDL | BDL | BDL | BDL |
| ***N-acyl taurine*** | | | | | | |
| *N-palmitoyl taurine* | BAL | BAL | BAL | BAL | BAL | BAL |
| *N-stearoyl taurine* | BAL | BAL | BAL | BAL | BAL | BAL |
| *N-oleoyl taurine* | BAL | BAL | BAL | BAL | BAL | BAL |
| *N-arachidonoyl taurine* | BDL | BDL | BDL | BDL | BDL | BDL |
| ***N-acyl tyrosine*** | | | | | | |
| *N-palmitoyl tyrosine* | | | | ↑↑ | ↑↑↑ | |
| *N-stearoyl tyrosine* | | | | | | |
| *N-oleoyl tyrosine* | | | | | ↑↑ | |
| *N-linoleoyl tyrosine* | | | | | ↑↑ | |
| *N-arachidonoyl tyrosine* | BAL | BAL | BAL | BAL | BAL | BAL |
| *N-docosahexaenoyl tyrosine* | BAL | BAL | BAL | BAL | BAL | BAL |
| ***N-acyl valine*** | | | | | | |
| *N-palmitoyl valine* | | | ↑↑↑ | ↑↑ | ↑↑↑ | |
| *N- stearoyl valine* | | | | | | |
| *N-oleoyl valine* | | | | | ↑↑↑ | |
| *N-linoleoyl valine* | | | | ↑↑ | | |
| *N-docosahexaenoyl valine* | BAL | BAL | BAL | BAL | BAL | BAL |
| **2-acyl-sn-glycerol** | | | | | | |
| 2-arachidonoyl-sn-glycerol | | ↓↓↓ | ↓↓ | ↓↓↓ | ↓ | ↑↑ |
| 2-palmitoyl-sn-glycerol | ↑ | ↓↓ | | ↓↓↓ | ↓↓ | |
| 2-oleoyl-sn-glycerol | | ↓↓↓↓ | | ↓↓↓↓ | | ↑↑↑↑ |
| 2-linoleoyl-sn-glycerol | ↑ | ↓↓↓ | | ↓↓↓↓ | ↓↓ | ↑↑ |
| **Free Fatty Acids** | | | | | | |
| Oleic acid | | | | ↓ | | ↑ |
| Linoleic acid | | | | | | |
| Arachidonic acid | | | | | | |
| Eicosapentaenoic acid | | | | ↓↓ | | ↑↑ |
| Docosahexaenoic acid | ↓ | | | | | |

Significance ($p<0.05$) is depicted by dark shades of color, while trending towards significance ($0.05 < p < 0.10$) is depicted by light shades of color. Direction of difference, male CB relative to male vehicle, is depicted by color and arrow direction. When mean CB treatment group concentrations are higher than mean vehicle group concentrations, the cell is green with up arrows. When mean CB group concentrations are lower than mean vehicle concentrations, the cell is orange with down arrows. Fold-change is represented by the number of arrows. 1 arrow represents a 1-1.49-fold change, 2 arrows represent a 1.5-1.99-fold difference, 3 arrows represent a 2-2.99-fold difference, 4 arrows represent a 3-9.99-fold difference, and 5 arrows represents a 10-fold or greater difference in CB relative to vehicle.

Supplemental Figure 1.6. Heatmaps of differences between lipids measured in the liver of male mice 2h after a 10mg/kg injection of CBD, THC, or THC+CBD relative to levels measured in the liver of male mice 2h after injection of vehicle.

| | Veh → CB | | | CBD → CB | | THC→ |
|---|---|---|---|---|---|---|
| | CBD | THC | THC+CBD | THC | THC + CBD | THC + CBD |
| ***N-acyl alanine*** | | | | | | |
| *N-palmitoyl alanine* | | ↑ | | ↑↑ | | ↓ |
| *N-stearoyl alanine* | | | | | | |
| *N-oleoyl alanine* | | | | | ↑ | |
| *N-linoleoyl alanine* | | ↑↑↑ | ↑ | ↑↑ | | ↓ |
| *N-arachidonoyl alanine* | | ↑↑ | | ↑↑ | | ↓ |
| *N-docosahexaenoyl alanine* | | ↑↑ | | ↑↑↑ | ↑↑ | ↓ |
| ***N-acyl ethanolamine*** | | | | | | |
| *N-arachidonoyl ethanolamine* | | ↑↑↑↑ | ↑↑↑↑ | ↑↑↑↑ | ↑↑↑↑ | |
| *N-palmitoyl ethanolamine* | | | | | | |
| *N-stearoyl ethanolamine* | ↓↓ | | | | | |
| *N-oleoyl ethanolamine* | | | | ↑↑ | ↑↑ | |
| *N-linoleoyl ethanolamine* | | ↑↑↑ | ↑↑↑ | ↑↑↑ | ↑↑↑ | |
| *N-docosahexaenoyl ethanolamine* | | | ↑↑↑↑↑ | | ↑↑↑↑↑ | ↑↑↑↑ |
| ***N-acyl GABA*** | | | | | | |
| *N-palmitoyl GABA* | | ↑↑↑ | | ↑↑↑↑ | | ↓↓↓ |
| *N-stearoyl GABA* | | | | | ↓↓ | |
| *N-oleoyl GABA* | | | | | | |
| *N-linoleoyl GABA* | BAL | BAL | BAL | ↑↑↑↑ | | |
| *N-arachidonoyl GABA* | BAL | BAL | BAL | BAL | BAL | |
| *N-docosahexaenoyl GABA* | BAL | BAL | BAL | ↑↑ | | |
| ***N-acyl glycine*** | | | | | | |
| *N-palmitoyl glycine* | | ↑↑ | | ↑↑ | | ↓↓ |
| *N-stearoyl glycine* | | ↑↑ | | | | ↓↓ |
| *N-oleoyl glycine* | | ↑↑ | ↑↑↑ | ↑ | ↑↑ | |
| *N-linoleoyl glycine* | | ↑↑↑ | ↑↑↑↑ | ↑↑ | ↑↑↑ | |
| *N-arachidonoyl glycine* | | ↑↑↑ | ↑↑ | ↑↑ | ↑ | |
| *N-docosahexaenoyl glycine* | | ↑↑↑ | ↑↑↑ | ↑↑↑ | ↑↑ | |
| ***N-acyl leucine*** | | | | | | |
| *N-palmitoyl leucine* | | ↑↑ | | ↑↑↑ | | ↓↓↓ |
| *N-stearoyl leucine* | ↓↓↓ | | | ↑↑↑ | | |
| *N-oleoyl leucine* | | | | ↑↑ | | |
| *N-linoleoyl leucine* | | ↑↑↑ | | ↑↑↑ | | ↓↓ |
| *N-docosahexaenoyl leucine* | | ↑↑ | | ↑↑↑↑ | | ↓↓ |
| ***N-acyl methionine*** | | | | | | |
| *N-palmitoyl methionine* | | ↑↑↑ | | ↑↑↑ | | ↓↓↓ |
| *N-stearoyl methionine* | | | | | | ↓↓↓ |
| *N-oleoyl methionine* | | ↑↑ | | ↑↑ | | |
| *N-linoleoyl methionine* | | ↑↑↑ | | ↑↑↑ | | ↓↓↓ |
| *N-arachidonoyl methionine* | | ↑↑ | | ↑↑ | | ↓↓ |
| *N-docosahexaenoyl methionine* | | ↑↑↑↑ | ↑↑↑ | ↑↑↑↑ | ↑↑↑ | |
| ***N- acyl phenylalanine*** | | | | | | |
| *N-palmitoyl phenylalanine* | | ↑ | | ↑↑ | | ↓↓ |
| *N-stearoyl phenylalanine* | | | | | | |
| *N-oleoyl phenylalanine* | | | | | | |
| *N-linoleoyl phenylalanine* | | ↑↑ | | ↑↑↑ | | ↓↓ |
| *N-arachidonoyl phenylalanine* | | ↑↑ | | ↑↑↑ | | ↓↓↓ |
| *N-docosahexaenoyl phenylalanine* | | ↑↑ | | ↑↑↑ | | ↓↓ |
| ***N- acyl proline*** | | | | | | |
| *N-palmitoyl proline* | | ↑↑↑ | ↑↑↑ | | | |
| *N-stearoyl proline* | BAL | BAL | BAL | BAL | BAL | BAL |
| *N-oleoyl proline* | BAL | BAL | BAL | BAL | BAL | BAL |
| *N-linoleoyl proline* | | BDL | BAL | BDL | BAL | BAL |
| *N-arachidonoyl proline* | BDL | BDL | BDL | BDL | BDL | BDL |
| *N-docosahexaenoyl proline* | BAL | BAL | BDL | BAL | BDL | BDL |

| | Veh → CB | | | CBD → CB | | THC→ |
|---|---|---|---|---|---|---|
| | CBD | THC | THC+CBD | THC | THC + CBD | THC + CBD |
| ***N-acyl serine*** | | | | | | |
| *N-palmitoyl serine* | | ↑↑↑ | | ↑↑ | | ↓↓ |
| *N-stearoyl serine* | | | | | | |
| *N-oleoyl serine* | | ↑↑↑↑ | ↑↑↑ | ↑↑ | | |
| *N-linoleoyl serine* | | ↑↑↑ | | ↑↑↑ | | |
| *N-arachidonoyl serine* | | ↑↑↑↑ | | | | |
| *N-docosahexaenoyl serine* | | | | ↑↑↑↑ | | |
| ***N-acyl taurine*** | | | | | | |
| *N-palmitoyl taurine* | | ↑↑ | ↑↑ | ↑ | ↑ | |
| *N-stearoyl taurine* | | | | | | |
| *N-oleoyl taurine* | | | ↑↑↑ | | ↑↑↑ | ↑↑↑ |
| *N-arachidonoyl taurine* | | ↑↑↑↑ | ↑↑↑↑ | ↑↑↑ | ↑↑↑ | |
| ***N-acyl tryptophan*** | | | | | | |
| *N-palmitoyl tryptophan* | BAL | BAL | BAL | BAL | BAL | BAL |
| *N-stearoyl tryptophan* | BAL | BAL | BAL | BAL | BAL | BAL |
| *N-oleoyl tryptophan* | BDL | BDL | BDL | BDL | BDL | BDL |
| *N-linoleoyl tryptophan* | BAL | BAL | BAL | BAL | | BAL |
| *N-arachidonoyl tryptophan* | BDL | BDL | BDL | BDL | BDL | BDL |
| *N-docosahexaenoyl tryptophan* | BDL | BDL | BDL | BDL | BDL | BDL |
| ***N-acyl tyrosine*** | | | | | | |
| *N-palmitoyl tyrosine* | | ↑↑ | | ↑↑↑ | | ↓↓↓ |
| *N-stearoyl tyrosine* | | ↑↑↑ | | ↑↑↑ | | |
| *N-oleoyl tyrosine* | | | | | | |
| *N-linoleoyl tyrosine* | | ↑↑↑ | | ↑↑↑ | | ↓↓ |
| *N-arachidonoyl tyrosine* | | ↑↑ | | ↑↑↑ | | ↓↓↓ |
| *N-docosahexaenoyl tyrosine* | | ↑↑ | | ↑↑↑ | | ↓↓ |
| ***N-acyl valine*** | | | | | | |
| *N-palmitoyl valine* | | ↑↑ | | ↑↑↑ | | ↓↓↓ |
| *N- stearoyl valine* | | | | ↑↑↑ | | |
| *N-oleoyl valine* | | | | ↑↑↑ | | |
| *N-linoleoyl valine* | | ↑↑↑ | | ↑↑↑↑ | | ↓↓ |
| *N-docosahexaenoyl valine* | | ↑↑↑ | | ↑↑↑↑ | | ↓↓ |
| **2-acyl-sn-glycerol** | | | | | | |
| 2-arachidonoyl-sn-glycerol | | | | | | |
| 2-palmitoyl-sn-glycerol | | ↑↑ | | | | |
| 2-oleoyl-sn-glycerol | | | | | | |
| 2-linoleoyl-sn-glycerol | | ↑↑ | ↑ | | | |
| **Free Fatty Acids** | | | | | | |
| Oleic acid | | | | | | ↑ |
| Linoleic acid | | ↑↑ | ↑↑ | ↑↑ | ↑↑ | |
| Arachidonic acid | | | ↑ | | ↑ | |
| Eicosapentaenoic acid | | ↑↑ | ↑↑↑ | ↑↑ | ↑↑↑ | ↑↑ |
| Docosahexaenoic acid | | ↑ | ↑↑ | ↑ | ↑↑ | ↑ |
| **Prostaglandins** | | | | | | |
| $PGE_2$ | | ↑↑↑ | ↑↑ | | | |
| $PGE_{1\alpha}$ | | | | | | ↓ |
| 6-keto-$PGF_{1\alpha}$ | BDL | BAL | BAL | BAL | BAL | BAL |

Significance ($p<0.05$) is depicted by dark shades of color, while trending towards significance ($0.05 < p < 0.10$) is depicted by light shades of color. Direction of difference, male CB relative to male vehicle, is depicted by color and arrow direction. When mean CB treatment group concentrations are higher than mean vehicle group concentrations, the cell is green with up arrows. When mean CB group concentrations are lower than mean vehicle concentrations, the cell is orange with down arrows. Fold-change is represented by the number of arrows. 1 arrow represents a 1-1.49-fold change, 2 arrows represent a 1.5-1.99-fold difference, 3 arrows represent a 2-2.99-fold difference, 4 arrows represent a 3-9.99-fold difference, and 5 arrows represents a 10-fold or greater difference in CB relative to vehicle.

Supplemental Figure 1.7. Analysis of *N*-acyl tyrosine lipids in plasma.

A

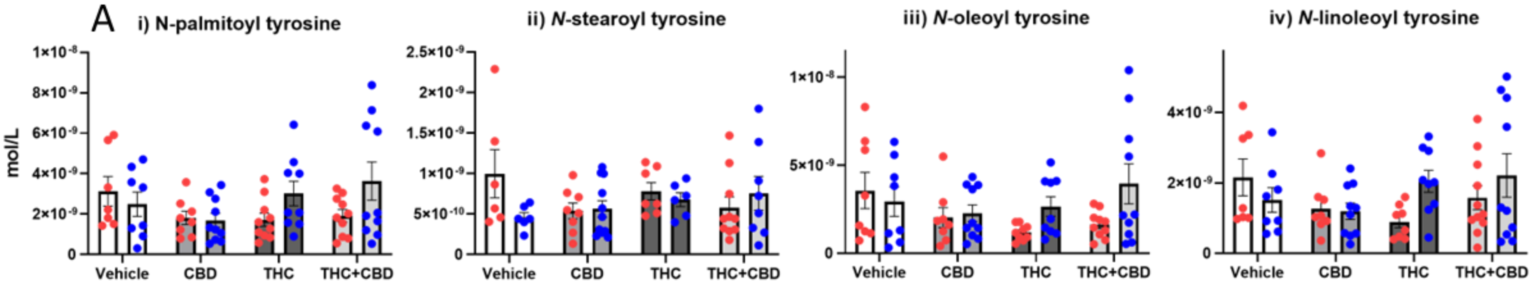

B

| | Female → Male | | | |
|---|---|---|---|---|
| | VEH | CBD | THC | T+C |
| PAL-Tyr | | | ↑↑ | ↑↑ |
| STE-Tyr | ↓↓↓ | | | |
| OLE-Tyr | | | ↑↑↑ | ↑↑↑ |
| LIN-Tyr | | | ↑↑↑ | |
| ARA-Tyr | BAL | BAL | BAL | BAL |
| DOC-Tyr | | BAL | BAL | BAL |

C

| | Female Veh → CB | | | CBD → CB | | THC |
|---|---|---|---|---|---|---|
| | CBD | THC | T+C | THC | T+C | T+C |
| PAL-Tyr | | ↓↓ | | | | |
| STE-Tyr | ↓↓ | | ↓↓ | | | |
| OLE-Tyr | | ↓↓↓↓ | ↓↓↓ | | | |
| LIN-Tyr | | ↓↓↓ | | | | |
| ARA-Tyr | BAL | BAL | BAL | BAL | BAL | BAL |
| DOC-Tyr | | | | | | |

D

| | Male Veh → CB | | | CBD → CB | | THC |
|---|---|---|---|---|---|---|
| | CBD | THC | T+C | THC | T+C | T+C |
| PAL-Tyr | | | | ↑↑ | ↑↑↑ | |
| STE-Tyr | | | | | | |
| OLE-Tyr | | | | | ↑↑ | |
| LIN-Tyr | | | | | ↑↑ | |
| ARA-Tyr | BAL | BAL | BAL | BAL | BAL | BAL |
| DOC-Tyr | BAL | BAL | BAL | BAL | BAL | BAL |

**A)** Average measured concentrations (mol/L) for each *N*-acyl tyrosine: i) *N*-palmitoyl tyrosine (PAL-tyr), ii) *N*-stearoyl tyrosine (PAL-tyr), iii) *N*-oleoyl tyrosine (OLE-tyr), iv) *N*-arachidonoyl tyrosine (ARA-tyr). Data are shown as SEM with individual measurements from male (blue) and female (red) animals to illustrate distribution. **B)** Heatmaps illustrating *N*-acyl tyrosines in males compared to (→) females across treatment groups. **C)** Heatmaps for changes in *N*-acyl tyrosine levels from female vehicle → CB treatment group; CBD → THC and T+C; and THC → T+C. **D)** Heatmaps for changes in *N*-acyl tyrosine levels from male vehicle → CB treatment group; CBD → THC and T+C; and THC → T+C. Green indicates significant increases, orange indicates significant decreases, arrows indicate effect size.

Supplemental Figure 1.8. Analysis of *N*-acyl valine lipids in plasma.

A

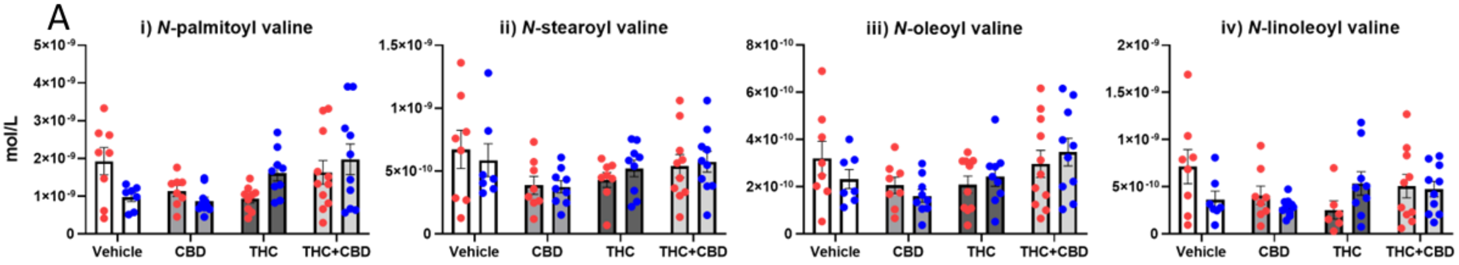

B

| | Female → Male | | | |
|---|---|---|---|---|
| | VEH | CBD | THC | T+C |
| PAL-Val | ↓↓ | ↓ | ↑↑ | |
| STE-Val | | | | |
| OLE-Val | | | | |
| LIN-Val | ↓↓ | | ↑↑↑ | |
| DOC-Val | BAL | BAL | BAL | BAL |

C

| | Female Veh → CB | | | CBD → CB | | THC |
|---|---|---|---|---|---|---|
| | CBD | THC | T+C | THC | T+C | T+C |
| PAL-Val | ↓↓ | ↓↓↓ | | | | ↑↑ |
| STE-Val | ↓↓ | ↓↓ | | | | |
| OLE-Val | | | | | | |
| LIN-Val | ↓↓ | ↓↓↓ | | | | |
| DOC-Val | BAL | BAL | BAL | BAL | BAL | BAL |

D

| | Male Veh → CB | | | CBD → CB | | THC |
|---|---|---|---|---|---|---|
| | CBD | THC | T+C | THC | T+C | T+C |
| PAL-Val | | | ↑↑↑ | ↑↑ | ↑↑ | |
| STE-Val | | | | | | |
| OLE-Val | | | | | ↑↑↑ | |
| LIN-Val | | | | ↑↑ | | |
| DOC-Val | BAL | BAL | BAL | BAL | BAL | BAL |

**A)** Average measured concentrations (mol/L) for each *N*-acyl valine species: i) *N*-palmitoyl valine (PAL-val), ii) *N*-stearoyl valine (STE-val), iii) *N*-oleoyl valine (OLE-val), iv) *N*-linoleoyl valine (OLE-val). Data are shown as SEM with individual measurements from male (blue) and female (red) animals to illustrate distribution. **B)** Heatmaps illustrating *N*-acyl valines in males compared to (→) females across treatment groups. **C)** Heatmaps for changes in *N*-acyl valine levels from female vehicle → CB treatment group; CBD → THC and T+C; and THC → T+C. **D)** Heatmaps for changes in *N*-acyl valine levels from male vehicle → CB treatment group; CBD → THC and T+C; and THC → T+C. Green indicates significant increases, orange indicates significant decreases, arrows indicate effect size.

Supplemental Table 2.1 Descriptive statistics for cannabinoids and metabolites (mol/L) measured in plasma collected from male and female adult mice 2h after receiving an i.p. injection of 10mg/kg CBD, THC, or THC+CBD

| | | N | Mean | Std. Deviation | Std. Error | 95% Confidence Interval for Mean | | Minimum | Maximum |
|---|---|---|---|---|---|---|---|---|---|
| | | | | | | Lower Bound | Upper Bound | | |
| CBD | CBD F | 9 | 7.1911E-07 | 2.18097E-07 | 7.26989E-08 | 5.5147E-07 | 8.8675E-07 | 4.59E-07 | 1.07E-06 |
| | THC+CBD F | 11 | 9.9966E-07 | 5.00166E-07 | 1.50806E-07 | 6.6365E-07 | 1.3357E-06 | 4.73E-08 | 1.58E-06 |
| | CBD M | 11 | 5.4234E-07 | 3.19807E-07 | 9.64253E-08 | 3.2749E-07 | 7.5719E-07 | 8.77E-08 | 1.03E-06 |
| | THC+CBD M | 10 | 8.7520E-07 | 1.22116E-07 | 3.86166E-08 | 7.8784E-07 | 9.6256E-07 | 7.33E-07 | 1.10E-06 |
| THC | THC F | 10 | 4.2692E-07 | 1.70934E-07 | 5.40541E-08 | 3.0464E-07 | 5.4920E-07 | 1.96E-07 | 7.14E-07 |
| | THC+CBD F | 11 | 9.0785E-07 | 4.60231E-07 | 1.38765E-07 | 5.9866E-07 | 1.2170E-06 | 6.43E-08 | 1.47E-06 |
| | THC M | 9 | 8.3593E-07 | 3.57003E-07 | 1.19001E-07 | 5.6152E-07 | 1.1104E-06 | 6.24E-08 | 1.15E-06 |
| | THC+CBD M | 10 | 7.8550E-07 | 1.39798E-07 | 4.42079E-08 | 6.8549E-07 | 8.8551E-07 | 4.55E-07 | 9.45E-07 |
| 11-OH-THC | THC F | 10 | 2.1668E-08 | 9.44366E-09 | 2.98635E-09 | 1.4912E-08 | 2.8424E-08 | 5.76E-09 | 3.54E-08 |
| | THC+CBD F | 10 | 8.7050E-08 | 2.42305E-08 | 7.66236E-09 | 6.9716E-08 | 1.0438E-07 | 3.51E-08 | 1.23E-07 |
| | THC M | 8 | 3.6127E-08 | 6.44722E-09 | 2.27944E-09 | 3.0737E-08 | 4.1517E-08 | 2.81E-08 | 5.04E-08 |
| | THC+CBD M | 9 | 2.9786E-08 | 5.73924E-09 | 1.91308E-09 | 2.5374E-08 | 3.4197E-08 | 2.17E-08 | 3.55E-08 |
| 7-OH-CBD | CBD F | 8 | 1.1600E-07 | 3.76148E-08 | 1.32988E-08 | 8.4549E-08 | 1.4744E-07 | 6.44E-08 | 1.58E-07 |
| | THC+CBD F | 11 | 2.2212E-07 | 9.98755E-08 | 3.01136E-08 | 1.5502E-07 | 2.8922E-07 | 2.77E-08 | 3.40E-07 |
| | CBD M | 10 | 4.1240E-08 | 1.90796E-08 | 6.03350E-09 | 2.7591E-08 | 5.4889E-08 | 1.20E-08 | 7.37E-08 |
| | THC+CBD M | 9 | 7.1179E-08 | 3.66500E-08 | 1.22167E-08 | 4.3008E-08 | 9.9351E-08 | 3.32E-08 | 1.23E-07 |
| 11-COOH-THC | THC F | 10 | 4.8340E-08 | 1.28266E-08 | 4.05614E-09 | 3.9164E-08 | 5.7515E-08 | 2.90E-08 | 7.13E-08 |
| | THC+CBD F | 10 | 2.1814E-08 | 3.94229E-09 | 1.24666E-09 | 1.8993E-08 | 2.4634E-08 | 1.52E-08 | 2.64E-08 |
| | THC M | 9 | 5.0348E-08 | 1.99716E-08 | 6.65720E-09 | 3.4996E-08 | 6.5699E-08 | 2.45E-08 | 8.58E-08 |
| | THC+CBD M | 10 | 2.9336E-08 | 1.13294E-08 | 3.58268E-09 | 2.1231E-08 | 3.7440E-08 | 1.57E-08 | 4.92E-08 |
| 7-COOH-CBD | CBD F | 8 | 2.1520E-07 | 3.48292E-08 | 1.23140E-08 | 1.8608E-07 | 2.4432E-07 | 1.58E-07 | 2.66E-07 |
| | THC+CBD F | 11 | 4.8284E-07 | 2.61022E-07 | 7.87011E-08 | 3.0748E-07 | 6.5820E-07 | 3.92E-08 | 9.83E-07 |
| | THC+CBD M | 7 | 4.4645E-08 | 2.34302E-08 | 8.85577E-09 | 2.2976E-08 | 6.6314E-08 | 2.31E-08 | 8.84E-08 |
| | CBD M | BDL | | | | | | | |

Supplemental Table 2.2 Descriptive statistics for cannabinoids and metabolites (mol/g) measured in liver collected from male and female adult mice 2h after receiving an i.p. injection of 10mg/kg CBD, THC, or THC+CBD

| | | N | Mean | Std. Deviation | Std. Error | 95% Confidence Interval for Mean | | Minimum | Maximum |
|---|---|---|---|---|---|---|---|---|---|
| | | | | | | Lower Bound | Upper Bound | | |
| CBD | CBD F | 8 | 7.3280E-10 | 3.83171E-10 | 1.35471E-10 | 4.1246E-10 | 1.0531E-09 | 2.17E-10 | 1.26E-09 |
| | THC+CBD F | 10 | 9.2245E-10 | 6.20166E-10 | 1.96114E-10 | 4.7881E-10 | 1.3661E-09 | 5.29E-11 | 1.76E-09 |
| | CBD M | 10 | 2.2352E-10 | 1.24194E-10 | 3.92737E-11 | 1.3468E-10 | 3.1237E-10 | 2.57E-11 | 4.11E-10 |
| | THC+CBD M | 11 | 9.9230E-10 | 5.10577E-10 | 1.53945E-10 | 6.4929E-10 | 1.3353E-09 | 4.16E-10 | 1.90E-09 |
| THC | THC F | 10 | 3.0885E-10 | 1.56840E-10 | 4.95973E-11 | 1.9665E-10 | 4.2105E-10 | 8.52E-11 | 6.02E-10 |
| | THC+CBD F | 10 | 1.4282E-09 | 7.61564E-10 | 2.40828E-10 | 8.8337E-10 | 1.9730E-09 | 8.73E-11 | 2.20E-09 |
| | THC M | 9 | 1.1505E-09 | 5.09231E-10 | 1.69744E-10 | 7.5909E-10 | 1.5420E-09 | 1.57E-10 | 1.80E-09 |
| | THC+CBD M | 11 | 2.1483E-09 | 9.34022E-10 | 2.81618E-10 | 1.5208E-09 | 2.7758E-09 | 1.21E-09 | 3.83E-09 |
| 11-OH-THC | THC F | 10 | 2.3771E-09 | 1.20257E-09 | 3.80287E-10 | 1.5168E-09 | 3.2374E-09 | 4.41E-10 | 4.60E-09 |
| | THC+CBD F | 10 | 5.6849E-09 | 2.29251E-09 | 7.24957E-10 | 4.0449E-09 | 7.3249E-09 | 2.52E-09 | 9.12E-09 |
| | THC M | 9 | 3.1110E-09 | 1.62293E-09 | 5.40976E-10 | 1.8635E-09 | 4.3585E-09 | 3.39E-10 | 5.35E-09 |
| | THC+CBD M | 10 | 2.0959E-09 | 5.04175E-10 | 1.59434E-10 | 1.7352E-09 | 2.4566E-09 | 1.52E-09 | 2.96E-09 |
| 7-OH-CBD | CBD F | 8 | 1.3794E-09 | 8.36298E-10 | 2.95676E-10 | 6.8020E-10 | 2.0785E-09 | 5.21E-10 | 3.05E-09 |
| | THC+CBD F | 10 | 2.4396E-09 | 1.55127E-09 | 4.90556E-10 | 1.3299E-09 | 3.5493E-09 | 3.19E-10 | 5.36E-09 |
| | CBD M | 10 | 4.9960E-10 | 2.23450E-10 | 7.06611E-11 | 3.3975E-10 | 6.5944E-10 | 1.45E-10 | 8.83E-10 |
| | THC+CBD M | 10 | 1.0894E-09 | 3.87531E-10 | 1.22548E-10 | 8.1220E-10 | 1.3666E-09 | 5.95E-10 | 1.67E-09 |
| 11-COOH-THC | THC F | 9 | 6.7764E-09 | 2.48008E-09 | 8.26694E-10 | 4.8700E-09 | 8.6827E-09 | 2.78E-09 | 1.16E-08 |
| | THC+CBD F | 10 | 3.6888E-09 | 9.31575E-10 | 2.94590E-10 | 3.0224E-09 | 4.3552E-09 | 2.25E-09 | 5.51E-09 |
| | THC M | 9 | 8.8798E-09 | 3.27596E-09 | 1.09199E-09 | 6.3617E-09 | 1.1398E-08 | 3.05E-09 | 1.47E-08 |
| | THC+CBD M | 10 | 4.6031E-09 | 1.43112E-09 | 4.52561E-10 | 3.5793E-09 | 5.6268E-09 | 3.36E-09 | 7.11E-09 |
| 7-COOH-CBD | CBD F | 8 | 1.0702E-09 | 4.44790E-10 | 1.57257E-10 | 6.9833E-10 | 1.4420E-09 | 6.27E-10 | 1.67E-09 |
| | THC+CBD F | 10 | 1.2655E-09 | 4.39904E-10 | 1.39110E-10 | 9.5077E-10 | 1.5801E-09 | 5.16E-10 | 1.93E-09 |
| | THC+CBD M | 6 | 5.9804E-10 | 2.20578E-10 | 9.00506E-11 | 3.6656E-10 | 8.2952E-10 | 3.71E-10 | 9.60E-10 |
| | CBD M | BAL | | | | | | | |

Supplemental Table 2.3 One-way ANOVAs for levels of cannabinoids and metabolites (mol/L) in plasma in each male and female treatment group separately

| | | Sum of Squares | df | Mean Square | F | Sig. |
|---|---|---|---|---|---|---|
| 7-OH-CBD | Between Groups | 1.98E-13 | 3 | 6.59E-14 | 18.129 | 0.0000003 |
| | Within Groups | 1.24E-13 | 34 | 3.64E-15 | | |
| | Total | 3.22E-13 | 37 | | | |
| 7-COOH-CBD | Between Groups | 8.74E-13 | 2 | 4.37E-13 | 14.497 | 0.0000844 |
| | Within Groups | 6.93E-13 | 23 | 3.01E-14 | | |
| | Total | 1.57E-12 | 25 | | | |
| 11-OH-THC | Between Groups | 2.58E-14 | 3 | 8.61E-15 | 42.778 | 0.0000000 |
| | Within Groups | 6.64E-15 | 33 | 2.01E-16 | | |
| | Total | 3.25E-14 | 36 | | | |
| 11-COOH-THC | Between Groups | 5.78E-15 | 3 | 1.93E-15 | 11.306 | 0.0000246 |
| | Within Groups | 5.97E-15 | 35 | 1.70E-16 | | |
| | Total | 1.17E-14 | 38 | | | |

Supplemental Table 2.4 One-way ANOVAs for levels of cannabinoids and metabolites (mol/g) in liver in each male and female treatment group separately

| | | Sum of Squares | df | Mean Square | F | Sig. |
|---|---|---|---|---|---|---|
| 7-OH-CBD | Between Groups | 1.98E-17 | 3 | 6.60E-18 | 7.910 | 0.00039 |
| | Within Groups | 2.84E-17 | 34 | 8.34E-19 | | |
| | Total | 4.81E-17 | 37 | | | |
| 7-COOH-CBD | Between Groups | 1.69E-18 | 2 | 8.43E-19 | 5.255 | 0.01411 |
| | Within Groups | 3.37E-18 | 21 | 1.60E-19 | | |
| | Total | 5.06E-18 | 23 | | | |
| 11-OH-THC | Between Groups | 8.02E-17 | 3 | 2.67E-17 | 11.181 | 0.00003 |
| | Within Groups | 8.37E-17 | 35 | 2.39E-18 | | |
| | Total | 1.64E-16 | 38 | | | |
| 11-COOH-THC | Between Groups | 1.53E-16 | 3 | 5.08E-17 | 10.717 | 0.00004 |
| | Within Groups | 1.61E-16 | 34 | 4.74E-18 | | |
| | Total | 3.14E-16 | 37 | | | |

#### Supplemental 2.5 Data analysis summary for Fisher's LSD post-hoc test for cannabinoids and cannabinoid metabolite ANOVAs in plasma

| Dependent Variable | | | Std. Error | Sig. | 95% Confidence Interval Lower Bound | 95% Confidence Interval Upper Bound |
|---|---|---|---|---|---|---|
| | CBD female | CBD in Tp C female | 1.45E-07 | 0.0572 | -5.70E-07 | 8.78E-09 |
| | | CBD male | 1.45E-07 | 0.2273 | -1.13E-07 | 4.66E-07 |
| | | CBD in TpC male | 1.48E-07 | 0.2964 | -4.52E-07 | 1.40E-07 |
| | | THC female | 1.48E-07 | 0.0528 | -3.58E-09 | 5.88E-07 |
| | | THC in TpC female | 1.45E-07 | 0.1977 | -4.78E-07 | 1.01E-07 |
| | | THC male | 1.52E-07 | 0.4454 | -4.20E-07 | 1.87E-07 |
| | CBD in Tp C female | THC in TpC male | 1.48E-07 | 0.6559 | -3.62E-07 | 2.29E-07 |
| | | CBD female | 1.45E-07 | 0.0572 | -8.78E-09 | 5.70E-07 |
| | | CBD male | 1.38E-07 | 0.0014 | 1.83E-07 | 7.32E-07 |
| | | CBD in TpC male | 1.41E-07 | 0.3807 | -1.57E-07 | 4.06E-07 |
| | | THC female | 1.41E-07 | 0.0001 | 2.91E-07 | 8.54E-07 |
| | | THC in TpC female | 1.38E-07 | 0.5071 | -1.83E-07 | 3.66E-07 |
| | | THC male | 1.45E-07 | 0.2631 | -1.26E-07 | 4.53E-07 |
| | | THC in TpC male | 1.41E-07 | 0.1335 | -6.71E-08 | 4.95E-07 |
| CBD | CBD male | CBD female | 1.45E-07 | 0.2273 | -4.66E-07 | 1.13E-07 |
| | | CBD in Tp C female | 1.38E-07 | 0.0014 | -7.32E-07 | -1.83E-07 |
| | | CBD in TpC male | 1.41E-07 | 0.0210 | -6.14E-07 | -5.16E-08 |
| | | THC female | 1.41E-07 | 0.4161 | -1.66E-07 | 3.97E-07 |
| | | THC in TpC female | 1.38E-07 | 0.0098 | -6.40E-07 | -9.10E-08 |
| | | THC male | 1.45E-07 | 0.0468 | -5.83E-07 | -4.26E-09 |
| | | THC in TpC male | 1.41E-07 | 0.0891 | -5.24E-07 | 3.81E-08 |
| | CBD in TpC male | CBD female | 1.48E-07 | 0.2964 | -1.40E-07 | 4.52E-07 |
| | | CBD in Tp C female | 1.41E-07 | 0.3807 | -4.06E-07 | 1.57E-07 |
| | | CBD male | 1.41E-07 | 0.0210 | 5.16E-08 | 6.14E-07 |
| | | THC female | 1.44E-07 | 0.0027 | 1.60E-07 | 7.36E-07 |
| | | THC in TpC female | 1.41E-07 | 0.8177 | -3.14E-07 | 2.49E-07 |
| | | THC male | 1.48E-07 | 0.7921 | -2.57E-07 | 3.35E-07 |
| | | THC in TpC male | 1.44E-07 | 0.5365 | -1.98E-07 | 3.78E-07 |
| | THC female | CBD female | 1.48E-07 | 0.0528 | -5.88E-07 | 3.58E-09 |
| | | CBD in Tp C female | 1.41E-07 | 0.0001 | -8.54E-07 | -2.91E-07 |
| | | CBD male | 1.41E-07 | 0.4161 | -3.97E-07 | 1.66E-07 |
| | | CBD in TpC male | 1.44E-07 | 0.0027 | -7.36E-07 | -1.60E-07 |
| THC | | THC in TpC female | 1.41E-07 | 0.0011 | -7.62E-07 | -2.00E-07 |
| | | THC male | 1.48E-07 | 0.0074 | -7.05E-07 | -1.13E-07 |
| | | THC in TpC male | 1.44E-07 | 0.0153 | -6.46E-07 | -7.07E-08 |
| | THC in TpC female | CBD female | 1.45E-07 | 0.1977 | -1.01E-07 | 4.78E-07 |
| | | CBD in Tp C female | 1.38E-07 | 0.5071 | -3.66E-07 | 1.83E-07 |

| | | | | | | |
|---|---|---|---|---|---|---|
| | | CBD male | 1.38E-07 | 0.0098 | 9.10E-08 | 6.40E-07 |
| | | CBD in TpC male | 1.41E-07 | 0.8177 | -2.49E-07 | 3.14E-07 |
| | | THC female | 1.41E-07 | 0.0011 | 2.00E-07 | 7.62E-07 |
| | | THC male | 1.45E-07 | 0.6218 | -2.17E-07 | 3.61E-07 |
| | | THC in TpC male | 1.41E-07 | 0.3888 | -1.59E-07 | 4.04E-07 |
| | THC male | CBD female | 1.52E-07 | 0.4454 | -1.87E-07 | 4.20E-07 |
| | | CBD in Tp C female | 1.45E-07 | 0.2631 | -4.53E-07 | 1.26E-07 |
| | | CBD male | 1.45E-07 | 0.0468 | 4.26E-09 | 5.83E-07 |
| | | CBD in TpC male | 1.48E-07 | 0.7921 | -3.35E-07 | 2.57E-07 |
| | | THC female | 1.48E-07 | 0.0074 | 1.13E-07 | 7.05E-07 |
| | | THC in TpC female | 1.45E-07 | 0.6218 | -3.61E-07 | 2.17E-07 |
| | | THC in TpC male | 1.48E-07 | 0.7350 | -2.45E-07 | 3.46E-07 |
| | THC in TpC male | CBD female | 1.48E-07 | 0.6559 | -2.29E-07 | 3.62E-07 |
| | | CBD in Tp C female | 1.41E-07 | 0.1335 | -4.95E-07 | 6.71E-08 |
| | | CBD male | 1.41E-07 | 0.0891 | -3.81E-08 | 5.24E-07 |
| | | CBD in TpC male | 1.44E-07 | 0.5365 | -3.78E-07 | 1.98E-07 |
| | | THC female | 1.44E-07 | 0.0153 | 7.07E-08 | 6.46E-07 |
| | | THC in TpC female | 1.41E-07 | 0.3888 | -4.04E-07 | 1.59E-07 |
| | | THC male | 1.48E-07 | 0.7350 | -3.46E-07 | 2.45E-07 |
| 11-OH-THC | THC female | THC in T+C Female | 6.34E-09 | 0.0000 | -7.83E-08 | -5.25E-08 |
| | | THC Male | 6.73E-09 | 0.0391 | -2.81E-08 | -7.69E-10 |
| | | THC in T+C Male | 6.52E-09 | 0.2217 | -2.14E-08 | 5.14E-09 |
| | THC in T+C Female | THC Female | 6.34E-09 | 0.0000 | 5.25E-08 | 7.83E-08 |
| | | THC Male | 6.73E-09 | 0.0000 | 3.72E-08 | 6.46E-08 |
| | | THC in T+C Male | 6.52E-09 | 0.0000 | 4.40E-08 | 7.05E-08 |
| | THC Male | THC Female | 6.73E-09 | 0.0391 | 7.69E-10 | 2.81E-08 |
| | | THC in T+C Female | 6.73E-09 | 0.0000 | -6.46E-08 | -3.72E-08 |
| | | THC in T+C Male | 6.89E-09 | 0.3643 | -7.68E-09 | 2.04E-08 |
| | THC in T+C Male | THC Female | 6.52E-09 | 0.2217 | -5.14E-09 | 2.14E-08 |
| | | THC in T+C Female | 6.52E-09 | 0.0000 | -7.05E-08 | -4.40E-08 |
| | | THC Male | 6.89E-09 | 0.3643 | -2.04E-08 | 7.68E-09 |
| 7-OH-CBD | CBD Female | CBD in T+C Female | 2.80E-08 | 0.0006 | -1.63E-07 | -4.92E-08 |
| | | CBD Male | 2.86E-08 | 0.0133 | 1.66E-08 | 1.33E-07 |
| | | CBD in T+C Male | 2.93E-08 | 0.1355 | -1.47E-08 | 1.04E-07 |
| | CBD in T+C Female | CBD Female | 2.80E-08 | 0.0006 | 4.92E-08 | 1.63E-07 |
| | | CBD Male | 2.64E-08 | 0.0000 | 1.27E-07 | 2.34E-07 |
| | | CBD in T+C Male | 2.71E-08 | 0.0000 | 9.58E-08 | 2.06E-07 |
| | CBD Male | CBD Female | 2.86E-08 | 0.0133 | -1.33E-07 | -1.66E-08 |
| | | CBD in T+C Female | 2.64E-08 | 0.0000 | -2.34E-07 | -1.27E-07 |
| | | CBD in T+C Male | 2.77E-08 | 0.2876 | -8.63E-08 | 2.64E-08 |
| | CBD in T+C Male | CBD Female | 2.93E-08 | 0.1355 | -1.04E-07 | 1.47E-08 |

| | | | | | | |
|---|---|---|---|---|---|---|
| | | CBD in T+C Female | 2.71E-08 | 0.0000 | -2.06E-07 | -9.58E-08 |
| | | CBD Male | 2.77E-08 | 0.2876 | -2.64E-08 | 8.63E-08 |
| 11-COOH-THC | THC Female | THC in T+C Female | 5.84E-09 | 0.0001 | 1.47E-08 | 3.84E-08 |
| | | THC Male | 6.00E-09 | 0.7398 | -1.42E-08 | 1.02E-08 |
| | | THC in T+C Male | 5.84E-09 | 0.0025 | 7.15E-09 | 3.09E-08 |
| | THC in T+C Female | THC Female | 5.84E-09 | 0.0001 | -3.84E-08 | -1.47E-08 |
| | | THC Male | 6.00E-09 | 0.0000 | -4.07E-08 | -1.64E-08 |
| | | THC in T+C Male | 5.84E-09 | 0.2061 | -1.94E-08 | 4.33E-09 |
| | THC Male | THC Female | 6.00E-09 | 0.7398 | -1.02E-08 | 1.42E-08 |
| | | THC in T+C Female | 6.00E-09 | 0.0000 | 1.64E-08 | 4.07E-08 |
| | | THC in T+C Male | 6.00E-09 | 0.0013 | 8.83E-09 | 3.32E-08 |
| | THC in T+C Male | THC Female | 5.84E-09 | 0.0025 | -3.09E-08 | -7.15E-09 |
| | | THC in T+C Female | 5.84E-09 | 0.2061 | -4.33E-09 | 1.94E-08 |
| | | THC Male | 6.00E-09 | 0.0013 | -3.32E-08 | -8.83E-09 |
| 7-COOH-CBD | CBD Female | CBD in T+C Female | 8.07E-08 | 0.0030 | -4.35E-07 | -1.01E-07 |
| | | CBD in T+C Male | 8.98E-08 | 0.0703 | -1.53E-08 | 3.56E-07 |
| | CBD in T+C Female | CBD Female | 8.07E-08 | 0.0030 | 1.01E-07 | 4.35E-07 |
| | | CBD in T+C Male | 8.39E-08 | 0.0000 | 2.65E-07 | 6.12E-07 |
| | CBD in T+C Male | CBD Female | 8.98E-08 | 0.0703 | -3.56E-07 | 1.53E-08 |
| | | CBD in T+C Female | 8.39E-08 | 0.0000 | -6.12E-07 | -2.65E-07 |

Supplemental 2.6 Data analysis summary for Fisher's LSD post-hoc test for cannabinoids and cannabinoid metabolite ANOVAs in liver

| | (I) TxSex | | Std. Error | Sig. | 95% Confidence Interval Lower Bound | Upper Bound |
|---|---|---|---|---|---|---|
| | CBD female | CBD in TpC female | 2.72748E-10 | 0.489 | -7.3350E-10 | 3.5419E-10 |
| | | CBD male | 2.72748E-10 | 0.066 | -3.4569E-11 | 1.0531E-09 |
| | | CBD in TpC male | 2.67181E-10 | 0.335 | -7.9224E-10 | 2.7324E-10 |
| | | THC female | 2.72748E-10 | 0.125 | -1.1990E-10 | 9.6779E-10 |
| | | THC in TpC female | 2.72748E-10 | 0.013 | -1.2392E-09 | -1.5152E-10 |
| | | THC male | 2.79401E-10 | 0.139 | -9.7484E-10 | 1.3938E-10 |
| | | THC in TpC male | 2.67181E-10 | 0.000 | -1.9482E-09 | -8.8274E-10 |
| | CBD in Tp C female | CBD female | 2.72748E-10 | 0.489 | -3.5419E-10 | 7.3350E-10 |
| | | CBD male | 2.57149E-10 | 0.008 | 1.8619E-10 | 1.2117E-09 |
| | | CBD in TpC male | 2.51237E-10 | 0.782 | -5.7080E-10 | 4.3111E-10 |
| | | THC female | 2.57149E-10 | 0.020 | 1.0086E-10 | 1.1263E-09 |
| | | THC in TpC female | 2.57149E-10 | 0.053 | -1.0184E-09 | 7.0331E-12 |
| | | THC male | 2.64196E-10 | 0.391 | -7.5486E-10 | 2.9872E-10 |
| | | THC in TpC male | 2.51237E-10 | 0.000 | -1.7268E-09 | -7.2488E-10 |
| CBD | CBD male | CBD female | 2.72748E-10 | 0.066 | -1.0531E-09 | 3.4569E-11 |
| | | CBD in TpC female | 2.57149E-10 | 0.008 | -1.2117E-09 | -1.8619E-10 |
| | | CBD in TpC male | 2.51237E-10 | 0.003 | -1.2697E-09 | -2.6782E-10 |
| | | THC female | 2.57149E-10 | 0.741 | -5.9807E-10 | 4.2741E-10 |
| | | THC in TpC female | 2.57149E-10 | 0.000 | -1.7174E-09 | -6.9190E-10 |
| | | THC male | 2.64196E-10 | 0.001 | -1.4538E-09 | -4.0021E-10 |
| | | THC in TpC male | 2.51237E-10 | 0.000 | -2.4257E-09 | -1.4238E-09 |
| | CBD in TpC male | CBD female | 2.67181E-10 | 0.335 | -2.7324E-10 | 7.9224E-10 |
| | | CBD in TpC female | 2.51237E-10 | 0.782 | -4.3111E-10 | 5.7080E-10 |
| | | CBD male | 2.51237E-10 | 0.003 | 2.6782E-10 | 1.2697E-09 |
| | | THC female | 2.51237E-10 | 0.008 | 1.8249E-10 | 1.1844E-09 |
| | | THC in TpC female | 2.51237E-10 | 0.087 | -9.3682E-10 | 6.5087E-11 |
| | | THC male | 2.58445E-10 | 0.542 | -6.7355E-10 | 3.5710E-10 |
| | | THC in TpC male | 2.45182E-10 | 0.000 | -1.6449E-09 | -6.6711E-10 |
| | THC female | CBD female | 2.72748E-10 | 0.125 | -9.6779E-10 | 1.1990E-10 |
| | | CBD in TpC female | 2.57149E-10 | 0.020 | -1.1263E-09 | -1.0086E-10 |
| | | CBD male | 2.57149E-10 | 0.741 | -4.2741E-10 | 5.9807E-10 |
| | | CBD in TpC male | 2.51237E-10 | 0.008 | -1.1844E-09 | -1.8249E-10 |
| | | THC in TpC female | 2.57149E-10 | 0.000 | -1.6321E-09 | -6.0657E-10 |
| | | THC male | 2.64196E-10 | 0.002 | -1.3685E-09 | -3.1488E-10 |
| | | THC in TpC male | 2.51237E-10 | 0.000 | -2.3404E-09 | -1.3385E-09 |
| | THC in TpC female | CBD female | 2.72748E-10 | 0.013 | 1.5152E-10 | 1.2392E-09 |
| | | CBD in Tp C female | 2.57149E-10 | 0.053 | -7.0331E-12 | 1.0184E-09 |
| | | CBD male | 2.57149E-10 | 0.000 | 6.9190E-10 | 1.7174E-09 |
| THC | | CBD in TpC male | 2.51237E-10 | 0.087 | -6.5087E-11 | 9.3682E-10 |
| | | THC female | 2.57149E-10 | 0.000 | 6.0657E-10 | 1.6321E-09 |
| | | THC male | 2.64196E-10 | 0.297 | -2.4915E-10 | 8.0443E-10 |
| | | THC in TpC male | 2.51237E-10 | 0.005 | -1.2211E-09 | -2.1917E-10 |
| | THC male | CBD female | 2.79401E-10 | 0.139 | -1.3938E-10 | 9.7484E-10 |
| | | CBD in Tp C female | 2.64196E-10 | 0.391 | -2.9872E-10 | 7.5486E-10 |
| | | CBD male | 2.64196E-10 | 0.001 | 4.0021E-10 | 1.4538E-09 |
| | | CBD in TpC male | 2.58445E-10 | 0.542 | -3.5710E-10 | 6.7355E-10 |
| | | THC female | 2.64196E-10 | 0.002 | 3.1488E-10 | 1.3685E-09 |
| | | THC in TpC female | 2.64196E-10 | 0.297 | -8.0443E-10 | 2.4915E-10 |
| | | THC in TpC male | 2.58445E-10 | 0.000 | -1.5131E-09 | -4.8243E-10 |
| | THC in TpC male | CBD female | 2.67181E-10 | 0.000 | 8.8274E-10 | 1.9482E-09 |

| | | | | | | |
|---|---|---|---|---|---|---|
| | | CBD in Tp C female | 2.51237E-10 | 0.000 | 7.2488E-10 | 1.7268E-09 |
| | | CBD male | 2.51237E-10 | 0.000 | 1.4238E-09 | 2.4257E-09 |
| | | CBD in TpC male | 2.45182E-10 | 0.000 | 6.6711E-10 | 1.6449E-09 |
| | | THC female | 2.51237E-10 | 0.000 | 1.3385E-09 | 2.3404E-09 |
| | | THC in TpC female | 2.51237E-10 | 0.005 | 2.1917E-10 | 1.2211E-09 |
| | | THC male | 2.58445E-10 | 0.000 | 4.8243E-10 | 1.5131E-09 |
| 11-OH-THC | THC Female | THC in T+C Female | 6.91479E-10 | 0.000 | -4.7116E-09 | -1.9040E-09 |
| | | THC Male | 7.10427E-10 | 0.309 | -2.1762E-09 | 7.0834E-10 |
| | | THC in T+C Male | 6.91479E-10 | 0.687 | -1.1226E-09 | 1.6850E-09 |
| | THC in T+C Female | THC Female | 6.91479E-10 | 0.000 | 1.9040E-09 | 4.7116E-09 |
| | | THC Male | 7.10427E-10 | 0.001 | 1.1316E-09 | 4.0161E-09 |
| | | THC in T+C Male | 6.91479E-10 | 0.000 | 2.1852E-09 | 4.9928E-09 |
| | THC Male | THC Female | 7.10427E-10 | 0.309 | -7.0834E-10 | 2.1762E-09 |
| | | THC in T+C Female | 7.10427E-10 | 0.001 | -4.0161E-09 | -1.1316E-09 |
| | | THC in T+C Male | 7.10427E-10 | 0.162 | -4.2714E-10 | 2.4574E-09 |
| | THC in T+C Male | THC Female | 6.91479E-10 | 0.687 | -1.6850E-09 | 1.1226E-09 |
| | | THC in T+C Female | 6.91479E-10 | 0.000 | -4.9928E-09 | -2.1852E-09 |
| | | THC Male | 7.10427E-10 | 0.162 | -2.4574E-09 | 4.2714E-10 |
| 7-OH-CBD | CBD Female | CBD in T+C Female | 4.33177E-10 | 0.020 | -1.9406E-09 | -1.7991E-10 |
| | | CBD Male | 4.33177E-10 | 0.050 | -5.5032E-13 | 1.7601E-09 |
| | | CBD in T+C Male | 4.33177E-10 | 0.508 | -5.9038E-10 | 1.1703E-09 |
| | CBD in T+C Female | CBD Female | 4.33177E-10 | 0.020 | 1.7991E-10 | 1.9406E-09 |
| | | CBD Male | 4.08403E-10 | 0.000 | 1.1100E-09 | 2.7700E-09 |
| | | CBD in T+C Male | 4.08403E-10 | 0.002 | 5.2020E-10 | 2.1802E-09 |
| | CBD Male | CBD Female | 4.33177E-10 | 0.050 | -1.7601E-09 | 5.5032E-13 |
| | | CBD in T+C Female | 4.08403E-10 | 0.000 | -2.7700E-09 | -1.1100E-09 |
| | | CBD in T+C Male | 4.08403E-10 | 0.158 | -1.4198E-09 | 2.4015E-10 |
| | CBD in T+C Male | CBD Female | 4.33177E-10 | 0.508 | -1.1703E-09 | 5.9038E-10 |
| | | CBD in T+C Female | 4.08403E-10 | 0.002 | -2.1802E-09 | -5.2020E-10 |
| | | CBD Male | 4.08403E-10 | 0.158 | -2.4015E-10 | 1.4198E-09 |
| 11-COOH-THC | THC Female | THC in T+C Female | 1.00078E-09 | 0.004 | 1.0537E-09 | 5.1214E-09 |
| | | THC Male | 1.02678E-09 | 0.048 | -4.1901E-09 | -1.6781E-11 |
| | | THC in T+C Male | 1.00078E-09 | 0.037 | 1.3944E-10 | 4.2071E-09 |
| | THC in T+C Female | THC Female | 1.00078E-09 | 0.004 | -5.1214E-09 | -1.0537E-09 |
| | | THC Male | 1.00078E-09 | 0.000 | -7.2248E-09 | -3.1571E-09 |
| | | THC in T+C Male | 9.74091E-10 | 0.355 | -2.8938E-09 | 1.0653E-09 |
| | THC Male | THC Female | 1.02678E-09 | 0.048 | 1.6781E-11 | 4.1901E-09 |
| | | THC in T+C Female | 1.00078E-09 | 0.000 | 3.1571E-09 | 7.2248E-09 |
| | | THC in T+C Male | 1.00078E-09 | 0.000 | 2.2429E-09 | 6.3106E-09 |
| | THC in T+C Male | THC Female | 1.00078E-09 | 0.037 | -4.2071E-09 | -1.3944E-10 |
| | | THC in T+C Female | 9.74091E-10 | 0.355 | -1.0653E-09 | 2.8938E-09 |
| | | THC Male | 1.00078E-09 | 0.000 | -6.3106E-09 | -2.2429E-09 |
| 7-COOH-CBD | CBD Female | CBD in T+C Female | 1.90013E-10 | 0.316 | -5.9042E-10 | 1.9988E-10 |
| | | CBD in TpC Male | 2.16339E-10 | 0.041 | 2.2246E-11 | 9.2205E-10 |
| | CBD in T+C Female | CBD Female | 1.90013E-10 | 0.316 | -1.9988E-10 | 5.9042E-10 |
| | | CBD in TpC Male | 2.06860E-10 | 0.004 | 2.3723E-10 | 1.0976E-09 |
| | CBD in TpC Male | CBD Female | 2.16339E-10 | 0.041 | -9.2205E-10 | -2.2246E-11 |
| | | CBD in T+C Female | 2.06860E-10 | 0.004 | -1.0976E-09 | -2.3723E-10 |

Supplemental Table 3.1 Descriptive statistics for *N*-acyl ethanolamines (mol/L) in plasma collected from male and female adult mice 2h after receiving an i.p. injection of 10mg/kg CBD, THC, or THC+CBD

| | | N | Mean | Std. Deviation | Std. Error | 95% Confidence Interval for Mean | | Minimum | Maximum |
|---|---|---|---|---|---|---|---|---|---|
| | | | | | | Lower Bound | Upper Bound | | |
| *N*-palmitoyl ethanolamine | Veh F | 7 | 5.4990E-08 | 6.69771E-09 | 2.53149E-09 | 4.8796E-08 | 6.1184E-08 | 4.56E-08 | 6.44E-08 |
| | CBD F | 8 | 6.0441E-08 | 6.23270E-09 | 2.20359E-09 | 5.5230E-08 | 6.5651E-08 | 4.66E-08 | 6.69E-08 |
| | THC F | 10 | 6.1098E-08 | 1.13741E-08 | 3.59682E-09 | 5.2962E-08 | 6.9235E-08 | 4.01E-08 | 7.54E-08 |
| | THC+CBD F | 11 | 6.5448E-08 | 1.57648E-08 | 4.75328E-09 | 5.4857E-08 | 7.6039E-08 | 4.98E-08 | 9.59E-08 |
| | Veh M | 8 | 6.0680E-08 | 5.37243E-09 | 1.89944E-09 | 5.6189E-08 | 6.5172E-08 | 5.53E-08 | 7.06E-08 |
| | CBD M | 11 | 6.3404E-08 | 1.10585E-08 | 3.33426E-09 | 5.5975E-08 | 7.0833E-08 | 4.96E-08 | 7.91E-08 |
| | THC M | 9 | 5.4108E-08 | 7.06537E-09 | 2.35512E-09 | 4.8677E-08 | 5.9539E-08 | 4.30E-08 | 6.59E-08 |
| | THC+CBD M | 10 | 6.6794E-08 | 1.17639E-08 | 3.72006E-09 | 5.8379E-08 | 7.5210E-08 | 4.85E-08 | 8.40E-08 |
| *N*-stearoyl ethanolamine | Veh F | 7 | 4.1945E-08 | 4.44183E-09 | 1.67886E-09 | 3.7837E-08 | 4.6053E-08 | 3.28E-08 | 4.50E-08 |
| | CBD F | 9 | 4.6161E-08 | 5.52247E-09 | 1.84082E-09 | 4.1917E-08 | 5.0406E-08 | 4.05E-08 | 5.35E-08 |
| | THC F | 10 | 4.6960E-08 | 8.99374E-09 | 2.84407E-09 | 4.0527E-08 | 5.3394E-08 | 3.42E-08 | 6.09E-08 |
| | THC+CBD F | 11 | 5.0331E-08 | 9.59002E-09 | 2.89150E-09 | 4.3888E-08 | 5.6774E-08 | 3.75E-08 | 6.85E-08 |
| | Veh M | 8 | 5.0575E-08 | 1.41775E-08 | 5.01249E-09 | 3.8722E-08 | 6.2427E-08 | 3.32E-08 | 7.10E-08 |
| | CBD M | 10 | 4.6526E-08 | 7.00865E-09 | 2.21633E-09 | 4.1512E-08 | 5.1540E-08 | 4.05E-08 | 5.99E-08 |
| | THC M | 9 | 4.4217E-08 | 6.23473E-09 | 2.07824E-09 | 3.9425E-08 | 4.9010E-08 | 3.59E-08 | 5.35E-08 |
| | THC+CBD M | 9 | 4.8631E-08 | 4.93799E-09 | 1.64600E-09 | 4.4835E-08 | 5.2426E-08 | 4.01E-08 | 5.59E-08 |
| *N*-oleoyl ethanolamine | Veh F | 7 | 6.0573E-08 | 9.79096E-09 | 3.70063E-09 | 5.1518E-08 | 6.9628E-08 | 4.88E-08 | 7.39E-08 |
| | CBD F | 9 | 6.9150E-08 | 1.10469E-08 | 3.68230E-09 | 6.0659E-08 | 7.7641E-08 | 5.19E-08 | 8.54E-08 |
| | THC F | 10 | 7.1852E-08 | 1.29739E-08 | 4.10270E-09 | 6.2571E-08 | 8.1133E-08 | 5.63E-08 | 9.72E-08 |
| | THC+CBD F | 10 | 7.7944E-08 | 1.54155E-08 | 4.87480E-09 | 6.6916E-08 | 8.8971E-08 | 5.56E-08 | 1.05E-07 |
| | Veh M | 8 | 6.7720E-08 | 1.39301E-08 | 4.92502E-09 | 5.6074E-08 | 7.9366E-08 | 4.99E-08 | 8.75E-08 |
| | CBD M | 11 | 6.7259E-08 | 1.97745E-08 | 5.96225E-09 | 5.3974E-08 | 8.0544E-08 | 4.47E-08 | 1.01E-07 |
| | THC M | 8 | 6.0536E-08 | 8.45671E-09 | 2.98990E-09 | 5.3466E-08 | 6.7606E-08 | 5.21E-08 | 7.43E-08 |
| | THC+CBD M | 10 | 7.7484E-08 | 1.29578E-08 | 4.09761E-09 | 6.8215E-08 | 8.6753E-08 | 5.35E-08 | 9.20E-08 |
| *N*-linoleoyl ethanolamine | Veh F | 7 | 3.2895E-08 | 5.75511E-09 | 2.17523E-09 | 2.7573E-08 | 3.8218E-08 | 2.57E-08 | 4.28E-08 |
| | CBD F | 9 | 3.1148E-08 | 4.87427E-09 | 1.62476E-09 | 2.7401E-08 | 3.4895E-08 | 2.58E-08 | 4.04E-08 |
| | THC F | 9 | 3.0986E-08 | 6.83197E-09 | 2.27732E-09 | 2.5735E-08 | 3.6238E-08 | 2.29E-08 | 4.37E-08 |

| | | | | | | | | |
|---|---|---|---|---|---|---|---|---|
| | THC+CBD F | 10 | 4.2388E-08 | 8.42062E-09 | 2.66283E-09 | 3.6365E-08 | 4.8412E-08 | 2.92E-08 | 5.57E-08 |
| | Veh M | 8 | 3.2232E-08 | 6.09248E-09 | 2.15402E-09 | 2.7138E-08 | 3.7325E-08 | 2.38E-08 | 4.02E-08 |
| | CBD M | 11 | 3.2294E-08 | 6.69580E-09 | 2.01886E-09 | 2.7796E-08 | 3.6793E-08 | 2.04E-08 | 4.55E-08 |
| | THC M | 8 | 3.4211E-08 | 5.36288E-09 | 1.89606E-09 | 2.9728E-08 | 3.8695E-08 | 2.80E-08 | 4.21E-08 |
| | THC+CBD M | 10 | 3.3806E-08 | 7.69707E-09 | 2.43403E-09 | 2.8300E-08 | 3.9312E-08 | 2.18E-08 | 4.73E-08 |
| *N*-arachidonoyl ethanolamine | Veh F | 7 | 6.9641E-09 | 1.44288E-09 | 5.45357E-10 | 5.6296E-09 | 8.2985E-09 | 4.42E-09 | 8.91E-09 |
| | CBD F | 8 | 6.5533E-09 | 1.04486E-09 | 3.69413E-10 | 5.6798E-09 | 7.4268E-09 | 5.31E-09 | 8.53E-09 |
| | THC F | 9 | 7.2002E-09 | 1.58246E-09 | 5.27488E-10 | 5.9838E-09 | 8.4166E-09 | 5.71E-09 | 1.04E-08 |
| | THC+CBD F | 11 | 7.2124E-09 | 2.06235E-09 | 6.21821E-10 | 5.8269E-09 | 8.5979E-09 | 4.24E-09 | 1.10E-08 |
| | Veh M | 8 | 7.8321E-09 | 1.19808E-09 | 4.23584E-10 | 6.8304E-09 | 8.8337E-09 | 5.95E-09 | 9.11E-09 |
| | CBD M | 11 | 7.3830E-09 | 1.79072E-09 | 5.39922E-10 | 6.1800E-09 | 8.5860E-09 | 3.87E-09 | 1.05E-08 |
| | THC M | 9 | 6.5205E-09 | 1.45229E-09 | 4.84098E-10 | 5.4042E-09 | 7.6368E-09 | 5.20E-09 | 9.27E-09 |
| | THC+CBD M | 10 | 7.0255E-09 | 2.04243E-09 | 6.45874E-10 | 5.5645E-09 | 8.4866E-09 | 5.10E-09 | 1.07E-08 |
| *N*-docosahexaenoyl ethanolamine | Veh F | 7 | 3.8643E-08 | 9.06794E-09 | 3.42736E-09 | 3.0256E-08 | 4.7029E-08 | 2.54E-08 | 5.26E-08 |
| | CBD F | 8 | 3.8522E-08 | 4.77030E-09 | 1.68656E-09 | 3.4534E-08 | 4.2510E-08 | 3.19E-08 | 4.48E-08 |
| | THC F | 9 | 4.3224E-08 | 1.57918E-08 | 5.26394E-09 | 3.1085E-08 | 5.5363E-08 | 2.15E-08 | 7.41E-08 |
| | THC+CBD F | 11 | 3.9580E-08 | 3.55607E-09 | 1.07220E-09 | 3.7191E-08 | 4.1969E-08 | 3.40E-08 | 4.47E-08 |
| | Veh M | 8 | 4.1491E-08 | 9.93533E-09 | 3.51267E-09 | 3.3185E-08 | 4.9797E-08 | 3.19E-08 | 5.76E-08 |
| | CBD M | 10 | 4.6398E-08 | 1.13288E-08 | 3.58247E-09 | 3.8294E-08 | 5.4503E-08 | 2.49E-08 | 7.05E-08 |
| | THC M | 9 | 5.0788E-08 | 1.36182E-08 | 4.53939E-09 | 4.0320E-08 | 6.1255E-08 | 3.00E-08 | 7.48E-08 |
| | THC+CBD M | 10 | 4.1226E-08 | 1.11788E-08 | 3.53504E-09 | 3.3229E-08 | 4.9223E-08 | 2.66E-08 | 6.17E-08 |

Supplemental Table 3.2 Descriptive statistics for *N*-acyl ethanolamines (mol/g) in liver collected from male and female adult mice 2h after receiving an i.p. injection of 10mg/kg CBD, THC, or THC+CBD

| | | N | Mean | Std. Deviation | Std. Error | 95% Confidence Interval for Mean Lower Bound | 95% Confidence Interval for Mean Upper Bound | Minimum | Maximum |
|---|---|---|---|---|---|---|---|---|---|
| *N*-palmitoyl ethanolamine | Veh F | 7 | 7.3285E-11 | 3.15039E-11 | 1.19073E-11 | 4.4148E-11 | 1.0242E-10 | 2.11E-11 | 1.13E-10 |
| | CBD F | 8 | 9.2554E-11 | 3.17534E-11 | 1.12265E-11 | 6.6008E-11 | 1.191E-10 | 4.96E-11 | 1.37E-10 |
| | THC F | 10 | 1.6638E-10 | 8.33701E-11 | 2.63639E-11 | 1.0674E-10 | 2.2602E-10 | 6.12E-11 | 2.89E-10 |
| | THC+CBD F | 10 | 1.5956E-10 | 7.53073E-11 | 2.38143E-11 | 1.0569E-10 | 2.1343E-10 | 4.63E-11 | 2.74E-10 |
| | Veh M | 7 | 1.1329E-10 | 5.0846E-11 | 1.9218E-11 | 6.6262E-11 | 1.6031E-10 | 3.85E-11 | 1.81E-10 |
| | CBD M | 9 | 7.9081E-11 | 2.41653E-11 | 8.05509E-12 | 6.0506E-11 | 9.7656E-11 | 3.66E-11 | 1.08E-10 |
| | THC M | 9 | 9.9125E-11 | 4.35726E-11 | 1.45242E-11 | 6.5632E-11 | 1.3262E-10 | 5.32E-11 | 1.86E-10 |
| | THC+CBD M | 11 | 9.9357E-11 | 5.46059E-11 | 1.64643E-11 | 6.2673E-11 | 1.3604E-10 | 3.79E-11 | 2E-10 |
| *N*-stearoyl ethanolamine | Veh F | 7 | 5.6153E-11 | 2.61055E-11 | 9.86697E-12 | 3.2009E-11 | 8.0296E-11 | 2.39E-11 | 9.82E-11 |
| | CBD F | 8 | 6.6432E-11 | 4.26153E-11 | 1.50668E-11 | 3.0805E-11 | 1.0206E-10 | 1.75E-11 | 1.46E-10 |
| | THC F | 9 | 1.2882E-10 | 3.74008E-11 | 1.24669E-11 | 1.0007E-10 | 1.5757E-10 | 8.10E-11 | 1.82E-10 |
| | THC+CBD F | 10 | 1.4671E-10 | 6.52160E-11 | 2.06231E-11 | 1.0006E-10 | 1.9337E-10 | 5.55E-11 | 2.66E-10 |
| | Veh M | 7 | 1.0342E-10 | 5.01658E-11 | 1.89609E-11 | 5.7022E-11 | 1.4981E-10 | 2.93E-11 | 1.69E-10 |
| | CBD M | 10 | 6.5517E-11 | 2.26230E-11 | 7.15403E-12 | 4.9333E-11 | 8.1700E-11 | 3.62E-11 | 9.71E-11 |
| | THC M | 8 | 7.9701E-11 | 3.53284E-11 | 1.24905E-11 | 5.0166E-11 | 1.0924E-10 | 4.18E-11 | 1.46E-10 |
| | THC+CBD M | 10 | 7.0183E-11 | 2.43422E-11 | 7.69767E-12 | 5.2770E-11 | 8.7596E-11 | 4.79E-11 | 1.25E-10 |
| *N*-oleoyl ethanolamine | Veh F | 7 | 3.3640E-11 | 1.26948E-11 | 4.79818E-12 | 2.1899E-11 | 4.5380E-11 | 1.10E-11 | 4.74E-11 |
| | CBD F | 8 | 3.7143E-11 | 1.97923E-11 | 6.99764E-12 | 2.0596E-11 | 5.3689E-11 | 1.65E-11 | 7.28E-11 |
| | THC F | 10 | 1.3302E-10 | 6.79700E-11 | 2.14940E-11 | 8.4395E-11 | 1.8164E-10 | 4.34E-11 | 2.20E-10 |
| | THC+CBD F | 9 | 1.5741E-10 | 7.77013E-11 | 2.59004E-11 | 9.7683E-11 | 2.1714E-10 | 5.00E-11 | 3.21E-10 |
| | Veh M | 7 | 8.2125E-11 | 3.25242E-11 | 1.22930E-11 | 5.2046E-11 | 1.1221E-10 | 3.33E-11 | 1.37E-10 |
| | CBD M | 10 | 6.9919E-11 | 2.07104E-11 | 6.54920E-12 | 5.5103E-11 | 8.4734E-11 | 2.94E-11 | 9.53E-11 |
| | THC M | 9 | 1.1061E-10 | 5.59223E-11 | 1.86408E-11 | 6.7620E-11 | 1.5359E-10 | 4.41E-11 | 1.85E-10 |
| | THC+CBD M | 11 | 1.1053E-10 | 5.01763E-11 | 1.51287E-11 | 7.6823E-11 | 1.4424E-10 | 4.38E-11 | 1.93E-10 |
| *N*-linoleoyl ethanolamine | Veh F | 8 | 2.6115E-11 | 8.61461E-12 | 3.04572E-12 | 1.8913E-11 | 3.3317E-11 | 1.23E-11 | 3.69E-11 |
| | CBD F | 7 | 2.2564E-11 | 1.29212E-11 | 4.88376E-12 | 1.0613E-11 | 3.4514E-11 | 7.56E-12 | 3.82E-11 |
| | THC F | 10 | 6.3409E-11 | 2.89278E-11 | 9.14777E-12 | 4.2716E-11 | 8.4103E-11 | 1.56E-11 | 9.75E-11 |

| | | | | | | | | | |
|---|---|---|---|---|---|---|---|---|---|
| | THC+CBD F | 9 | 1.4031E-10 | 1.07695E-10 | 3.58984E-11 | 5.7526E-11 | 2.2309E-10 | 2.87E-11 | 4.01E-10 |
| | Veh M | 7 | 3.1560E-11 | 1.23675E-11 | 4.67446E-12 | 2.0122E-11 | 4.2998E-11 | 1.27E-11 | 4.95E-11 |
| | CBD M | 10 | 3.6493E-11 | 8.38725E-12 | 2.65228E-12 | 3.0493E-11 | 4.2493E-11 | 2.33E-11 | 4.72E-11 |
| | THC M | 8 | 8.9480E-11 | 1.03726E-11 | 3.66726E-12 | 8.0809E-11 | 9.8152E-11 | 7.09E-11 | 1.02E-10 |
| | THC+CBD M | 10 | 8.8984E-11 | 3.24842E-11 | 1.02724E-11 | 6.5747E-11 | 1.1222E-10 | 4.24E-11 | 1.41E-10 |
| *N*-arachidonoyl ethanolamine | Veh F | 8 | 2.7101E-12 | 1.06116E-12 | 3.75176E-13 | 1.8230E-12 | 3.5973E-12 | 1.25E-12 | 3.84E-12 |
| | CBD F | 7 | 2.9020E-12 | 1.42166E-12 | 5.37336E-13 | 1.5871E-12 | 4.2168E-12 | 1.12E-12 | 5.16E-12 |
| | THC F | 10 | 8.2878E-12 | 2.70213E-12 | 8.54490E-13 | 6.3549E-12 | 1.0221E-11 | 3.04E-12 | 1.15E-11 |
| | THC+CBD F | 9 | 1.4850E-11 | 1.27143E-11 | 4.23809E-12 | 5.0770E-12 | 2.4623E-11 | 4.25E-12 | 4.77E-11 |
| | Veh M | 7 | 2.3995E-12 | 1.18773E-12 | 4.48920E-13 | 1.3010E-12 | 3.4980E-12 | 1.18E-12 | 4.10E-12 |
| | CBD M | 10 | 3.2431E-12 | 1.04431E-12 | 3.30238E-13 | 2.4961E-12 | 3.9902E-12 | 1.52E-12 | 4.68E-12 |
| | THC M | 9 | 1.2307E-11 | 4.33079E-12 | 1.44360E-12 | 8.9784E-12 | 1.5636E-11 | 7.43E-12 | 1.95E-11 |
| | THC+CBD M | 11 | 1.0981E-11 | 3.78367E-12 | 1.14082E-12 | 8.4395E-12 | 1.3523E-11 | 3.75E-12 | 1.76E-11 |
| *N*-docosahexaenoyl ethanolamine | Veh F | 8 | 4.8097E-11 | 3.44001E-11 | 1.21623E-11 | 1.9338E-11 | 7.6856E-11 | 1.35E-11 | 1.05E-10 |
| | CBD F | 8 | 9.7888E-11 | 6.41639E-11 | 2.26854E-11 | 4.4246E-11 | 1.5153E-10 | 2.86E-11 | 2.09E-10 |
| | THC F | 9 | 1.7427E-11 | 9.94607E-12 | 3.31536E-12 | 9.7820E-12 | 2.5072E-11 | 6.96E-12 | 3.65E-11 |
| | THC+CBD F | 9 | 1.9518E-11 | 1.04210E-11 | 3.47367E-12 | 1.1508E-11 | 2.7528E-11 | 9.18E-12 | 4.30E-11 |
| | Veh M | 7 | 6.3017E-12 | 2.09571E-12 | 7.92104E-13 | 4.3635E-12 | 8.2399E-12 | 3.40E-12 | 8.49E-12 |
| | CBD M | 10 | 9.1694E-12 | 3.93477E-12 | 1.24428E-12 | 6.3546E-12 | 1.1984E-11 | 3.23E-12 | 1.47E-11 |
| | THC M | 9 | 1.9953E-11 | 1.11136E-11 | 3.70452E-12 | 1.1410E-11 | 2.8496E-11 | 8.02E-12 | 3.66E-11 |
| | THC+CBD M | 11 | 9.8299E-11 | 5.44843E-11 | 1.64276E-11 | 6.1696E-11 | 1.3490E-10 | 3.56E-11 | 1.87E-10 |

Supplemental Table 3.3 Independent t-test comparing male and female levels of N-acyl ethanolamines measured in plasma collected from adult mice 2h after an i.p. injection of 10mg/kg CBD, THC, or THC+CBD.

| | | | Levene's Test for Equality of Variances | | t-test for Equality of Means | | | | | | |
|---|---|---|---|---|---|---|---|---|---|---|---|
| | | | | | | | Significance | | | 95% Confidence Interval of the Difference | |
| | | | F | Sig. | t | df | One-Sided p | Two-Sided p | Std. Error Difference | Lower | Upper |
| Veh | *N*-palmitoyl ethanolamine | Equal variances assumed | 0.419 | 0.529 | -1.826 | 13 | 0.045 | 0.091 | 3.11588E-09 | -1.24216E-08 | 1.04135E-09 |
| | | Equal variances not assumed | | | -1.798 | 11.526 | 0.049 | 0.098 | 3.16486E-09 | -1.26173E-08 | 1.23711E-09 |
| | *N*-stearoyl ethanolamine | Equal variances assumed | 12.507 | 0.004 | -1.539 | 13 | 0.074 | 0.148 | 5.60621E-09 | -2.07413E-08 | 3.48166E-09 |
| | | Equal variances not assumed | | | -1.633 | 8.533 | 0.069 | 0.139 | 5.28617E-09 | -2.06884E-08 | 3.42877E-09 |
| | *N*-oleoyl ethanolamine | Equal variances assumed | 1.135 | 0.306 | -1.132 | 13 | 0.139 | 0.278 | 6.31179E-09 | -2.07832E-08 | 6.48841E-09 |
| | | Equal variances not assumed | | | -1.160 | 12.491 | 0.134 | 0.268 | 6.16040E-09 | -2.05115E-08 | 6.21675E-09 |
| | *N*-linoleoyl ethanolamine | Equal variances assumed | 0.079 | 0.783 | 0.216 | 13 | 0.416 | 0.832 | 3.07380E-09 | -5.97689E-09 | 7.30420E-09 |
| | | Equal variances not assumed | | | 0.217 | 12.902 | 0.416 | 0.832 | 3.06127E-09 | -5.95491E-09 | 7.28223E-09 |
| | *N*-arachidonoyl ethanolamine | Equal variances assumed | 0.003 | 0.960 | -1.274 | 13 | 0.113 | 0.225 | 6.81472E-10 | -2.34023E-09 | 6.04232E-10 |
| | | Equal variances not assumed | | | -1.257 | 11.756 | 0.117 | 0.233 | 6.90534E-10 | -2.37602E-09 | 6.40020E-10 |
| | *N*-docosahexaenoyl ethanolamine | Equal variances assumed | 0.585 | 0.458 | -0.577 | 13 | 0.287 | 0.574 | 4.93990E-09 | -1.35205E-08 | 7.82351E-09 |
| | | Equal variances not assumed | | | -0.580 | 12.964 | 0.286 | 0.572 | 4.90771E-09 | -1.34539E-08 | 7.75694E-09 |
| CBD | *N*-palmitoyl ethanolamine | Equal variances assumed | 5.230 | 0.035 | -0.680 | 17 | 0.253 | 0.506 | 4.35719E-09 | -1.21564E-08 | 6.22933E-09 |
| | | Equal variances not assumed | | | -0.742 | 16.222 | 0.234 | 0.469 | 3.99664E-09 | -1.14266E-08 | 5.49953E-09 |

| | | | | | | | | | | | |
|---|---|---|---|---|---|---|---|---|---|---|---|
| | *N*-stearoyl ethanolamine | Equal variances assumed | 0.423 | 0.524 | -0.125 | 17 | 0.451 | 0.902 | 2.91888E-09 | -6.52299E-09 | 5.79360E-09 |
| | | Equal variances not assumed | | | -0.127 | 16.739 | 0.450 | 0.901 | 2.88110E-09 | -6.45052E-09 | 5.72114E-09 |
| | *N*-oleoyl ethanolamine | Equal variances assumed | 3.594 | 0.074 | 0.255 | 18 | 0.401 | 0.801 | 7.40567E-09 | -1.36676E-08 | 1.74499E-08 |
| | | Equal variances not assumed | | | 0.270 | 16.147 | 0.395 | 0.791 | 7.00769E-09 | -1.29535E-08 | 1.67358E-08 |
| | *N*-linoleoyl ethanolamine | Equal variances assumed | 0.958 | 0.341 | -0.428 | 18 | 0.337 | 0.674 | 2.67676E-09 | -6.76987E-09 | 4.47746E-09 |
| | | Equal variances not assumed | | | -0.442 | 17.810 | 0.332 | 0.664 | 2.59145E-09 | -6.59482E-09 | 4.30240E-09 |
| | *N*-arachidonoyl ethanolamine | Equal variances assumed | 1.173 | 0.294 | -1.168 | 17 | 0.129 | 0.259 | 7.10157E-10 | -2.32800E-09 | 6.68597E-10 |
| | | Equal variances not assumed | | | -1.268 | 16.415 | 0.111 | 0.222 | 6.54203E-10 | -2.21371E-09 | 5.54301E-10 |
| | *N*-docosahexaenoyl ethanolamine | Equal variances assumed | 1.226 | 0.284 | -1.832 | 16 | 0.043 | 0.086 | 4.29921E-09 | -1.69907E-08 | 1.23716E-09 |
| | | Equal variances not assumed | | | -1.989 | 12.634 | 0.034 | 0.069 | 3.95962E-09 | -1.64563E-08 | 7.02770E-10 |
| THC | *N*-palmitoyl ethanolamine | Equal variances assumed | 1.948 | 0.181 | 1.586 | 17 | 0.066 | 0.131 | 4.40664E-09 | -2.30699E-09 | 1.62874E-08 |
| | | Equal variances not assumed | | | 1.626 | 15.223 | 0.062 | 0.124 | 4.29927E-09 | -2.16178E-09 | 1.61422E-08 |
| | *N*-stearoyl ethanolamine | Equal variances assumed | 0.838 | 0.373 | 0.764 | 17 | 0.228 | 0.456 | 3.59195E-09 | -4.83534E-09 | 1.03214E-08 |
| | | Equal variances not assumed | | | 0.779 | 16.034 | 0.224 | 0.447 | 3.52247E-09 | -4.72299E-09 | 1.02090E-08 |
| | *N*-oleoyl ethanolamine | Equal variances assumed | 1.021 | 0.327 | 2.125 | 16 | 0.025 | 0.049 | 5.32382E-09 | 2.94446E-11 | 2.26014E-08 |
| | | Equal variances not assumed | | | 2.229 | 15.483 | 0.021 | 0.041 | 5.07658E-09 | 5.24306E-10 | 2.21066E-08 |
| | *N*-linoleoyl ethanolamine | Equal variances assumed | 0.459 | 0.508 | -1.072 | 15 | 0.150 | 0.301 | 3.00777E-09 | -9.63601E-09 | 3.18580E-09 |
| | | Equal variances not assumed | | | -1.088 | 14.805 | 0.147 | 0.294 | 2.96332E-09 | -9.54853E-09 | 3.09832E-09 |

| | | | | | | | | | | | |
|---|---|---|---|---|---|---|---|---|---|---|---|
| | *N*-arachidonoyl ethanolamine | Equal variances assumed | 0.005 | 0.944 | 0.949 | 16 | 0.178 | 0.357 | 7.15957E-10 | -8.38034E-10 | 2.19749E-09 |
| | | Equal variances not assumed | | | 0.949 | 15.884 | 0.178 | 0.357 | 7.15957E-10 | -8.38939E-10 | 2.19839E-09 |
| | *N*-docosahexaenoyl ethanolamine | Equal variances assumed | 0.346 | 0.564 | -1.088 | 16 | 0.146 | 0.293 | 6.95091E-09 | -2.22988E-08 | 7.17177E-09 |
| | | Equal variances not assumed | | | -1.088 | 15.662 | 0.146 | 0.293 | 6.95091E-09 | -2.23247E-08 | 7.19769E-09 |
| THC+CBD | *N*-palmitoyl ethanolamine | Equal variances assumed | 0.146 | 0.707 | -0.220 | 19 | 0.414 | 0.828 | 6.12262E-09 | -1.41616E-08 | 1.14680E-08 |
| | | Equal variances not assumed | | | -0.223 | 18.352 | 0.413 | 0.826 | 6.03593E-09 | -1.40104E-08 | 1.13168E-08 |
| | *N*-stearoyl ethanolamine | Equal variances assumed | 3.145 | 0.093 | 0.481 | 18 | 0.318 | 0.637 | 3.53713E-09 | -5.73095E-09 | 9.13151E-09 |
| | | Equal variances not assumed | | | 0.511 | 15.497 | 0.308 | 0.617 | 3.32717E-09 | -5.37167E-09 | 8.77222E-09 |
| | *N*-oleoyl ethanolamine | Equal variances assumed | 0.173 | 0.682 | 0.072 | 18 | 0.472 | 0.943 | 6.36821E-09 | -1.29193E-08 | 1.38389E-08 |
| | | Equal variances not assumed | | | 0.072 | 17.483 | 0.472 | 0.943 | 6.36821E-09 | -1.29477E-08 | 1.38673E-08 |
| | *N*-linoleoyl ethanolamine | Equal variances assumed | 0.023 | 0.880 | 2.379 | 18 | 0.014 | 0.029 | 3.60766E-09 | 1.00300E-09 | 1.61618E-08 |
| | | Equal variances not assumed | | | 2.379 | 17.857 | 0.014 | 0.029 | 3.60766E-09 | 9.98633E-10 | 1.61662E-08 |
| | *N*-arachidonoyl ethanolamine | Equal variances assumed | 0.045 | 0.835 | 0.208 | 19 | 0.419 | 0.837 | 8.96993E-10 | -1.69062E-09 | 2.06424E-09 |
| | | Equal variances not assumed | | | 0.208 | 18.845 | 0.419 | 0.837 | 8.96557E-10 | -1.69075E-09 | 2.06437E-09 |
| | *N*-docosahexaenoyl ethanolamine | Equal variances assumed | 6.987 | 0.016 | -0.464 | 19 | 0.324 | 0.648 | 3.54560E-09 | -9.06738E-09 | 5.77469E-09 |
| | | Equal variances not assumed | | | -0.446 | 10.651 | 0.332 | 0.665 | 3.69407E-09 | -9.80957E-09 | 6.51687E-09 |

#### Supplemental Table 3.4 Independent t-test comparing male and female levels of N-acyl ethanolamines measured in liver collected from adult mice 2h after an i.p. injection of 10mg/kg CBD, THC, or THC+CBD.

| | | | Levene's Test for Equality of Variances | | t-test for Equality of Means | | | | | | | |
|---|---|---|---|---|---|---|---|---|---|---|---|---|
| | | | | | | | Significance | | | | 95% Confidence Interval of the Difference | |
| | | | F | Sig. | t | df | One-Sided p | Two-Sided p | Mean Difference | Std. Error Difference | Lower | Upper |
| Veh | *N*-palmitoyl ethanolamine | Equal variances assumed | 3.108 | 0.103 | -1.769 | 12 | 0.051 | 0.102 | -4.00017E-11 | 2.26079E-11 | -8.92600E-11 | 9.25664E-12 |
| | | Equal variances not assumed | | | -1.769 | 10.015 | 0.054 | 0.107 | -4.00017E-11 | 2.26079E-11 | -9.03649E-11 | 1.03615E-11 |
| | *N*-stearoyl ethanolamine | Equal variances assumed | 2.955 | 0.111 | -2.211 | 12 | 0.024 | 0.047 | -4.72646E-11 | 2.13746E-11 | -9.38358E-11 | -6.93422E-13 |
| | | Equal variances not assumed | | | -2.211 | 9.028 | 0.027 | 0.054 | -4.72646E-11 | 2.13746E-11 | -9.55947E-11 | 1.06551E-12 |
| | *N*-oleoyl ethanolamine | Equal variances assumed | 2.189 | 0.165 | -3.674 | 12 | 0.002 | 0.003 | -4.84856E-11 | 1.31962E-11 | -7.72377E-11 | -1.97336E-11 |
| | | Equal variances not assumed | | | -3.674 | 7.787 | 0.003 | 0.007 | -4.84856E-11 | 1.31962E-11 | -7.90618E-11 | -1.79094E-11 |
| | *N*-linoleoyl ethanolamine | Equal variances assumed | 1.328 | 0.270 | -1.001 | 13 | 0.168 | 0.335 | -5.44529E-12 | 5.44176E-12 | -1.72015E-11 | 6.31091E-12 |
| | | Equal variances not assumed | | | -0.976 | 10.547 | 0.175 | 0.351 | -5.44529E-12 | 5.57916E-12 | -1.77897E-11 | 6.89907E-12 |
| | *N*-arachidonoyl ethanolamine | Equal variances assumed | 0.544 | 0.474 | 0.535 | 13 | 0.301 | 0.602 | 3.10601E-13 | 5.80355E-13 | -9.43181E-13 | 1.56438E-12 |
| | | Equal variances not assumed | | | 0.531 | 12.205 | 0.303 | 0.605 | 3.10601E-13 | 5.85052E-13 | -9.61748E-13 | 1.58295E-12 |
| | *N*-docosahexaenoyl ethanolamine | Equal variances assumed | 15.231 | 0.002 | 3.194 | 13 | 0.004 | 0.007 | 4.17956E-11 | 1.30851E-11 | 1.35269E-11 | 7.00643E-11 |
| | | Equal variances not assumed | | | 3.429 | 7.059 | 0.005 | 0.011 | 4.17956E-11 | 1.21880E-11 | 1.30246E-11 | 7.05666E-11 |
| CBD | *N*-palmitoyl ethanolamine | Equal variances assumed | 0.272 | 0.610 | 0.992 | 15 | 0.169 | 0.337 | 1.34735E-11 | 1.35880E-11 | -1.54886E-11 | 4.24356E-11 |
| | | Equal variances not assumed | | | 0.975 | 13.039 | 0.174 | 0.347 | 1.34735E-11 | 1.38174E-11 | -1.63680E-11 | 4.33151E-11 |
| | *N*-stearoyl ethanolamine | Equal variances assumed | 2.020 | 0.174 | 0.059 | 16 | 0.477 | 0.954 | 9.15535E-13 | 1.56059E-11 | -3.21675E-11 | 3.39985E-11 |

| | | | | | | | | | | | |
|---|---|---|---|---|---|---|---|---|---|---|---|
| | | Equal variances not assumed | | | 0.055 | 10.112 | 0.479 | 0.957 | 9.15535E-13 | 1.66790E-11 | -3.61916E-11 | 3.80227E-11 |
| | *N*-oleoyl ethanolamine | Equal variances assumed | 0.033 | 0.858 | -3.402 | 16 | 0.002 | 0.004 | -3.27760E-11 | 9.63570E-12 | -5.32027E-11 | -1.23492E-11 |
| | | Equal variances not assumed | | | -3.420 | 15.427 | 0.002 | 0.004 | -3.27760E-11 | 9.58431E-12 | -5.31553E-11 | -1.23967E-11 |
| | *N*-linoleoyl ethanolamine | Equal variances assumed | 2.459 | 0.138 | -2.707 | 15 | 0.008 | 0.016 | -1.39292E-11 | 5.14482E-12 | -2.48951E-11 | -2.96325E-12 |
| | | Equal variances not assumed | | | -2.506 | 9.510 | 0.016 | 0.032 | -1.39292E-11 | 5.55749E-12 | -2.63991E-11 | -1.45922E-12 |
| | *N*-arachidonoyl ethanolamine | Equal variances assumed | 0.838 | 0.375 | -0.572 | 15 | 0.288 | 0.576 | -3.41176E-13 | 5.96027E-13 | -1.61158E-12 | 9.29226E-13 |
| | | Equal variances not assumed | | | -0.541 | 10.399 | 0.300 | 0.600 | -3.41176E-13 | 6.30704E-13 | -1.73919E-12 | 1.05684E-12 |
| | *N*-docosahexaenoyl ethanolamine | Equal variances assumed | 17.317 | 0.001 | 4.396 | 16 | 0.000 | 0.000 | 8.87189E-11 | 2.01799E-11 | 4.59395E-11 | 1.31498E-10 |
| | | Equal variances not assumed | | | 3.905 | 7.042 | 0.003 | 0.006 | 8.87189E-11 | 2.27195E-11 | 3.50610E-11 | 1.42377E-10 |
| | *N*-palmitoyl ethanolamine | Equal variances assumed | 8.452 | 0.010 | 2.165 | 17 | 0.022 | 0.045 | 6.72556E-11 | 3.10716E-11 | 1.70029E-12 | 1.32811E-10 |
| | | Equal variances not assumed | | | 2.234 | 13.856 | 0.021 | 0.042 | 6.72556E-11 | 3.01000E-11 | 2.63468E-12 | 1.31877E-10 |
| | *N*-stearoyl ethanolamine | Equal variances assumed | 0.202 | 0.660 | 2.773 | 15 | 0.007 | 0.014 | 4.91183E-11 | 1.77107E-11 | 1.13688E-11 | 8.68678E-11 |
| | | Equal variances not assumed | | | 2.783 | 14.930 | 0.007 | 0.014 | 4.91183E-11 | 1.76476E-11 | 1.14880E-11 | 8.67487E-11 |
| THC | *N*-oleoyl ethanolamine | Equal variances assumed | 1.335 | 0.264 | 0.779 | 17 | 0.223 | 0.446 | 2.24123E-11 | 2.87581E-11 | -3.82621E-11 | 8.30866E-11 |
| | | Equal variances not assumed | | | 0.788 | 16.884 | 0.221 | 0.442 | 2.24123E-11 | 2.84512E-11 | -3.76459E-11 | 8.24704E-11 |
| | *N*-linoleoyl ethanolamine | Equal variances assumed | 7.846 | 0.013 | -2.415 | 16 | 0.014 | 0.028 | -2.60709E-11 | 1.07935E-11 | -4.89522E-11 | -3.18965E-12 |
| | | Equal variances not assumed | | | -2.645 | 11.736 | 0.011 | 0.022 | -2.60709E-11 | 9.85548E-12 | -4.75980E-11 | -4.54391E-12 |
| | *N*-arachidonoyl ethanolamine | Equal variances assumed | 2.906 | 0.106 | -2.456 | 17 | 0.013 | 0.025 | -4.01950E-12 | 1.63688E-12 | -7.47301E-12 | -5.65986E-13 |
| | | Equal variances | | | -2.396 | 13.153 | 0.016 | 0.032 | -4.01950E-12 | 1.67753E-12 | -7.63932E-12 | -3.99675E-13 |

| | | | | | | | | | | | | |
|---|---|---|---|---|---|---|---|---|---|---|---|---|
| | | not assumed | | | | | | | | | | |
| | *N*-docosahexaenoyl ethanolamine | Equal variances assumed | 0.939 | 0.347 | -0.508 | 16 | 0.309 | 0.618 | -2.52575E-12 | 4.97142E-12 | -1.30647E-11 | 8.01319E-12 |
| | | Equal variances not assumed | | | -0.508 | 15.807 | 0.309 | 0.618 | -2.52575E-12 | 4.97142E-12 | -1.30752E-11 | 8.02367E-12 |
| | *N*-palmitoyl ethanolamine | Equal variances assumed | 1.083 | 0.311 | 2.112 | 19 | 0.024 | 0.048 | 6.02045E-11 | 2.85036E-11 | 5.45699E-13 | 1.19863E-10 |
| | | Equal variances not assumed | | | 2.079 | 16.307 | 0.027 | 0.054 | 6.02045E-11 | 2.89515E-11 | -1.07635E-12 | 1.21485E-10 |
| | *N*-stearoyl ethanolamine | Equal variances assumed | 5.843 | 0.026 | 3.477 | 18 | 0.001 | 0.003 | 7.65318E-11 | 2.20129E-11 | 3.02844E-11 | 1.22779E-10 |
| | | Equal variances not assumed | | | 3.477 | 11.460 | 0.002 | 0.005 | 7.65318E-11 | 2.20129E-11 | 2.83179E-11 | 1.24746E-10 |
| | *N*-oleoyl ethanolamine | Equal variances assumed | 0.748 | 0.398 | 1.632 | 18 | 0.060 | 0.120 | 4.68778E-11 | 2.87168E-11 | -1.34540E-11 | 1.07209E-10 |
| | | Equal variances not assumed | | | 1.563 | 13.164 | 0.071 | 0.142 | 4.68778E-11 | 2.99952E-11 | -1.78408E-11 | 1.11596E-10 |
| THC+CBD | *N*-linoleoyl ethanolamine | Equal variances assumed | 3.305 | 0.087 | 1.440 | 17 | 0.084 | 0.168 | 5.13238E-11 | 3.56396E-11 | -2.38692E-11 | 1.26517E-10 |
| | | Equal variances not assumed | | | 1.375 | 9.308 | 0.101 | 0.201 | 5.13238E-11 | 3.73392E-11 | -3.27189E-11 | 1.35366E-10 |
| | *N*-arachidonoyl ethanolamine | Equal variances assumed | 1.957 | 0.179 | 0.964 | 18 | 0.174 | 0.348 | 3.86862E-12 | 4.01510E-12 | -4.56679E-12 | 1.23040E-11 |
| | | Equal variances not assumed | | | 0.881 | 9.163 | 0.200 | 0.401 | 3.86862E-12 | 4.38895E-12 | -6.03304E-12 | 1.37703E-11 |
| | *N*-docosahexaenoyl ethanolamine | Equal variances assumed | 17.208 | 0.001 | -4.254 | 18 | 0.000 | 0.000 | -7.87808E-11 | 1.85181E-11 | -1.17686E-10 | -3.98758E-11 |
| | | Equal variances not assumed | | | -4.692 | 10.887 | 0.000 | 0.001 | -7.87808E-11 | 1.67909E-11 | -1.15784E-10 | -4.17775E-11 |

Supplemental Table 3.5 One-way ANOVA for levels of *N*-acyl ethanolamines in plasma in each male and female treatment group separately

| | | Sum of Squares | df | Mean Square | F | Sig. |
|---|---|---|---|---|---|---|
| *N*-palmitoyl ethanolamine | Between Groups | 1.29E-15 | 7 | 1.85E-16 | 1.680 | 0.1293167 |
| | Within Groups | 7.26E-15 | 66 | 1.10E-16 | | |
| | Total | 8.55E-15 | 73 | | | |
| *N*-stearoyl ethanolamine | Between Groups | 5.05E-16 | 7 | 7.21E-17 | 1.074 | 0.3904188 |
| | Within Groups | 4.37E-15 | 65 | 6.72E-17 | | |
| | Total | 4.87E-15 | 72 | | | |
| *N*-oleoyl ethanolamine | Between Groups | 2.69E-15 | 7 | 3.84E-16 | 1.998 | 0.0687669 |
| | Within Groups | 1.25E-14 | 65 | 1.92E-16 | | |
| | Total | 1.52E-14 | 72 | | | |
| *N*-linoleoyl ethanolamine | Between Groups | 9.24E-16 | 7 | 1.32E-16 | 2.970 | 0.0092458 |
| | Within Groups | 2.84E-15 | 64 | 4.44E-17 | | |
| | Total | 3.77E-15 | 71 | | | |
| *N*-arachidonoyl ethanolamine | Between Groups | 1.10E-17 | 7 | 1.57E-18 | 0.570 | 0.7778803 |
| | Within Groups | 1.79E-16 | 65 | 2.76E-18 | | |
| | Total | 1.90E-16 | 72 | | | |
| *N*-docosahexaenoyl ethanolamine | Between Groups | 1.12E-15 | 7 | 1.60E-16 | 1.420 | 0.2127996 |
| | Within Groups | 7.23E-15 | 64 | 1.13E-16 | | |
| | Total | 8.35E-15 | 71 | | | |

#### Supplemental Table 3.6 One-way ANOVA for levels of *N*-acyl ethanolamines in liver in each male and female treatment group separately

| | | Sum of Squares | df | Mean Square | F | Sig. |
|---|---|---|---|---|---|---|
| *N*-palmitoyl ethanolamine | Between Groups | 7.87E-20 | 7 | 1.12E-20 | 3.692 | 0.00210 |
| | Within Groups | 1.92E-19 | 63 | 3.04E-21 | | |
| | Total | 2.70E-19 | 70 | | | |
| *N*-stearoyl ethanolamine | Between Groups | 7.02E-20 | 7 | 1.00E-20 | 6.112 | 0.00002 |
| | Within Groups | 1.00E-19 | 61 | 1.64E-21 | | |
| | Total | 1.70E-19 | 68 | | | |
| *N*-oleoyl ethanolamine | Between Groups | 1.15E-19 | 7 | 1.64E-20 | 6.717 | 0.00001 |
| | Within Groups | 1.54E-19 | 63 | 2.44E-21 | | |
| | Total | 2.69E-19 | 70 | | | |
| *N*-linoleoyl ethanolamine | Between Groups | 1.02E-19 | 7 | 1.46E-20 | 7.841 | 0.00000 |
| | Within Groups | 1.14E-19 | 61 | 1.86E-21 | | |
| | Total | 2.16E-19 | 68 | | | |
| *N*-arachidonoyl ethanolamine | Between Groups | 1.53E-21 | 7 | 2.18E-22 | 8.135 | 0.00000 |
| | Within Groups | 1.69E-21 | 63 | 2.68E-23 | | |
| | Total | 3.22E-21 | 70 | | | |
| *N*-docosahexaenoyl ethanolamine | Between Groups | 9.41E-20 | 7 | 1.34E-20 | 12.169 | 0.00000 |
| | Within Groups | 6.96E-20 | 63 | 1.10E-21 | | |
| | Total | 1.64E-19 | 70 | | | |

#### Supplemental Table 3.7 Fisher's LSD post-hoc test for plasma *N*-acyl ethanolamine ANOVA

| Dependent Variable | | | Std. Error | Sig. | 95% Confidence Interval Lower Bound | 95% Confidence Interval Upper Bound |
|---|---|---|---|---|---|---|
| *N*-palmitoyl ethanolamine | VehFemale | CBDfemale | 5.43E-09 | 0.3190 | -1.63E-08 | 5.39E-09 |
| | | THCfemale | 5.17E-09 | 0.2415 | -1.64E-08 | 4.21E-09 |
| | | TpCfemale | 5.07E-09 | 0.0431 | -2.06E-08 | -3.33E-10 |
| | | VehMale | 5.43E-09 | 0.2984 | -1.65E-08 | 5.15E-09 |
| | | CBDmale | 5.07E-09 | 0.1018 | -1.85E-08 | 1.71E-09 |
| | | THCmale | 5.29E-09 | 0.8680 | -9.67E-09 | 1.14E-08 |
| | | TpCmale | 5.17E-09 | 0.0256 | -2.21E-08 | -1.48E-09 |
| | CBDfemale | VehFemale | 5.43E-09 | 0.3190 | -5.39E-09 | 1.63E-08 |
| | | THCfemale | 4.98E-09 | 0.8952 | -1.06E-08 | 9.28E-09 |
| | | TpCfemale | 4.87E-09 | 0.3080 | -1.47E-08 | 4.72E-09 |
| | | VehMale | 5.24E-09 | 0.9637 | -1.07E-08 | 1.02E-08 |
| | | CBDmale | 4.87E-09 | 0.5452 | -1.27E-08 | 6.77E-09 |
| | | THCmale | 5.10E-09 | 0.2184 | -3.84E-09 | 1.65E-08 |
| | | TpCmale | 4.98E-09 | 0.2060 | -1.63E-08 | 3.58E-09 |
| | THCfemale | VehFemale | 5.17E-09 | 0.2415 | -4.21E-09 | 1.64E-08 |
| | | CBDfemale | 4.98E-09 | 0.8952 | -9.28E-09 | 1.06E-08 |
| | | TpCfemale | 4.58E-09 | 0.3461 | -1.35E-08 | 4.80E-09 |
| | | VehMale | 4.98E-09 | 0.9333 | -9.51E-09 | 1.04E-08 |
| | | CBDmale | 4.58E-09 | 0.6165 | -1.15E-08 | 6.84E-09 |
| | | THCmale | 4.82E-09 | 0.1517 | -2.63E-09 | 1.66E-08 |
| | | TpCmale | 4.69E-09 | 0.2289 | -1.51E-08 | 3.67E-09 |
| | TpCfemale | VehFemale | 5.07E-09 | 0.0431 | 3.33E-10 | 2.06E-08 |
| | | CBDfemale | 4.87E-09 | 0.3080 | -4.72E-09 | 1.47E-08 |
| | | THCfemale | 4.58E-09 | 0.3461 | -4.80E-09 | 1.35E-08 |
| | | VehMale | 4.87E-09 | 0.3315 | -4.96E-09 | 1.45E-08 |
| | | CBDmale | 4.47E-09 | 0.6492 | -6.89E-09 | 1.10E-08 |
| | | THCmale | 4.71E-09 | 0.0190 | 1.93E-09 | 2.08E-08 |
| | | TpCmale | 4.58E-09 | 0.7698 | -1.05E-08 | 7.80E-09 |
| | VehMale | VehFemale | 5.43E-09 | 0.2984 | -5.15E-09 | 1.65E-08 |
| | | CBDfemale | 5.24E-09 | 0.9637 | -1.02E-08 | 1.07E-08 |
| | | THCfemale | 4.98E-09 | 0.9333 | -1.04E-08 | 9.51E-09 |
| | | TpCfemale | 4.87E-09 | 0.3315 | -1.45E-08 | 4.96E-09 |
| | | CBDmale | 4.87E-09 | 0.5781 | -1.25E-08 | 7.01E-09 |
| | | THCmale | 5.10E-09 | 0.2017 | -3.60E-09 | 1.67E-08 |
| | | TpCmale | 4.98E-09 | 0.2234 | -1.60E-08 | 3.82E-09 |
| | CBDmale | VehFemale | 5.07E-09 | 0.1018 | -1.71E-09 | 1.85E-08 |
| | | CBDfemale | 4.87E-09 | 0.5452 | -6.77E-09 | 1.27E-08 |
| | | THCfemale | 4.58E-09 | 0.6165 | -6.84E-09 | 1.15E-08 |
| | | TpCfemale | 4.47E-09 | 0.6492 | -1.10E-08 | 6.89E-09 |
| | | VehMale | 4.87E-09 | 0.5781 | -7.01E-09 | 1.25E-08 |
| | | THCmale | 4.71E-09 | 0.0528 | -1.16E-10 | 1.87E-08 |
| | | TpCmale | 4.58E-09 | 0.4621 | -1.25E-08 | 5.76E-09 |
| | THCmale | VehFemale | 5.29E-09 | 0.8680 | -1.14E-08 | 9.67E-09 |
| | | CBDfemale | 5.10E-09 | 0.2184 | -1.65E-08 | 3.84E-09 |
| | | THCfemale | 4.82E-09 | 0.1517 | -1.66E-08 | 2.63E-09 |
| | | TpCfemale | 4.71E-09 | 0.0190 | -2.08E-08 | -1.93E-09 |
| | | VehMale | 5.10E-09 | 0.2017 | -1.67E-08 | 3.60E-09 |
| | | CBDmale | 4.71E-09 | 0.0528 | -1.87E-08 | 1.16E-10 |
| | | TpCmale | 4.82E-09 | 0.0105 | -2.23E-08 | -3.06E-09 |
| | TpCmale | VehFemale | 5.17E-09 | 0.0256 | 1.48E-09 | 2.21E-08 |
| | | CBDfemale | 4.98E-09 | 0.2060 | -3.58E-09 | 1.63E-08 |
| | | THCfemale | 4.69E-09 | 0.2289 | -3.67E-09 | 1.51E-08 |
| | | TpCfemale | 4.58E-09 | 0.7698 | -7.80E-09 | 1.05E-08 |
| | | VehMale | 4.98E-09 | 0.2234 | -3.82E-09 | 1.60E-08 |
| | | CBDmale | 4.58E-09 | 0.4621 | -5.76E-09 | 1.25E-08 |
| | | THCmale | 4.82E-09 | 0.0105 | 3.06E-09 | 2.23E-08 |
| *N*-stearoyl ethanolamine | VehFemale | CBDfemale | 4.13E-09 | 0.3110 | -1.25E-08 | 4.03E-09 |

| | | | | | | |
|---|---|---|---|---|---|---|
| | | THCfemale | 4.04E-09 | 0.2187 | -1.31E-08 | 3.05E-09 |
| | | TpCfemale | 3.96E-09 | 0.0381 | -1.63E-08 | -4.73E-10 |
| | | VehMale | 4.24E-09 | 0.0460 | -1.71E-08 | -1.59E-10 |
| | | CBDmale | 4.04E-09 | 0.2608 | -1.26E-08 | 3.48E-09 |
| | | THCmale | 4.13E-09 | 0.5840 | -1.05E-08 | 5.98E-09 |
| | | TpCmale | 4.13E-09 | 0.1103 | -1.49E-08 | 1.56E-09 |
| | CBDfemale | VehFemale | 4.13E-09 | 0.3110 | -4.03E-09 | 1.25E-08 |
| | | THCfemale | 3.77E-09 | 0.8326 | -8.32E-09 | 6.72E-09 |
| | | TpCfemale | 3.68E-09 | 0.2618 | -1.15E-08 | 3.19E-09 |
| | | VehMale | 3.98E-09 | 0.2718 | -1.24E-08 | 3.54E-09 |
| | | CBDmale | 3.77E-09 | 0.9231 | -7.88E-09 | 7.16E-09 |
| | | THCmale | 3.86E-09 | 0.6165 | -5.77E-09 | 9.66E-09 |
| | | TpCmale | 3.86E-09 | 0.5250 | -1.02E-08 | 5.25E-09 |
| | THCfemale | VehFemale | 4.04E-09 | 0.2187 | -3.05E-09 | 1.31E-08 |
| | | CBDfemale | 3.77E-09 | 0.8326 | -6.72E-09 | 8.32E-09 |
| | | TpCfemale | 3.58E-09 | 0.3500 | -1.05E-08 | 3.78E-09 |
| | | VehMale | 3.89E-09 | 0.3559 | -1.14E-08 | 4.15E-09 |
| | | CBDmale | 3.66E-09 | 0.9060 | -6.89E-09 | 7.75E-09 |
| | | THCmale | 3.77E-09 | 0.4689 | -4.78E-09 | 1.03E-08 |
| | | TpCmale | 3.77E-09 | 0.6588 | -9.19E-09 | 5.85E-09 |
| | TpCfemale | VehFemale | 3.96E-09 | 0.0381 | 4.73E-10 | 1.63E-08 |
| | | CBDfemale | 3.68E-09 | 0.2618 | -3.19E-09 | 1.15E-08 |
| | | THCfemale | 3.58E-09 | 0.3500 | -3.78E-09 | 1.05E-08 |
| | | VehMale | 3.81E-09 | 0.9491 | -7.85E-09 | 7.36E-09 |
| | | CBDmale | 3.58E-09 | 0.2919 | -3.35E-09 | 1.10E-08 |
| | | THCmale | 3.68E-09 | 0.1018 | -1.24E-09 | 1.35E-08 |
| | | TpCmale | 3.68E-09 | 0.6459 | -5.66E-09 | 9.06E-09 |
| | VehMale | VehFemale | 4.24E-09 | 0.0460 | 1.59E-10 | 1.71E-08 |
| | | CBDfemale | 3.98E-09 | 0.2718 | -3.54E-09 | 1.24E-08 |
| | | THCfemale | 3.89E-09 | 0.3559 | -4.15E-09 | 1.14E-08 |
| | | TpCfemale | 3.81E-09 | 0.9491 | -7.36E-09 | 7.85E-09 |
| | | CBDmale | 3.89E-09 | 0.3015 | -3.71E-09 | 1.18E-08 |
| | | THCmale | 3.98E-09 | 0.1152 | -1.60E-09 | 1.43E-08 |
| | | TpCmale | 3.98E-09 | 0.6270 | -6.01E-09 | 9.90E-09 |
| | CBDmale | VehFemale | 4.04E-09 | 0.2608 | -3.48E-09 | 1.26E-08 |
| | | CBDfemale | 3.77E-09 | 0.9231 | -7.16E-09 | 7.88E-09 |
| | | THCfemale | 3.66E-09 | 0.9060 | -7.75E-09 | 6.89E-09 |
| | | TpCfemale | 3.58E-09 | 0.2919 | -1.10E-08 | 3.35E-09 |
| | | VehMale | 3.89E-09 | 0.3015 | -1.18E-08 | 3.71E-09 |
| | | THCmale | 3.77E-09 | 0.5419 | -5.21E-09 | 9.83E-09 |
| | | TpCmale | 3.77E-09 | 0.5781 | -9.62E-09 | 5.42E-09 |
| | THCmale | VehFemale | 4.13E-09 | 0.5840 | -5.98E-09 | 1.05E-08 |
| | | CBDfemale | 3.86E-09 | 0.6165 | -9.66E-09 | 5.77E-09 |
| | | THCfemale | 3.77E-09 | 0.4689 | -1.03E-08 | 4.78E-09 |
| | | TpCfemale | 3.68E-09 | 0.1018 | -1.35E-08 | 1.24E-09 |
| | | VehMale | 3.98E-09 | 0.1152 | -1.43E-08 | 1.60E-09 |
| | | CBDmale | 3.77E-09 | 0.5419 | -9.83E-09 | 5.21E-09 |
| | | TpCmale | 3.86E-09 | 0.2575 | -1.21E-08 | 3.30E-09 |
| | TpCmale | VehFemale | 4.13E-09 | 0.1103 | -1.56E-09 | 1.49E-08 |
| | | CBDfemale | 3.86E-09 | 0.5250 | -5.25E-09 | 1.02E-08 |
| | | THCfemale | 3.77E-09 | 0.6588 | -5.85E-09 | 9.19E-09 |
| | | TpCfemale | 3.68E-09 | 0.6459 | -9.06E-09 | 5.66E-09 |
| | | VehMale | 3.98E-09 | 0.6270 | -9.90E-09 | 6.01E-09 |
| | | CBDmale | 3.77E-09 | 0.5781 | -5.42E-09 | 9.62E-09 |
| | | THCmale | 3.86E-09 | 0.2575 | -3.30E-09 | 1.21E-08 |
| *N*-oleoyl ethanolamine | VehFemale | CBDfemale | 6.98E-09 | 0.2239 | -2.25E-08 | 5.37E-09 |
| | | THCfemale | 6.83E-09 | 0.1035 | -2.49E-08 | 2.36E-09 |
| | | TpCfemale | 6.83E-09 | 0.0134 | -3.10E-08 | -3.73E-09 |
| | | VehMale | 7.17E-09 | 0.3227 | -2.15E-08 | 7.18E-09 |
| | | CBDmale | 6.70E-09 | 0.3221 | -2.01E-08 | 6.70E-09 |

| | | | | | | |
|---|---|---|---|---|---|---|
| | | THCmale | 7.17E-09 | 0.9960 | -1.43E-08 | 1.44E-08 |
| | | TpCmale | 6.83E-09 | 0.0159 | -3.06E-08 | -3.27E-09 |
| | CBDfemale | VehFemale | 6.98E-09 | 0.2239 | -5.37E-09 | 2.25E-08 |
| | | THCfemale | 6.37E-09 | 0.6728 | -1.54E-08 | 1.00E-08 |
| | | TpCfemale | 6.37E-09 | 0.1720 | -2.15E-08 | 3.92E-09 |
| | | VehMale | 6.73E-09 | 0.8325 | -1.20E-08 | 1.49E-08 |
| | | CBDmale | 6.23E-09 | 0.7624 | -1.05E-08 | 1.43E-08 |
| | | THCmale | 6.73E-09 | 0.2054 | -4.84E-09 | 2.21E-08 |
| | | TpCmale | 6.37E-09 | 0.1952 | -2.11E-08 | 4.38E-09 |
| | THCfemale | VehFemale | 6.83E-09 | 0.1035 | -2.36E-09 | 2.49E-08 |
| | | CBDfemale | 6.37E-09 | 0.6728 | -1.00E-08 | 1.54E-08 |
| | | TpCfemale | 6.20E-09 | 0.3293 | -1.85E-08 | 6.29E-09 |
| | | VehMale | 6.57E-09 | 0.5319 | -9.00E-09 | 1.73E-08 |
| | | CBDmale | 6.06E-09 | 0.4509 | -7.50E-09 | 1.67E-08 |
| | | THCmale | 6.57E-09 | 0.0900 | -1.81E-09 | 2.44E-08 |
| | | TpCmale | 6.20E-09 | 0.3669 | -1.80E-08 | 6.75E-09 |
| | TpCfemale | VehFemale | 6.83E-09 | 0.0134 | 3.73E-09 | 3.10E-08 |
| | | CBDfemale | 6.37E-09 | 0.1720 | -3.92E-09 | 2.15E-08 |
| | | THCfemale | 6.20E-09 | 0.3293 | -6.29E-09 | 1.85E-08 |
| | | VehMale | 6.57E-09 | 0.1248 | -2.91E-09 | 2.34E-08 |
| | | CBDmale | 6.06E-09 | 0.0823 | -1.41E-09 | 2.28E-08 |
| | | THCmale | 6.57E-09 | 0.0102 | 4.28E-09 | 3.05E-08 |
| | | TpCmale | 6.20E-09 | 0.9411 | -1.19E-08 | 1.28E-08 |
| | VehMale | VehFemale | 7.17E-09 | 0.3227 | -7.18E-09 | 2.15E-08 |
| | | CBDfemale | 6.73E-09 | 0.8325 | -1.49E-08 | 1.20E-08 |
| | | THCfemale | 6.57E-09 | 0.5319 | -1.73E-08 | 9.00E-09 |
| | | TpCfemale | 6.57E-09 | 0.1248 | -2.34E-08 | 2.91E-09 |
| | | CBDmale | 6.44E-09 | 0.9431 | -1.24E-08 | 1.33E-08 |
| | | THCmale | 6.93E-09 | 0.3037 | -6.66E-09 | 2.10E-08 |
| | | TpCmale | 6.57E-09 | 0.1423 | -2.29E-08 | 3.37E-09 |
| | CBDmale | VehFemale | 6.70E-09 | 0.3221 | -6.70E-09 | 2.01E-08 |
| | | CBDfemale | 6.23E-09 | 0.7624 | -1.43E-08 | 1.05E-08 |
| | | THCfemale | 6.06E-09 | 0.4509 | -1.67E-08 | 7.50E-09 |
| | | TpCfemale | 6.06E-09 | 0.0823 | -2.28E-08 | 1.41E-09 |
| | | VehMale | 6.44E-09 | 0.9431 | -1.33E-08 | 1.24E-08 |
| | | THCmale | 6.44E-09 | 0.3004 | -6.14E-09 | 1.96E-08 |
| | | TpCmale | 6.06E-09 | 0.0961 | -2.23E-08 | 1.87E-09 |
| | THCmale | VehFemale | 7.17E-09 | 0.9960 | -1.44E-08 | 1.43E-08 |
| | | CBDfemale | 6.73E-09 | 0.2054 | -2.21E-08 | 4.84E-09 |
| | | THCfemale | 6.57E-09 | 0.0900 | -2.44E-08 | 1.81E-09 |
| | | TpCfemale | 6.57E-09 | 0.0102 | -3.05E-08 | -4.28E-09 |
| | | VehMale | 6.93E-09 | 0.3037 | -2.10E-08 | 6.66E-09 |
| | | CBDmale | 6.44E-09 | 0.3004 | -1.96E-08 | 6.14E-09 |
| | | TpCmale | 6.57E-09 | 0.0122 | -3.01E-08 | -3.82E-09 |
| | TpCmale | VehFemale | 6.83E-09 | 0.0159 | 3.27E-09 | 3.06E-08 |
| | | CBDfemale | 6.37E-09 | 0.1952 | -4.38E-09 | 2.11E-08 |
| | | THCfemale | 6.20E-09 | 0.3669 | -6.75E-09 | 1.80E-08 |
| | | TpCfemale | 6.20E-09 | 0.9411 | -1.28E-08 | 1.19E-08 |
| | | VehMale | 6.57E-09 | 0.1423 | -3.37E-09 | 2.29E-08 |
| | | CBDmale | 6.06E-09 | 0.0961 | -1.87E-09 | 2.23E-08 |
| | | THCmale | 6.57E-09 | 0.0122 | 3.82E-09 | 3.01E-08 |
| *N*-linoleoyl ethanolamine | VehFemale | CBDfemale | 3.36E-09 | 0.6047 | -4.96E-09 | 8.46E-09 |
| | | THCfemale | 3.36E-09 | 0.5717 | -4.80E-09 | 8.62E-09 |
| | | TpCfemale | 3.28E-09 | 0.0053 | -1.61E-08 | -2.93E-09 |
| | | VehMale | 3.45E-09 | 0.8480 | -6.23E-09 | 7.55E-09 |
| | | CBDmale | 3.22E-09 | 0.8526 | -5.84E-09 | 7.04E-09 |
| | | THCmale | 3.45E-09 | 0.7041 | -8.21E-09 | 5.58E-09 |
| | | TpCmale | 3.28E-09 | 0.7825 | -7.47E-09 | 5.65E-09 |
| | CBDfemale | VehFemale | 3.36E-09 | 0.6047 | -8.46E-09 | 4.96E-09 |
| | | THCfemale | 3.14E-09 | 0.9591 | -6.11E-09 | 6.44E-09 |

| | | | | | | |
|---|---|---|---|---|---|---|
| | | TpCfemale | 3.06E-09 | 0.0005 | -1.74E-08 | -5.12E-09 |
| | | VehMale | 3.24E-09 | 0.7390 | -7.55E-09 | 5.39E-09 |
| | | CBDmale | 3.00E-09 | 0.7033 | -7.13E-09 | 4.84E-09 |
| | | THCmale | 3.24E-09 | 0.3478 | -9.53E-09 | 3.41E-09 |
| | | TpCmale | 3.06E-09 | 0.3887 | -8.78E-09 | 3.46E-09 |
| | THCfemale | VehFemale | 3.36E-09 | 0.5717 | -8.62E-09 | 4.80E-09 |
| | | CBDfemale | 3.14E-09 | 0.9591 | -6.44E-09 | 6.11E-09 |
| | | TpCfemale | 3.06E-09 | 0.0004 | -1.75E-08 | -5.28E-09 |
| | | VehMale | 3.24E-09 | 0.7018 | -7.72E-09 | 5.22E-09 |
| | | CBDmale | 3.00E-09 | 0.6639 | -7.29E-09 | 4.68E-09 |
| | | THCmale | 3.24E-09 | 0.3231 | -9.70E-09 | 3.24E-09 |
| | | TpCmale | 3.06E-09 | 0.3606 | -8.94E-09 | 3.30E-09 |
| | TpCfemale | VehFemale | 3.28E-09 | 0.0053 | 2.93E-09 | 1.61E-08 |
| | | CBDfemale | 3.06E-09 | 0.0005 | 5.12E-09 | 1.74E-08 |
| | | THCfemale | 3.06E-09 | 0.0004 | 5.28E-09 | 1.75E-08 |
| | | VehMale | 3.16E-09 | 0.0021 | 3.84E-09 | 1.65E-08 |
| | | CBDmale | 2.91E-09 | 0.0009 | 4.28E-09 | 1.59E-08 |
| | | THCmale | 3.16E-09 | 0.0120 | 1.86E-09 | 1.45E-08 |
| | | TpCmale | 2.98E-09 | 0.0054 | 2.63E-09 | 1.45E-08 |
| | VehMale | VehFemale | 3.45E-09 | 0.8480 | -7.55E-09 | 6.23E-09 |
| | | CBDfemale | 3.24E-09 | 0.7390 | -5.39E-09 | 7.55E-09 |
| | | THCfemale | 3.24E-09 | 0.7018 | -5.22E-09 | 7.72E-09 |
| | | TpCfemale | 3.16E-09 | 0.0021 | -1.65E-08 | -3.84E-09 |
| | | CBDmale | 3.10E-09 | 0.9840 | -6.25E-09 | 6.12E-09 |
| | | THCmale | 3.33E-09 | 0.5546 | -8.64E-09 | 4.68E-09 |
| | | TpCmale | 3.16E-09 | 0.6203 | -7.89E-09 | 4.74E-09 |
| | CBDmale | VehFemale | 3.22E-09 | 0.8526 | -7.04E-09 | 5.84E-09 |
| | | CBDfemale | 3.00E-09 | 0.7033 | -4.84E-09 | 7.13E-09 |
| | | THCfemale | 3.00E-09 | 0.6639 | -4.68E-09 | 7.29E-09 |
| | | TpCfemale | 2.91E-09 | 0.0009 | -1.59E-08 | -4.28E-09 |
| | | VehMale | 3.10E-09 | 0.9840 | -6.12E-09 | 6.25E-09 |
| | | THCmale | 3.10E-09 | 0.5381 | -8.10E-09 | 4.27E-09 |
| | | TpCmale | 2.91E-09 | 0.6055 | -7.33E-09 | 4.31E-09 |
| | THCmale | VehFemale | 3.45E-09 | 0.7041 | -5.58E-09 | 8.21E-09 |
| | | CBDfemale | 3.24E-09 | 0.3478 | -3.41E-09 | 9.53E-09 |
| | | THCfemale | 3.24E-09 | 0.3231 | -3.24E-09 | 9.70E-09 |
| | | TpCfemale | 3.16E-09 | 0.0120 | -1.45E-08 | -1.86E-09 |
| | | VehMale | 3.33E-09 | 0.5546 | -4.68E-09 | 8.64E-09 |
| | | CBDmale | 3.10E-09 | 0.5381 | -4.27E-09 | 8.10E-09 |
| | | TpCmale | 3.16E-09 | 0.8983 | -5.91E-09 | 6.72E-09 |
| | TpCmale | VehFemale | 3.28E-09 | 0.7825 | -5.65E-09 | 7.47E-09 |
| | | CBDfemale | 3.06E-09 | 0.3887 | -3.46E-09 | 8.78E-09 |
| | | THCfemale | 3.06E-09 | 0.3606 | -3.30E-09 | 8.94E-09 |
| | | TpCfemale | 2.98E-09 | 0.0054 | -1.45E-08 | -2.63E-09 |
| | | VehMale | 3.16E-09 | 0.6203 | -4.74E-09 | 7.89E-09 |
| | | CBDmale | 2.91E-09 | 0.6055 | -4.31E-09 | 7.33E-09 |
| | | THCmale | 3.16E-09 | 0.8983 | -6.72E-09 | 5.91E-09 |
| *N*-arachidonoyl ethanolamine | VehFemale | CBDfemale | 8.59E-10 | 0.6343 | -1.31E-09 | 2.13E-09 |
| | | THCfemale | 8.37E-10 | 0.7787 | -1.91E-09 | 1.44E-09 |
| | | TpCfemale | 8.03E-10 | 0.7581 | -1.85E-09 | 1.36E-09 |
| | | VehMale | 8.59E-10 | 0.3162 | -2.58E-09 | 8.48E-10 |
| | | CBDmale | 8.03E-10 | 0.6036 | -2.02E-09 | 1.18E-09 |
| | | THCmale | 8.37E-10 | 0.5979 | -1.23E-09 | 2.11E-09 |
| | | TpCmale | 8.18E-10 | 0.9403 | -1.70E-09 | 1.57E-09 |
| | CBDfemale | VehFemale | 8.59E-10 | 0.6343 | -2.13E-09 | 1.31E-09 |
| | | THCfemale | 8.07E-10 | 0.4256 | -2.26E-09 | 9.65E-10 |
| | | TpCfemale | 7.72E-10 | 0.3962 | -2.20E-09 | 8.82E-10 |
| | | VehMale | 8.30E-10 | 0.1284 | -2.94E-09 | 3.79E-10 |
| | | CBDmale | 7.72E-10 | 0.2862 | -2.37E-09 | 7.11E-10 |
| | | THCmale | 8.07E-10 | 0.9677 | -1.58E-09 | 1.64E-09 |

| | | | | | | |
|---|---|---|---|---|---|---|
| | | TpCmale | 7.88E-10 | 0.5509 | -2.05E-09 | 1.10E-09 |
| | THCfemale | VehFemale | 8.37E-10 | 0.7787 | -1.44E-09 | 1.91E-09 |
| | | CBDfemale | 8.07E-10 | 0.4256 | -9.65E-10 | 2.26E-09 |
| | | TpCfemale | 7.46E-10 | 0.9871 | -1.50E-09 | 1.48E-09 |
| | | VehMale | 8.07E-10 | 0.4364 | -2.24E-09 | 9.80E-10 |
| | | CBDmale | 7.46E-10 | 0.8073 | -1.67E-09 | 1.31E-09 |
| | | THCmale | 7.83E-10 | 0.3884 | -8.84E-10 | 2.24E-09 |
| | | TpCmale | 7.63E-10 | 0.8196 | -1.35E-09 | 1.70E-09 |
| | TpCfemale | VehFemale | 8.03E-10 | 0.7581 | -1.36E-09 | 1.85E-09 |
| | | CBDfemale | 7.72E-10 | 0.3962 | -8.82E-10 | 2.20E-09 |
| | | THCfemale | 7.46E-10 | 0.9871 | -1.48E-09 | 1.50E-09 |
| | | VehMale | 7.72E-10 | 0.4248 | -2.16E-09 | 9.21E-10 |
| | | CBDmale | 7.08E-10 | 0.8103 | -1.58E-09 | 1.24E-09 |
| | | THCmale | 7.46E-10 | 0.3574 | -7.99E-10 | 2.18E-09 |
| | | TpCmale | 7.26E-10 | 0.7976 | -1.26E-09 | 1.64E-09 |
| | VehMale | VehFemale | 8.59E-10 | 0.3162 | -8.48E-10 | 2.58E-09 |
| | | CBDfemale | 8.30E-10 | 0.1284 | -3.79E-10 | 2.94E-09 |
| | | THCfemale | 8.07E-10 | 0.4364 | -9.80E-10 | 2.24E-09 |
| | | TpCfemale | 7.72E-10 | 0.4248 | -9.21E-10 | 2.16E-09 |
| | | CBDmale | 7.72E-10 | 0.5626 | -1.09E-09 | 1.99E-09 |
| | | THCmale | 8.07E-10 | 0.1089 | -3.00E-10 | 2.92E-09 |
| | | TpCmale | 7.88E-10 | 0.3097 | -7.67E-10 | 2.38E-09 |
| | CBDmale | VehFemale | 8.03E-10 | 0.6036 | -1.18E-09 | 2.02E-09 |
| | | CBDfemale | 7.72E-10 | 0.2862 | -7.11E-10 | 2.37E-09 |
| | | THCfemale | 7.46E-10 | 0.8073 | -1.31E-09 | 1.67E-09 |
| | | TpCfemale | 7.08E-10 | 0.8103 | -1.24E-09 | 1.58E-09 |
| | | VehMale | 7.72E-10 | 0.5626 | -1.99E-09 | 1.09E-09 |
| | | THCmale | 7.46E-10 | 0.2521 | -6.28E-10 | 2.35E-09 |
| | | TpCmale | 7.26E-10 | 0.6239 | -1.09E-09 | 1.81E-09 |
| | THCmale | VehFemale | 8.37E-10 | 0.5979 | -2.11E-09 | 1.23E-09 |
| | | CBDfemale | 8.07E-10 | 0.9677 | -1.64E-09 | 1.58E-09 |
| | | THCfemale | 7.83E-10 | 0.3884 | -2.24E-09 | 8.84E-10 |
| | | TpCfemale | 7.46E-10 | 0.3574 | -2.18E-09 | 7.99E-10 |
| | | VehMale | 8.07E-10 | 0.1089 | -2.92E-09 | 3.00E-10 |
| | | CBDmale | 7.46E-10 | 0.2521 | -2.35E-09 | 6.28E-10 |
| | | TpCmale | 7.63E-10 | 0.5103 | -2.03E-09 | 1.02E-09 |
| | TpCmale | VehFemale | 8.18E-10 | 0.9403 | -1.57E-09 | 1.70E-09 |
| | | CBDfemale | 7.88E-10 | 0.5509 | -1.10E-09 | 2.05E-09 |
| | | THCfemale | 7.63E-10 | 0.8196 | -1.70E-09 | 1.35E-09 |
| | | TpCfemale | 7.26E-10 | 0.7976 | -1.64E-09 | 1.26E-09 |
| | | VehMale | 7.88E-10 | 0.3097 | -2.38E-09 | 7.67E-10 |
| | | CBDmale | 7.26E-10 | 0.6239 | -1.81E-09 | 1.09E-09 |
| | | THCmale | 7.63E-10 | 0.5103 | -1.02E-09 | 2.03E-09 |
| | VehFemale | CBDfemale | 5.50E-09 | 0.9825 | -1.09E-08 | 1.11E-08 |
| | | THCfemale | 5.36E-09 | 0.3955 | -1.53E-08 | 6.12E-09 |
| | | TpCfemale | 5.14E-09 | 0.8559 | -1.12E-08 | 9.33E-09 |
| | | VehMale | 5.50E-09 | 0.6063 | -1.38E-08 | 8.14E-09 |
| | | CBDmale | 5.24E-09 | 0.1435 | -1.82E-08 | 2.71E-09 |
| | | THCmale | 5.36E-09 | 0.0267 | -2.28E-08 | -1.45E-09 |
| | | TpCmale | 5.24E-09 | 0.6235 | -1.30E-08 | 7.88E-09 |
| | CBDfemale | VehFemale | 5.50E-09 | 0.9825 | -1.11E-08 | 1.09E-08 |
| *N*-docosahexaenoyl ethanolamine | | THCfemale | 5.16E-09 | 0.3659 | -1.50E-08 | 5.61E-09 |
| | | TpCfemale | 4.94E-09 | 0.8310 | -1.09E-08 | 8.81E-09 |
| | | VehMale | 5.31E-09 | 0.5782 | -1.36E-08 | 7.65E-09 |
| | | CBDmale | 5.04E-09 | 0.1231 | -1.79E-08 | 2.19E-09 |
| | | THCmale | 5.16E-09 | 0.0205 | -2.26E-08 | -1.95E-09 |
| | | TpCmale | 5.04E-09 | 0.5935 | -1.28E-08 | 7.37E-09 |
| | THCfemale | VehFemale | 5.36E-09 | 0.3955 | -6.12E-09 | 1.53E-08 |
| | | CBDfemale | 5.16E-09 | 0.3659 | -5.61E-09 | 1.50E-08 |
| | | TpCfemale | 4.78E-09 | 0.4483 | -5.90E-09 | 1.32E-08 |

| | | | | | |
|---|---|---|---|---|---|
| | VehMale | 5.16E-09 | 0.7383 | -8.58E-09 | 1.20E-08 |
| | CBDmale | 4.88E-09 | 0.5180 | -1.29E-08 | 6.58E-09 |
| | THCmale | 5.01E-09 | 0.1360 | -1.76E-08 | 2.44E-09 |
| | TpCmale | 4.88E-09 | 0.6838 | -7.76E-09 | 1.18E-08 |
| TpCfemale | VehFemale | 5.14E-09 | 0.8559 | -9.33E-09 | 1.12E-08 |
| | CBDfemale | 4.94E-09 | 0.8310 | -8.81E-09 | 1.09E-08 |
| | THCfemale | 4.78E-09 | 0.4483 | -1.32E-08 | 5.90E-09 |
| | VehMale | 4.94E-09 | 0.7000 | -1.18E-08 | 7.95E-09 |
| | CBDmale | 4.64E-09 | 0.1469 | -1.61E-08 | 2.46E-09 |
| | THCmale | 4.78E-09 | 0.0221 | -2.08E-08 | -1.67E-09 |
| | TpCmale | 4.64E-09 | 0.7241 | -1.09E-08 | 7.63E-09 |
| VehMale | VehFemale | 5.50E-09 | 0.6063 | -8.14E-09 | 1.38E-08 |
| | CBDfemale | 5.31E-09 | 0.5782 | -7.65E-09 | 1.36E-08 |
| | THCfemale | 5.16E-09 | 0.7383 | -1.20E-08 | 8.58E-09 |
| | TpCfemale | 4.94E-09 | 0.7000 | -7.95E-09 | 1.18E-08 |
| | CBDmale | 5.04E-09 | 0.3340 | -1.50E-08 | 5.16E-09 |
| | THCmale | 5.16E-09 | 0.0765 | -1.96E-08 | 1.02E-09 |
| | TpCmale | 5.04E-09 | 0.9582 | -9.81E-09 | 1.03E-08 |
| CBDmale | VehFemale | 5.24E-09 | 0.1435 | -2.71E-09 | 1.82E-08 |
| | CBDfemale | 5.04E-09 | 0.1231 | -2.19E-09 | 1.79E-08 |
| | THCfemale | 4.88E-09 | 0.5180 | -6.58E-09 | 1.29E-08 |
| | TpCfemale | 4.64E-09 | 0.1469 | -2.46E-09 | 1.61E-08 |
| | VehMale | 5.04E-09 | 0.3340 | -5.16E-09 | 1.50E-08 |
| | THCmale | 4.88E-09 | 0.3721 | -1.41E-08 | 5.37E-09 |
| | TpCmale | 4.75E-09 | 0.2805 | -4.32E-09 | 1.47E-08 |
| THCmale | VehFemale | 5.36E-09 | 0.0267 | 1.45E-09 | 2.28E-08 |
| | CBDfemale | 5.16E-09 | 0.0205 | 1.95E-09 | 2.26E-08 |
| | THCfemale | 5.01E-09 | 0.1360 | -2.44E-09 | 1.76E-08 |
| | TpCfemale | 4.78E-09 | 0.0221 | 1.67E-09 | 2.08E-08 |
| | VehMale | 5.16E-09 | 0.0765 | -1.02E-09 | 1.96E-08 |
| | CBDmale | 4.88E-09 | 0.3721 | -5.37E-09 | 1.41E-08 |
| | TpCmale | 4.88E-09 | 0.0546 | -1.93E-10 | 1.93E-08 |
| TpCmale | VehFemale | 5.24E-09 | 0.6235 | -7.88E-09 | 1.30E-08 |
| | CBDfemale | 5.04E-09 | 0.5935 | -7.37E-09 | 1.28E-08 |
| | THCfemale | 4.88E-09 | 0.6838 | -1.18E-08 | 7.76E-09 |
| | TpCfemale | 4.64E-09 | 0.7241 | -7.63E-09 | 1.09E-08 |
| | VehMale | 5.04E-09 | 0.9582 | -1.03E-08 | 9.81E-09 |
| | CBDmale | 4.75E-09 | 0.2805 | -1.47E-08 | 4.32E-09 |
| | THCmale | 4.88E-09 | 0.0546 | -1.93E-08 | 1.93E-10 |

#### Supplemental Table 3.8 Fisher's LSD post-hoc test for liver *N*-acyl ethanolamine ANOVA

| | | | | | 95% Confidence Interval | |
|---|---|---|---|---|---|---|
| | | | Std. Error | Sig. | Lower Bound | Upper Bound |
| *N*-palmitoyl ethanolamine | VehFemale | CBDfemale | 2.85565E-11 | 0.502 | -7.6335E-11 | 3.7796E-11 |
| | | THCfemale | 2.71912E-11 | 0.001 | -1.4743E-10 | -3.8759E-11 |
| | | TpCfemale | 2.71912E-11 | 0.002 | -1.4061E-10 | -3.1940E-11 |
| | | VehMale | 2.94930E-11 | 0.180 | -9.8939E-11 | 1.8935E-11 |
| | | CBDmale | 2.78063E-11 | 0.836 | -6.1363E-11 | 4.9770E-11 |
| | | THCmale | 2.78063E-11 | 0.356 | -8.1407E-11 | 2.9726E-11 |
| | | TpCmale | 2.66774E-11 | 0.332 | -7.9383E-11 | 2.7238E-11 |
| | CBDfemale | VehFemale | 2.85565E-11 | 0.502 | -3.7796E-11 | 7.6335E-11 |
| | | THCfemale | 2.61724E-11 | 0.006 | -1.2613E-10 | -2.1525E-11 |
| | | TpCfemale | 2.61724E-11 | 0.013 | -1.1931E-10 | -1.4706E-11 |
| | | VehMale | 2.85565E-11 | 0.471 | -7.7798E-11 | 3.6334E-11 |
| | | CBDmale | 2.68109E-11 | 0.617 | -4.0104E-11 | 6.7051E-11 |
| | | THCmale | 2.68109E-11 | 0.807 | -6.0148E-11 | 4.7007E-11 |
| | | TpCmale | 2.56382E-11 | 0.792 | -5.8037E-11 | 4.4431E-11 |
| | THCfemale | VehFemale | 2.71912E-11 | 0.001 | 3.8759E-11 | 1.4743E-10 |
| | | CBDfemale | 2.61724E-11 | 0.006 | 2.1525E-11 | 1.2613E-10 |
| | | TpCfemale | 2.46756E-11 | 0.783 | -4.2491E-11 | 5.6129E-11 |
| | | VehMale | 2.71912E-11 | 0.055 | -1.2429E-12 | 1.0743E-10 |
| | | CBDmale | 2.53518E-11 | 0.001 | 3.6638E-11 | 1.3796E-10 |
| | | THCmale | 2.53518E-11 | 0.010 | 1.6594E-11 | 1.1792E-10 |
| | | TpCmale | 2.41083E-11 | 0.007 | 1.8847E-11 | 1.1520E-10 |
| | TpCfemale | VehFemale | 2.71912E-11 | 0.002 | 3.1940E-11 | 1.4061E-10 |
| | | CBDfemale | 2.61724E-11 | 0.013 | 1.4706E-11 | 1.1931E-10 |
| | | THCfemale | 2.46756E-11 | 0.783 | -5.6129E-11 | 4.2491E-11 |
| | | VehMale | 2.71912E-11 | 0.094 | -8.0618E-12 | 1.0061E-10 |
| | | CBDmale | 2.53518E-11 | 0.002 | 2.9819E-11 | 1.3114E-10 |
| | | THCmale | 2.53518E-11 | 0.020 | 9.7753E-12 | 1.1110E-10 |
| | | TpCmale | 2.41083E-11 | 0.015 | 1.2028E-11 | 1.0838E-10 |
| | VehMale | VehFemale | 2.94930E-11 | 0.180 | -1.8935E-11 | 9.8939E-11 |
| | | CBDfemale | 2.85565E-11 | 0.471 | -3.6334E-11 | 7.7798E-11 |
| | | THCfemale | 2.71912E-11 | 0.055 | -1.0743E-10 | 1.2429E-12 |
| | | TpCfemale | 2.71912E-11 | 0.094 | -1.0061E-10 | 8.0618E-12 |
| | | CBDmale | 2.78063E-11 | 0.223 | -2.1361E-11 | 8.9772E-11 |
| | | THCmale | 2.78063E-11 | 0.612 | -4.1405E-11 | 6.9728E-11 |
| | | TpCmale | 2.66774E-11 | 0.603 | -3.9382E-11 | 6.7240E-11 |
| | CBDmale | VehFemale | 2.78063E-11 | 0.836 | -4.9770E-11 | 6.1363E-11 |
| | | CBDfemale | 2.68109E-11 | 0.617 | -6.7051E-11 | 4.0104E-11 |
| | | THCfemale | 2.53518E-11 | 0.001 | -1.3796E-10 | -3.6638E-11 |
| | | TpCfemale | 2.53518E-11 | 0.002 | -1.3114E-10 | -2.9819E-11 |
| | | VehMale | 2.78063E-11 | 0.223 | -8.9772E-11 | 2.1361E-11 |
| | | THCmale | 2.60104E-11 | 0.444 | -7.2022E-11 | 3.1933E-11 |
| | | TpCmale | 2.47999E-11 | 0.417 | -6.9835E-11 | 2.9282E-11 |
| | THCmale | VehFemale | 2.78063E-11 | 0.356 | -2.9726E-11 | 8.1407E-11 |
| | | CBDfemale | 2.68109E-11 | 0.807 | -4.7007E-11 | 6.0148E-11 |
| | | THCfemale | 2.53518E-11 | 0.010 | -1.1792E-10 | -1.6594E-11 |
| | | TpCfemale | 2.53518E-11 | 0.020 | -1.1110E-10 | -9.7753E-12 |
| | | VehMale | 2.78063E-11 | 0.612 | -6.9728E-11 | 4.1405E-11 |
| | | CBDmale | 2.60104E-11 | 0.444 | -3.1933E-11 | 7.2022E-11 |
| | | TpCmale | 2.47999E-11 | 0.993 | -4.9791E-11 | 4.9326E-11 |
| | TpCmale | VehFemale | 2.66774E-11 | 0.332 | -2.7238E-11 | 7.9383E-11 |
| | | CBDfemale | 2.56382E-11 | 0.792 | -4.4431E-11 | 5.8037E-11 |
| | | THCfemale | 2.41083E-11 | 0.007 | -1.1520E-10 | -1.8847E-11 |
| | | TpCfemale | 2.41083E-11 | 0.015 | -1.0838E-10 | -1.2028E-11 |
| | | VehMale | 2.66774E-11 | 0.603 | -6.7240E-11 | 3.9382E-11 |
| | | CBDmale | 2.47999E-11 | 0.417 | -2.9282E-11 | 6.9835E-11 |

| | | | | | | |
|---|---|---|---|---|---|---|
| | | THCmale | 2.47999E-11 | 0.993 | -4.9326E-11 | 4.9791E-11 |
| | VehFemale | CBDfemale | 2.09597E-11 | 0.626 | -5.2191E-11 | 3.1632E-11 |
| | | THCfemale | 2.04091E-11 | 0.001 | -1.1348E-10 | -3.1856E-11 |
| | | TpCfemale | 1.99577E-11 | 0.000 | -1.3047E-10 | -5.0654E-11 |
| | | VehMale | 2.16471E-11 | 0.033 | -9.0551E-11 | -3.9785E-12 |
| | | CBDmale | 1.99577E-11 | 0.641 | -4.9272E-11 | 3.0544E-11 |
| | | THCmale | 2.09597E-11 | 0.266 | -6.5460E-11 | 1.8363E-11 |
| | | TpCmale | 1.99577E-11 | 0.485 | -5.3938E-11 | 2.5878E-11 |
| | CBDfemale | VehFemale | 2.09597E-11 | 0.626 | -3.1632E-11 | 5.2191E-11 |
| | | THCfemale | 1.96785E-11 | 0.002 | -1.0174E-10 | -2.3038E-11 |
| | | TpCfemale | 1.92099E-11 | 0.000 | -1.1870E-10 | -4.1870E-11 |
| | | VehMale | 2.09597E-11 | 0.083 | -7.8897E-11 | 4.9264E-12 |
| | | CBDmale | 1.92099E-11 | 0.962 | -3.7497E-11 | 3.9328E-11 |
| | | THCmale | 2.02490E-11 | 0.515 | -5.3759E-11 | 2.7222E-11 |
| | | TpCmale | 1.92099E-11 | 0.846 | -4.2163E-11 | 3.4662E-11 |
| | THCfemale | VehFemale | 2.04091E-11 | 0.001 | 3.1856E-11 | 1.1348E-10 |
| | | CBDfemale | 1.96785E-11 | 0.002 | 2.3038E-11 | 1.0174E-10 |
| | | TpCfemale | 1.86076E-11 | 0.340 | -5.5103E-11 | 1.9313E-11 |
| | | VehMale | 2.04091E-11 | 0.218 | -1.5409E-11 | 6.6213E-11 |
| | | CBDmale | 1.86076E-11 | 0.001 | 2.6095E-11 | 1.0051E-10 |
| | | THCmale | 1.96785E-11 | 0.015 | 9.7687E-12 | 8.8468E-11 |
| | | TpCmale | 1.86076E-11 | 0.003 | 2.1428E-11 | 9.5845E-11 |
| | TpCfemale | VehFemale | 1.99577E-11 | 0.000 | 5.0654E-11 | 1.3047E-10 |
| | | CBDfemale | 1.92099E-11 | 0.000 | 4.1870E-11 | 1.1870E-10 |
| | | THCfemale | 1.86076E-11 | 0.340 | -1.9313E-11 | 5.5103E-11 |
| | | VehMale | 1.99577E-11 | 0.034 | 3.3896E-12 | 8.3205E-11 |
| | | CBDmale | 1.81113E-11 | 0.000 | 4.4982E-11 | 1.1741E-10 |
| | | THCmale | 1.92099E-11 | 0.001 | 2.8601E-11 | 1.0543E-10 |
| | | TpCmale | 1.81113E-11 | 0.000 | 4.0316E-11 | 1.1275E-10 |
| *N*-stearoyl ethanolamine | VehMale | VehFemale | 2.16471E-11 | 0.033 | 3.9785E-12 | 9.0551E-11 |
| | | CBDfemale | 2.09597E-11 | 0.083 | -4.9264E-12 | 7.8897E-11 |
| | | THCfemale | 2.04091E-11 | 0.218 | -6.6213E-11 | 1.5409E-11 |
| | | TpCfemale | 1.99577E-11 | 0.034 | -8.3205E-11 | -3.3896E-12 |
| | | CBDmale | 1.99577E-11 | 0.062 | -2.0071E-12 | 7.7808E-11 |
| | | THCmale | 2.09597E-11 | 0.262 | -1.8195E-11 | 6.5628E-11 |
| | | TpCmale | 1.99577E-11 | 0.101 | -6.6734E-12 | 7.3142E-11 |
| | CBDmale | VehFemale | 1.99577E-11 | 0.641 | -3.0544E-11 | 4.9272E-11 |
| | | CBDfemale | 1.92099E-11 | 0.962 | -3.9328E-11 | 3.7497E-11 |
| | | THCfemale | 1.86076E-11 | 0.001 | -1.0051E-10 | -2.6095E-11 |
| | | TpCfemale | 1.81113E-11 | 0.000 | -1.1741E-10 | -4.4982E-11 |
| | | VehMale | 1.99577E-11 | 0.062 | -7.7808E-11 | 2.0071E-12 |
| | | THCmale | 1.92099E-11 | 0.463 | -5.2597E-11 | 2.4228E-11 |
| | | TpCmale | 1.81113E-11 | 0.798 | -4.0882E-11 | 3.1549E-11 |
| | THCmale | VehFemale | 2.09597E-11 | 0.266 | -1.8363E-11 | 6.5460E-11 |
| | | CBDfemale | 2.02490E-11 | 0.515 | -2.7222E-11 | 5.3759E-11 |
| | | THCfemale | 1.96785E-11 | 0.015 | -8.8468E-11 | -9.7687E-12 |
| | | TpCfemale | 1.92099E-11 | 0.001 | -1.0543E-10 | -2.8601E-11 |
| | | VehMale | 2.09597E-11 | 0.262 | -6.5628E-11 | 1.8195E-11 |
| | | CBDmale | 1.92099E-11 | 0.463 | -2.4228E-11 | 5.2597E-11 |
| | | TpCmale | 1.92099E-11 | 0.622 | -2.8895E-11 | 4.7931E-11 |
| | TpCmale | VehFemale | 1.99577E-11 | 0.485 | -2.5878E-11 | 5.3938E-11 |
| | | CBDfemale | 1.92099E-11 | 0.846 | -3.4662E-11 | 4.2163E-11 |
| | | THCfemale | 1.86076E-11 | 0.003 | -9.5845E-11 | -2.1428E-11 |
| | | TpCfemale | 1.81113E-11 | 0.000 | -1.1275E-10 | -4.0316E-11 |
| | | VehMale | 1.99577E-11 | 0.101 | -7.3142E-11 | 6.6734E-12 |
| | | CBDmale | 1.81113E-11 | 0.798 | -3.1549E-11 | 4.0882E-11 |
| | | THCmale | 1.92099E-11 | 0.622 | -4.7931E-11 | 2.8895E-11 |
| *N*-oleoyl ethanolamine | VehFemale | CBDfemale | 2.55875E-11 | 0.892 | -5.4636E-11 | 4.7630E-11 |
| | | THCfemale | 2.43642E-11 | 0.000 | -1.4807E-10 | -5.0691E-11 |

| | | | | | | |
|---|---|---|---|---|---|---|
| | | TpCfemale | 2.49153E-11 | 0.000 | -1.7356E-10 | -7.3981E-11 |
| | | VehMale | 2.64267E-11 | 0.071 | -1.0130E-10 | 4.3239E-12 |
| | | CBDmale | 2.43642E-11 | 0.141 | -8.4967E-11 | 1.2409E-11 |
| | | THCmale | 2.49153E-11 | 0.003 | -1.2676E-10 | -2.7177E-11 |
| | | TpCmale | 2.39038E-11 | 0.002 | -1.2466E-10 | -2.9124E-11 |
| | CBDfemale | VehFemale | 2.55875E-11 | 0.892 | -4.7630E-11 | 5.4636E-11 |
| | | THCfemale | 2.34514E-11 | 0.000 | -1.4274E-10 | -4.9012E-11 |
| | | TpCfemale | 2.40234E-11 | 0.000 | -1.6827E-10 | -7.2260E-11 |
| | | VehMale | 2.55875E-11 | 0.084 | -9.6115E-11 | 6.1499E-12 |
| | | CBDmale | 2.34514E-11 | 0.167 | -7.9640E-11 | 1.4088E-11 |
| | | THCmale | 2.40234E-11 | 0.003 | -1.2147E-10 | -2.5456E-11 |
| | | TpCmale | 2.29727E-11 | 0.002 | -1.1930E-10 | -2.7482E-11 |
| | THCfemale | VehFemale | 2.43642E-11 | 0.000 | 5.0691E-11 | 1.4807E-10 |
| | | CBDfemale | 2.34514E-11 | 0.000 | 4.9012E-11 | 1.4274E-10 |
| | | TpCfemale | 2.27160E-11 | 0.287 | -6.9786E-11 | 2.1003E-11 |
| | | VehMale | 2.43642E-11 | 0.041 | 2.2049E-12 | 9.9581E-11 |
| | | CBDmale | 2.21102E-11 | 0.006 | 1.8916E-11 | 1.0728E-10 |
| | | THCmale | 2.27160E-11 | 0.328 | -2.2982E-11 | 6.7807E-11 |
| | | TpCmale | 2.16018E-11 | 0.302 | -2.0681E-11 | 6.5654E-11 |
| | TpCfemale | VehFemale | 2.49153E-11 | 0.000 | 7.3981E-11 | 1.7356E-10 |
| | | CBDfemale | 2.40234E-11 | 0.000 | 7.2260E-11 | 1.6827E-10 |
| | | THCfemale | 2.27160E-11 | 0.287 | -2.1003E-11 | 6.9786E-11 |
| | | VehMale | 2.49153E-11 | 0.004 | 2.5495E-11 | 1.2507E-10 |
| | | CBDmale | 2.27160E-11 | 0.000 | 4.2097E-11 | 1.3289E-10 |
| | | THCmale | 2.33062E-11 | 0.049 | 2.3002E-13 | 9.3377E-11 |
| | | TpCmale | 2.22215E-11 | 0.039 | 2.4715E-12 | 9.1284E-11 |
| | VehMale | VehFemale | 2.64267E-11 | 0.071 | -4.3239E-12 | 1.0130E-10 |
| | | CBDfemale | 2.55875E-11 | 0.084 | -6.1499E-12 | 9.6115E-11 |
| | | THCfemale | 2.43642E-11 | 0.041 | -9.9581E-11 | -2.2049E-12 |
| | | TpCfemale | 2.49153E-11 | 0.004 | -1.2507E-10 | -2.5495E-11 |
| | | CBDmale | 2.43642E-11 | 0.618 | -3.6481E-11 | 6.0895E-11 |
| | | THCmale | 2.49153E-11 | 0.257 | -7.8270E-11 | 2.1309E-11 |
| | | TpCmale | 2.39038E-11 | 0.239 | -7.6175E-11 | 1.9361E-11 |
| | CBDmale | VehFemale | 2.43642E-11 | 0.141 | -1.2409E-11 | 8.4967E-11 |
| | | CBDfemale | 2.34514E-11 | 0.167 | -1.4088E-11 | 7.9640E-11 |
| | | THCfemale | 2.21102E-11 | 0.006 | -1.0728E-10 | -1.8916E-11 |
| | | TpCfemale | 2.27160E-11 | 0.000 | -1.3289E-10 | -4.2097E-11 |
| | | VehMale | 2.43642E-11 | 0.618 | -6.0895E-11 | 3.6481E-11 |
| | | THCmale | 2.27160E-11 | 0.078 | -8.6082E-11 | 4.7069E-12 |
| | | TpCmale | 2.16018E-11 | 0.065 | -8.3781E-11 | 2.5545E-12 |
| | THCmale | VehFemale | 2.49153E-11 | 0.003 | 2.7177E-11 | 1.2676E-10 |
| | | CBDfemale | 2.40234E-11 | 0.003 | 2.5456E-11 | 1.2147E-10 |
| | | THCfemale | 2.27160E-11 | 0.328 | -6.7807E-11 | 2.2982E-11 |
| | | TpCfemale | 2.33062E-11 | 0.049 | -9.3377E-11 | -2.3002E-13 |
| | | VehMale | 2.49153E-11 | 0.257 | -2.1309E-11 | 7.8270E-11 |
| | | CBDmale | 2.27160E-11 | 0.078 | -4.7069E-12 | 8.6082E-11 |
| | | TpCmale | 2.22215E-11 | 0.997 | -4.4332E-11 | 4.4480E-11 |
| | TpCmale | VehFemale | 2.39038E-11 | 0.002 | 2.9124E-11 | 1.2466E-10 |
| | | CBDfemale | 2.29727E-11 | 0.002 | 2.7482E-11 | 1.1930E-10 |
| | | THCfemale | 2.16018E-11 | 0.302 | -6.5654E-11 | 2.0681E-11 |
| | | TpCfemale | 2.22215E-11 | 0.039 | -9.1284E-11 | -2.4715E-12 |
| | | VehMale | 2.39038E-11 | 0.239 | -1.9361E-11 | 7.6175E-11 |
| | | CBDmale | 2.16018E-11 | 0.065 | -2.5545E-12 | 8.3781E-11 |
| | | THCmale | 2.22215E-11 | 0.997 | -4.4480E-11 | 4.4332E-11 |
| | VehFemale | CBDfemale | 2.23384E-11 | 0.874 | -4.1117E-11 | 4.8219E-11 |
| | | THCfemale | 2.04735E-11 | 0.073 | -7.8234E-11 | 3.6446E-12 |
| *N*-linoleoyl ethanolamine | | TpCfemale | 2.09729E-11 | 0.000 | -1.5613E-10 | -7.2256E-11 |
| | | VehMale | 2.23384E-11 | 0.808 | -5.0114E-11 | 3.9223E-11 |
| | | CBDmale | 2.04735E-11 | 0.614 | -5.1317E-11 | 3.0561E-11 |

| | | | | | | |
|---|---|---|---|---|---|---|
| | | THCmale | 2.15809E-11 | 0.005 | -1.0652E-10 | -2.0212E-11 |
| | | TpCmale | 2.04735E-11 | 0.003 | -1.0381E-10 | -2.1930E-11 |
| | CBDfemale | VehFemale | 2.23384E-11 | 0.874 | -4.8219E-11 | 4.1117E-11 |
| | | THCfemale | 2.12704E-11 | 0.059 | -8.3378E-11 | 1.6871E-12 |
| | | TpCfemale | 2.17515E-11 | 0.000 | -1.6124E-10 | -7.4250E-11 |
| | | VehMale | 2.30710E-11 | 0.698 | -5.5130E-11 | 3.7137E-11 |
| | | CBDmale | 2.12704E-11 | 0.515 | -5.6462E-11 | 2.8604E-11 |
| | | THCmale | 2.23384E-11 | 0.004 | -1.1159E-10 | -2.2248E-11 |
| | | TpCmale | 2.12704E-11 | 0.003 | -1.0895E-10 | -2.3888E-11 |
| | THCfemale | VehFemale | 2.04735E-11 | 0.073 | -3.6446E-12 | 7.8234E-11 |
| | | CBDfemale | 2.12704E-11 | 0.059 | -1.6871E-12 | 8.3378E-11 |
| | | TpCfemale | 1.98315E-11 | 0.000 | -1.1655E-10 | -3.7243E-11 |
| | | VehMale | 2.12704E-11 | 0.139 | -1.0683E-11 | 7.4382E-11 |
| | | CBDmale | 1.93026E-11 | 0.168 | -1.1681E-11 | 6.5514E-11 |
| | | THCmale | 2.04735E-11 | 0.208 | -6.7010E-11 | 1.4868E-11 |
| | | TpCmale | 1.93026E-11 | 0.190 | -6.4173E-11 | 1.3023E-11 |
| | TpCfemale | VehFemale | 2.09729E-11 | 0.000 | 7.2256E-11 | 1.5613E-10 |
| | | CBDfemale | 2.17515E-11 | 0.000 | 7.4250E-11 | 1.6124E-10 |
| | | THCfemale | 1.98315E-11 | 0.000 | 3.7243E-11 | 1.1655E-10 |
| | | VehMale | 2.17515E-11 | 0.000 | 6.5253E-11 | 1.5224E-10 |
| | | CBDmale | 1.98315E-11 | 0.000 | 6.4160E-11 | 1.4347E-10 |
| | | THCmale | 2.09729E-11 | 0.018 | 8.8900E-12 | 9.2766E-11 |
| | | TpCmale | 1.98315E-11 | 0.012 | 1.1668E-11 | 9.0979E-11 |
| | VehMale | VehFemale | 2.23384E-11 | 0.808 | -3.9223E-11 | 5.0114E-11 |
| | | CBDfemale | 2.30710E-11 | 0.698 | -3.7137E-11 | 5.5130E-11 |
| | | THCfemale | 2.12704E-11 | 0.139 | -7.4382E-11 | 1.0683E-11 |
| | | TpCfemale | 2.17515E-11 | 0.000 | -1.5224E-10 | -6.5253E-11 |
| | | CBDmale | 2.12704E-11 | 0.817 | -4.7466E-11 | 3.7600E-11 |
| | | THCmale | 2.23384E-11 | 0.012 | -1.0259E-10 | -1.3252E-11 |
| | | TpCmale | 2.12704E-11 | 0.009 | -9.9957E-11 | -1.4892E-11 |
| | CBDmale | VehFemale | 2.04735E-11 | 0.614 | -3.0561E-11 | 5.1317E-11 |
| | | CBDfemale | 2.12704E-11 | 0.515 | -2.8604E-11 | 5.6462E-11 |
| | | THCfemale | 1.93026E-11 | 0.168 | -6.5514E-11 | 1.1681E-11 |
| | | TpCfemale | 1.98315E-11 | 0.000 | -1.4347E-10 | -6.4160E-11 |
| | | VehMale | 2.12704E-11 | 0.817 | -3.7600E-11 | 4.7466E-11 |
| | | THCmale | 2.04735E-11 | 0.012 | -9.3927E-11 | -1.2048E-11 |
| | | TpCmale | 1.93026E-11 | 0.009 | -9.1089E-11 | -1.3894E-11 |
| | THCmale | VehFemale | 2.15809E-11 | 0.005 | 2.0212E-11 | 1.0652E-10 |
| | | CBDfemale | 2.23384E-11 | 0.004 | 2.2248E-11 | 1.1159E-10 |
| | | THCfemale | 2.04735E-11 | 0.208 | -1.4868E-11 | 6.7010E-11 |
| | | TpCfemale | 2.09729E-11 | 0.018 | -9.2766E-11 | -8.8900E-12 |
| | | VehMale | 2.23384E-11 | 0.012 | 1.3252E-11 | 1.0259E-10 |
| | | CBDmale | 2.04735E-11 | 0.012 | 1.2048E-11 | 9.3927E-11 |
| | | TpCmale | 2.04735E-11 | 0.981 | -4.0443E-11 | 4.1435E-11 |
| | TpCmale | VehFemale | 2.04735E-11 | 0.003 | 2.1930E-11 | 1.0381E-10 |
| | | CBDfemale | 2.12704E-11 | 0.003 | 2.3888E-11 | 1.0895E-10 |
| | | THCfemale | 1.93026E-11 | 0.190 | -1.3023E-11 | 6.4173E-11 |
| | | TpCfemale | 1.98315E-11 | 0.012 | -9.0979E-11 | -1.1668E-11 |
| | | VehMale | 2.12704E-11 | 0.009 | 1.4892E-11 | 9.9957E-11 |
| | | CBDmale | 1.93026E-11 | 0.009 | 1.3894E-11 | 9.1089E-11 |
| | | THCmale | 2.04735E-11 | 0.981 | -4.1435E-11 | 4.0443E-11 |
| *N*-arachidonoyl ethanolamine | VehFemale | CBDfemale | 2.68090E-12 | 0.943 | -5.5492E-12 | 5.1655E-12 |
| | | THCfemale | 2.45708E-12 | 0.027 | -1.0488E-11 | -6.6765E-13 |
| | | TpCfemale | 2.51702E-12 | 0.000 | -1.7170E-11 | -7.1101E-12 |
| | | VehMale | 2.68090E-12 | 0.908 | -5.0467E-12 | 5.6679E-12 |
| | | CBDmale | 2.45708E-12 | 0.829 | -5.4431E-12 | 4.3771E-12 |
| | | THCmale | 2.51702E-12 | 0.000 | -1.4627E-11 | -4.5674E-12 |
| | | TpCmale | 2.40693E-12 | 0.001 | -1.3081E-11 | -3.4614E-12 |
| | CBDfemale | VehFemale | 2.68090E-12 | 0.943 | -5.1655E-12 | 5.5492E-12 |

| | | | | | |
|---|---|---|---|---|---|
| | THCfemale | 2.55273E-12 | 0.039 | -1.0487E-11 | -2.8467E-13 |
| | TpCfemale | 2.61047E-12 | 0.000 | -1.7165E-11 | -6.7315E-12 |
| | VehMale | 2.76882E-12 | 0.857 | -5.0306E-12 | 6.0355E-12 |
| | CBDmale | 2.55273E-12 | 0.894 | -5.4424E-12 | 4.7600E-12 |
| | THCmale | 2.61047E-12 | 0.001 | -1.4622E-11 | -4.1888E-12 |
| | TpCmale | 2.50449E-12 | 0.002 | -1.3084E-11 | -3.0746E-12 |
| THCfemale | VehFemale | 2.45708E-12 | 0.027 | 6.6765E-13 | 1.0488E-11 |
| | CBDfemale | 2.55273E-12 | 0.039 | 2.8467E-13 | 1.0487E-11 |
| | TpCfemale | 2.38004E-12 | 0.008 | -1.1318E-11 | -1.8061E-12 |
| | VehMale | 2.55273E-12 | 0.024 | 7.8712E-13 | 1.0990E-11 |
| | CBDmale | 2.31656E-12 | 0.033 | 4.1544E-13 | 9.6740E-12 |
| | THCmale | 2.38004E-12 | 0.096 | -8.7756E-12 | 7.3663E-13 |
| | TpCmale | 2.26330E-12 | 0.238 | -7.2164E-12 | 1.8293E-12 |
| TpCfemale | VehFemale | 2.51702E-12 | 0.000 | 7.1101E-12 | 1.7170E-11 |
| | CBDfemale | 2.61047E-12 | 0.000 | 6.7315E-12 | 1.7165E-11 |
| | THCfemale | 2.38004E-12 | 0.008 | 1.8061E-12 | 1.1318E-11 |
| | VehMale | 2.61047E-12 | 0.000 | 7.2339E-12 | 1.7667E-11 |
| | CBDmale | 2.38004E-12 | 0.000 | 6.8508E-12 | 1.6363E-11 |
| | THCmale | 2.44187E-12 | 0.302 | -2.3370E-12 | 7.4224E-12 |
| | TpCmale | 2.32823E-12 | 0.102 | -7.8398E-13 | 8.5212E-12 |
| VehMale | VehFemale | 2.68090E-12 | 0.908 | -5.6679E-12 | 5.0467E-12 |
| | CBDfemale | 2.76882E-12 | 0.857 | -6.0355E-12 | 5.0306E-12 |
| | THCfemale | 2.55273E-12 | 0.024 | -1.0990E-11 | -7.8712E-13 |
| | TpCfemale | 2.61047E-12 | 0.000 | -1.7667E-11 | -7.2339E-12 |
| | CBDmale | 2.55273E-12 | 0.742 | -5.9448E-12 | 4.2576E-12 |
| | THCmale | 2.61047E-12 | 0.000 | -1.5124E-11 | -4.6912E-12 |
| | TpCmale | 2.50449E-12 | 0.001 | -1.3587E-11 | -3.5771E-12 |
| CBDmale | VehFemale | 2.45708E-12 | 0.829 | -4.3771E-12 | 5.4431E-12 |
| | CBDfemale | 2.55273E-12 | 0.894 | -4.7600E-12 | 5.4424E-12 |
| | THCfemale | 2.31656E-12 | 0.033 | -9.6740E-12 | -4.1544E-13 |
| | TpCfemale | 2.38004E-12 | 0.000 | -1.6363E-11 | -6.8508E-12 |
| | VehMale | 2.55273E-12 | 0.742 | -4.2576E-12 | 5.9448E-12 |
| | THCmale | 2.38004E-12 | 0.000 | -1.3820E-11 | -4.3081E-12 |
| | TpCmale | 2.26330E-12 | 0.001 | -1.2261E-11 | -3.2154E-12 |
| THCmale | VehFemale | 2.51702E-12 | 0.000 | 4.5674E-12 | 1.4627E-11 |
| | CBDfemale | 2.61047E-12 | 0.001 | 4.1888E-12 | 1.4622E-11 |
| | THCfemale | 2.38004E-12 | 0.096 | -7.3663E-13 | 8.7756E-12 |
| | TpCfemale | 2.44187E-12 | 0.302 | -7.4224E-12 | 2.3370E-12 |
| | VehMale | 2.61047E-12 | 0.000 | 4.6912E-12 | 1.5124E-11 |
| | CBDmale | 2.38004E-12 | 0.000 | 4.3081E-12 | 1.3820E-11 |
| | TpCmale | 2.32823E-12 | 0.571 | -3.3267E-12 | 5.9785E-12 |
| TpCmale | VehFemale | 2.40693E-12 | 0.001 | 3.4614E-12 | 1.3081E-11 |
| | CBDfemale | 2.50449E-12 | 0.002 | 3.0746E-12 | 1.3084E-11 |
| | THCfemale | 2.26330E-12 | 0.238 | -1.8293E-12 | 7.2164E-12 |
| | TpCfemale | 2.32823E-12 | 0.102 | -8.5212E-12 | 7.8398E-13 |
| | VehMale | 2.50449E-12 | 0.001 | 3.5771E-12 | 1.3587E-11 |
| | CBDmale | 2.26330E-12 | 0.001 | 3.2154E-12 | 1.2261E-11 |
| | THCmale | 2.32823E-12 | 0.571 | -5.9785E-12 | 3.3267E-12 |
| *N*-docosahexaenoyl ethanolamine VehFemale | CBDfemale | 1.66192E-11 | 0.004 | -8.3002E-11 | -1.6580E-11 |
| | THCfemale | 1.61510E-11 | 0.062 | -1.6051E-12 | 6.2945E-11 |
| | TpCfemale | 1.61510E-11 | 0.082 | -3.6961E-12 | 6.0854E-11 |
| | VehMale | 1.72025E-11 | 0.018 | 7.4191E-12 | 7.6172E-11 |
| | CBDmale | 1.57664E-11 | 0.016 | 7.4213E-12 | 7.0434E-11 |
| | THCmale | 1.61510E-11 | 0.086 | -4.1309E-12 | 6.0419E-11 |
| | TpCmale | 1.54446E-11 | 0.002 | -8.1065E-11 | -1.9338E-11 |
| CBDfemale | VehFemale | 1.66192E-11 | 0.004 | 1.6580E-11 | 8.3002E-11 |
| | THCfemale | 1.61510E-11 | 0.000 | 4.8186E-11 | 1.1274E-10 |
| | TpCfemale | 1.61510E-11 | 0.000 | 4.6095E-11 | 1.1065E-10 |
| | VehMale | 1.72025E-11 | 0.000 | 5.7210E-11 | 1.2596E-10 |
| | CBDmale | 1.57664E-11 | 0.000 | 5.7212E-11 | 1.2023E-10 |

| | | | | | |
|---|---|---|---|---|---|
| | THCmale | 1.61510E-11 | 0.000 | 4.5660E-11 | 1.1021E-10 |
| | TpCmale | 1.54446E-11 | 0.979 | -3.1274E-11 | 3.0453E-11 |
| THCfemale | VehFemale | 1.61510E-11 | 0.062 | -6.2945E-11 | 1.6051E-12 |
| | CBDfemale | 1.61510E-11 | 0.000 | -1.1274E-10 | -4.8186E-11 |
| | TpCfemale | 1.56687E-11 | 0.894 | -3.3402E-11 | 2.9221E-11 |
| | VehMale | 1.67506E-11 | 0.509 | -2.2348E-11 | 4.4599E-11 |
| | CBDmale | 1.52720E-11 | 0.591 | -2.2261E-11 | 3.8777E-11 |
| | THCmale | 1.56687E-11 | 0.872 | -3.3837E-11 | 2.8786E-11 |
| | TpCmale | 1.49396E-11 | 0.000 | -1.1073E-10 | -5.1017E-11 |
| TpCfemale | VehFemale | 1.61510E-11 | 0.082 | -6.0854E-11 | 3.6961E-12 |
| | CBDfemale | 1.61510E-11 | 0.000 | -1.1065E-10 | -4.6095E-11 |
| | THCfemale | 1.56687E-11 | 0.894 | -2.9221E-11 | 3.3402E-11 |
| | VehMale | 1.67506E-11 | 0.433 | -2.0257E-11 | 4.6690E-11 |
| | CBDmale | 1.52720E-11 | 0.500 | -2.0170E-11 | 4.0867E-11 |
| | THCmale | 1.56687E-11 | 0.978 | -3.1746E-11 | 3.0877E-11 |
| | TpCmale | 1.49396E-11 | 0.000 | -1.0864E-10 | -4.8927E-11 |
| VehMale | VehFemale | 1.72025E-11 | 0.018 | -7.6172E-11 | -7.4191E-12 |
| | CBDfemale | 1.72025E-11 | 0.000 | -1.2596E-10 | -5.7210E-11 |
| | THCfemale | 1.67506E-11 | 0.509 | -4.4599E-11 | 2.2348E-11 |
| | TpCfemale | 1.67506E-11 | 0.433 | -4.6690E-11 | 2.0257E-11 |
| | CBDmale | 1.63801E-11 | 0.862 | -3.5601E-11 | 2.9865E-11 |
| | THCmale | 1.67506E-11 | 0.418 | -4.7125E-11 | 1.9822E-11 |
| | TpCmale | 1.60706E-11 | 0.000 | -1.2411E-10 | -5.9883E-11 |
| CBDmale | VehFemale | 1.57664E-11 | 0.016 | -7.0434E-11 | -7.4213E-12 |
| | CBDfemale | 1.57664E-11 | 0.000 | -1.2023E-10 | -5.7212E-11 |
| | THCfemale | 1.52720E-11 | 0.591 | -3.8777E-11 | 2.2261E-11 |
| | TpCfemale | 1.52720E-11 | 0.500 | -4.0867E-11 | 2.0170E-11 |
| | VehMale | 1.63801E-11 | 0.862 | -2.9865E-11 | 3.5601E-11 |
| | THCmale | 1.52720E-11 | 0.483 | -4.1302E-11 | 1.9735E-11 |
| | TpCmale | 1.45229E-11 | 0.000 | -1.1815E-10 | -6.0108E-11 |
| THCmale | VehFemale | 1.61510E-11 | 0.086 | -6.0419E-11 | 4.1309E-12 |
| | CBDfemale | 1.61510E-11 | 0.000 | -1.1021E-10 | -4.5660E-11 |
| | THCfemale | 1.56687E-11 | 0.872 | -2.8786E-11 | 3.3837E-11 |
| | TpCfemale | 1.56687E-11 | 0.978 | -3.0877E-11 | 3.1746E-11 |
| | VehMale | 1.67506E-11 | 0.418 | -1.9822E-11 | 4.7125E-11 |
| | CBDmale | 1.52720E-11 | 0.483 | -1.9735E-11 | 4.1302E-11 |
| | TpCmale | 1.49396E-11 | 0.000 | -1.0820E-10 | -4.8492E-11 |
| TpCmale | VehFemale | 1.54446E-11 | 0.002 | 1.9338E-11 | 8.1065E-11 |
| | CBDfemale | 1.54446E-11 | 0.979 | -3.0453E-11 | 3.1274E-11 |
| | THCfemale | 1.49396E-11 | 0.000 | 5.1017E-11 | 1.1073E-10 |
| | TpCfemale | 1.49396E-11 | 0.000 | 4.8927E-11 | 1.0864E-10 |
| | VehMale | 1.60706E-11 | 0.000 | 5.9883E-11 | 1.2411E-10 |
| | CBDmale | 1.45229E-11 | 0.000 | 6.0108E-11 | 1.1815E-10 |
| | THCmale | 1.49396E-11 | 0.000 | 4.8492E-11 | 1.0820E-10 |

Supplemental Table 4.1 Descriptive statistics for 2-acyl glycerols (mol/L) in plasma collected from male and female adult mice 2h after receiving an i.p. injection of 10mg/kg CBD, THC, or THC+CBD

| | | N | Mean | Std. Deviation | Std. Error | 95% Confidence Interval for Mean | | Minimum | Maximum |
|---|---|---|---|---|---|---|---|---|---|
| | | | | | | Lower Bound | Upper Bound | | |
| 2-palmitoyl glycerol | Veh F | 7 | 1.5245E-05 | 5.89390E-06 | 2.22768E-06 | 9.7944E-06 | 2.0696E-05 | 3.68E-06 | 2.18E-05 |
| | CBD F | 9 | 1.8507E-05 | 1.33716E-05 | 4.45720E-06 | 8.2288E-06 | 2.8785E-05 | 5.25E-06 | 4.27E-05 |
| | THC F | 10 | 1.5679E-05 | 4.72406E-06 | 1.49388E-06 | 1.2300E-05 | 1.9059E-05 | 8.24E-06 | 2.21E-05 |
| | THC+CBD F | 10 | 1.1126E-05 | 3.51834E-06 | 1.11260E-06 | 8.6096E-06 | 1.3643E-05 | 5.15E-06 | 1.74E-05 |
| | Veh M | 8 | 1.3702E-05 | 5.60833E-06 | 1.98284E-06 | 9.0134E-06 | 1.8391E-05 | 5.86E-06 | 2.18E-05 |
| | CBD M | 10 | 1.9483E-05 | 5.04523E-06 | 1.59544E-06 | 1.5874E-05 | 2.3092E-05 | 1.12E-05 | 2.66E-05 |
| | THC M | 9 | 7.7991E-06 | 4.02281E-06 | 1.34094E-06 | 4.7069E-06 | 1.0891E-05 | 2.64E-06 | 1.44E-05 |
| | THC+CBD M | 9 | 1.2516E-05 | 5.28308E-06 | 1.76103E-06 | 8.4550E-06 | 1.6577E-05 | 4.83E-06 | 2.15E-05 |
| 2-oleoyl glycerol | Veh F | 7 | 9.0239E-05 | 2.34389E-05 | 8.85909E-06 | 6.8562E-05 | 1.1192E-04 | 6.62E-05 | 1.39E-04 |
| | CBD F | 9 | 1.3901E-04 | 1.06524E-04 | 3.55081E-05 | 5.7128E-05 | 2.2089E-04 | 2.07E-05 | 3.32E-04 |
| | THC F | 10 | 9.5045E-05 | 5.93326E-05 | 1.87626E-05 | 5.2601E-05 | 1.3749E-04 | 3.12E-05 | 1.72E-04 |
| | THC+CBD F | 11 | 5.5985E-05 | 2.40184E-05 | 7.24181E-06 | 3.9849E-05 | 7.2121E-05 | 2.29E-05 | 9.77E-05 |
| | Veh M | 8 | 1.4141E-04 | 7.83050E-05 | 2.76850E-05 | 7.5947E-05 | 2.0688E-04 | 4.36E-05 | 2.38E-04 |
| | CBD M | 11 | 1.6676E-04 | 1.03892E-04 | 3.13245E-05 | 9.6964E-05 | 2.3655E-04 | 4.21E-05 | 3.30E-04 |
| | THC M | 8 | 3.0231E-05 | 1.64842E-05 | 5.82803E-06 | 1.6450E-05 | 4.4013E-05 | 3.03E-06 | 5.47E-05 |
| | THC+CBD M | 10 | 1.3724E-04 | 1.01558E-04 | 3.21154E-05 | 6.4586E-05 | 2.0989E-04 | 1.18E-05 | 3.04E-04 |
| 2-linoleoyl glycerol | Veh F | 8 | 6.5390E-04 | 2.61715E-04 | 9.25302E-05 | 4.3510E-04 | 8.7269E-04 | 3.01E-04 | 1.17E-03 |
| | CBD F | 9 | 7.2952E-04 | 7.37512E-04 | 2.45837E-04 | 1.6261E-04 | 1.2964E-03 | 1.22E-04 | 2.01E-03 |
| | THC F | 9 | 2.6313E-04 | 1.20446E-04 | 4.01488E-05 | 1.7055E-04 | 3.5572E-04 | 4.25E-05 | 5.03E-04 |
| | THC+CBD F | 11 | 3.2753E-04 | 1.14923E-04 | 3.46506E-05 | 2.5032E-04 | 4.0473E-04 | 1.62E-04 | 4.99E-04 |
| | Veh M | 8 | 6.3538E-04 | 2.93370E-04 | 1.03722E-04 | 3.9011E-04 | 8.8064E-04 | 2.90E-04 | 1.12E-03 |
| | CBD M | 10 | 9.3972E-04 | 2.31826E-04 | 7.33099E-05 | 7.7388E-04 | 1.1056E-03 | 4.89E-04 | 1.25E-03 |
| | THC M | 9 | 2.7119E-04 | 1.73775E-04 | 5.79251E-05 | 1.3762E-04 | 4.0477E-04 | 3.72E-05 | 4.70E-04 |
| | THC+CBD M | 10 | 5.2519E-04 | 2.96883E-04 | 9.38828E-05 | 3.1281E-04 | 7.3756E-04 | 1.16E-04 | 9.22E-04 |
| 2-arachidonoyl glycerol | Veh F | 8 | 7.0291E-06 | 2.55236E-06 | 9.02394E-07 | 4.8952E-06 | 9.1629E-06 | 2.32E-06 | 1.02E-05 |
| | CBD F | 8 | 7.3132E-06 | 4.98934E-06 | 1.76400E-06 | 3.1420E-06 | 1.1484E-05 | 1.48E-06 | 1.72E-05 |
| | THC F | 10 | 4.8034E-06 | 3.64830E-06 | 1.15370E-06 | 2.1936E-06 | 7.4133E-06 | 9.08E-07 | 1.13E-05 |
| | THC+CBD F | 11 | 6.1314E-06 | 2.15981E-06 | 6.51208E-07 | 4.6804E-06 | 7.5824E-06 | 3.13E-06 | 9.45E-06 |
| | Veh M | 8 | 1.2133E-05 | 7.10492E-06 | 2.51197E-06 | 6.1930E-06 | 1.8073E-05 | 5.68E-06 | 2.35E-05 |
| | CBD M | 10 | 1.1281E-05 | 4.13014E-06 | 1.30607E-06 | 8.3264E-06 | 1.4235E-05 | 4.98E-06 | 1.79E-05 |
| | THC M | 9 | 4.4196E-06 | 2.72185E-06 | 9.07282E-07 | 2.3274E-06 | 6.5118E-06 | 1.19E-06 | 8.53E-06 |
| | THC+CBD M | 10 | 7.8863E-06 | 3.93430E-06 | 1.24413E-06 | 5.0719E-06 | 1.0701E-05 | 2.69E-06 | 1.34E-05 |

Supplemental Table 4.2 Descriptive statistics for 2-acyl glycerols (mol/g) in liver collected from male and female adult mice 2h after receiving an i.p. injection of 10mg/kg CBD, THC, or THC+CBD

| | | N | Mean | Std. Deviation | Std. Error | 95% Confidence Interval for Mean Lower Bound | 95% Confidence Interval for Mean Upper Bound | Minimum | Maximum |
|---|---|---|---|---|---|---|---|---|---|
| 2-palmitoyl glycerol | Veh F | 8 | 1.4863E-09 | 6.66235E-10 | 2.35550E-10 | 9.2928E-10 | 2.0433E-09 | 7.04E-10 | 2.47E-09 |
| | CBD F | 7 | 1.6485E-09 | 9.17628E-10 | 3.46831E-10 | 7.9985E-10 | 2.4972E-09 | 7.93E-10 | 3.52E-09 |
| | THC F | 9 | 2.4416E-09 | 1.03900E-09 | 3.46332E-10 | 1.6429E-09 | 3.2402E-09 | 1.33E-09 | 4.69E-09 |
| | THC+CBD F | 10 | 3.2352E-09 | 9.00946E-10 | 2.84904E-10 | 2.5907E-09 | 3.8797E-09 | 2.09E-09 | 4.77E-09 |
| | Veh M | 7 | 1.9459E-09 | 7.84574E-10 | 2.96541E-10 | 1.2202E-09 | 2.6715E-09 | 9.58E-10 | 3.45E-09 |
| | CBD M | 9 | 2.5528E-09 | 1.71132E-09 | 5.70439E-10 | 1.2374E-09 | 3.8683E-09 | 1.17E-09 | 6.48E-09 |
| | THC M | 9 | 2.9947E-09 | 1.34893E-09 | 4.49643E-10 | 1.9579E-09 | 4.0316E-09 | 1.45E-09 | 5.32E-09 |
| | THC+CBD M | 11 | 2.7397E-09 | 1.16974E-09 | 3.52690E-10 | 1.9539E-09 | 3.5256E-09 | 1.25E-09 | 4.85E-09 |
| 2-oleoyl glycerol | Veh F | 8 | 9.8120E-09 | 2.04234E-09 | 7.22077E-10 | 8.1045E-09 | 1.1519E-08 | 7.16E-09 | 1.31E-08 |
| | CBD F | 8 | 1.0653E-08 | 4.30537E-09 | 1.52218E-09 | 7.0532E-09 | 1.4252E-08 | 2.76E-09 | 1.62E-08 |
| | THC F | 10 | 3.2945E-08 | 1.18175E-08 | 3.73703E-09 | 2.4491E-08 | 4.1399E-08 | 1.25E-08 | 4.75E-08 |
| | THC+CBD F | 10 | 3.6401E-08 | 1.68174E-08 | 5.31814E-09 | 2.4370E-08 | 4.8431E-08 | 1.94E-08 | 6.75E-08 |
| | Veh M | 7 | 2.4032E-08 | 1.28237E-08 | 4.84689E-09 | 1.2172E-08 | 3.5892E-08 | 9.91E-09 | 4.21E-08 |
| | CBD M | 10 | 2.6167E-08 | 1.23076E-08 | 3.89199E-09 | 1.7363E-08 | 3.4972E-08 | 4.08E-09 | 4.17E-08 |
| | THC M | 8 | 2.8483E-08 | 1.68135E-08 | 5.94448E-09 | 1.4427E-08 | 4.2540E-08 | 5.34E-09 | 5.23E-08 |
| | THC+CBD M | 10 | 3.3725E-08 | 8.10014E-09 | 2.56149E-09 | 2.7931E-08 | 3.9520E-08 | 2.05E-08 | 4.36E-08 |
| 2-linoleoyl glycerol | Veh F | 8 | 9.2713E-09 | 1.57389E-09 | 5.56455E-10 | 7.9555E-09 | 1.0587E-08 | 6.73E-09 | 1.17E-08 |
| | CBD F | 8 | 1.0633E-08 | 4.66860E-09 | 1.65060E-09 | 6.7295E-09 | 1.4536E-08 | 4.23E-09 | 1.71E-08 |
| | THC F | 9 | 1.8118E-08 | 5.74259E-09 | 1.91420E-09 | 1.3704E-08 | 2.2532E-08 | 8.94E-09 | 2.72E-08 |
| | THC+CBD F | 10 | 3.2237E-08 | 1.34733E-08 | 4.26062E-09 | 2.2598E-08 | 4.1875E-08 | 1.24E-08 | 5.07E-08 |
| | Veh M | 7 | 1.7238E-08 | 2.49382E-09 | 9.42576E-10 | 1.4931E-08 | 1.9544E-08 | 1.43E-08 | 2.15E-08 |
| | CBD M | 9 | 2.1934E-08 | 4.81822E-09 | 1.60607E-09 | 1.8230E-08 | 2.5638E-08 | 1.69E-08 | 3.08E-08 |
| | THC M | 9 | 2.9194E-08 | 1.44472E-08 | 4.81573E-09 | 1.8089E-08 | 4.0299E-08 | 7.49E-09 | 5.39E-08 |
| | THC+CBD M | 11 | 2.5501E-08 | 1.28789E-08 | 3.88312E-09 | 1.6849E-08 | 3.4153E-08 | 1.14E-08 | 4.80E-08 |
| 2-arachidonoyl glycerol | Veh F | 7 | 1.1099E-08 | 1.64879E-09 | 6.23185E-10 | 9.5744E-09 | 1.2624E-08 | 9.37E-09 | 1.40E-08 |
| | CBD F | 8 | 1.4034E-08 | 4.15972E-09 | 1.47068E-09 | 1.0556E-08 | 1.7512E-08 | 8.21E-09 | 2.14E-08 |
| | THC F | 10 | 2.7176E-08 | 6.70955E-09 | 2.12175E-09 | 2.2376E-08 | 3.1976E-08 | 1.76E-08 | 3.80E-08 |
| | THC+CBD F | 10 | 3.4101E-08 | 1.34945E-08 | 4.26734E-09 | 2.4448E-08 | 4.3754E-08 | 1.26E-08 | 5.72E-08 |
| | Veh M | 7 | 2.2630E-08 | 6.72115E-09 | 2.54036E-09 | 1.6414E-08 | 2.8846E-08 | 1.63E-08 | 3.33E-08 |
| | CBD M | 9 | 2.4148E-08 | 6.48673E-09 | 2.16224E-09 | 1.9162E-08 | 2.9134E-08 | 1.14E-08 | 3.31E-08 |
| | THC M | 9 | 2.8984E-08 | 9.66924E-09 | 3.22308E-09 | 2.1552E-08 | 3.6416E-08 | 1.23E-08 | 4.57E-08 |
| | THC+CBD M | 11 | 2.4406E-08 | 9.98520E-09 | 3.01065E-09 | 1.7698E-08 | 3.1114E-08 | 1.50E-08 | 4.20E-08 |

Supplemental Table 4.3 Independent t-test comparing male and female levels of 2-acyl glycerols measured in plasma collected from adult mice 2h after an i.p. injection of 10mg/kg CBD, THC, or THC+CBD.

| | | | Levene's Test for Equality of Variances | | t-test for Equality of Means | | | | | | |
|---|---|---|---|---|---|---|---|---|---|---|---|
| | | | | | | | Significance | | | 95% Confidence Interval of the Difference | |
| | | | F | Sig. | t | df | One-Sided p | Two-Sided p | Std. Error Difference | Lower | Upper |
| Veh | 2-palmitoyl glycerol | Equal variances assumed | 0.130 | 0.724 | 0.519 | 13 | 0.306 | 0.612 | 2.97171E-06 | -4.87674E-06 | 7.96326E-06 |
| | | Equal variances not assumed | | | 0.517 | 12.531 | 0.307 | 0.614 | 2.98232E-06 | -4.92424E-06 | 8.01076E-06 |
| | 2-oleoyl glycerol | Equal variances assumed | 13.132 | 0.003 | -1.658 | 13 | 0.061 | 0.121 | 3.08593E-05 | -1.17840E-04 | 1.54951E-05 |
| | | Equal variances not assumed | | | -1.760 | 8.404 | 0.057 | 0.115 | 2.90679E-05 | -1.17646E-04 | 1.53014E-05 |
| | 2-linoleoyl glycerol | Equal variances assumed | 0.137 | 0.717 | 0.133 | 14 | 0.448 | 0.896 | 1.38997E-04 | -2.79600E-04 | 3.16637E-04 |
| | | Equal variances not assumed | | | 0.133 | 13.821 | 0.448 | 0.896 | 1.38997E-04 | -2.79962E-04 | 3.16998E-04 |
| | 2-arachidonoyl glycerol | Equal variances assumed | 5.951 | 0.029 | -1.912 | 14 | 0.038 | 0.077 | 2.66914E-06 | -1.08285E-05 | 6.20951E-07 |
| | | Equal variances not assumed | | | -1.912 | 8.777 | 0.044 | 0.089 | 2.66914E-06 | -1.11652E-05 | 9.57681E-07 |
| CBD | 2-palmitoyl glycerol | Equal variances assumed | 6.716 | 0.019 | -0.215 | 17 | 0.416 | 0.832 | 4.53960E-06 | -1.05533E-05 | 8.60212E-06 |
| | | Equal variances not assumed | | | -0.206 | 10.035 | 0.420 | 0.841 | 4.73414E-06 | -1.15190E-05 | 9.56774E-06 |
| | 2-oleoyl glycerol | Equal variances assumed | 0.106 | 0.749 | -0.588 | 18 | 0.282 | 0.564 | 4.72254E-05 | -1.26967E-04 | 7.14668E-05 |
| | | Equal variances not assumed | | | -0.586 | 17.041 | 0.283 | 0.566 | 4.73503E-05 | -1.27632E-04 | 7.21322E-05 |
| | 2-linoleoyl glycerol | Equal variances assumed | 10.537 | 0.005 | -0.858 | 17 | 0.201 | 0.403 | 2.45038E-04 | -7.27187E-04 | 3.06782E-04 |
| | | Equal variances not assumed | | | -0.819 | 9.420 | 0.216 | 0.433 | 2.56535E-04 | -7.86607E-04 | 3.66201E-04 |

| | | | | | | | | | | | |
|---|---|---|---|---|---|---|---|---|---|---|---|
| | 2-arachidonoyl glycerol | Equal variances assumed | 0.131 | 0.723 | -1.848 | 16 | 0.042 | 0.083 | 2.14694E-06 | -8.51907E-06 | 5.83568E-07 |
| | | Equal variances not assumed | | | -1.808 | 13.600 | 0.046 | 0.093 | 2.19488E-06 | -8.68833E-06 | 7.52834E-07 |
| THC | 2-palmitoyl glycerol | Equal variances assumed | 0.094 | 0.763 | 3.891 | 17 | 0.001 | 0.001 | 2.02533E-06 | 3.60731E-06 | 1.21534E-05 |
| | | Equal variances not assumed | | | 3.926 | 16.960 | 0.001 | 0.001 | 2.00743E-06 | 3.64429E-06 | 1.21165E-05 |
| | 2-oleoyl glycerol | Equal variances assumed | 21.850 | 0.000 | 2.982 | 16 | 0.004 | 0.009 | 2.17323E-05 | 1.87427E-05 | 1.10884E-04 |
| | | Equal variances not assumed | | | 3.299 | 10.693 | 0.004 | 0.007 | 1.96469E-05 | 2.14184E-05 | 1.08208E-04 |
| | 2-linoleoyl glycerol | Equal variances assumed | 2.569 | 0.128 | -0.114 | 16 | 0.455 | 0.910 | 7.04787E-05 | -1.57468E-04 | 1.41348E-04 |
| | | Equal variances not assumed | | | -0.114 | 14.245 | 0.455 | 0.911 | 7.04787E-05 | -1.58978E-04 | 1.42858E-04 |
| | 2-arachidonoyl glycerol | Equal variances assumed | 0.328 | 0.574 | 0.257 | 17 | 0.400 | 0.800 | 1.49118E-06 | -2.76227E-06 | 3.52994E-06 |
| | | Equal variances not assumed | | | 0.262 | 16.482 | 0.398 | 0.797 | 1.46771E-06 | -2.72019E-06 | 3.48786E-06 |
| THC+CBD | 2-palmitoyl glycerol | Equal variances assumed | 1.211 | 0.286 | -0.682 | 17 | 0.252 | 0.505 | 2.03871E-06 | -5.69083E-06 | 2.91179E-06 |
| | | Equal variances not assumed | | | -0.667 | 13.718 | 0.258 | 0.516 | 2.08305E-06 | -5.86583E-06 | 3.08680E-06 |
| | 2-oleoyl glycerol | Equal variances assumed | 12.733 | 0.002 | -2.581 | 19 | 0.009 | 0.018 | 3.14748E-05 | -1.47129E-04 | -1.53736E-05 |
| | | Equal variances not assumed | | | -2.468 | 9.915 | 0.017 | 0.033 | 3.29218E-05 | -1.54690E-04 | -7.81205E-06 |
| | 2-linoleoyl glycerol | Equal variances assumed | 8.755 | 0.008 | -2.050 | 19 | 0.027 | 0.054 | 9.64240E-05 | -3.99475E-04 | 4.16061E-06 |
| | | Equal variances not assumed | | | -1.975 | 11.428 | 0.036 | 0.073 | 1.00073E-04 | -4.16914E-04 | 2.15999E-05 |
| | 2-arachidonoyl glycerol | Equal variances assumed | 4.447 | 0.048 | -1.284 | 19 | 0.107 | 0.215 | 1.36692E-06 | -4.61590E-06 | 1.10608E-06 |
| | | Equal variances not assumed | | | -1.250 | 13.683 | 0.116 | 0.232 | 1.40426E-06 | -4.77331E-06 | 1.26349E-06 |

Supplemental Table 4.4 Independent t-test comparing male and female levels of 2-acyl glycerols measured in liver collected from adult mice 2h after an i.p. injection of 10mg/kg CBD, THC, or THC+CBD.

| | | | Levene's Test for Equality of Variances | | t-test for Equality of Means | | | | | | | |
|---|---|---|---|---|---|---|---|---|---|---|---|---|
| | | | | | | | Significance | | | | 95% Confidence Interval of the Difference | |
| | | | F | Sig. | t | df | One-Sided p | Two-Sided p | Mean Difference | Std. Error Difference | Lower | Upper |
| Veh | 2-palmitoyl glycerol | Equal variances assumed | 0.020 | 0.890 | -1.228 | 13 | 0.121 | 0.241 | -4.59589E-10 | 3.74324E-10 | -1.26827E-09 | 3.49089E-10 |
| | | Equal variances not assumed | | | -1.214 | 11.900 | 0.124 | 0.248 | -4.59589E-10 | 3.78709E-10 | -1.28550E-09 | 3.66320E-10 |
| | 2-oleoyl glycerol | Equal variances assumed | 37.829 | 0.000 | -3.108 | 13 | 0.004 | 0.008 | -1.42201E-08 | 4.57509E-09 | -2.41040E-08 | -4.33621E-09 |
| | | Equal variances not assumed | | | -2.902 | 6.267 | 0.013 | 0.026 | -1.42201E-08 | 4.90038E-09 | -2.60883E-08 | -2.35190E-09 |
| | 2-linoleoyl glycerol | Equal variances assumed | 1.029 | 0.329 | -7.507 | 13 | 0.000 | 0.000 | -7.96649E-09 | 1.06119E-09 | -1.02591E-08 | -5.67392E-09 |
| | | Equal variances not assumed | | | -7.278 | 9.882 | 0.000 | 0.000 | -7.96649E-09 | 1.09457E-09 | -1.04093E-08 | -5.52368E-09 |
| | 2-arachidonoyl glycerol | Equal variances assumed | 9.471 | 0.010 | -4.408 | 12 | 0.000 | 0.001 | -1.15307E-08 | 2.61568E-09 | -1.72298E-08 | -5.83167E-09 |
| | | Equal variances not assumed | | | -4.408 | 6.720 | 0.002 | 0.003 | -1.15307E-08 | 2.61568E-09 | -1.77686E-08 | -5.29293E-09 |
| | 2-palmitoyl glycerol | Equal variances assumed | 1.695 | 0.214 | -1.258 | 14 | 0.114 | 0.229 | -9.04322E-10 | 7.18794E-10 | -2.44598E-09 | 6.37337E-10 |
| | | Equal variances not assumed | | | -1.355 | 12.695 | 0.100 | 0.199 | -9.04322E-10 | 6.67602E-10 | -2.35012E-09 | 5.41476E-10 |
| | 2-oleoyl glycerol | Equal variances assumed | 7.714 | 0.013 | -3.386 | 16 | 0.002 | 0.004 | -1.55148E-08 | 4.58212E-09 | -2.52285E-08 | -5.80117E-09 |
| | | Equal variances not assumed | | | -3.713 | 11.615 | 0.002 | 0.003 | -1.55148E-08 | 4.17907E-09 | -2.46539E-08 | -6.37579E-09 |
| | 2-linoleoyl glycerol | Equal variances assumed | 0.034 | 0.856 | -4.898 | 15 | 0.000 | 0.000 | -1.13015E-08 | 2.30759E-09 | -1.62200E-08 | -6.38298E-09 |
| | | Equal variances not assumed | | | -4.907 | 14.868 | 0.000 | 0.000 | -1.13015E-08 | 2.30303E-09 | -1.62141E-08 | -6.38891E-09 |
| | 2-arachidonoyl glycerol | Equal variances assumed | 0.715 | 0.411 | -3.768 | 15 | 0.001 | 0.002 | -1.01137E-08 | 2.68426E-09 | -1.58350E-08 | -4.39234E-09 |

| | | | | | | | | | | | | |
|---|---|---|---|---|---|---|---|---|---|---|---|---|
| | | Equal variances not assumed | | | -3.868 | 13.751 | 0.001 | 0.002 | -1.01137E-08 | 2.61499E-09 | -1.57318E-08 | -4.49553E-09 |
| THC | 2-palmitoyl glycerol | Equal variances assumed | 0.454 | 0.510 | -0.975 | 16 | 0.172 | 0.344 | -5.53160E-10 | 5.67560E-10 | -1.75633E-09 | 6.50014E-10 |
| | | Equal variances not assumed | | | -0.975 | 15.021 | 0.173 | 0.345 | -5.53160E-10 | 5.67560E-10 | -1.76274E-09 | 6.56418E-10 |
| | 2-oleoyl glycerol | Equal variances assumed | 0.809 | 0.382 | 0.661 | 16 | 0.259 | 0.518 | 4.46161E-09 | 6.74557E-09 | -9.83836E-09 | 1.87616E-08 |
| | | Equal variances not assumed | | | 0.635 | 12.150 | 0.268 | 0.537 | 4.46161E-09 | 7.02156E-09 | -1.08161E-08 | 1.97393E-08 |
| | 2-linoleoyl glycerol | Equal variances assumed | 4.963 | 0.041 | -2.137 | 16 | 0.024 | 0.048 | -1.10765E-08 | 5.18222E-09 | -2.20623E-08 | -9.06743E-11 |
| | | Equal variances not assumed | | | -2.137 | 10.466 | 0.029 | 0.057 | -1.10765E-08 | 5.18222E-09 | -2.25538E-08 | 4.00853E-10 |
| | 2-arachidonoyl glycerol | Equal variances assumed | 0.596 | 0.451 | -0.478 | 17 | 0.319 | 0.639 | -1.80797E-09 | 3.78414E-09 | -9.79181E-09 | 6.17587E-09 |
| | | Equal variances not assumed | | | -0.469 | 14.085 | 0.323 | 0.647 | -1.80797E-09 | 3.85876E-09 | -1.00795E-08 | 6.46358E-09 |
| THC+CBD | 2-palmitoyl glycerol | Equal variances assumed | 1.374 | 0.256 | 1.079 | 19 | 0.147 | 0.294 | 4.95491E-10 | 4.59224E-10 | -4.65676E-10 | 1.45666E-09 |
| | | Equal variances not assumed | | | 1.093 | 18.538 | 0.144 | 0.288 | 4.95491E-10 | 4.53388E-10 | -4.55064E-10 | 1.44605E-09 |
| | 2-oleoyl glycerol | Equal variances assumed | 6.072 | 0.024 | 0.453 | 18 | 0.328 | 0.656 | 2.67563E-09 | 5.90287E-09 | -9.72583E-09 | 1.50771E-08 |
| | | Equal variances not assumed | | | 0.453 | 12.963 | 0.329 | 0.658 | 2.67563E-09 | 5.90287E-09 | -1.00805E-08 | 1.54317E-08 |
| | 2-linoleoyl glycerol | Equal variances assumed | 0.394 | 0.537 | 1.171 | 19 | 0.128 | 0.256 | 6.73558E-09 | 5.75166E-09 | -5.30278E-09 | 1.87739E-08 |
| | | Equal variances not assumed | | | 1.168 | 18.607 | 0.129 | 0.257 | 6.73558E-09 | 5.76468E-09 | -5.34730E-09 | 1.88185E-08 |
| | 2-arachidonoyl glycerol | Equal variances assumed | 1.187 | 0.290 | 1.884 | 19 | 0.037 | 0.075 | 9.69499E-09 | 5.14643E-09 | -1.07660E-09 | 2.04666E-08 |
| | | Equal variances not assumed | | | 1.856 | 16.508 | 0.041 | 0.081 | 9.69499E-09 | 5.22247E-09 | -1.34851E-09 | 2.07385E-08 |

Supplemental Table 4.5 One-way ANOVA for 2-acyl glycerols measured in plasma collected from adult mice 2h after an i.p. injection of 10mg/kg CBD, THC, or THC+CBD

| | | Sum of Squares | df | Mean Square | F | Sig. |
|---|---|---|---|---|---|---|
| 2-palmitoyl glycerol | Between Groups | 9.66E-10 | 7 | 1.38E-10 | 3.207 | 0.0056379 |
| | Within Groups | 2.75E-09 | 64 | 4.30E-11 | | |
| | Total | 3.72E-09 | 71 | | | |
| 2-oleoyl glycerol | Between Groups | 1.46E-07 | 7 | 2.09E-08 | 3.653 | 0.0021651 |
| | Within Groups | 3.77E-07 | 66 | 5.71E-09 | | |
| | Total | 5.23E-07 | 73 | | | |
| 2-linoleoyl glycerol | Between Groups | 3.94E-06 | 7 | 5.62E-07 | 5.155 | 0.0001022 |
| | Within Groups | 7.20E-06 | 66 | 1.09E-07 | | |
| | Total | 1.11E-05 | 73 | | | |
| 2-arachidonoyl glycerol | Between Groups | 4.97E-10 | 7 | 7.10E-11 | 4.290 | 0.0005821 |
| | Within Groups | 1.09E-09 | 66 | 1.65E-11 | | |
| | Total | 1.59E-09 | 73 | | | |

Supplemental Table 4.6 One-way ANOVA for 2-acyl glycerols measured in liver collected from adult mice 2h after an i.p. injection of 10mg/kg CBD, THC, or THC+CBD

| | | Sum of Squares | df | Mean Square | F | Sig. |
|---|---|---|---|---|---|---|
| 2-palmitoyl glycerol | Between Groups | 2.36E-17 | 7 | 3.37E-18 | 2.626 | 0.01924 |
| | Within Groups | 7.95E-17 | 62 | 1.28E-18 | | |
| | Total | 1.03E-16 | 69 | | | |
| 2-oleoyl glycerol | Between Groups | 6.22E-15 | 7 | 8.88E-16 | 6.301 | 0.00001 |
| | Within Groups | 8.88E-15 | 63 | 1.41E-16 | | |
| | Total | 1.51E-14 | 70 | | | |
| 2-linoleoyl glycerol | Between Groups | 4.23E-15 | 7 | 6.04E-16 | 6.773 | 0.00001 |
| | Within Groups | 5.62E-15 | 63 | 8.92E-17 | | |
| | Total | 9.85E-15 | 70 | | | |
| 2-arachidonoyl glycerol | Between Groups | 3.32E-15 | 7 | 4.74E-16 | 6.588 | 0.00001 |
| | Within Groups | 4.53E-15 | 63 | 7.20E-17 | | |
| | Total | 7.85E-15 | 70 | | | |

#### Supplemental Table 4.7 Fisher's LSD post-hoc test for plasma 2-acyl glycerol ANOVA

| | | | Std. Error | Sig. | 95% Confidence Interval Lower Bound | Upper Bound |
|---|---|---|---|---|---|---|
| 2-palmitoyl glycerol | VehFemale | CBDfemale | 3.31E-06 | 0.3274 | -9.86E-06 | 3.34E-06 |
| | | THCfemale | 3.23E-06 | 0.8936 | -6.89E-06 | 6.02E-06 |
| | | TpCfemale | 3.23E-06 | 0.2071 | -2.34E-06 | 1.06E-05 |
| | | VehMale | 3.39E-06 | 0.6509 | -5.24E-06 | 8.32E-06 |
| | | CBDmale | 3.23E-06 | 0.1945 | -1.07E-05 | 2.22E-06 |
| | | THCmale | 3.31E-06 | 0.0277 | 8.43E-07 | 1.40E-05 |
| | | TpCmale | 3.31E-06 | 0.4120 | -3.87E-06 | 9.33E-06 |
| | CBDfemale | VehFemale | 3.31E-06 | 0.3274 | -3.34E-06 | 9.86E-06 |
| | | THCfemale | 3.01E-06 | 0.3516 | -3.19E-06 | 8.85E-06 |
| | | TpCfemale | 3.01E-06 | 0.0171 | 1.36E-06 | 1.34E-05 |
| | | VehMale | 3.19E-06 | 0.1365 | -1.56E-06 | 1.12E-05 |
| | | CBDmale | 3.01E-06 | 0.7472 | -7.00E-06 | 5.04E-06 |
| | | THCmale | 3.09E-06 | 0.0010 | 4.53E-06 | 1.69E-05 |
| | | TpCmale | 3.09E-06 | 0.0571 | -1.85E-07 | 1.22E-05 |
| | THCfemale | VehFemale | 3.23E-06 | 0.8936 | -6.02E-06 | 6.89E-06 |
| | | CBDfemale | 3.01E-06 | 0.3516 | -8.85E-06 | 3.19E-06 |
| | | TpCfemale | 2.93E-06 | 0.1255 | -1.31E-06 | 1.04E-05 |
| | | VehMale | 3.11E-06 | 0.5273 | -4.24E-06 | 8.19E-06 |
| | | CBDmale | 2.93E-06 | 0.1994 | -9.66E-06 | 2.06E-06 |
| | | THCmale | 3.01E-06 | 0.0111 | 1.86E-06 | 1.39E-05 |
| | | TpCmale | 3.01E-06 | 0.2978 | -2.86E-06 | 9.18E-06 |
| | TpCfemale | VehFemale | 3.23E-06 | 0.2071 | -1.06E-05 | 2.34E-06 |
| | | CBDfemale | 3.01E-06 | 0.0171 | -1.34E-05 | -1.36E-06 |
| | | THCfemale | 2.93E-06 | 0.1255 | -1.04E-05 | 1.31E-06 |
| | | VehMale | 3.11E-06 | 0.4108 | -8.79E-06 | 3.64E-06 |
| | | CBDmale | 2.93E-06 | 0.0059 | -1.42E-05 | -2.50E-06 |
| | | THCmale | 3.01E-06 | 0.2737 | -2.69E-06 | 9.35E-06 |
| | | TpCmale | 3.01E-06 | 0.6463 | -7.41E-06 | 4.63E-06 |
| | VehMale | VehFemale | 3.39E-06 | 0.6509 | -8.32E-06 | 5.24E-06 |
| | | CBDfemale | 3.19E-06 | 0.1365 | -1.12E-05 | 1.56E-06 |
| | | THCfemale | 3.11E-06 | 0.5273 | -8.19E-06 | 4.24E-06 |
| | | TpCfemale | 3.11E-06 | 0.4108 | -3.64E-06 | 8.79E-06 |
| | | CBDmale | 3.11E-06 | 0.0678 | -1.20E-05 | 4.34E-07 |
| | | THCmale | 3.19E-06 | 0.0686 | -4.64E-07 | 1.23E-05 |
| | | TpCmale | 3.19E-06 | 0.7110 | -5.18E-06 | 7.55E-06 |
| | CBDmale | VehFemale | 3.23E-06 | 0.1945 | -2.22E-06 | 1.07E-05 |
| | | CBDfemale | 3.01E-06 | 0.7472 | -5.04E-06 | 7.00E-06 |
| | | THCfemale | 2.93E-06 | 0.1994 | -2.06E-06 | 9.66E-06 |
| | | TpCfemale | 2.93E-06 | 0.0059 | 2.50E-06 | 1.42E-05 |
| | | VehMale | 3.11E-06 | 0.0678 | -4.34E-07 | 1.20E-05 |
| | | THCmale | 3.01E-06 | 0.0003 | 5.66E-06 | 1.77E-05 |
| | | TpCmale | 3.01E-06 | 0.0240 | 9.47E-07 | 1.30E-05 |
| | THCmale | VehFemale | 3.31E-06 | 0.0277 | -1.40E-05 | -8.43E-07 |
| | | CBDfemale | 3.09E-06 | 0.0010 | -1.69E-05 | -4.53E-06 |
| | | THCfemale | 3.01E-06 | 0.0111 | -1.39E-05 | -1.86E-06 |
| | | TpCfemale | 3.01E-06 | 0.2737 | -9.35E-06 | 2.69E-06 |
| | | VehMale | 3.19E-06 | 0.0686 | -1.23E-05 | 4.64E-07 |
| | | CBDmale | 3.01E-06 | 0.0003 | -1.77E-05 | -5.66E-06 |
| | | TpCmale | 3.09E-06 | 0.1320 | -1.09E-05 | 1.46E-06 |
| | TpCmale | VehFemale | 3.31E-06 | 0.4120 | -9.33E-06 | 3.87E-06 |
| | | CBDfemale | 3.09E-06 | 0.0571 | -1.22E-05 | 1.85E-07 |
| | | THCfemale | 3.01E-06 | 0.2978 | -9.18E-06 | 2.86E-06 |
| | | TpCfemale | 3.01E-06 | 0.6463 | -4.63E-06 | 7.41E-06 |
| | | VehMale | 3.19E-06 | 0.7110 | -7.55E-06 | 5.18E-06 |
| | | CBDmale | 3.01E-06 | 0.0240 | -1.30E-05 | -9.47E-07 |
| | | THCmale | 3.09E-06 | 0.1320 | -1.46E-06 | 1.09E-05 |
| 2-oleoyl glycerol | VehFemale | CBDfemale | 3.81E-05 | 0.2049 | -1.25E-04 | 2.73E-05 |

| | | | | | | |
|---|---|---|---|---|---|---|
| | | THCfemale | 3.73E-05 | 0.8977 | -7.92E-05 | 6.96E-05 |
| | | TpCfemale | 3.65E-05 | 0.3520 | -3.87E-05 | 1.07E-04 |
| | | VehMale | 3.91E-05 | 0.1954 | -1.29E-04 | 2.69E-05 |
| | | CBDmale | 3.65E-05 | 0.0401 | -1.49E-04 | -3.55E-06 |
| | | THCmale | 3.91E-05 | 0.1298 | -1.81E-05 | 1.38E-04 |
| | | TpCmale | 3.73E-05 | 0.2115 | -1.21E-04 | 2.74E-05 |
| | CBDfemale | VehFemale | 3.81E-05 | 0.2049 | -2.73E-05 | 1.25E-04 |
| | | THCfemale | 3.47E-05 | 0.2100 | -2.54E-05 | 1.13E-04 |
| | | TpCfemale | 3.40E-05 | 0.0172 | 1.52E-05 | 1.51E-04 |
| | | VehMale | 3.67E-05 | 0.9481 | -7.57E-05 | 7.09E-05 |
| | | CBDmale | 3.40E-05 | 0.4170 | -9.56E-05 | 4.01E-05 |
| | | THCmale | 3.67E-05 | 0.0043 | 3.54E-05 | 1.82E-04 |
| | | TpCmale | 3.47E-05 | 0.9594 | -6.76E-05 | 7.11E-05 |
| | THCfemale | VehFemale | 3.73E-05 | 0.8977 | -6.96E-05 | 7.92E-05 |
| | | CBDfemale | 3.47E-05 | 0.2100 | -1.13E-04 | 2.54E-05 |
| | | TpCfemale | 3.30E-05 | 0.2412 | -2.69E-05 | 1.05E-04 |
| | | VehMale | 3.59E-05 | 0.2005 | -1.18E-04 | 2.52E-05 |
| | | CBDmale | 3.30E-05 | 0.0335 | -1.38E-04 | -5.77E-06 |
| | | THCmale | 3.59E-05 | 0.0752 | -6.77E-06 | 1.36E-04 |
| | | TpCmale | 3.38E-05 | 0.2164 | -1.10E-04 | 2.53E-05 |
| | TpCfemale | VehFemale | 3.65E-05 | 0.3520 | -1.07E-04 | 3.87E-05 |
| | | CBDfemale | 3.40E-05 | 0.0172 | -1.51E-04 | -1.52E-05 |
| | | THCfemale | 3.30E-05 | 0.2412 | -1.05E-04 | 2.69E-05 |
| | | VehMale | 3.51E-05 | 0.0177 | -1.56E-04 | -1.53E-05 |
| | | CBDmale | 3.22E-05 | 0.0010 | -1.75E-04 | -4.64E-05 |
| | | THCmale | 3.51E-05 | 0.4660 | -4.44E-05 | 9.59E-05 |
| | | TpCmale | 3.30E-05 | 0.0165 | -1.47E-04 | -1.53E-05 |
| | VehMale | VehFemale | 3.91E-05 | 0.1954 | -2.69E-05 | 1.29E-04 |
| | | CBDfemale | 3.67E-05 | 0.9481 | -7.09E-05 | 7.57E-05 |
| | | THCfemale | 3.59E-05 | 0.2005 | -2.52E-05 | 1.18E-04 |
| | | TpCfemale | 3.51E-05 | 0.0177 | 1.53E-05 | 1.56E-04 |
| | | CBDmale | 3.51E-05 | 0.4730 | -9.55E-05 | 4.48E-05 |
| | | THCmale | 3.78E-05 | 0.0045 | 3.57E-05 | 1.87E-04 |
| | | TpCmale | 3.59E-05 | 0.9077 | -6.74E-05 | 7.58E-05 |
| | CBDmale | VehFemale | 3.65E-05 | 0.0401 | 3.55E-06 | 1.49E-04 |
| | | CBDfemale | 3.40E-05 | 0.4170 | -4.01E-05 | 9.56E-05 |
| | | THCfemale | 3.30E-05 | 0.0335 | 5.77E-06 | 1.38E-04 |
| | | TpCfemale | 3.22E-05 | 0.0010 | 4.64E-05 | 1.75E-04 |
| | | VehMale | 3.51E-05 | 0.4730 | -4.48E-05 | 9.55E-05 |
| | | THCmale | 3.51E-05 | 0.0002 | 6.64E-05 | 2.07E-04 |
| | | TpCmale | 3.30E-05 | 0.3746 | -3.64E-05 | 9.55E-05 |
| | THCmale | VehFemale | 3.91E-05 | 0.1298 | -1.38E-04 | 1.81E-05 |
| | | CBDfemale | 3.67E-05 | 0.0043 | -1.82E-04 | -3.54E-05 |
| | | THCfemale | 3.59E-05 | 0.0752 | -1.36E-04 | 6.77E-06 |
| | | TpCfemale | 3.51E-05 | 0.4660 | -9.59E-05 | 4.44E-05 |
| | | VehMale | 3.78E-05 | 0.0045 | -1.87E-04 | -3.57E-05 |
| | | CBDmale | 3.51E-05 | 0.0002 | -2.07E-04 | -6.64E-05 |
| | | TpCmale | 3.59E-05 | 0.0040 | -1.79E-04 | -3.54E-05 |
| | TpCmale | VehFemale | 3.73E-05 | 0.2115 | -2.74E-05 | 1.21E-04 |
| | | CBDfemale | 3.47E-05 | 0.9594 | -7.11E-05 | 6.76E-05 |
| | | THCfemale | 3.38E-05 | 0.2164 | -2.53E-05 | 1.10E-04 |
| | | TpCfemale | 3.30E-05 | 0.0165 | 1.53E-05 | 1.47E-04 |
| | | VehMale | 3.59E-05 | 0.9077 | -7.58E-05 | 6.74E-05 |
| | | CBDmale | 3.30E-05 | 0.3746 | -9.55E-05 | 3.64E-05 |
| | | THCmale | 3.59E-05 | 0.0040 | 3.54E-05 | 1.79E-04 |
| | VehFemale | CBDfemale | 1.60E-04 | 0.6391 | -3.96E-04 | 2.45E-04 |
| | | THCfemale | 1.60E-04 | 0.0176 | 7.03E-05 | 7.11E-04 |
| 2-linoleoyl glycerol | | TpCfemale | 1.53E-04 | 0.0372 | 2.00E-05 | 6.33E-04 |
| | | VehMale | 1.65E-04 | 0.9111 | -3.11E-04 | 3.48E-04 |
| | | CBDmale | 1.57E-04 | 0.0726 | -5.99E-04 | 2.70E-05 |

| | | | | | | |
|---|---|---|---|---|---|---|
| | | THCmale | 1.60E-04 | 0.0200 | 6.23E-05 | 7.03E-04 |
| | | TpCmale | 1.57E-04 | 0.4143 | -1.84E-04 | 4.42E-04 |
| | CBDfemale | VehFemale | 1.60E-04 | 0.6391 | -2.45E-04 | 3.96E-04 |
| | | THCfemale | 1.56E-04 | 0.0039 | 1.56E-04 | 7.77E-04 |
| | | TpCfemale | 1.48E-04 | 0.0086 | 1.06E-04 | 6.98E-04 |
| | | VehMale | 1.60E-04 | 0.5595 | -2.26E-04 | 4.15E-04 |
| | | CBDmale | 1.52E-04 | 0.1707 | -5.13E-04 | 9.28E-05 |
| | | THCmale | 1.56E-04 | 0.0045 | 1.47E-04 | 7.69E-04 |
| | | TpCmale | 1.52E-04 | 0.1828 | -9.87E-05 | 5.07E-04 |
| | THCfemale | VehFemale | 1.60E-04 | 0.0176 | -7.11E-04 | -7.03E-05 |
| | | CBDfemale | 1.56E-04 | 0.0039 | -7.77E-04 | -1.56E-04 |
| | | TpCfemale | 1.48E-04 | 0.6659 | -3.61E-04 | 2.32E-04 |
| | | VehMale | 1.60E-04 | 0.0235 | -6.93E-04 | -5.18E-05 |
| | | CBDmale | 1.52E-04 | 0.0000 | -9.80E-04 | -3.74E-04 |
| | | THCmale | 1.56E-04 | 0.9589 | -3.19E-04 | 3.03E-04 |
| | | TpCmale | 1.52E-04 | 0.0889 | -5.65E-04 | 4.09E-05 |
| | TpCfemale | VehFemale | 1.53E-04 | 0.0372 | -6.33E-04 | -2.00E-05 |
| | | CBDfemale | 1.48E-04 | 0.0086 | -6.98E-04 | -1.06E-04 |
| | | THCfemale | 1.48E-04 | 0.6659 | -2.32E-04 | 3.61E-04 |
| | | VehMale | 1.53E-04 | 0.0490 | -6.14E-04 | -1.43E-06 |
| | | CBDmale | 1.44E-04 | 0.0001 | -9.00E-04 | -3.24E-04 |
| | | THCmale | 1.48E-04 | 0.7056 | -2.40E-04 | 3.53E-04 |
| | | TpCmale | 1.44E-04 | 0.1754 | -4.86E-04 | 9.05E-05 |
| | VehMale | VehFemale | 1.65E-04 | 0.9111 | -3.48E-04 | 3.11E-04 |
| | | CBDfemale | 1.60E-04 | 0.5595 | -4.15E-04 | 2.26E-04 |
| | | THCfemale | 1.60E-04 | 0.0235 | 5.18E-05 | 6.93E-04 |
| | | TpCfemale | 1.53E-04 | 0.0490 | 1.43E-06 | 6.14E-04 |
| | | CBDmale | 1.57E-04 | 0.0563 | -6.17E-04 | 8.46E-06 |
| | | THCmale | 1.60E-04 | 0.0265 | 4.38E-05 | 6.85E-04 |
| | | TpCmale | 1.57E-04 | 0.4843 | -2.03E-04 | 4.23E-04 |
| | CBDmale | VehFemale | 1.57E-04 | 0.0726 | -2.70E-05 | 5.99E-04 |
| | | CBDfemale | 1.52E-04 | 0.1707 | -9.28E-05 | 5.13E-04 |
| | | THCfemale | 1.52E-04 | 0.0000 | 3.74E-04 | 9.80E-04 |
| | | TpCfemale | 1.44E-04 | 0.0001 | 3.24E-04 | 9.00E-04 |
| | | VehMale | 1.57E-04 | 0.0563 | -8.46E-06 | 6.17E-04 |
| | | THCmale | 1.52E-04 | 0.0000 | 3.66E-04 | 9.72E-04 |
| | | TpCmale | 1.48E-04 | 0.0066 | 1.20E-04 | 7.09E-04 |
| | THCmale | VehFemale | 1.60E-04 | 0.0200 | -7.03E-04 | -6.23E-05 |
| | | CBDfemale | 1.56E-04 | 0.0045 | -7.69E-04 | -1.47E-04 |
| | | THCfemale | 1.56E-04 | 0.9589 | -3.03E-04 | 3.19E-04 |
| | | TpCfemale | 1.48E-04 | 0.7056 | -3.53E-04 | 2.40E-04 |
| | | VehMale | 1.60E-04 | 0.0265 | -6.85E-04 | -4.38E-05 |
| | | CBDmale | 1.52E-04 | 0.0000 | -9.72E-04 | -3.66E-04 |
| | | TpCmale | 1.52E-04 | 0.0989 | -5.57E-04 | 4.90E-05 |
| | TpCmale | VehFemale | 1.57E-04 | 0.4143 | -4.42E-04 | 1.84E-04 |
| | | CBDfemale | 1.52E-04 | 0.1828 | -5.07E-04 | 9.87E-05 |
| | | THCfemale | 1.52E-04 | 0.0889 | -4.09E-05 | 5.65E-04 |
| | | TpCfemale | 1.44E-04 | 0.1754 | -9.05E-05 | 4.86E-04 |
| | | VehMale | 1.57E-04 | 0.4843 | -4.23E-04 | 2.03E-04 |
| | | CBDmale | 1.48E-04 | 0.0066 | -7.09E-04 | -1.20E-04 |
| | | THCmale | 1.52E-04 | 0.0989 | -4.90E-05 | 5.57E-04 |
| 2-arachidonoyl glycerol | VehFemale | CBDfemale | 2.03E-06 | 0.8893 | -4.34E-06 | 3.78E-06 |
| | | THCfemale | 1.93E-06 | 0.2528 | -1.63E-06 | 6.08E-06 |
| | | TpCfemale | 1.89E-06 | 0.6364 | -2.88E-06 | 4.67E-06 |
| | | VehMale | 2.03E-06 | 0.0145 | -9.16E-06 | -1.04E-06 |
| | | CBDmale | 1.93E-06 | 0.0310 | -8.10E-06 | -4.00E-07 |
| | | THCmale | 1.98E-06 | 0.1913 | -1.34E-06 | 6.56E-06 |
| | | TpCmale | 1.93E-06 | 0.6582 | -4.71E-06 | 2.99E-06 |
| | CBDfemale | VehFemale | 2.03E-06 | 0.8893 | -3.78E-06 | 4.34E-06 |
| | | THCfemale | 1.93E-06 | 0.1978 | -1.34E-06 | 6.36E-06 |

| | | | | | |
|---|---|---|---|---|---|
| | TpCfemale | 1.89E-06 | 0.5339 | -2.59E-06 | 4.95E-06 |
| | VehMale | 2.03E-06 | 0.0207 | -8.88E-06 | -7.59E-07 |
| | CBDmale | 1.93E-06 | 0.0437 | -7.82E-06 | -1.16E-07 |
| | THCmale | 1.98E-06 | 0.1479 | -1.05E-06 | 6.84E-06 |
| | TpCmale | 1.93E-06 | 0.7673 | -4.42E-06 | 3.28E-06 |
| THCfemale | VehFemale | 1.93E-06 | 0.2528 | -6.08E-06 | 1.63E-06 |
| | CBDfemale | 1.93E-06 | 0.1978 | -6.36E-06 | 1.34E-06 |
| | TpCfemale | 1.78E-06 | 0.4575 | -4.88E-06 | 2.22E-06 |
| | VehMale | 1.93E-06 | 0.0003 | -1.12E-05 | -3.48E-06 |
| | CBDmale | 1.82E-06 | 0.0007 | -1.01E-05 | -2.85E-06 |
| | THCmale | 1.87E-06 | 0.8379 | -3.35E-06 | 4.11E-06 |
| | TpCmale | 1.82E-06 | 0.0948 | -6.71E-06 | 5.49E-07 |
| TpCfemale | VehFemale | 1.89E-06 | 0.6364 | -4.67E-06 | 2.88E-06 |
| | CBDfemale | 1.89E-06 | 0.5339 | -4.95E-06 | 2.59E-06 |
| | THCfemale | 1.78E-06 | 0.4575 | -2.22E-06 | 4.88E-06 |
| | VehMale | 1.89E-06 | 0.0023 | -9.77E-06 | -2.23E-06 |
| | CBDmale | 1.78E-06 | 0.0051 | -8.70E-06 | -1.60E-06 |
| | THCmale | 1.83E-06 | 0.3525 | -1.94E-06 | 5.36E-06 |
| | TpCmale | 1.78E-06 | 0.3270 | -5.30E-06 | 1.79E-06 |
| VehMale | VehFemale | 2.03E-06 | 0.0145 | 1.04E-06 | 9.16E-06 |
| | CBDfemale | 2.03E-06 | 0.0207 | 7.59E-07 | 8.88E-06 |
| | THCfemale | 1.93E-06 | 0.0003 | 3.48E-06 | 1.12E-05 |
| | TpCfemale | 1.89E-06 | 0.0023 | 2.23E-06 | 9.77E-06 |
| | CBDmale | 1.93E-06 | 0.6602 | -3.00E-06 | 4.70E-06 |
| | THCmale | 1.98E-06 | 0.0002 | 3.77E-06 | 1.17E-05 |
| | TpCmale | 1.93E-06 | 0.0312 | 3.95E-07 | 8.10E-06 |
| CBDmale | VehFemale | 1.93E-06 | 0.0310 | 4.00E-07 | 8.10E-06 |
| | CBDfemale | 1.93E-06 | 0.0437 | 1.16E-07 | 7.82E-06 |
| | THCfemale | 1.82E-06 | 0.0007 | 2.85E-06 | 1.01E-05 |
| | TpCfemale | 1.78E-06 | 0.0051 | 1.60E-06 | 8.70E-06 |
| | VehMale | 1.93E-06 | 0.6602 | -4.70E-06 | 3.00E-06 |
| | THCmale | 1.87E-06 | 0.0005 | 3.13E-06 | 1.06E-05 |
| | TpCmale | 1.82E-06 | 0.0664 | -2.37E-07 | 7.03E-06 |
| THCmale | VehFemale | 1.98E-06 | 0.1913 | -6.56E-06 | 1.34E-06 |
| | CBDfemale | 1.98E-06 | 0.1479 | -6.84E-06 | 1.05E-06 |
| | THCfemale | 1.87E-06 | 0.8379 | -4.11E-06 | 3.35E-06 |
| | TpCfemale | 1.83E-06 | 0.3525 | -5.36E-06 | 1.94E-06 |
| | VehMale | 1.98E-06 | 0.0002 | -1.17E-05 | -3.77E-06 |
| | CBDmale | 1.87E-06 | 0.0005 | -1.06E-05 | -3.13E-06 |
| | TpCmale | 1.87E-06 | 0.0680 | -7.20E-06 | 2.64E-07 |
| TpCmale | VehFemale | 1.93E-06 | 0.6582 | -2.99E-06 | 4.71E-06 |
| | CBDfemale | 1.93E-06 | 0.7673 | -3.28E-06 | 4.42E-06 |
| | THCfemale | 1.82E-06 | 0.0948 | -5.49E-07 | 6.71E-06 |
| | TpCfemale | 1.78E-06 | 0.3270 | -1.79E-06 | 5.30E-06 |
| | VehMale | 1.93E-06 | 0.0312 | -8.10E-06 | -3.95E-07 |
| | CBDmale | 1.82E-06 | 0.0664 | -7.03E-06 | 2.37E-07 |
| | THCmale | 1.87E-06 | 0.0680 | -2.64E-07 | 7.20E-06 |

#### Supplemental Table 4.8 Fisher's LSD post-hoc test for liver 2-acyl glycerol ANOVA

| | | | | | 95% Confidence Interval | |
|---|---|---|---|---|---|---|
| | | | Std. Error | Sig. | Lower Bound | Upper Bound |
| 2-palmitoyl glycerol | VehFemale | CBDfemale | 5.85919E-10 | 0.783 | -1.3335E-09 | 1.0090E-09 |
| | | THCfemale | 5.50103E-10 | 0.087 | -2.0549E-09 | 1.4434E-10 |
| | | TpCfemale | 5.37004E-10 | 0.002 | -2.8224E-09 | -6.7548E-10 |
| | | VehMale | 5.85919E-10 | 0.436 | -1.6308E-09 | 7.1165E-10 |
| | | CBDmale | 5.50103E-10 | 0.057 | -2.1662E-09 | 3.3070E-11 |
| | | THCmale | 5.50103E-10 | 0.008 | -2.6081E-09 | -4.0882E-10 |
| | | TpCmale | 5.26043E-10 | 0.020 | -2.3050E-09 | -2.0190E-10 |
| | CBDfemale | VehFemale | 5.85919E-10 | 0.783 | -1.0090E-09 | 1.3335E-09 |
| | | THCfemale | 5.70526E-10 | 0.169 | -1.9335E-09 | 3.4741E-10 |
| | | TpCfemale | 5.57907E-10 | 0.006 | -2.7019E-09 | -4.7145E-10 |
| | | VehMale | 6.05135E-10 | 0.625 | -1.5070E-09 | 9.1231E-10 |
| | | CBDmale | 5.70526E-10 | 0.118 | -2.0448E-09 | 2.3614E-10 |
| | | THCmale | 5.70526E-10 | 0.021 | -2.4867E-09 | -2.0575E-10 |
| | | TpCmale | 5.47365E-10 | 0.051 | -2.1854E-09 | 2.9712E-12 |
| | THCfemale | VehFemale | 5.50103E-10 | 0.087 | -1.4434E-10 | 2.0549E-09 |
| | | CBDfemale | 5.70526E-10 | 0.169 | -3.4741E-10 | 1.9335E-09 |
| | | TpCfemale | 5.20166E-10 | 0.132 | -1.8334E-09 | 2.4616E-10 |
| | | VehMale | 5.70526E-10 | 0.388 | -6.4475E-10 | 1.6362E-09 |
| | | CBDmale | 5.33679E-10 | 0.836 | -1.1781E-09 | 9.5554E-10 |
| | | THCmale | 5.33679E-10 | 0.304 | -1.6200E-09 | 5.1365E-10 |
| | | TpCmale | 5.08843E-10 | 0.560 | -1.3153E-09 | 7.1902E-10 |
| | TpCfemale | VehFemale | 5.37004E-10 | 0.002 | 6.7548E-10 | 2.8224E-09 |
| | | CBDfemale | 5.57907E-10 | 0.006 | 4.7145E-10 | 2.7019E-09 |
| | | THCfemale | 5.20166E-10 | 0.132 | -2.4616E-10 | 1.8334E-09 |
| | | VehMale | 5.57907E-10 | 0.024 | 1.7411E-10 | 2.4046E-09 |
| | | CBDmale | 5.20166E-10 | 0.194 | -3.5743E-10 | 1.7222E-09 |
| | | THCmale | 5.20166E-10 | 0.645 | -7.9932E-10 | 1.2803E-09 |
| | | TpCmale | 4.94652E-10 | 0.320 | -4.9330E-10 | 1.4843E-09 |
| | VehMale | VehFemale | 5.85919E-10 | 0.436 | -7.1165E-10 | 1.6308E-09 |
| | | CBDfemale | 6.05135E-10 | 0.625 | -9.1231E-10 | 1.5070E-09 |

| | | | | | | |
|---|---|---|---|---|---|---|
| | | THCfemale | 5.70526E-10 | 0.388 | -1.6362E-09 | 6.4475E-10 |
| | | TpCfemale | 5.57907E-10 | 0.024 | -2.4046E-09 | -1.7411E-10 |
| | | CBDmale | 5.70526E-10 | 0.292 | -1.7474E-09 | 5.3348E-10 |
| | | THCmale | 5.70526E-10 | 0.071 | -2.1893E-09 | 9.1591E-11 |
| | | TpCmale | 5.47365E-10 | 0.152 | -1.8880E-09 | 3.0031E-10 |
| | CBDmale | VehFemale | 5.50103E-10 | 0.057 | -3.3070E-11 | 2.1662E-09 |
| | | CBDfemale | 5.70526E-10 | 0.118 | -2.3614E-10 | 2.0448E-09 |
| | | THCfemale | 5.33679E-10 | 0.836 | -9.5554E-10 | 1.1781E-09 |
| | | TpCfemale | 5.20166E-10 | 0.194 | -1.7222E-09 | 3.5743E-10 |
| | | VehMale | 5.70526E-10 | 0.292 | -5.3348E-10 | 1.7474E-09 |
| | | THCmale | 5.33679E-10 | 0.411 | -1.5087E-09 | 6.2492E-10 |
| | | TpCmale | 5.08843E-10 | 0.715 | -1.2040E-09 | 8.3029E-10 |
| | THCmale | VehFemale | 5.50103E-10 | 0.008 | 4.0882E-10 | 2.6081E-09 |
| | | CBDfemale | 5.70526E-10 | 0.021 | 2.0575E-10 | 2.4867E-09 |
| | | THCfemale | 5.33679E-10 | 0.304 | -5.1365E-10 | 1.6200E-09 |
| | | TpCfemale | 5.20166E-10 | 0.645 | -1.2803E-09 | 7.9932E-10 |
| | | VehMale | 5.70526E-10 | 0.071 | -9.1591E-11 | 2.1893E-09 |
| | | CBDmale | 5.33679E-10 | 0.411 | -6.2492E-10 | 1.5087E-09 |
| | | TpCmale | 5.08843E-10 | 0.618 | -7.6214E-10 | 1.2722E-09 |
| | TpCmale | VehFemale | 5.26043E-10 | 0.020 | 2.0190E-10 | 2.3050E-09 |
| | | CBDfemale | 5.47365E-10 | 0.051 | -2.9712E-12 | 2.1854E-09 |
| | | THCfemale | 5.08843E-10 | 0.560 | -7.1902E-10 | 1.3153E-09 |
| | | TpCfemale | 4.94652E-10 | 0.320 | -1.4843E-09 | 4.9330E-10 |
| | | VehMale | 5.47365E-10 | 0.152 | -3.0031E-10 | 1.8880E-09 |
| | | CBDmale | 5.08843E-10 | 0.715 | -8.3029E-10 | 1.2040E-09 |
| | | THCmale | 5.08843E-10 | 0.618 | -1.2722E-09 | 7.6214E-10 |
| | VehFemale | CBDfemale | 5.93637E-09 | 0.888 | -1.2704E-08 | 1.1022E-08 |
| | | THCfemale | 5.63174E-09 | 0.000 | -3.4387E-08 | -1.1879E-08 |
| | | TpCfemale | 5.63174E-09 | 0.000 | -3.7843E-08 | -1.5335E-08 |
| 2-oleoyl glycerol | | VehMale | 6.14473E-09 | 0.024 | -2.6499E-08 | -1.9408E-09 |
| | | CBDmale | 5.63174E-09 | 0.005 | -2.7610E-08 | -5.1014E-09 |
| | | THCmale | 5.93637E-09 | 0.003 | -3.0534E-08 | -6.8086E-09 |

| | | | | | |
|---|---|---|---|---|---|
| | TpCmale | 5.63174E-09 | 0.000 | -3.5167E-08 | -1.2659E-08 |
| CBDfemale | VehFemale | 5.93637E-09 | 0.888 | -1.1022E-08 | 1.2704E-08 |
| | THCfemale | 5.63174E-09 | 0.000 | -3.3547E-08 | -1.1038E-08 |
| | TpCfemale | 5.63174E-09 | 0.000 | -3.7002E-08 | -1.4494E-08 |
| | VehMale | 6.14473E-09 | 0.033 | -2.5659E-08 | -1.1002E-09 |
| | CBDmale | 5.63174E-09 | 0.008 | -2.6769E-08 | -4.2607E-09 |
| | THCmale | 5.93637E-09 | 0.004 | -2.9694E-08 | -5.9679E-09 |
| | TpCmale | 5.63174E-09 | 0.000 | -3.4327E-08 | -1.1819E-08 |
| THCfemale | VehFemale | 5.63174E-09 | 0.000 | 1.1879E-08 | 3.4387E-08 |
| | CBDfemale | 5.63174E-09 | 0.000 | 1.1038E-08 | 3.3547E-08 |
| | TpCfemale | 5.30965E-09 | 0.517 | -1.4066E-08 | 7.1546E-09 |
| | VehMale | 5.85095E-09 | 0.133 | -2.7792E-09 | 2.0605E-08 |
| | CBDmale | 5.30965E-09 | 0.206 | -3.8329E-09 | 1.7388E-08 |
| | THCmale | 5.63174E-09 | 0.431 | -6.7925E-09 | 1.5716E-08 |
| | TpCmale | 5.30965E-09 | 0.884 | -1.1391E-08 | 9.8302E-09 |
| TpCfemale | VehFemale | 5.63174E-09 | 0.000 | 1.5335E-08 | 3.7843E-08 |
| | CBDfemale | 5.63174E-09 | 0.000 | 1.4494E-08 | 3.7002E-08 |
| | THCfemale | 5.30965E-09 | 0.517 | -7.1546E-09 | 1.4066E-08 |
| | VehMale | 5.85095E-09 | 0.038 | 6.7667E-10 | 2.4061E-08 |
| | CBDmale | 5.30965E-09 | 0.058 | -3.7703E-10 | 2.0844E-08 |
| | THCmale | 5.63174E-09 | 0.165 | -3.3366E-09 | 1.9172E-08 |
| | TpCmale | 5.30965E-09 | 0.616 | -7.9349E-09 | 1.3286E-08 |
| VehMale | VehFemale | 6.14473E-09 | 0.024 | 1.9408E-09 | 2.6499E-08 |
| | CBDfemale | 6.14473E-09 | 0.033 | 1.1002E-09 | 2.5659E-08 |
| | THCfemale | 5.85095E-09 | 0.133 | -2.0605E-08 | 2.7792E-09 |
| | TpCfemale | 5.85095E-09 | 0.038 | -2.4061E-08 | -6.7667E-10 |
| | CBDmale | 5.85095E-09 | 0.716 | -1.3828E-08 | 9.5568E-09 |
| | THCmale | 6.14473E-09 | 0.471 | -1.6731E-08 | 7.8279E-09 |
| | TpCmale | 5.85095E-09 | 0.103 | -2.1385E-08 | 1.9990E-09 |
| CBDmale | VehFemale | 5.63174E-09 | 0.005 | 5.1014E-09 | 2.7610E-08 |
| | CBDfemale | 5.63174E-09 | 0.008 | 4.2607E-09 | 2.6769E-08 |
| | THCfemale | 5.30965E-09 | 0.206 | -1.7388E-08 | 3.8329E-09 |

| | | | | | | |
|---|---|---|---|---|---|---|
| | | TpCfemale | 5.30965E-09 | 0.058 | -2.0844E-08 | 3.7703E-10 |
| | | VehMale | 5.85095E-09 | 0.716 | -9.5568E-09 | 1.3828E-08 |
| | | THCmale | 5.63174E-09 | 0.682 | -1.3570E-08 | 8.9382E-09 |
| | | TpCmale | 5.30965E-09 | 0.160 | -1.8168E-08 | 3.0527E-09 |
| | THCmale | VehFemale | 5.93637E-09 | 0.003 | 6.8086E-09 | 3.0534E-08 |
| | | CBDfemale | 5.93637E-09 | 0.004 | 5.9679E-09 | 2.9694E-08 |
| | | THCfemale | 5.63174E-09 | 0.431 | -1.5716E-08 | 6.7925E-09 |
| | | TpCfemale | 5.63174E-09 | 0.165 | -1.9172E-08 | 3.3366E-09 |
| | | VehMale | 6.14473E-09 | 0.471 | -7.8279E-09 | 1.6731E-08 |
| | | CBDmale | 5.63174E-09 | 0.682 | -8.9382E-09 | 1.3570E-08 |
| | | TpCmale | 5.63174E-09 | 0.356 | -1.6496E-08 | 6.0122E-09 |
| | TpCmale | VehFemale | 5.63174E-09 | 0.000 | 1.2659E-08 | 3.5167E-08 |
| | | CBDfemale | 5.63174E-09 | 0.000 | 1.1819E-08 | 3.4327E-08 |
| | | THCfemale | 5.30965E-09 | 0.884 | -9.8302E-09 | 1.1391E-08 |
| | | TpCfemale | 5.30965E-09 | 0.616 | -1.3286E-08 | 7.9349E-09 |
| | | VehMale | 5.85095E-09 | 0.103 | -1.9990E-09 | 2.1385E-08 |
| | | CBDmale | 5.30965E-09 | 0.160 | -3.0527E-09 | 1.8168E-08 |
| | | THCmale | 5.63174E-09 | 0.356 | -6.0122E-09 | 1.6496E-08 |
| 2-linoleoyl glycerol | VehFemale | CBDfemale | 4.72201E-09 | 0.774 | -1.0797E-08 | 8.0749E-09 |
| | | THCfemale | 4.58897E-09 | 0.058 | -1.8017E-08 | 3.2381E-10 |
| | | TpCfemale | 4.47969E-09 | 0.000 | -3.1917E-08 | -1.4013E-08 |
| | | VehMale | 4.88775E-09 | 0.108 | -1.7734E-08 | 1.8009E-09 |
| | | CBDmale | 4.58897E-09 | 0.008 | -2.1833E-08 | -3.4924E-09 |
| | | THCmale | 4.58897E-09 | 0.000 | -2.9093E-08 | -1.0753E-08 |
| | | TpCmale | 4.38826E-09 | 0.000 | -2.4999E-08 | -7.4605E-09 |
| | CBDfemale | VehFemale | 4.72201E-09 | 0.774 | -8.0749E-09 | 1.0797E-08 |
| | | THCfemale | 4.58897E-09 | 0.108 | -1.6656E-08 | 1.6851E-09 |
| | | TpCfemale | 4.47969E-09 | 0.000 | -3.0556E-08 | -1.2652E-08 |
| | | VehMale | 4.88775E-09 | 0.181 | -1.6373E-08 | 3.1622E-09 |
| | | CBDmale | 4.58897E-09 | 0.017 | -2.0472E-08 | -2.1312E-09 |
| | | THCmale | 4.58897E-09 | 0.000 | -2.7732E-08 | -9.3914E-09 |
| | | TpCmale | 4.38826E-09 | 0.001 | -2.3638E-08 | -6.0992E-09 |

| | | | | | |
|---|---|---|---|---|---|
| THCfemale | VehFemale | 4.58897E-09 | 0.058 | -3.2381E-10 | 1.8017E-08 |
| | CBDfemale | 4.58897E-09 | 0.108 | -1.6851E-09 | 1.6656E-08 |
| | TpCfemale | 4.33923E-09 | 0.002 | -2.2790E-08 | -5.4475E-09 |
| | VehMale | 4.75934E-09 | 0.854 | -8.6308E-09 | 1.0391E-08 |
| | CBDmale | 4.45196E-09 | 0.395 | -1.2713E-08 | 5.0803E-09 |
| | THCmale | 4.45196E-09 | 0.016 | -1.9973E-08 | -2.1800E-09 |
| | TpCmale | 4.24477E-09 | 0.087 | -1.5866E-08 | 1.0993E-09 |
| TpCfemale | VehFemale | 4.47969E-09 | 0.000 | 1.4013E-08 | 3.1917E-08 |
| | CBDfemale | 4.47969E-09 | 0.000 | 1.2652E-08 | 3.0556E-08 |
| | THCfemale | 4.33923E-09 | 0.002 | 5.4475E-09 | 2.2790E-08 |
| | VehMale | 4.65407E-09 | 0.002 | 5.6984E-09 | 2.4299E-08 |
| | CBDmale | 4.33923E-09 | 0.021 | 1.6313E-09 | 1.8974E-08 |
| | THCmale | 4.33923E-09 | 0.486 | -5.6290E-09 | 1.1714E-08 |
| | TpCmale | 4.12639E-09 | 0.108 | -1.5104E-09 | 1.4982E-08 |
| VehMale | VehFemale | 4.88775E-09 | 0.108 | -1.8009E-09 | 1.7734E-08 |
| | CBDfemale | 4.88775E-09 | 0.181 | -3.1622E-09 | 1.6373E-08 |
| | THCfemale | 4.75934E-09 | 0.854 | -1.0391E-08 | 8.6308E-09 |
| | TpCfemale | 4.65407E-09 | 0.002 | -2.4299E-08 | -5.6984E-09 |
| | CBDmale | 4.75934E-09 | 0.328 | -1.4207E-08 | 4.8145E-09 |
| | THCmale | 4.75934E-09 | 0.015 | -2.1467E-08 | -2.4457E-09 |
| | TpCmale | 4.56613E-09 | 0.075 | -1.7388E-08 | 8.6145E-10 |
| CBDmale | VehFemale | 4.58897E-09 | 0.008 | 3.4924E-09 | 2.1833E-08 |
| | CBDfemale | 4.58897E-09 | 0.017 | 2.1312E-09 | 2.0472E-08 |
| | THCfemale | 4.45196E-09 | 0.395 | -5.0803E-09 | 1.2713E-08 |
| | TpCfemale | 4.33923E-09 | 0.021 | -1.8974E-08 | -1.6313E-09 |
| | VehMale | 4.75934E-09 | 0.328 | -4.8145E-09 | 1.4207E-08 |
| | THCmale | 4.45196E-09 | 0.108 | -1.6157E-08 | 1.6363E-09 |
| | TpCmale | 4.24477E-09 | 0.404 | -1.2049E-08 | 4.9156E-09 |
| THCmale | VehFemale | 4.58897E-09 | 0.000 | 1.0753E-08 | 2.9093E-08 |
| | CBDfemale | 4.58897E-09 | 0.000 | 9.3914E-09 | 2.7732E-08 |
| | THCfemale | 4.45196E-09 | 0.016 | 2.1800E-09 | 1.9973E-08 |
| | TpCfemale | 4.33923E-09 | 0.486 | -1.1714E-08 | 5.6290E-09 |

| | | | | | | |
|---|---|---|---|---|---|---|
| | | VehMale | 4.75934E-09 | 0.015 | 2.4457E-09 | 2.1467E-08 |
| | | CBDmale | 4.45196E-09 | 0.108 | -1.6363E-09 | 1.6157E-08 |
| | | TpCmale | 4.24477E-09 | 0.388 | -4.7892E-09 | 1.2176E-08 |
| | TpCmale | VehFemale | 4.38826E-09 | 0.000 | 7.4605E-09 | 2.4999E-08 |
| | | CBDfemale | 4.38826E-09 | 0.001 | 6.0992E-09 | 2.3638E-08 |
| | | THCfemale | 4.24477E-09 | 0.087 | -1.0993E-09 | 1.5866E-08 |
| | | TpCfemale | 4.12639E-09 | 0.108 | -1.4982E-08 | 1.5104E-09 |
| | | VehMale | 4.56613E-09 | 0.075 | -8.6145E-10 | 1.7388E-08 |
| | | CBDmale | 4.24477E-09 | 0.404 | -4.9156E-09 | 1.2049E-08 |
| | | THCmale | 4.24477E-09 | 0.388 | -1.2176E-08 | 4.7892E-09 |
| | VehFemale | CBDfemale | 4.39066E-09 | 0.506 | -1.1709E-08 | 5.8392E-09 |
| | | THCfemale | 4.18075E-09 | 0.000 | -2.4431E-08 | -7.7222E-09 |
| | | TpCfemale | 4.18075E-09 | 0.000 | -3.1356E-08 | -1.4647E-08 |
| | | VehMale | 4.53466E-09 | 0.013 | -2.0593E-08 | -2.4689E-09 |
| | | CBDmale | 4.27532E-09 | 0.003 | -2.1592E-08 | -4.5050E-09 |
| | | THCmale | 4.27532E-09 | 0.000 | -2.6428E-08 | -9.3412E-09 |
| | | TpCmale | 4.10175E-09 | 0.002 | -2.1503E-08 | -5.1101E-09 |
| | CBDfemale | VehFemale | 4.39066E-09 | 0.506 | -5.8392E-09 | 1.1709E-08 |
| | | THCfemale | 4.02411E-09 | 0.002 | -2.1183E-08 | -5.1004E-09 |
| | | TpCfemale | 4.02411E-09 | 0.000 | -2.8108E-08 | -1.2025E-08 |
| | | VehMale | 4.39066E-09 | 0.055 | -1.7370E-08 | 1.7816E-10 |
| 2-arachidonoyl glycerol | | CBDmale | 4.12227E-09 | 0.017 | -1.8351E-08 | -1.8760E-09 |
| | | THCmale | 4.12227E-09 | 0.001 | -2.3188E-08 | -6.7122E-09 |
| | | TpCmale | 3.94198E-09 | 0.011 | -1.8249E-08 | -2.4945E-09 |
| | THCfemale | VehFemale | 4.18075E-09 | 0.000 | 7.7222E-09 | 2.4431E-08 |
| | | CBDfemale | 4.02411E-09 | 0.002 | 5.1004E-09 | 2.1183E-08 |
| | | TpCfemale | 3.79397E-09 | 0.073 | -1.4507E-08 | 6.5664E-10 |
| | | VehMale | 4.18075E-09 | 0.281 | -3.8085E-09 | 1.2901E-08 |
| | | CBDmale | 3.89793E-09 | 0.440 | -4.7612E-09 | 1.0818E-08 |
| | | THCmale | 3.89793E-09 | 0.644 | -9.5974E-09 | 5.9814E-09 |
| | | TpCmale | 3.70674E-09 | 0.458 | -4.6373E-09 | 1.0177E-08 |
| | TpCfemale | VehFemale | 4.18075E-09 | 0.000 | 1.4647E-08 | 3.1356E-08 |

| | | | | | |
|---|---|---|---|---|---|
| | CBDfemale | 4.02411E-09 | 0.000 | 1.2025E-08 | 2.8108E-08 |
| | THCfemale | 3.79397E-09 | 0.073 | -6.5664E-10 | 1.4507E-08 |
| | VehMale | 4.18075E-09 | 0.008 | 3.1165E-09 | 1.9826E-08 |
| | CBDmale | 3.89793E-09 | 0.013 | 2.1638E-09 | 1.7743E-08 |
| | THCmale | 3.89793E-09 | 0.194 | -2.6724E-09 | 1.2906E-08 |
| | TpCmale | 3.70674E-09 | 0.011 | 2.2877E-09 | 1.7102E-08 |
| VehMale | VehFemale | 4.53466E-09 | 0.013 | 2.4689E-09 | 2.0593E-08 |
| | CBDfemale | 4.39066E-09 | 0.055 | -1.7816E-10 | 1.7370E-08 |
| | THCfemale | 4.18075E-09 | 0.281 | -1.2901E-08 | 3.8085E-09 |
| | TpCfemale | 4.18075E-09 | 0.008 | -1.9826E-08 | -3.1165E-09 |
| | CBDmale | 4.27532E-09 | 0.724 | -1.0061E-08 | 7.0257E-09 |
| | THCmale | 4.27532E-09 | 0.142 | -1.4898E-08 | 2.1895E-09 |
| | TpCmale | 4.10175E-09 | 0.666 | -9.9727E-09 | 6.4207E-09 |
| CBDmale | VehFemale | 4.27532E-09 | 0.003 | 4.5050E-09 | 2.1592E-08 |
| | CBDfemale | 4.12227E-09 | 0.017 | 1.8760E-09 | 1.8351E-08 |
| | THCfemale | 3.89793E-09 | 0.440 | -1.0818E-08 | 4.7612E-09 |
| | TpCfemale | 3.89793E-09 | 0.013 | -1.7743E-08 | -2.1638E-09 |
| | VehMale | 4.27532E-09 | 0.724 | -7.0257E-09 | 1.0061E-08 |
| | THCmale | 3.99919E-09 | 0.231 | -1.2828E-08 | 3.1556E-09 |
| | TpCmale | 3.81308E-09 | 0.946 | -7.8781E-09 | 7.3616E-09 |
| THCmale | VehFemale | 4.27532E-09 | 0.000 | 9.3412E-09 | 2.6428E-08 |
| | CBDfemale | 4.12227E-09 | 0.001 | 6.7122E-09 | 2.3188E-08 |
| | THCfemale | 3.89793E-09 | 0.644 | -5.9814E-09 | 9.5974E-09 |
| | TpCfemale | 3.89793E-09 | 0.194 | -1.2906E-08 | 2.6724E-09 |
| | VehMale | 4.27532E-09 | 0.142 | -2.1895E-09 | 1.4898E-08 |
| | CBDmale | 3.99919E-09 | 0.231 | -3.1556E-09 | 1.2828E-08 |
| | TpCmale | 3.81308E-09 | 0.234 | -3.0419E-09 | 1.2198E-08 |
| TpCmale | VehFemale | 4.10175E-09 | 0.002 | 5.1101E-09 | 2.1503E-08 |
| | CBDfemale | 3.94198E-09 | 0.011 | 2.4945E-09 | 1.8249E-08 |
| | THCfemale | 3.70674E-09 | 0.458 | -1.0177E-08 | 4.6373E-09 |
| | TpCfemale | 3.70674E-09 | 0.011 | -1.7102E-08 | -2.2877E-09 |
| | VehMale | 4.10175E-09 | 0.666 | -6.4207E-09 | 9.9727E-09 |

| | | | | | |
|---|---|---|---|---|---|
| | CBDmale | 3.81308E-09 | 0.946 | -7.3616E-09 | 7.8781E-09 |
| | THCmale | 3.81308E-09 | 0.234 | -1.2198E-08 | 3.0419E-09 |

Supplemental Table 5.1 Descriptive statistics for free fatty acids (mol/L) in plasma collected from male and female adult mice 2h after receiving an i.p. injection of 10mg/kg CBD, THC, or THC+CBD

| | | N | Mean | Std. Deviation | Std. Error | 95% Confidence Interval for Mean | | Minimum | Maximum |
|---|---|---|---|---|---|---|---|---|---|
| | | | | | | Lower Bound | Upper Bound | | |
| Oleic acid | Veh F | 7 | 1.1345E-08 | 1.34335E-09 | 5.07740E-10 | 1.0102E-08 | 1.2587E-08 | 9.30E-09 | 1.30E-08 |
| | CBD F | 8 | 1.1673E-08 | 4.12417E-09 | 1.45811E-09 | 8.2252E-09 | 1.5121E-08 | 6.86E-09 | 1.95E-08 |
| | THC F | 10 | 1.2646E-08 | 3.30983E-09 | 1.04666E-09 | 1.0279E-08 | 1.5014E-08 | 8.60E-09 | 1.75E-08 |
| | THC+CBD F | 10 | 1.1687E-08 | 2.03697E-09 | 6.44148E-10 | 1.0230E-08 | 1.3144E-08 | 8.97E-09 | 1.46E-08 |
| | Veh M | 8 | 1.3188E-08 | 6.17394E-09 | 2.18282E-09 | 8.0265E-09 | 1.8350E-08 | 6.13E-09 | 2.20E-08 |
| | CBD M | 11 | 1.5242E-08 | 7.73395E-09 | 2.33187E-09 | 1.0046E-08 | 2.0438E-08 | 5.79E-09 | 2.90E-08 |
| | THC M | 9 | 1.0403E-08 | 4.39515E-09 | 1.46505E-09 | 7.0246E-09 | 1.3781E-08 | 4.62E-09 | 1.75E-08 |
| | THC+CBD M | 10 | 1.4424E-08 | 5.62128E-09 | 1.77761E-09 | 1.0403E-08 | 1.8445E-08 | 4.78E-09 | 2.18E-08 |
| Linoleic acid | Veh F | 7 | 2.1456E-08 | 3.69340E-09 | 1.39597E-09 | 1.8040E-08 | 2.4872E-08 | 1.62E-08 | 2.69E-08 |
| | CBD F | 9 | 2.0457E-08 | 1.07722E-08 | 3.59072E-09 | 1.2177E-08 | 2.8737E-08 | 1.00E-08 | 3.93E-08 |
| | THC F | 10 | 1.5462E-08 | 3.79085E-09 | 1.19877E-09 | 1.2751E-08 | 1.8174E-08 | 9.85E-09 | 1.92E-08 |
| | THC+CBD F | 11 | 1.9123E-08 | 4.90707E-09 | 1.47954E-09 | 1.5827E-08 | 2.2420E-08 | 1.03E-08 | 2.50E-08 |
| | Veh M | 8 | 1.7321E-08 | 3.09462E-09 | 1.09411E-09 | 1.4734E-08 | 1.9908E-08 | 1.33E-08 | 2.18E-08 |
| | CBD M | 11 | 2.0904E-08 | 6.05113E-09 | 1.82448E-09 | 1.6839E-08 | 2.4970E-08 | 1.43E-08 | 3.23E-08 |
| | THC M | 9 | 1.7335E-08 | 6.44419E-09 | 2.14806E-09 | 1.2382E-08 | 2.2288E-08 | 7.09E-09 | 2.82E-08 |
| | THC+CBD M | 10 | 1.8054E-08 | 5.24000E-09 | 1.65703E-09 | 1.4305E-08 | 2.1802E-08 | 9.74E-09 | 2.63E-08 |
| Arachidonic acid | Veh F | 7 | 1.1310E-08 | 2.19855E-09 | 8.30972E-10 | 9.2767E-09 | 1.3343E-08 | 8.93E-09 | 1.44E-08 |
| | CBD F | 8 | 9.6424E-09 | 3.50476E-09 | 1.23912E-09 | 6.7123E-09 | 1.2572E-08 | 5.56E-09 | 1.51E-08 |
| | THC F | 9 | 1.4046E-08 | 2.75936E-09 | 9.19788E-10 | 1.1925E-08 | 1.6167E-08 | 8.83E-09 | 1.78E-08 |
| | THC+CBD F | 11 | 1.5018E-08 | 5.96448E-09 | 1.79836E-09 | 1.1011E-08 | 1.9025E-08 | 6.85E-09 | 2.44E-08 |
| | Veh M | 8 | 1.2597E-08 | 2.26373E-09 | 8.00351E-10 | 1.0705E-08 | 1.4490E-08 | 9.28E-09 | 1.56E-08 |
| | CBD M | 11 | 1.3073E-08 | 6.55932E-09 | 1.97771E-09 | 8.6666E-09 | 1.7480E-08 | 6.00E-09 | 2.54E-08 |
| | THC M | 9 | 1.0969E-08 | 5.97759E-09 | 1.99253E-09 | 6.3744E-09 | 1.5564E-08 | 3.22E-09 | 2.23E-08 |
| | THC+CBD M | 9 | 1.0369E-08 | 3.41759E-09 | 1.13920E-09 | 7.7420E-09 | 1.2996E-08 | 6.68E-09 | 1.66E-08 |
| Docosahexaenoic acid | Veh F | 7 | 4.9316E-08 | 1.34631E-08 | 5.08856E-09 | 3.6865E-08 | 6.1767E-08 | 2.67E-08 | 6.28E-08 |
| | CBD F | 9 | 3.2624E-08 | 1.58340E-08 | 5.27799E-09 | 2.0453E-08 | 4.4795E-08 | 1.78E-08 | 5.71E-08 |
| | THC F | 9 | 4.4969E-08 | 1.64024E-08 | 5.46745E-09 | 3.2362E-08 | 5.7577E-08 | 2.02E-08 | 6.95E-08 |

| | | | | | | | | | |
|---|---|---|---|---|---|---|---|---|---|
| | THC+CBD F | 10 | 5.2835E-08 | 1.77762E-08 | 5.62132E-09 | 4.0119E-08 | 6.5552E-08 | 2.03E-08 | 7.54E-08 |
| | Veh M | 7 | 5.4424E-08 | 1.45919E-08 | 5.51523E-09 | 4.0929E-08 | 6.7920E-08 | 3.86E-08 | 7.47E-08 |
| | CBD M | 9 | 4.1067E-08 | 6.58537E-09 | 2.19512E-09 | 3.6005E-08 | 4.6129E-08 | 3.10E-08 | 4.88E-08 |
| | THC M | 8 | 4.3087E-08 | 9.92416E-09 | 3.50872E-09 | 3.4791E-08 | 5.1384E-08 | 2.63E-08 | 5.62E-08 |
| | THC+CBD M | 9 | 4.7141E-08 | 1.64557E-08 | 5.48523E-09 | 3.4492E-08 | 5.9790E-08 | 3.41E-08 | 8.85E-08 |
| Eicosapentaenoic acid | Veh F | 7 | 8.7174E-09 | 1.44948E-09 | 5.47851E-10 | 7.3768E-09 | 1.0058E-08 | 6.59E-09 | 1.05E-08 |
| | CBD F | 8 | 7.5958E-09 | 1.38453E-09 | 4.89506E-10 | 6.4383E-09 | 8.7533E-09 | 5.61E-09 | 9.58E-09 |
| | THC F | 9 | 7.2286E-09 | 1.90011E-09 | 6.33368E-10 | 5.7681E-09 | 8.6892E-09 | 4.34E-09 | 1.03E-08 |
| | THC+CBD F | 10 | 8.2107E-09 | 3.18934E-09 | 1.00856E-09 | 5.9292E-09 | 1.0492E-08 | 5.35E-09 | 1.63E-08 |
| | Veh M | 8 | 1.1916E-08 | 3.93813E-09 | 1.39234E-09 | 8.6234E-09 | 1.5208E-08 | 5.43E-09 | 1.70E-08 |
| | CBD M | 11 | 1.4074E-08 | 7.73375E-09 | 2.33181E-09 | 8.8785E-09 | 1.9270E-08 | 5.19E-09 | 2.72E-08 |
| | THC M | 9 | 8.9537E-09 | 5.29835E-09 | 1.76612E-09 | 4.8811E-09 | 1.3026E-08 | 9.80E-10 | 1.74E-08 |
| | THC+CBD M | 10 | 1.3976E-08 | 6.80424E-09 | 2.15169E-09 | 9.1083E-09 | 1.8843E-08 | 5.57E-09 | 2.63E-08 |

**Supplemental Table 5.2 Descriptive statistics for free fatty acids (mol/g) in liver collected from male and female adult mice 2h after receiving an i.p. injection of 10mg/kg CBD, THC, or THC+CBD**

| | | N | Mean | Std. Deviation | Std. Error | 95% Confidence Interval for Mean | | Minimum | Maximum |
|---|---|---|---|---|---|---|---|---|---|
| | | | | | | Lower Bound | Upper Bound | | |
| Oleic acid | Veh F | 8 | 3.7853E-10 | 1.90506E-10 | 6.73541E-11 | 2.1927E-10 | 5.3780E-10 | 1.03E-10 | 6.37E-10 |
| | CBD F | 8 | 4.4804E-10 | 2.85056E-10 | 1.00782E-10 | 2.0973E-10 | 6.8636E-10 | 1.80E-10 | 9.20E-10 |
| | THC F | 9 | 6.7502E-10 | 1.89646E-10 | 6.32152E-11 | 5.2925E-10 | 8.2080E-10 | 3.47E-10 | 9.49E-10 |
| | THC+CBD F | 10 | 7.2876E-10 | 2.86963E-10 | 9.07455E-11 | 5.2348E-10 | 9.3404E-10 | 4.46E-10 | 1.24E-09 |
| | Veh M | 7 | 6.1036E-10 | 3.18064E-10 | 1.20217E-10 | 3.1620E-10 | 9.0452E-10 | 3.55E-10 | 1.07E-09 |
| | CBD M | 10 | 5.8313E-10 | 2.51709E-10 | 7.95975E-11 | 4.0307E-10 | 7.6320E-10 | 1.47E-10 | 9.09E-10 |
| | THC M | 8 | 4.9148E-10 | 2.09328E-10 | 7.40086E-11 | 3.1648E-10 | 6.6648E-10 | 2.07E-10 | 8.14E-10 |
| | THC+CBD M | 10 | 7.0061E-10 | 2.51925E-10 | 7.96656E-11 | 5.2039E-10 | 8.8082E-10 | 3.99E-10 | 1.20E-09 |
| Linoleic acid | Veh F | 8 | 9.9989E-10 | 4.23453E-10 | 1.49713E-10 | 6.4588E-10 | 1.3539E-09 | 4.63E-10 | 1.70E-09 |
| | CBD F | 7 | 8.5520E-10 | 2.69473E-10 | 1.01851E-10 | 6.0598E-10 | 1.1044E-09 | 5.12E-10 | 1.20E-09 |
| | THC F | 10 | 1.5282E-09 | 5.26079E-10 | 1.66361E-10 | 1.1519E-09 | 1.9045E-09 | 7.15E-10 | 2.23E-09 |
| | THC+CBD F | 10 | 2.4151E-09 | 8.07180E-10 | 2.55253E-10 | 1.8377E-09 | 2.9925E-09 | 1.47E-09 | 3.90E-09 |
| | Veh M | 7 | 1.1199E-09 | 4.47337E-10 | 1.69077E-10 | 7.0622E-10 | 1.5337E-09 | 5.34E-10 | 1.75E-09 |
| | CBD M | 10 | 1.1479E-09 | 4.49383E-10 | 1.42108E-10 | 8.2646E-10 | 1.4694E-09 | 5.75E-10 | 1.92E-09 |
| | THC M | 9 | 1.7544E-09 | 8.23086E-10 | 2.74362E-10 | 1.1217E-09 | 2.3871E-09 | 6.56E-10 | 2.83E-09 |
| | THC+CBD M | 10 | 1.9578E-09 | 5.72233E-10 | 1.80956E-10 | 1.5484E-09 | 2.3671E-09 | 1.20E-09 | 3.22E-09 |
| Arachidonic acid | Veh F | 7 | 2.5199E-09 | 9.34892E-10 | 3.53356E-10 | 1.6553E-09 | 3.3846E-09 | 9.44E-10 | 3.66E-09 |
| | CBD F | 8 | 3.9955E-09 | 1.57145E-09 | 5.55592E-10 | 2.6818E-09 | 5.3093E-09 | 2.02E-09 | 6.80E-09 |
| | THC F | 10 | 5.6953E-09 | 1.08131E-09 | 3.41939E-10 | 4.9218E-09 | 6.4688E-09 | 4.48E-09 | 7.61E-09 |
| | THC+CBD F | 10 | 6.5562E-09 | 2.10311E-09 | 6.65063E-10 | 5.0518E-09 | 8.0607E-09 | 4.47E-09 | 1.03E-08 |
| | Veh M | 7 | 5.0634E-09 | 2.08584E-09 | 7.88374E-10 | 3.1344E-09 | 6.9925E-09 | 2.74E-09 | 7.71E-09 |
| | CBD M | 10 | 5.0180E-09 | 1.58239E-09 | 5.00395E-10 | 3.8860E-09 | 6.1499E-09 | 2.72E-09 | 7.38E-09 |
| | THC M | 9 | 6.2097E-09 | 2.11043E-09 | 7.03478E-10 | 4.5875E-09 | 7.8319E-09 | 3.26E-09 | 9.65E-09 |
| | THC+CBD M | 11 | 6.7841E-09 | 1.39418E-09 | 4.20360E-10 | 5.8475E-09 | 7.7207E-09 | 5.50E-09 | 9.53E-09 |
| Docosahexaenoic acid | Veh F | 8 | 1.4605E-09 | 4.57479E-10 | 1.61743E-10 | 1.0780E-09 | 1.8430E-09 | 8.60E-10 | 2.11E-09 |
| | CBD F | 8 | 2.3135E-09 | 8.24989E-10 | 2.91678E-10 | 1.6238E-09 | 3.0032E-09 | 1.05E-09 | 3.52E-09 |
| | THC F | 9 | 2.5809E-09 | 4.94578E-10 | 1.64859E-10 | 2.2008E-09 | 2.9611E-09 | 2.09E-09 | 3.57E-09 |
| | THC+CBD F | 10 | 3.4623E-09 | 7.64638E-10 | 2.41800E-10 | 2.9153E-09 | 4.0093E-09 | 2.41E-09 | 4.66E-09 |
| | Veh M | 6 | 2.0366E-09 | 4.29451E-10 | 1.75323E-10 | 1.5859E-09 | 2.4873E-09 | 1.21E-09 | 2.34E-09 |
| | CBD M | 9 | 2.2404E-09 | 3.46376E-10 | 1.15459E-10 | 1.9741E-09 | 2.5066E-09 | 1.76E-09 | 2.83E-09 |
| | THC M | 8 | 2.9059E-09 | 5.62623E-10 | 1.98917E-10 | 2.4355E-09 | 3.3763E-09 | 1.98E-09 | 3.72E-09 |
| | THC+CBD M | 11 | 3.9769E-09 | 1.32223E-09 | 3.98667E-10 | 3.0886E-09 | 4.8652E-09 | 1.80E-09 | 5.56E-09 |
| Eicosapentaenoic acid | Veh F | 8 | 4.4640E-10 | 1.52569E-10 | 5.39411E-11 | 3.1885E-10 | 5.7395E-10 | 2.97E-10 | 6.66E-10 |
| | CBD F | 7 | 4.1595E-10 | 2.00481E-10 | 7.57746E-11 | 2.3053E-10 | 6.0136E-10 | 1.74E-10 | 8.09E-10 |
| | THC F | 9 | 7.9031E-10 | 1.76884E-10 | 5.89613E-11 | 6.5435E-10 | 9.2628E-10 | 4.96E-10 | 9.73E-10 |
| | THC+CBD F | 10 | 1.2843E-09 | 4.79945E-10 | 1.51772E-10 | 9.4092E-10 | 1.6276E-09 | 6.25E-10 | 1.96E-09 |
| | Veh M | 7 | 9.8819E-10 | 3.18688E-10 | 1.20453E-10 | 6.9345E-10 | 1.2829E-09 | 4.59E-10 | 1.45E-09 |
| | CBD M | 9 | 8.9119E-10 | 2.66696E-10 | 8.88985E-11 | 6.8619E-10 | 1.0962E-09 | 4.19E-10 | 1.22E-09 |
| | THC M | 9 | 1.6436E-09 | 9.01184E-10 | 3.00395E-10 | 9.5092E-10 | 2.3363E-09 | 4.34E-10 | 2.99E-09 |
| | THC+CBD M | 11 | 2.4930E-09 | 9.30057E-10 | 2.80423E-10 | 1.8682E-09 | 3.1179E-09 | 1.35E-09 | 4.01E-09 |

Supplemental Table 5.3 Independent t-test comparing male and female levels of free fatty acids measured in plasma collected from adult mice 2h after an i.p. injection of 10mg/kg CBD, THC, or THC+CBD

| | | | Levene's Test for Equality of Variances | | t-test for Equality of Means | | | | | | |
|---|---|---|---|---|---|---|---|---|---|---|---|
| | | | | | | | Significance | | | 95% Confidence Interval of the Difference | |
| | | | F | Sig. | t | df | One-Sided p | Two-Sided p | Std. Error Difference | Lower | Upper |
| Veh | Oleic acid | Equal variances assumed | 15.993 | 0.002 | -0.771 | 13 | 0.227 | 0.455 | 2.39182E-09 | -7.01064E-09 | 3.32380E-09 |
| | | Equal variances not assumed | | | -0.823 | 7.752 | 0.218 | 0.435 | 2.24109E-09 | -7.04037E-09 | 3.35353E-09 |
| | Linoleic acid | Equal variances assumed | 0.109 | 0.746 | 2.361 | 13 | 0.017 | 0.035 | 1.75147E-09 | 3.50914E-10 | 7.91857E-09 |
| | | Equal variances not assumed | | | 2.331 | 11.814 | 0.019 | 0.038 | 1.77365E-09 | 2.63549E-10 | 8.00594E-09 |
| | Arachidonic acid | Equal variances assumed | 0.008 | 0.931 | -1.114 | 13 | 0.143 | 0.286 | 1.15614E-09 | -3.78511E-09 | 1.21029E-09 |
| | | Equal variances not assumed | | | -1.116 | 12.831 | 0.142 | 0.285 | 1.15372E-09 | -3.78322E-09 | 1.20840E-09 |
| | Docosahexaenoic acid | Equal variances assumed | 0.007 | 0.933 | -0.681 | 12 | 0.254 | 0.509 | 7.50408E-09 | -2.14585E-08 | 1.12414E-08 |
| | | Equal variances not assumed | | | -0.681 | 11.923 | 0.255 | 0.509 | 7.50408E-09 | -2.14702E-08 | 1.12531E-08 |
| | Eicosapentaenoic acid | Equal variances assumed | 6.451 | 0.025 | -2.024 | 13 | 0.032 | 0.064 | 1.58006E-09 | -6.61185E-09 | 2.15186E-10 |
| | | Equal variances not assumed | | | -2.138 | 9.081 | 0.030 | 0.061 | 1.49625E-09 | -6.57846E-09 | 1.81797E-10 |
| CBD | Oleic acid | Equal variances assumed | 4.172 | 0.057 | -1.183 | 17 | 0.127 | 0.253 | 3.01808E-09 | -9.93663E-09 | 2.79858E-09 |
| | | Equal variances not assumed | | | -1.298 | 15.880 | 0.106 | 0.213 | 2.75022E-09 | -9.40281E-09 | 2.26476E-09 |
| | Linoleic acid | Equal variances assumed | 1.878 | 0.187 | -0.117 | 18 | 0.454 | 0.908 | 3.81161E-09 | -8.45541E-09 | 7.56037E-09 |
| | | Equal variances not assumed | | | -0.111 | 12.023 | 0.457 | 0.913 | 4.02766E-09 | -9.22117E-09 | 8.32613E-09 |

| | | | | | | | | | | | |
|---|---|---|---|---|---|---|---|---|---|---|---|
| | Arachidonic acid | Equal variances assumed | 6.951 | 0.017 | -1.340 | 17 | 0.099 | 0.198 | 2.56055E-09 | -8.83315E-09 | 1.97141E-09 |
| | | Equal variances not assumed | | | -1.470 | 15.893 | 0.081 | 0.161 | 2.33383E-09 | -8.38107E-09 | 1.51932E-09 |
| | Docosahexaenoic acid | Equal variances assumed | 8.879 | 0.009 | -1.477 | 16 | 0.080 | 0.159 | 5.71627E-09 | -2.05613E-08 | 3.67463E-09 |
| | | Equal variances not assumed | | | -1.477 | 10.687 | 0.084 | 0.169 | 5.71627E-09 | -2.10698E-08 | 4.18318E-09 |
| | Eicosapentaenoic acid | Equal variances assumed | 12.217 | 0.003 | -2.325 | 17 | 0.016 | 0.033 | 2.78689E-09 | -1.23581E-08 | -5.98471E-10 |
| | | Equal variances not assumed | | | -2.719 | 10.871 | 0.010 | 0.020 | 2.38264E-09 | -1.17301E-08 | -1.22651E-09 |
| | Oleic acid | Equal variances assumed | 1.677 | 0.213 | 1.265 | 17 | 0.111 | 0.223 | 1.77299E-09 | -1.49745E-09 | 5.98390E-09 |
| | | Equal variances not assumed | | | 1.246 | 14.819 | 0.116 | 0.232 | 1.80052E-09 | -1.59857E-09 | 6.08503E-09 |
| | Linoleic acid | Equal variances assumed | 1.691 | 0.211 | -0.782 | 17 | 0.222 | 0.445 | 2.39411E-09 | -6.92385E-09 | 3.17840E-09 |
| | | Equal variances not assumed | | | -0.761 | 12.667 | 0.230 | 0.460 | 2.45993E-09 | -7.20131E-09 | 3.45587E-09 |
| | Arachidonic acid | Equal variances assumed | 4.070 | 0.061 | 1.402 | 16 | 0.090 | 0.180 | 2.19458E-09 | -1.57582E-09 | 7.72879E-09 |
| THC | | Equal variances not assumed | | | 1.402 | 11.261 | 0.094 | 0.188 | 2.19458E-09 | -1.74011E-09 | 7.89309E-09 |
| | Docosahexaenoic acid | Equal variances assumed | 2.452 | 0.138 | 0.281 | 15 | 0.391 | 0.782 | 6.68811E-09 | -1.23734E-08 | 1.61374E-08 |
| | | Equal variances not assumed | | | 0.290 | 13.357 | 0.388 | 0.776 | 6.49647E-09 | -1.21147E-08 | 1.58787E-08 |
| | Eicosapentaenoic acid | Equal variances assumed | 6.124 | 0.025 | -0.919 | 16 | 0.186 | 0.372 | 1.87625E-09 | -5.70260E-09 | 2.25236E-09 |
| | | Equal variances not assumed | | | -0.919 | 10.024 | 0.190 | 0.379 | 1.87625E-09 | -5.90430E-09 | 2.45406E-09 |
| | Oleic acid | Equal variances assumed | 7.473 | 0.014 | -1.448 | 18 | 0.082 | 0.165 | 1.89072E-09 | -6.70915E-09 | 1.23534E-09 |
| THC+CBD | | Equal variances not assumed | | | -1.448 | 11.324 | 0.087 | 0.175 | 1.89072E-09 | -6.88388E-09 | 1.41007E-09 |

| | | | | | | | | | | |
|---|---|---|---|---|---|---|---|---|---|---|
| Linoleic acid | Equal variances assumed | 0.026 | 0.873 | 0.483 | 19 | 0.317 | 0.635 | 2.21415E-09 | -3.56451E-09 | 5.70402E-09 |
| | Equal variances not assumed | | | 0.482 | 18.492 | 0.318 | 0.636 | 2.22144E-09 | -3.58842E-09 | 5.72793E-09 |
| Arachidonic acid | Equal variances assumed | 4.851 | 0.041 | 2.071 | 18 | 0.027 | 0.053 | 2.24531E-09 | -6.81753E-11 | 9.36626E-09 |
| | Equal variances not assumed | | | 2.184 | 16.346 | 0.022 | 0.044 | 2.12882E-09 | 1.43897E-10 | 9.15419E-09 |
| Docosahexaenoic acid | Equal variances assumed | 0.741 | 0.401 | 0.722 | 17 | 0.240 | 0.480 | 7.88789E-09 | -1.09474E-08 | 2.23366E-08 |
| | Equal variances not assumed | | | 0.725 | 16.980 | 0.239 | 0.478 | 7.85411E-09 | -1.08776E-08 | 2.22668E-08 |
| Eicosapentaenoic acid | Equal variances assumed | 6.031 | 0.024 | -2.426 | 18 | 0.013 | 0.026 | 2.37633E-09 | -1.07575E-08 | -7.72508E-10 |
| | Equal variances not assumed | | | -2.426 | 12.773 | 0.015 | 0.031 | 2.37633E-09 | -1.09081E-08 | -6.21937E-10 |

Supplemental Table 5.4 Independent t-test comparing male and female levels of free fatty acids measured in liver collected from adult mice 2h after an i.p. injection of 10mg/kg CBD, THC, or THC+CBD

| | | | Levene's Test for Equality of Variances | | t-test for Equality of Means | | | | | | | |
|---|---|---|---|---|---|---|---|---|---|---|---|---|
| | | | | | | | Significance | | | | 95% Confidence Interval of the Difference | |
| | | | F | Sig. | t | df | One-Sided p | Two-Sided p | Mean Difference | Std. Error Difference | Lower | Upper |
| | Oleic acid | Equal variances assumed | 2.917 | 0.111 | -1.740 | 13 | 0.053 | 0.105 | -2.31825E-10 | 1.33196E-10 | -5.19577E-10 | 5.59271E-11 |
| | | Equal variances not assumed | | | -1.682 | 9.551 | 0.062 | 0.125 | -2.31825E-10 | 1.37799E-10 | -5.40827E-10 | 7.71772E-11 |
| | Linoleic acid | Equal variances assumed | 0.174 | 0.683 | -0.534 | 13 | 0.301 | 0.603 | -1.20049E-10 | 2.24947E-10 | -6.06018E-10 | 3.65920E-10 |
| | | Equal variances not assumed | | | -0.532 | 12.507 | 0.302 | 0.604 | -1.20049E-10 | 2.25834E-10 | -6.09897E-10 | 3.69799E-10 |
| | Arachidonic acid | Equal variances assumed | 16.072 | 0.002 | -2.944 | 12 | 0.006 | 0.012 | -2.54350E-09 | 8.63941E-10 | -4.42587E-09 | -6.61138E-10 |
| Veh | | Equal variances not assumed | | | -2.944 | 8.317 | 0.009 | 0.018 | -2.54350E-09 | 8.63941E-10 | -4.52261E-09 | -5.64400E-10 |
| | Docosahexaenoic acid | Equal variances assumed | 0.549 | 0.473 | -2.392 | 12 | 0.017 | 0.034 | -5.76140E-10 | 2.40876E-10 | -1.10096E-09 | -5.13167E-11 |
| | | Equal variances not assumed | | | -2.415 | 11.291 | 0.017 | 0.034 | -5.76140E-10 | 2.38535E-10 | -1.09951E-09 | -5.27727E-11 |
| | Eicosapentaenoic acid | Equal variances assumed | 2.778 | 0.119 | -4.295 | 13 | 0.000 | 0.001 | -5.41783E-10 | 1.26147E-10 | -8.14307E-10 | -2.69259E-10 |
| | | Equal variances not assumed | | | -4.105 | 8.360 | 0.002 | 0.003 | -5.41783E-10 | 1.31979E-10 | -8.43864E-10 | -2.39702E-10 |
| | Oleic acid | Equal variances assumed | 0.154 | 0.700 | -1.067 | 16 | 0.151 | 0.302 | -1.35088E-10 | 1.26560E-10 | -4.03383E-10 | 1.33207E-10 |
| | | Equal variances not assumed | | | -1.052 | 14.169 | 0.155 | 0.310 | -1.35088E-10 | 1.28425E-10 | -4.10224E-10 | 1.40048E-10 |
| CBD | Linoleic acid | Equal variances assumed | 2.123 | 0.166 | -1.533 | 15 | 0.073 | 0.146 | -2.92732E-10 | 1.90999E-10 | -6.99836E-10 | 1.14371E-10 |
| | | Equal variances not assumed | | | -1.674 | 14.774 | 0.058 | 0.115 | -2.92732E-10 | 1.74838E-10 | -6.65888E-10 | 8.04233E-11 |
| | Arachidonic acid | Equal variances assumed | 0.187 | 0.671 | -1.366 | 16 | 0.095 | 0.191 | -1.02242E-09 | 7.48327E-10 | -2.60880E-09 | 5.63967E-10 |

| | | | | | | | | | | | | |
|---|---|---|---|---|---|---|---|---|---|---|---|---|
| | | Equal variances not assumed | | | -1.367 | 15.189 | 0.096 | 0.191 | -1.02242E-09 | 7.47715E-10 | -2.61441E-09 | 5.69575E-10 |
| | Docosahexaenoic acid | Equal variances assumed | 4.236 | 0.057 | 0.244 | 15 | 0.405 | 0.811 | 7.30937E-11 | 3.00168E-10 | -5.66700E-10 | 7.12887E-10 |
| | | Equal variances not assumed | | | 0.233 | 9.169 | 0.410 | 0.821 | 7.30937E-11 | 3.13698E-10 | -6.34557E-10 | 7.80744E-10 |
| | Eicosapentaenoic acid | Equal variances assumed | 1.297 | 0.274 | -3.920 | 14 | 0.001 | 0.002 | -4.75240E-10 | 1.21231E-10 | -7.35255E-10 | -2.15226E-10 |
| | | Equal variances not assumed | | | -4.068 | 13.997 | 0.001 | 0.001 | -4.75240E-10 | 1.16811E-10 | -7.25780E-10 | -2.24701E-10 |
| | Oleic acid | Equal variances assumed | 0.008 | 0.931 | 1.897 | 15 | 0.039 | 0.077 | 1.83540E-10 | 9.67321E-11 | -2.26395E-11 | 3.89720E-10 |
| | | Equal variances not assumed | | | 1.886 | 14.286 | 0.040 | 0.080 | 1.83540E-10 | 9.73316E-11 | -2.48236E-11 | 3.91904E-10 |
| | Linoleic acid | Equal variances assumed | 4.265 | 0.055 | -0.722 | 17 | 0.240 | 0.480 | -2.26219E-10 | 3.13426E-10 | -8.87491E-10 | 4.35053E-10 |
| | | Equal variances not assumed | | | -0.705 | 13.359 | 0.246 | 0.493 | -2.26219E-10 | 3.20859E-10 | -9.17504E-10 | 4.65066E-10 |
| | Arachidonic acid | Equal variances assumed | 3.888 | 0.065 | -0.679 | 17 | 0.253 | 0.506 | -5.14428E-10 | 7.57074E-10 | -2.11171E-09 | 1.08286E-09 |
| THC | | Equal variances not assumed | | | -0.658 | 11.649 | 0.262 | 0.524 | -5.14428E-10 | 7.82179E-10 | -2.22437E-09 | 1.19551E-09 |
| | Docosahexaenoic acid | Equal variances assumed | 0.079 | 0.782 | -1.268 | 15 | 0.112 | 0.224 | -3.24979E-10 | 2.56283E-10 | -8.71233E-10 | 2.21275E-10 |
| | | Equal variances not assumed | | | -1.258 | 14.099 | 0.114 | 0.229 | -3.24979E-10 | 2.58354E-10 | -8.78730E-10 | 2.28771E-10 |
| | Eicosapentaenoic acid | Equal variances assumed | 12.945 | 0.002 | -2.787 | 16 | 0.007 | 0.013 | -8.53325E-10 | 3.06126E-10 | -1.50228E-09 | -2.04366E-10 |
| | | Equal variances not assumed | | | -2.787 | 8.615 | 0.011 | 0.022 | -8.53325E-10 | 3.06126E-10 | -1.55057E-09 | -1.56079E-10 |
| | Oleic acid | Equal variances assumed | 0.124 | 0.729 | 0.233 | 18 | 0.409 | 0.818 | 2.81553E-11 | 1.20753E-10 | -2.25538E-10 | 2.81849E-10 |
| THC+CBD | | Equal variances not assumed | | | 0.233 | 17.703 | 0.409 | 0.818 | 2.81553E-11 | 1.20753E-10 | -2.25843E-10 | 2.82154E-10 |
| | Linoleic acid | Equal variances assumed | 2.458 | 0.134 | 1.462 | 18 | 0.081 | 0.161 | 4.57326E-10 | 3.12888E-10 | -2.00028E-10 | 1.11468E-09 |
| | | Equal variances | | | 1.462 | 16.222 | 0.081 | 0.163 | 4.57326E-10 | 3.12888E-10 | -2.05230E-10 | 1.11988E-09 |

| | | | | | | | | | | | |
|---|---|---|---|---|---|---|---|---|---|---|---|
| | not assumed | | | | | | | | | | |
| Arachidonic acid | Equal variances assumed | 5.385 | 0.032 | -0.295 | 19 | 0.385 | 0.771 | -2.27852E-10 | 7.71547E-10 | -1.84272E-09 | 1.38702E-09 |
| | Equal variances not assumed | | | -0.290 | 15.413 | 0.388 | 0.776 | -2.27852E-10 | 7.86773E-10 | -1.90091E-09 | 1.44521E-09 |
| Docosahexaenoic acid | Equal variances assumed | 3.923 | 0.062 | -1.076 | 19 | 0.148 | 0.295 | -5.14591E-10 | 4.78056E-10 | -1.51517E-09 | 4.85991E-10 |
| | Equal variances not assumed | | | -1.104 | 16.265 | 0.143 | 0.286 | -5.14591E-10 | 4.66264E-10 | -1.50172E-09 | 4.72538E-10 |
| Eicosapentaenoic acid | Equal variances assumed | 5.947 | 0.025 | -3.683 | 19 | 0.001 | 0.002 | -1.20879E-09 | 3.28245E-10 | -1.89581E-09 | -5.21761E-10 |
| | Equal variances not assumed | | | -3.791 | 15.262 | 0.001 | 0.002 | -1.20879E-09 | 3.18860E-10 | -1.88741E-09 | -5.30166E-10 |

Supplemental Table 5.5 One-way ANOVA for free fatty acids measured in plasma collected from adult mice 2h after an i.p. injection of 10mg/kg CBD, THC, or THC+CBD

| | | Sum of Squares | df | Mean Square | F | Sig. |
|---|---|---|---|---|---|---|
| Oleic acid | Between Groups | 1.82E-16 | 7 | 2.60E-17 | 1.075 | 0.3893495 |
| | Within Groups | 1.57E-15 | 65 | 2.41E-17 | | |
| | Total | 1.75E-15 | 72 | | | |
| Linoleic acid | Between Groups | 2.77E-16 | 7 | 3.96E-17 | 1.109 | 0.3677163 |
| | Within Groups | 2.39E-15 | 67 | 3.57E-17 | | |
| | Total | 2.67E-15 | 74 | | | |
| Arachidonic acid | Between Groups | 2.29E-16 | 7 | 3.27E-17 | 1.519 | 0.177 |
| | Within Groups | 1.38E-15 | 64 | 2.15E-17 | | |
| | Total | 1.61E-15 | 71 | | | |
| Docosahexaenoic acid | Between Groups | 2.94E-15 | 7 | 4.20E-16 | 2.005 | 0.0692505 |
| | Within Groups | 1.26E-14 | 60 | 2.09E-16 | | |
| | Total | 1.55E-14 | 67 | | | |
| Eicosapentaenoic acid | Between Groups | 5.34E-16 | 7 | 7.62E-17 | 3.264 | 0.0050088 |
| | Within Groups | 1.49E-15 | 64 | 2.33E-17 | | |
| | Total | 2.03E-15 | 71 | | | |

Supplemental Table 5.6 One-way ANOVA for free fatty acids measured in liver collected from adult mice 2h after an i.p. injection of 10mg/kg CBD, THC, or THC+CBD

| | | Sum of Squares | df | Mean Square | F | Sig. |
|---|---|---|---|---|---|---|
| Oleic acid | Between Groups | 9.79E-19 | 7 | 1.40E-19 | 2.219 | 0.04445 |
| | Within Groups | 3.91E-18 | 62 | 6.30E-20 | | |
| | Total | 4.89E-18 | 69 | | | |
| Linoleic acid | Between Groups | 1.82E-17 | 7 | 2.60E-18 | 7.637 | 0.00000 |
| | Within Groups | 2.14E-17 | 63 | 3.40E-19 | | |
| | Total | 3.96E-17 | 70 | | | |
| Arachidonic acid | Between Groups | 1.17E-16 | 7 | 1.68E-17 | 6.074 | 0.00002 |
| | Within Groups | 1.77E-16 | 64 | 2.76E-18 | | |
| | Total | 2.94E-16 | 71 | | | |
| Docosahexaenoic acid | Between Groups | 4.22E-17 | 7 | 6.03E-18 | 10.504 | 0.00000 |
| | Within Groups | 3.50E-17 | 61 | 5.74E-19 | | |
| | Total | 7.73E-17 | 68 | | | |
| Eicosapentaenoic acid | Between Groups | 3.18E-17 | 7 | 4.54E-18 | 14.765 | 0.00000 |
| | Within Groups | 1.91E-17 | 62 | 3.07E-19 | | |
| | Total | 5.08E-17 | 69 | | | |

#### Supplemental Table 5.7 Fisher's LSD post-hoc test for plasma free fatty acid ANOVA

| Dependent Variable | | | Std. Error | Sig. | 95% Confidence Interval Lower Bound | Upper Bound |
|---|---|---|---|---|---|---|
| Oleic acid | VehFemale | CBDfemale | 2.54E-09 | 0.90 | -5.41E-09 | 4.75E-09 |
| | | THCfemale | 2.42E-09 | 0.59 | -6.14E-09 | 3.53E-09 |
| | | TpCfemale | 2.42E-09 | 0.89 | -5.18E-09 | 4.49E-09 |
| | | VehMale | 2.54E-09 | 0.47 | -6.92E-09 | 3.24E-09 |
| | | CBDmale | 2.38E-09 | 0.11 | -8.64E-09 | 8.48E-10 |
| | | THCmale | 2.48E-09 | 0.71 | -4.00E-09 | 5.89E-09 |
| | | TpCmale | 2.42E-09 | 0.21 | -7.92E-09 | 1.76E-09 |
| | CBDfemale | VehFemale | 2.54E-09 | 0.90 | -4.75E-09 | 5.41E-09 |
| | | THCfemale | 2.33E-09 | 0.68 | -5.63E-09 | 3.68E-09 |
| | | TpCfemale | 2.33E-09 | 1.00 | -4.67E-09 | 4.64E-09 |
| | | VehMale | 2.46E-09 | 0.54 | -6.42E-09 | 3.39E-09 |
| | | CBDmale | 2.28E-09 | 0.12 | -8.13E-09 | 9.91E-10 |
| | | THCmale | 2.39E-09 | 0.60 | -3.50E-09 | 6.04E-09 |
| | | TpCmale | 2.33E-09 | 0.24 | -7.41E-09 | 1.90E-09 |
| | THCfemale | VehFemale | 2.42E-09 | 0.59 | -3.53E-09 | 6.14E-09 |
| | | CBDfemale | 2.33E-09 | 0.68 | -3.68E-09 | 5.63E-09 |
| | | TpCfemale | 2.20E-09 | 0.66 | -3.43E-09 | 5.35E-09 |
| | | VehMale | 2.33E-09 | 0.82 | -5.20E-09 | 4.11E-09 |
| | | CBDmale | 2.15E-09 | 0.23 | -6.88E-09 | 1.69E-09 |
| | | THCmale | 2.26E-09 | 0.32 | -2.27E-09 | 6.75E-09 |
| | | TpCmale | 2.20E-09 | 0.42 | -6.17E-09 | 2.61E-09 |
| | TpCfemale | VehFemale | 2.42E-09 | 0.89 | -4.49E-09 | 5.18E-09 |
| | | CBDfemale | 2.33E-09 | 1.00 | -4.64E-09 | 4.67E-09 |
| | | THCfemale | 2.20E-09 | 0.66 | -5.35E-09 | 3.43E-09 |
| | | VehMale | 2.33E-09 | 0.52 | -6.16E-09 | 3.15E-09 |
| | | CBDmale | 2.15E-09 | 0.10 | -7.84E-09 | 7.33E-10 |
| | | THCmale | 2.26E-09 | 0.57 | -3.23E-09 | 5.79E-09 |
| | | TpCmale | 2.20E-09 | 0.22 | -7.13E-09 | 1.65E-09 |
| | VehMale | VehFemale | 2.54E-09 | 0.47 | -3.24E-09 | 6.92E-09 |
| | | CBDfemale | 2.46E-09 | 0.54 | -3.39E-09 | 6.42E-09 |
| | | THCfemale | 2.33E-09 | 0.82 | -4.11E-09 | 5.20E-09 |
| | | TpCfemale | 2.33E-09 | 0.52 | -3.15E-09 | 6.16E-09 |
| | | CBDmale | 2.28E-09 | 0.37 | -6.61E-09 | 2.51E-09 |
| | | THCmale | 2.39E-09 | 0.25 | -1.98E-09 | 7.55E-09 |
| | | TpCmale | 2.33E-09 | 0.60 | -5.89E-09 | 3.42E-09 |
| | CBDmale | VehFemale | 2.38E-09 | 0.11 | -8.48E-10 | 8.64E-09 |
| | | CBDfemale | 2.28E-09 | 0.12 | -9.91E-10 | 8.13E-09 |
| | | THCfemale | 2.15E-09 | 0.23 | -1.69E-09 | 6.88E-09 |
| | | TpCfemale | 2.15E-09 | 0.10 | -7.33E-10 | 7.84E-09 |
| | | VehMale | 2.28E-09 | 0.37 | -2.51E-09 | 6.61E-09 |
| | | THCmale | 2.21E-09 | 0.03 | 4.28E-10 | 9.25E-09 |
| | | TpCmale | 2.15E-09 | 0.70 | -3.47E-09 | 5.11E-09 |
| | THCmale | VehFemale | 2.48E-09 | 0.71 | -5.89E-09 | 4.00E-09 |
| | | CBDfemale | 2.39E-09 | 0.60 | -6.04E-09 | 3.50E-09 |
| | | THCfemale | 2.26E-09 | 0.32 | -6.75E-09 | 2.27E-09 |
| | | TpCfemale | 2.26E-09 | 0.57 | -5.79E-09 | 3.23E-09 |
| | | VehMale | 2.39E-09 | 0.25 | -7.55E-09 | 1.98E-09 |
| | | CBDmale | 2.21E-09 | 0.03 | -9.25E-09 | -4.28E-10 |
| | | TpCmale | 2.26E-09 | 0.08 | -8.53E-09 | 4.88E-10 |
| | TpCmale | VehFemale | 2.42E-09 | 0.21 | -1.76E-09 | 7.92E-09 |
| | | CBDfemale | 2.33E-09 | 0.24 | -1.90E-09 | 7.41E-09 |
| | | THCfemale | 2.20E-09 | 0.42 | -2.61E-09 | 6.17E-09 |
| | | TpCfemale | 2.20E-09 | 0.22 | -1.65E-09 | 7.13E-09 |
| | | VehMale | 2.33E-09 | 0.60 | -3.42E-09 | 5.89E-09 |
| | | CBDmale | 2.15E-09 | 0.70 | -5.11E-09 | 3.47E-09 |
| | | THCmale | 2.26E-09 | 0.08 | -4.88E-10 | 8.53E-09 |
| Linoleic acid | VehFemale | CBDfemale | 3.01E-09 | 0.74 | -5.01E-09 | 7.01E-09 |

| | | | | | | |
|---|---|---|---|---|---|---|
| | | THCfemale | 2.95E-09 | 0.05 | 1.15E-10 | 1.19E-08 |
| | | TpCfemale | 2.89E-09 | 0.42 | -3.43E-09 | 8.10E-09 |
| | | VehMale | 3.09E-09 | 0.19 | -2.04E-09 | 1.03E-08 |
| | | CBDmale | 2.89E-09 | 0.85 | -5.22E-09 | 6.32E-09 |
| | | THCmale | 3.01E-09 | 0.18 | -1.89E-09 | 1.01E-08 |
| | | TpCmale | 2.95E-09 | 0.25 | -2.48E-09 | 9.28E-09 |
| | CBDfemale | VehFemale | 3.01E-09 | 0.74 | -7.01E-09 | 5.01E-09 |
| | | THCfemale | 2.75E-09 | 0.07 | -4.86E-10 | 1.05E-08 |
| | | TpCfemale | 2.69E-09 | 0.62 | -4.03E-09 | 6.69E-09 |
| | | VehMale | 2.90E-09 | 0.28 | -2.66E-09 | 8.93E-09 |
| | | CBDmale | 2.69E-09 | 0.87 | -5.81E-09 | 4.91E-09 |
| | | THCmale | 2.82E-09 | 0.27 | -2.50E-09 | 8.74E-09 |
| | | TpCmale | 2.75E-09 | 0.38 | -3.08E-09 | 7.88E-09 |
| | THCfemale | VehFemale | 2.95E-09 | 0.05 | -1.19E-08 | -1.15E-10 |
| | | CBDfemale | 2.75E-09 | 0.07 | -1.05E-08 | 4.86E-10 |
| | | TpCfemale | 2.61E-09 | 0.17 | -8.87E-09 | 1.55E-09 |
| | | VehMale | 2.83E-09 | 0.51 | -7.52E-09 | 3.80E-09 |
| | | CBDmale | 2.61E-09 | 0.04 | -1.07E-08 | -2.30E-10 |
| | | THCmale | 2.75E-09 | 0.50 | -7.35E-09 | 3.61E-09 |
| | | TpCmale | 2.67E-09 | 0.34 | -7.93E-09 | 2.74E-09 |
| | TpCfemale | VehFemale | 2.89E-09 | 0.42 | -8.10E-09 | 3.43E-09 |
| | | CBDfemale | 2.69E-09 | 0.62 | -6.69E-09 | 4.03E-09 |
| | | THCfemale | 2.61E-09 | 0.17 | -1.55E-09 | 8.87E-09 |
| | | VehMale | 2.78E-09 | 0.52 | -3.74E-09 | 7.34E-09 |
| | | CBDmale | 2.55E-09 | 0.49 | -6.87E-09 | 3.31E-09 |
| | | THCmale | 2.69E-09 | 0.51 | -3.57E-09 | 7.15E-09 |
| | | TpCmale | 2.61E-09 | 0.68 | -4.14E-09 | 6.28E-09 |
| | VehMale | VehFemale | 3.09E-09 | 0.19 | -1.03E-08 | 2.04E-09 |
| | | CBDfemale | 2.90E-09 | 0.28 | -8.93E-09 | 2.66E-09 |
| | | THCfemale | 2.83E-09 | 0.51 | -3.80E-09 | 7.52E-09 |
| | | TpCfemale | 2.78E-09 | 0.52 | -7.34E-09 | 3.74E-09 |
| | | CBDmale | 2.78E-09 | 0.20 | -9.13E-09 | 1.96E-09 |
| | | THCmale | 2.90E-09 | 1.00 | -5.81E-09 | 5.78E-09 |
| | | TpCmale | 2.83E-09 | 0.80 | -6.39E-09 | 4.93E-09 |
| | CBDmale | VehFemale | 2.89E-09 | 0.85 | -6.32E-09 | 5.22E-09 |
| | | CBDfemale | 2.69E-09 | 0.87 | -4.91E-09 | 5.81E-09 |
| | | THCfemale | 2.61E-09 | 0.04 | 2.30E-10 | 1.07E-08 |
| | | TpCfemale | 2.55E-09 | 0.49 | -3.31E-09 | 6.87E-09 |
| | | VehMale | 2.78E-09 | 0.20 | -1.96E-09 | 9.13E-09 |
| | | THCmale | 2.69E-09 | 0.19 | -1.79E-09 | 8.93E-09 |
| | | TpCmale | 2.61E-09 | 0.28 | -2.36E-09 | 8.06E-09 |
| | THCmale | VehFemale | 3.01E-09 | 0.18 | -1.01E-08 | 1.89E-09 |
| | | CBDfemale | 2.82E-09 | 0.27 | -8.74E-09 | 2.50E-09 |
| | | THCfemale | 2.75E-09 | 0.50 | -3.61E-09 | 7.35E-09 |
| | | TpCfemale | 2.69E-09 | 0.51 | -7.15E-09 | 3.57E-09 |
| | | VehMale | 2.90E-09 | 1.00 | -5.78E-09 | 5.81E-09 |
| | | CBDmale | 2.69E-09 | 0.19 | -8.93E-09 | 1.79E-09 |
| | | TpCmale | 2.75E-09 | 0.79 | -6.20E-09 | 4.76E-09 |
| | TpCmale | VehFemale | 2.95E-09 | 0.25 | -9.28E-09 | 2.48E-09 |
| | | CBDfemale | 2.75E-09 | 0.38 | -7.88E-09 | 3.08E-09 |
| | | THCfemale | 2.67E-09 | 0.34 | -2.74E-09 | 7.93E-09 |
| | | TpCfemale | 2.61E-09 | 0.68 | -6.28E-09 | 4.14E-09 |
| | | VehMale | 2.83E-09 | 0.80 | -4.93E-09 | 6.39E-09 |
| | | CBDmale | 2.61E-09 | 0.28 | -8.06E-09 | 2.36E-09 |
| | | THCmale | 2.75E-09 | 0.79 | -4.76E-09 | 6.20E-09 |
| Arachidonic acid | VehFemale | CBDfemale | 2.40070E-09 | 0.49 | -3.1284E-09 | 6.4635E-09 |
| | | THCfemale | 2.33763E-09 | 0.25 | -7.4057E-09 | 1.9342E-09 |
| | | TpCfemale | 2.24273E-09 | 0.10 | -8.1884E-09 | 7.7235E-10 |
| | | VehMale | 2.40070E-09 | 0.59 | -6.0834E-09 | 3.5085E-09 |
| | | CBDmale | 2.24273E-09 | 0.43 | -6.2437E-09 | 2.7171E-09 |

| | | | | | | |
|---|---|---|---|---|---|---|
| | | THCmale | 2.33763E-09 | 0.88 | -4.3292E-09 | 5.0107E-09 |
| | | TpCmale | 2.33763E-09 | 0.69 | -3.7289E-09 | 5.6110E-09 |
| | CBDfemale | VehFemale | 2.40070E-09 | 0.49 | -6.4635E-09 | 3.1284E-09 |
| | | THCfemale | 2.25395E-09 | 0.06 | -8.9061E-09 | 9.9467E-11 |
| | | TpCfemale | 2.15537E-09 | 0.02 | -9.6815E-09 | -1.0698E-09 |
| | | VehMale | 2.31930E-09 | 0.21 | -7.5883E-09 | 1.6783E-09 |
| | | CBDmale | 2.15537E-09 | 0.12 | -7.7367E-09 | 8.7497E-10 |
| | | THCmale | 2.25395E-09 | 0.56 | -5.8296E-09 | 3.1760E-09 |
| | | TpCmale | 2.25395E-09 | 0.75 | -5.2294E-09 | 3.7762E-09 |
| | THCfemale | VehFemale | 2.33763E-09 | 0.25 | -1.9342E-09 | 7.4057E-09 |
| | | CBDfemale | 2.25395E-09 | 0.06 | -9.9467E-11 | 8.9061E-09 |
| | | TpCfemale | 2.08489E-09 | 0.64 | -5.1373E-09 | 3.1928E-09 |
| | | VehMale | 2.25395E-09 | 0.52 | -3.0545E-09 | 5.9511E-09 |
| | | CBDmale | 2.08489E-09 | 0.64 | -3.1926E-09 | 5.1375E-09 |
| | | THCmale | 2.18665E-09 | 0.16 | -1.2919E-09 | 7.4448E-09 |
| | | TpCmale | 2.18665E-09 | 0.10 | -6.9159E-10 | 8.0451E-09 |
| | TpCfemale | VehFemale | 2.24273E-09 | 0.10 | -7.7235E-10 | 8.1884E-09 |
| | | CBDfemale | 2.15537E-09 | 0.02 | 1.0698E-09 | 9.6815E-09 |
| | | THCfemale | 2.08489E-09 | 0.64 | -3.1928E-09 | 5.1373E-09 |
| | | VehMale | 2.15537E-09 | 0.27 | -1.8852E-09 | 6.7265E-09 |
| | | CBDmale | 1.97790E-09 | 0.33 | -2.0066E-09 | 5.8961E-09 |
| | | THCmale | 2.08489E-09 | 0.06 | -1.1627E-10 | 8.2138E-09 |
| | | TpCmale | 2.08489E-09 | 0.03 | 4.8399E-10 | 8.8141E-09 |
| | VehMale | VehFemale | 2.40070E-09 | 0.59 | -3.5085E-09 | 6.0834E-09 |
| | | CBDfemale | 2.31930E-09 | 0.21 | -1.6783E-09 | 7.5883E-09 |
| | | THCfemale | 2.25395E-09 | 0.52 | -5.9511E-09 | 3.0545E-09 |
| | | TpCfemale | 2.15537E-09 | 0.27 | -6.7265E-09 | 1.8852E-09 |
| | | CBDmale | 2.15537E-09 | 0.83 | -4.7817E-09 | 3.8300E-09 |
| | | THCmale | 2.25395E-09 | 0.47 | -2.8746E-09 | 6.1310E-09 |
| | | TpCmale | 2.25395E-09 | 0.33 | -2.2743E-09 | 6.7312E-09 |
| | CBDmale | VehFemale | 2.24273E-09 | 0.43 | -2.7171E-09 | 6.2437E-09 |
| | | CBDfemale | 2.15537E-09 | 0.12 | -8.7497E-10 | 7.7367E-09 |
| | | THCfemale | 2.08489E-09 | 0.64 | -5.1375E-09 | 3.1926E-09 |
| | | TpCfemale | 1.97790E-09 | 0.33 | -5.8961E-09 | 2.0066E-09 |
| | | VehMale | 2.15537E-09 | 0.83 | -3.8300E-09 | 4.7817E-09 |
| | | THCmale | 2.08489E-09 | 0.32 | -2.0610E-09 | 6.2691E-09 |
| | | TpCmale | 2.08489E-09 | 0.20 | -1.4607E-09 | 6.8694E-09 |
| | THCmale | VehFemale | 2.33763E-09 | 0.88 | -5.0107E-09 | 4.3292E-09 |
| | | CBDfemale | 2.25395E-09 | 0.56 | -3.1760E-09 | 5.8296E-09 |
| | | THCfemale | 2.18665E-09 | 0.16 | -7.4448E-09 | 1.2919E-09 |
| | | TpCfemale | 2.08489E-09 | 0.06 | -8.2138E-09 | 1.1627E-10 |
| | | VehMale | 2.25395E-09 | 0.47 | -6.1310E-09 | 2.8746E-09 |
| | | CBDmale | 2.08489E-09 | 0.32 | -6.2691E-09 | 2.0610E-09 |
| | | TpCmale | 2.18665E-09 | 0.78 | -3.7681E-09 | 4.9686E-09 |
| | TpCmale | VehFemale | 2.33763E-09 | 0.69 | -5.6110E-09 | 3.7289E-09 |
| | | CBDfemale | 2.25395E-09 | 0.75 | -3.7762E-09 | 5.2294E-09 |
| | | THCfemale | 2.18665E-09 | 0.10 | -8.0451E-09 | 6.9159E-10 |
| | | TpCfemale | 2.08489E-09 | 0.03 | -8.8141E-09 | -4.8399E-10 |
| | | VehMale | 2.25395E-09 | 0.33 | -6.7312E-09 | 2.2743E-09 |
| | | CBDmale | 2.08489E-09 | 0.20 | -6.8694E-09 | 1.4607E-09 |
| | | THCmale | 2.18665E-09 | 0.78 | -4.9686E-09 | 3.7681E-09 |
| | VehFemale | CBDfemale | 7.29E-09 | 0.03 | 2.10E-09 | 3.13E-08 |
| | | THCfemale | 7.29E-09 | 0.55 | -1.02E-08 | 1.89E-08 |
| | | TpCfemale | 7.13E-09 | 0.62 | -1.78E-08 | 1.07E-08 |
| Docosahexaenoic acid | | VehMale | 7.74E-09 | 0.51 | -2.06E-08 | 1.04E-08 |
| | | CBDmale | 7.29E-09 | 0.26 | -6.34E-09 | 2.28E-08 |
| | | THCmale | 7.49E-09 | 0.41 | -8.76E-09 | 2.12E-08 |
| | | TpCmale | 7.29E-09 | 0.77 | -1.24E-08 | 1.68E-08 |
| | CBDfemale | VehFemale | 7.29E-09 | 0.03 | -3.13E-08 | -2.10E-09 |

| | | | | | | |
|---|---|---|---|---|---|---|
| | | THCfemale | 6.82E-09 | 0.08 | -2.60E-08 | 1.30E-09 |
| | | TpCfemale | 6.65E-09 | 0.00 | -3.35E-08 | -6.91E-09 |
| | | VehMale | 7.29E-09 | 0.00 | -3.64E-08 | -7.21E-09 |
| | | CBDmale | 6.82E-09 | 0.22 | -2.21E-08 | 5.20E-09 |
| | | THCmale | 7.03E-09 | 0.14 | -2.45E-08 | 3.60E-09 |
| | | TpCmale | 6.82E-09 | 0.04 | -2.82E-08 | -8.69E-10 |
| | THCfemale | VehFemale | 7.29E-09 | 0.55 | -1.89E-08 | 1.02E-08 |
| | | CBDfemale | 6.82E-09 | 0.08 | -1.30E-09 | 2.60E-08 |
| | | TpCfemale | 6.65E-09 | 0.24 | -2.12E-08 | 5.44E-09 |
| | | VehMale | 7.29E-09 | 0.20 | -2.40E-08 | 5.14E-09 |
| | | CBDmale | 6.82E-09 | 0.57 | -9.75E-09 | 1.76E-08 |
| | | THCmale | 7.03E-09 | 0.79 | -1.22E-08 | 1.60E-08 |
| | | TpCmale | 6.82E-09 | 0.75 | -1.58E-08 | 1.15E-08 |
| | TpCfemale | VehFemale | 7.13E-09 | 0.62 | -1.07E-08 | 1.78E-08 |
| | | CBDfemale | 6.65E-09 | 0.00 | 6.91E-09 | 3.35E-08 |
| | | THCfemale | 6.65E-09 | 0.24 | -5.44E-09 | 2.12E-08 |
| | | VehMale | 7.13E-09 | 0.82 | -1.59E-08 | 1.27E-08 |
| | | CBDmale | 6.65E-09 | 0.08 | -1.53E-09 | 2.51E-08 |
| | | THCmale | 6.87E-09 | 0.16 | -3.99E-09 | 2.35E-08 |
| | | TpCmale | 6.65E-09 | 0.40 | -7.61E-09 | 1.90E-08 |
| | VehMale | VehFemale | 7.74E-09 | 0.51 | -1.04E-08 | 2.06E-08 |
| | | CBDfemale | 7.29E-09 | 0.00 | 7.21E-09 | 3.64E-08 |
| | | THCfemale | 7.29E-09 | 0.20 | -5.14E-09 | 2.40E-08 |
| | | TpCfemale | 7.13E-09 | 0.82 | -1.27E-08 | 1.59E-08 |
| | | CBDmale | 7.29E-09 | 0.07 | -1.23E-09 | 2.79E-08 |
| | | THCmale | 7.49E-09 | 0.14 | -3.65E-09 | 2.63E-08 |
| | | TpCmale | 7.29E-09 | 0.32 | -7.31E-09 | 2.19E-08 |
| | CBDmale | VehFemale | 7.29E-09 | 0.26 | -2.28E-08 | 6.34E-09 |
| | | CBDfemale | 6.82E-09 | 0.22 | -5.20E-09 | 2.21E-08 |
| | | THCfemale | 6.82E-09 | 0.57 | -1.76E-08 | 9.75E-09 |
| | | TpCfemale | 6.65E-09 | 0.08 | -2.51E-08 | 1.53E-09 |
| | | VehMale | 7.29E-09 | 0.07 | -2.79E-08 | 1.23E-09 |
| | | THCmale | 7.03E-09 | 0.77 | -1.61E-08 | 1.20E-08 |
| | | TpCmale | 6.82E-09 | 0.38 | -1.97E-08 | 7.57E-09 |
| | THCmale | VehFemale | 7.49E-09 | 0.41 | -2.12E-08 | 8.76E-09 |
| | | CBDfemale | 7.03E-09 | 0.14 | -3.60E-09 | 2.45E-08 |
| | | THCfemale | 7.03E-09 | 0.79 | -1.60E-08 | 1.22E-08 |
| | | TpCfemale | 6.87E-09 | 0.16 | -2.35E-08 | 3.99E-09 |
| | | VehMale | 7.49E-09 | 0.14 | -2.63E-08 | 3.65E-09 |
| | | CBDmale | 7.03E-09 | 0.77 | -1.20E-08 | 1.61E-08 |
| | | TpCmale | 7.03E-09 | 0.57 | -1.81E-08 | 1.00E-08 |
| | TpCmale | VehFemale | 7.29E-09 | 0.77 | -1.68E-08 | 1.24E-08 |
| | | CBDfemale | 6.82E-09 | 0.04 | 8.69E-10 | 2.82E-08 |
| | | THCfemale | 6.82E-09 | 0.75 | -1.15E-08 | 1.58E-08 |
| | | TpCfemale | 6.65E-09 | 0.40 | -1.90E-08 | 7.61E-09 |
| | | VehMale | 7.29E-09 | 0.32 | -2.19E-08 | 7.31E-09 |
| | | CBDmale | 6.82E-09 | 0.38 | -7.57E-09 | 1.97E-08 |
| | | THCmale | 7.03E-09 | 0.57 | -1.00E-08 | 1.81E-08 |
| Eicosapentaenoic acid | VehFemale | CBDfemale | 2.50E-09 | 0.66 | -3.87E-09 | 6.12E-09 |
| | | THCfemale | 2.44E-09 | 0.54 | -3.38E-09 | 6.35E-09 |
| | | TpCfemale | 2.38E-09 | 0.83 | -4.25E-09 | 5.26E-09 |
| | | VehMale | 2.50E-09 | 0.21 | -8.19E-09 | 1.80E-09 |
| | | CBDmale | 2.34E-09 | 0.03 | -1.00E-08 | -6.89E-10 |
| | | THCmale | 2.44E-09 | 0.92 | -5.10E-09 | 4.63E-09 |
| | | TpCmale | 2.38E-09 | 0.03 | -1.00E-08 | -5.01E-10 |
| | CBDfemale | VehFemale | 2.50E-09 | 0.66 | -6.12E-09 | 3.87E-09 |
| | | THCfemale | 2.35E-09 | 0.88 | -4.32E-09 | 5.06E-09 |
| | | TpCfemale | 2.29E-09 | 0.79 | -5.19E-09 | 3.96E-09 |
| | | VehMale | 2.42E-09 | 0.08 | -9.15E-09 | 5.07E-10 |
| | | CBDmale | 2.25E-09 | 0.01 | -1.10E-08 | -1.99E-09 |

| | | | | | |
|---|---|---|---|---|---|
| | THCmale | 2.35E-09 | 0.57 | -6.05E-09 | 3.33E-09 |
| | TpCmale | 2.29E-09 | 0.01 | -1.10E-08 | -1.80E-09 |
| THCfemale | VehFemale | 2.44E-09 | 0.54 | -6.35E-09 | 3.38E-09 |
| | CBDfemale | 2.35E-09 | 0.88 | -5.06E-09 | 4.32E-09 |
| | TpCfemale | 2.22E-09 | 0.66 | -5.42E-09 | 3.45E-09 |
| | VehMale | 2.35E-09 | 0.05 | -9.38E-09 | 3.58E-12 |
| | CBDmale | 2.17E-09 | 0.00 | -1.12E-08 | -2.51E-09 |
| | THCmale | 2.28E-09 | 0.45 | -6.28E-09 | 2.83E-09 |
| | TpCmale | 2.22E-09 | 0.00 | -1.12E-08 | -2.31E-09 |
| TpCfemale | VehFemale | 2.38E-09 | 0.83 | -5.26E-09 | 4.25E-09 |
| | CBDfemale | 2.29E-09 | 0.79 | -3.96E-09 | 5.19E-09 |
| | THCfemale | 2.22E-09 | 0.66 | -3.45E-09 | 5.42E-09 |
| | VehMale | 2.29E-09 | 0.11 | -8.28E-09 | 8.74E-10 |
| | CBDmale | 2.11E-09 | 0.01 | -1.01E-08 | -1.65E-09 |
| | THCmale | 2.22E-09 | 0.74 | -5.18E-09 | 3.69E-09 |
| | TpCmale | 2.16E-09 | 0.01 | -1.01E-08 | -1.45E-09 |
| VehMale | VehFemale | 2.50E-09 | 0.21 | -1.80E-09 | 8.19E-09 |
| | CBDfemale | 2.42E-09 | 0.08 | -5.07E-10 | 9.15E-09 |
| | THCfemale | 2.35E-09 | 0.05 | -3.58E-12 | 9.38E-09 |
| | TpCfemale | 2.29E-09 | 0.11 | -8.74E-10 | 8.28E-09 |
| | CBDmale | 2.25E-09 | 0.34 | -6.64E-09 | 2.33E-09 |
| | THCmale | 2.35E-09 | 0.21 | -1.73E-09 | 7.65E-09 |
| | TpCmale | 2.29E-09 | 0.37 | -6.64E-09 | 2.52E-09 |
| CBDmale | VehFemale | 2.34E-09 | 0.03 | 6.89E-10 | 1.00E-08 |
| | CBDfemale | 2.25E-09 | 0.01 | 1.99E-09 | 1.10E-08 |
| | THCfemale | 2.17E-09 | 0.00 | 2.51E-09 | 1.12E-08 |
| | TpCfemale | 2.11E-09 | 0.01 | 1.65E-09 | 1.01E-08 |
| | VehMale | 2.25E-09 | 0.34 | -2.33E-09 | 6.64E-09 |
| | THCmale | 2.17E-09 | 0.02 | 7.81E-10 | 9.46E-09 |
| | TpCmale | 2.11E-09 | 0.96 | -4.12E-09 | 4.32E-09 |
| THCmale | VehFemale | 2.44E-09 | 0.92 | -4.63E-09 | 5.10E-09 |
| | CBDfemale | 2.35E-09 | 0.57 | -3.33E-09 | 6.05E-09 |
| | THCfemale | 2.28E-09 | 0.45 | -2.83E-09 | 6.28E-09 |
| | TpCfemale | 2.22E-09 | 0.74 | -3.69E-09 | 5.18E-09 |
| | VehMale | 2.35E-09 | 0.21 | -7.65E-09 | 1.73E-09 |
| | CBDmale | 2.17E-09 | 0.02 | -9.46E-09 | -7.81E-10 |
| | TpCmale | 2.22E-09 | 0.03 | -9.46E-09 | -5.87E-10 |
| TpCmale | VehFemale | 2.38E-09 | 0.03 | 5.01E-10 | 1.00E-08 |
| | CBDfemale | 2.29E-09 | 0.01 | 1.80E-09 | 1.10E-08 |
| | THCfemale | 2.22E-09 | 0.00 | 2.31E-09 | 1.12E-08 |
| | TpCfemale | 2.16E-09 | 0.01 | 1.45E-09 | 1.01E-08 |
| | VehMale | 2.29E-09 | 0.37 | -2.52E-09 | 6.64E-09 |
| | CBDmale | 2.11E-09 | 0.96 | -4.32E-09 | 4.12E-09 |
| | THCmale | 2.22E-09 | 0.03 | 5.87E-10 | 9.46E-09 |

Supplemental Table 5.8 Fisher's LSD post-hoc test for liver free fatty acid ANOVA

| Dependent Variable | (I) drugVsex | | Std. Error | Sig. | 95% Confidence Interval Lower Bound | Upper Bound |
|---|---|---|---|---|---|---|
| Oleic acid | VehFemale | CBDfemale | 1.25512E-10 | 0.582 | -3.2041E-10 | 1.8138E-10 |
| | | THCfemale | 1.21976E-10 | 0.018 | -5.4032E-10 | -5.2662E-11 |
| | | TpCfemale | 1.19071E-10 | 0.005 | -5.8825E-10 | -1.1221E-10 |
| | | VehMale | 1.29918E-10 | 0.079 | -4.9153E-10 | 2.7877E-11 |
| | | CBDmale | 1.19071E-10 | 0.091 | -4.4262E-10 | 3.3420E-11 |
| | | THCmale | 1.25512E-10 | 0.372 | -3.6384E-10 | 1.3795E-10 |
| | | TpCmale | 1.19071E-10 | 0.009 | -5.6010E-10 | -8.4056E-11 |
| | CBDfemale | VehFemale | 1.25512E-10 | 0.582 | -1.8138E-10 | 3.2041E-10 |
| | | THCfemale | 1.21976E-10 | 0.068 | -4.7080E-10 | 1.6850E-11 |
| | | TpCfemale | 1.19071E-10 | 0.022 | -5.1874E-10 | -4.2699E-11 |
| | | VehMale | 1.29918E-10 | 0.216 | -4.2201E-10 | 9.7389E-11 |
| | | CBDmale | 1.19071E-10 | 0.261 | -3.7311E-10 | 1.0293E-10 |
| | | THCmale | 1.25512E-10 | 0.730 | -2.9433E-10 | 2.0746E-10 |
| | | TpCmale | 1.19071E-10 | 0.038 | -4.9058E-10 | -1.4544E-11 |
| | THCfemale | VehFemale | 1.21976E-10 | 0.018 | 5.2662E-11 | 5.4032E-10 |
| | | CBDfemale | 1.21976E-10 | 0.068 | -1.6850E-11 | 4.7080E-10 |
| | | TpCfemale | 1.15338E-10 | 0.643 | -2.8430E-10 | 1.7681E-10 |
| | | VehMale | 1.26504E-10 | 0.611 | -1.8821E-10 | 3.1754E-10 |
| | | CBDmale | 1.15338E-10 | 0.429 | -1.3867E-10 | 3.2244E-10 |
| | | THCmale | 1.21976E-10 | 0.137 | -6.0286E-11 | 4.2737E-10 |
| | | TpCmale | 1.15338E-10 | 0.825 | -2.5614E-10 | 2.0497E-10 |
| | TpCfemale | VehFemale | 1.19071E-10 | 0.005 | 1.1221E-10 | 5.8825E-10 |
| | | CBDfemale | 1.19071E-10 | 0.022 | 4.2699E-11 | 5.1874E-10 |
| | | THCfemale | 1.15338E-10 | 0.643 | -1.7681E-10 | 2.8430E-10 |
| | | VehMale | 1.23706E-10 | 0.342 | -1.2888E-10 | 3.6569E-10 |
| | | CBDmale | 1.12262E-10 | 0.199 | -7.8777E-11 | 3.7004E-10 |
| | | THCmale | 1.19071E-10 | 0.051 | -7.3744E-13 | 4.7530E-10 |
| | | TpCmale | 1.12262E-10 | 0.803 | -1.9625E-10 | 2.5256E-10 |
| | VehMale | VehFemale | 1.29918E-10 | 0.079 | -2.7877E-11 | 4.9153E-10 |
| | | CBDfemale | 1.29918E-10 | 0.216 | -9.7389E-11 | 4.2201E-10 |

| | | | | | | |
|---|---|---|---|---|---|---|
| | | THCfemale | 1.26504E-10 | 0.611 | -3.1754E-10 | 1.8821E-10 |
| | | TpCfemale | 1.23706E-10 | 0.342 | -3.6569E-10 | 1.2888E-10 |
| | | CBDmale | 1.23706E-10 | 0.827 | -2.2006E-10 | 2.7451E-10 |
| | | THCmale | 1.29918E-10 | 0.364 | -1.4083E-10 | 3.7858E-10 |
| | | TpCmale | 1.23706E-10 | 0.468 | -3.3754E-10 | 1.5703E-10 |
| | CBDmale | VehFemale | 1.19071E-10 | 0.091 | -3.3420E-11 | 4.4262E-10 |
| | | CBDfemale | 1.19071E-10 | 0.261 | -1.0293E-10 | 3.7311E-10 |
| | | THCfemale | 1.15338E-10 | 0.429 | -3.2244E-10 | 1.3867E-10 |
| | | TpCfemale | 1.12262E-10 | 0.199 | -3.7004E-10 | 7.8777E-11 |
| | | VehMale | 1.23706E-10 | 0.827 | -2.7451E-10 | 2.2006E-10 |
| | | THCmale | 1.19071E-10 | 0.444 | -1.4637E-10 | 3.2967E-10 |
| | | TpCmale | 1.12262E-10 | 0.299 | -3.4188E-10 | 1.0693E-10 |
| | THCmale | VehFemale | 1.25512E-10 | 0.372 | -1.3795E-10 | 3.6384E-10 |
| | | CBDfemale | 1.25512E-10 | 0.730 | -2.0746E-10 | 2.9433E-10 |
| | | THCfemale | 1.21976E-10 | 0.137 | -4.2737E-10 | 6.0286E-11 |
| | | TpCfemale | 1.19071E-10 | 0.051 | -4.7530E-10 | 7.3744E-13 |
| | | VehMale | 1.29918E-10 | 0.364 | -3.7858E-10 | 1.4083E-10 |
| | | CBDmale | 1.19071E-10 | 0.444 | -3.2967E-10 | 1.4637E-10 |
| | | TpCmale | 1.19071E-10 | 0.084 | -4.4715E-10 | 2.8893E-11 |
| | TpCmale | VehFemale | 1.19071E-10 | 0.009 | 8.4056E-11 | 5.6010E-10 |
| | | CBDfemale | 1.19071E-10 | 0.038 | 1.4544E-11 | 4.9058E-10 |
| | | THCfemale | 1.15338E-10 | 0.825 | -2.0497E-10 | 2.5614E-10 |
| | | TpCfemale | 1.12262E-10 | 0.803 | -2.5256E-10 | 1.9625E-10 |
| | | VehMale | 1.23706E-10 | 0.468 | -1.5703E-10 | 3.3754E-10 |
| | | CBDmale | 1.12262E-10 | 0.299 | -1.0693E-10 | 3.4188E-10 |
| | | THCmale | 1.19071E-10 | 0.084 | -2.8893E-11 | 4.4715E-10 |
| | VehFemale | CBDfemale | 3.01855E-10 | 0.633 | -4.5852E-10 | 7.4790E-10 |
| | | THCfemale | 2.76655E-10 | 0.061 | -1.0812E-09 | 2.4537E-11 |
| | | TpCfemale | 2.76655E-10 | 0.000 | -1.9681E-09 | -8.6237E-10 |
| Linoleic acid | | VehMale | 3.01855E-10 | 0.692 | -7.2326E-10 | 4.8316E-10 |
| | | CBDmale | 2.76655E-10 | 0.594 | -7.0089E-10 | 4.0481E-10 |
| | | THCmale | 2.83403E-10 | 0.010 | -1.3209E-09 | -1.8820E-10 |

| | | | | | |
|---|---|---|---|---|---|
| | TpCmale | 2.76655E-10 | 0.001 | -1.5107E-09 | -4.0504E-10 |
| CBDfemale | VehFemale | 3.01855E-10 | 0.633 | -7.4790E-10 | 4.5852E-10 |
| | THCfemale | 2.87423E-10 | 0.022 | -1.2474E-09 | -9.8636E-11 |
| | TpCfemale | 2.87423E-10 | 0.000 | -2.1343E-09 | -9.8554E-10 |
| | VehMale | 3.11754E-10 | 0.399 | -8.8773E-10 | 3.5825E-10 |
| | CBDmale | 2.87423E-10 | 0.312 | -8.6710E-10 | 2.8164E-10 |
| | THCmale | 2.93925E-10 | 0.003 | -1.4866E-09 | -3.1186E-10 |
| | TpCmale | 2.87423E-10 | 0.000 | -1.6770E-09 | -5.2821E-10 |
| THCfemale | VehFemale | 2.76655E-10 | 0.061 | -2.4537E-11 | 1.0812E-09 |
| | CBDfemale | 2.87423E-10 | 0.022 | 9.8636E-11 | 1.2474E-09 |
| | TpCfemale | 2.60832E-10 | 0.001 | -1.4081E-09 | -3.6567E-10 |
| | VehMale | 2.87423E-10 | 0.160 | -1.6611E-10 | 9.8263E-10 |
| | CBDmale | 2.60832E-10 | 0.150 | -1.4096E-10 | 9.0151E-10 |
| | THCmale | 2.67980E-10 | 0.402 | -7.6173E-10 | 3.0930E-10 |
| | TpCmale | 2.60832E-10 | 0.105 | -9.5081E-10 | 9.1654E-11 |
| TpCfemale | VehFemale | 2.76655E-10 | 0.000 | 8.6237E-10 | 1.9681E-09 |
| | CBDfemale | 2.87423E-10 | 0.000 | 9.8554E-10 | 2.1343E-09 |
| | THCfemale | 2.60832E-10 | 0.001 | 3.6567E-10 | 1.4081E-09 |
| | VehMale | 2.87423E-10 | 0.000 | 7.2080E-10 | 1.8695E-09 |
| | CBDmale | 2.60832E-10 | 0.000 | 7.4595E-10 | 1.7884E-09 |
| | THCmale | 2.67980E-10 | 0.016 | 1.2517E-10 | 1.1962E-09 |
| | TpCmale | 2.60832E-10 | 0.084 | -6.3906E-11 | 9.7856E-10 |
| VehMale | VehFemale | 3.01855E-10 | 0.692 | -4.8316E-10 | 7.2326E-10 |
| | CBDfemale | 3.11754E-10 | 0.399 | -3.5825E-10 | 8.8773E-10 |
| | THCfemale | 2.87423E-10 | 0.160 | -9.8263E-10 | 1.6611E-10 |
| | TpCfemale | 2.87423E-10 | 0.000 | -1.8695E-09 | -7.2080E-10 |
| | CBDmale | 2.87423E-10 | 0.923 | -6.0236E-10 | 5.4638E-10 |
| | THCmale | 2.93925E-10 | 0.035 | -1.2218E-09 | -4.7122E-11 |
| | TpCmale | 2.87423E-10 | 0.005 | -1.4122E-09 | -2.6347E-10 |
| CBDmale | VehFemale | 2.76655E-10 | 0.594 | -4.0481E-10 | 7.0089E-10 |
| | CBDfemale | 2.87423E-10 | 0.312 | -2.8164E-10 | 8.6710E-10 |
| | THCfemale | 2.60832E-10 | 0.150 | -9.0151E-10 | 1.4096E-10 |

| | | | | | | |
|---|---|---|---|---|---|---|
| | | TpCfemale | 2.60832E-10 | 0.000 | -1.7884E-09 | -7.4595E-10 |
| | | VehMale | 2.87423E-10 | 0.923 | -5.4638E-10 | 6.0236E-10 |
| | | THCmale | 2.67980E-10 | 0.027 | -1.1420E-09 | -7.0978E-11 |
| | | TpCmale | 2.60832E-10 | 0.003 | -1.3311E-09 | -2.8862E-10 |
| | THCmale | VehFemale | 2.83403E-10 | 0.010 | 1.8820E-10 | 1.3209E-09 |
| | | CBDfemale | 2.93925E-10 | 0.003 | 3.1186E-10 | 1.4866E-09 |
| | | THCfemale | 2.67980E-10 | 0.402 | -3.0930E-10 | 7.6173E-10 |
| | | TpCfemale | 2.67980E-10 | 0.016 | -1.1962E-09 | -1.2517E-10 |
| | | VehMale | 2.93925E-10 | 0.035 | 4.7122E-11 | 1.2218E-09 |
| | | CBDmale | 2.67980E-10 | 0.027 | 7.0978E-11 | 1.1420E-09 |
| | | TpCmale | 2.67980E-10 | 0.451 | -7.3887E-10 | 3.3216E-10 |
| | TpCmale | VehFemale | 2.76655E-10 | 0.001 | 4.0504E-10 | 1.5107E-09 |
| | | CBDfemale | 2.87423E-10 | 0.000 | 5.2821E-10 | 1.6770E-09 |
| | | THCfemale | 2.60832E-10 | 0.105 | -9.1654E-11 | 9.5081E-10 |
| | | TpCfemale | 2.60832E-10 | 0.084 | -9.7856E-10 | 6.3906E-11 |
| | | VehMale | 2.87423E-10 | 0.005 | 2.6347E-10 | 1.4122E-09 |
| | | CBDmale | 2.60832E-10 | 0.003 | 2.8862E-10 | 1.3311E-09 |
| | | THCmale | 2.67980E-10 | 0.451 | -3.3216E-10 | 7.3887E-10 |
| | VehFemale | CBDfemale | 8.59647E-10 | 0.091 | -3.1929E-09 | 2.4174E-10 |
| | | THCfemale | 8.18547E-10 | 0.000 | -4.8106E-09 | -1.5401E-09 |
| | | TpCfemale | 8.18547E-10 | 0.000 | -5.6715E-09 | -2.4011E-09 |
| | | VehMale | 8.87839E-10 | 0.006 | -4.3172E-09 | -7.6984E-10 |
| | | CBDmale | 8.18547E-10 | 0.003 | -4.1333E-09 | -8.6278E-10 |
| | | THCmale | 8.37063E-10 | 0.000 | -5.3620E-09 | -2.0176E-09 |
| | | TpCmale | 8.03081E-10 | 0.000 | -5.8685E-09 | -2.6598E-09 |
| Arachidonic acid | CBDfemale | VehFemale | 8.59647E-10 | 0.091 | -2.4174E-10 | 3.1929E-09 |
| | | THCfemale | 7.87879E-10 | 0.035 | -3.2737E-09 | -1.2578E-10 |
| | | TpCfemale | 7.87879E-10 | 0.002 | -4.1347E-09 | -9.8673E-10 |
| | | VehMale | 8.59647E-10 | 0.219 | -2.7852E-09 | 6.4944E-10 |
| | | CBDmale | 7.87879E-10 | 0.199 | -2.5964E-09 | 5.5155E-10 |
| | | THCmale | 8.07098E-10 | 0.008 | -3.8265E-09 | -6.0181E-10 |
| | | TpCmale | 7.71798E-10 | 0.001 | -4.3304E-09 | -1.2467E-09 |

| | | | | | |
|---|---|---|---|---|---|
| THCfemale | VehFemale | 8.18547E-10 | 0.000 | 1.5401E-09 | 4.8106E-09 |
| | CBDfemale | 7.87879E-10 | 0.035 | 1.2578E-10 | 3.2737E-09 |
| | TpCfemale | 7.42820E-10 | 0.251 | -2.3449E-09 | 6.2300E-10 |
| | VehMale | 8.18547E-10 | 0.443 | -1.0034E-09 | 2.2671E-09 |
| | CBDmale | 7.42820E-10 | 0.365 | -8.0662E-10 | 2.1613E-09 |
| | THCmale | 7.63174E-10 | 0.503 | -2.0390E-09 | 1.0102E-09 |
| | TpCmale | 7.25741E-10 | 0.138 | -2.5386E-09 | 3.6103E-10 |
| TpCfemale | VehFemale | 8.18547E-10 | 0.000 | 2.4011E-09 | 5.6715E-09 |
| | CBDfemale | 7.87879E-10 | 0.002 | 9.8673E-10 | 4.1347E-09 |
| | THCfemale | 7.42820E-10 | 0.251 | -6.2300E-10 | 2.3449E-09 |
| | VehMale | 8.18547E-10 | 0.073 | -1.4244E-10 | 3.1280E-09 |
| | CBDmale | 7.42820E-10 | 0.042 | 5.4333E-11 | 3.0222E-09 |
| | THCmale | 7.63174E-10 | 0.651 | -1.1781E-09 | 1.8711E-09 |
| | TpCmale | 7.25741E-10 | 0.755 | -1.6777E-09 | 1.2220E-09 |
| VehMale | VehFemale | 8.87839E-10 | 0.006 | 7.6984E-10 | 4.3172E-09 |
| | CBDfemale | 8.59647E-10 | 0.219 | -6.4944E-10 | 2.7852E-09 |
| | THCfemale | 8.18547E-10 | 0.443 | -2.2671E-09 | 1.0034E-09 |
| | TpCfemale | 8.18547E-10 | 0.073 | -3.1280E-09 | 1.4244E-10 |
| | CBDmale | 8.18547E-10 | 0.956 | -1.5897E-09 | 1.6807E-09 |
| | THCmale | 8.37063E-10 | 0.176 | -2.8185E-09 | 5.2595E-10 |
| | TpCmale | 8.03081E-10 | 0.036 | -3.3250E-09 | -1.1631E-10 |
| CBDmale | VehFemale | 8.18547E-10 | 0.003 | 8.6278E-10 | 4.1333E-09 |
| | CBDfemale | 7.87879E-10 | 0.199 | -5.5155E-10 | 2.5964E-09 |
| | THCfemale | 7.42820E-10 | 0.365 | -2.1613E-09 | 8.0662E-10 |
| | TpCfemale | 7.42820E-10 | 0.042 | -3.0222E-09 | -5.4333E-11 |
| | VehMale | 8.18547E-10 | 0.956 | -1.6807E-09 | 1.5897E-09 |
| | THCmale | 7.63174E-10 | 0.123 | -2.7164E-09 | 3.3286E-10 |
| | TpCmale | 7.25741E-10 | 0.018 | -3.2160E-09 | -3.1630E-10 |
| THCmale | VehFemale | 8.37063E-10 | 0.000 | 2.0176E-09 | 5.3620E-09 |
| | CBDfemale | 8.07098E-10 | 0.008 | 6.0181E-10 | 3.8265E-09 |
| | THCfemale | 7.63174E-10 | 0.503 | -1.0102E-09 | 2.0390E-09 |
| | TpCfemale | 7.63174E-10 | 0.651 | -1.8711E-09 | 1.1781E-09 |

| | | | | | | |
|---|---|---|---|---|---|---|
| | | VehMale | 8.37063E-10 | 0.176 | -5.2595E-10 | 2.8185E-09 |
| | | CBDmale | 7.63174E-10 | 0.123 | -3.3286E-10 | 2.7164E-09 |
| | | TpCmale | 7.46562E-10 | 0.445 | -2.0658E-09 | 9.1705E-10 |
| | TpCmale | VehFemale | 8.03081E-10 | 0.000 | 2.6598E-09 | 5.8685E-09 |
| | | CBDfemale | 7.71798E-10 | 0.001 | 1.2467E-09 | 4.3304E-09 |
| | | THCfemale | 7.25741E-10 | 0.138 | -3.6103E-10 | 2.5386E-09 |
| | | TpCfemale | 7.25741E-10 | 0.755 | -1.2220E-09 | 1.6777E-09 |
| | | VehMale | 8.03081E-10 | 0.036 | 1.1631E-10 | 3.3250E-09 |
| | | CBDmale | 7.25741E-10 | 0.018 | 3.1630E-10 | 3.2160E-09 |
| | | THCmale | 7.46562E-10 | 0.445 | -9.1705E-10 | 2.0658E-09 |
| | VehFemale | CBDfemale | 3.78894E-10 | 0.028 | -1.6106E-09 | -9.5338E-11 |
| | | THCfemale | 3.68219E-10 | 0.003 | -1.8567E-09 | -3.8414E-10 |
| | | TpCfemale | 3.59450E-10 | 0.000 | -2.7206E-09 | -1.2831E-09 |
| | | VehMale | 4.09252E-10 | 0.164 | -1.3945E-09 | 2.4221E-10 |
| | | CBDmale | 3.68219E-10 | 0.038 | -1.5162E-09 | -4.3591E-11 |
| | | THCmale | 3.78894E-10 | 0.000 | -2.2031E-09 | -6.8778E-10 |
| | | TpCmale | 3.52114E-10 | 0.000 | -3.2205E-09 | -1.8123E-09 |
| | CBDfemale | VehFemale | 3.78894E-10 | 0.028 | 9.5338E-11 | 1.6106E-09 |
| | | THCfemale | 3.68219E-10 | 0.470 | -1.0038E-09 | 4.6884E-10 |
| | | TpCfemale | 3.59450E-10 | 0.002 | -1.8676E-09 | -4.3009E-10 |
| | | VehMale | 4.09252E-10 | 0.501 | -5.4151E-10 | 1.0952E-09 |
| Docosahexaenoic acid | | CBDmale | 3.68219E-10 | 0.843 | -6.6321E-10 | 8.0939E-10 |
| | | THCmale | 3.78894E-10 | 0.123 | -1.3501E-09 | 1.6521E-10 |
| | | TpCmale | 3.52114E-10 | 0.000 | -2.3675E-09 | -9.5935E-10 |
| | THCfemale | VehFemale | 3.68219E-10 | 0.003 | 3.8414E-10 | 1.8567E-09 |
| | | CBDfemale | 3.68219E-10 | 0.470 | -4.6884E-10 | 1.0038E-09 |
| | | TpCfemale | 3.48180E-10 | 0.014 | -1.5776E-09 | -1.8517E-10 |
| | | VehMale | 3.99389E-10 | 0.178 | -2.5432E-10 | 1.3429E-09 |
| | | CBDmale | 3.57225E-10 | 0.344 | -3.7376E-10 | 1.0549E-09 |
| | | THCmale | 3.68219E-10 | 0.381 | -1.0613E-09 | 4.1132E-10 |
| | | TpCmale | 3.40600E-10 | 0.000 | -2.0771E-09 | -7.1492E-10 |
| | TpCfemale | VehFemale | 3.59450E-10 | 0.000 | 1.2831E-09 | 2.7206E-09 |

| | | | | | |
|---|---|---|---|---|---|
| | CBDfemale | 3.59450E-10 | 0.002 | 4.3009E-10 | 1.8676E-09 |
| | THCfemale | 3.48180E-10 | 0.014 | 1.8517E-10 | 1.5776E-09 |
| | VehMale | 3.91320E-10 | 0.001 | 6.4321E-10 | 2.2082E-09 |
| | CBDmale | 3.48180E-10 | 0.001 | 5.2572E-10 | 1.9182E-09 |
| | THCmale | 3.59450E-10 | 0.127 | -1.6235E-10 | 1.2752E-09 |
| | TpCmale | 3.31101E-10 | 0.125 | -1.1767E-09 | 1.4749E-10 |
| VehMale | VehFemale | 4.09252E-10 | 0.164 | -2.4221E-10 | 1.3945E-09 |
| | CBDfemale | 4.09252E-10 | 0.501 | -1.0952E-09 | 5.4151E-10 |
| | THCfemale | 3.99389E-10 | 0.178 | -1.3429E-09 | 2.5432E-10 |
| | TpCfemale | 3.91320E-10 | 0.001 | -2.2082E-09 | -6.4321E-10 |
| | CBDmale | 3.99389E-10 | 0.612 | -1.0024E-09 | 5.9488E-10 |
| | THCmale | 4.09252E-10 | 0.038 | -1.6876E-09 | -5.0932E-11 |
| | TpCmale | 3.84592E-10 | 0.000 | -2.7093E-09 | -1.1713E-09 |
| CBDmale | VehFemale | 3.68219E-10 | 0.038 | 4.3591E-11 | 1.5162E-09 |
| | CBDfemale | 3.68219E-10 | 0.843 | -8.0939E-10 | 6.6321E-10 |
| | THCfemale | 3.57225E-10 | 0.344 | -1.0549E-09 | 3.7376E-10 |
| | TpCfemale | 3.48180E-10 | 0.001 | -1.9182E-09 | -5.2572E-10 |
| | VehMale | 3.99389E-10 | 0.612 | -5.9488E-10 | 1.0024E-09 |
| | THCmale | 3.68219E-10 | 0.076 | -1.4018E-09 | 7.0766E-11 |
| | TpCmale | 3.40600E-10 | 0.000 | -2.4176E-09 | -1.0555E-09 |
| THCmale | VehFemale | 3.78894E-10 | 0.000 | 6.8778E-10 | 2.2031E-09 |
| | CBDfemale | 3.78894E-10 | 0.123 | -1.6521E-10 | 1.3501E-09 |
| | THCfemale | 3.68219E-10 | 0.381 | -4.1132E-10 | 1.0613E-09 |
| | TpCfemale | 3.59450E-10 | 0.127 | -1.2752E-09 | 1.6235E-10 |
| | VehMale | 4.09252E-10 | 0.038 | 5.0932E-11 | 1.6876E-09 |
| | CBDmale | 3.68219E-10 | 0.076 | -7.0766E-11 | 1.4018E-09 |
| | TpCmale | 3.52114E-10 | 0.003 | -1.7751E-09 | -3.6691E-10 |
| TpCmale | VehFemale | 3.52114E-10 | 0.000 | 1.8123E-09 | 3.2205E-09 |
| | CBDfemale | 3.52114E-10 | 0.000 | 9.5935E-10 | 2.3675E-09 |
| | THCfemale | 3.40600E-10 | 0.000 | 7.1492E-10 | 2.0771E-09 |
| | TpCfemale | 3.31101E-10 | 0.125 | -1.4749E-10 | 1.1767E-09 |
| | VehMale | 3.84592E-10 | 0.000 | 1.1713E-09 | 2.7093E-09 |

| | | | | | | |
|---|---|---|---|---|---|---|
| | | CBDmale | 3.40600E-10 | 0.000 | 1.0555E-09 | 2.4176E-09 |
| | | THCmale | 3.52114E-10 | 0.003 | 3.6691E-10 | 1.7751E-09 |
| | VehFemale | CBDfemale | 2.86905E-10 | 0.916 | -5.4306E-10 | 6.0397E-10 |
| | | THCfemale | 2.69367E-10 | 0.206 | -8.8236E-10 | 1.9455E-10 |
| | | TpCfemale | 2.62953E-10 | 0.002 | -1.3635E-09 | -3.1222E-10 |
| | | VehMale | 2.86905E-10 | 0.064 | -1.1153E-09 | 3.1731E-11 |
| | | CBDmale | 2.69367E-10 | 0.104 | -9.8324E-10 | 9.3673E-11 |
| | | THCmale | 2.69367E-10 | 0.000 | -1.7357E-09 | -6.5878E-10 |
| | | TpCmale | 2.57586E-10 | 0.000 | -2.5615E-09 | -1.5317E-09 |
| | CBDfemale | VehFemale | 2.86905E-10 | 0.916 | -6.0397E-10 | 5.4306E-10 |
| | | THCfemale | 2.79368E-10 | 0.185 | -9.3281E-10 | 1.8408E-10 |
| | | TpCfemale | 2.73188E-10 | 0.002 | -1.4144E-09 | -3.2221E-10 |
| | | VehMale | 2.96314E-10 | 0.058 | -1.1646E-09 | 2.0084E-11 |
| | | CBDmale | 2.79368E-10 | 0.094 | -1.0337E-09 | 8.3207E-11 |
| | | THCmale | 2.79368E-10 | 0.000 | -1.7861E-09 | -6.6924E-10 |
| | | TpCmale | 2.68026E-10 | 0.000 | -2.6129E-09 | -1.5413E-09 |
| | THCfemale | VehFemale | 2.69367E-10 | 0.206 | -1.9455E-10 | 8.8236E-10 |
| Eicosapentaenoic acid | | CBDfemale | 2.79368E-10 | 0.185 | -1.8408E-10 | 9.3281E-10 |
| | | TpCfemale | 2.54707E-10 | 0.057 | -1.0031E-09 | 1.5207E-11 |
| | | VehMale | 2.79368E-10 | 0.481 | -7.5632E-10 | 3.6057E-10 |
| | | CBDmale | 2.61324E-10 | 0.701 | -6.2326E-10 | 4.2150E-10 |
| | | THCmale | 2.61324E-10 | 0.002 | -1.3757E-09 | -3.3095E-10 |
| | | TpCmale | 2.49163E-10 | 0.000 | -2.2008E-09 | -1.2047E-09 |
| | TpCfemale | VehFemale | 2.62953E-10 | 0.002 | 3.1222E-10 | 1.3635E-09 |
| | | CBDfemale | 2.73188E-10 | 0.002 | 3.2221E-10 | 1.4144E-09 |
| | | THCfemale | 2.54707E-10 | 0.057 | -1.5207E-11 | 1.0031E-09 |
| | | VehMale | 2.73188E-10 | 0.283 | -2.5003E-10 | 8.4216E-10 |
| | | CBDmale | 2.54707E-10 | 0.128 | -1.1608E-10 | 9.0222E-10 |
| | | THCmale | 2.54707E-10 | 0.163 | -8.6853E-10 | 1.4977E-10 |
| | | TpCmale | 2.42214E-10 | 0.000 | -1.6930E-09 | -7.2461E-10 |
| | VehMale | VehFemale | 2.86905E-10 | 0.064 | -3.1731E-11 | 1.1153E-09 |
| | | CBDfemale | 2.96314E-10 | 0.058 | -2.0084E-11 | 1.1646E-09 |

| | | | | | |
|---|---|---|---|---|---|
| | THCfemale | 2.79368E-10 | 0.481 | -3.6057E-10 | 7.5632E-10 |
| | TpCfemale | 2.73188E-10 | 0.283 | -8.4216E-10 | 2.5003E-10 |
| | CBDmale | 2.79368E-10 | 0.730 | -4.6145E-10 | 6.5545E-10 |
| | THCmale | 2.79368E-10 | 0.022 | -1.2139E-09 | -9.7002E-11 |
| | TpCmale | 2.68026E-10 | 0.000 | -2.0406E-09 | -9.6908E-10 |
| CBDmale | VehFemale | 2.69367E-10 | 0.104 | -9.3673E-11 | 9.8324E-10 |
| | CBDfemale | 2.79368E-10 | 0.094 | -8.3207E-11 | 1.0337E-09 |
| | THCfemale | 2.61324E-10 | 0.701 | -4.2150E-10 | 6.2326E-10 |
| | TpCfemale | 2.54707E-10 | 0.128 | -9.0222E-10 | 1.1608E-10 |
| | VehMale | 2.79368E-10 | 0.730 | -6.5545E-10 | 4.6145E-10 |
| | THCmale | 2.61324E-10 | 0.005 | -1.2748E-09 | -2.3007E-10 |
| | TpCmale | 2.49163E-10 | 0.000 | -2.0999E-09 | -1.1038E-09 |
| THCmale | VehFemale | 2.69367E-10 | 0.000 | 6.5878E-10 | 1.7357E-09 |
| | CBDfemale | 2.79368E-10 | 0.000 | 6.6924E-10 | 1.7861E-09 |
| | THCfemale | 2.61324E-10 | 0.002 | 3.3095E-10 | 1.3757E-09 |
| | TpCfemale | 2.54707E-10 | 0.163 | -1.4977E-10 | 8.6853E-10 |
| | VehMale | 2.79368E-10 | 0.022 | 9.7002E-11 | 1.2139E-09 |
| | CBDmale | 2.61324E-10 | 0.005 | 2.3007E-10 | 1.2748E-09 |
| | TpCmale | 2.49163E-10 | 0.001 | -1.3475E-09 | -3.5134E-10 |
| TpCmale | VehFemale | 2.57586E-10 | 0.000 | 1.5317E-09 | 2.5615E-09 |
| | CBDfemale | 2.68026E-10 | 0.000 | 1.5413E-09 | 2.6129E-09 |
| | THCfemale | 2.49163E-10 | 0.000 | 1.2047E-09 | 2.2008E-09 |
| | TpCfemale | 2.42214E-10 | 0.000 | 7.2461E-10 | 1.6930E-09 |
| | VehMale | 2.68026E-10 | 0.000 | 9.6908E-10 | 2.0406E-09 |
| | CBDmale | 2.49163E-10 | 0.000 | 1.1038E-09 | 2.0999E-09 |
| | THCmale | 2.49163E-10 | 0.001 | 3.5134E-10 | 1.3475E-09 |

Supplemental Table 6.1 Descriptive statistics for *N*-acyl phenylalanines (mol/L) in plasma collected from male and female adult mice 2h after receiving an i.p. injection of 10mg/kg CBD, THC, or THC+CBD

| | | N | Mean | Std. Deviation | Std. Error | 95% Confidence Interval for Mean | | Minimum | Maximum |
|---|---|---|---|---|---|---|---|---|---|
| | | | | | | Lower Bound | Upper Bound | | |
| *N*-palmitoyl phenylalanine | Veh F | 7 | 4.2419E-09 | 2.41742E-09 | 9.13700E-10 | 2.0062E-09 | 6.4777E-09 | 1.76E-09 | 8.68E-09 |
| | CBD F | 8 | 2.8001E-09 | 1.64771E-09 | 5.82554E-10 | 1.4226E-09 | 4.1776E-09 | 1.28E-09 | 6.02E-09 |
| | THC F | 9 | 2.7470E-09 | 1.25763E-09 | 4.19211E-10 | 1.7803E-09 | 3.7137E-09 | 8.62E-10 | 5.25E-09 |
| | THC+CBD F | 10 | 4.5160E-09 | 2.61848E-09 | 8.28036E-10 | 2.6428E-09 | 6.3891E-09 | 1.97E-09 | 9.31E-09 |
| | Veh M | 8 | 3.6007E-09 | 2.29429E-09 | 8.11156E-10 | 1.6827E-09 | 5.5188E-09 | 8.62E-10 | 7.38E-09 |
| | CBD M | 11 | 3.3432E-09 | 2.24792E-09 | 6.77773E-10 | 1.8330E-09 | 4.8534E-09 | 9.14E-10 | 7.00E-09 |
| | THC M | 9 | 5.2164E-09 | 3.46996E-09 | 1.15665E-09 | 2.5492E-09 | 7.8837E-09 | 1.48E-09 | 1.07E-08 |
| | THC+CBD M | 10 | 6.7420E-09 | 5.09872E-09 | 1.61236E-09 | 3.0946E-09 | 1.0389E-08 | 1.48E-09 | 1.44E-08 |
| *N*-stearoyl phenylalanine | Veh F | 7 | 1.7913E-09 | 1.93182E-09 | 7.30158E-10 | 4.6451E-12 | 3.5779E-09 | 2.38E-10 | 5.44E-09 |
| | CBD F | 8 | 7.0578E-10 | 3.33749E-10 | 1.17998E-10 | 4.2676E-10 | 9.8480E-10 | 2.86E-10 | 1.32E-09 |
| | THC F | 7 | 6.4112E-10 | 3.04585E-10 | 1.15122E-10 | 3.5943E-10 | 9.2282E-10 | 3.45E-10 | 1.23E-09 |
| | THC+CBD F | 10 | 1.1543E-09 | 6.28637E-10 | 1.98792E-10 | 7.0458E-10 | 1.6040E-09 | 3.04E-10 | 2.21E-09 |
| | Veh M | 6 | 9.7488E-10 | 7.10981E-10 | 2.90257E-10 | 2.2875E-10 | 1.7210E-09 | 3.44E-10 | 2.27E-09 |
| | CBD M | 10 | 1.0709E-09 | 5.67362E-10 | 1.79416E-10 | 6.6501E-10 | 1.4767E-09 | 5.40E-10 | 2.03E-09 |
| | THC M | 9 | 1.6203E-09 | 1.00307E-09 | 3.34357E-10 | 8.4924E-10 | 2.3913E-09 | 3.95E-10 | 2.95E-09 |
| | THC+CBD M | 10 | 2.0582E-09 | 1.35920E-09 | 4.29816E-10 | 1.0859E-09 | 3.0305E-09 | 4.07E-10 | 4.41E-09 |
| *N*-oleoyl phenylalanine | Veh F | 8 | 4.0565E-09 | 2.37645E-09 | 8.40202E-10 | 2.0698E-09 | 6.0433E-09 | 1.32E-09 | 8.28E-09 |
| | CBD F | 8 | 2.5702E-09 | 1.17462E-09 | 4.15292E-10 | 1.5882E-09 | 3.5523E-09 | 1.35E-09 | 4.14E-09 |
| | THC F | 10 | 2.2803E-09 | 1.04662E-09 | 3.30971E-10 | 1.5316E-09 | 3.0290E-09 | 1.14E-09 | 4.13E-09 |
| | THC+CBD F | 10 | 3.3725E-09 | 1.35526E-09 | 4.28570E-10 | 2.4030E-09 | 4.3420E-09 | 1.03E-09 | 5.15E-09 |
| | Veh M | 8 | 3.2532E-09 | 2.24628E-09 | 7.94180E-10 | 1.3753E-09 | 5.1311E-09 | 5.08E-10 | 6.76E-09 |
| | CBD M | 10 | 2.3656E-09 | 1.02347E-09 | 3.23649E-10 | 1.6335E-09 | 3.0978E-09 | 1.01E-09 | 4.10E-09 |
| | THC M | 9 | 3.1044E-09 | 1.53030E-09 | 5.10101E-10 | 1.9281E-09 | 4.2807E-09 | 1.12E-09 | 5.31E-09 |
| | THC+CBD M | 10 | 5.0983E-09 | 3.33769E-09 | 1.05547E-09 | 2.7107E-09 | 7.4860E-09 | 1.76E-09 | 1.13E-08 |
| *N*-linoleoyl phenylalanine | Veh F | 8 | 3.2730E-09 | 1.69954E-09 | 6.00876E-10 | 1.8521E-09 | 4.6938E-09 | 9.99E-10 | 5.71E-09 |
| | CBD F | 8 | 2.0024E-09 | 1.05591E-09 | 3.73319E-10 | 1.1196E-09 | 2.8852E-09 | 9.17E-10 | 3.61E-09 |
| | THC F | 9 | 1.9420E-09 | 9.34153E-10 | 3.11384E-10 | 1.2239E-09 | 2.6600E-09 | 6.14E-10 | 3.67E-09 |
| | THC+CBD F | 10 | 2.6719E-09 | 1.46085E-09 | 4.61960E-10 | 1.6269E-09 | 3.7169E-09 | 1.04E-09 | 5.24E-09 |
| | Veh M | 8 | 2.0659E-09 | 8.54271E-10 | 3.02030E-10 | 1.3517E-09 | 2.7801E-09 | 1.19E-09 | 3.54E-09 |
| | CBD M | 11 | 1.7685E-09 | 8.54321E-10 | 2.57588E-10 | 1.1945E-09 | 2.3424E-09 | 5.17E-10 | 3.14E-09 |
| | THC M | 9 | 2.5375E-09 | 1.09628E-09 | 3.65425E-10 | 1.6949E-09 | 3.3802E-09 | 1.09E-09 | 4.07E-09 |
| | THC+CBD M | 9 | 2.5845E-09 | 1.28070E-09 | 4.26901E-10 | 1.6001E-09 | 3.5689E-09 | 1.10E-09 | 4.47E-09 |
| *N*-arachidonoyl phenylalanine | Veh F | 7 | 1.0416E-09 | 4.63971E-10 | 1.75364E-10 | 6.1250E-10 | 1.4707E-09 | 3.13E-10 | 1.66E-09 |
| | CBD F | 7 | 4.6798E-10 | 1.67576E-10 | 6.33378E-11 | 3.1300E-10 | 6.2296E-10 | 2.17E-10 | 6.93E-10 |
| | THC F | 7 | 5.4570E-10 | 2.07946E-10 | 7.85961E-11 | 3.5339E-10 | 7.3802E-10 | 1.71E-10 | 8.19E-10 |
| | THC+CBD F | 10 | 7.8365E-10 | 4.15633E-10 | 1.31435E-10 | 4.8633E-10 | 1.0810E-09 | 2.81E-10 | 1.51E-09 |
| | Veh M | 6 | 5.4449E-10 | 2.55380E-10 | 1.04259E-10 | 2.7648E-10 | 8.1249E-10 | 2.63E-10 | 9.20E-10 |
| | CBD M | 9 | 5.0844E-10 | 1.35183E-10 | 4.50610E-11 | 4.0453E-10 | 6.1235E-10 | 2.63E-10 | 7.05E-10 |
| | THC M | 7 | 5.1104E-10 | 1.13228E-10 | 4.27961E-11 | 4.0632E-10 | 6.1576E-10 | 3.75E-10 | 6.81E-10 |
| | THC+CBD M | 9 | 5.9614E-10 | 3.66585E-10 | 1.22195E-10 | 3.1436E-10 | 8.7792E-10 | 1.86E-10 | 1.13E-09 |
| | Veh F | 5 | 6.8633E-10 | 3.17888E-10 | 1.42164E-10 | 2.9162E-10 | 1.0810E-09 | 4.38E-10 | 1.21E-09 |

| | | | | | | | | | |
|---|---|---|---|---|---|---|---|---|---|
| *N*-docosahexaenoyl phenylalanine | CBD F | 8 | 3.9863E-10 | 2.06946E-10 | 7.31666E-11 | 2.2562E-10 | 5.7164E-10 | 2.01E-10 | 7.27E-10 |
| | THC F | 8 | 4.4460E-10 | 2.55337E-10 | 9.02753E-11 | 2.3114E-10 | 6.5807E-10 | 2.35E-10 | 8.43E-10 |
| | THC+CBD F | 9 | 9.4948E-10 | 7.74338E-10 | 2.58113E-10 | 3.5427E-10 | 1.5447E-09 | 2.72E-10 | 2.36E-09 |
| | Veh M | 7 | 4.8020E-10 | 2.50275E-10 | 9.45949E-11 | 2.4874E-10 | 7.1167E-10 | 1.65E-10 | 9.21E-10 |
| | CBD M | 9 | 3.6853E-10 | 1.83485E-10 | 6.11617E-11 | 2.2749E-10 | 5.0956E-10 | 2.01E-10 | 7.25E-10 |
| | THC M | 7 | 6.6539E-10 | 2.59628E-10 | 9.81303E-11 | 4.2527E-10 | 9.0551E-10 | 3.65E-10 | 1.11E-09 |
| | THC+CBD M | 7 | 3.4921E-10 | 1.81090E-10 | 6.84454E-11 | 1.8173E-10 | 5.1669E-10 | 2.20E-10 | 7.52E-10 |

Supplemental Table 6.2 Descriptive statistics for *N*-acyl phenylalanines (mol/g) in liver collected from male and female adult mice 2h after receiving an i.p. injection of 10mg/kg CBD, THC, or THC+CBD

| | | N | Mean | Std. Deviation | Std. Error | 95% Confidence Interval for Mean | | Minimum | Maximum |
|---|---|---|---|---|---|---|---|---|---|
| | | | | | | Lower Bound | Upper Bound | | |
| *N*-palmitoyl phenylalanine | Veh F | 8 | 3.7691E-11 | 1.46756E-11 | 5.18861E-12 | 2.5422E-11 | 4.9960E-11 | 1.86E-11 | 5.71E-11 |
| | CBD F | 7 | 4.6477E-11 | 2.55925E-11 | 9.67304E-12 | 2.2807E-11 | 7.0146E-11 | 2.34E-11 | 9.10E-11 |
| | THC F | 10 | 4.5592E-11 | 1.66741E-11 | 5.27282E-12 | 3.3664E-11 | 5.7520E-11 | 2.73E-11 | 7.61E-11 |
| | THC+CBD F | 10 | 6.0121E-11 | 1.80579E-11 | 5.71042E-12 | 4.7204E-11 | 7.3039E-11 | 2.57E-11 | 8.43E-11 |
| | Veh M | 7 | 3.1528E-11 | 1.36350E-11 | 5.15355E-12 | 1.8918E-11 | 4.4139E-11 | 1.40E-11 | 5.21E-11 |
| | CBD M | 10 | 2.4264E-11 | 6.40805E-12 | 2.02640E-12 | 1.9680E-11 | 2.8848E-11 | 1.70E-11 | 3.63E-11 |
| | THC M | 9 | 4.6396E-11 | 1.42909E-11 | 4.76365E-12 | 3.5411E-11 | 5.7381E-11 | 2.99E-11 | 7.29E-11 |
| | THC+CBD M | 10 | 2.4351E-11 | 1.12894E-11 | 3.57002E-12 | 1.6275E-11 | 3.2427E-11 | 1.53E-11 | 5.30E-11 |
| *N*-stearoyl phenylalanine | Veh F | 8 | 5.1157E-12 | 1.83728E-12 | 6.49578E-13 | 3.5797E-12 | 6.6518E-12 | 2.26E-12 | 7.49E-12 |
| | CBD F | 7 | 4.6175E-12 | 1.93602E-12 | 7.31746E-13 | 2.8270E-12 | 6.4080E-12 | 2.17E-12 | 8.14E-12 |
| | THC F | 10 | 5.4025E-12 | 2.74652E-12 | 8.68525E-13 | 3.4378E-12 | 7.3672E-12 | 2.17E-12 | 9.80E-12 |
| | THC+CBD F | 9 | 5.8880E-12 | 1.49057E-12 | 4.96858E-13 | 4.7423E-12 | 7.0338E-12 | 3.02E-12 | 8.54E-12 |
| | Veh M | 7 | 3.4060E-12 | 1.10328E-12 | 4.17002E-13 | 2.3856E-12 | 4.4264E-12 | 2.06E-12 | 4.87E-12 |
| | CBD M | 9 | 2.9024E-12 | 1.40884E-12 | 4.69613E-13 | 1.8194E-12 | 3.9853E-12 | 8.33E-13 | 4.95E-12 |
| | THC M | 9 | 3.5308E-12 | 1.13079E-12 | 3.76931E-13 | 2.6616E-12 | 4.4000E-12 | 2.09E-12 | 5.33E-12 |
| | THC+CBD M | 10 | 2.7897E-12 | 9.62643E-13 | 3.04414E-13 | 2.1011E-12 | 3.4784E-12 | 1.40E-12 | 4.06E-12 |
| *N*-oleoyl phenylalanine | Veh F | 8 | 2.5729E-11 | 9.79233E-12 | 3.46211E-12 | 1.7543E-11 | 3.3916E-11 | 1.67E-11 | 4.31E-11 |
| | CBD F | 7 | 2.8014E-11 | 1.09536E-11 | 4.14007E-12 | 1.7883E-11 | 3.8144E-11 | 1.31E-11 | 4.20E-11 |
| | THC F | 10 | 4.5560E-11 | 1.95772E-11 | 6.19084E-12 | 3.1555E-11 | 5.9565E-11 | 1.81E-11 | 8.06E-11 |
| | THC+CBD F | 10 | 5.3682E-11 | 1.35304E-11 | 4.27870E-12 | 4.4003E-11 | 6.3361E-11 | 3.29E-11 | 7.89E-11 |
| | Veh M | 7 | 2.1180E-11 | 8.86553E-12 | 3.35086E-12 | 1.2981E-11 | 2.9379E-11 | 1.18E-11 | 3.47E-11 |
| | CBD M | 10 | 1.7768E-11 | 5.24435E-12 | 1.65841E-12 | 1.4017E-11 | 2.1520E-11 | 9.25E-12 | 2.44E-11 |
| | THC M | 8 | 2.6354E-11 | 7.52392E-12 | 2.66011E-12 | 2.0064E-11 | 3.2644E-11 | 1.49E-11 | 3.52E-11 |
| | THC+CBD M | 10 | 2.2149E-11 | 7.03909E-12 | 2.22596E-12 | 1.7113E-11 | 2.7184E-11 | 1.38E-11 | 3.69E-11 |
| *N*-linoleoyl phenylalanine | Veh F | 8 | 2.0533E-11 | 7.96714E-12 | 2.81681E-12 | 1.3872E-11 | 2.7193E-11 | 1.08E-11 | 3.43E-11 |
| | CBD F | 7 | 2.4197E-11 | 1.72402E-11 | 6.51618E-12 | 8.2524E-12 | 4.0141E-11 | 3.60E-12 | 4.88E-11 |
| | THC F | 9 | 2.2002E-11 | 8.58741E-12 | 2.86247E-12 | 1.5401E-11 | 2.8603E-11 | 8.09E-12 | 3.57E-11 |
| | THC+CBD F | 9 | 4.5811E-11 | 8.44998E-12 | 2.81666E-12 | 3.9316E-11 | 5.2306E-11 | 3.12E-11 | 5.73E-11 |
| | Veh M | 7 | 1.3156E-11 | 5.95313E-12 | 2.25007E-12 | 7.6503E-12 | 1.8662E-11 | 5.31E-12 | 2.02E-11 |
| | CBD M | 9 | 8.8434E-12 | 2.30215E-12 | 7.67384E-13 | 7.0738E-12 | 1.0613E-11 | 5.62E-12 | 1.34E-11 |
| | THC M | 9 | 2.4832E-11 | 8.92946E-12 | 2.97649E-12 | 1.7968E-11 | 3.1696E-11 | 1.50E-11 | 3.90E-11 |
| | THC+CBD M | 10 | 1.2709E-11 | 7.22657E-12 | 2.28524E-12 | 7.5390E-12 | 1.7878E-11 | 4.31E-12 | 2.46E-11 |
| *N*-arachidonoyl phenylalanine | Veh F | 8 | 2.4323E-11 | 1.39432E-11 | 4.92968E-12 | 1.2666E-11 | 3.5980E-11 | 9.22E-12 | 4.83E-11 |
| | CBD F | 7 | 3.7661E-11 | 2.43727E-11 | 9.21200E-12 | 1.5120E-11 | 6.0202E-11 | 1.60E-11 | 8.21E-11 |
| | THC F | 9 | 3.3098E-11 | 8.68278E-12 | 2.89426E-12 | 2.6424E-11 | 3.9773E-11 | 1.86E-11 | 4.60E-11 |
| | THC+CBD F | 10 | 4.6355E-11 | 1.53551E-11 | 4.85572E-12 | 3.5371E-11 | 5.7339E-11 | 2.21E-11 | 6.95E-11 |
| | Veh M | 7 | 2.0627E-11 | 1.16529E-11 | 4.40437E-12 | 9.8498E-12 | 3.1404E-11 | 7.20E-12 | 3.85E-11 |
| | CBD M | 10 | 1.4867E-11 | 7.03843E-12 | 2.22575E-12 | 9.8318E-12 | 1.9902E-11 | 7.36E-12 | 2.81E-11 |
| | THC M | 9 | 3.4111E-11 | 1.12969E-11 | 3.76564E-12 | 2.5428E-11 | 4.2795E-11 | 1.66E-11 | 5.13E-11 |
| | THC+CBD M | 10 | 1.6277E-11 | 9.60332E-12 | 3.03684E-12 | 9.4074E-12 | 2.3147E-11 | 5.98E-12 | 3.47E-11 |
| | Veh F | 8 | 1.4922E-11 | 6.94805E-12 | 2.45651E-12 | 9.1132E-12 | 2.0731E-11 | 3.84E-12 | 2.34E-11 |

| | | | | | | | | | |
|---|---|---|---|---|---|---|---|---|---|
| *N*-docosahexaenoyl phenylalanine | CBD F | 8 | 2.8870E-11 | 2.41250E-11 | 8.52948E-12 | 8.7013E-12 | 4.9039E-11 | 5.10E-12 | 7.57E-11 |
| | THC F | 10 | 2.5334E-11 | 1.00059E-11 | 3.16414E-12 | 1.8176E-11 | 3.2492E-11 | 1.30E-11 | 4.08E-11 |
| | THC+CBD F | 10 | 3.4961E-11 | 1.45411E-11 | 4.59829E-12 | 2.4559E-11 | 4.5363E-11 | 1.35E-11 | 6.09E-11 |
| | Veh M | 7 | 1.2529E-11 | 4.52454E-12 | 1.71011E-12 | 8.3444E-12 | 1.6713E-11 | 5.66E-12 | 1.84E-11 |
| | CBD M | 9 | 9.4167E-12 | 3.17080E-12 | 1.05693E-12 | 6.9794E-12 | 1.1854E-11 | 5.41E-12 | 1.57E-11 |
| | THC M | 9 | 2.2790E-11 | 1.07021E-11 | 3.56738E-12 | 1.4563E-11 | 3.1016E-11 | 9.68E-12 | 4.23E-11 |
| | THC+CBD M | 10 | 1.2018E-11 | 7.68108E-12 | 2.42897E-12 | 6.5234E-12 | 1.7513E-11 | 5.68E-12 | 3.00E-11 |

Supplemental Table 6.3 Independent t-test comparing male and female levels of *N*-acyl phenylalanines measured in plasma collected from adult mice 2h after an i.p. injection of 10mg/kg CBD, THC, or THC+CBD

| | | | Levene's Test for Equality of Variances | | t-test for Equality of Means | | | | | | |
|---|---|---|---|---|---|---|---|---|---|---|---|
| | | | | | | | Significance | | | 95% Confidence Interval of the Difference | |
| | | | F | Sig. | t | df | One-Sided p | Two-Sided p | Std. Error Difference | Lower | Upper |
| Veh | *N*-palmitoyl phenylalanine | Equal variances assumed | 0.035 | 0.855 | 0.527 | 13 | 0.304 | 0.607 | 1.21724E-09 | -1.98848E-09 | 3.27087E-09 |
| | | Equal variances not assumed | | | 0.525 | 12.519 | 0.304 | 0.609 | 1.22181E-09 | -2.00871E-09 | 3.29110E-09 |
| | *N*-stearoyl phenylalanine | Equal variances assumed | 5.983 | 0.032 | 0.975 | 11 | 0.175 | 0.351 | 8.37367E-10 | -1.02664E-09 | 2.65943E-09 |
| | | Equal variances not assumed | | | 1.039 | 7.812 | 0.165 | 0.330 | 7.85735E-10 | -1.00313E-09 | 2.63592E-09 |
| | *N*-oleoyl phenylalanine | Equal variances assumed | 0.023 | 0.881 | 0.695 | 14 | 0.249 | 0.499 | 1.15614E-09 | -1.67635E-09 | 3.28299E-09 |
| | | Equal variances not assumed | | | 0.695 | 13.956 | 0.249 | 0.499 | 1.15614E-09 | -1.67709E-09 | 3.28373E-09 |
| | *N*-linoleoyl phenylalanine | Equal variances assumed | 3.120 | 0.099 | 1.795 | 14 | 0.047 | 0.094 | 6.72514E-10 | -2.35343E-10 | 2.64945E-09 |
| | | Equal variances not assumed | | | 1.795 | 10.325 | 0.051 | 0.102 | 6.72514E-10 | -2.85031E-10 | 2.69914E-09 |
| | *N*-arachidonoyl phenylalanine | Equal variances assumed | 1.566 | 0.237 | 2.330 | 11 | 0.020 | 0.040 | 2.13354E-10 | 2.75229E-11 | 9.66701E-10 |
| | | Equal variances not assumed | | | 2.437 | 9.558 | 0.018 | 0.036 | 2.04016E-10 | 3.96713E-11 | 9.54552E-10 |
| | *N*-docosahexaenoyl phenylalanine | Equal variances assumed | 0.238 | 0.636 | 1.260 | 10 | 0.118 | 0.236 | 1.63536E-10 | -1.58251E-10 | 5.70512E-10 |
| | | Equal variances not assumed | | | 1.207 | 7.364 | 0.132 | 0.265 | 1.70759E-10 | -1.93643E-10 | 6.05904E-10 |
| CBD | *N*-palmitoyl phenylalanine | Equal variances assumed | 3.677 | 0.072 | -0.578 | 17 | 0.285 | 0.571 | 9.39758E-10 | -2.52579E-09 | 1.43964E-09 |
| | | Equal variances not assumed | | | -0.608 | 16.988 | 0.276 | 0.551 | 8.93726E-10 | -2.42877E-09 | 1.34262E-09 |

| | | | | | | | | | | | |
|---|---|---|---|---|---|---|---|---|---|---|---|
| | *N*-stearoyl phenylalanine | Equal variances assumed | 4.241 | 0.056 | -1.606 | 16 | 0.064 | 0.128 | 2.27388E-10 | -8.47143E-10 | 1.16938E-10 |
| | | Equal variances not assumed | | | -1.700 | 14.888 | 0.055 | 0.110 | 2.14741E-10 | -8.23111E-10 | 9.29057E-11 |
| | *N*-oleoyl phenylalanine | Equal variances assumed | 0.668 | 0.426 | 0.395 | 16 | 0.349 | 0.698 | 5.18065E-10 | -8.93652E-10 | 1.30284E-09 |
| | | Equal variances not assumed | | | 0.389 | 14.053 | 0.352 | 0.703 | 5.26514E-10 | -9.24262E-10 | 1.33346E-09 |
| | *N*-linoleoyl phenylalanine | Equal variances assumed | 1.270 | 0.275 | 0.534 | 17 | 0.300 | 0.600 | 4.37971E-10 | -6.90133E-10 | 1.15794E-09 |
| | | Equal variances not assumed | | | 0.516 | 13.163 | 0.307 | 0.615 | 4.53562E-10 | -7.44721E-10 | 1.21253E-09 |
| | *N*-arachidonoyl phenylalanine | Equal variances assumed | 0.723 | 0.410 | -0.536 | 14 | 0.300 | 0.601 | 7.55552E-11 | -2.02513E-10 | 1.21587E-10 |
| | | Equal variances not assumed | | | -0.521 | 11.417 | 0.306 | 0.613 | 7.77314E-11 | -2.10789E-10 | 1.29863E-10 |
| | *N*-docosahexaenoyl phenylalanine | Equal variances assumed | 0.087 | 0.772 | 0.318 | 15 | 0.377 | 0.755 | 9.46489E-11 | -1.71638E-10 | 2.31841E-10 |
| | | Equal variances not assumed | | | 0.316 | 14.154 | 0.378 | 0.757 | 9.53631E-11 | -1.74224E-10 | 2.34427E-10 |
| THC | *N*-palmitoyl phenylalanine | Equal variances assumed | 21.342 | 0.000 | -2.007 | 16 | 0.031 | 0.062 | 1.23028E-09 | -5.07750E-09 | 1.38655E-10 |
| | | Equal variances not assumed | | | -2.007 | 10.066 | 0.036 | 0.072 | 1.23028E-09 | -5.20822E-09 | 2.69375E-10 |
| | *N*-stearoyl phenylalanine | Equal variances assumed | 12.086 | 0.004 | -2.478 | 14 | 0.013 | 0.027 | 3.95114E-10 | -1.82657E-09 | -1.31705E-10 |
| | | Equal variances not assumed | | | -2.769 | 9.825 | 0.010 | 0.020 | 3.53621E-10 | -1.76896E-09 | -1.89318E-10 |
| | *N*-oleoyl phenylalanine | Equal variances assumed | 2.572 | 0.127 | -1.383 | 17 | 0.092 | 0.185 | 5.95887E-10 | -2.08132E-09 | 4.33102E-10 |
| | | Equal variances not assumed | | | -1.355 | 13.955 | 0.098 | 0.197 | 6.08067E-10 | -2.12868E-09 | 4.80457E-10 |
| | *N*-linoleoyl phenylalanine | Equal variances assumed | 0.683 | 0.421 | -1.241 | 16 | 0.116 | 0.233 | 4.80100E-10 | -1.61334E-09 | 4.22193E-10 |
| | | Equal variances not assumed | | | -1.241 | 15.607 | 0.117 | 0.233 | 4.80100E-10 | -1.61543E-09 | 4.24280E-10 |

| | | | | | | | | | | | |
|---|---|---|---|---|---|---|---|---|---|---|---|
| | *N*-arachidonoyl phenylalanine | Equal variances assumed | 1.400 | 0.260 | 0.387 | 12 | 0.353 | 0.705 | 8.94922E-11 | -1.60322E-10 | 2.29652E-10 |
| | | Equal variances not assumed | | | 0.387 | 9.270 | 0.354 | 0.707 | 8.94922E-11 | -1.66884E-10 | 2.36214E-10 |
| | *N*-docosahexaenoyl phenylalanine | Equal variances assumed | 0.109 | 0.746 | -1.658 | 13 | 0.061 | 0.121 | 1.33179E-10 | -5.08502E-10 | 6.69300E-11 |
| | | Equal variances not assumed | | | -1.656 | 12.673 | 0.061 | 0.122 | 1.33339E-10 | -5.09604E-10 | 6.80319E-11 |
| THC+CBD | *N*-palmitoyl phenylalanine | Equal variances assumed | 15.436 | 0.001 | -1.228 | 18 | 0.118 | 0.235 | 1.81255E-09 | -6.03409E-09 | 1.58197E-09 |
| | | Equal variances not assumed | | | -1.228 | 13.439 | 0.120 | 0.240 | 1.81255E-09 | -6.12888E-09 | 1.67677E-09 |
| | *N*-stearoyl phenylalanine | Equal variances assumed | 9.111 | 0.007 | -1.909 | 18 | 0.036 | 0.072 | 4.73561E-10 | -1.89888E-09 | 9.09546E-11 |
| | | Equal variances not assumed | | | -1.909 | 12.682 | 0.040 | 0.079 | 4.73561E-10 | -1.92964E-09 | 1.21721E-10 |
| | *N*-oleoyl phenylalanine | Equal variances assumed | 13.850 | 0.002 | -1.515 | 18 | 0.074 | 0.147 | 1.13916E-09 | -4.11910E-09 | 6.67489E-10 |
| | | Equal variances not assumed | | | -1.515 | 11.889 | 0.078 | 0.156 | 1.13916E-09 | -4.21039E-09 | 7.58787E-10 |
| | *N*-linoleoyl phenylalanine | Equal variances assumed | 0.292 | 0.596 | 0.138 | 17 | 0.446 | 0.892 | 6.33610E-10 | -1.24940E-09 | 1.42420E-09 |
| | | Equal variances not assumed | | | 0.139 | 16.993 | 0.446 | 0.891 | 6.29008E-10 | -1.23973E-09 | 1.41453E-09 |
| | *N*-arachidonoyl phenylalanine | Equal variances assumed | 0.445 | 0.514 | 1.038 | 17 | 0.157 | 0.314 | 1.80716E-10 | -1.93765E-10 | 5.68788E-10 |
| | | Equal variances not assumed | | | 1.045 | 16.997 | 0.155 | 0.311 | 1.79462E-10 | -1.91126E-10 | 5.66150E-10 |
| | *N*-docosahexaenoyl phenylalanine | Equal variances assumed | 9.467 | 0.008 | 1.994 | 14 | 0.033 | 0.066 | 3.00975E-10 | -4.52513E-11 | 1.24580E-09 |
| | | Equal variances not assumed | | | 2.248 | 9.105 | 0.025 | 0.051 | 2.67034E-10 | -2.73919E-12 | 1.20329E-09 |

Supplemental Table 6.4 Independent t-test comparing male and female levels of *N*-acyl phenylalanines measured in liver collected from adult mice 2h after an i.p. injection of 10mg/kg CBD, THC, or THC+CBD

| | | | Levene's Test for Equality of Variances | | t-test for Equality of Means | | | | | | | |
|---|---|---|---|---|---|---|---|---|---|---|---|---|
| | | | | | | | Significance | | | | 95% Confidence Interval of the Difference | |
| | | | F | Sig. | t | df | One-Sided p | Two-Sided p | Mean Difference | Std. Error Difference | Lower | Upper |
| Veh | *N*-palmitoyl phenylalanine | Equal variances assumed | 0.727 | 0.409 | 0.838 | 13 | 0.209 | 0.417 | 6.16263E-12 | 7.35169E-12 | -9.71973E-12 | 2.20450E-11 |
| | | Equal variances not assumed | | | 0.843 | 12.936 | 0.207 | 0.415 | 6.16263E-12 | 7.31306E-12 | -9.64422E-12 | 2.19695E-11 |
| | *N*-stearoyl phenylalanine | Equal variances assumed | 1.776 | 0.205 | 2.142 | 13 | 0.026 | 0.052 | 1.70974E-12 | 7.98342E-13 | -1.49739E-14 | 3.43445E-12 |
| | | Equal variances not assumed | | | 2.215 | 11.650 | 0.024 | 0.047 | 1.70974E-12 | 7.71908E-13 | 2.22747E-14 | 3.39720E-12 |
| | *N*-oleoyl phenylalanine | Equal variances assumed | 0.144 | 0.710 | 0.938 | 13 | 0.183 | 0.366 | 4.54939E-12 | 4.85252E-12 | -5.93385E-12 | 1.50326E-11 |
| | | Equal variances not assumed | | | 0.944 | 12.974 | 0.181 | 0.362 | 4.54939E-12 | 4.81814E-12 | -5.86165E-12 | 1.49604E-11 |
| | *N*-linoleoyl phenylalanine | Equal variances assumed | 0.632 | 0.441 | 2.005 | 13 | 0.033 | 0.066 | 7.37664E-12 | 3.67918E-12 | -5.71757E-13 | 1.53250E-11 |
| | | Equal variances not assumed | | | 2.046 | 12.734 | 0.031 | 0.062 | 7.37664E-12 | 3.60517E-12 | -4.28409E-13 | 1.51817E-11 |
| | *N*-arachidonoyl phenylalanine | Equal variances assumed | 0.316 | 0.584 | 0.552 | 13 | 0.295 | 0.590 | 3.69609E-12 | 6.69535E-12 | -1.07683E-11 | 1.81605E-11 |
| | | Equal variances not assumed | | | 0.559 | 12.984 | 0.293 | 0.586 | 3.69609E-12 | 6.61062E-12 | -1.05871E-11 | 1.79793E-11 |
| | *N*-docosahexaenoyl phenylalanine | Equal variances assumed | 2.132 | 0.168 | 0.777 | 13 | 0.226 | 0.451 | 2.39302E-12 | 3.08117E-12 | -4.26344E-12 | 9.04948E-12 |
| | | Equal variances not assumed | | | 0.799 | 12.111 | 0.220 | 0.439 | 2.39302E-12 | 2.99314E-12 | -4.12189E-12 | 8.90793E-12 |
| CBD | *N*-palmitoyl phenylalanine | Equal variances assumed | 17.850 | 0.001 | 2.662 | 15 | 0.009 | 0.018 | 2.22129E-11 | 8.34323E-12 | 4.42973E-12 | 3.99961E-11 |
| | | Equal variances not assumed | | | 2.248 | 6.530 | 0.031 | 0.062 | 2.22129E-11 | 9.88302E-12 | -1.50221E-12 | 4.59280E-11 |
| | *N*-stearoyl phenylalanine | Equal variances assumed | 0.103 | 0.753 | 2.056 | 14 | 0.029 | 0.059 | 1.71511E-12 | 8.34273E-13 | -7.42309E-14 | 3.50444E-12 |

| | | | | | | | | | | | | |
|---|---|---|---|---|---|---|---|---|---|---|---|---|
| | | Equal variances not assumed | | | 1.973 | 10.610 | 0.038 | 0.075 | 1.71511E-12 | 8.69476E-13 | -2.07208E-13 | 3.63742E-12 |
| | *N*-oleoyl phenylalanine | Equal variances assumed | 7.332 | 0.016 | 2.589 | 15 | 0.010 | 0.021 | 1.02452E-11 | 3.95764E-12 | 1.80969E-12 | 1.86807E-11 |
| | | Equal variances not assumed | | | 2.297 | 7.944 | 0.025 | 0.051 | 1.02452E-11 | 4.45987E-12 | -5.19952E-14 | 2.05424E-11 |
| | *N*-linoleoyl phenylalanine | Equal variances assumed | 23.481 | 0.000 | 2.668 | 14 | 0.009 | 0.018 | 1.53536E-11 | 5.75500E-12 | 3.01033E-12 | 2.76968E-11 |
| | | Equal variances not assumed | | | 2.340 | 6.167 | 0.028 | 0.057 | 1.53536E-11 | 6.56121E-12 | -5.96504E-13 | 3.13037E-11 |
| | *N*-arachidonoyl phenylalanine | Equal variances assumed | 10.302 | 0.006 | 2.829 | 15 | 0.006 | 0.013 | 2.27945E-11 | 8.05755E-12 | 5.62024E-12 | 3.99688E-11 |
| | | Equal variances not assumed | | | 2.405 | 6.706 | 0.024 | 0.049 | 2.27945E-11 | 9.47708E-12 | 1.83876E-13 | 4.54051E-11 |
| | *N*-docosahexaenoyl phenylalanine | Equal variances assumed | 11.259 | 0.004 | 2.406 | 15 | 0.015 | 0.029 | 1.94537E-11 | 8.08676E-12 | 2.21718E-12 | 3.66902E-11 |
| | | Equal variances not assumed | | | 2.263 | 7.215 | 0.028 | 0.057 | 1.94537E-11 | 8.59472E-12 | -7.47381E-13 | 3.96548E-11 |
| | *N*-palmitoyl phenylalanine | Equal variances assumed | 0.131 | 0.722 | -0.112 | 17 | 0.456 | 0.912 | -8.04248E-13 | 7.16681E-12 | -1.59249E-11 | 1.43164E-11 |
| | | Equal variances not assumed | | | -0.113 | 16.969 | 0.456 | 0.911 | -8.04248E-13 | 7.10598E-12 | -1.57986E-11 | 1.41901E-11 |
| | *N*-stearoyl phenylalanine | Equal variances assumed | 11.626 | 0.003 | 1.900 | 17 | 0.037 | 0.074 | 1.87169E-12 | 9.84944E-13 | -2.06363E-13 | 3.94974E-12 |
| | | Equal variances not assumed | | | 1.977 | 12.222 | 0.036 | 0.071 | 1.87169E-12 | 9.46791E-13 | -1.87049E-13 | 3.93043E-12 |
| THC | *N*-oleoyl phenylalanine | Equal variances assumed | 7.686 | 0.014 | 2.612 | 16 | 0.009 | 0.019 | 1.92060E-11 | 7.35388E-12 | 3.61644E-12 | 3.47955E-11 |
| | | Equal variances not assumed | | | 2.850 | 12.100 | 0.007 | 0.015 | 1.92060E-11 | 6.73815E-12 | 4.53821E-12 | 3.38737E-11 |
| | *N*-linoleoyl phenylalanine | Equal variances assumed | 0.107 | 0.748 | -0.685 | 16 | 0.252 | 0.503 | -2.82972E-12 | 4.12955E-12 | -1.15840E-11 | 5.92454E-12 |
| | | Equal variances not assumed | | | -0.685 | 15.976 | 0.252 | 0.503 | -2.82972E-12 | 4.12955E-12 | -1.15851E-11 | 5.92562E-12 |
| | *N*-arachidonoyl phenylalanine | Equal variances assumed | 1.064 | 0.318 | -0.213 | 16 | 0.417 | 0.834 | -1.01283E-12 | 4.74940E-12 | -1.10811E-11 | 9.05544E-12 |
| | | Equal variances | | | -0.213 | 15.007 | 0.417 | 0.834 | -1.01283E-12 | 4.74940E-12 | -1.11355E-11 | 9.10988E-12 |

| | | | | | | | | | | | | |
|---|---|---|---|---|---|---|---|---|---|---|---|---|
| | | not assumed | | | | | | | | | | |
| | *N*-docosahexaenoyl phenylalanine | Equal variances assumed | 0.000 | 0.988 | 0.536 | 17 | 0.300 | 0.599 | 2.54424E-12 | 4.75062E-12 | -7.47869E-12 | 1.25672E-11 |
| | | Equal variances not assumed | | | 0.534 | 16.475 | 0.300 | 0.601 | 2.54424E-12 | 4.76844E-12 | -7.54076E-12 | 1.26292E-11 |
| THC+CBD | *N*-palmitoyl phenylalanine | Equal variances assumed | 1.807 | 0.196 | 5.312 | 18 | 0.000 | 0.000 | 3.57706E-11 | 6.73454E-12 | 2.16219E-11 | 4.99194E-11 |
| | | Equal variances not assumed | | | 5.312 | 15.103 | 0.000 | 0.000 | 3.57706E-11 | 6.73454E-12 | 2.14248E-11 | 5.01164E-11 |
| | *N*-stearoyl phenylalanine | Equal variances assumed | 0.337 | 0.569 | 5.441 | 17 | 0.000 | 0.000 | 3.09828E-12 | 5.69472E-13 | 1.89680E-12 | 4.29976E-12 |
| | | Equal variances not assumed | | | 5.317 | 13.449 | 0.000 | 0.000 | 3.09828E-12 | 5.82697E-13 | 1.84370E-12 | 4.35286E-12 |
| | *N*-oleoyl phenylalanine | Equal variances assumed | 2.546 | 0.128 | 6.538 | 18 | 0.000 | 0.000 | 3.15333E-11 | 4.82308E-12 | 2.14003E-11 | 4.16662E-11 |
| | | Equal variances not assumed | | | 6.538 | 13.539 | 0.000 | 0.000 | 3.15333E-11 | 4.82308E-12 | 2.11556E-11 | 4.19109E-11 |
| | *N*-linoleoyl phenylalanine | Equal variances assumed | 0.061 | 0.808 | 9.206 | 17 | 0.000 | 0.000 | 3.31025E-11 | 3.59587E-12 | 2.55159E-11 | 4.06891E-11 |
| | | Equal variances not assumed | | | 9.126 | 15.882 | 0.000 | 0.000 | 3.31025E-11 | 3.62711E-12 | 2.54087E-11 | 4.07963E-11 |
| | *N*-arachidonoyl phenylalanine | Equal variances assumed | 2.548 | 0.128 | 5.252 | 18 | 0.000 | 0.000 | 3.00779E-11 | 5.72717E-12 | 1.80455E-11 | 4.21102E-11 |
| | | Equal variances not assumed | | | 5.252 | 15.106 | 0.000 | 0.000 | 3.00779E-11 | 5.72717E-12 | 1.78782E-11 | 4.22775E-11 |
| | *N*-docosahexaenoyl phenylalanine | Equal variances assumed | 4.071 | 0.059 | 4.412 | 18 | 0.000 | 0.000 | 2.29432E-11 | 5.20041E-12 | 1.20175E-11 | 3.38688E-11 |
| | | Equal variances not assumed | | | 4.412 | 13.660 | 0.000 | 0.001 | 2.29432E-11 | 5.20041E-12 | 1.17633E-11 | 3.41231E-11 |

Supplemental Table 6.5 One-way ANOVA for *N*-acyl phenylalanines measured in plasma collected from adult mice 2h after an i.p. injection of 10mg/kg CBD, THC, or THC+CBD

| | | Sum of Squares | df | Mean Square | F | Sig. |
|---|---|---|---|---|---|---|
| *N*-palmitoyl phenylalanine | Between Groups | 1.21E-16 | 7 | 1.72E-17 | 2.018 | 0.0662397 |
| | Within Groups | 5.46E-16 | 64 | 8.53E-18 | | |
| | Total | 6.67E-16 | 71 | | | |
| *N*-stearoyl phenylalanine | Between Groups | 1.56E-17 | 7 | 2.23E-18 | 2.288 | 0.0393395 |
| | Within Groups | 5.74E-17 | 59 | 9.73E-19 | | |
| | Total | 7.30E-17 | 66 | | | |
| *N*-oleoyl phenylalanine | Between Groups | 6.06E-17 | 7 | 8.66E-18 | 2.352 | 0.0332511 |
| | Within Groups | 2.39E-16 | 65 | 3.68E-18 | | |
| | Total | 3.00E-16 | 72 | | | |
| *N*-linoleoyl phenylalanine | Between Groups | 1.55E-17 | 7 | 2.21E-18 | 1.585 | 0.1561326 |
| | Within Groups | 8.94E-17 | 64 | 1.40E-18 | | |
| | Total | 1.05E-16 | 71 | | | |
| *N*-arachidonoyl phenylalanine | Between Groups | 1.94E-18 | 7 | 2.77E-19 | 3.058 | 0.0087374 |
| | Within Groups | 4.90E-18 | 54 | 9.07E-20 | | |
| | Total | 6.84E-18 | 61 | | | |
| *N*-docosahexaenoyl phenylalanine | Between Groups | 2.50E-18 | 7 | 3.58E-19 | 2.581 | 0.0231319 |
| | Within Groups | 7.20E-18 | 52 | 1.39E-19 | | |
| | Total | 9.71E-18 | 59 | | | |

Supplemental Table 6.6 One-way ANOVA for *N*-acyl phenylalanines measured in liver collected from adult mice 2h after an i.p. injection of 10mg/kg CBD, THC, or THC+CBD

| | | Sum of Squares | df | Mean Square | F | Sig. |
|---|---|---|---|---|---|---|
| *N*-palmitoyl phenylalanine | Between Groups | 1.05E-20 | 7 | 1.50E-21 | 6.229 | 0.00001 |
| | Within Groups | 1.51E-20 | 63 | 2.40E-22 | | |
| | Total | 2.56E-20 | 70 | | | |
| *N*-stearoyl phenylalanine | Between Groups | 9.15E-23 | 7 | 1.31E-23 | 4.596 | 0.00036 |
| | Within Groups | 1.74E-22 | 61 | 2.84E-24 | | |
| | Total | 2.65E-22 | 68 | | | |
| *N*-oleoyl phenylalanine | Between Groups | 1.09E-20 | 7 | 1.56E-21 | 12.023 | 0.00000 |
| | Within Groups | 8.05E-21 | 62 | 1.30E-22 | | |
| | Total | 1.90E-20 | 69 | | | |
| *N*-linoleoyl phenylalanine | Between Groups | 8.18E-21 | 7 | 1.17E-21 | 14.758 | 0.00000 |
| | Within Groups | 4.75E-21 | 60 | 7.92E-23 | | |
| | Total | 1.29E-20 | 67 | | | |
| *N*-arachidonoyl phenylalanine | Between Groups | 8.17E-21 | 7 | 1.17E-21 | 6.727 | 0.00001 |
| | Within Groups | 1.08E-20 | 62 | 1.74E-22 | | |
| | Total | 1.89E-20 | 69 | | | |
| *N*-docosahexaenoyl phenylalanine | Between Groups | 5.45E-21 | 7 | 7.79E-22 | 5.533 | 0.00005 |
| | Within Groups | 8.87E-21 | 63 | 1.41E-22 | | |
| | Total | 1.43E-20 | 70 | | | |

#### Supplemental Table 6.7 Fisher's LSD post-hoc test for plasma *N*-acyl phenylalanine ANOVA

| Dependent Variable | | | Std. Error | Sig. | 95% Confidence Interval Lower Bound | 95% Confidence Interval Upper Bound |
|---|---|---|---|---|---|---|
| *N*-palmitoyl phenylalanine | VehFemale | CBDfemale | 1.51E-09 | 0.3438 | -1.58E-09 | 4.46E-09 |
| | | THCfemale | 1.47E-09 | 0.3137 | -1.45E-09 | 4.44E-09 |
| | | TpCfemale | 1.44E-09 | 0.8496 | -3.15E-09 | 2.60E-09 |
| | | VehMale | 1.51E-09 | 0.6729 | -2.38E-09 | 3.66E-09 |
| | | CBDmale | 1.41E-09 | 0.5268 | -1.92E-09 | 3.72E-09 |
| | | THCmale | 1.47E-09 | 0.5104 | -3.92E-09 | 1.97E-09 |
| | | TpCmale | 1.44E-09 | 0.0872 | -5.38E-09 | 3.76E-10 |
| | CBDfemale | VehFemale | 1.51E-09 | 0.3438 | -4.46E-09 | 1.58E-09 |
| | | THCfemale | 1.42E-09 | 0.9702 | -2.78E-09 | 2.89E-09 |
| | | TpCfemale | 1.39E-09 | 0.2201 | -4.48E-09 | 1.05E-09 |
| | | VehMale | 1.46E-09 | 0.5855 | -3.72E-09 | 2.12E-09 |
| | | CBDmale | 1.36E-09 | 0.6904 | -3.25E-09 | 2.17E-09 |
| | | THCmale | 1.42E-09 | 0.0935 | -5.25E-09 | 4.19E-10 |
| | | TpCmale | 1.39E-09 | 0.0060 | -6.71E-09 | -1.17E-09 |
| | THCfemale | VehFemale | 1.47E-09 | 0.3137 | -4.44E-09 | 1.45E-09 |
| | | CBDfemale | 1.42E-09 | 0.9702 | -2.89E-09 | 2.78E-09 |
| | | TpCfemale | 1.34E-09 | 0.1922 | -4.45E-09 | 9.12E-10 |
| | | VehMale | 1.42E-09 | 0.5496 | -3.69E-09 | 1.98E-09 |
| | | CBDmale | 1.31E-09 | 0.6513 | -3.22E-09 | 2.03E-09 |
| | | THCmale | 1.38E-09 | 0.0776 | -5.22E-09 | 2.82E-10 |
| | | TpCmale | 1.34E-09 | 0.0041 | -6.68E-09 | -1.31E-09 |
| | TpCfemale | VehFemale | 1.44E-09 | 0.8496 | -2.60E-09 | 3.15E-09 |
| | | CBDfemale | 1.39E-09 | 0.2201 | -1.05E-09 | 4.48E-09 |
| | | THCfemale | 1.34E-09 | 0.1922 | -9.12E-10 | 4.45E-09 |
| | | VehMale | 1.39E-09 | 0.5113 | -1.85E-09 | 3.68E-09 |
| | | CBDmale | 1.28E-09 | 0.3616 | -1.38E-09 | 3.72E-09 |
| | | THCmale | 1.34E-09 | 0.6036 | -3.38E-09 | 1.98E-09 |
| | | TpCmale | 1.31E-09 | 0.0932 | -4.84E-09 | 3.84E-10 |
| | VehMale | VehFemale | 1.51E-09 | 0.6729 | -3.66E-09 | 2.38E-09 |
| | | CBDfemale | 1.46E-09 | 0.5855 | -2.12E-09 | 3.72E-09 |
| | | THCfemale | 1.42E-09 | 0.5496 | -1.98E-09 | 3.69E-09 |
| | | TpCfemale | 1.39E-09 | 0.5113 | -3.68E-09 | 1.85E-09 |
| | | CBDmale | 1.36E-09 | 0.8501 | -2.45E-09 | 2.97E-09 |
| | | THCmale | 1.42E-09 | 0.2592 | -4.45E-09 | 1.22E-09 |
| | | TpCmale | 1.39E-09 | 0.0268 | -5.91E-09 | -3.73E-10 |
| | CBDmale | VehFemale | 1.41E-09 | 0.5268 | -3.72E-09 | 1.92E-09 |
| | | CBDfemale | 1.36E-09 | 0.6904 | -2.17E-09 | 3.25E-09 |
| | | THCfemale | 1.31E-09 | 0.6513 | -2.03E-09 | 3.22E-09 |
| | | TpCfemale | 1.28E-09 | 0.3616 | -3.72E-09 | 1.38E-09 |
| | | VehMale | 1.36E-09 | 0.8501 | -2.97E-09 | 2.45E-09 |
| | | THCmale | 1.31E-09 | 0.1585 | -4.50E-09 | 7.50E-10 |
| | | TpCmale | 1.28E-09 | 0.0098 | -5.95E-09 | -8.49E-10 |
| | THCmale | VehFemale | 1.47E-09 | 0.5104 | -1.97E-09 | 3.92E-09 |
| | | CBDfemale | 1.42E-09 | 0.0935 | -4.19E-10 | 5.25E-09 |
| | | THCfemale | 1.38E-09 | 0.0776 | -2.82E-10 | 5.22E-09 |
| | | TpCfemale | 1.34E-09 | 0.6036 | -1.98E-09 | 3.38E-09 |
| | | VehMale | 1.42E-09 | 0.2592 | -1.22E-09 | 4.45E-09 |
| | | CBDmale | 1.31E-09 | 0.1585 | -7.50E-10 | 4.50E-09 |
| | | TpCmale | 1.34E-09 | 0.2599 | -4.21E-09 | 1.16E-09 |
| | TpCmale | VehFemale | 1.44E-09 | 0.0872 | -3.76E-10 | 5.38E-09 |
| | | CBDfemale | 1.39E-09 | 0.0060 | 1.17E-09 | 6.71E-09 |
| | | THCfemale | 1.34E-09 | 0.0041 | 1.31E-09 | 6.68E-09 |
| | | TpCfemale | 1.31E-09 | 0.0932 | -3.84E-10 | 4.84E-09 |
| | | VehMale | 1.39E-09 | 0.0268 | 3.73E-10 | 5.91E-09 |
| | | CBDmale | 1.28E-09 | 0.0098 | 8.49E-10 | 5.95E-09 |
| | | THCmale | 1.34E-09 | 0.2599 | -1.16E-09 | 4.21E-09 |
| *N*-stearoyl phenylalanine | VehFemale | CBDfemale | 5.10E-10 | 0.0376 | 6.42E-11 | 2.11E-09 |

| | | | | | | |
|---|---|---|---|---|---|---|
| | | THCfemale | 5.27E-10 | 0.0331 | 9.53E-11 | 2.20E-09 |
| | | TpCfemale | 4.86E-10 | 0.1951 | -3.36E-10 | 1.61E-09 |
| | | VehMale | 5.49E-10 | 0.1421 | -2.82E-10 | 1.91E-09 |
| | | CBDmale | 4.86E-10 | 0.1436 | -2.52E-10 | 1.69E-09 |
| | | THCmale | 4.97E-10 | 0.7320 | -8.23E-10 | 1.17E-09 |
| | | TpCmale | 4.86E-10 | 0.5849 | -1.24E-09 | 7.06E-10 |
| | CBDfemale | VehFemale | 5.10E-10 | 0.0376 | -2.11E-09 | -6.42E-11 |
| | | THCfemale | 5.10E-10 | 0.8996 | -9.57E-10 | 1.09E-09 |
| | | TpCfemale | 4.68E-10 | 0.3416 | -1.38E-09 | 4.88E-10 |
| | | VehMale | 5.33E-10 | 0.6153 | -1.33E-09 | 7.97E-10 |
| | | CBDmale | 4.68E-10 | 0.4382 | -1.30E-09 | 5.71E-10 |
| | | THCmale | 4.79E-10 | 0.0612 | -1.87E-09 | 4.44E-11 |
| | | TpCmale | 4.68E-10 | 0.0054 | -2.29E-09 | -4.16E-10 |
| | THCfemale | VehFemale | 5.27E-10 | 0.0331 | -2.20E-09 | -9.53E-11 |
| | | CBDfemale | 5.10E-10 | 0.8996 | -1.09E-09 | 9.57E-10 |
| | | TpCfemale | 4.86E-10 | 0.2953 | -1.49E-09 | 4.59E-10 |
| | | VehMale | 5.49E-10 | 0.5453 | -1.43E-09 | 7.64E-10 |
| | | CBDmale | 4.86E-10 | 0.3802 | -1.40E-09 | 5.43E-10 |
| | | THCmale | 4.97E-10 | 0.0535 | -1.97E-09 | 1.54E-11 |
| | | TpCmale | 4.86E-10 | 0.0050 | -2.39E-09 | -4.45E-10 |
| | TpCfemale | VehFemale | 4.86E-10 | 0.1951 | -1.61E-09 | 3.36E-10 |
| | | CBDfemale | 4.68E-10 | 0.3416 | -4.88E-10 | 1.38E-09 |
| | | THCfemale | 4.86E-10 | 0.2953 | -4.59E-10 | 1.49E-09 |
| | | VehMale | 5.09E-10 | 0.7259 | -8.40E-10 | 1.20E-09 |
| | | CBDmale | 4.41E-10 | 0.8507 | -7.99E-10 | 9.66E-10 |
| | | THCmale | 4.53E-10 | 0.3080 | -1.37E-09 | 4.41E-10 |
| | | TpCmale | 4.41E-10 | 0.0449 | -1.79E-09 | -2.14E-11 |
| | VehMale | VehFemale | 5.49E-10 | 0.1421 | -1.91E-09 | 2.82E-10 |
| | | CBDfemale | 5.33E-10 | 0.6153 | -7.97E-10 | 1.33E-09 |
| | | THCfemale | 5.49E-10 | 0.5453 | -7.64E-10 | 1.43E-09 |
| | | TpCfemale | 5.09E-10 | 0.7259 | -1.20E-09 | 8.40E-10 |
| | | CBDmale | 5.09E-10 | 0.8511 | -1.12E-09 | 9.23E-10 |
| | | THCmale | 5.20E-10 | 0.2193 | -1.69E-09 | 3.95E-10 |
| | | TpCmale | 5.09E-10 | 0.0376 | -2.10E-09 | -6.43E-11 |
| | CBDmale | VehFemale | 4.86E-10 | 0.1436 | -1.69E-09 | 2.52E-10 |
| | | CBDfemale | 4.68E-10 | 0.4382 | -5.71E-10 | 1.30E-09 |
| | | THCfemale | 4.86E-10 | 0.3802 | -5.43E-10 | 1.40E-09 |
| | | TpCfemale | 4.41E-10 | 0.8507 | -9.66E-10 | 7.99E-10 |
| | | VehMale | 5.09E-10 | 0.8511 | -9.23E-10 | 1.12E-09 |
| | | THCmale | 4.53E-10 | 0.2302 | -1.46E-09 | 3.57E-10 |
| | | TpCmale | 4.41E-10 | 0.0290 | -1.87E-09 | -1.05E-10 |
| | THCmale | VehFemale | 4.97E-10 | 0.7320 | -1.17E-09 | 8.23E-10 |
| | | CBDfemale | 4.79E-10 | 0.0612 | -4.44E-11 | 1.87E-09 |
| | | THCfemale | 4.97E-10 | 0.0535 | -1.54E-11 | 1.97E-09 |
| | | TpCfemale | 4.53E-10 | 0.3080 | -4.41E-10 | 1.37E-09 |
| | | VehMale | 5.20E-10 | 0.2193 | -3.95E-10 | 1.69E-09 |
| | | CBDmale | 4.53E-10 | 0.2302 | -3.57E-10 | 1.46E-09 |
| | | TpCmale | 4.53E-10 | 0.3377 | -1.34E-09 | 4.69E-10 |
| | TpCmale | VehFemale | 4.86E-10 | 0.5849 | -7.06E-10 | 1.24E-09 |
| | | CBDfemale | 4.68E-10 | 0.0054 | 4.16E-10 | 2.29E-09 |
| | | THCfemale | 4.86E-10 | 0.0050 | 4.45E-10 | 2.39E-09 |
| | | TpCfemale | 4.41E-10 | 0.0449 | 2.14E-11 | 1.79E-09 |
| | | VehMale | 5.09E-10 | 0.0376 | 6.43E-11 | 2.10E-09 |
| | | CBDmale | 4.41E-10 | 0.0290 | 1.05E-10 | 1.87E-09 |
| | | THCmale | 4.53E-10 | 0.3377 | -4.69E-10 | 1.34E-09 |
| *N*-oleoyl phenylalanine | VehFemale | CBDfemale | 9.59E-10 | 0.1262 | -4.30E-10 | 3.40E-09 |
| | | THCfemale | 9.10E-10 | 0.0553 | -4.16E-11 | 3.59E-09 |
| | | TpCfemale | 9.10E-10 | 0.4551 | -1.13E-09 | 2.50E-09 |
| | | VehMale | 9.59E-10 | 0.4055 | -1.11E-09 | 2.72E-09 |
| | | CBDmale | 9.10E-10 | 0.0677 | -1.27E-10 | 3.51E-09 |

| | | | | | | |
|---|---|---|---|---|---|---|
| | | THCmale | 9.32E-10 | 0.3110 | -9.10E-10 | 2.81E-09 |
| | | TpCmale | 9.10E-10 | 0.2566 | -2.86E-09 | 7.76E-10 |
| | CBDfemale | VehFemale | 9.59E-10 | 0.1262 | -3.40E-09 | 4.30E-10 |
| | | THCfemale | 9.10E-10 | 0.7511 | -1.53E-09 | 2.11E-09 |
| | | TpCfemale | 9.10E-10 | 0.3813 | -2.62E-09 | 1.02E-09 |
| | | VehMale | 9.59E-10 | 0.4791 | -2.60E-09 | 1.23E-09 |
| | | CBDmale | 9.10E-10 | 0.8229 | -1.61E-09 | 2.02E-09 |
| | | THCmale | 9.32E-10 | 0.5687 | -2.40E-09 | 1.33E-09 |
| | | TpCmale | 9.10E-10 | 0.0071 | -4.35E-09 | -7.10E-10 |
| | THCfemale | VehFemale | 9.10E-10 | 0.0553 | -3.59E-09 | 4.16E-11 |
| | | CBDfemale | 9.10E-10 | 0.7511 | -2.11E-09 | 1.53E-09 |
| | | TpCfemale | 8.58E-10 | 0.2076 | -2.81E-09 | 6.22E-10 |
| | | VehMale | 9.10E-10 | 0.2891 | -2.79E-09 | 8.45E-10 |
| | | CBDmale | 8.58E-10 | 0.9211 | -1.80E-09 | 1.63E-09 |
| | | THCmale | 8.82E-10 | 0.3534 | -2.58E-09 | 9.37E-10 |
| | | TpCmale | 8.58E-10 | 0.0017 | -4.53E-09 | -1.10E-09 |
| | TpCfemale | VehFemale | 9.10E-10 | 0.4551 | -2.50E-09 | 1.13E-09 |
| | | CBDfemale | 9.10E-10 | 0.3813 | -1.02E-09 | 2.62E-09 |
| | | THCfemale | 8.58E-10 | 0.2076 | -6.22E-10 | 2.81E-09 |
| | | VehMale | 9.10E-10 | 0.8961 | -1.70E-09 | 1.94E-09 |
| | | CBDmale | 8.58E-10 | 0.2449 | -7.07E-10 | 2.72E-09 |
| | | THCmale | 8.82E-10 | 0.7620 | -1.49E-09 | 2.03E-09 |
| | | TpCmale | 8.58E-10 | 0.0485 | -3.44E-09 | -1.20E-11 |
| | VehMale | VehFemale | 9.59E-10 | 0.4055 | -2.72E-09 | 1.11E-09 |
| | | CBDfemale | 9.59E-10 | 0.4791 | -1.23E-09 | 2.60E-09 |
| | | THCfemale | 9.10E-10 | 0.2891 | -8.45E-10 | 2.79E-09 |
| | | TpCfemale | 9.10E-10 | 0.8961 | -1.94E-09 | 1.70E-09 |
| | | CBDmale | 9.10E-10 | 0.3331 | -9.30E-10 | 2.71E-09 |
| | | THCmale | 9.32E-10 | 0.8737 | -1.71E-09 | 2.01E-09 |
| | | TpCmale | 9.10E-10 | 0.0467 | -3.66E-09 | -2.74E-11 |
| | CBDmale | VehFemale | 9.10E-10 | 0.0677 | -3.51E-09 | 1.27E-10 |
| | | CBDfemale | 9.10E-10 | 0.8229 | -2.02E-09 | 1.61E-09 |
| | | THCfemale | 8.58E-10 | 0.9211 | -1.63E-09 | 1.80E-09 |
| | | TpCfemale | 8.58E-10 | 0.2449 | -2.72E-09 | 7.07E-10 |
| | | VehMale | 9.10E-10 | 0.3331 | -2.71E-09 | 9.30E-10 |
| | | THCmale | 8.82E-10 | 0.4051 | -2.50E-09 | 1.02E-09 |
| | | TpCmale | 8.58E-10 | 0.0022 | -4.45E-09 | -1.02E-09 |
| | THCmale | VehFemale | 9.32E-10 | 0.3110 | -2.81E-09 | 9.10E-10 |
| | | CBDfemale | 9.32E-10 | 0.5687 | -1.33E-09 | 2.40E-09 |
| | | THCfemale | 8.82E-10 | 0.3534 | -9.37E-10 | 2.58E-09 |
| | | TpCfemale | 8.82E-10 | 0.7620 | -2.03E-09 | 1.49E-09 |
| | | VehMale | 9.32E-10 | 0.8737 | -2.01E-09 | 1.71E-09 |
| | | CBDmale | 8.82E-10 | 0.4051 | -1.02E-09 | 2.50E-09 |
| | | TpCmale | 8.82E-10 | 0.0271 | -3.75E-09 | -2.33E-10 |
| | TpCmale | VehFemale | 9.10E-10 | 0.2566 | -7.76E-10 | 2.86E-09 |
| | | CBDfemale | 9.10E-10 | 0.0071 | 7.10E-10 | 4.35E-09 |
| | | THCfemale | 8.58E-10 | 0.0017 | 1.10E-09 | 4.53E-09 |
| | | TpCfemale | 8.58E-10 | 0.0485 | 1.20E-11 | 3.44E-09 |
| | | VehMale | 9.10E-10 | 0.0467 | 2.74E-11 | 3.66E-09 |
| | | CBDmale | 8.58E-10 | 0.0022 | 1.02E-09 | 4.45E-09 |
| | | THCmale | 8.82E-10 | 0.0271 | 2.33E-10 | 3.75E-09 |
| *N*-linoleoyl phenylalanine | VehFemale | CBDfemale | 5.91E-10 | 0.0353 | 9.03E-11 | 2.45E-09 |
| | | THCfemale | 5.74E-10 | 0.0236 | 1.84E-10 | 2.48E-09 |
| | | TpCfemale | 5.60E-10 | 0.2875 | -5.19E-10 | 1.72E-09 |
| | | VehMale | 5.91E-10 | 0.0452 | 2.68E-11 | 2.39E-09 |
| | | CBDmale | 5.49E-10 | 0.0079 | 4.08E-10 | 2.60E-09 |
| | | THCmale | 5.74E-10 | 0.2048 | -4.12E-10 | 1.88E-09 |
| | | TpCmale | 5.74E-10 | 0.2349 | -4.59E-10 | 1.84E-09 |
| | CBDfemale | VehFemale | 5.91E-10 | 0.0353 | -2.45E-09 | -9.03E-11 |
| | | THCfemale | 5.74E-10 | 0.9165 | -1.09E-09 | 1.21E-09 |

| | | | | | | |
|---|---|---|---|---|---|---|
| | | TpCfemale | 5.60E-10 | 0.2367 | -1.79E-09 | 4.50E-10 |
| | | VehMale | 5.91E-10 | 0.9147 | -1.24E-09 | 1.12E-09 |
| | | CBDmale | 5.49E-10 | 0.6715 | -8.63E-10 | 1.33E-09 |
| | | THCmale | 5.74E-10 | 0.3548 | -1.68E-09 | 6.12E-10 |
| | | TpCmale | 5.74E-10 | 0.3145 | -1.73E-09 | 5.65E-10 |
| | THCfemale | VehFemale | 5.74E-10 | 0.0236 | -2.48E-09 | -1.84E-10 |
| | | CBDfemale | 5.74E-10 | 0.9165 | -1.21E-09 | 1.09E-09 |
| | | TpCfemale | 5.43E-10 | 0.1835 | -1.81E-09 | 3.55E-10 |
| | | VehMale | 5.74E-10 | 0.8297 | -1.27E-09 | 1.02E-09 |
| | | CBDmale | 5.31E-10 | 0.7450 | -8.87E-10 | 1.23E-09 |
| | | THCmale | 5.57E-10 | 0.2890 | -1.71E-09 | 5.17E-10 |
| | | TpCmale | 5.57E-10 | 0.2530 | -1.76E-09 | 4.70E-10 |
| | TpCfemale | VehFemale | 5.60E-10 | 0.2875 | -1.72E-09 | 5.19E-10 |
| | | CBDfemale | 5.60E-10 | 0.2367 | -4.50E-10 | 1.79E-09 |
| | | THCfemale | 5.43E-10 | 0.1835 | -3.55E-10 | 1.81E-09 |
| | | VehMale | 5.60E-10 | 0.2837 | -5.14E-10 | 1.73E-09 |
| | | CBDmale | 5.16E-10 | 0.0849 | -1.28E-10 | 1.93E-09 |
| | | THCmale | 5.43E-10 | 0.8053 | -9.50E-10 | 1.22E-09 |
| | | TpCmale | 5.43E-10 | 0.8726 | -9.97E-10 | 1.17E-09 |
| | VehMale | VehFemale | 5.91E-10 | 0.0452 | -2.39E-09 | -2.68E-11 |
| | | CBDfemale | 5.91E-10 | 0.9147 | -1.12E-09 | 1.24E-09 |
| | | THCfemale | 5.74E-10 | 0.8297 | -1.02E-09 | 1.27E-09 |
| | | TpCfemale | 5.60E-10 | 0.2837 | -1.73E-09 | 5.14E-10 |
| | | CBDmale | 5.49E-10 | 0.5899 | -7.99E-10 | 1.39E-09 |
| | | THCmale | 5.74E-10 | 0.4145 | -1.62E-09 | 6.75E-10 |
| | | TpCmale | 5.74E-10 | 0.3698 | -1.67E-09 | 6.28E-10 |
| | CBDmale | VehFemale | 5.49E-10 | 0.0079 | -2.60E-09 | -4.08E-10 |
| | | CBDfemale | 5.49E-10 | 0.6715 | -1.33E-09 | 8.63E-10 |
| | | THCfemale | 5.31E-10 | 0.7450 | -1.23E-09 | 8.87E-10 |
| | | TpCfemale | 5.16E-10 | 0.0849 | -1.93E-09 | 1.28E-10 |
| | | VehMale | 5.49E-10 | 0.5899 | -1.39E-09 | 7.99E-10 |
| | | THCmale | 5.31E-10 | 0.1525 | -1.83E-09 | 2.92E-10 |
| | | TpCmale | 5.31E-10 | 0.1293 | -1.88E-09 | 2.45E-10 |
| | THCmale | VehFemale | 5.74E-10 | 0.2048 | -1.88E-09 | 4.12E-10 |
| | | CBDfemale | 5.74E-10 | 0.3548 | -6.12E-10 | 1.68E-09 |
| | | THCfemale | 5.57E-10 | 0.2890 | -5.17E-10 | 1.71E-09 |
| | | TpCfemale | 5.43E-10 | 0.8053 | -1.22E-09 | 9.50E-10 |
| | | VehMale | 5.74E-10 | 0.4145 | -6.75E-10 | 1.62E-09 |
| | | CBDmale | 5.31E-10 | 0.1525 | -2.92E-10 | 1.83E-09 |
| | | TpCmale | 5.57E-10 | 0.9331 | -1.16E-09 | 1.07E-09 |
| | TpCmale | VehFemale | 5.74E-10 | 0.2349 | -1.84E-09 | 4.59E-10 |
| | | CBDfemale | 5.74E-10 | 0.3145 | -5.65E-10 | 1.73E-09 |
| | | THCfemale | 5.57E-10 | 0.2530 | -4.70E-10 | 1.76E-09 |
| | | TpCfemale | 5.43E-10 | 0.8726 | -1.17E-09 | 9.97E-10 |
| | | VehMale | 5.74E-10 | 0.3698 | -6.28E-10 | 1.67E-09 |
| | | CBDmale | 5.31E-10 | 0.1293 | -2.45E-10 | 1.88E-09 |
| | | THCmale | 5.57E-10 | 0.9331 | -1.07E-09 | 1.16E-09 |
| *N*-arachidonoyl phenylalanine | VehFemale | CBDfemale | 1.61E-10 | 0.0008 | 2.51E-10 | 8.96E-10 |
| | | THCfemale | 1.61E-10 | 0.0033 | 1.73E-10 | 8.19E-10 |
| | | TpCfemale | 1.48E-10 | 0.0879 | -3.96E-11 | 5.56E-10 |
| | | VehMale | 1.68E-10 | 0.0045 | 1.61E-10 | 8.33E-10 |
| | | CBDmale | 1.52E-10 | 0.0009 | 2.29E-10 | 8.37E-10 |
| | | THCmale | 1.61E-10 | 0.0017 | 2.08E-10 | 8.53E-10 |
| | | TpCmale | 1.52E-10 | 0.0049 | 1.41E-10 | 7.50E-10 |
| | CBDfemale | VehFemale | 1.61E-10 | 0.0008 | -8.96E-10 | -2.51E-10 |
| | | THCfemale | 1.61E-10 | 0.6312 | -4.00E-10 | 2.45E-10 |
| | | TpCfemale | 1.48E-10 | 0.0380 | -6.13E-10 | -1.81E-11 |
| | | VehMale | 1.68E-10 | 0.6498 | -4.12E-10 | 2.59E-10 |
| | | CBDmale | 1.52E-10 | 0.7908 | -3.45E-10 | 2.64E-10 |
| | | THCmale | 1.61E-10 | 0.7901 | -3.66E-10 | 2.80E-10 |

| | | | | | | |
|---|---|---|---|---|---|---|
| | | TpCmale | 1.52E-10 | 0.4022 | -4.32E-10 | 1.76E-10 |
| | THCfemale | VehFemale | 1.61E-10 | 0.0033 | -8.19E-10 | -1.73E-10 |
| | | CBDfemale | 1.61E-10 | 0.6312 | -2.45E-10 | 4.00E-10 |
| | | TpCfemale | 1.48E-10 | 0.1147 | -5.36E-10 | 5.96E-11 |
| | | VehMale | 1.68E-10 | 0.9942 | -3.35E-10 | 3.37E-10 |
| | | CBDmale | 1.52E-10 | 0.8070 | -2.67E-10 | 3.42E-10 |
| | | THCmale | 1.61E-10 | 0.8303 | -2.88E-10 | 3.57E-10 |
| | | TpCmale | 1.52E-10 | 0.7410 | -3.55E-10 | 2.54E-10 |
| | TpCfemale | VehFemale | 1.48E-10 | 0.0879 | -5.56E-10 | 3.96E-11 |
| | | CBDfemale | 1.48E-10 | 0.0380 | 1.81E-11 | 6.13E-10 |
| | | THCfemale | 1.48E-10 | 0.1147 | -5.96E-11 | 5.36E-10 |
| | | VehMale | 1.56E-10 | 0.1300 | -7.27E-11 | 5.51E-10 |
| | | CBDmale | 1.38E-10 | 0.0518 | -2.24E-12 | 5.53E-10 |
| | | THCmale | 1.48E-10 | 0.0718 | -2.50E-11 | 5.70E-10 |
| | | TpCmale | 1.38E-10 | 0.1811 | -8.99E-11 | 4.65E-10 |
| | VehMale | VehFemale | 1.68E-10 | 0.0045 | -8.33E-10 | -1.61E-10 |
| | | CBDfemale | 1.68E-10 | 0.6498 | -2.59E-10 | 4.12E-10 |
| | | THCfemale | 1.68E-10 | 0.9942 | -3.37E-10 | 3.35E-10 |
| | | TpCfemale | 1.56E-10 | 0.1300 | -5.51E-10 | 7.27E-11 |
| | | CBDmale | 1.59E-10 | 0.8212 | -2.82E-10 | 3.54E-10 |
| | | THCmale | 1.68E-10 | 0.8425 | -3.03E-10 | 3.69E-10 |
| | | TpCmale | 1.59E-10 | 0.7461 | -3.70E-10 | 2.67E-10 |
| | CBDmale | VehFemale | 1.52E-10 | 0.0009 | -8.37E-10 | -2.29E-10 |
| | | CBDfemale | 1.52E-10 | 0.7908 | -2.64E-10 | 3.45E-10 |
| | | THCfemale | 1.52E-10 | 0.8070 | -3.42E-10 | 2.67E-10 |
| | | TpCfemale | 1.38E-10 | 0.0518 | -5.53E-10 | 2.24E-12 |
| | | VehMale | 1.59E-10 | 0.8212 | -3.54E-10 | 2.82E-10 |
| | | THCmale | 1.52E-10 | 0.9864 | -3.07E-10 | 3.02E-10 |
| | | TpCmale | 1.42E-10 | 0.5394 | -3.72E-10 | 1.97E-10 |
| | THCmale | VehFemale | 1.61E-10 | 0.0017 | -8.53E-10 | -2.08E-10 |
| | | CBDfemale | 1.61E-10 | 0.7901 | -2.80E-10 | 3.66E-10 |
| | | THCfemale | 1.61E-10 | 0.8303 | -3.57E-10 | 2.88E-10 |
| | | TpCfemale | 1.48E-10 | 0.0718 | -5.70E-10 | 2.50E-11 |
| | | VehMale | 1.68E-10 | 0.8425 | -3.69E-10 | 3.03E-10 |
| | | CBDmale | 1.52E-10 | 0.9864 | -3.02E-10 | 3.07E-10 |
| | | TpCmale | 1.52E-10 | 0.5773 | -3.89E-10 | 2.19E-10 |
| | TpCmale | VehFemale | 1.52E-10 | 0.0049 | -7.50E-10 | -1.41E-10 |
| | | CBDfemale | 1.52E-10 | 0.4022 | -1.76E-10 | 4.32E-10 |
| | | THCfemale | 1.52E-10 | 0.7410 | -2.54E-10 | 3.55E-10 |
| | | TpCfemale | 1.38E-10 | 0.1811 | -4.65E-10 | 8.99E-11 |
| | | VehMale | 1.59E-10 | 0.7461 | -2.67E-10 | 3.70E-10 |
| | | CBDmale | 1.42E-10 | 0.5394 | -1.97E-10 | 3.72E-10 |
| | | THCmale | 1.52E-10 | 0.5773 | -2.19E-10 | 3.89E-10 |
| *N*-docosahexaenoyl phenylalanine | VehFemale | CBDfemale | 2.12E-10 | 0.1810 | -1.38E-10 | 7.13E-10 |
| | | THCfemale | 2.12E-10 | 0.2598 | -1.84E-10 | 6.68E-10 |
| | | TpCfemale | 2.08E-10 | 0.2106 | -6.80E-10 | 1.53E-10 |
| | | VehMale | 2.18E-10 | 0.3486 | -2.31E-10 | 6.43E-10 |
| | | CBDmale | 2.08E-10 | 0.1319 | -9.88E-11 | 7.34E-10 |
| | | THCmale | 2.18E-10 | 0.9238 | -4.16E-10 | 4.58E-10 |
| | | TpCmale | 2.18E-10 | 0.1279 | -1.00E-10 | 7.74E-10 |
| | CBDfemale | VehFemale | 2.12E-10 | 0.1810 | -7.13E-10 | 1.38E-10 |
| | | THCfemale | 1.86E-10 | 0.8058 | -4.19E-10 | 3.27E-10 |
| | | TpCfemale | 1.81E-10 | 0.0036 | -9.14E-10 | -1.88E-10 |
| | | VehMale | 1.93E-10 | 0.6737 | -4.68E-10 | 3.05E-10 |
| | | CBDmale | 1.81E-10 | 0.8685 | -3.33E-10 | 3.93E-10 |
| | | THCmale | 1.93E-10 | 0.1720 | -6.53E-10 | 1.20E-10 |
| | | TpCmale | 1.93E-10 | 0.7985 | -3.37E-10 | 4.36E-10 |
| | THCfemale | VehFemale | 2.12E-10 | 0.2598 | -6.68E-10 | 1.84E-10 |
| | | CBDfemale | 1.86E-10 | 0.8058 | -3.27E-10 | 4.19E-10 |
| | | TpCfemale | 1.81E-10 | 0.0073 | -8.68E-10 | -1.42E-10 |

| | | | | | |
|---|---|---|---|---|---|
| | VehMale | 1.93E-10 | 0.8541 | -4.22E-10 | 3.51E-10 |
| | CBDmale | 1.81E-10 | 0.6757 | -2.87E-10 | 4.39E-10 |
| | THCmale | 1.93E-10 | 0.2570 | -6.07E-10 | 1.66E-10 |
| | TpCmale | 1.93E-10 | 0.6225 | -2.91E-10 | 4.82E-10 |
| TpCfemale | VehFemale | 2.08E-10 | 0.2106 | -1.53E-10 | 6.80E-10 |
| | CBDfemale | 1.81E-10 | 0.0036 | 1.88E-10 | 9.14E-10 |
| | THCfemale | 1.81E-10 | 0.0073 | 1.42E-10 | 8.68E-10 |
| | VehMale | 1.88E-10 | 0.0155 | 9.29E-11 | 8.46E-10 |
| | CBDmale | 1.75E-10 | 0.0017 | 2.29E-10 | 9.33E-10 |
| | THCmale | 1.88E-10 | 0.1359 | -9.23E-11 | 6.60E-10 |
| | TpCmale | 1.88E-10 | 0.0023 | 2.24E-10 | 9.77E-10 |
| VehMale | VehFemale | 2.18E-10 | 0.3486 | -6.43E-10 | 2.31E-10 |
| | CBDfemale | 1.93E-10 | 0.6737 | -3.05E-10 | 4.68E-10 |
| | THCfemale | 1.93E-10 | 0.8541 | -3.51E-10 | 4.22E-10 |
| | TpCfemale | 1.88E-10 | 0.0155 | -8.46E-10 | -9.29E-11 |
| | CBDmale | 1.88E-10 | 0.5542 | -2.65E-10 | 4.88E-10 |
| | THCmale | 1.99E-10 | 0.3562 | -5.84E-10 | 2.14E-10 |
| | TpCmale | 1.99E-10 | 0.5131 | -2.68E-10 | 5.30E-10 |
| CBDmale | VehFemale | 2.08E-10 | 0.1319 | -7.34E-10 | 9.88E-11 |
| | CBDfemale | 1.81E-10 | 0.8685 | -3.93E-10 | 3.33E-10 |
| | THCfemale | 1.81E-10 | 0.6757 | -4.39E-10 | 2.87E-10 |
| | TpCfemale | 1.75E-10 | 0.0017 | -9.33E-10 | -2.29E-10 |
| | VehMale | 1.88E-10 | 0.5542 | -4.88E-10 | 2.65E-10 |
| | THCmale | 1.88E-10 | 0.1196 | -6.73E-10 | 7.95E-11 |
| | TpCmale | 1.88E-10 | 0.9184 | -3.57E-10 | 3.96E-10 |
| THCmale | VehFemale | 2.18E-10 | 0.9238 | -4.58E-10 | 4.16E-10 |
| | CBDfemale | 1.93E-10 | 0.1720 | -1.20E-10 | 6.53E-10 |
| | THCfemale | 1.93E-10 | 0.2570 | -1.66E-10 | 6.07E-10 |
| | TpCfemale | 1.88E-10 | 0.1359 | -6.60E-10 | 9.23E-11 |
| | VehMale | 1.99E-10 | 0.3562 | -2.14E-10 | 5.84E-10 |
| | CBDmale | 1.88E-10 | 0.1196 | -7.95E-11 | 6.73E-10 |
| | TpCmale | 1.99E-10 | 0.1181 | -8.30E-11 | 7.15E-10 |
| TpCmale | VehFemale | 2.18E-10 | 0.1279 | -7.74E-10 | 1.00E-10 |
| | CBDfemale | 1.93E-10 | 0.7985 | -4.36E-10 | 3.37E-10 |
| | THCfemale | 1.93E-10 | 0.6225 | -4.82E-10 | 2.91E-10 |
| | TpCfemale | 1.88E-10 | 0.0023 | -9.77E-10 | -2.24E-10 |
| | VehMale | 1.99E-10 | 0.5131 | -5.30E-10 | 2.68E-10 |
| | CBDmale | 1.88E-10 | 0.9184 | -3.96E-10 | 3.57E-10 |
| | THCmale | 1.99E-10 | 0.1181 | -7.15E-10 | 8.30E-11 |

#### Supplemental Table 6.8 Fisher's LSD post-hoc test for liver *N*-acyl phenylalanine ANOVA

| Dependent Variable | | | Std. Error | Sig. | 95% Confidence Interval Lower Bound | Upper Bound |
|---|---|---|---|---|---|---|
| | VehFemale | CBDfemale | 8.02326E-12 | 0.278 | -2.4819E-11 | 7.2476E-12 |
| | | THCfemale | 7.35344E-12 | 0.287 | -2.2596E-11 | 6.7934E-12 |
| | | TpCfemale | 7.35344E-12 | 0.003 | -3.7125E-11 | -7.7358E-12 |
| | | VehMale | 8.02326E-12 | 0.445 | -9.8706E-12 | 2.2196E-11 |
| | | CBDmale | 7.35344E-12 | 0.073 | -1.2673E-12 | 2.8122E-11 |
| | | THCmale | 7.53282E-12 | 0.252 | -2.3759E-11 | 6.3476E-12 |
| | | TpCmale | 7.35344E-12 | 0.074 | -1.3545E-12 | 2.8035E-11 |
| | CBDfemale | VehFemale | 8.02326E-12 | 0.278 | -7.2476E-12 | 2.4819E-11 |
| | | THCfemale | 7.63967E-12 | 0.908 | -1.4382E-11 | 1.6151E-11 |
| | | TpCfemale | 7.63967E-12 | 0.079 | -2.8912E-11 | 1.6218E-12 |
| | | VehMale | 8.28639E-12 | 0.076 | -1.6108E-12 | 3.1507E-11 |
| | | CBDmale | 7.63967E-12 | 0.005 | 6.9462E-12 | 3.7480E-11 |
| | | THCmale | 7.81248E-12 | 0.992 | -1.5532E-11 | 1.5692E-11 |
| | | TpCmale | 7.63967E-12 | 0.005 | 6.8591E-12 | 3.7392E-11 |
| | THCfemale | VehFemale | 7.35344E-12 | 0.287 | -6.7934E-12 | 2.2596E-11 |
| | | CBDfemale | 7.63967E-12 | 0.908 | -1.6151E-11 | 1.4382E-11 |
| | | TpCfemale | 6.93289E-12 | 0.040 | -2.8383E-11 | -6.7491E-13 |
| | | VehMale | 7.63967E-12 | 0.070 | -1.2028E-12 | 2.9331E-11 |
| | | CBDmale | 6.93289E-12 | 0.003 | 7.4743E-12 | 3.5183E-11 |
| | | THCmale | 7.12287E-12 | 0.910 | -1.5038E-11 | 1.3430E-11 |
| | | TpCmale | 6.93289E-12 | 0.003 | 7.3872E-12 | 3.5096E-11 |
| | TpCfemale | VehFemale | 7.35344E-12 | 0.003 | 7.7358E-12 | 3.7125E-11 |
| | | CBDfemale | 7.63967E-12 | 0.079 | -1.6218E-12 | 2.8912E-11 |
| | | THCfemale | 6.93289E-12 | 0.040 | 6.7491E-13 | 2.8383E-11 |
| | | VehMale | 7.63967E-12 | 0.000 | 1.3326E-11 | 4.3860E-11 |
| | | CBDmale | 6.93289E-12 | 0.000 | 2.2003E-11 | 4.9712E-11 |
| | | THCmale | 7.12287E-12 | 0.059 | -5.0898E-13 | 2.7959E-11 |
| | | TpCmale | 6.93289E-12 | 0.000 | 2.1916E-11 | 4.9625E-11 |
| *N*-palmitoyl phenylalanine | VehMale | VehFemale | 8.02326E-12 | 0.445 | -2.2196E-11 | 9.8706E-12 |
| | | CBDfemale | 8.28639E-12 | 0.076 | -3.1507E-11 | 1.6108E-12 |
| | | THCfemale | 7.63967E-12 | 0.070 | -2.9331E-11 | 1.2028E-12 |
| | | TpCfemale | 7.63967E-12 | 0.000 | -4.3860E-11 | -1.3326E-11 |
| | | CBDmale | 7.63967E-12 | 0.345 | -8.0020E-12 | 2.2531E-11 |
| | | THCmale | 7.81248E-12 | 0.062 | -3.0480E-11 | 7.4386E-13 |
| | | TpCmale | 7.63967E-12 | 0.351 | -8.0891E-12 | 2.2444E-11 |
| | CBDmale | VehFemale | 7.35344E-12 | 0.073 | -2.8122E-11 | 1.2673E-12 |
| | | CBDfemale | 7.63967E-12 | 0.005 | -3.7480E-11 | -6.9462E-12 |
| | | THCfemale | 6.93289E-12 | 0.003 | -3.5183E-11 | -7.4743E-12 |
| | | TpCfemale | 6.93289E-12 | 0.000 | -4.9712E-11 | -2.2003E-11 |
| | | VehMale | 7.63967E-12 | 0.345 | -2.2531E-11 | 8.0020E-12 |
| | | THCmale | 7.12287E-12 | 0.003 | -3.6367E-11 | -7.8989E-12 |
| | | TpCmale | 6.93289E-12 | 0.990 | -1.3941E-11 | 1.3767E-11 |
| | THCmale | VehFemale | 7.53282E-12 | 0.252 | -6.3476E-12 | 2.3759E-11 |
| | | CBDfemale | 7.81248E-12 | 0.992 | -1.5692E-11 | 1.5532E-11 |
| | | THCfemale | 7.12287E-12 | 0.910 | -1.3430E-11 | 1.5038E-11 |
| | | TpCfemale | 7.12287E-12 | 0.059 | -2.7959E-11 | 5.0898E-13 |
| | | VehMale | 7.81248E-12 | 0.062 | -7.4386E-13 | 3.0480E-11 |
| | | CBDmale | 7.12287E-12 | 0.003 | 7.8989E-12 | 3.6367E-11 |
| | | TpCmale | 7.12287E-12 | 0.003 | 7.8118E-12 | 3.6280E-11 |
| | TpCmale | VehFemale | 7.35344E-12 | 0.074 | -2.8035E-11 | 1.3545E-12 |
| | | CBDfemale | 7.63967E-12 | 0.005 | -3.7392E-11 | -6.8591E-12 |
| | | THCfemale | 6.93289E-12 | 0.003 | -3.5096E-11 | -7.3872E-12 |
| | | TpCfemale | 6.93289E-12 | 0.000 | -4.9625E-11 | -2.1916E-11 |
| | | VehMale | 7.63967E-12 | 0.351 | -2.2444E-11 | 8.0891E-12 |
| | | CBDmale | 6.93289E-12 | 0.990 | -1.3767E-11 | 1.3941E-11 |
| | | THCmale | 7.12287E-12 | 0.003 | -3.6280E-11 | -7.8118E-12 |
| *N*-stearoyl phenylalanine | VehFemale | CBDfemale | 8.72931E-13 | 0.570 | -1.2473E-12 | 2.2438E-12 |

| | | | | | | |
|---|---|---|---|---|---|---|
| | | THCfemale | 8.00055E-13 | 0.721 | -1.8866E-12 | 1.3131E-12 |
| | | TpCfemale | 8.19571E-13 | 0.350 | -2.4111E-12 | 8.6656E-13 |
| | | VehMale | 8.72931E-13 | 0.055 | -3.5794E-14 | 3.4553E-12 |
| | | CBDmale | 8.19571E-13 | 0.009 | 5.7454E-13 | 3.8522E-12 |
| | | THCmale | 8.19571E-13 | 0.058 | -5.3896E-14 | 3.2238E-12 |
| | | TpCmale | 8.00055E-13 | 0.005 | 7.2620E-13 | 3.9258E-12 |
| | CBDfemale | VehFemale | 8.72931E-13 | 0.570 | -2.2438E-12 | 1.2473E-12 |
| | | THCfemale | 8.31197E-13 | 0.349 | -2.4471E-12 | 8.7706E-13 |
| | | TpCfemale | 8.49998E-13 | 0.140 | -2.9702E-12 | 4.2913E-13 |
| | | VehMale | 9.01559E-13 | 0.184 | -5.9131E-13 | 3.0142E-12 |
| | | CBDmale | 8.49998E-13 | 0.048 | 1.5430E-14 | 3.4148E-12 |
| | | THCmale | 8.49998E-13 | 0.206 | -6.1301E-13 | 2.7863E-12 |
| | | TpCmale | 8.31197E-13 | 0.032 | 1.6565E-13 | 3.4898E-12 |
| | THCfemale | VehFemale | 8.00055E-13 | 0.721 | -1.3131E-12 | 1.8866E-12 |
| | | CBDfemale | 8.31197E-13 | 0.349 | -8.7706E-13 | 2.4471E-12 |
| | | TpCfemale | 7.74968E-13 | 0.533 | -2.0352E-12 | 1.0641E-12 |
| | | VehMale | 8.31197E-13 | 0.019 | 3.3441E-13 | 3.6586E-12 |
| | | CBDmale | 7.74968E-13 | 0.002 | 9.5048E-13 | 4.0498E-12 |
| | | THCmale | 7.74968E-13 | 0.019 | 3.2204E-13 | 3.4213E-12 |
| | | TpCmale | 7.54299E-13 | 0.001 | 1.1044E-12 | 4.1211E-12 |
| | TpCfemale | VehFemale | 8.19571E-13 | 0.350 | -8.6656E-13 | 2.4111E-12 |
| | | CBDfemale | 8.49998E-13 | 0.140 | -4.2913E-13 | 2.9702E-12 |
| | | THCfemale | 7.74968E-13 | 0.533 | -1.0641E-12 | 2.0352E-12 |
| | | VehMale | 8.49998E-13 | 0.005 | 7.8234E-13 | 4.1817E-12 |
| | | CBDmale | 7.95101E-13 | 0.000 | 1.3957E-12 | 4.5756E-12 |
| | | THCmale | 7.95101E-13 | 0.004 | 7.6731E-13 | 3.9471E-12 |
| | | TpCmale | 7.74968E-13 | 0.000 | 1.5486E-12 | 4.6479E-12 |
| | VehMale | VehFemale | 8.72931E-13 | 0.055 | -3.4553E-12 | 3.5794E-14 |
| | | CBDfemale | 9.01559E-13 | 0.184 | -3.0142E-12 | 5.9131E-13 |
| | | THCfemale | 8.31197E-13 | 0.019 | -3.6586E-12 | -3.3441E-13 |
| | | TpCfemale | 8.49998E-13 | 0.005 | -4.1817E-12 | -7.8234E-13 |
| | | CBDmale | 8.49998E-13 | 0.556 | -1.1960E-12 | 2.2033E-12 |
| | | THCmale | 8.49998E-13 | 0.884 | -1.8245E-12 | 1.5749E-12 |
| | | TpCmale | 8.31197E-13 | 0.461 | -1.0458E-12 | 2.2783E-12 |
| | CBDmale | VehFemale | 8.19571E-13 | 0.009 | -3.8522E-12 | -5.7454E-13 |
| | | CBDfemale | 8.49998E-13 | 0.048 | -3.4148E-12 | -1.5430E-14 |
| | | THCfemale | 7.74968E-13 | 0.002 | -4.0498E-12 | -9.5048E-13 |
| | | TpCfemale | 7.95101E-13 | 0.000 | -4.5756E-12 | -1.3957E-12 |
| | | VehMale | 8.49998E-13 | 0.556 | -2.2033E-12 | 1.1960E-12 |
| | | THCmale | 7.95101E-13 | 0.432 | -2.2183E-12 | 9.6146E-13 |
| | | TpCmale | 7.74968E-13 | 0.885 | -1.4370E-12 | 1.6623E-12 |
| | THCmale | VehFemale | 8.19571E-13 | 0.058 | -3.2238E-12 | 5.3896E-14 |
| | | CBDfemale | 8.49998E-13 | 0.206 | -2.7863E-12 | 6.1301E-13 |
| | | THCfemale | 7.74968E-13 | 0.019 | -3.4213E-12 | -3.2204E-13 |
| | | TpCfemale | 7.95101E-13 | 0.004 | -3.9471E-12 | -7.6731E-13 |
| | | VehMale | 8.49998E-13 | 0.884 | -1.5749E-12 | 1.8245E-12 |
| | | CBDmale | 7.95101E-13 | 0.432 | -9.6146E-13 | 2.2183E-12 |
| | | TpCmale | 7.74968E-13 | 0.343 | -8.0858E-13 | 2.2907E-12 |
| | TpCmale | VehFemale | 8.00055E-13 | 0.005 | -3.9258E-12 | -7.2620E-13 |
| | | CBDfemale | 8.31197E-13 | 0.032 | -3.4898E-12 | -1.6565E-13 |
| | | THCfemale | 7.54299E-13 | 0.001 | -4.1211E-12 | -1.1044E-12 |
| | | TpCfemale | 7.74968E-13 | 0.000 | -4.6479E-12 | -1.5486E-12 |
| | | VehMale | 8.31197E-13 | 0.461 | -2.2783E-12 | 1.0458E-12 |
| | | CBDmale | 7.74968E-13 | 0.885 | -1.6623E-12 | 1.4370E-12 |
| | | THCmale | 7.74968E-13 | 0.343 | -2.2907E-12 | 8.0858E-13 |
| *N*-oleoyl phenylalanine | VehFemale | CBDfemale | 5.89711E-12 | 0.700 | -1.4072E-11 | 9.5040E-12 |
| | | THCfemale | 5.40479E-12 | 0.001 | -3.0635E-11 | -9.0267E-12 |
| | | TpCfemale | 5.40479E-12 | 0.000 | -3.8756E-11 | -1.7148E-11 |
| | | VehMale | 5.89711E-12 | 0.443 | -7.2388E-12 | 1.6338E-11 |
| | | CBDmale | 5.40479E-12 | 0.146 | -2.8430E-12 | 1.8765E-11 |

| | | | | | | |
|---|---|---|---|---|---|---|
| | | THCmale | 5.69715E-12 | 0.913 | -1.2013E-11 | 1.0764E-11 |
| | | TpCmale | 5.40479E-12 | 0.510 | -7.2232E-12 | 1.4385E-11 |
| | CBDfemale | VehFemale | 5.89711E-12 | 0.700 | -9.5040E-12 | 1.4072E-11 |
| | | THCfemale | 5.61518E-12 | 0.003 | -2.8771E-11 | -6.3219E-12 |
| | | TpCfemale | 5.61518E-12 | 0.000 | -3.6893E-11 | -1.4444E-11 |
| | | VehMale | 6.09051E-12 | 0.266 | -5.3412E-12 | 1.9008E-11 |
| | | CBDmale | 5.61518E-12 | 0.073 | -9.7938E-13 | 2.1470E-11 |
| | | THCmale | 5.89711E-12 | 0.779 | -1.0129E-11 | 1.3448E-11 |
| | | TpCmale | 5.61518E-12 | 0.300 | -5.3596E-12 | 1.7090E-11 |
| | THCfemale | VehFemale | 5.40479E-12 | 0.001 | 9.0267E-12 | 3.0635E-11 |
| | | CBDfemale | 5.61518E-12 | 0.003 | 6.3219E-12 | 2.8771E-11 |
| | | TpCfemale | 5.09569E-12 | 0.116 | -1.8308E-11 | 2.0644E-12 |
| | | VehMale | 5.61518E-12 | 0.000 | 1.3156E-11 | 3.5605E-11 |
| | | CBDmale | 5.09569E-12 | 0.000 | 1.7606E-11 | 3.7978E-11 |
| | | THCmale | 5.40479E-12 | 0.001 | 8.4019E-12 | 3.0010E-11 |
| | | TpCmale | 5.09569E-12 | 0.000 | 1.3225E-11 | 3.3598E-11 |
| | TpCfemale | VehFemale | 5.40479E-12 | 0.000 | 1.7148E-11 | 3.8756E-11 |
| | | CBDfemale | 5.61518E-12 | 0.000 | 1.4444E-11 | 3.6893E-11 |
| | | THCfemale | 5.09569E-12 | 0.116 | -2.0644E-12 | 1.8308E-11 |
| | | VehMale | 5.61518E-12 | 0.000 | 2.1277E-11 | 4.3726E-11 |
| | | CBDmale | 5.09569E-12 | 0.000 | 2.5727E-11 | 4.6100E-11 |
| | | THCmale | 5.40479E-12 | 0.000 | 1.6524E-11 | 3.8132E-11 |
| | | TpCmale | 5.09569E-12 | 0.000 | 2.1347E-11 | 4.1719E-11 |
| | VehMale | VehFemale | 5.89711E-12 | 0.443 | -1.6338E-11 | 7.2388E-12 |
| | | CBDfemale | 6.09051E-12 | 0.266 | -1.9008E-11 | 5.3412E-12 |
| | | THCfemale | 5.61518E-12 | 0.000 | -3.5605E-11 | -1.3156E-11 |
| | | TpCfemale | 5.61518E-12 | 0.000 | -4.3726E-11 | -2.1277E-11 |
| | | CBDmale | 5.61518E-12 | 0.546 | -7.8129E-12 | 1.4636E-11 |
| | | THCmale | 5.89711E-12 | 0.384 | -1.6962E-11 | 6.6140E-12 |
| | | TpCmale | 5.61518E-12 | 0.864 | -1.2193E-11 | 1.0256E-11 |
| | CBDmale | VehFemale | 5.40479E-12 | 0.146 | -1.8765E-11 | 2.8430E-12 |
| | | CBDfemale | 5.61518E-12 | 0.073 | -2.1470E-11 | 9.7938E-13 |
| | | THCfemale | 5.09569E-12 | 0.000 | -3.7978E-11 | -1.7606E-11 |
| | | TpCfemale | 5.09569E-12 | 0.000 | -4.6100E-11 | -2.5727E-11 |
| | | VehMale | 5.61518E-12 | 0.546 | -1.4636E-11 | 7.8129E-12 |
| | | THCmale | 5.40479E-12 | 0.117 | -1.9390E-11 | 2.2183E-12 |
| | | TpCmale | 5.09569E-12 | 0.393 | -1.4566E-11 | 5.8059E-12 |
| | THCmale | VehFemale | 5.69715E-12 | 0.913 | -1.0764E-11 | 1.2013E-11 |
| | | CBDfemale | 5.89711E-12 | 0.779 | -1.3448E-11 | 1.0129E-11 |
| | | THCfemale | 5.40479E-12 | 0.001 | -3.0010E-11 | -8.4019E-12 |
| | | TpCfemale | 5.40479E-12 | 0.000 | -3.8132E-11 | -1.6524E-11 |
| | | VehMale | 5.89711E-12 | 0.384 | -6.6140E-12 | 1.6962E-11 |
| | | CBDmale | 5.40479E-12 | 0.117 | -2.2183E-12 | 1.9390E-11 |
| | | TpCmale | 5.40479E-12 | 0.439 | -6.5985E-12 | 1.5010E-11 |
| | TpCmale | VehFemale | 5.40479E-12 | 0.510 | -1.4385E-11 | 7.2232E-12 |
| | | CBDfemale | 5.61518E-12 | 0.300 | -1.7090E-11 | 5.3596E-12 |
| | | THCfemale | 5.09569E-12 | 0.000 | -3.3598E-11 | -1.3225E-11 |
| | | TpCfemale | 5.09569E-12 | 0.000 | -4.1719E-11 | -2.1347E-11 |
| | | VehMale | 5.61518E-12 | 0.864 | -1.0256E-11 | 1.2193E-11 |
| | | CBDmale | 5.09569E-12 | 0.393 | -5.8059E-12 | 1.4566E-11 |
| | | THCmale | 5.40479E-12 | 0.439 | -1.5010E-11 | 6.5985E-12 |
| *N*-linoleoyl phenylalanine | VehFemale | CBDfemale | 4.60578E-12 | 0.429 | -1.2877E-11 | 5.5486E-12 |
| | | THCfemale | 4.32424E-12 | 0.735 | -1.0119E-11 | 7.1801E-12 |
| | | TpCfemale | 4.32424E-12 | 0.000 | -3.3928E-11 | -1.6629E-11 |
| | | VehMale | 4.60578E-12 | 0.114 | -1.8363E-12 | 1.6590E-11 |
| | | CBDmale | 4.32424E-12 | 0.009 | 3.0395E-12 | 2.0339E-11 |
| | | THCmale | 4.32424E-12 | 0.324 | -1.2949E-11 | 4.3504E-12 |
| | | TpCmale | 4.22127E-12 | 0.069 | -6.1969E-13 | 1.6268E-11 |
| | CBDfemale | VehFemale | 4.60578E-12 | 0.429 | -5.5486E-12 | 1.2877E-11 |
| | | THCfemale | 4.48478E-12 | 0.626 | -6.7763E-12 | 1.1166E-11 |

| | | | | | | |
|---|---|---|---|---|---|---|
| | | TpCfemale | 4.48478E-12 | 0.000 | -3.0585E-11 | -1.2643E-11 |
| | | VehMale | 4.75683E-12 | 0.024 | 1.5258E-12 | 2.0556E-11 |
| | | CBDmale | 4.48478E-12 | 0.001 | 6.3827E-12 | 2.4324E-11 |
| | | THCmale | 4.48478E-12 | 0.888 | -9.6060E-12 | 8.3358E-12 |
| | | TpCmale | 4.38558E-12 | 0.011 | 2.7159E-12 | 2.0261E-11 |
| | THCfemale | VehFemale | 4.32424E-12 | 0.735 | -7.1801E-12 | 1.0119E-11 |
| | | CBDfemale | 4.48478E-12 | 0.626 | -1.1166E-11 | 6.7763E-12 |
| | | TpCfemale | 4.19513E-12 | 0.000 | -3.2200E-11 | -1.5417E-11 |
| | | VehMale | 4.48478E-12 | 0.053 | -1.2458E-13 | 1.7817E-11 |
| | | CBDmale | 4.19513E-12 | 0.003 | 4.7675E-12 | 2.1550E-11 |
| | | THCmale | 4.19513E-12 | 0.503 | -1.1221E-11 | 5.5618E-12 |
| | | TpCmale | 4.08891E-12 | 0.027 | 1.1148E-12 | 1.7473E-11 |
| | TpCfemale | VehFemale | 4.32424E-12 | 0.000 | 1.6629E-11 | 3.3928E-11 |
| | | CBDfemale | 4.48478E-12 | 0.000 | 1.2643E-11 | 3.0585E-11 |
| | | THCfemale | 4.19513E-12 | 0.000 | 1.5417E-11 | 3.2200E-11 |
| | | VehMale | 4.48478E-12 | 0.000 | 2.3684E-11 | 4.1626E-11 |
| | | CBDmale | 4.19513E-12 | 0.000 | 2.8576E-11 | 4.5359E-11 |
| | | THCmale | 4.19513E-12 | 0.000 | 1.2587E-11 | 2.9370E-11 |
| | | TpCmale | 4.08891E-12 | 0.000 | 2.4923E-11 | 4.1282E-11 |
| | VehMale | VehFemale | 4.60578E-12 | 0.114 | -1.6590E-11 | 1.8363E-12 |
| | | CBDfemale | 4.75683E-12 | 0.024 | -2.0556E-11 | -1.5258E-12 |
| | | THCfemale | 4.48478E-12 | 0.053 | -1.7817E-11 | 1.2458E-13 |
| | | TpCfemale | 4.48478E-12 | 0.000 | -4.1626E-11 | -2.3684E-11 |
| | | CBDmale | 4.48478E-12 | 0.340 | -4.6583E-12 | 1.3284E-11 |
| | | THCmale | 4.48478E-12 | 0.012 | -2.0647E-11 | -2.7051E-12 |
| | | TpCmale | 4.38558E-12 | 0.919 | -8.3250E-12 | 9.2199E-12 |
| | CBDmale | VehFemale | 4.32424E-12 | 0.009 | -2.0339E-11 | -3.0395E-12 |
| | | CBDfemale | 4.48478E-12 | 0.001 | -2.4324E-11 | -6.3827E-12 |
| | | THCfemale | 4.19513E-12 | 0.003 | -2.1550E-11 | -4.7675E-12 |
| | | TpCfemale | 4.19513E-12 | 0.000 | -4.5359E-11 | -2.8576E-11 |
| | | VehMale | 4.48478E-12 | 0.340 | -1.3284E-11 | 4.6583E-12 |
| | | THCmale | 4.19513E-12 | 0.000 | -2.4380E-11 | -7.5972E-12 |
| | | TpCmale | 4.08891E-12 | 0.348 | -1.2044E-11 | 4.3139E-12 |
| | THCmale | VehFemale | 4.32424E-12 | 0.324 | -4.3504E-12 | 1.2949E-11 |
| | | CBDfemale | 4.48478E-12 | 0.888 | -8.3358E-12 | 9.6060E-12 |
| | | THCfemale | 4.19513E-12 | 0.503 | -5.5618E-12 | 1.1221E-11 |
| | | TpCfemale | 4.19513E-12 | 0.000 | -2.9370E-11 | -1.2587E-11 |
| | | VehMale | 4.48478E-12 | 0.012 | 2.7051E-12 | 2.0647E-11 |
| | | CBDmale | 4.19513E-12 | 0.000 | 7.5972E-12 | 2.4380E-11 |
| | | TpCmale | 4.08891E-12 | 0.004 | 3.9445E-12 | 2.0303E-11 |
| | TpCmale | VehFemale | 4.22127E-12 | 0.069 | -1.6268E-11 | 6.1969E-13 |
| | | CBDfemale | 4.38558E-12 | 0.011 | -2.0261E-11 | -2.7159E-12 |
| | | THCfemale | 4.08891E-12 | 0.027 | -1.7473E-11 | -1.1148E-12 |
| | | TpCfemale | 4.08891E-12 | 0.000 | -4.1282E-11 | -2.4923E-11 |
| | | VehMale | 4.38558E-12 | 0.919 | -9.2199E-12 | 8.3250E-12 |
| | | CBDmale | 4.08891E-12 | 0.348 | -4.3139E-12 | 1.2044E-11 |
| | | THCmale | 4.08891E-12 | 0.004 | -2.0303E-11 | -3.9445E-12 |
| | VehFemale | CBDfemale | 6.81864E-12 | 0.055 | -2.6969E-11 | 2.9201E-13 |
| | | THCfemale | 6.40183E-12 | 0.175 | -2.1572E-11 | 4.0217E-12 |
| | | TpCfemale | 6.24939E-12 | 0.001 | -3.4524E-11 | -9.5397E-12 |
| | | VehMale | 6.81864E-12 | 0.590 | -9.9342E-12 | 1.7326E-11 |
| | | CBDmale | 6.24939E-12 | 0.135 | -3.0361E-12 | 2.1949E-11 |
| | | THCmale | 6.40183E-12 | 0.131 | -2.2585E-11 | 3.0089E-12 |
| *N*-arachidonoyl phenylalanine | | TpCmale | 6.24939E-12 | 0.203 | -4.4465E-12 | 2.0538E-11 |
| | CBDfemale | VehFemale | 6.81864E-12 | 0.055 | -2.9201E-13 | 2.6969E-11 |
| | | THCfemale | 6.63951E-12 | 0.494 | -8.7093E-12 | 1.7835E-11 |
| | | TpCfemale | 6.49265E-12 | 0.185 | -2.1672E-11 | 4.2848E-12 |
| | | VehMale | 7.04226E-12 | 0.019 | 2.9571E-12 | 3.1112E-11 |
| | | CBDmale | 6.49265E-12 | 0.001 | 9.8159E-12 | 3.5773E-11 |
| | | THCmale | 6.63951E-12 | 0.595 | -9.7221E-12 | 1.6822E-11 |

| | | | | | | |
|---|---|---|---|---|---|---|
| | | TpCmale | 6.49265E-12 | 0.002 | 8.4055E-12 | 3.4363E-11 |
| | THCfemale | VehFemale | 6.40183E-12 | 0.175 | -4.0217E-12 | 2.1572E-11 |
| | | CBDfemale | 6.63951E-12 | 0.494 | -1.7835E-11 | 8.7093E-12 |
| | | TpCfemale | 6.05343E-12 | 0.032 | -2.5357E-11 | -1.1560E-12 |
| | | VehMale | 6.63951E-12 | 0.065 | -8.0074E-13 | 2.5744E-11 |
| | | CBDmale | 6.05343E-12 | 0.004 | 6.1310E-12 | 3.0332E-11 |
| | | THCmale | 6.21069E-12 | 0.871 | -1.3428E-11 | 1.1402E-11 |
| | | TpCmale | 6.05343E-12 | 0.007 | 4.7205E-12 | 2.8922E-11 |
| | TpCfemale | VehFemale | 6.24939E-12 | 0.001 | 9.5397E-12 | 3.4524E-11 |
| | | CBDfemale | 6.49265E-12 | 0.185 | -4.2848E-12 | 2.1672E-11 |
| | | THCfemale | 6.05343E-12 | 0.032 | 1.1560E-12 | 2.5357E-11 |
| | | VehMale | 6.49265E-12 | 0.000 | 1.2750E-11 | 3.8707E-11 |
| | | CBDmale | 5.89198E-12 | 0.000 | 1.9710E-11 | 4.3266E-11 |
| | | THCmale | 6.05343E-12 | 0.047 | 1.4320E-13 | 2.4344E-11 |
| | | TpCmale | 5.89198E-12 | 0.000 | 1.8300E-11 | 4.1856E-11 |
| | VehMale | VehFemale | 6.81864E-12 | 0.590 | -1.7326E-11 | 9.9342E-12 |
| | | CBDfemale | 7.04226E-12 | 0.019 | -3.1112E-11 | -2.9571E-12 |
| | | THCfemale | 6.63951E-12 | 0.065 | -2.5744E-11 | 8.0074E-13 |
| | | TpCfemale | 6.49265E-12 | 0.000 | -3.8707E-11 | -1.2750E-11 |
| | | CBDmale | 6.49265E-12 | 0.378 | -7.2185E-12 | 1.8739E-11 |
| | | THCmale | 6.63951E-12 | 0.047 | -2.6756E-11 | -2.1210E-13 |
| | | TpCmale | 6.49265E-12 | 0.505 | -8.6289E-12 | 1.7328E-11 |
| | CBDmale | VehFemale | 6.24939E-12 | 0.135 | -2.1949E-11 | 3.0361E-12 |
| | | CBDfemale | 6.49265E-12 | 0.001 | -3.5773E-11 | -9.8159E-12 |
| | | THCfemale | 6.05343E-12 | 0.004 | -3.0332E-11 | -6.1310E-12 |
| | | TpCfemale | 5.89198E-12 | 0.000 | -4.3266E-11 | -1.9710E-11 |
| | | VehMale | 6.49265E-12 | 0.378 | -1.8739E-11 | 7.2185E-12 |
| | | THCmale | 6.05343E-12 | 0.002 | -3.1345E-11 | -7.1438E-12 |
| | | TpCmale | 5.89198E-12 | 0.812 | -1.3188E-11 | 1.0367E-11 |
| | THCmale | VehFemale | 6.40183E-12 | 0.131 | -3.0089E-12 | 2.2585E-11 |
| | | CBDfemale | 6.63951E-12 | 0.595 | -1.6822E-11 | 9.7221E-12 |
| | | THCfemale | 6.21069E-12 | 0.871 | -1.1402E-11 | 1.3428E-11 |
| | | TpCfemale | 6.05343E-12 | 0.047 | -2.4344E-11 | -1.4320E-13 |
| | | VehMale | 6.63951E-12 | 0.047 | 2.1210E-13 | 2.6756E-11 |
| | | CBDmale | 6.05343E-12 | 0.002 | 7.1438E-12 | 3.1345E-11 |
| | | TpCmale | 6.05343E-12 | 0.005 | 5.7334E-12 | 2.9935E-11 |
| | TpCmale | VehFemale | 6.24939E-12 | 0.203 | -2.0538E-11 | 4.4465E-12 |
| | | CBDfemale | 6.49265E-12 | 0.002 | -3.4363E-11 | -8.4055E-12 |
| | | THCfemale | 6.05343E-12 | 0.007 | -2.8922E-11 | -4.7205E-12 |
| | | TpCfemale | 5.89198E-12 | 0.000 | -4.1856E-11 | -1.8300E-11 |
| | | VehMale | 6.49265E-12 | 0.505 | -1.7328E-11 | 8.6289E-12 |
| | | CBDmale | 5.89198E-12 | 0.812 | -1.0367E-11 | 1.3188E-11 |
| | | THCmale | 6.05343E-12 | 0.005 | -2.9935E-11 | -5.7334E-12 |
| *N*-docosahexaenoyl phenylalanine | VehFemale | CBDfemale | 5.93170E-12 | 0.022 | -2.5802E-11 | -2.0949E-12 |
| | | THCfemale | 5.62730E-12 | 0.069 | -2.1657E-11 | 8.3338E-13 |
| | | TpCfemale | 5.62730E-12 | 0.001 | -3.1285E-11 | -8.7942E-12 |
| | | VehMale | 6.13989E-12 | 0.698 | -9.8766E-12 | 1.4663E-11 |
| | | CBDmale | 5.76457E-12 | 0.343 | -6.0143E-12 | 1.7025E-11 |
| | | THCmale | 5.76457E-12 | 0.177 | -1.9387E-11 | 3.6519E-12 |
| | | TpCmale | 5.62730E-12 | 0.608 | -8.3415E-12 | 1.4149E-11 |
| | CBDfemale | VehFemale | 5.93170E-12 | 0.022 | 2.0949E-12 | 2.5802E-11 |
| | | THCfemale | 5.62730E-12 | 0.532 | -7.7087E-12 | 1.4782E-11 |
| | | TpCfemale | 5.62730E-12 | 0.283 | -1.7336E-11 | 5.1543E-12 |
| | | VehMale | 6.13989E-12 | 0.010 | 4.0719E-12 | 2.8611E-11 |
| | | CBDmale | 5.76457E-12 | 0.001 | 7.9341E-12 | 3.0973E-11 |
| | | THCmale | 5.76457E-12 | 0.296 | -5.4388E-12 | 1.7600E-11 |
| | | TpCmale | 5.62730E-12 | 0.004 | 5.6069E-12 | 2.8097E-11 |
| | THCfemale | VehFemale | 5.62730E-12 | 0.069 | -8.3338E-13 | 2.1657E-11 |
| | | CBDfemale | 5.62730E-12 | 0.532 | -1.4782E-11 | 7.7087E-12 |
| | | TpCfemale | 5.30547E-12 | 0.074 | -2.0230E-11 | 9.7459E-13 |

| | | | | | | |
|---|---|---|---|---|---|---|
| | | VehMale | 5.84634E-12 | 0.032 | 1.1219E-12 | 2.4488E-11 |
| | | CBDmale | 5.45085E-12 | 0.005 | 5.0245E-12 | 2.6810E-11 |
| | | THCmale | 5.45085E-12 | 0.642 | -8.3484E-12 | 1.3437E-11 |
| | | TpCmale | 5.30547E-12 | 0.015 | 2.7135E-12 | 2.3918E-11 |
| | TpCfemale | VehFemale | 5.62730E-12 | 0.001 | 8.7942E-12 | 3.1285E-11 |
| | | CBDfemale | 5.62730E-12 | 0.283 | -5.1543E-12 | 1.7336E-11 |
| | | THCfemale | 5.30547E-12 | 0.074 | -9.7459E-13 | 2.0230E-11 |
| | | VehMale | 5.84634E-12 | 0.000 | 1.0749E-11 | 3.4115E-11 |
| | | CBDmale | 5.45085E-12 | 0.000 | 1.4652E-11 | 3.6437E-11 |
| | | THCmale | 5.45085E-12 | 0.029 | 1.2791E-12 | 2.3064E-11 |
| | | TpCmale | 5.30547E-12 | 0.000 | 1.2341E-11 | 3.3545E-11 |
| | VehMale | VehFemale | 6.13989E-12 | 0.698 | -1.4663E-11 | 9.8766E-12 |
| | | CBDfemale | 6.13989E-12 | 0.010 | -2.8611E-11 | -4.0719E-12 |
| | | THCfemale | 5.84634E-12 | 0.032 | -2.4488E-11 | -1.1219E-12 |
| | | TpCfemale | 5.84634E-12 | 0.000 | -3.4115E-11 | -1.0749E-11 |
| | | CBDmale | 5.97859E-12 | 0.604 | -8.8350E-12 | 1.5059E-11 |
| | | THCmale | 5.97859E-12 | 0.091 | -2.2208E-11 | 1.6866E-12 |
| | | TpCmale | 5.84634E-12 | 0.931 | -1.1172E-11 | 1.2194E-11 |
| | CBDmale | VehFemale | 5.76457E-12 | 0.343 | -1.7025E-11 | 6.0143E-12 |
| | | CBDfemale | 5.76457E-12 | 0.001 | -3.0973E-11 | -7.9341E-12 |
| | | THCfemale | 5.45085E-12 | 0.005 | -2.6810E-11 | -5.0245E-12 |
| | | TpCfemale | 5.45085E-12 | 0.000 | -3.6437E-11 | -1.4652E-11 |
| | | VehMale | 5.97859E-12 | 0.604 | -1.5059E-11 | 8.8350E-12 |
| | | THCmale | 5.59246E-12 | 0.020 | -2.4549E-11 | -2.1973E-12 |
| | | TpCmale | 5.45085E-12 | 0.635 | -1.3494E-11 | 8.2912E-12 |
| | THCmale | VehFemale | 5.76457E-12 | 0.177 | -3.6519E-12 | 1.9387E-11 |
| | | CBDfemale | 5.76457E-12 | 0.296 | -1.7600E-11 | 5.4388E-12 |
| | | THCfemale | 5.45085E-12 | 0.642 | -1.3437E-11 | 8.3484E-12 |
| | | TpCfemale | 5.45085E-12 | 0.029 | -2.3064E-11 | -1.2791E-12 |
| | | VehMale | 5.97859E-12 | 0.091 | -1.6866E-12 | 2.2208E-11 |
| | | CBDmale | 5.59246E-12 | 0.020 | 2.1973E-12 | 2.4549E-11 |
| | | TpCmale | 5.45085E-12 | 0.053 | -1.2127E-13 | 2.1664E-11 |
| | TpCmale | VehFemale | 5.62730E-12 | 0.608 | -1.4149E-11 | 8.3415E-12 |
| | | CBDfemale | 5.62730E-12 | 0.004 | -2.8097E-11 | -5.6069E-12 |
| | | THCfemale | 5.30547E-12 | 0.015 | -2.3918E-11 | -2.7135E-12 |
| | | TpCfemale | 5.30547E-12 | 0.000 | -3.3545E-11 | -1.2341E-11 |
| | | VehMale | 5.84634E-12 | 0.931 | -1.2194E-11 | 1.1172E-11 |
| | | CBDmale | 5.45085E-12 | 0.635 | -8.2912E-12 | 1.3494E-11 |
| | | THCmale | 5.45085E-12 | 0.053 | -2.1664E-11 | 1.2127E-13 |

Supplemental Table 7.1 Descriptive statistics for *N*-acyl tyrosines (mol/L) in plasma collected from male and female adult mice 2h after receiving an i.p. injection of 10mg/kg CBD, THC, or THC+CBD

| | | N | Mean | Std. Deviation | Std. Error | 95% Confidence Interval for Mean | | Minimum | Maximum |
|---|---|---|---|---|---|---|---|---|---|
| | | | | | | Lower Bound | Upper Bound | | |
| *N*-palmitoyl tyrosine | Veh F | 7 | 3.1235E-09 | 1.92576E-09 | 7.27867E-10 | 1.3425E-09 | 4.9045E-09 | 1.43E-09 | 5.91E-09 |
| | CBD F | 8 | 1.8132E-09 | 9.28899E-10 | 3.28415E-10 | 1.0366E-09 | 2.5898E-09 | 7.97E-10 | 3.58E-09 |
| | THC F | 10 | 1.7192E-09 | 1.02320E-09 | 3.23564E-10 | 9.8720E-10 | 2.4511E-09 | 5.85E-10 | 3.73E-09 |
| | THC+CBD F | 10 | 1.9183E-09 | 9.66519E-10 | 3.05640E-10 | 1.2269E-09 | 2.6097E-09 | 5.66E-10 | 3.26E-09 |
| | Veh M | 8 | 2.4883E-09 | 1.68731E-09 | 5.96553E-10 | 1.0777E-09 | 3.8990E-09 | 3.12E-10 | 4.70E-09 |
| | CBD M | 10 | 1.6967E-09 | 1.07146E-09 | 3.38825E-10 | 9.3020E-10 | 2.4632E-09 | 5.36E-10 | 3.44E-09 |
| | THC M | 9 | 3.0184E-09 | 1.82466E-09 | 6.08220E-10 | 1.6159E-09 | 4.4210E-09 | 8.95E-10 | 6.42E-09 |
| | THC+CBD M | 10 | 3.6314E-09 | 2.98714E-09 | 9.44617E-10 | 1.4945E-09 | 5.7683E-09 | 5.34E-10 | 8.38E-09 |
| *N*-stearoyl tyrosine | Veh F | 6 | 9.9851E-10 | 7.30365E-10 | 2.98170E-10 | 2.3204E-10 | 1.7650E-09 | 4.05E-10 | 2.29E-09 |
| | CBD F | 8 | 5.4023E-10 | 2.74716E-10 | 9.71268E-11 | 3.1056E-10 | 7.6990E-10 | 1.35E-10 | 9.80E-10 |
| | THC F | 7 | 7.8143E-10 | 2.84667E-10 | 1.07594E-10 | 5.1815E-10 | 1.0447E-09 | 4.78E-10 | 1.14E-09 |
| | THC+CBD F | 10 | 5.8059E-10 | 4.15929E-10 | 1.31528E-10 | 2.8305E-10 | 8.7812E-10 | 1.80E-10 | 1.47E-09 |
| | Veh M | 6 | 4.5913E-10 | 1.45629E-10 | 5.94529E-11 | 3.0630E-10 | 6.1196E-10 | 2.34E-10 | 6.42E-10 |
| | CBD M | 11 | 5.6368E-10 | 3.41352E-10 | 1.02921E-10 | 3.3436E-10 | 7.9301E-10 | 2.13E-10 | 1.08E-09 |
| | THC M | 6 | 6.7816E-10 | 2.08605E-10 | 8.51627E-11 | 4.5924E-10 | 8.9708E-10 | 3.98E-10 | 9.40E-10 |
| | THC+CBD M | 8 | 7.5679E-10 | 5.86577E-10 | 2.07386E-10 | 2.6639E-10 | 1.2472E-09 | 1.14E-10 | 1.80E-09 |
| *N*-oleoyl tyrosine | Veh F | 8 | 3.5666E-09 | 2.90663E-09 | 1.02765E-09 | 1.1366E-09 | 5.9966E-09 | 7.19E-10 | 8.31E-09 |
| | CBD F | 8 | 2.0241E-09 | 1.59980E-09 | 5.65615E-10 | 6.8664E-10 | 3.3616E-09 | 4.16E-10 | 5.50E-09 |
| | THC F | 9 | 1.1866E-09 | 4.49309E-10 | 1.49770E-10 | 8.4123E-10 | 1.5320E-09 | 5.43E-10 | 1.79E-09 |
| | THC+CBD F | 10 | 1.6324E-09 | 7.61794E-10 | 2.40900E-10 | 1.0875E-09 | 2.1774E-09 | 5.12E-10 | 2.81E-09 |
| | Veh M | 8 | 2.9333E-09 | 2.37229E-09 | 8.38733E-10 | 9.4998E-10 | 4.9166E-09 | 3.17E-10 | 6.33E-09 |
| | CBD M | 10 | 2.2738E-09 | 1.48494E-09 | 4.69580E-10 | 1.2116E-09 | 3.3361E-09 | 5.21E-10 | 4.34E-09 |
| | THC M | 9 | 2.6597E-09 | 1.65948E-09 | 5.53159E-10 | 1.3841E-09 | 3.9353E-09 | 7.82E-10 | 5.16E-09 |
| | THC+CBD M | 10 | 3.9561E-09 | 3.60189E-09 | 1.13902E-09 | 1.3794E-09 | 6.5327E-09 | 5.30E-10 | 1.04E-08 |
| N-linoleoyl tyrosine | Veh F | 7 | 2.1562E-09 | 1.38733E-09 | 5.24363E-10 | 8.7311E-10 | 3.4392E-09 | 9.73E-10 | 4.19E-09 |
| | CBD F | 8 | 1.2701E-09 | 7.42394E-10 | 2.62476E-10 | 6.4946E-10 | 1.8908E-09 | 3.63E-10 | 2.84E-09 |
| | THC F | 8 | 8.9080E-10 | 4.68621E-10 | 1.65683E-10 | 4.9903E-10 | 1.2826E-09 | 3.96E-10 | 1.60E-09 |
| | THC+CBD F | 11 | 1.5775E-09 | 1.12025E-09 | 3.37767E-10 | 8.2491E-10 | 2.3301E-09 | 1.60E-10 | 3.81E-09 |
| | Veh M | 8 | 1.5174E-09 | 9.84641E-10 | 3.48123E-10 | 6.9422E-10 | 2.3406E-09 | 5.54E-10 | 3.44E-09 |
| | CBD M | 10 | 1.1943E-09 | 7.32789E-10 | 2.31728E-10 | 6.7005E-10 | 1.7185E-09 | 2.54E-10 | 2.41E-09 |
| | THC M | 9 | 2.0434E-09 | 9.29684E-10 | 3.09895E-10 | 1.3288E-09 | 2.7580E-09 | 4.47E-10 | 3.32E-09 |
| | THC+CBD M | 10 | 2.2120E-09 | 1.95442E-09 | 6.18042E-10 | 8.1390E-10 | 3.6101E-09 | 3.27E-10 | 5.02E-09 |
| *N*-arachidonoyl tyrosine | Veh F | BAL | | | | | | | |
| | CBD F | BAL | | | | | | | |
| | THC F | BAL | | | | | | | |
| | THC+CBD F | BAL | | | | | | | |
| | Veh M | BAL | | | | | | | |
| | CBD M | BAL | | | | | | | |
| | THC M | BAL | | | | | | | |
| | THC+CBD M | BAL | | | | | | | |
| | Veh F | 5 | 5.8876E-10 | 3.24985E-10 | 1.45338E-10 | 1.8524E-10 | 9.9228E-10 | 2.75E-10 | 9.96E-10 |

| | | | | | | | | | |
|---|---|---|---|---|---|---|---|---|---|
| *N*-docosahexaenoyl tyrosine | CBD F | 6 | 4.6083E-10 | 1.98123E-10 | 8.08833E-11 | 2.5292E-10 | 6.6875E-10 | 2.91E-10 | 7.64E-10 |
| | THC F | BAL | | | | | | | |
| | THC+CBD F | 5 | 6.0562E-10 | 3.57435E-10 | 1.59850E-10 | 1.6181E-10 | 1.0494E-09 | 3.27E-10 | 1.21E-09 |
| | Veh M | 4 | 4.6424E-10 | 2.16058E-10 | 1.08029E-10 | 1.2044E-10 | 8.0804E-10 | 2.71E-10 | 7.00E-10 |
| | CBD M | BAL | | | | | | | |
| | THC M | BAL | | | | | | | |
| | THC+CBD M | BAL | | | | | | | |

Supplemental Table 7.2 Descriptive statistics for *N*-acyl tyrosines (mol/g) in liver collected from male and female adult mice 2h after receiving an i.p. injection of 10mg/kg CBD, THC, or THC+CBD

| | | N | Mean | Std. Deviation | Std. Error | 95% Confidence Interval for Mean | | Minimum | Maximum |
|---|---|---|---|---|---|---|---|---|---|
| | | | | | | Lower Bound | Upper Bound | | |
| *N*-palmitoyl tyrosine | Veh F | 8 | 1.0519E-10 | 4.50453E-11 | 1.59259E-11 | 6.7533E-11 | 1.4285E-10 | 5.48E-11 | 1.87E-10 |
| | CBD F | 7 | 1.0687E-10 | 5.92043E-11 | 2.23771E-11 | 5.2113E-11 | 1.6162E-10 | 3.78E-11 | 1.79E-10 |
| | THC F | 10 | 1.2083E-10 | 4.83355E-11 | 1.52850E-11 | 8.6253E-11 | 1.5541E-10 | 5.78E-11 | 2.11E-10 |
| | THC+CBD F | 9 | 1.5210E-10 | 4.68588E-11 | 1.56196E-11 | 1.1608E-10 | 1.8812E-10 | 6.85E-11 | 2.29E-10 |
| | Veh M | 7 | 6.7155E-11 | 1.72213E-11 | 6.50904E-12 | 5.1228E-11 | 8.3082E-11 | 3.67E-11 | 8.55E-11 |
| | CBD M | 9 | 4.9279E-11 | 1.22649E-11 | 4.08829E-12 | 3.9851E-11 | 5.8706E-11 | 3.35E-11 | 7.58E-11 |
| | THC M | 9 | 1.2207E-10 | 4.62387E-11 | 1.54129E-11 | 8.6533E-11 | 1.5762E-10 | 7.29E-11 | 1.98E-10 |
| | THC+CBD M | 10 | 5.8255E-11 | 1.84197E-11 | 5.82481E-12 | 4.5079E-11 | 7.1432E-11 | 3.63E-11 | 1.00E-10 |
| *N*-stearoyl tyrosine | Veh F | 8 | 6.5183E-12 | 3.53122E-12 | 1.24848E-12 | 3.5661E-12 | 9.4705E-12 | 1.53E-12 | 1.10E-11 |
| | CBD F | 8 | 1.3055E-11 | 1.25975E-11 | 4.45388E-12 | 2.5228E-12 | 2.3586E-11 | 1.96E-12 | 3.74E-11 |
| | THC F | 9 | 7.6993E-12 | 3.86813E-12 | 1.28938E-12 | 4.7260E-12 | 1.0673E-11 | 2.70E-12 | 1.51E-11 |
| | THC+CBD F | 10 | 1.2451E-11 | 5.55686E-12 | 1.75723E-12 | 8.4759E-12 | 1.6426E-11 | 5.27E-12 | 2.33E-11 |
| | Veh M | 7 | 2.6762E-12 | 6.40117E-13 | 2.41941E-13 | 2.0842E-12 | 3.2682E-12 | 1.72E-12 | 3.23E-12 |
| | CBD M | 9 | 2.9706E-12 | 1.03054E-12 | 3.43515E-13 | 2.1784E-12 | 3.7627E-12 | 1.04E-12 | 4.57E-12 |
| | THC M | 9 | 7.3961E-12 | 3.35963E-12 | 1.11988E-12 | 4.8137E-12 | 9.9786E-12 | 2.49E-12 | 1.34E-11 |
| | THC+CBD M | 9 | 4.3593E-12 | 3.27834E-12 | 1.09278E-12 | 1.8393E-12 | 6.8792E-12 | 1.31E-12 | 1.05E-11 |
| *N*-oleoyl tyrosine | Veh F | 7 | 1.1177E-10 | 3.94609E-11 | 1.49148E-11 | 7.5278E-11 | 1.4827E-10 | 5.39E-11 | 1.60E-10 |
| | CBD F | 7 | 1.1735E-10 | 5.92511E-11 | 2.23948E-11 | 6.2554E-11 | 1.7215E-10 | 3.78E-11 | 2.09E-10 |
| | THC F | 10 | 2.3288E-10 | 1.26046E-10 | 3.98591E-11 | 1.4271E-10 | 3.2304E-10 | 1.00E-10 | 4.78E-10 |
| | THC+CBD F | 9 | 2.3172E-10 | 7.55625E-11 | 2.51875E-11 | 1.7364E-10 | 2.8980E-10 | 1.32E-10 | 3.58E-10 |
| | Veh M | 7 | 9.3699E-11 | 4.67107E-11 | 1.76550E-11 | 5.0499E-11 | 1.3690E-10 | 4.64E-11 | 1.74E-10 |
| | CBD M | 10 | 7.0879E-11 | 2.53196E-11 | 8.00675E-12 | 5.2766E-11 | 8.8991E-11 | 3.05E-11 | 9.99E-11 |
| | THC M | 8 | 1.1730E-10 | 3.85399E-11 | 1.36259E-11 | 8.5084E-11 | 1.4952E-10 | 6.92E-11 | 1.74E-10 |
| | THC+CBD M | 10 | 1.0655E-10 | 3.46903E-11 | 1.09700E-11 | 8.1730E-11 | 1.3136E-10 | 4.99E-11 | 1.69E-10 |
| N-linoleoyl tyrosine | Veh F | 7 | 1.0031E-10 | 3.29829E-11 | 1.24664E-11 | 6.9807E-11 | 1.3081E-10 | 6.01E-11 | 1.59E-10 |
| | CBD F | 7 | 1.2051E-10 | 8.26399E-11 | 3.12349E-11 | 4.4082E-11 | 1.9694E-10 | 2.50E-11 | 2.06E-10 |
| | THC F | 9 | 1.2085E-10 | 4.47788E-11 | 1.49263E-11 | 8.6433E-11 | 1.5527E-10 | 4.23E-11 | 1.83E-10 |
| | THC+CBD F | 10 | 2.3908E-10 | 9.27282E-11 | 2.93232E-11 | 1.7274E-10 | 3.0541E-10 | 9.88E-11 | 4.19E-10 |
| | Veh M | 7 | 5.8376E-11 | 1.82189E-11 | 6.88609E-12 | 4.1526E-11 | 7.5226E-11 | 2.90E-11 | 7.96E-11 |
| | CBD M | 9 | 4.7797E-11 | 1.73907E-11 | 5.79689E-12 | 3.4429E-11 | 6.1164E-11 | 2.61E-11 | 8.65E-11 |
| | THC M | 9 | 1.3437E-10 | 4.63351E-11 | 1.54450E-11 | 9.8758E-11 | 1.6999E-10 | 8.47E-11 | 2.09E-10 |
| | THC+CBD M | 10 | 6.8392E-11 | 2.60118E-11 | 8.22565E-12 | 4.9784E-11 | 8.7000E-11 | 4.19E-11 | 1.24E-10 |
| *N*-arachidonoyl tyrosine | Veh F | 8 | 4.6887E-11 | 2.72221E-11 | 9.62446E-12 | 2.4129E-11 | 6.9646E-11 | 1.78E-11 | 8.85E-11 |
| | CBD F | 7 | 6.2197E-11 | 2.97740E-11 | 1.12535E-11 | 3.4660E-11 | 8.9733E-11 | 2.47E-11 | 1.02E-10 |
| | THC F | 10 | 6.3003E-11 | 2.34114E-11 | 7.40333E-12 | 4.6255E-11 | 7.9750E-11 | 3.21E-11 | 9.90E-11 |
| | THC+CBD F | 9 | 7.4745E-11 | 1.98213E-11 | 6.60709E-12 | 5.9509E-11 | 8.9981E-11 | 3.11E-11 | 1.04E-10 |
| | Veh M | 7 | 3.1026E-11 | 1.01211E-11 | 3.82542E-12 | 2.1665E-11 | 4.0386E-11 | 1.60E-11 | 4.17E-11 |
| | CBD M | 9 | 2.2099E-11 | 5.96223E-12 | 1.98741E-12 | 1.7516E-11 | 2.6682E-11 | 1.44E-11 | 3.07E-11 |
| | THC M | 9 | 5.8938E-11 | 2.07462E-11 | 6.91541E-12 | 4.2991E-11 | 7.4885E-11 | 3.43E-11 | 8.69E-11 |
| | THC+CBD M | 10 | 2.8466E-11 | 1.21966E-11 | 3.85692E-12 | 1.9741E-11 | 3.7191E-11 | 1.46E-11 | 5.15E-11 |
| | Veh F | 8 | 8.1091E-11 | 3.82709E-11 | 1.35308E-11 | 4.9096E-11 | 1.1309E-10 | 3.84E-11 | 1.31E-10 |

| | | | | | | | | | |
|---|---|---|---|---|---|---|---|---|---|
| *N*-docosahexaenoyl tyrosine | CBD F | 7 | 9.8547E-11 | 6.16883E-11 | 2.33160E-11 | 4.1495E-11 | 1.5560E-10 | 3.47E-11 | 1.74E-10 |
| | THC F | 10 | 1.2058E-10 | 5.22603E-11 | 1.65262E-11 | 8.3192E-11 | 1.5796E-10 | 4.29E-11 | 2.18E-10 |
| | THC+CBD F | 9 | 1.5721E-10 | 5.20055E-11 | 1.73352E-11 | 1.1723E-10 | 1.9718E-10 | 6.82E-11 | 2.39E-10 |
| | Veh M | 7 | 6.7354E-11 | 2.22402E-11 | 8.40600E-12 | 4.6785E-11 | 8.7922E-11 | 2.82E-11 | 9.72E-11 |
| | CBD M | 9 | 4.2635E-11 | 9.07662E-12 | 3.02554E-12 | 3.5658E-11 | 4.9612E-11 | 3.10E-11 | 5.54E-11 |
| | THC M | 9 | 1.1737E-10 | 4.82174E-11 | 1.60725E-11 | 8.0307E-11 | 1.5443E-10 | 6.19E-11 | 1.96E-10 |
| | THC+CBD M | 10 | 7.2863E-11 | 3.59697E-11 | 1.13746E-11 | 4.7132E-11 | 9.8595E-11 | 4.39E-11 | 1.54E-10 |

Supplemental Table 7.3 Independent t-test comparing male and female levels of *N*-acyl tyrosines measured in plasma collected from adult mice 2h after an i.p. injection of 10mg/kg CBD, THC, or THC+CBD

| | | | Levene's Test for Equality of Variances | | t-test for Equality of Means | | | | | | |
|---|---|---|---|---|---|---|---|---|---|---|---|
| | | | | | | | Significance | | | 95% Confidence Interval of the Difference | |
| | | | F | Sig. | t | df | One-Sided p | Two-Sided p | Std. Error Difference | Lower | Upper |
| Veh | *N*-palmitoyl tyrosine | Equal variances assumed | 0.047 | 0.833 | 0.681 | 13 | 0.254 | 0.508 | 9.32254E-10 | -1.37884E-09 | 2.64918E-09 |
| | | Equal variances not assumed | | | 0.675 | 12.092 | 0.256 | 0.512 | 9.41098E-10 | -1.41358E-09 | 2.68393E-09 |
| | *N*-stearoyl tyrosine | Equal variances assumed | 7.968 | 0.018 | 1.774 | 10 | 0.053 | 0.106 | 3.04040E-10 | -1.38065E-10 | 1.21682E-09 |
| | | Equal variances not assumed | | | 1.774 | 5.397 | 0.066 | 0.132 | 3.04040E-10 | -2.25203E-10 | 1.30396E-09 |
| | *N*-oleoyl tyrosine | Equal variances assumed | 0.535 | 0.477 | 0.477 | 14 | 0.320 | 0.640 | 1.32648E-09 | -2.21171E-09 | 3.47831E-09 |
| | | Equal variances not assumed | | | 0.477 | 13.460 | 0.320 | 0.641 | 1.32648E-09 | -2.22246E-09 | 3.48906E-09 |
| | *N*-linoleoyl tyrosine | Equal variances assumed | 3.746 | 0.075 | 1.039 | 13 | 0.159 | 0.318 | 6.14636E-10 | -6.89067E-10 | 1.96662E-09 |
| | | Equal variances not assumed | | | 1.015 | 10.677 | 0.166 | 0.333 | 6.29402E-10 | -7.51662E-10 | 2.02921E-09 |
| | *N*-docosahexaenoyl tyrosine | Equal variances assumed | 3.636 | 0.098 | 0.655 | 7 | 0.267 | 0.534 | 1.90160E-10 | -3.25137E-10 | 5.74179E-10 |
| | | Equal variances not assumed | | | 0.688 | 6.852 | 0.257 | 0.514 | 1.81089E-10 | -3.05567E-10 | 5.54609E-10 |
| CBD | *N*-palmitoyl tyrosine | Equal variances assumed | 0.896 | 0.358 | 0.243 | 16 | 0.406 | 0.811 | 4.79827E-10 | -9.00648E-10 | 1.13373E-09 |
| | | Equal variances not assumed | | | 0.247 | 15.858 | 0.404 | 0.808 | 4.71867E-10 | -8.84503E-10 | 1.11758E-09 |
| | *N*-stearoyl tyrosine | Equal variances assumed | 1.284 | 0.273 | -0.160 | 17 | 0.437 | 0.875 | 1.46657E-10 | -3.32872E-10 | 2.85966E-10 |
| | | Equal variances not assumed | | | -0.166 | 16.757 | 0.435 | 0.870 | 1.41515E-10 | -3.22353E-10 | 2.75448E-10 |

| | | | | | | | | | | | |
|---|---|---|---|---|---|---|---|---|---|---|---|
| | *N*-oleoyl tyrosine | Equal variances assumed | 0.414 | 0.529 | -0.343 | 16 | 0.368 | 0.736 | 7.28707E-10 | -1.79450E-09 | 1.29508E-09 |
| | | Equal variances not assumed | | | -0.340 | 14.586 | 0.369 | 0.739 | 7.35137E-10 | -1.82051E-09 | 1.32108E-09 |
| | *N*-linoleoyl tyrosine | Equal variances assumed | 0.093 | 0.764 | 0.217 | 16 | 0.415 | 0.831 | 3.49593E-10 | -6.65242E-10 | 8.16966E-10 |
| | | Equal variances not assumed | | | 0.217 | 15.052 | 0.416 | 0.831 | 3.50131E-10 | -6.70198E-10 | 8.21922E-10 |
| THC | *N*-palmitoyl tyrosine | Equal variances assumed | 5.528 | 0.031 | -1.942 | 17 | 0.034 | 0.069 | 6.69159E-10 | -2.71109E-09 | 1.12511E-10 |
| | | Equal variances not assumed | | | -1.886 | 12.294 | 0.042 | 0.083 | 6.88931E-10 | -2.79638E-09 | 1.97794E-10 |
| | *N*-stearoyl tyrosine | Equal variances assumed | 3.071 | 0.107 | 0.734 | 11 | 0.239 | 0.478 | 1.40725E-10 | -2.06469E-10 | 4.13000E-10 |
| | | Equal variances not assumed | | | 0.753 | 10.791 | 0.234 | 0.468 | 1.37219E-10 | -1.99468E-10 | 4.06000E-10 |
| | *N*-oleoyl tyrosine | Equal variances assumed | 40.101 | 0.000 | -2.571 | 16 | 0.010 | 0.021 | 5.73076E-10 | -2.68798E-09 | -2.58244E-10 |
| | | Equal variances not assumed | | | -2.571 | 9.167 | 0.015 | 0.030 | 5.73076E-10 | -2.76592E-09 | -1.80307E-10 |
| | *N*-linoleoyl tyrosine | Equal variances assumed | 1.965 | 0.181 | -3.160 | 15 | 0.003 | 0.006 | 3.64742E-10 | -1.93005E-09 | -3.75190E-10 |
| | | Equal variances not assumed | | | -3.280 | 12.097 | 0.003 | 0.007 | 3.51405E-10 | -1.91758E-09 | -3.87657E-10 |
| THC+CBD | *N*-palmitoyl tyrosine | Equal variances assumed | 32.470 | 0.000 | -1.726 | 18 | 0.051 | 0.102 | 9.92832E-10 | -3.79902E-09 | 3.72710E-10 |
| | | Equal variances not assumed | | | -1.726 | 10.864 | 0.056 | 0.113 | 9.92832E-10 | -3.90170E-09 | 4.75398E-10 |
| | *N*-stearoyl tyrosine | Equal variances assumed | 1.061 | 0.318 | -0.746 | 16 | 0.233 | 0.466 | 2.36145E-10 | -6.76803E-10 | 3.24408E-10 |
| | | Equal variances not assumed | | | -0.717 | 12.225 | 0.243 | 0.487 | 2.45578E-10 | -7.10175E-10 | 3.57780E-10 |
| | *N*-oleoyl tyrosine | Equal variances assumed | 23.683 | 0.000 | -1.996 | 18 | 0.031 | 0.061 | 1.16421E-09 | -4.76955E-09 | 1.22294E-10 |
| | | Equal variances not assumed | | | -1.996 | 9.804 | 0.037 | 0.074 | 1.16421E-09 | -4.92472E-09 | 2.77464E-10 |

| | | | | | | | | | | |
|---|---|---|---|---|---|---|---|---|---|---|
| *N*-linoleoyl tyrosine | Equal variances assumed | 10.703 | 0.004 | -0.924 | 19 | 0.184 | 0.367 | 6.86672E-10 | -2.07173E-09 | 8.02715E-10 |
| | Equal variances not assumed | | | -0.901 | 14.051 | 0.191 | 0.383 | 7.04317E-10 | -2.14460E-09 | 8.75590E-10 |

Supplemental Table 7.4 Independent t-test comparing male and female levels of *N*-acyl tyrosines measured in liver collected from adult mice 2h after an i.p. injection of 10mg/kg CBD, THC, or THC+CBD

| | | | Levene's Test for Equality of Variances | | t-test for Equality of Means | | | | | | | |
|---|---|---|---|---|---|---|---|---|---|---|---|---|
| | | | | | | | Significance | | | | 95% Confidence Interval of the Difference | |
| | | | F | Sig. | t | df | One-Sided p | Two-Sided p | Mean Difference | Std. Error Difference | Lower | Upper |
| Veh | *N*-palmitoyl tyrosine | Equal variances assumed | 5.373 | 0.037 | 2.096 | 13 | 0.028 | 0.056 | 3.80366E-11 | 1.81472E-11 | -1.16797E-12 | 7.72411E-11 |
| | | Equal variances not assumed | | | 2.211 | 9.233 | 0.027 | 0.054 | 3.80366E-11 | 1.72047E-11 | -7.33769E-13 | 7.68069E-11 |
| | *N*-stearoyl tyrosine | Equal variances assumed | 19.723 | 0.001 | 2.825 | 13 | 0.007 | 0.014 | 3.84212E-12 | 1.35983E-12 | 9.04382E-13 | 6.77986E-12 |
| | | Equal variances not assumed | | | 3.021 | 7.523 | 0.009 | 0.018 | 3.84212E-12 | 1.27170E-12 | 8.76907E-13 | 6.80734E-12 |
| | *N*-oleoyl tyrosine | Equal variances assumed | 0.151 | 0.704 | 0.782 | 12 | 0.225 | 0.449 | 1.80748E-11 | 2.31117E-11 | -3.22813E-11 | 6.84308E-11 |
| | | Equal variances not assumed | | | 0.782 | 11.674 | 0.225 | 0.450 | 1.80748E-11 | 2.31117E-11 | -3.24376E-11 | 6.85872E-11 |
| | *N*-linoleoyl tyrosine | Equal variances assumed | 1.175 | 0.300 | 2.944 | 12 | 0.006 | 0.012 | 4.19345E-11 | 1.42418E-11 | 1.09043E-11 | 7.29646E-11 |
| | | Equal variances not assumed | | | 2.944 | 9.350 | 0.008 | 0.016 | 4.19345E-11 | 1.42418E-11 | 9.90001E-12 | 7.39689E-11 |
| | *N*-arachidonoyl tyrosine | Equal variances assumed | 6.778 | 0.022 | 1.451 | 13 | 0.085 | 0.171 | 1.58618E-11 | 1.09337E-11 | -7.75891E-12 | 3.94826E-11 |
| | | Equal variances not assumed | | | 1.532 | 9.121 | 0.080 | 0.160 | 1.58618E-11 | 1.03568E-11 | -7.51973E-12 | 3.92434E-11 |
| | *N*-docosahexaenoyl tyrosine | Equal variances assumed | 4.323 | 0.058 | 0.832 | 13 | 0.210 | 0.420 | 1.37372E-11 | 1.65045E-11 | -2.19185E-11 | 4.93930E-11 |
| | | Equal variances not assumed | | | 0.862 | 11.455 | 0.203 | 0.406 | 1.37372E-11 | 1.59293E-11 | -2.11537E-11 | 4.86282E-11 |
| CBD | *N*-palmitoyl tyrosine | Equal variances assumed | 20.903 | 0.000 | 2.868 | 14 | 0.006 | 0.012 | 5.75895E-11 | 2.00834E-11 | 1.45148E-11 | 1.00664E-10 |
| | | Equal variances not assumed | | | 2.532 | 6.402 | 0.021 | 0.042 | 5.75895E-11 | 2.27475E-11 | 2.76441E-12 | 1.12415E-10 |
| | *N*-stearoyl tyrosine | Equal variances assumed | 12.735 | 0.003 | 2.402 | 15 | 0.015 | 0.030 | 1.00840E-11 | 4.19758E-12 | 1.13702E-12 | 1.90309E-11 |

| | | | | | | | | | | | | |
|---|---|---|---|---|---|---|---|---|---|---|---|---|
| | | Equal variances not assumed | | | 2.257 | 7.083 | 0.029 | 0.058 | 1.00840E-11 | 4.46711E-12 | -4.53938E-13 | 2.06218E-11 |
| | *N*-oleoyl tyrosine | Equal variances assumed | 5.369 | 0.035 | 2.230 | 15 | 0.021 | 0.041 | 4.64740E-11 | 2.08435E-11 | 2.04701E-12 | 9.09010E-11 |
| | | Equal variances not assumed | | | 1.954 | 7.550 | 0.044 | 0.089 | 4.64740E-11 | 2.37831E-11 | -8.94457E-12 | 1.01893E-10 |
| | *N*-linoleoyl tyrosine | Equal variances assumed | 54.791 | 0.000 | 2.592 | 14 | 0.011 | 0.021 | 7.27139E-11 | 2.80575E-11 | 1.25366E-11 | 1.32891E-10 |
| | | Equal variances not assumed | | | 2.289 | 6.415 | 0.030 | 0.059 | 7.27139E-11 | 3.17683E-11 | -3.81821E-12 | 1.49246E-10 |
| | *N*-arachidonoyl tyrosine | Equal variances assumed | 18.665 | 0.001 | 3.977 | 14 | 0.001 | 0.001 | 4.00975E-11 | 1.00821E-11 | 1.84736E-11 | 6.17213E-11 |
| | | Equal variances not assumed | | | 3.509 | 6.375 | 0.006 | 0.011 | 4.00975E-11 | 1.14277E-11 | 1.25294E-11 | 6.76656E-11 |
| | *N*-docosahexaenoyl tyrosine | Equal variances assumed | 40.189 | 0.000 | 2.708 | 14 | 0.008 | 0.017 | 5.59121E-11 | 2.06435E-11 | 1.16363E-11 | 1.00188E-10 |
| | | Equal variances not assumed | | | 2.378 | 6.202 | 0.027 | 0.054 | 5.59121E-11 | 2.35115E-11 | -1.16619E-12 | 1.12990E-10 |
| | *N*-palmitoyl tyrosine | Equal variances assumed | 0.054 | 0.819 | -0.057 | 17 | 0.478 | 0.955 | -1.24442E-12 | 2.17606E-11 | -4.71552E-11 | 4.46664E-11 |
| | | Equal variances not assumed | | | -0.057 | 16.923 | 0.477 | 0.955 | -1.24442E-12 | 2.17069E-11 | -4.70578E-11 | 4.45689E-11 |
| | *N*-stearoyl tyrosine | Equal variances assumed | 0.210 | 0.653 | 0.177 | 16 | 0.431 | 0.861 | 3.03132E-13 | 1.70781E-12 | -3.31726E-12 | 3.92353E-12 |
| | | Equal variances not assumed | | | 0.177 | 15.692 | 0.431 | 0.861 | 3.03132E-13 | 1.70781E-12 | -3.32304E-12 | 3.92931E-12 |
| THC | *N*-oleoyl tyrosine | Equal variances assumed | 8.735 | 0.009 | 2.488 | 16 | 0.012 | 0.024 | 1.15572E-10 | 4.64432E-11 | 1.71169E-11 | 2.14027E-10 |
| | | Equal variances not assumed | | | 2.744 | 11.033 | 0.010 | 0.019 | 1.15572E-10 | 4.21238E-11 | 2.28917E-11 | 2.08252E-10 |
| | *N*-linoleoyl tyrosine | Equal variances assumed | 0.144 | 0.709 | -0.630 | 16 | 0.269 | 0.538 | -1.35214E-11 | 2.14789E-11 | -5.90546E-11 | 3.20118E-11 |
| | | Equal variances not assumed | | | -0.630 | 15.981 | 0.269 | 0.538 | -1.35214E-11 | 2.14789E-11 | -5.90589E-11 | 3.20162E-11 |
| | *N*-arachidonoyl tyrosine | Equal variances assumed | 0.078 | 0.784 | 0.399 | 17 | 0.348 | 0.695 | 4.06453E-12 | 1.01989E-11 | -1.74532E-11 | 2.55822E-11 |
| | | Equal variances | | | 0.401 | 16.999 | 0.347 | 0.693 | 4.06453E-12 | 1.01308E-11 | -1.73096E-11 | 2.54387E-11 |

| | | | | | | | | | | | | |
|---|---|---|---|---|---|---|---|---|---|---|---|---|
| | | not assumed | | | | | | | | | | |
| | *N*-docosahexaenoyl tyrosine | Equal variances assumed | 0.030 | 0.865 | 0.138 | 17 | 0.446 | 0.892 | 3.20594E-12 | 2.31564E-11 | -4.56497E-11 | 5.20616E-11 |
| | | Equal variances not assumed | | | 0.139 | 16.984 | 0.446 | 0.891 | 3.20594E-12 | 2.30529E-11 | -4.54351E-11 | 5.18469E-11 |
| | *N*-palmitoyl tyrosine | Equal variances assumed | 3.563 | 0.076 | 5.865 | 17 | 0.000 | 0.000 | 9.38434E-11 | 1.60019E-11 | 6.00824E-11 | 1.27604E-10 |
| | | Equal variances not assumed | | | 5.629 | 10.204 | 0.000 | 0.000 | 9.38434E-11 | 1.66703E-11 | 5.68001E-11 | 1.30887E-10 |
| | *N*-stearoyl tyrosine | Equal variances assumed | 2.294 | 0.148 | 3.807 | 17 | 0.001 | 0.001 | 8.09176E-12 | 2.12577E-12 | 3.60679E-12 | 1.25767E-11 |
| | | Equal variances not assumed | | | 3.910 | 14.815 | 0.001 | 0.001 | 8.09176E-12 | 2.06931E-12 | 3.67632E-12 | 1.25072E-11 |
| | *N*-oleoyl tyrosine | Equal variances assumed | 2.889 | 0.107 | 4.725 | 17 | 0.000 | 0.000 | 1.25172E-10 | 2.64903E-11 | 6.92819E-11 | 1.81061E-10 |
| | | Equal variances not assumed | | | 4.556 | 10.972 | 0.000 | 0.001 | 1.25172E-10 | 2.74727E-11 | 6.46857E-11 | 1.85657E-10 |
| THC+CBD | *N*-linoleoyl tyrosine | Equal variances assumed | 8.531 | 0.009 | 5.604 | 18 | 0.000 | 0.000 | 1.70685E-10 | 3.04551E-11 | 1.06701E-10 | 2.34669E-10 |
| | | Equal variances not assumed | | | 5.604 | 10.408 | 0.000 | 0.000 | 1.70685E-10 | 3.04551E-11 | 1.03185E-10 | 2.38185E-10 |
| | *N*-arachidonoyl tyrosine | Equal variances assumed | 0.268 | 0.612 | 6.203 | 17 | 0.000 | 0.000 | 4.62795E-11 | 7.46039E-12 | 3.05395E-11 | 6.20196E-11 |
| | | Equal variances not assumed | | | 6.049 | 13.036 | 0.000 | 0.000 | 4.62795E-11 | 7.65046E-12 | 2.97563E-11 | 6.28027E-11 |
| | *N*-docosahexaenoyl tyrosine | Equal variances assumed | 0.894 | 0.358 | 4.149 | 17 | 0.000 | 0.001 | 8.43425E-11 | 2.03296E-11 | 4.14508E-11 | 1.27234E-10 |
| | | Equal variances not assumed | | | 4.068 | 14.056 | 0.001 | 0.001 | 8.43425E-11 | 2.07338E-11 | 3.98895E-11 | 1.28796E-10 |

Supplemental Table 7.5 One-way ANOVA for *N*-acyl tyrosines measured in plasma collected from adult mice 2h after an i.p. injection of 10mg/kg CBD, THC, or THC+CBD

| | | Sum of Squares | df | Mean Square | F | Sig. |
|---|---|---|---|---|---|---|
| *N*-palmitoyl tyrosine | Between Groups | 3.70E-17 | 7 | 5.28E-18 | 1.845 | 0.0938589 |
| | Within Groups | 1.83E-16 | 64 | 2.86E-18 | | |
| | Total | 2.20E-16 | 71 | | | |
| *N*-stearoyl tyrosine | Between Groups | 1.39E-18 | 7 | 1.98E-19 | 1.173 | 0.3338795 |
| | Within Groups | 9.14E-18 | 54 | 1.69E-19 | | |
| | Total | 1.05E-17 | 61 | | | |
| *N*-oleoyl tyrosine | Between Groups | 5.74E-17 | 7 | 8.20E-18 | 1.861 | 0.0908976 |
| | Within Groups | 2.82E-16 | 64 | 4.41E-18 | | |
| | Total | 3.39E-16 | 71 | | | |
| *N*-linoleoyl tyrosine | Between Groups | 1.43E-17 | 7 | 2.04E-18 | 1.559 | 0.1642926 |
| | Within Groups | 8.24E-17 | 63 | 1.31E-18 | | |
| | Total | 9.67E-17 | 70 | | | |
| *N*-docosahexaenoyl tyrosine | Between Groups | 2.10E-19 | 7 | 3.00E-20 | 0.442 | 0.8660404 |
| | Within Groups | 1.63E-18 | 24 | 6.80E-20 | | |
| | Total | 1.84E-18 | 31 | | | |

Supplemental Table 7.6 One-way ANOVA for *N*-acyl tyrosines measured in liver collected from adult mice 2h after an i.p. injection of 10mg/kg CBD, THC, or THC+CBD

| | | Sum of Squares | df | Mean Square | F | Sig. |
|---|---|---|---|---|---|---|
| *N*-palmitoyl tyrosine | Between Groups | 8.16E-20 | 7 | 1.17E-20 | 7.329 | 0.00000 |
| | Within Groups | 9.70E-20 | 61 | 1.59E-21 | | |
| | Total | 1.79E-19 | 68 | | | |
| *N*-stearoyl tyrosine | Between Groups | 9.33E-22 | 7 | 1.33E-22 | 4.559 | 0.00038 |
| | Within Groups | 1.78E-21 | 61 | 2.92E-23 | | |
| | Total | 2.72E-21 | 68 | | | |
| *N*-oleoyl tyrosine | Between Groups | 2.49E-19 | 7 | 3.56E-20 | 8.234 | 0.00000 |
| | Within Groups | 2.59E-19 | 60 | 4.32E-21 | | |
| | Total | 5.08E-19 | 67 | | | |
| *N*-linoleoyl tyrosine | Between Groups | 2.44E-19 | 7 | 3.49E-20 | 12.410 | 0.00000 |
| | Within Groups | 1.69E-19 | 60 | 2.81E-21 | | |
| | Total | 4.13E-19 | 67 | | | |
| *N*-arachidonoyl tyrosine | Between Groups | 2.30E-20 | 7 | 3.29E-21 | 8.275 | 0.00000 |
| | Within Groups | 2.43E-20 | 61 | 3.98E-22 | | |
| | Total | 4.73E-20 | 68 | | | |
| *N*-docosahexaenoyl tyrosine | Between Groups | 8.24E-20 | 7 | 1.18E-20 | 6.345 | 0.00001 |
| | Within Groups | 1.13E-19 | 61 | 1.86E-21 | | |
| | Total | 1.96E-19 | 68 | | | |

#### Supplemental Table 7.7 Fisher's LSD post-hoc test for plasma *N*-acyl tyrosine ANOVA

| Dependent Variable | | | Std. Error | Sig. | 95% Confidence Interval | |
|---|---|---|---|---|---|---|
| | | | | | Lower Bound | Upper Bound |
| *N*-palmitoyl tyrosine | VehFemale | CBDfemale | 8.76E-10 | 0.1396 | -4.40E-10 | 3.06E-09 |
| | | THCfemale | 8.34E-10 | 0.0971 | -2.62E-10 | 3.07E-09 |
| | | TpCfemale | 8.34E-10 | 0.1533 | -4.61E-10 | 2.87E-09 |
| | | VehMale | 8.76E-10 | 0.4710 | -1.11E-09 | 2.39E-09 |
| | | CBDmale | 8.34E-10 | 0.0920 | -2.39E-10 | 3.09E-09 |
| | | THCmale | 8.53E-10 | 0.9023 | -1.60E-09 | 1.81E-09 |
| | | TpCmale | 8.34E-10 | 0.5447 | -2.17E-09 | 1.16E-09 |
| | CBDfemale | VehFemale | 8.76E-10 | 0.1396 | -3.06E-09 | 4.40E-10 |
| | | THCfemale | 8.03E-10 | 0.9071 | -1.51E-09 | 1.70E-09 |
| | | TpCfemale | 8.03E-10 | 0.8963 | -1.71E-09 | 1.50E-09 |
| | | VehMale | 8.46E-10 | 0.4279 | -2.37E-09 | 1.02E-09 |
| | | CBDmale | 8.03E-10 | 0.8850 | -1.49E-09 | 1.72E-09 |
| | | THCmale | 8.22E-10 | 0.1477 | -2.85E-09 | 4.38E-10 |
| | | TpCmale | 8.03E-10 | 0.0269 | -3.42E-09 | -2.14E-10 |
| | THCfemale | VehFemale | 8.34E-10 | 0.0971 | -3.07E-09 | 2.62E-10 |
| | | CBDfemale | 8.03E-10 | 0.9071 | -1.70E-09 | 1.51E-09 |
| | | TpCfemale | 7.57E-10 | 0.7934 | -1.71E-09 | 1.31E-09 |
| | | VehMale | 8.03E-10 | 0.3416 | -2.37E-09 | 8.35E-10 |
| | | CBDmale | 7.57E-10 | 0.9764 | -1.49E-09 | 1.53E-09 |
| | | THCmale | 7.78E-10 | 0.0996 | -2.85E-09 | 2.54E-10 |
| | | TpCmale | 7.57E-10 | 0.0140 | -3.42E-09 | -4.00E-10 |
| | TpCfemale | VehFemale | 8.34E-10 | 0.1533 | -2.87E-09 | 4.61E-10 |
| | | CBDfemale | 8.03E-10 | 0.8963 | -1.50E-09 | 1.71E-09 |
| | | THCfemale | 7.57E-10 | 0.7934 | -1.31E-09 | 1.71E-09 |
| | | VehMale | 8.03E-10 | 0.4802 | -2.17E-09 | 1.03E-09 |
| | | CBDmale | 7.57E-10 | 0.7707 | -1.29E-09 | 1.73E-09 |
| | | THCmale | 7.78E-10 | 0.1620 | -2.65E-09 | 4.53E-10 |
| | | TpCmale | 7.57E-10 | 0.0270 | -3.23E-09 | -2.01E-10 |
| | VehMale | VehFemale | 8.76E-10 | 0.4710 | -2.39E-09 | 1.11E-09 |
| | | CBDfemale | 8.46E-10 | 0.4279 | -1.02E-09 | 2.37E-09 |
| | | THCfemale | 8.03E-10 | 0.3416 | -8.35E-10 | 2.37E-09 |
| | | TpCfemale | 8.03E-10 | 0.4802 | -1.03E-09 | 2.17E-09 |
| | | CBDmale | 8.03E-10 | 0.3278 | -8.12E-10 | 2.40E-09 |
| | | THCmale | 8.22E-10 | 0.5215 | -2.17E-09 | 1.11E-09 |
| | | TpCmale | 8.03E-10 | 0.1593 | -2.75E-09 | 4.61E-10 |
| | CBDmale | VehFemale | 8.34E-10 | 0.0920 | -3.09E-09 | 2.39E-10 |
| | | CBDfemale | 8.03E-10 | 0.8850 | -1.72E-09 | 1.49E-09 |
| | | THCfemale | 7.57E-10 | 0.9764 | -1.53E-09 | 1.49E-09 |
| | | TpCfemale | 7.57E-10 | 0.7707 | -1.73E-09 | 1.29E-09 |
| | | VehMale | 8.03E-10 | 0.3278 | -2.40E-09 | 8.12E-10 |
| | | THCmale | 7.78E-10 | 0.0940 | -2.88E-09 | 2.32E-10 |
| | | TpCmale | 7.57E-10 | 0.0130 | -3.45E-09 | -4.23E-10 |
| | THCmale | VehFemale | 8.53E-10 | 0.9023 | -1.81E-09 | 1.60E-09 |
| | | CBDfemale | 8.22E-10 | 0.1477 | -4.38E-10 | 2.85E-09 |
| | | THCfemale | 7.78E-10 | 0.0996 | -2.54E-10 | 2.85E-09 |
| | | TpCfemale | 7.78E-10 | 0.1620 | -4.53E-10 | 2.65E-09 |
| | | VehMale | 8.22E-10 | 0.5215 | -1.11E-09 | 2.17E-09 |
| | | CBDmale | 7.78E-10 | 0.0940 | -2.32E-10 | 2.88E-09 |
| | | TpCmale | 7.78E-10 | 0.4335 | -2.17E-09 | 9.41E-10 |
| | TpCmale | VehFemale | 8.34E-10 | 0.5447 | -1.16E-09 | 2.17E-09 |
| | | CBDfemale | 8.03E-10 | 0.0269 | 2.14E-10 | 3.42E-09 |
| | | THCfemale | 7.57E-10 | 0.0140 | 4.00E-10 | 3.42E-09 |
| | | TpCfemale | 7.57E-10 | 0.0270 | 2.01E-10 | 3.23E-09 |
| | | VehMale | 8.03E-10 | 0.1593 | -4.61E-10 | 2.75E-09 |
| | | CBDmale | 7.57E-10 | 0.0130 | 4.23E-10 | 3.45E-09 |
| | | THCmale | 7.78E-10 | 0.4335 | -9.41E-10 | 2.17E-09 |
| *N*-stearoyl tyrosine | VehFemale | CBDfemale | 2.22E-10 | 0.0439 | 1.29E-11 | 9.04E-10 |

| | | | | | | |
|---|---|---|---|---|---|---|
| | | THCfemale | 2.29E-10 | 0.3470 | -2.42E-10 | 6.76E-10 |
| | | TpCfemale | 2.12E-10 | 0.0543 | -7.92E-12 | 8.44E-10 |
| | | VehMale | 2.37E-10 | 0.0271 | 6.33E-11 | 1.02E-09 |
| | | CBDmale | 2.09E-10 | 0.0420 | 1.63E-11 | 8.53E-10 |
| | | THCmale | 2.37E-10 | 0.1830 | -1.56E-10 | 7.96E-10 |
| | | TpCmale | 2.22E-10 | 0.2814 | -2.04E-10 | 6.87E-10 |
| | CBDfemale | VehFemale | 2.22E-10 | 0.0439 | -9.04E-10 | -1.29E-11 |
| | | THCfemale | 2.13E-10 | 0.2622 | -6.68E-10 | 1.86E-10 |
| | | TpCfemale | 1.95E-10 | 0.8369 | -4.32E-10 | 3.51E-10 |
| | | VehMale | 2.22E-10 | 0.7165 | -3.64E-10 | 5.26E-10 |
| | | CBDmale | 1.91E-10 | 0.9028 | -4.07E-10 | 3.60E-10 |
| | | THCmale | 2.22E-10 | 0.5373 | -5.83E-10 | 3.07E-10 |
| | | TpCmale | 2.06E-10 | 0.2970 | -6.29E-10 | 1.96E-10 |
| | THCfemale | VehFemale | 2.29E-10 | 0.3470 | -6.76E-10 | 2.42E-10 |
| | | CBDfemale | 2.13E-10 | 0.2622 | -1.86E-10 | 6.68E-10 |
| | | TpCfemale | 2.03E-10 | 0.3262 | -2.06E-10 | 6.07E-10 |
| | | VehMale | 2.29E-10 | 0.1647 | -1.36E-10 | 7.81E-10 |
| | | CBDmale | 1.99E-10 | 0.2784 | -1.81E-10 | 6.16E-10 |
| | | THCmale | 2.29E-10 | 0.6536 | -3.56E-10 | 5.62E-10 |
| | | TpCmale | 2.13E-10 | 0.9083 | -4.02E-10 | 4.51E-10 |
| | TpCfemale | VehFemale | 2.12E-10 | 0.0543 | -8.44E-10 | 7.92E-12 |
| | | CBDfemale | 1.95E-10 | 0.8369 | -3.51E-10 | 4.32E-10 |
| | | THCfemale | 2.03E-10 | 0.3262 | -6.07E-10 | 2.06E-10 |
| | | VehMale | 2.12E-10 | 0.5698 | -3.04E-10 | 5.47E-10 |
| | | CBDmale | 1.80E-10 | 0.9254 | -3.43E-10 | 3.77E-10 |
| | | THCmale | 2.12E-10 | 0.6478 | -5.23E-10 | 3.28E-10 |
| | | TpCmale | 1.95E-10 | 0.3705 | -5.67E-10 | 2.15E-10 |
| | VehMale | VehFemale | 2.37E-10 | 0.0271 | -1.02E-09 | -6.33E-11 |
| | | CBDfemale | 2.22E-10 | 0.7165 | -5.26E-10 | 3.64E-10 |
| | | THCfemale | 2.29E-10 | 0.1647 | -7.81E-10 | 1.36E-10 |
| | | TpCfemale | 2.12E-10 | 0.5698 | -5.47E-10 | 3.04E-10 |
| | | CBDmale | 2.09E-10 | 0.6185 | -5.23E-10 | 3.14E-10 |
| | | THCmale | 2.37E-10 | 0.3605 | -6.95E-10 | 2.57E-10 |
| | | TpCmale | 2.22E-10 | 0.1859 | -7.43E-10 | 1.48E-10 |
| | CBDmale | VehFemale | 2.09E-10 | 0.0420 | -8.53E-10 | -1.63E-11 |
| | | CBDfemale | 1.91E-10 | 0.9028 | -3.60E-10 | 4.07E-10 |
| | | THCfemale | 1.99E-10 | 0.2784 | -6.16E-10 | 1.81E-10 |
| | | TpCfemale | 1.80E-10 | 0.9254 | -3.77E-10 | 3.43E-10 |
| | | VehMale | 2.09E-10 | 0.6185 | -3.14E-10 | 5.23E-10 |
| | | THCmale | 2.09E-10 | 0.5857 | -5.33E-10 | 3.04E-10 |
| | | TpCmale | 1.91E-10 | 0.3168 | -5.76E-10 | 1.90E-10 |
| | THCmale | VehFemale | 2.37E-10 | 0.1830 | -7.96E-10 | 1.56E-10 |
| | | CBDfemale | 2.22E-10 | 0.5373 | -3.07E-10 | 5.83E-10 |
| | | THCfemale | 2.29E-10 | 0.6536 | -5.62E-10 | 3.56E-10 |
| | | TpCfemale | 2.12E-10 | 0.6478 | -3.28E-10 | 5.23E-10 |
| | | VehMale | 2.37E-10 | 0.3605 | -2.57E-10 | 6.95E-10 |
| | | CBDmale | 2.09E-10 | 0.5857 | -3.04E-10 | 5.33E-10 |
| | | TpCmale | 2.22E-10 | 0.7248 | -5.24E-10 | 3.67E-10 |
| | TpCmale | VehFemale | 2.22E-10 | 0.2814 | -6.87E-10 | 2.04E-10 |
| | | CBDfemale | 2.06E-10 | 0.2970 | -1.96E-10 | 6.29E-10 |
| | | THCfemale | 2.13E-10 | 0.9083 | -4.51E-10 | 4.02E-10 |
| | | TpCfemale | 1.95E-10 | 0.3705 | -2.15E-10 | 5.67E-10 |
| | | VehMale | 2.22E-10 | 0.1859 | -1.48E-10 | 7.43E-10 |
| | | CBDmale | 1.91E-10 | 0.3168 | -1.90E-10 | 5.76E-10 |
| | | THCmale | 2.22E-10 | 0.7248 | -3.67E-10 | 5.24E-10 |
| *N*-oleoyl tyrosine | VehFemale | CBDfemale | 1.05E-09 | 0.1465 | -5.54E-10 | 3.64E-09 |
| | | THCfemale | 1.02E-09 | 0.0228 | 3.43E-10 | 4.42E-09 |
| | | TpCfemale | 9.96E-10 | 0.0564 | -5.47E-11 | 3.92E-09 |
| | | VehMale | 1.05E-09 | 0.5483 | -1.46E-09 | 2.73E-09 |
| | | CBDmale | 9.96E-10 | 0.1988 | -6.96E-10 | 3.28E-09 |

| | | | | | | |
|---|---|---|---|---|---|---|
| | | THCmale | 1.02E-09 | 0.3772 | -1.13E-09 | 2.94E-09 |
| | | TpCmale | 9.96E-10 | 0.6969 | -2.38E-09 | 1.60E-09 |
| | CBDfemale | VehFemale | 1.05E-09 | 0.1465 | -3.64E-09 | 5.54E-10 |
| | | THCfemale | 1.02E-09 | 0.4146 | -1.20E-09 | 2.87E-09 |
| | | TpCfemale | 9.96E-10 | 0.6953 | -1.60E-09 | 2.38E-09 |
| | | VehMale | 1.05E-09 | 0.3895 | -3.01E-09 | 1.19E-09 |
| | | CBDmale | 9.96E-10 | 0.8028 | -2.24E-09 | 1.74E-09 |
| | | THCmale | 1.02E-09 | 0.5353 | -2.67E-09 | 1.40E-09 |
| | | TpCmale | 9.96E-10 | 0.0567 | -3.92E-09 | 5.69E-11 |
| | THCfemale | VehFemale | 1.02E-09 | 0.0228 | -4.42E-09 | -3.43E-10 |
| | | CBDfemale | 1.02E-09 | 0.4146 | -2.87E-09 | 1.20E-09 |
| | | TpCfemale | 9.64E-10 | 0.6454 | -2.37E-09 | 1.48E-09 |
| | | VehMale | 1.02E-09 | 0.0916 | -3.78E-09 | 2.91E-10 |
| | | CBDmale | 9.64E-10 | 0.2638 | -3.01E-09 | 8.39E-10 |
| | | THCmale | 9.89E-10 | 0.1414 | -3.45E-09 | 5.03E-10 |
| | | TpCmale | 9.64E-10 | 0.0055 | -4.70E-09 | -8.43E-10 |
| | TpCfemale | VehFemale | 9.96E-10 | 0.0564 | -3.92E-09 | 5.47E-11 |
| | | CBDfemale | 9.96E-10 | 0.6953 | -2.38E-09 | 1.60E-09 |
| | | THCfemale | 9.64E-10 | 0.6454 | -1.48E-09 | 2.37E-09 |
| | | VehMale | 9.96E-10 | 0.1960 | -3.29E-09 | 6.88E-10 |
| | | CBDmale | 9.39E-10 | 0.4969 | -2.52E-09 | 1.23E-09 |
| | | THCmale | 9.64E-10 | 0.2908 | -2.95E-09 | 8.99E-10 |
| | | TpCmale | 9.39E-10 | 0.0160 | -4.20E-09 | -4.49E-10 |
| | VehMale | VehFemale | 1.05E-09 | 0.5483 | -2.73E-09 | 1.46E-09 |
| | | CBDfemale | 1.05E-09 | 0.3895 | -1.19E-09 | 3.01E-09 |
| | | THCfemale | 1.02E-09 | 0.0916 | -2.91E-10 | 3.78E-09 |
| | | TpCfemale | 9.96E-10 | 0.1960 | -6.88E-10 | 3.29E-09 |
| | | CBDmale | 9.96E-10 | 0.5101 | -1.33E-09 | 2.65E-09 |
| | | THCmale | 1.02E-09 | 0.7894 | -1.76E-09 | 2.31E-09 |
| | | TpCmale | 9.96E-10 | 0.3081 | -3.01E-09 | 9.66E-10 |
| | CBDmale | VehFemale | 9.96E-10 | 0.1988 | -3.28E-09 | 6.96E-10 |
| | | CBDfemale | 9.96E-10 | 0.8028 | -1.74E-09 | 2.24E-09 |
| | | THCfemale | 9.64E-10 | 0.2638 | -8.39E-10 | 3.01E-09 |
| | | TpCfemale | 9.39E-10 | 0.4969 | -1.23E-09 | 2.52E-09 |
| | | VehMale | 9.96E-10 | 0.5101 | -2.65E-09 | 1.33E-09 |
| | | THCmale | 9.64E-10 | 0.6904 | -2.31E-09 | 1.54E-09 |
| | | TpCmale | 9.39E-10 | 0.0778 | -3.56E-09 | 1.93E-10 |
| | THCmale | VehFemale | 1.02E-09 | 0.3772 | -2.94E-09 | 1.13E-09 |
| | | CBDfemale | 1.02E-09 | 0.5353 | -1.40E-09 | 2.67E-09 |
| | | THCfemale | 9.89E-10 | 0.1414 | -5.03E-10 | 3.45E-09 |
| | | TpCfemale | 9.64E-10 | 0.2908 | -8.99E-10 | 2.95E-09 |
| | | VehMale | 1.02E-09 | 0.7894 | -2.31E-09 | 1.76E-09 |
| | | CBDmale | 9.64E-10 | 0.6904 | -1.54E-09 | 2.31E-09 |
| | | TpCmale | 9.64E-10 | 0.1836 | -3.22E-09 | 6.30E-10 |
| | TpCmale | VehFemale | 9.96E-10 | 0.6969 | -1.60E-09 | 2.38E-09 |
| | | CBDfemale | 9.96E-10 | 0.0567 | -5.69E-11 | 3.92E-09 |
| | | THCfemale | 9.64E-10 | 0.0055 | 8.43E-10 | 4.70E-09 |
| | | TpCfemale | 9.39E-10 | 0.0160 | 4.49E-10 | 4.20E-09 |
| | | VehMale | 9.96E-10 | 0.3081 | -9.66E-10 | 3.01E-09 |
| | | CBDmale | 9.39E-10 | 0.0778 | -1.93E-10 | 3.56E-09 |
| | | THCmale | 9.64E-10 | 0.1836 | -6.30E-10 | 3.22E-09 |
| *N*-linoleoyl tyrosine | VehFemale | CBDfemale | 5.92E-10 | 0.1394 | -2.97E-10 | 2.07E-09 |
| | | THCfemale | 5.92E-10 | 0.0364 | 8.25E-11 | 2.45E-09 |
| | | TpCfemale | 5.53E-10 | 0.2993 | -5.26E-10 | 1.68E-09 |
| | | VehMale | 5.92E-10 | 0.2846 | -5.44E-10 | 1.82E-09 |
| | | CBDmale | 5.64E-10 | 0.0928 | -1.64E-10 | 2.09E-09 |
| | | THCmale | 5.76E-10 | 0.8455 | -1.04E-09 | 1.26E-09 |
| | | TpCmale | 5.64E-10 | 0.9214 | -1.18E-09 | 1.07E-09 |
| | CBDfemale | VehFemale | 5.92E-10 | 0.1394 | -2.07E-09 | 2.97E-10 |
| | | THCfemale | 5.72E-10 | 0.5095 | -7.63E-10 | 1.52E-09 |

| | | | | | | |
|---|---|---|---|---|---|---|
| | | TpCfemale | 5.31E-10 | 0.5650 | -1.37E-09 | 7.55E-10 |
| | | VehMale | 5.72E-10 | 0.6669 | -1.39E-09 | 8.95E-10 |
| | | CBDmale | 5.42E-10 | 0.8892 | -1.01E-09 | 1.16E-09 |
| | | THCmale | 5.56E-10 | 0.1690 | -1.88E-09 | 3.37E-10 |
| | | TpCmale | 5.42E-10 | 0.0874 | -2.03E-09 | 1.42E-10 |
| | THCfemale | VehFemale | 5.92E-10 | 0.0364 | -2.45E-09 | -8.25E-11 |
| | | CBDfemale | 5.72E-10 | 0.5095 | -1.52E-09 | 7.63E-10 |
| | | TpCfemale | 5.31E-10 | 0.2010 | -1.75E-09 | 3.75E-10 |
| | | VehMale | 5.72E-10 | 0.2774 | -1.77E-09 | 5.16E-10 |
| | | CBDmale | 5.42E-10 | 0.5779 | -1.39E-09 | 7.81E-10 |
| | | THCmale | 5.56E-10 | 0.0422 | -2.26E-09 | -4.21E-11 |
| | | TpCmale | 5.42E-10 | 0.0177 | -2.41E-09 | -2.37E-10 |
| | TpCfemale | VehFemale | 5.53E-10 | 0.2993 | -1.68E-09 | 5.26E-10 |
| | | CBDfemale | 5.31E-10 | 0.5650 | -7.55E-10 | 1.37E-09 |
| | | THCfemale | 5.31E-10 | 0.2010 | -3.75E-10 | 1.75E-09 |
| | | VehMale | 5.31E-10 | 0.9103 | -1.00E-09 | 1.12E-09 |
| | | CBDmale | 5.00E-10 | 0.4460 | -6.15E-10 | 1.38E-09 |
| | | THCmale | 5.14E-10 | 0.3682 | -1.49E-09 | 5.61E-10 |
| | | TpCmale | 5.00E-10 | 0.2088 | -1.63E-09 | 3.64E-10 |
| | VehMale | VehFemale | 5.92E-10 | 0.2846 | -1.82E-09 | 5.44E-10 |
| | | CBDfemale | 5.72E-10 | 0.6669 | -8.95E-10 | 1.39E-09 |
| | | THCfemale | 5.72E-10 | 0.2774 | -5.16E-10 | 1.77E-09 |
| | | TpCfemale | 5.31E-10 | 0.9103 | -1.12E-09 | 1.00E-09 |
| | | CBDmale | 5.42E-10 | 0.5535 | -7.61E-10 | 1.41E-09 |
| | | THCmale | 5.56E-10 | 0.3475 | -1.64E-09 | 5.85E-10 |
| | | TpCmale | 5.42E-10 | 0.2051 | -1.78E-09 | 3.89E-10 |
| | CBDmale | VehFemale | 5.64E-10 | 0.0928 | -2.09E-09 | 1.64E-10 |
| | | CBDfemale | 5.42E-10 | 0.8892 | -1.16E-09 | 1.01E-09 |
| | | THCfemale | 5.42E-10 | 0.5779 | -7.81E-10 | 1.39E-09 |
| | | TpCfemale | 5.00E-10 | 0.4460 | -1.38E-09 | 6.15E-10 |
| | | VehMale | 5.42E-10 | 0.5535 | -1.41E-09 | 7.61E-10 |
| | | THCmale | 5.25E-10 | 0.1111 | -1.90E-09 | 2.01E-10 |
| | | TpCmale | 5.11E-10 | 0.0510 | -2.04E-09 | 4.34E-12 |
| | THCmale | VehFemale | 5.76E-10 | 0.8455 | -1.26E-09 | 1.04E-09 |
| | | CBDfemale | 5.56E-10 | 0.1690 | -3.37E-10 | 1.88E-09 |
| | | THCfemale | 5.56E-10 | 0.0422 | 4.21E-11 | 2.26E-09 |
| | | TpCfemale | 5.14E-10 | 0.3682 | -5.61E-10 | 1.49E-09 |
| | | VehMale | 5.56E-10 | 0.3475 | -5.85E-10 | 1.64E-09 |
| | | CBDmale | 5.25E-10 | 0.1111 | -2.01E-10 | 1.90E-09 |
| | | TpCmale | 5.25E-10 | 0.7494 | -1.22E-09 | 8.82E-10 |
| | TpCmale | VehFemale | 5.64E-10 | 0.9214 | -1.07E-09 | 1.18E-09 |
| | | CBDfemale | 5.42E-10 | 0.0874 | -1.42E-10 | 2.03E-09 |
| | | THCfemale | 5.42E-10 | 0.0177 | 2.37E-10 | 2.41E-09 |
| | | TpCfemale | 5.00E-10 | 0.2088 | -3.64E-10 | 1.63E-09 |
| | | VehMale | 5.42E-10 | 0.2051 | -3.89E-10 | 1.78E-09 |
| | | CBDmale | 5.11E-10 | 0.0510 | -4.34E-12 | 2.04E-09 |
| | | THCmale | 5.25E-10 | 0.7494 | -8.82E-10 | 1.22E-09 |
| *N*-docosahexaenoyl tyrosine | VehFemale | CBDfemale | 1.58E-10 | 0.4257 | -1.98E-10 | 4.54E-10 |
| | | TpCfemale | 1.65E-10 | 0.9194 | -3.57E-10 | 3.23E-10 |
| | | VehMale | 1.75E-10 | 0.4833 | -2.36E-10 | 4.85E-10 |
| | CBDfemale | VehFemale | 1.58E-10 | 0.4257 | -4.54E-10 | 1.98E-10 |
| | | TpCfemale | 1.58E-10 | 0.3681 | -4.71E-10 | 1.81E-10 |
| | | VehMale | 1.68E-10 | 0.9840 | -3.51E-10 | 3.44E-10 |
| | TpCfemale | VehFemale | 1.65E-10 | 0.9194 | -3.23E-10 | 3.57E-10 |
| | | CBDfemale | 1.58E-10 | 0.3681 | -1.81E-10 | 4.71E-10 |
| | | VehMale | 1.75E-10 | 0.4267 | -2.20E-10 | 5.02E-10 |
| | VehMale | VehFemale | 1.75E-10 | 0.4833 | -4.85E-10 | 2.36E-10 |
| | | CBDfemale | 1.68E-10 | 0.9840 | -3.44E-10 | 3.51E-10 |
| | | TpCfemale | 1.75E-10 | 0.4267 | -5.02E-10 | 2.20E-10 |

#### Supplemental Table 7.8 Fisher's LSD post-hoc test for liver *N*-acyl tyrosine ANOVA

| Dependent Variable | | | Std. Error | Sig. | 95% Confidence Interval Lower Bound | 95% Confidence Interval Upper Bound |
|---|---|---|---|---|---|---|
| *N*-palmitoyl tyrosine | VehFemale | CBDfemale | 2.06348E-11 | 0.936 | -4.2938E-11 | 3.9586E-11 |
| | | THCfemale | 1.89121E-11 | 0.412 | -5.3456E-11 | 2.2179E-11 |
| | | TpCfemale | 1.93735E-11 | 0.018 | -8.5647E-11 | -8.1673E-12 |
| | | VehMale | 2.06348E-11 | 0.070 | -3.2253E-12 | 7.9298E-11 |
| | | CBDmale | 1.93735E-11 | 0.005 | 1.7173E-11 | 9.4653E-11 |
| | | THCmale | 1.93735E-11 | 0.387 | -5.5623E-11 | 2.1857E-11 |
| | | TpCmale | 1.89121E-11 | 0.016 | 9.1193E-12 | 8.4754E-11 |
| | CBDfemale | VehFemale | 2.06348E-11 | 0.936 | -3.9586E-11 | 4.2938E-11 |
| | | THCfemale | 1.96483E-11 | 0.480 | -5.3251E-11 | 2.5327E-11 |
| | | TpCfemale | 2.00927E-11 | 0.028 | -8.5408E-11 | -5.0527E-12 |
| | | VehMale | 2.13115E-11 | 0.067 | -2.9022E-12 | 8.2328E-11 |
| | | CBDmale | 2.00927E-11 | 0.006 | 1.7412E-11 | 9.7767E-11 |
| | | THCmale | 2.00927E-11 | 0.452 | -5.5384E-11 | 2.4971E-11 |
| | | TpCmale | 1.96483E-11 | 0.016 | 9.3236E-12 | 8.7902E-11 |
| | THCfemale | VehFemale | 1.89121E-11 | 0.412 | -2.2179E-11 | 5.3456E-11 |
| | | CBDfemale | 1.96483E-11 | 0.480 | -2.5327E-11 | 5.3251E-11 |
| | | TpCfemale | 1.83191E-11 | 0.093 | -6.7900E-11 | 5.3629E-12 |
| | | VehMale | 1.96483E-11 | 0.008 | 1.4386E-11 | 9.2964E-11 |
| | | CBDmale | 1.83191E-11 | 0.000 | 3.4920E-11 | 1.0818E-10 |
| | | THCmale | 1.83191E-11 | 0.946 | -3.7876E-11 | 3.5387E-11 |
| | | TpCmale | 1.78305E-11 | 0.001 | 2.6921E-11 | 9.8229E-11 |
| | TpCfemale | VehFemale | 1.93735E-11 | 0.018 | 8.1673E-12 | 8.5647E-11 |
| | | CBDfemale | 2.00927E-11 | 0.028 | 5.0527E-12 | 8.5408E-11 |
| | | THCfemale | 1.83191E-11 | 0.093 | -5.3629E-12 | 6.7900E-11 |
| | | VehMale | 2.00927E-11 | 0.000 | 4.4766E-11 | 1.2512E-10 |
| | | CBDmale | 1.87950E-11 | 0.000 | 6.5237E-11 | 1.4040E-10 |
| | | THCmale | 1.87950E-11 | 0.115 | -7.5590E-12 | 6.7607E-11 |
| | | TpCmale | 1.83191E-11 | 0.000 | 5.7212E-11 | 1.3047E-10 |
| | VehMale | VehFemale | 2.06348E-11 | 0.070 | -7.9298E-11 | 3.2253E-12 |
| | | CBDfemale | 2.13115E-11 | 0.067 | -8.2328E-11 | 2.9022E-12 |
| | | THCfemale | 1.96483E-11 | 0.008 | -9.2964E-11 | -1.4386E-11 |
| | | TpCfemale | 2.00927E-11 | 0.000 | -1.2512E-10 | -4.4766E-11 |
| | | CBDmale | 2.00927E-11 | 0.377 | -2.2301E-11 | 5.8054E-11 |
| | | THCmale | 2.00927E-11 | 0.008 | -9.5097E-11 | -1.4742E-11 |
| | | TpCmale | 1.96483E-11 | 0.652 | -3.0389E-11 | 4.8189E-11 |
| | CBDmale | VehFemale | 1.93735E-11 | 0.005 | -9.4653E-11 | -1.7173E-11 |
| | | CBDfemale | 2.00927E-11 | 0.006 | -9.7767E-11 | -1.7412E-11 |
| | | THCfemale | 1.83191E-11 | 0.000 | -1.0818E-10 | -3.4920E-11 |
| | | TpCfemale | 1.87950E-11 | 0.000 | -1.4040E-10 | -6.5237E-11 |
| | | VehMale | 2.00927E-11 | 0.377 | -5.8054E-11 | 2.2301E-11 |
| | | THCmale | 1.87950E-11 | 0.000 | -1.1038E-10 | -3.5213E-11 |
| | | TpCmale | 1.83191E-11 | 0.626 | -4.5608E-11 | 2.7655E-11 |
| | THCmale | VehFemale | 1.93735E-11 | 0.387 | -2.1857E-11 | 5.5623E-11 |
| | | CBDfemale | 2.00927E-11 | 0.452 | -2.4971E-11 | 5.5384E-11 |
| | | THCfemale | 1.83191E-11 | 0.946 | -3.5387E-11 | 3.7876E-11 |
| | | TpCfemale | 1.87950E-11 | 0.115 | -6.7607E-11 | 7.5590E-12 |
| | | VehMale | 2.00927E-11 | 0.008 | 1.4742E-11 | 9.5097E-11 |
| | | CBDmale | 1.87950E-11 | 0.000 | 3.5213E-11 | 1.1038E-10 |
| | | TpCmale | 1.83191E-11 | 0.001 | 2.7188E-11 | 1.0045E-10 |
| | TpCmale | VehFemale | 1.89121E-11 | 0.016 | -8.4754E-11 | -9.1193E-12 |
| | | CBDfemale | 1.96483E-11 | 0.016 | -8.7902E-11 | -9.3236E-12 |
| | | THCfemale | 1.78305E-11 | 0.001 | -9.8229E-11 | -2.6921E-11 |
| | | TpCfemale | 1.83191E-11 | 0.000 | -1.3047E-10 | -5.7212E-11 |
| | | VehMale | 1.96483E-11 | 0.652 | -4.8189E-11 | 3.0389E-11 |
| | | CBDmale | 1.83191E-11 | 0.626 | -2.7655E-11 | 4.5608E-11 |
| | | THCmale | 1.83191E-11 | 0.001 | -1.0045E-10 | -2.7188E-11 |
| *N*-stearoyl tyrosine | VehFemale | CBDfemale | 2.70322E-12 | 0.019 | -1.1942E-11 | -1.1308E-12 |

| | | | | | | |
|---|---|---|---|---|---|---|
| | | THCfemale | 2.62705E-12 | 0.655 | -6.4341E-12 | 4.0722E-12 |
| | | TpCfemale | 2.56450E-12 | 0.024 | -1.1061E-11 | -8.0471E-13 |
| | | VehMale | 2.79809E-12 | 0.175 | -1.7530E-12 | 9.4373E-12 |
| | | CBDmale | 2.62705E-12 | 0.182 | -1.7054E-12 | 8.8009E-12 |
| | | THCmale | 2.62705E-12 | 0.739 | -6.1309E-12 | 4.3753E-12 |
| | | TpCmale | 2.62705E-12 | 0.414 | -3.0941E-12 | 7.4121E-12 |
| | CBDfemale | VehFemale | 2.70322E-12 | 0.019 | 1.1308E-12 | 1.1942E-11 |
| | | THCfemale | 2.62705E-12 | 0.046 | 1.0213E-13 | 1.0608E-11 |
| | | TpCfemale | 2.56450E-12 | 0.815 | -4.5246E-12 | 5.7315E-12 |
| | | VehMale | 2.79809E-12 | 0.000 | 4.7832E-12 | 1.5973E-11 |
| | | CBDmale | 2.62705E-12 | 0.000 | 4.8308E-12 | 1.5337E-11 |
| | | THCmale | 2.62705E-12 | 0.035 | 4.0526E-13 | 1.0912E-11 |
| | | TpCmale | 2.62705E-12 | 0.002 | 3.4421E-12 | 1.3948E-11 |
| | THCfemale | VehFemale | 2.62705E-12 | 0.655 | -4.0722E-12 | 6.4341E-12 |
| | | CBDfemale | 2.62705E-12 | 0.046 | -1.0608E-11 | -1.0213E-13 |
| | | TpCfemale | 2.48408E-12 | 0.060 | -9.7190E-12 | 2.1545E-13 |
| | | VehMale | 2.72459E-12 | 0.070 | -4.2508E-13 | 1.0471E-11 |
| | | CBDmale | 2.54862E-12 | 0.068 | -3.6757E-13 | 9.8250E-12 |
| | | THCmale | 2.54862E-12 | 0.906 | -4.7931E-12 | 5.3994E-12 |
| | | TpCmale | 2.54862E-12 | 0.195 | -1.7563E-12 | 8.4362E-12 |
| | TpCfemale | VehFemale | 2.56450E-12 | 0.024 | 8.0471E-13 | 1.1061E-11 |
| | | CBDfemale | 2.56450E-12 | 0.815 | -5.7315E-12 | 4.5246E-12 |
| | | THCfemale | 2.48408E-12 | 0.060 | -2.1545E-13 | 9.7190E-12 |
| | | VehMale | 2.66432E-12 | 0.001 | 4.4472E-12 | 1.5102E-11 |
| | | CBDmale | 2.48408E-12 | 0.000 | 4.5133E-12 | 1.4448E-11 |
| | | THCmale | 2.48408E-12 | 0.046 | 8.7683E-14 | 1.0022E-11 |
| | | TpCmale | 2.48408E-12 | 0.002 | 3.1245E-12 | 1.3059E-11 |
| | VehMale | VehFemale | 2.79809E-12 | 0.175 | -9.4373E-12 | 1.7530E-12 |
| | | CBDfemale | 2.79809E-12 | 0.000 | -1.5973E-11 | -4.7832E-12 |
| | | THCfemale | 2.72459E-12 | 0.070 | -1.0471E-11 | 4.2508E-13 |
| | | TpCfemale | 2.66432E-12 | 0.001 | -1.5102E-11 | -4.4472E-12 |
| | | CBDmale | 2.72459E-12 | 0.914 | -5.7425E-12 | 5.1538E-12 |
| | | THCmale | 2.72459E-12 | 0.088 | -1.0168E-11 | 7.2821E-13 |
| | | TpCmale | 2.72459E-12 | 0.539 | -7.1312E-12 | 3.7651E-12 |
| | CBDmale | VehFemale | 2.62705E-12 | 0.182 | -8.8009E-12 | 1.7054E-12 |
| | | CBDfemale | 2.62705E-12 | 0.000 | -1.5337E-11 | -4.8308E-12 |
| | | THCfemale | 2.54862E-12 | 0.068 | -9.8250E-12 | 3.6757E-13 |
| | | TpCfemale | 2.48408E-12 | 0.000 | -1.4448E-11 | -4.5133E-12 |
| | | VehMale | 2.72459E-12 | 0.914 | -5.1538E-12 | 5.7425E-12 |
| | | THCmale | 2.54862E-12 | 0.088 | -9.5218E-12 | 6.7070E-13 |
| | | TpCmale | 2.54862E-12 | 0.588 | -6.4850E-12 | 3.7075E-12 |
| | THCmale | VehFemale | 2.62705E-12 | 0.739 | -4.3753E-12 | 6.1309E-12 |
| | | CBDfemale | 2.62705E-12 | 0.035 | -1.0912E-11 | -4.0526E-13 |
| | | THCfemale | 2.54862E-12 | 0.906 | -5.3994E-12 | 4.7931E-12 |
| | | TpCfemale | 2.48408E-12 | 0.046 | -1.0022E-11 | -8.7683E-14 |
| | | VehMale | 2.72459E-12 | 0.088 | -7.2821E-13 | 1.0168E-11 |
| | | CBDmale | 2.54862E-12 | 0.088 | -6.7070E-13 | 9.5218E-12 |
| | | TpCmale | 2.54862E-12 | 0.238 | -2.0594E-12 | 8.1331E-12 |
| | TpCmale | VehFemale | 2.62705E-12 | 0.414 | -7.4121E-12 | 3.0941E-12 |
| | | CBDfemale | 2.62705E-12 | 0.002 | -1.3948E-11 | -3.4421E-12 |
| | | THCfemale | 2.54862E-12 | 0.195 | -8.4362E-12 | 1.7563E-12 |
| | | TpCfemale | 2.48408E-12 | 0.002 | -1.3059E-11 | -3.1245E-12 |
| | | VehMale | 2.72459E-12 | 0.539 | -3.7651E-12 | 7.1312E-12 |
| | | CBDmale | 2.54862E-12 | 0.588 | -3.7075E-12 | 6.4850E-12 |
| | | THCmale | 2.54862E-12 | 0.238 | -8.1331E-12 | 2.0594E-12 |
| *N*-oleoyl tyrosine | VehFemale | CBDfemale | 3.51298E-11 | 0.874 | -7.5849E-11 | 6.4691E-11 |
| | | THCfemale | 3.23880E-11 | 0.000 | -1.8589E-10 | -5.6317E-11 |
| | | TpCfemale | 3.31206E-11 | 0.001 | -1.8620E-10 | -5.3693E-11 |
| | | VehMale | 3.51298E-11 | 0.609 | -5.2195E-11 | 8.8345E-11 |
| | | CBDmale | 3.23880E-11 | 0.212 | -2.3891E-11 | 1.0568E-10 |

| | | | | | | |
|---|---|---|---|---|---|---|
| | | THCmale | 3.40142E-11 | 0.871 | -7.3569E-11 | 6.2508E-11 |
| | | TpCmale | 3.23880E-11 | 0.872 | -5.9558E-11 | 7.0013E-11 |
| | CBDfemale | VehFemale | 3.51298E-11 | 0.874 | -6.4691E-11 | 7.5849E-11 |
| | | THCfemale | 3.23880E-11 | 0.001 | -1.8031E-10 | -5.0738E-11 |
| | | TpCfemale | 3.31206E-11 | 0.001 | -1.8062E-10 | -4.8114E-11 |
| | | VehMale | 3.51298E-11 | 0.503 | -4.6616E-11 | 9.3924E-11 |
| | | CBDmale | 3.23880E-11 | 0.157 | -1.8312E-11 | 1.1126E-10 |
| | | THCmale | 3.40142E-11 | 0.999 | -6.7991E-11 | 6.8087E-11 |
| | | TpCmale | 3.23880E-11 | 0.740 | -5.3979E-11 | 7.5592E-11 |
| | THCfemale | VehFemale | 3.23880E-11 | 0.000 | 5.6317E-11 | 1.8589E-10 |
| | | CBDfemale | 3.23880E-11 | 0.001 | 5.0738E-11 | 1.8031E-10 |
| | | TpCfemale | 3.01971E-11 | 0.970 | -5.9244E-11 | 6.1562E-11 |
| | | VehMale | 3.23880E-11 | 0.000 | 7.4392E-11 | 2.0396E-10 |
| | | CBDmale | 2.93917E-11 | 0.000 | 1.0321E-10 | 2.2079E-10 |
| | | THCmale | 3.11746E-11 | 0.000 | 5.3214E-11 | 1.7793E-10 |
| | | TpCmale | 2.93917E-11 | 0.000 | 6.7538E-11 | 1.8512E-10 |
| | TpCfemale | VehFemale | 3.31206E-11 | 0.001 | 5.3693E-11 | 1.8620E-10 |
| | | CBDfemale | 3.31206E-11 | 0.001 | 4.8114E-11 | 1.8062E-10 |
| | | THCfemale | 3.01971E-11 | 0.970 | -6.1562E-11 | 5.9244E-11 |
| | | VehMale | 3.31206E-11 | 0.000 | 7.1768E-11 | 2.0427E-10 |
| | | CBDmale | 3.01971E-11 | 0.000 | 1.0044E-10 | 2.2124E-10 |
| | | THCmale | 3.19350E-11 | 0.001 | 5.0534E-11 | 1.7829E-10 |
| | | TpCmale | 3.01971E-11 | 0.000 | 6.4768E-11 | 1.8557E-10 |
| | VehMale | VehFemale | 3.51298E-11 | 0.609 | -8.8345E-11 | 5.2195E-11 |
| | | CBDfemale | 3.51298E-11 | 0.503 | -9.3924E-11 | 4.6616E-11 |
| | | THCfemale | 3.23880E-11 | 0.000 | -2.0396E-10 | -7.4392E-11 |
| | | TpCfemale | 3.31206E-11 | 0.000 | -2.0427E-10 | -7.1768E-11 |
| | | CBDmale | 3.23880E-11 | 0.484 | -4.1965E-11 | 8.7606E-11 |
| | | THCmale | 3.40142E-11 | 0.490 | -9.1644E-11 | 4.4433E-11 |
| | | TpCmale | 3.23880E-11 | 0.693 | -7.7633E-11 | 5.1939E-11 |
| | CBDmale | VehFemale | 3.23880E-11 | 0.212 | -1.0568E-10 | 2.3891E-11 |
| | | CBDfemale | 3.23880E-11 | 0.157 | -1.1126E-10 | 1.8312E-11 |
| | | THCfemale | 2.93917E-11 | 0.000 | -2.2079E-10 | -1.0321E-10 |
| | | TpCfemale | 3.01971E-11 | 0.000 | -2.2124E-10 | -1.0044E-10 |
| | | VehMale | 3.23880E-11 | 0.484 | -8.7606E-11 | 4.1965E-11 |
| | | THCmale | 3.11746E-11 | 0.142 | -1.0878E-10 | 1.5933E-11 |
| | | TpCmale | 2.93917E-11 | 0.230 | -9.4460E-11 | 2.3125E-11 |
| | THCmale | VehFemale | 3.40142E-11 | 0.871 | -6.2508E-11 | 7.3569E-11 |
| | | CBDfemale | 3.40142E-11 | 0.999 | -6.8087E-11 | 6.7991E-11 |
| | | THCfemale | 3.11746E-11 | 0.000 | -1.7793E-10 | -5.3214E-11 |
| | | TpCfemale | 3.19350E-11 | 0.001 | -1.7829E-10 | -5.0534E-11 |
| | | VehMale | 3.40142E-11 | 0.490 | -4.4433E-11 | 9.1644E-11 |
| | | CBDmale | 3.11746E-11 | 0.142 | -1.5933E-11 | 1.0878E-10 |
| | | TpCmale | 3.11746E-11 | 0.731 | -5.1600E-11 | 7.3117E-11 |
| | TpCmale | VehFemale | 3.23880E-11 | 0.872 | -7.0013E-11 | 5.9558E-11 |
| | | CBDfemale | 3.23880E-11 | 0.740 | -7.5592E-11 | 5.3979E-11 |
| | | THCfemale | 2.93917E-11 | 0.000 | -1.8512E-10 | -6.7538E-11 |
| | | TpCfemale | 3.01971E-11 | 0.000 | -1.8557E-10 | -6.4768E-11 |
| | | VehMale | 3.23880E-11 | 0.693 | -5.1939E-11 | 7.7633E-11 |
| | | CBDmale | 2.93917E-11 | 0.230 | -2.3125E-11 | 9.4460E-11 |
| | | THCmale | 3.11746E-11 | 0.731 | -7.3117E-11 | 5.1600E-11 |
| *N*-linoleoyl tyrosine | VehFemale | CBDfemale | 2.83353E-11 | 0.479 | -7.6879E-11 | 3.6479E-11 |
| | | THCfemale | 2.67148E-11 | 0.445 | -7.3980E-11 | 3.2895E-11 |
| | | TpCfemale | 2.61239E-11 | 0.000 | -1.9102E-10 | -8.6511E-11 |
| | | VehMale | 2.83353E-11 | 0.144 | -1.4745E-11 | 9.8614E-11 |
| | | CBDmale | 2.67148E-11 | 0.054 | -9.2380E-13 | 1.0595E-10 |
| | | THCmale | 2.67148E-11 | 0.207 | -8.7501E-11 | 1.9374E-11 |
| | | TpCmale | 2.61239E-11 | 0.227 | -2.0337E-11 | 8.4174E-11 |
| | CBDfemale | VehFemale | 2.83353E-11 | 0.479 | -3.6479E-11 | 7.6879E-11 |
| | | THCfemale | 2.67148E-11 | 0.990 | -5.3780E-11 | 5.3095E-11 |

| | | | | | | |
|---|---|---|---|---|---|---|
| | | TpCfemale | 2.61239E-11 | 0.000 | -1.7082E-10 | -6.6311E-11 |
| | | VehMale | 2.83353E-11 | 0.032 | 5.4554E-12 | 1.1881E-10 |
| | | CBDmale | 2.67148E-11 | 0.008 | 1.9276E-11 | 1.2615E-10 |
| | | THCmale | 2.67148E-11 | 0.606 | -6.7301E-11 | 3.9574E-11 |
| | | TpCmale | 2.61239E-11 | 0.051 | -1.3684E-13 | 1.0437E-10 |
| | THCfemale | VehFemale | 2.67148E-11 | 0.445 | -3.2895E-11 | 7.3980E-11 |
| | | CBDfemale | 2.67148E-11 | 0.990 | -5.3095E-11 | 5.3780E-11 |
| | | TpCfemale | 2.43567E-11 | 0.000 | -1.6694E-10 | -6.9503E-11 |
| | | VehMale | 2.67148E-11 | 0.023 | 9.0393E-12 | 1.1591E-10 |
| | | CBDmale | 2.49894E-11 | 0.005 | 2.3070E-11 | 1.2304E-10 |
| | | THCmale | 2.49894E-11 | 0.590 | -6.3508E-11 | 3.6465E-11 |
| | | TpCmale | 2.43567E-11 | 0.035 | 3.7405E-12 | 1.0118E-10 |
| | TpCfemale | VehFemale | 2.61239E-11 | 0.000 | 8.6511E-11 | 1.9102E-10 |
| | | CBDfemale | 2.61239E-11 | 0.000 | 6.6311E-11 | 1.7082E-10 |
| | | THCfemale | 2.43567E-11 | 0.000 | 6.9503E-11 | 1.6694E-10 |
| | | VehMale | 2.61239E-11 | 0.000 | 1.2845E-10 | 2.3296E-10 |
| | | CBDmale | 2.43567E-11 | 0.000 | 1.4256E-10 | 2.4000E-10 |
| | | THCmale | 2.43567E-11 | 0.000 | 5.5982E-11 | 1.5342E-10 |
| | | TpCmale | 2.37071E-11 | 0.000 | 1.2326E-10 | 2.1811E-10 |
| | VehMale | VehFemale | 2.83353E-11 | 0.144 | -9.8614E-11 | 1.4745E-11 |
| | | CBDfemale | 2.83353E-11 | 0.032 | -1.1881E-10 | -5.4554E-12 |
| | | THCfemale | 2.67148E-11 | 0.023 | -1.1591E-10 | -9.0393E-12 |
| | | TpCfemale | 2.61239E-11 | 0.000 | -2.3296E-10 | -1.2845E-10 |
| | | CBDmale | 2.67148E-11 | 0.694 | -4.2858E-11 | 6.4017E-11 |
| | | THCmale | 2.67148E-11 | 0.006 | -1.2944E-10 | -2.2561E-11 |
| | | TpCmale | 2.61239E-11 | 0.703 | -6.2271E-11 | 4.2240E-11 |
| | CBDmale | VehFemale | 2.67148E-11 | 0.054 | -1.0595E-10 | 9.2380E-13 |
| | | CBDfemale | 2.67148E-11 | 0.008 | -1.2615E-10 | -1.9276E-11 |
| | | THCfemale | 2.49894E-11 | 0.005 | -1.2304E-10 | -2.3070E-11 |
| | | TpCfemale | 2.43567E-11 | 0.000 | -2.4000E-10 | -1.4256E-10 |
| | | VehMale | 2.67148E-11 | 0.694 | -6.4017E-11 | 4.2858E-11 |
| | | THCmale | 2.49894E-11 | 0.001 | -1.3656E-10 | -3.6591E-11 |
| | | TpCmale | 2.43567E-11 | 0.401 | -6.9316E-11 | 2.8125E-11 |
| | THCmale | VehFemale | 2.67148E-11 | 0.207 | -1.9374E-11 | 8.7501E-11 |
| | | CBDfemale | 2.67148E-11 | 0.606 | -3.9574E-11 | 6.7301E-11 |
| | | THCfemale | 2.49894E-11 | 0.590 | -3.6465E-11 | 6.3508E-11 |
| | | TpCfemale | 2.43567E-11 | 0.000 | -1.5342E-10 | -5.5982E-11 |
| | | VehMale | 2.67148E-11 | 0.006 | 2.2561E-11 | 1.2944E-10 |
| | | CBDmale | 2.49894E-11 | 0.001 | 3.6591E-11 | 1.3656E-10 |
| | | TpCmale | 2.43567E-11 | 0.009 | 1.7262E-11 | 1.1470E-10 |
| | TpCmale | VehFemale | 2.61239E-11 | 0.227 | -8.4174E-11 | 2.0337E-11 |
| | | CBDfemale | 2.61239E-11 | 0.051 | -1.0437E-10 | 1.3684E-13 |
| | | THCfemale | 2.43567E-11 | 0.035 | -1.0118E-10 | -3.7405E-12 |
| | | TpCfemale | 2.37071E-11 | 0.000 | -2.1811E-10 | -1.2326E-10 |
| | | VehMale | 2.61239E-11 | 0.703 | -4.2240E-11 | 6.2271E-11 |
| | | CBDmale | 2.43567E-11 | 0.401 | -2.8125E-11 | 6.9316E-11 |
| | | THCmale | 2.43567E-11 | 0.009 | -1.1470E-10 | -1.7262E-11 |
| *N*-arachidonoyl tyrosine | VehFemale | CBDfemale | 1.03219E-11 | 0.143 | -3.5949E-11 | 5.3308E-12 |
| | | THCfemale | 9.46021E-12 | 0.094 | -3.5032E-11 | 2.8015E-12 |
| | | TpCfemale | 9.69098E-12 | 0.006 | -4.7236E-11 | -8.4794E-12 |
| | | VehMale | 1.03219E-11 | 0.130 | -4.7782E-12 | 3.6502E-11 |
| | | CBDmale | 9.69098E-12 | 0.013 | 5.4100E-12 | 4.4167E-11 |
| | | THCmale | 9.69098E-12 | 0.218 | -3.1429E-11 | 7.3275E-12 |
| | | TpCmale | 9.46021E-12 | 0.056 | -4.9506E-13 | 3.7339E-11 |
| | CBDfemale | VehFemale | 1.03219E-11 | 0.143 | -5.3308E-12 | 3.5949E-11 |
| | | THCfemale | 9.82844E-12 | 0.935 | -2.0459E-11 | 1.8847E-11 |
| | | TpCfemale | 1.00508E-11 | 0.217 | -3.2646E-11 | 7.5492E-12 |
| | | VehMale | 1.06604E-11 | 0.005 | 9.8542E-12 | 5.2488E-11 |
| | | CBDmale | 1.00508E-11 | 0.000 | 2.0000E-11 | 6.0195E-11 |
| | | THCmale | 1.00508E-11 | 0.747 | -1.6839E-11 | 2.3356E-11 |

| | | | | | | |
|---|---|---|---|---|---|---|
| | | TpCmale | 9.82844E-12 | 0.001 | 1.4078E-11 | 5.3384E-11 |
| | THCfemale | VehFemale | 9.46021E-12 | 0.094 | -2.8015E-12 | 3.5032E-11 |
| | | CBDfemale | 9.82844E-12 | 0.935 | -1.8847E-11 | 2.0459E-11 |
| | | TpCfemale | 9.16357E-12 | 0.205 | -3.0066E-11 | 6.5813E-12 |
| | | VehMale | 9.82844E-12 | 0.002 | 1.2324E-11 | 5.1630E-11 |
| | | CBDmale | 9.16357E-12 | 0.000 | 2.2580E-11 | 5.9227E-11 |
| | | THCmale | 9.16357E-12 | 0.659 | -1.4259E-11 | 2.2388E-11 |
| | | TpCmale | 8.91917E-12 | 0.000 | 1.6702E-11 | 5.2372E-11 |
| | TpCfemale | VehFemale | 9.69098E-12 | 0.006 | 8.4794E-12 | 4.7236E-11 |
| | | CBDfemale | 1.00508E-11 | 0.217 | -7.5492E-12 | 3.2646E-11 |
| | | THCfemale | 9.16357E-12 | 0.205 | -6.5813E-12 | 3.0066E-11 |
| | | VehMale | 1.00508E-11 | 0.000 | 2.3622E-11 | 6.3817E-11 |
| | | CBDmale | 9.40163E-12 | 0.000 | 3.3846E-11 | 7.1446E-11 |
| | | THCmale | 9.40163E-12 | 0.098 | -2.9928E-12 | 3.4607E-11 |
| | | TpCmale | 9.16357E-12 | 0.000 | 2.7956E-11 | 6.4603E-11 |
| | VehMale | VehFemale | 1.03219E-11 | 0.130 | -3.6502E-11 | 4.7782E-12 |
| | | CBDfemale | 1.06604E-11 | 0.005 | -5.2488E-11 | -9.8542E-12 |
| | | THCfemale | 9.82844E-12 | 0.002 | -5.1630E-11 | -1.2324E-11 |
| | | TpCfemale | 1.00508E-11 | 0.000 | -6.3817E-11 | -2.3622E-11 |
| | | CBDmale | 1.00508E-11 | 0.378 | -1.1171E-11 | 2.9024E-11 |
| | | THCmale | 1.00508E-11 | 0.007 | -4.8010E-11 | -7.8149E-12 |
| | | TpCmale | 9.82844E-12 | 0.795 | -1.7093E-11 | 2.2213E-11 |
| | CBDmale | VehFemale | 9.69098E-12 | 0.013 | -4.4167E-11 | -5.4100E-12 |
| | | CBDfemale | 1.00508E-11 | 0.000 | -6.0195E-11 | -2.0000E-11 |
| | | THCfemale | 9.16357E-12 | 0.000 | -5.9227E-11 | -2.2580E-11 |
| | | TpCfemale | 9.40163E-12 | 0.000 | -7.1446E-11 | -3.3846E-11 |
| | | VehMale | 1.00508E-11 | 0.378 | -2.9024E-11 | 1.1171E-11 |
| | | THCmale | 9.40163E-12 | 0.000 | -5.5639E-11 | -1.8039E-11 |
| | | TpCmale | 9.16357E-12 | 0.490 | -2.4690E-11 | 1.1957E-11 |
| | THCmale | VehFemale | 9.69098E-12 | 0.218 | -7.3275E-12 | 3.1429E-11 |
| | | CBDfemale | 1.00508E-11 | 0.747 | -2.3356E-11 | 1.6839E-11 |
| | | THCfemale | 9.16357E-12 | 0.659 | -2.2388E-11 | 1.4259E-11 |
| | | TpCfemale | 9.40163E-12 | 0.098 | -3.4607E-11 | 2.9928E-12 |
| | | VehMale | 1.00508E-11 | 0.007 | 7.8149E-12 | 4.8010E-11 |
| | | CBDmale | 9.40163E-12 | 0.000 | 1.8039E-11 | 5.5639E-11 |
| | | TpCmale | 9.16357E-12 | 0.001 | 1.2149E-11 | 4.8796E-11 |
| | TpCmale | VehFemale | 9.46021E-12 | 0.056 | -3.7339E-11 | 4.9506E-13 |
| | | CBDfemale | 9.82844E-12 | 0.001 | -5.3384E-11 | -1.4078E-11 |
| | | THCfemale | 8.91917E-12 | 0.000 | -5.2372E-11 | -1.6702E-11 |
| | | TpCfemale | 9.16357E-12 | 0.000 | -6.4603E-11 | -2.7956E-11 |
| | | VehMale | 9.82844E-12 | 0.795 | -2.2213E-11 | 1.7093E-11 |
| | | CBDmale | 9.16357E-12 | 0.490 | -1.1957E-11 | 2.4690E-11 |
| | | THCmale | 9.16357E-12 | 0.001 | -4.8796E-11 | -1.2149E-11 |
| *N*-docosahexaenoyl tyrosine | VehFemale | CBDfemale | 2.22924E-11 | 0.437 | -6.2033E-11 | 2.7120E-11 |
| | | THCfemale | 2.04313E-11 | 0.058 | -8.0340E-11 | 1.3695E-12 |
| | | TpCfemale | 2.09297E-11 | 0.001 | -1.1797E-10 | -3.4263E-11 |
| | | VehMale | 2.22924E-11 | 0.540 | -3.0839E-11 | 5.8314E-11 |
| | | CBDmale | 2.09297E-11 | 0.071 | -3.3958E-12 | 8.0307E-11 |
| | | THCmale | 2.09297E-11 | 0.088 | -7.8131E-11 | 5.5721E-12 |
| | | TpCmale | 2.04313E-11 | 0.689 | -3.2628E-11 | 4.9082E-11 |
| | CBDfemale | VehFemale | 2.22924E-11 | 0.437 | -2.7120E-11 | 6.2033E-11 |
| | | THCfemale | 2.12266E-11 | 0.303 | -6.4474E-11 | 2.0416E-11 |
| | | TpCfemale | 2.17068E-11 | 0.009 | -1.0206E-10 | -1.5253E-11 |
| | | VehMale | 2.30235E-11 | 0.180 | -1.4845E-11 | 7.7232E-11 |
| | | CBDmale | 2.17068E-11 | 0.012 | 1.2507E-11 | 9.9318E-11 |
| | | THCmale | 2.17068E-11 | 0.389 | -6.2229E-11 | 2.4582E-11 |
| | | TpCmale | 2.12266E-11 | 0.231 | -1.6761E-11 | 6.8129E-11 |
| | THCfemale | VehFemale | 2.04313E-11 | 0.058 | -1.3695E-12 | 8.0340E-11 |
| | | CBDfemale | 2.12266E-11 | 0.303 | -2.0416E-11 | 6.4474E-11 |
| | | TpCfemale | 1.97907E-11 | 0.069 | -7.6204E-11 | 2.9444E-12 |

| | | | | | |
|---|---|---|---|---|---|
| | VehMale | 2.12266E-11 | 0.015 | 1.0777E-11 | 9.5668E-11 |
| | CBDmale | 1.97907E-11 | 0.000 | 3.8367E-11 | 1.1752E-10 |
| | THCmale | 1.97907E-11 | 0.872 | -3.6368E-11 | 4.2780E-11 |
| | TpCmale | 1.92629E-11 | 0.016 | 9.1945E-12 | 8.6231E-11 |
| TpCfemale | VehFemale | 2.09297E-11 | 0.001 | 3.4263E-11 | 1.1797E-10 |
| | CBDfemale | 2.17068E-11 | 0.009 | 1.5253E-11 | 1.0206E-10 |
| | THCfemale | 1.97907E-11 | 0.069 | -2.9444E-12 | 7.6204E-11 |
| | VehMale | 2.17068E-11 | 0.000 | 4.6447E-11 | 1.3326E-10 |
| | CBDmale | 2.03048E-11 | 0.000 | 7.3969E-11 | 1.5517E-10 |
| | THCmale | 2.03048E-11 | 0.054 | -7.6650E-13 | 8.0438E-11 |
| | TpCmale | 1.97907E-11 | 0.000 | 4.4769E-11 | 1.2392E-10 |
| VehMale | VehFemale | 2.22924E-11 | 0.540 | -5.8314E-11 | 3.0839E-11 |
| | CBDfemale | 2.30235E-11 | 0.180 | -7.7232E-11 | 1.4845E-11 |
| | THCfemale | 2.12266E-11 | 0.015 | -9.5668E-11 | -1.0777E-11 |
| | TpCfemale | 2.17068E-11 | 0.000 | -1.3326E-10 | -4.6447E-11 |
| | CBDmale | 2.17068E-11 | 0.259 | -1.8687E-11 | 6.8124E-11 |
| | THCmale | 2.17068E-11 | 0.025 | -9.3422E-11 | -6.6114E-12 |
| | TpCmale | 2.12266E-11 | 0.796 | -4.7955E-11 | 3.6935E-11 |
| CBDmale | VehFemale | 2.09297E-11 | 0.071 | -8.0307E-11 | 3.3958E-12 |
| | CBDfemale | 2.17068E-11 | 0.012 | -9.9318E-11 | -1.2507E-11 |
| | THCfemale | 1.97907E-11 | 0.000 | -1.1752E-10 | -3.8367E-11 |
| | TpCfemale | 2.03048E-11 | 0.000 | -1.5517E-10 | -7.3969E-11 |
| | VehMale | 2.17068E-11 | 0.259 | -6.8124E-11 | 1.8687E-11 |
| | THCmale | 2.03048E-11 | 0.000 | -1.1534E-10 | -3.4133E-11 |
| | TpCmale | 1.97907E-11 | 0.132 | -6.9802E-11 | 9.3456E-12 |
| THCmale | VehFemale | 2.09297E-11 | 0.088 | -5.5721E-12 | 7.8131E-11 |
| | CBDfemale | 2.17068E-11 | 0.389 | -2.4582E-11 | 6.2229E-11 |
| | THCfemale | 1.97907E-11 | 0.872 | -4.2780E-11 | 3.6368E-11 |
| | TpCfemale | 2.03048E-11 | 0.054 | -8.0438E-11 | 7.6650E-13 |
| | VehMale | 2.17068E-11 | 0.025 | 6.6114E-12 | 9.3422E-11 |
| | CBDmale | 2.03048E-11 | 0.000 | 3.4133E-11 | 1.1534E-10 |
| | TpCmale | 1.97907E-11 | 0.028 | 4.9330E-12 | 8.4081E-11 |
| TpCmale | VehFemale | 2.04313E-11 | 0.689 | -4.9082E-11 | 3.2628E-11 |
| | CBDfemale | 2.12266E-11 | 0.231 | -6.8129E-11 | 1.6761E-11 |
| | THCfemale | 1.92629E-11 | 0.016 | -8.6231E-11 | -9.1945E-12 |
| | TpCfemale | 1.97907E-11 | 0.000 | -1.2392E-10 | -4.4769E-11 |
| | VehMale | 2.12266E-11 | 0.796 | -3.6935E-11 | 4.7955E-11 |
| | CBDmale | 1.97907E-11 | 0.132 | -9.3456E-12 | 6.9802E-11 |
| | THCmale | 1.97907E-11 | 0.028 | -8.4081E-11 | -4.9330E-12 |

Supplemental Table 8.1 Descriptive statistics for *N*-acyl valines (mol/L) in plasma collected from male and female adult mice 2h after receiving an i.p. injection of 10mg/kg CBD, THC, or THC+CBD

| | | | | | | 95% Confidence Interval for Mean | | | |
|---|---|---|---|---|---|---|---|---|---|
| | | N | Mean | Std. Deviation | Std. Error | Lower Bound | Upper Bound | Minimum | Maximum |
| *N*-palmitoyl valine | Veh F | 8 | 1.9275E-09 | 1.02209E-09 | 3.61363E-10 | 1.0730E-09 | 2.7820E-09 | 4.15E-10 | 3.33E-09 |
| | CBD F | 8 | 1.1389E-09 | 4.05250E-10 | 1.43277E-10 | 8.0009E-10 | 1.4777E-09 | 4.56E-10 | 1.76E-09 |
| | THC F | 9 | 9.3458E-10 | 3.31965E-10 | 1.10655E-10 | 6.7941E-10 | 1.1898E-09 | 4.16E-10 | 1.47E-09 |
| | THC+CBD F | 11 | 1.6311E-09 | 1.03426E-09 | 3.11842E-10 | 9.3631E-10 | 2.3260E-09 | 2.99E-10 | 3.32E-09 |
| | Veh M | 7 | 9.7884E-10 | 3.14883E-10 | 1.19015E-10 | 6.8762E-10 | 1.2701E-09 | 5.16E-10 | 1.32E-09 |
| | CBD M | 10 | 8.7216E-10 | 3.34199E-10 | 1.05683E-10 | 6.3309E-10 | 1.1112E-09 | 4.50E-10 | 1.48E-09 |
| | THC M | 9 | 1.6122E-09 | 6.19428E-10 | 2.06476E-10 | 1.1360E-09 | 2.0883E-09 | 8.29E-10 | 2.69E-09 |
| | THC+CBD M | 10 | 1.9776E-09 | 1.28636E-09 | 4.06783E-10 | 1.0574E-09 | 2.8979E-09 | 5.66E-10 | 3.90E-09 |
| *N*-stearoyl valine | Veh F | 8 | 6.7157E-10 | 4.28254E-10 | 1.51411E-10 | 3.1354E-10 | 1.0296E-09 | 1.27E-10 | 1.36E-09 |
| | CBD F | 8 | 3.8691E-10 | 2.01109E-10 | 7.11027E-11 | 2.1878E-10 | 5.5504E-10 | 1.20E-10 | 7.32E-10 |
| | THC F | 8 | 4.2937E-10 | 1.65943E-10 | 5.86697E-11 | 2.9064E-10 | 5.6811E-10 | 6.97E-11 | 5.99E-10 |
| | THC+CBD F | 10 | 5.3934E-10 | 2.88352E-10 | 9.11848E-11 | 3.3306E-10 | 7.4561E-10 | 1.35E-10 | 1.06E-09 |
| | Veh M | 7 | 5.8493E-10 | 3.46319E-10 | 1.30896E-10 | 2.6464E-10 | 9.0522E-10 | 3.26E-10 | 1.28E-09 |
| | CBD M | 9 | 3.7434E-10 | 1.44369E-10 | 4.81231E-11 | 2.6337E-10 | 4.8532E-10 | 1.52E-10 | 6.08E-10 |
| | THC M | 9 | 5.2107E-10 | 2.01333E-10 | 6.71109E-11 | 3.6631E-10 | 6.7583E-10 | 2.17E-10 | 7.54E-10 |
| | THC+CBD M | 10 | 5.7356E-10 | 2.58103E-10 | 8.16193E-11 | 3.8893E-10 | 7.5820E-10 | 1.49E-10 | 1.06E-09 |
| *N*-oleoyl valine | Veh F | 8 | 3.2126E-10 | 2.00733E-10 | 7.09699E-11 | 1.5345E-10 | 4.8908E-10 | 5.29E-11 | 6.90E-10 |
| | CBD F | 8 | 2.0600E-10 | 9.76868E-11 | 3.45375E-11 | 1.2433E-10 | 2.8767E-10 | 6.67E-11 | 3.68E-10 |
| | THC F | 10 | 2.0990E-10 | 1.14000E-10 | 3.60501E-11 | 1.2835E-10 | 2.9145E-10 | 3.72E-11 | 3.46E-10 |
| | THC+CBD F | 11 | 2.9696E-10 | 1.89322E-10 | 5.70826E-11 | 1.6977E-10 | 4.2415E-10 | 6.59E-11 | 6.17E-10 |
| | Veh M | 7 | 2.3266E-10 | 1.06801E-10 | 4.03670E-11 | 1.3388E-10 | 3.3143E-10 | 1.13E-10 | 4.01E-10 |
| | CBD M | 9 | 1.6132E-10 | 8.04274E-11 | 2.68091E-11 | 9.9495E-11 | 2.2314E-10 | 3.79E-11 | 2.98E-10 |
| | THC M | 9 | 2.4304E-10 | 1.21296E-10 | 4.04320E-11 | 1.4980E-10 | 3.3628E-10 | 5.20E-11 | 4.85E-10 |
| | THC+CBD M | 10 | 3.4706E-10 | 1.84906E-10 | 5.84723E-11 | 2.1479E-10 | 4.7934E-10 | 1.04E-10 | 6.16E-10 |
| *N*-linoleoyl valine | Veh F | 8 | 7.1330E-10 | 5.09461E-10 | 1.80122E-10 | 2.8738E-10 | 1.1392E-09 | 9.59E-11 | 1.69E-09 |
| | CBD F | 8 | 4.0948E-10 | 2.76971E-10 | 9.79241E-11 | 1.7793E-10 | 6.4104E-10 | 8.43E-11 | 9.36E-10 |
| | THC F | 6 | 2.5341E-10 | 2.37200E-10 | 9.68365E-11 | 4.4796E-12 | 5.0233E-10 | 3.03E-11 | 6.99E-10 |
| | THC+CBD F | 10 | 5.0807E-10 | 3.98199E-10 | 1.25922E-10 | 2.2322E-10 | 7.9293E-10 | 6.16E-11 | 1.27E-09 |
| | Veh M | 7 | 3.6451E-10 | 2.35263E-10 | 8.89210E-11 | 1.4693E-10 | 5.8210E-10 | 9.44E-11 | 8.10E-10 |
| | CBD M | 10 | 2.8544E-10 | 9.29720E-11 | 2.94003E-11 | 2.1893E-10 | 3.5195E-10 | 1.42E-10 | 4.74E-10 |
| | THC M | 9 | 5.3501E-10 | 3.70330E-10 | 1.23443E-10 | 2.5035E-10 | 8.1967E-10 | 7.56E-11 | 1.18E-09 |
| | THC+CBD M | 10 | 4.7655E-10 | 2.58456E-10 | 8.17308E-11 | 2.9166E-10 | 6.6144E-10 | 1.23E-10 | 8.26E-10 |
| *N*-docosahexaenoyl valine | Veh F | BAL | | | | | | | |
| | CBD F | BAL | | | | | | | |
| | THC F | BAL | | | | | | | |
| | THC+CBD F | 4 | 2.2650E-10 | 2.26331E-10 | 1.13165E-10 | -1.3364E-10 | 5.8664E-10 | 7.25E-11 | 5.61E-10 |
| | Veh M | BDL | | | | | | | |
| | CBD M | BAL | | | | | | | |
| | THC M | BAL | | | | | | | |
| | THC+CBD M | BAL | | | | | | | |

Supplemental Table 8.2 Descriptive statistics for *N*-acyl valines (mol/g) in liver collected from male and female adult mice 2h after receiving an i.p. injection of 10mg/kg CBD, THC, or THC+CBD

| | | N | Mean | Std. Deviation | Std. Error | 95% Confidence Interval for Mean | | Minimum | Maximum |
|---|---|---|---|---|---|---|---|---|---|
| | | | | | | Lower Bound | Upper Bound | | |
| *N*-palmitoyl valine | Veh F | 8 | 6.3925E-11 | 3.67529E-11 | 1.29941E-11 | 3.3199E-11 | 9.4652E-11 | 1.70E-11 | 1.25E-10 |
| | CBD F | 8 | 9.5808E-11 | 7.52153E-11 | 2.65926E-11 | 3.2926E-11 | 1.5869E-10 | 2.42E-11 | 2.18E-10 |
| | THC F | 10 | 8.3140E-11 | 3.37345E-11 | 1.06678E-11 | 5.9008E-11 | 1.0727E-10 | 3.50E-11 | 1.20E-10 |
| | THC+CBD F | 10 | 1.2620E-10 | 3.73514E-11 | 1.18115E-11 | 9.9481E-11 | 1.5292E-10 | 6.77E-11 | 1.89E-10 |
| | Veh M | 7 | 6.6595E-11 | 3.37728E-11 | 1.27649E-11 | 3.5360E-11 | 9.7830E-11 | 2.15E-11 | 1.27E-10 |
| | CBD M | 9 | 3.7265E-11 | 1.43263E-11 | 4.77542E-12 | 2.6253E-11 | 4.8277E-11 | 2.18E-11 | 6.67E-11 |
| | THC M | 9 | 1.1013E-10 | 4.90763E-11 | 1.63588E-11 | 7.2408E-11 | 1.4786E-10 | 5.07E-11 | 1.99E-10 |
| | THC+CBD M | 10 | 4.8676E-11 | 1.94626E-11 | 6.15461E-12 | 3.4753E-11 | 6.2599E-11 | 2.91E-11 | 9.27E-11 |
| *N*-stearoyl valine | Veh F | 7 | 2.0595E-12 | 1.35236E-12 | 5.11144E-13 | 8.0875E-13 | 3.3102E-12 | 5.96E-13 | 3.93E-12 |
| | CBD F | 5 | 4.7042E-12 | 4.52467E-12 | 2.02350E-12 | -9.1388E-13 | 1.0322E-11 | 6.38E-13 | 1.17E-11 |
| | THC F | 9 | 3.1319E-12 | 1.30056E-12 | 4.33522E-13 | 2.1322E-12 | 4.1316E-12 | 1.20E-12 | 5.53E-12 |
| | THC+CBD F | 10 | 3.4450E-12 | 1.39671E-12 | 4.41679E-13 | 2.4459E-12 | 4.4442E-12 | 1.73E-12 | 5.91E-12 |
| | Veh M | 5 | 3.0207E-12 | 3.60272E-12 | 1.61119E-12 | -1.4527E-12 | 7.4940E-12 | 7.24E-13 | 9.32E-12 |
| | CBD M | 7 | 1.1564E-12 | 5.10489E-13 | 1.92947E-13 | 6.8432E-13 | 1.6286E-12 | 5.35E-13 | 2.02E-12 |
| | THC M | 9 | 3.1576E-12 | 2.18819E-12 | 7.29397E-13 | 1.4756E-12 | 4.8396E-12 | 3.80E-13 | 7.04E-12 |
| | THC+CBD M | 9 | 1.8042E-12 | 1.19850E-12 | 3.99500E-13 | 8.8295E-13 | 2.7254E-12 | 6.84E-13 | 4.41E-12 |
| *N*-oleoyl valine | Veh F | 8 | 5.9187E-12 | 3.52668E-12 | 1.24687E-12 | 2.9703E-12 | 8.8670E-12 | 1.35E-12 | 1.19E-11 |
| | CBD F | 7 | 5.3123E-12 | 2.72648E-12 | 1.03051E-12 | 2.7908E-12 | 7.8339E-12 | 2.45E-12 | 1.11E-11 |
| | THC F | 10 | 9.3277E-12 | 5.81898E-12 | 1.84012E-12 | 5.1651E-12 | 1.3490E-11 | 2.39E-12 | 1.78E-11 |
| | THC+CBD F | 10 | 1.2286E-11 | 3.44279E-12 | 1.08871E-12 | 9.8236E-12 | 1.4749E-11 | 8.79E-12 | 1.80E-11 |
| | Veh M | 7 | 4.9139E-12 | 1.78581E-12 | 6.74974E-13 | 3.2623E-12 | 6.5655E-12 | 2.55E-12 | 8.17E-12 |
| | CBD M | 9 | 2.9164E-12 | 7.86008E-13 | 2.62003E-13 | 2.3123E-12 | 3.5206E-12 | 2.00E-12 | 4.24E-12 |
| | THC M | 8 | 6.4173E-12 | 2.70754E-12 | 9.57260E-13 | 4.1537E-12 | 8.6809E-12 | 3.39E-12 | 1.08E-11 |
| | THC+CBD M | 10 | 5.0538E-12 | 1.75795E-12 | 5.55912E-13 | 3.7962E-12 | 6.3114E-12 | 2.47E-12 | 8.22E-12 |
| *N*-linoleoyl valine | Veh F | 8 | 1.9255E-11 | 1.21883E-11 | 4.30922E-12 | 9.0657E-12 | 2.9445E-11 | 5.67E-12 | 3.81E-11 |
| | CBD F | 8 | 3.0268E-11 | 3.06155E-11 | 1.08242E-11 | 4.6728E-12 | 5.5863E-11 | 2.13E-12 | 8.27E-11 |
| | THC F | 10 | 2.1365E-11 | 9.62365E-12 | 3.04326E-12 | 1.4481E-11 | 2.8250E-11 | 6.54E-12 | 3.89E-11 |
| | THC+CBD F | 10 | 4.4672E-11 | 1.53028E-11 | 4.83916E-12 | 3.3725E-11 | 5.5619E-11 | 2.13E-11 | 6.95E-11 |
| | Veh M | 7 | 1.1103E-11 | 6.60126E-12 | 2.49504E-12 | 4.9977E-12 | 1.7208E-11 | 2.86E-12 | 2.36E-11 |
| | CBD M | 9 | 8.1967E-12 | 3.84325E-12 | 1.28108E-12 | 5.2425E-12 | 1.1151E-11 | 4.86E-12 | 1.75E-11 |
| | THC M | 9 | 2.6484E-11 | 1.06849E-11 | 3.56162E-12 | 1.8271E-11 | 3.4697E-11 | 9.74E-12 | 4.01E-11 |
| | THC+CBD M | 10 | 1.3568E-11 | 7.33928E-12 | 2.32088E-12 | 8.3181E-12 | 1.8819E-11 | 4.99E-12 | 2.64E-11 |
| *N*-docosahexaenoyl valine | Veh F | 7 | 4.3789E-12 | 2.57828E-12 | 9.74498E-13 | 1.9944E-12 | 6.7634E-12 | 1.11E-12 | 7.86E-12 |
| | CBD F | 8 | 1.2588E-11 | 1.28529E-11 | 4.54420E-12 | 1.8427E-12 | 2.3333E-11 | 2.20E-12 | 3.49E-11 |
| | THC F | 10 | 1.3490E-11 | 6.82620E-12 | 2.15863E-12 | 8.6072E-12 | 1.8374E-11 | 1.67E-12 | 2.27E-11 |
| | THC+CBD F | 10 | 1.8591E-11 | 9.20833E-12 | 2.91193E-12 | 1.2004E-11 | 2.5179E-11 | 5.98E-12 | 3.43E-11 |
| | Veh M | 7 | 7.4869E-12 | 4.98390E-12 | 1.88374E-12 | 2.8776E-12 | 1.2096E-11 | 1.05E-12 | 1.69E-11 |
| | CBD M | 9 | 4.0816E-12 | 1.75013E-12 | 5.83378E-13 | 2.7364E-12 | 5.4269E-12 | 2.18E-12 | 7.41E-12 |
| | THC M | 9 | 1.5549E-11 | 8.93520E-12 | 2.97840E-12 | 8.6803E-12 | 2.2417E-11 | 5.13E-12 | 2.83E-11 |
| | THC+CBD M | 10 | 8.2981E-12 | 6.10497E-12 | 1.93056E-12 | 3.9308E-12 | 1.2665E-11 | 1.95E-12 | 1.99E-11 |

Supplemental Table 8.3 Independent t-test comparing male and female levels of *N*-acyl valines measured in plasma collected from adult mice 2h after an i.p. injection of 10mg/kg CBD, THC, or THC+CBD

| | | | Levene's Test for Equality of Variances | | t-test for Equality of Means | | | | | | |
|---|---|---|---|---|---|---|---|---|---|---|---|
| | | | | | | | Significance | | | 95% Confidence Interval of the Difference | |
| | | | F | Sig. | t | df | One-Sided p | Two-Sided p | Std. Error Difference | Lower | Upper |
| Veh | *N*-palmitoyl valine | Equal variances assumed | 7.618 | 0.016 | 2.350 | 13 | 0.018 | 0.035 | 4.03647E-10 | 7.66606E-11 | 1.82071E-09 |
| | | Equal variances not assumed | | | 2.494 | 8.484 | 0.018 | 0.036 | 3.80457E-10 | 7.99947E-11 | 1.81738E-09 |
| | *N*-stearoyl valine | Equal variances assumed | 0.381 | 0.548 | 0.426 | 13 | 0.338 | 0.677 | 2.03174E-10 | -3.52293E-10 | 5.25568E-10 |
| | | Equal variances not assumed | | | 0.433 | 12.940 | 0.336 | 0.672 | 2.00148E-10 | -3.45958E-10 | 5.19233E-10 |
| | *N*-oleoyl valine | Equal variances assumed | 1.998 | 0.181 | 1.043 | 13 | 0.158 | 0.316 | 8.49808E-11 | -9.49839E-11 | 2.72196E-10 |
| | | Equal variances not assumed | | | 1.085 | 10.928 | 0.151 | 0.301 | 8.16469E-11 | -9.12430E-11 | 2.68455E-10 |
| | *N*-linoleoyl valine | Equal variances assumed | 2.131 | 0.168 | 1.658 | 13 | 0.061 | 0.121 | 2.10423E-10 | -1.05807E-10 | 8.03374E-10 |
| | | Equal variances not assumed | | | 1.736 | 10.126 | 0.056 | 0.113 | 2.00875E-10 | -9.80393E-11 | 7.95607E-10 |
| CBD | *N*-palmitoyl valine | Equal variances assumed | 0.465 | 0.505 | 1.532 | 16 | 0.072 | 0.145 | 1.74074E-10 | -1.02291E-10 | 6.35750E-10 |
| | | Equal variances not assumed | | | 1.498 | 13.566 | 0.078 | 0.157 | 1.78037E-10 | -1.16272E-10 | 6.49732E-10 |
| | *N*-stearoyl valine | Equal variances assumed | 1.192 | 0.292 | 0.149 | 15 | 0.442 | 0.883 | 8.41488E-11 | -1.66793E-10 | 1.91925E-10 |
| | | Equal variances not assumed | | | 0.146 | 12.573 | 0.443 | 0.886 | 8.58570E-11 | -1.73559E-10 | 1.98690E-10 |
| | *N*-oleoyl valine | Equal variances assumed | 0.470 | 0.503 | 1.034 | 15 | 0.159 | 0.317 | 4.31975E-11 | -4.73876E-11 | 1.36759E-10 |
| | | Equal variances not assumed | | | 1.022 | 13.643 | 0.162 | 0.325 | 4.37215E-11 | -4.93183E-11 | 1.38690E-10 |
| | *N*-linoleoyl valine | Equal variances assumed | 5.300 | 0.035 | 1.334 | 16 | 0.100 | 0.201 | 9.29808E-11 | -7.30684E-11 | 3.21152E-10 |

| | | | | | | | | | | | |
|---|---|---|---|---|---|---|---|---|---|---|---|
| | | Equal variances not assumed | | | 1.213 | 8.267 | 0.129 | 0.259 | 1.02242E-10 | -1.10412E-10 | 3.58496E-10 |
| THC | *N*-palmitoyl valine | Equal variances assumed | 2.822 | 0.112 | -2.893 | 16 | 0.005 | 0.011 | 2.34258E-10 | -1.17420E-09 | -1.80988E-10 |
| | | Equal variances not assumed | | | -2.893 | 12.245 | 0.007 | 0.013 | 2.34258E-10 | -1.18687E-09 | -1.68320E-10 |
| | *N*-stearoyl valine | Equal variances assumed | 1.021 | 0.328 | -1.016 | 15 | 0.163 | 0.326 | 9.02140E-11 | -2.83985E-10 | 1.00588E-10 |
| | | Equal variances not assumed | | | -1.029 | 14.933 | 0.160 | 0.320 | 8.91403E-11 | -2.81771E-10 | 9.83739E-11 |
| | *N*-oleoyl valine | Equal variances assumed | 0.362 | 0.555 | -0.614 | 17 | 0.274 | 0.547 | 5.39830E-11 | -1.47036E-10 | 8.07524E-11 |
| | | Equal variances not assumed | | | -0.612 | 16.504 | 0.274 | 0.549 | 5.41697E-11 | -1.47692E-10 | 8.14085E-11 |
| | *N*-linoleoyl valine | Equal variances assumed | 0.991 | 0.338 | -1.641 | 13 | 0.062 | 0.125 | 1.71623E-10 | -6.52371E-10 | 8.91679E-11 |
| | | Equal variances not assumed | | | -1.795 | 12.999 | 0.048 | 0.096 | 1.56893E-10 | -6.20551E-10 | 5.73481E-11 |
| THC+CBD | *N*-palmitoyl valine | Equal variances assumed | 1.118 | 0.304 | -0.683 | 19 | 0.251 | 0.503 | 5.07069E-10 | -1.40782E-09 | 7.14798E-10 |
| | | Equal variances not assumed | | | -0.676 | 17.307 | 0.254 | 0.508 | 5.12560E-10 | -1.42646E-09 | 7.33437E-10 |
| | *N*-stearoyl valine | Equal variances assumed | 0.161 | 0.693 | -0.280 | 18 | 0.391 | 0.783 | 1.22378E-10 | -2.91334E-10 | 2.22879E-10 |
| | | Equal variances not assumed | | | -0.280 | 17.783 | 0.391 | 0.783 | 1.22378E-10 | -2.91559E-10 | 2.23104E-10 |
| | *N*-oleoyl valine | Equal variances assumed | 0.042 | 0.840 | -0.612 | 19 | 0.274 | 0.548 | 8.18122E-11 | -2.21339E-10 | 1.21130E-10 |
| | | Equal variances not assumed | | | -0.613 | 18.889 | 0.274 | 0.547 | 8.17155E-11 | -2.21205E-10 | 1.20996E-10 |
| | *N*-linoleoyl valine | Equal variances assumed | 2.076 | 0.167 | 0.210 | 18 | 0.418 | 0.836 | 1.50121E-10 | -2.83873E-10 | 3.46911E-10 |
| | | Equal variances not assumed | | | 0.210 | 15.440 | 0.418 | 0.836 | 1.50121E-10 | -2.87663E-10 | 3.50701E-10 |

Supplemental Table 8.4 Independent t-test comparing male and female levels of *N*-acyl valines measured in liver collected from adult mice 2h after an i.p. injection of 10mg/kg CBD, THC, or THC+CBD

| | | | Levene's Test for Equality of Variances | | t-test for Equality of Means | | | | | | | |
|---|---|---|---|---|---|---|---|---|---|---|---|---|
| | | | | | | | Significance | | | | 95% Confidence Interval of the Difference | |
| | | | F | Sig. | t | df | One-Sided p | Two-Sided p | Mean Difference | Std. Error Difference | Lower | Upper |
| Veh | *N*-palmitoyl valine | Equal variances assumed | 0.217 | 0.649 | -0.146 | 13 | 0.443 | 0.886 | -2.66941E-12 | 1.83257E-11 | -4.22597E-11 | 3.69209E-11 |
| | | Equal variances not assumed | | | -0.147 | 12.954 | 0.443 | 0.886 | -2.66941E-12 | 1.82151E-11 | -4.20348E-11 | 3.66960E-11 |
| | *N*-stearoyl valine | Equal variances assumed | 2.466 | 0.147 | -0.655 | 10 | 0.264 | 0.528 | -9.61190E-13 | 1.46843E-12 | -4.23306E-12 | 2.31068E-12 |
| | | Equal variances not assumed | | | -0.569 | 4.813 | 0.298 | 0.595 | -9.61190E-13 | 1.69032E-12 | -5.35752E-12 | 3.43514E-12 |
| | *N*-oleoyl valine | Equal variances assumed | 3.082 | 0.103 | 0.679 | 13 | 0.254 | 0.509 | 1.00480E-12 | 1.47923E-12 | -2.19089E-12 | 4.20048E-12 |
| | | Equal variances not assumed | | | 0.709 | 10.638 | 0.247 | 0.494 | 1.00480E-12 | 1.41784E-12 | -2.12886E-12 | 4.13845E-12 |
| | *N*-linoleoyl valine | Equal variances assumed | 2.984 | 0.108 | 1.574 | 13 | 0.070 | 0.139 | 8.15255E-12 | 5.17817E-12 | -3.03420E-12 | 1.93393E-11 |
| | | Equal variances not assumed | | | 1.637 | 11.033 | 0.065 | 0.130 | 8.15255E-12 | 4.97942E-12 | -2.80303E-12 | 1.91081E-11 |
| | *N*-docosahexaenoyl valine | Equal variances assumed | 1.067 | 0.322 | -1.465 | 12 | 0.084 | 0.169 | -3.10798E-12 | 2.12088E-12 | -7.72897E-12 | 1.51301E-12 |
| | | Equal variances not assumed | | | -1.465 | 8.997 | 0.088 | 0.177 | -3.10798E-12 | 2.12088E-12 | -7.90600E-12 | 1.69003E-12 |
| CBD | *N*-palmitoyl valine | Equal variances assumed | 12.016 | 0.003 | 2.298 | 15 | 0.018 | 0.036 | 5.85424E-11 | 2.54794E-11 | 4.23431E-12 | 1.12850E-10 |
| | | Equal variances not assumed | | | 2.167 | 7.452 | 0.032 | 0.065 | 5.85424E-11 | 2.70180E-11 | -4.56785E-12 | 1.21653E-10 |
| | *N*-stearoyl valine | Equal variances assumed | 18.307 | 0.002 | 2.097 | 10 | 0.031 | 0.062 | 3.54780E-12 | 1.69154E-12 | -2.21174E-13 | 7.31678E-12 |
| | | Equal variances not assumed | | | 1.745 | 4.073 | 0.077 | 0.155 | 3.54780E-12 | 2.03267E-12 | -2.05622E-12 | 9.15182E-12 |
| | *N*-oleoyl valine | Equal variances assumed | 2.491 | 0.137 | 2.527 | 14 | 0.012 | 0.024 | 2.39590E-12 | 9.48035E-13 | 3.62570E-13 | 4.42923E-12 |

| | | | | | | | | | | | | |
|---|---|---|---|---|---|---|---|---|---|---|---|---|
| | | Equal variances not assumed | | | 2.253 | 6.780 | 0.030 | 0.060 | 2.39590E-12 | 1.06330E-12 | -1.35070E-13 | 4.92687E-12 |
| | *N*-linoleoyl valine | Equal variances assumed | 13.443 | 0.002 | 2.153 | 15 | 0.024 | 0.048 | 2.20713E-11 | 1.02537E-11 | 2.16200E-13 | 4.39265E-11 |
| | | Equal variances not assumed | | | 2.025 | 7.196 | 0.041 | 0.081 | 2.20713E-11 | 1.08998E-11 | -3.56067E-12 | 4.77034E-11 |
| | *N*-docosahexaenoyl valine | Equal variances assumed | 14.445 | 0.002 | 1.973 | 15 | 0.034 | 0.067 | 8.50637E-12 | 4.31139E-12 | -6.83142E-13 | 1.76959E-11 |
| | | Equal variances not assumed | | | 1.857 | 7.231 | 0.052 | 0.104 | 8.50637E-12 | 4.58149E-12 | -2.25740E-12 | 1.92701E-11 |
| | *N*-palmitoyl valine | Equal variances assumed | 0.749 | 0.399 | -1.410 | 17 | 0.088 | 0.177 | -2.69920E-11 | 1.91433E-11 | -6.73808E-11 | 1.33967E-11 |
| | | Equal variances not assumed | | | -1.382 | 14.000 | 0.094 | 0.189 | -2.69920E-11 | 1.95298E-11 | -6.88791E-11 | 1.48951E-11 |
| | *N*-stearoyl valine | Equal variances assumed | 2.735 | 0.118 | -0.030 | 16 | 0.488 | 0.976 | -2.57071E-14 | 8.48505E-13 | -1.82446E-12 | 1.77304E-12 |
| | | Equal variances not assumed | | | -0.030 | 13.025 | 0.488 | 0.976 | -2.57071E-14 | 8.48505E-13 | -1.85843E-12 | 1.80702E-12 |
| | *N*-oleoyl valine | Equal variances assumed | 7.752 | 0.013 | 1.301 | 16 | 0.106 | 0.212 | 2.91043E-12 | 2.23765E-12 | -1.83319E-12 | 7.65404E-12 |
| THC | | Equal variances not assumed | | | 1.403 | 13.280 | 0.092 | 0.184 | 2.91043E-12 | 2.07422E-12 | -1.56107E-12 | 7.38193E-12 |
| | *N*-linoleoyl valine | Equal variances assumed | 0.113 | 0.741 | -1.099 | 17 | 0.144 | 0.287 | -5.11856E-12 | 4.65758E-12 | -1.49452E-11 | 4.70808E-12 |
| | | Equal variances not assumed | | | -1.093 | 16.248 | 0.145 | 0.291 | -5.11856E-12 | 4.68472E-12 | -1.50374E-11 | 4.80032E-12 |
| | *N*-docosahexaenoyl valine | Equal variances assumed | 0.549 | 0.469 | -0.568 | 17 | 0.289 | 0.578 | -2.05818E-12 | 3.62484E-12 | -9.70593E-12 | 5.58957E-12 |
| | | Equal variances not assumed | | | -0.560 | 14.946 | 0.292 | 0.584 | -2.05818E-12 | 3.67839E-12 | -9.90095E-12 | 5.78459E-12 |
| | *N*-palmitoyl valine | Equal variances assumed | 2.941 | 0.104 | 5.821 | 18 | 0.000 | 0.000 | 7.75241E-11 | 1.33188E-11 | 4.95422E-11 | 1.05506E-10 |
| THC+CBD | | Equal variances not assumed | | | 5.821 | 13.552 | 0.000 | 0.000 | 7.75241E-11 | 1.33188E-11 | 4.88691E-11 | 1.06179E-10 |
| | *N*-stearoyl valine | Equal variances assumed | 0.277 | 0.606 | 2.732 | 17 | 0.007 | 0.014 | 1.64082E-12 | 6.00610E-13 | 3.73647E-13 | 2.90800E-12 |
| | | Equal variances | | | 2.755 | 16.971 | 0.007 | 0.014 | 1.64082E-12 | 5.95550E-13 | 3.84158E-13 | 2.89749E-12 |

| | | | | | | | | | | | |
|---|---|---|---|---|---|---|---|---|---|---|---|
| | not assumed | | | | | | | | | | |
| *N*-oleoyl valine | Equal variances assumed | 7.181 | 0.015 | 5.917 | 18 | 0.000 | 0.000 | 7.23266E-12 | 1.22242E-12 | 4.66445E-12 | 9.80088E-12 |
| | Equal variances not assumed | | | 5.917 | 13.394 | 0.000 | 0.000 | 7.23266E-12 | 1.22242E-12 | 4.59966E-12 | 9.86567E-12 |
| *N*-linoleoyl valine | Equal variances assumed | 2.904 | 0.106 | 5.795 | 18 | 0.000 | 0.000 | 3.11034E-11 | 5.36694E-12 | 1.98279E-11 | 4.23789E-11 |
| | Equal variances not assumed | | | 5.795 | 12.932 | 0.000 | 0.000 | 3.11034E-11 | 5.36694E-12 | 1.95026E-11 | 4.27041E-11 |
| *N*-docosahexaenoyl valine | Equal variances assumed | 2.053 | 0.169 | 2.946 | 18 | 0.004 | 0.009 | 1.02932E-11 | 3.49377E-12 | 2.95308E-12 | 1.76333E-11 |
| | Equal variances not assumed | | | 2.946 | 15.631 | 0.005 | 0.010 | 1.02932E-11 | 3.49377E-12 | 2.87252E-12 | 1.77139E-11 |

Supplemental Table 8.5 One-way ANOVA for *N*-acyl valines measured in plasma collected from adult mice 2h after an i.p. injection of 10mg/kg CBD, THC, or THC+CBD

| | | Sum of Squares | df | Mean Square | F | Sig. |
|---|---|---|---|---|---|---|
| *N*-palmitoyl valine | Between Groups | 1.31E-17 | 7 | 1.87E-18 | 3.019 | 0.0083405 |
| | Within Groups | 3.96E-17 | 64 | 6.19E-19 | | |
| | Total | 5.27E-17 | 71 | | | |
| *N*-stearoyl valine | Between Groups | 6.37E-19 | 7 | 9.10E-20 | 1.286 | 0.2725814 |
| | Within Groups | 4.32E-18 | 61 | 7.08E-20 | | |
| | Total | 4.96E-18 | 68 | | | |
| *N*-oleoyl valine | Between Groups | 2.63E-19 | 7 | 3.75E-20 | 1.752 | 0.1126974 |
| | Within Groups | 1.37E-18 | 64 | 2.14E-20 | | |
| | Total | 1.63E-18 | 71 | | | |
| *N*-linoleoyl valine | Between Groups | 1.23E-18 | 7 | 1.75E-19 | 1.705 | 0.1250120 |
| | Within Groups | 6.17E-18 | 60 | 1.03E-19 | | |
| | Total | 7.40E-18 | 67 | | | |

**Supplemental Table 8.6 One-way ANOVA for *N*-acyl valines measured in liver collected from adult mice 2h after an i.p. injection of 10mg/kg CBD, THC, or THC+CBD**

| | | Sum of Squares | df | Mean Square | F | Sig. |
|---|---|---|---|---|---|---|
| *N*-palmitoyl valine | Between Groups | 6.12E-20 | 7 | 8.74E-21 | 5.344 | 0.00008 |
| | Within Groups | 1.03E-19 | 63 | 1.64E-21 | | |
| | Total | 1.64E-19 | 70 | | | |
| *N*-stearoyl valine | Between Groups | 5.63E-23 | 7 | 8.04E-24 | 1.874 | 0.09236 |
| | Within Groups | 2.27E-22 | 53 | 4.29E-24 | | |
| | Total | 2.83E-22 | 60 | | | |
| *N*-oleoyl valine | Between Groups | 5.78E-22 | 7 | 8.26E-23 | 7.799 | 0.00000 |
| | Within Groups | 6.46E-22 | 61 | 1.06E-23 | | |
| | Total | 1.22E-21 | 68 | | | |
| *N*-linoleoyl valine | Between Groups | 9.18E-21 | 7 | 1.31E-21 | 6.709 | 0.00001 |
| | Within Groups | 1.23E-20 | 63 | 1.96E-22 | | |
| | Total | 2.15E-20 | 70 | | | |
| *N*-docosahexaenoyl valine | Between Groups | 1.74E-21 | 7 | 2.49E-22 | 4.373 | 0.00054 |
| | Within Groups | 3.53E-21 | 62 | 5.69E-23 | | |
| | Total | 5.27E-21 | 69 | | | |

#### Supplemental Table 8.7 Fisher's LSD post-hoc test for plasma *N*-acyl valine ANOVA

| Dependent Variable | | | Std. Error | Sig. | 95% Confidence Interval Lower Bound | Upper Bound |
|---|---|---|---|---|---|---|
| *N*-palmitoyl valine | VehFemale | CBDfemale | 3.93E-10 | 0.0492 | 2.89E-12 | 1.57E-09 |
| | | THCfemale | 3.82E-10 | 0.0116 | 2.29E-10 | 1.76E-09 |
| | | TpCfemale | 3.66E-10 | 0.4205 | -4.34E-10 | 1.03E-09 |
| | | VehMale | 4.07E-10 | 0.0230 | 1.35E-10 | 1.76E-09 |
| | | CBDmale | 3.73E-10 | 0.0062 | 3.10E-10 | 1.80E-09 |
| | | THCmale | 3.82E-10 | 0.4124 | -4.48E-10 | 1.08E-09 |
| | | TpCmale | 3.73E-10 | 0.8936 | -7.96E-10 | 6.95E-10 |
| | CBDfemale | VehFemale | 3.93E-10 | 0.0492 | -1.57E-09 | -2.89E-12 |
| | | THCfemale | 3.82E-10 | 0.5948 | -5.59E-10 | 9.68E-10 |
| | | TpCfemale | 3.66E-10 | 0.1828 | -1.22E-09 | 2.38E-10 |
| | | VehMale | 4.07E-10 | 0.6955 | -6.53E-10 | 9.73E-10 |
| | | CBDmale | 3.73E-10 | 0.4773 | -4.79E-10 | 1.01E-09 |
| | | THCmale | 3.82E-10 | 0.2202 | -1.24E-09 | 2.90E-10 |
| | | TpCmale | 3.73E-10 | 0.0280 | -1.58E-09 | -9.33E-11 |
| | THCfemale | VehFemale | 3.82E-10 | 0.0116 | -1.76E-09 | -2.29E-10 |
| | | CBDfemale | 3.82E-10 | 0.5948 | -9.68E-10 | 5.59E-10 |
| | | TpCfemale | 3.54E-10 | 0.0532 | -1.40E-09 | 9.77E-12 |
| | | VehMale | 3.96E-10 | 0.9115 | -8.36E-10 | 7.48E-10 |
| | | CBDmale | 3.61E-10 | 0.8634 | -6.60E-10 | 7.84E-10 |
| | | THCmale | 3.71E-10 | 0.0723 | -1.42E-09 | 6.32E-11 |
| | | TpCmale | 3.61E-10 | 0.0053 | -1.77E-09 | -3.21E-10 |
| | TpCfemale | VehFemale | 3.66E-10 | 0.4205 | -1.03E-09 | 4.34E-10 |
| | | CBDfemale | 3.66E-10 | 0.1828 | -2.38E-10 | 1.22E-09 |
| | | THCfemale | 3.54E-10 | 0.0532 | -9.77E-12 | 1.40E-09 |
| | | VehMale | 3.80E-10 | 0.0912 | -1.08E-10 | 1.41E-09 |
| | | CBDmale | 3.44E-10 | 0.0308 | 7.23E-11 | 1.45E-09 |
| | | THCmale | 3.54E-10 | 0.9574 | -6.87E-10 | 7.25E-10 |
| | | TpCmale | 3.44E-10 | 0.3172 | -1.03E-09 | 3.40E-10 |
| | VehMale | VehFemale | 4.07E-10 | 0.0230 | -1.76E-09 | -1.35E-10 |
| | | CBDfemale | 4.07E-10 | 0.6955 | -9.73E-10 | 6.53E-10 |
| | | THCfemale | 3.96E-10 | 0.9115 | -7.48E-10 | 8.36E-10 |
| | | TpCfemale | 3.80E-10 | 0.0912 | -1.41E-09 | 1.08E-10 |
| | | CBDmale | 3.88E-10 | 0.7841 | -6.68E-10 | 8.81E-10 |
| | | THCmale | 3.96E-10 | 0.1151 | -1.43E-09 | 1.59E-10 |
| | | TpCmale | 3.88E-10 | 0.0123 | -1.77E-09 | -2.24E-10 |
| | CBDmale | VehFemale | 3.73E-10 | 0.0062 | -1.80E-09 | -3.10E-10 |
| | | CBDfemale | 3.73E-10 | 0.4773 | -1.01E-09 | 4.79E-10 |
| | | THCfemale | 3.61E-10 | 0.8634 | -7.84E-10 | 6.60E-10 |
| | | TpCfemale | 3.44E-10 | 0.0308 | -1.45E-09 | -7.23E-11 |
| | | VehMale | 3.88E-10 | 0.7841 | -8.81E-10 | 6.68E-10 |
| | | THCmale | 3.61E-10 | 0.0447 | -1.46E-09 | -1.80E-11 |
| | | TpCmale | 3.52E-10 | 0.0025 | -1.81E-09 | -4.03E-10 |
| | THCmale | VehFemale | 3.82E-10 | 0.4124 | -1.08E-09 | 4.48E-10 |
| | | CBDfemale | 3.82E-10 | 0.2202 | -2.90E-10 | 1.24E-09 |
| | | THCfemale | 3.71E-10 | 0.0723 | -6.32E-11 | 1.42E-09 |
| | | TpCfemale | 3.54E-10 | 0.9574 | -7.25E-10 | 6.87E-10 |
| | | VehMale | 3.96E-10 | 0.1151 | -1.59E-10 | 1.43E-09 |
| | | CBDmale | 3.61E-10 | 0.0447 | 1.80E-11 | 1.46E-09 |
| | | TpCmale | 3.61E-10 | 0.3157 | -1.09E-09 | 3.57E-10 |
| | TpCmale | VehFemale | 3.73E-10 | 0.8936 | -6.95E-10 | 7.96E-10 |
| | | CBDfemale | 3.73E-10 | 0.0280 | 9.33E-11 | 1.58E-09 |
| | | THCfemale | 3.61E-10 | 0.0053 | 3.21E-10 | 1.77E-09 |
| | | TpCfemale | 3.44E-10 | 0.3172 | -3.40E-10 | 1.03E-09 |
| | | VehMale | 3.88E-10 | 0.0123 | 2.24E-10 | 1.77E-09 |
| | | CBDmale | 3.52E-10 | 0.0025 | 4.03E-10 | 1.81E-09 |
| | | THCmale | 3.61E-10 | 0.3157 | -3.57E-10 | 1.09E-09 |
| *N*-stearoyl valine | VehFemale | CBDfemale | 1.33E-10 | 0.0364 | 1.86E-11 | 5.51E-10 |

| | | | | | | |
|---|---|---|---|---|---|---|
| | | THCfemale | 1.33E-10 | 0.0736 | -2.38E-11 | 5.08E-10 |
| | | TpCfemale | 1.26E-10 | 0.2989 | -1.20E-10 | 3.85E-10 |
| | | VehMale | 1.38E-10 | 0.5316 | -1.89E-10 | 3.62E-10 |
| | | CBDmale | 1.29E-10 | 0.0249 | 3.87E-11 | 5.56E-10 |
| | | THCmale | 1.29E-10 | 0.2489 | -1.08E-10 | 4.09E-10 |
| | | TpCmale | 1.26E-10 | 0.4404 | -1.54E-10 | 3.50E-10 |
| | CBDfemale | VehFemale | 1.33E-10 | 0.0364 | -5.51E-10 | -1.86E-11 |
| | | THCfemale | 1.33E-10 | 0.7507 | -3.08E-10 | 2.24E-10 |
| | | TpCfemale | 1.26E-10 | 0.2318 | -4.05E-10 | 9.99E-11 |
| | | VehMale | 1.38E-10 | 0.1555 | -4.73E-10 | 7.73E-11 |
| | | CBDmale | 1.29E-10 | 0.9229 | -2.46E-10 | 2.71E-10 |
| | | THCmale | 1.29E-10 | 0.3035 | -3.93E-10 | 1.24E-10 |
| | | TpCmale | 1.26E-10 | 0.1443 | -4.39E-10 | 6.57E-11 |
| | THCfemale | VehFemale | 1.33E-10 | 0.0736 | -5.08E-10 | 2.38E-11 |
| | | CBDfemale | 1.33E-10 | 0.7507 | -2.24E-10 | 3.08E-10 |
| | | TpCfemale | 1.26E-10 | 0.3870 | -3.62E-10 | 1.42E-10 |
| | | VehMale | 1.38E-10 | 0.2630 | -4.31E-10 | 1.20E-10 |
| | | CBDmale | 1.29E-10 | 0.6719 | -2.03E-10 | 3.14E-10 |
| | | THCmale | 1.29E-10 | 0.4809 | -3.50E-10 | 1.67E-10 |
| | | TpCmale | 1.26E-10 | 0.2577 | -3.97E-10 | 1.08E-10 |
| | TpCfemale | VehFemale | 1.26E-10 | 0.2989 | -3.85E-10 | 1.20E-10 |
| | | CBDfemale | 1.26E-10 | 0.2318 | -9.99E-11 | 4.05E-10 |
| | | THCfemale | 1.26E-10 | 0.3870 | -1.42E-10 | 3.62E-10 |
| | | VehMale | 1.31E-10 | 0.7292 | -3.08E-10 | 2.17E-10 |
| | | CBDmale | 1.22E-10 | 0.1821 | -7.95E-11 | 4.09E-10 |
| | | THCmale | 1.22E-10 | 0.8817 | -2.26E-10 | 2.63E-10 |
| | | TpCmale | 1.19E-10 | 0.7746 | -2.72E-10 | 2.04E-10 |
| | VehMale | VehFemale | 1.38E-10 | 0.5316 | -3.62E-10 | 1.89E-10 |
| | | CBDfemale | 1.38E-10 | 0.1555 | -7.73E-11 | 4.73E-10 |
| | | THCfemale | 1.38E-10 | 0.2630 | -1.20E-10 | 4.31E-10 |
| | | TpCfemale | 1.31E-10 | 0.7292 | -2.17E-10 | 3.08E-10 |
| | | CBDmale | 1.34E-10 | 0.1215 | -5.75E-11 | 4.79E-10 |
| | | THCmale | 1.34E-10 | 0.6356 | -2.04E-10 | 3.32E-10 |
| | | TpCmale | 1.31E-10 | 0.9312 | -2.51E-10 | 2.74E-10 |
| | CBDmale | VehFemale | 1.29E-10 | 0.0249 | -5.56E-10 | -3.87E-11 |
| | | CBDfemale | 1.29E-10 | 0.9229 | -2.71E-10 | 2.46E-10 |
| | | THCfemale | 1.29E-10 | 0.6719 | -3.14E-10 | 2.03E-10 |
| | | TpCfemale | 1.22E-10 | 0.1821 | -4.09E-10 | 7.95E-11 |
| | | VehMale | 1.34E-10 | 0.1215 | -4.79E-10 | 5.75E-11 |
| | | THCmale | 1.25E-10 | 0.2466 | -3.98E-10 | 1.04E-10 |
| | | TpCmale | 1.22E-10 | 0.1083 | -4.44E-10 | 4.52E-11 |
| | THCmale | VehFemale | 1.29E-10 | 0.2489 | -4.09E-10 | 1.08E-10 |
| | | CBDfemale | 1.29E-10 | 0.3035 | -1.24E-10 | 3.93E-10 |
| | | THCfemale | 1.29E-10 | 0.4809 | -1.67E-10 | 3.50E-10 |
| | | TpCfemale | 1.22E-10 | 0.8817 | -2.63E-10 | 2.26E-10 |
| | | VehMale | 1.34E-10 | 0.6356 | -3.32E-10 | 2.04E-10 |
| | | CBDmale | 1.25E-10 | 0.2466 | -1.04E-10 | 3.98E-10 |
| | | TpCmale | 1.22E-10 | 0.6692 | -2.97E-10 | 1.92E-10 |
| | TpCmale | VehFemale | 1.26E-10 | 0.4404 | -3.50E-10 | 1.54E-10 |
| | | CBDfemale | 1.26E-10 | 0.1443 | -6.57E-11 | 4.39E-10 |
| | | THCfemale | 1.26E-10 | 0.2577 | -1.08E-10 | 3.97E-10 |
| | | TpCfemale | 1.19E-10 | 0.7746 | -2.04E-10 | 2.72E-10 |
| | | VehMale | 1.31E-10 | 0.9312 | -2.74E-10 | 2.51E-10 |
| | | CBDmale | 1.22E-10 | 0.1083 | -4.52E-11 | 4.44E-10 |
| | | THCmale | 1.22E-10 | 0.6692 | -1.92E-10 | 2.97E-10 |
| *N*-oleoyl valine | VehFemale | CBDfemale | 7.32E-11 | 0.1200 | -3.09E-11 | 2.61E-10 |
| | | THCfemale | 6.94E-11 | 0.1135 | -2.73E-11 | 2.50E-10 |
| | | TpCfemale | 6.80E-11 | 0.7219 | -1.11E-10 | 1.60E-10 |
| | | VehMale | 7.57E-11 | 0.2463 | -6.27E-11 | 2.40E-10 |
| | | CBDmale | 7.11E-11 | 0.0279 | 1.79E-11 | 3.02E-10 |

| | | | | | | |
|---|---|---|---|---|---|---|
| | | THCmale | 7.11E-11 | 0.2753 | -6.38E-11 | 2.20E-10 |
| | | TpCmale | 6.94E-11 | 0.7113 | -1.64E-10 | 1.13E-10 |
| | CBDfemale | VehFemale | 7.32E-11 | 0.1200 | -2.61E-10 | 3.09E-11 |
| | | THCfemale | 6.94E-11 | 0.9554 | -1.43E-10 | 1.35E-10 |
| | | TpCfemale | 6.80E-11 | 0.1856 | -2.27E-10 | 4.49E-11 |
| | | VehMale | 7.57E-11 | 0.7260 | -1.78E-10 | 1.25E-10 |
| | | CBDmale | 7.11E-11 | 0.5319 | -9.73E-11 | 1.87E-10 |
| | | THCmale | 7.11E-11 | 0.6042 | -1.79E-10 | 1.05E-10 |
| | | TpCmale | 6.94E-11 | 0.0462 | -2.80E-10 | -2.42E-12 |
| | THCfemale | VehFemale | 6.94E-11 | 0.1135 | -2.50E-10 | 2.73E-11 |
| | | CBDfemale | 6.94E-11 | 0.9554 | -1.35E-10 | 1.43E-10 |
| | | TpCfemale | 6.39E-11 | 0.1780 | -2.15E-10 | 4.06E-11 |
| | | VehMale | 7.21E-11 | 0.7533 | -1.67E-10 | 1.21E-10 |
| | | CBDmale | 6.72E-11 | 0.4725 | -8.57E-11 | 1.83E-10 |
| | | THCmale | 6.72E-11 | 0.6237 | -1.67E-10 | 1.01E-10 |
| | | TpCmale | 6.54E-11 | 0.0400 | -2.68E-10 | -6.46E-12 |
| | TpCfemale | VehFemale | 6.80E-11 | 0.7219 | -1.60E-10 | 1.11E-10 |
| | | CBDfemale | 6.80E-11 | 0.1856 | -4.49E-11 | 2.27E-10 |
| | | THCfemale | 6.39E-11 | 0.1780 | -4.06E-11 | 2.15E-10 |
| | | VehMale | 7.07E-11 | 0.3668 | -7.70E-11 | 2.06E-10 |
| | | CBDmale | 6.58E-11 | 0.0432 | 4.28E-12 | 2.67E-10 |
| | | THCmale | 6.58E-11 | 0.4153 | -7.74E-11 | 1.85E-10 |
| | | TpCmale | 6.39E-11 | 0.4360 | -1.78E-10 | 7.76E-11 |
| | VehMale | VehFemale | 7.57E-11 | 0.2463 | -2.40E-10 | 6.27E-11 |
| | | CBDfemale | 7.57E-11 | 0.7260 | -1.25E-10 | 1.78E-10 |
| | | THCfemale | 7.21E-11 | 0.7533 | -1.21E-10 | 1.67E-10 |
| | | TpCfemale | 7.07E-11 | 0.3668 | -2.06E-10 | 7.70E-11 |
| | | CBDmale | 7.37E-11 | 0.3369 | -7.59E-11 | 2.19E-10 |
| | | THCmale | 7.37E-11 | 0.8885 | -1.58E-10 | 1.37E-10 |
| | | TpCmale | 7.21E-11 | 0.1175 | -2.58E-10 | 2.96E-11 |
| | CBDmale | VehFemale | 7.11E-11 | 0.0279 | -3.02E-10 | -1.79E-11 |
| | | CBDfemale | 7.11E-11 | 0.5319 | -1.87E-10 | 9.73E-11 |
| | | THCfemale | 6.72E-11 | 0.4725 | -1.83E-10 | 8.57E-11 |
| | | TpCfemale | 6.58E-11 | 0.0432 | -2.67E-10 | -4.28E-12 |
| | | VehMale | 7.37E-11 | 0.3369 | -2.19E-10 | 7.59E-11 |
| | | THCmale | 6.90E-11 | 0.2404 | -2.19E-10 | 5.61E-11 |
| | | TpCmale | 6.72E-11 | 0.0075 | -3.20E-10 | -5.15E-11 |
| | THCmale | VehFemale | 7.11E-11 | 0.2753 | -2.20E-10 | 6.38E-11 |
| | | CBDfemale | 7.11E-11 | 0.6042 | -1.05E-10 | 1.79E-10 |
| | | THCfemale | 6.72E-11 | 0.6237 | -1.01E-10 | 1.67E-10 |
| | | TpCfemale | 6.58E-11 | 0.4153 | -1.85E-10 | 7.74E-11 |
| | | VehMale | 7.37E-11 | 0.8885 | -1.37E-10 | 1.58E-10 |
| | | CBDmale | 6.90E-11 | 0.2404 | -5.61E-11 | 2.19E-10 |
| | | TpCmale | 6.72E-11 | 0.1267 | -2.38E-10 | 3.03E-11 |
| | TpCmale | VehFemale | 6.94E-11 | 0.7113 | -1.13E-10 | 1.64E-10 |
| | | CBDfemale | 6.94E-11 | 0.0462 | 2.42E-12 | 2.80E-10 |
| | | THCfemale | 6.54E-11 | 0.0400 | 6.46E-12 | 2.68E-10 |
| | | TpCfemale | 6.39E-11 | 0.4360 | -7.76E-11 | 1.78E-10 |
| | | VehMale | 7.21E-11 | 0.1175 | -2.96E-11 | 2.58E-10 |
| | | CBDmale | 6.72E-11 | 0.0075 | 5.15E-11 | 3.20E-10 |
| | | THCmale | 6.72E-11 | 0.1267 | -3.03E-11 | 2.38E-10 |
| *N*-linoleoyl valine | VehFemale | CBDfemale | 1.60E-10 | 0.0629 | -1.69E-11 | 6.25E-10 |
| | | THCfemale | 1.73E-10 | 0.0101 | 1.13E-10 | 8.06E-10 |
| | | TpCfemale | 1.52E-10 | 0.1824 | -9.91E-11 | 5.10E-10 |
| | | VehMale | 1.66E-10 | 0.0398 | 1.68E-11 | 6.81E-10 |
| | | CBDmale | 1.52E-10 | 0.0066 | 1.24E-10 | 7.32E-10 |
| | | THCmale | 1.56E-10 | 0.2571 | -1.33E-10 | 4.90E-10 |
| | | TpCmale | 1.52E-10 | 0.1249 | -6.75E-11 | 5.41E-10 |
| | CBDfemale | VehFemale | 1.60E-10 | 0.0629 | -6.25E-10 | 1.69E-11 |
| | | THCfemale | 1.73E-10 | 0.3711 | -1.90E-10 | 5.03E-10 |

| | | | | | |
|---|---|---|---|---|---|
| | TpCfemale | 1.52E-10 | 0.5194 | -4.03E-10 | 2.06E-10 |
| | VehMale | 1.66E-10 | 0.7874 | -2.87E-10 | 3.77E-10 |
| | CBDmale | 1.52E-10 | 0.4180 | -1.80E-10 | 4.28E-10 |
| | THCmale | 1.56E-10 | 0.4237 | -4.37E-10 | 1.86E-10 |
| | TpCmale | 1.52E-10 | 0.6609 | -3.71E-10 | 2.37E-10 |
| THCfemale | VehFemale | 1.73E-10 | 0.0101 | -8.06E-10 | -1.13E-10 |
| | CBDfemale | 1.73E-10 | 0.3711 | -5.03E-10 | 1.90E-10 |
| | TpCfemale | 1.66E-10 | 0.1294 | -5.86E-10 | 7.66E-11 |
| | VehMale | 1.78E-10 | 0.5358 | -4.68E-10 | 2.46E-10 |
| | CBDmale | 1.66E-10 | 0.8473 | -3.63E-10 | 2.99E-10 |
| | THCmale | 1.69E-10 | 0.1009 | -6.20E-10 | 5.65E-11 |
| | TpCmale | 1.66E-10 | 0.1829 | -5.54E-10 | 1.08E-10 |
| TpCfemale | VehFemale | 1.52E-10 | 0.1824 | -5.10E-10 | 9.91E-11 |
| | CBDfemale | 1.52E-10 | 0.5194 | -2.06E-10 | 4.03E-10 |
| | THCfemale | 1.66E-10 | 0.1294 | -7.66E-11 | 5.86E-10 |
| | VehMale | 1.58E-10 | 0.3673 | -1.73E-10 | 4.60E-10 |
| | CBDmale | 1.43E-10 | 0.1258 | -6.42E-11 | 5.10E-10 |
| | THCmale | 1.47E-10 | 0.8556 | -3.22E-10 | 2.68E-10 |
| | TpCmale | 1.43E-10 | 0.8268 | -2.55E-10 | 3.18E-10 |
| VehMale | VehFemale | 1.66E-10 | 0.0398 | -6.81E-10 | -1.68E-11 |
| | CBDfemale | 1.66E-10 | 0.7874 | -3.77E-10 | 2.87E-10 |
| | THCfemale | 1.78E-10 | 0.5358 | -2.46E-10 | 4.68E-10 |
| | TpCfemale | 1.58E-10 | 0.3673 | -4.60E-10 | 1.73E-10 |
| | CBDmale | 1.58E-10 | 0.6187 | -2.37E-10 | 3.95E-10 |
| | THCmale | 1.62E-10 | 0.2957 | -4.94E-10 | 1.53E-10 |
| | TpCmale | 1.58E-10 | 0.4811 | -4.28E-10 | 2.04E-10 |
| CBDmale | VehFemale | 1.52E-10 | 0.0066 | -7.32E-10 | -1.24E-10 |
| | CBDfemale | 1.52E-10 | 0.4180 | -4.28E-10 | 1.80E-10 |
| | THCfemale | 1.66E-10 | 0.8473 | -2.99E-10 | 3.63E-10 |
| | TpCfemale | 1.43E-10 | 0.1258 | -5.10E-10 | 6.42E-11 |
| | VehMale | 1.58E-10 | 0.6187 | -3.95E-10 | 2.37E-10 |
| | THCmale | 1.47E-10 | 0.0955 | -5.44E-10 | 4.52E-11 |
| | TpCmale | 1.43E-10 | 0.1877 | -4.78E-10 | 9.58E-11 |
| THCmale | VehFemale | 1.56E-10 | 0.2571 | -4.90E-10 | 1.33E-10 |
| | CBDfemale | 1.56E-10 | 0.4237 | -1.86E-10 | 4.37E-10 |
| | THCfemale | 1.69E-10 | 0.1009 | -5.65E-11 | 6.20E-10 |
| | TpCfemale | 1.47E-10 | 0.8556 | -2.68E-10 | 3.22E-10 |
| | VehMale | 1.62E-10 | 0.2957 | -1.53E-10 | 4.94E-10 |
| | CBDmale | 1.47E-10 | 0.0955 | -4.52E-11 | 5.44E-10 |
| | TpCmale | 1.47E-10 | 0.6930 | -2.36E-10 | 3.53E-10 |
| TpCmale | VehFemale | 1.52E-10 | 0.1249 | -5.41E-10 | 6.75E-11 |
| | CBDfemale | 1.52E-10 | 0.6609 | -2.37E-10 | 3.71E-10 |
| | THCfemale | 1.66E-10 | 0.1829 | -1.08E-10 | 5.54E-10 |
| | TpCfemale | 1.43E-10 | 0.8268 | -3.18E-10 | 2.55E-10 |
| | VehMale | 1.58E-10 | 0.4811 | -2.04E-10 | 4.28E-10 |
| | CBDmale | 1.43E-10 | 0.1877 | -9.58E-11 | 4.78E-10 |
| | THCmale | 1.47E-10 | 0.6930 | -3.53E-10 | 2.36E-10 |

#### Supplemental Table 8.8 Fisher's LSD post-hoc test for liver *N*-acyl valine ANOVA

| Dependent Variable | | | Std. Error | Sig. | 95% Confidence Interval Lower Bound | 95% Confidence Interval Upper Bound |
|---|---|---|---|---|---|---|
| *N*-palmitoyl valine | VehFemale | CBDfemale | 2.02188E-11 | 0.120 | -7.2286E-11 | 8.5218E-12 |
| | | THCfemale | 1.91812E-11 | 0.320 | -5.7545E-11 | 1.9116E-11 |
| | | TpCfemale | 1.91812E-11 | 0.002 | -1.0061E-10 | -2.3944E-11 |
| | | VehMale | 2.09285E-11 | 0.899 | -4.4492E-11 | 3.9153E-11 |
| | | CBDmale | 1.96491E-11 | 0.180 | -1.2606E-11 | 6.5926E-11 |
| | | THCmale | 1.96491E-11 | 0.022 | -8.5472E-11 | -6.9407E-12 |
| | | TpCmale | 1.91812E-11 | 0.430 | -2.3081E-11 | 5.3580E-11 |
| | CBDfemale | VehFemale | 2.02188E-11 | 0.120 | -8.5218E-12 | 7.2286E-11 |
| | | THCfemale | 1.91812E-11 | 0.511 | -2.5663E-11 | 5.0999E-11 |
| | | TpCfemale | 1.91812E-11 | 0.118 | -6.8723E-11 | 7.9381E-12 |
| | | VehMale | 2.09285E-11 | 0.168 | -1.2609E-11 | 7.1035E-11 |
| | | CBDmale | 1.96491E-11 | 0.004 | 1.9277E-11 | 9.7808E-11 |
| | | THCmale | 1.96491E-11 | 0.469 | -5.3590E-11 | 2.4942E-11 |
| | | TpCmale | 1.91812E-11 | 0.017 | 8.8009E-12 | 8.5462E-11 |
| | THCfemale | VehFemale | 1.91812E-11 | 0.320 | -1.9116E-11 | 5.7545E-11 |
| | | CBDfemale | 1.91812E-11 | 0.511 | -5.0999E-11 | 2.5663E-11 |
| | | TpCfemale | 1.80842E-11 | 0.020 | -7.9199E-11 | -6.9220E-12 |
| | | VehMale | 1.99279E-11 | 0.410 | -2.3278E-11 | 5.6368E-11 |
| | | CBDmale | 1.85798E-11 | 0.016 | 8.7457E-12 | 8.3003E-11 |
| | | THCmale | 1.85798E-11 | 0.151 | -6.4121E-11 | 1.0137E-11 |
| | | TpCmale | 1.80842E-11 | 0.061 | -1.6749E-12 | 7.0602E-11 |
| | TpCfemale | VehFemale | 1.91812E-11 | 0.002 | 2.3944E-11 | 1.0061E-10 |
| | | CBDfemale | 1.91812E-11 | 0.118 | -7.9381E-12 | 6.8723E-11 |
| | | THCfemale | 1.80842E-11 | 0.020 | 6.9220E-12 | 7.9199E-11 |
| | | VehMale | 1.99279E-11 | 0.004 | 1.9783E-11 | 9.9428E-11 |
| | | CBDmale | 1.85798E-11 | 0.000 | 5.1806E-11 | 1.2606E-10 |
| | | THCmale | 1.85798E-11 | 0.390 | -2.1060E-11 | 5.3197E-11 |
| | | TpCmale | 1.80842E-11 | 0.000 | 4.1386E-11 | 1.1366E-10 |
| | VehMale | VehFemale | 2.09285E-11 | 0.899 | -3.9153E-11 | 4.4492E-11 |
| | | CBDfemale | 2.09285E-11 | 0.168 | -7.1035E-11 | 1.2609E-11 |
| | | THCfemale | 1.99279E-11 | 0.410 | -5.6368E-11 | 2.3278E-11 |
| | | TpCfemale | 1.99279E-11 | 0.004 | -9.9428E-11 | -1.9783E-11 |
| | | CBDmale | 2.03786E-11 | 0.155 | -1.1394E-11 | 7.0053E-11 |
| | | THCmale | 2.03786E-11 | 0.037 | -8.4260E-11 | -2.8135E-12 |
| | | TpCmale | 1.99279E-11 | 0.372 | -2.1904E-11 | 5.7741E-11 |
| | CBDmale | VehFemale | 1.96491E-11 | 0.180 | -6.5926E-11 | 1.2606E-11 |
| | | CBDfemale | 1.96491E-11 | 0.004 | -9.7808E-11 | -1.9277E-11 |
| | | THCfemale | 1.85798E-11 | 0.016 | -8.3003E-11 | -8.7457E-12 |
| | | TpCfemale | 1.85798E-11 | 0.000 | -1.2606E-10 | -5.1806E-11 |
| | | VehMale | 2.03786E-11 | 0.155 | -7.0053E-11 | 1.1394E-11 |
| | | THCmale | 1.90625E-11 | 0.000 | -1.1096E-10 | -3.4773E-11 |
| | | TpCmale | 1.85798E-11 | 0.541 | -4.8540E-11 | 2.5718E-11 |
| | THCmale | VehFemale | 1.96491E-11 | 0.022 | 6.9407E-12 | 8.5472E-11 |
| | | CBDfemale | 1.96491E-11 | 0.469 | -2.4942E-11 | 5.3590E-11 |
| | | THCfemale | 1.85798E-11 | 0.151 | -1.0137E-11 | 6.4121E-11 |
| | | TpCfemale | 1.85798E-11 | 0.390 | -5.3197E-11 | 2.1060E-11 |
| | | VehMale | 2.03786E-11 | 0.037 | 2.8135E-12 | 8.4260E-11 |
| | | CBDmale | 1.90625E-11 | 0.000 | 3.4773E-11 | 1.1096E-10 |
| | | TpCmale | 1.85798E-11 | 0.002 | 2.4327E-11 | 9.8584E-11 |
| | TpCmale | VehFemale | 1.91812E-11 | 0.430 | -5.3580E-11 | 2.3081E-11 |
| | | CBDfemale | 1.91812E-11 | 0.017 | -8.5462E-11 | -8.8009E-12 |
| | | THCfemale | 1.80842E-11 | 0.061 | -7.0602E-11 | 1.6749E-12 |
| | | TpCfemale | 1.80842E-11 | 0.000 | -1.1366E-10 | -4.1386E-11 |
| | | VehMale | 1.99279E-11 | 0.372 | -5.7741E-11 | 2.1904E-11 |
| | | CBDmale | 1.85798E-11 | 0.541 | -2.5718E-11 | 4.8540E-11 |
| | | THCmale | 1.85798E-11 | 0.002 | -9.8584E-11 | -2.4327E-11 |
| *N*-stearoyl valine | VehFemale | CBDfemale | 1.21242E-12 | 0.034 | -5.0766E-12 | -2.1296E-13 |

| | | | | | | |
|---|---|---|---|---|---|---|
| | | THCfemale | 1.04349E-12 | 0.309 | -3.1654E-12 | 1.0206E-12 |
| | | TpCfemale | 1.02040E-12 | 0.180 | -3.4322E-12 | 6.6113E-13 |
| | | VehMale | 1.21242E-12 | 0.431 | -3.3930E-12 | 1.4706E-12 |
| | | CBDmale | 1.10678E-12 | 0.418 | -1.3169E-12 | 3.1230E-12 |
| | | THCmale | 1.04349E-12 | 0.297 | -3.1911E-12 | 9.9485E-13 |
| | | TpCmale | 1.04349E-12 | 0.808 | -1.8377E-12 | 2.3482E-12 |
| | CBDfemale | VehFemale | 1.21242E-12 | 0.034 | 2.1296E-13 | 5.0766E-12 |
| | | THCfemale | 1.15493E-12 | 0.179 | -7.4413E-13 | 3.8888E-12 |
| | | TpCfemale | 1.13412E-12 | 0.272 | -1.0155E-12 | 3.5340E-12 |
| | | VehMale | 1.30956E-12 | 0.204 | -9.4307E-13 | 4.3102E-12 |
| | | CBDmale | 1.21242E-12 | 0.005 | 1.1160E-12 | 5.9796E-12 |
| | | THCmale | 1.15493E-12 | 0.186 | -7.6983E-13 | 3.8631E-12 |
| | | TpCmale | 1.15493E-12 | 0.015 | 5.8356E-13 | 5.2165E-12 |
| | THCfemale | VehFemale | 1.04349E-12 | 0.309 | -1.0206E-12 | 3.1654E-12 |
| | | CBDfemale | 1.15493E-12 | 0.179 | -3.8888E-12 | 7.4413E-13 |
| | | TpCfemale | 9.51376E-13 | 0.743 | -2.2214E-12 | 1.5951E-12 |
| | | VehMale | 1.15493E-12 | 0.924 | -2.2053E-12 | 2.4277E-12 |
| | | CBDmale | 1.04349E-12 | 0.064 | -1.1753E-13 | 4.0684E-12 |
| | | THCmale | 9.76091E-13 | 0.979 | -1.9835E-12 | 1.9321E-12 |
| | | TpCmale | 9.76091E-13 | 0.180 | -6.3010E-13 | 3.2855E-12 |
| | TpCfemale | VehFemale | 1.02040E-12 | 0.180 | -6.6113E-13 | 3.4322E-12 |
| | | CBDfemale | 1.13412E-12 | 0.272 | -3.5340E-12 | 1.0155E-12 |
| | | THCfemale | 9.51376E-13 | 0.743 | -1.5951E-12 | 2.2214E-12 |
| | | VehMale | 1.13412E-12 | 0.710 | -1.8504E-12 | 2.6991E-12 |
| | | CBDmale | 1.02040E-12 | 0.029 | 2.4191E-13 | 4.3352E-12 |
| | | THCmale | 9.51376E-13 | 0.764 | -1.6208E-12 | 2.1956E-12 |
| | | TpCmale | 9.51376E-13 | 0.090 | -2.6740E-13 | 3.5490E-12 |
| | VehMale | VehFemale | 1.21242E-12 | 0.431 | -1.4706E-12 | 3.3930E-12 |
| | | CBDfemale | 1.30956E-12 | 0.204 | -4.3102E-12 | 9.4307E-13 |
| | | THCfemale | 1.15493E-12 | 0.924 | -2.4277E-12 | 2.2053E-12 |
| | | TpCfemale | 1.13412E-12 | 0.710 | -2.6991E-12 | 1.8504E-12 |
| | | CBDmale | 1.21242E-12 | 0.130 | -5.6758E-13 | 4.2960E-12 |
| | | THCmale | 1.15493E-12 | 0.906 | -2.4534E-12 | 2.1796E-12 |
| | | TpCmale | 1.15493E-12 | 0.297 | -1.1000E-12 | 3.5330E-12 |
| | CBDmale | VehFemale | 1.10678E-12 | 0.418 | -3.1230E-12 | 1.3169E-12 |
| | | CBDfemale | 1.21242E-12 | 0.005 | -5.9796E-12 | -1.1160E-12 |
| | | THCfemale | 1.04349E-12 | 0.064 | -4.0684E-12 | 1.1753E-13 |
| | | TpCfemale | 1.02040E-12 | 0.029 | -4.3352E-12 | -2.4191E-13 |
| | | VehMale | 1.21242E-12 | 0.130 | -4.2960E-12 | 5.6758E-13 |
| | | THCmale | 1.04349E-12 | 0.061 | -4.0941E-12 | 9.1818E-14 |
| | | TpCmale | 1.04349E-12 | 0.537 | -2.7407E-12 | 1.4452E-12 |
| | THCmale | VehFemale | 1.04349E-12 | 0.297 | -9.9485E-13 | 3.1911E-12 |
| | | CBDfemale | 1.15493E-12 | 0.186 | -3.8631E-12 | 7.6983E-13 |
| | | THCfemale | 9.76091E-13 | 0.979 | -1.9321E-12 | 1.9835E-12 |
| | | TpCfemale | 9.51376E-13 | 0.764 | -2.1956E-12 | 1.6208E-12 |
| | | VehMale | 1.15493E-12 | 0.906 | -2.1796E-12 | 2.4534E-12 |
| | | CBDmale | 1.04349E-12 | 0.061 | -9.1818E-14 | 4.0941E-12 |
| | | TpCmale | 9.76091E-13 | 0.171 | -6.0440E-13 | 3.3112E-12 |
| | TpCmale | VehFemale | 1.04349E-12 | 0.808 | -2.3482E-12 | 1.8377E-12 |
| | | CBDfemale | 1.15493E-12 | 0.015 | -5.2165E-12 | -5.8356E-13 |
| | | THCfemale | 9.76091E-13 | 0.180 | -3.2855E-12 | 6.3010E-13 |
| | | TpCfemale | 9.51376E-13 | 0.090 | -3.5490E-12 | 2.6740E-13 |
| | | VehMale | 1.15493E-12 | 0.297 | -3.5330E-12 | 1.1000E-12 |
| | | CBDmale | 1.04349E-12 | 0.537 | -1.4452E-12 | 2.7407E-12 |
| | | THCmale | 9.76091E-13 | 0.171 | -3.3112E-12 | 6.0440E-13 |
| *N*-oleoyl valine | VehFemale | CBDfemale | 1.68461E-12 | 0.720 | -2.7623E-12 | 3.9749E-12 |
| | | THCfemale | 1.54397E-12 | 0.031 | -6.4964E-12 | -3.2170E-13 |
| | | TpCfemale | 1.54397E-12 | 0.000 | -9.4552E-12 | -3.2804E-12 |
| | | VehMale | 1.68461E-12 | 0.553 | -2.3638E-12 | 4.3734E-12 |
| | | CBDmale | 1.58164E-12 | 0.062 | -1.6045E-13 | 6.1649E-12 |

| | | | | | | |
|---|---|---|---|---|---|---|
| | | THCmale | 1.62749E-12 | 0.760 | -3.7530E-12 | 2.7557E-12 |
| | | TpCmale | 1.54397E-12 | 0.577 | -2.2225E-12 | 3.9522E-12 |
| | CBDfemale | VehFemale | 1.68461E-12 | 0.720 | -3.9749E-12 | 2.7623E-12 |
| | | THCfemale | 1.60407E-12 | 0.015 | -7.2229E-12 | -8.0786E-13 |
| | | TpCfemale | 1.60407E-12 | 0.000 | -1.0182E-11 | -3.7666E-12 |
| | | VehMale | 1.73986E-12 | 0.820 | -3.0806E-12 | 3.8775E-12 |
| | | CBDmale | 1.64036E-12 | 0.149 | -8.8420E-13 | 5.6760E-12 |
| | | THCmale | 1.68461E-12 | 0.514 | -4.4736E-12 | 2.2636E-12 |
| | | TpCmale | 1.60407E-12 | 0.872 | -2.9490E-12 | 3.4661E-12 |
| | THCfemale | VehFemale | 1.54397E-12 | 0.031 | 3.2170E-13 | 6.4964E-12 |
| | | CBDfemale | 1.60407E-12 | 0.015 | 8.0786E-13 | 7.2229E-12 |
| | | TpCfemale | 1.45567E-12 | 0.046 | -5.8695E-12 | -4.7919E-14 |
| | | VehMale | 1.60407E-12 | 0.008 | 1.2063E-12 | 7.6214E-12 |
| | | CBDmale | 1.49556E-12 | 0.000 | 3.4207E-12 | 9.4019E-12 |
| | | THCmale | 1.54397E-12 | 0.064 | -1.7694E-13 | 5.9978E-12 |
| | | TpCmale | 1.45567E-12 | 0.005 | 1.3631E-12 | 7.1847E-12 |
| | TpCfemale | VehFemale | 1.54397E-12 | 0.000 | 3.2804E-12 | 9.4552E-12 |
| | | CBDfemale | 1.60407E-12 | 0.000 | 3.7666E-12 | 1.0182E-11 |
| | | THCfemale | 1.45567E-12 | 0.046 | 4.7919E-14 | 5.8695E-12 |
| | | VehMale | 1.60407E-12 | 0.000 | 4.1650E-12 | 1.0580E-11 |
| | | CBDmale | 1.49556E-12 | 0.000 | 6.3795E-12 | 1.2361E-11 |
| | | THCmale | 1.54397E-12 | 0.000 | 2.7818E-12 | 8.9565E-12 |
| | | TpCmale | 1.45567E-12 | 0.000 | 4.3219E-12 | 1.0143E-11 |
| | VehMale | VehFemale | 1.68461E-12 | 0.553 | -4.3734E-12 | 2.3638E-12 |
| | | CBDfemale | 1.73986E-12 | 0.820 | -3.8775E-12 | 3.0806E-12 |
| | | THCfemale | 1.60407E-12 | 0.008 | -7.6214E-12 | -1.2063E-12 |
| | | TpCfemale | 1.60407E-12 | 0.000 | -1.0580E-11 | -4.1650E-12 |
| | | CBDmale | 1.64036E-12 | 0.228 | -1.2827E-12 | 5.2775E-12 |
| | | THCmale | 1.68461E-12 | 0.376 | -4.8720E-12 | 1.8652E-12 |
| | | TpCmale | 1.60407E-12 | 0.931 | -3.3475E-12 | 3.0676E-12 |
| | CBDmale | VehFemale | 1.58164E-12 | 0.062 | -6.1649E-12 | 1.6045E-13 |
| | | CBDfemale | 1.64036E-12 | 0.149 | -5.6760E-12 | 8.8420E-13 |
| | | THCfemale | 1.49556E-12 | 0.000 | -9.4019E-12 | -3.4207E-12 |
| | | TpCfemale | 1.49556E-12 | 0.000 | -1.2361E-11 | -6.3795E-12 |
| | | VehMale | 1.64036E-12 | 0.228 | -5.2775E-12 | 1.2827E-12 |
| | | THCmale | 1.58164E-12 | 0.031 | -6.6636E-12 | -3.3820E-13 |
| | | TpCmale | 1.49556E-12 | 0.158 | -5.1279E-12 | 8.5320E-13 |
| | THCmale | VehFemale | 1.62749E-12 | 0.760 | -2.7557E-12 | 3.7530E-12 |
| | | CBDfemale | 1.68461E-12 | 0.514 | -2.2636E-12 | 4.4736E-12 |
| | | THCfemale | 1.54397E-12 | 0.064 | -5.9978E-12 | 1.7694E-13 |
| | | TpCfemale | 1.54397E-12 | 0.000 | -8.9565E-12 | -2.7818E-12 |
| | | VehMale | 1.68461E-12 | 0.376 | -1.8652E-12 | 4.8720E-12 |
| | | CBDmale | 1.58164E-12 | 0.031 | 3.3820E-13 | 6.6636E-12 |
| | | TpCmale | 1.54397E-12 | 0.381 | -1.7238E-12 | 4.4509E-12 |
| | TpCmale | VehFemale | 1.54397E-12 | 0.577 | -3.9522E-12 | 2.2225E-12 |
| | | CBDfemale | 1.60407E-12 | 0.872 | -3.4661E-12 | 2.9490E-12 |
| | | THCfemale | 1.45567E-12 | 0.005 | -7.1847E-12 | -1.3631E-12 |
| | | TpCfemale | 1.45567E-12 | 0.000 | -1.0143E-11 | -4.3219E-12 |
| | | VehMale | 1.60407E-12 | 0.931 | -3.0676E-12 | 3.3475E-12 |
| | | CBDmale | 1.49556E-12 | 0.158 | -8.5320E-13 | 5.1279E-12 |
| | | THCmale | 1.54397E-12 | 0.381 | -4.4509E-12 | 1.7238E-12 |
| *N*-linoleoyl valine | VehFemale | CBDfemale | 6.99203E-12 | 0.120 | -2.4985E-11 | 2.9598E-12 |
| | | THCfemale | 6.63322E-12 | 0.751 | -1.5365E-11 | 1.1146E-11 |
| | | TpCfemale | 6.63322E-12 | 0.000 | -3.8672E-11 | -1.2161E-11 |
| | | VehMale | 7.23743E-12 | 0.264 | -6.3103E-12 | 2.2615E-11 |
| | | CBDmale | 6.79503E-12 | 0.109 | -2.5201E-12 | 2.4637E-11 |
| | | THCmale | 6.79503E-12 | 0.291 | -2.0807E-11 | 6.3504E-12 |
| | | TpCmale | 6.63322E-12 | 0.394 | -7.5684E-12 | 1.8943E-11 |
| | CBDfemale | VehFemale | 6.99203E-12 | 0.120 | -2.9598E-12 | 2.4985E-11 |
| | | THCfemale | 6.63322E-12 | 0.184 | -4.3526E-12 | 2.2158E-11 |

| | | | | | | |
|---|---|---|---|---|---|---|
| | | TpCfemale | 6.63322E-12 | 0.034 | -2.7659E-11 | -1.1483E-12 |
| | | VehMale | 7.23743E-12 | 0.010 | 4.7023E-12 | 3.3628E-11 |
| | | CBDmale | 6.79503E-12 | 0.002 | 8.4926E-12 | 3.5650E-11 |
| | | THCmale | 6.79503E-12 | 0.580 | -9.7945E-12 | 1.7363E-11 |
| | | TpCmale | 6.63322E-12 | 0.014 | 3.4443E-12 | 2.9955E-11 |
| | THCfemale | VehFemale | 6.63322E-12 | 0.751 | -1.1146E-11 | 1.5365E-11 |
| | | CBDfemale | 6.63322E-12 | 0.184 | -2.2158E-11 | 4.3526E-12 |
| | | TpCfemale | 6.25386E-12 | 0.000 | -3.5804E-11 | -1.0809E-11 |
| | | VehMale | 6.89142E-12 | 0.141 | -3.5091E-12 | 2.4034E-11 |
| | | CBDmale | 6.42523E-12 | 0.045 | 3.2869E-13 | 2.6008E-11 |
| | | THCmale | 6.42523E-12 | 0.429 | -1.7958E-11 | 7.7212E-12 |
| | | TpCmale | 6.25386E-12 | 0.217 | -4.7005E-12 | 2.0294E-11 |
| | TpCfemale | VehFemale | 6.63322E-12 | 0.000 | 1.2161E-11 | 3.8672E-11 |
| | | CBDfemale | 6.63322E-12 | 0.034 | 1.1483E-12 | 2.7659E-11 |
| | | THCfemale | 6.25386E-12 | 0.000 | 1.0809E-11 | 3.5804E-11 |
| | | VehMale | 6.89142E-12 | 0.000 | 1.9797E-11 | 4.7340E-11 |
| | | CBDmale | 6.42523E-12 | 0.000 | 2.3635E-11 | 4.9315E-11 |
| | | THCmale | 6.42523E-12 | 0.006 | 5.3482E-12 | 3.1028E-11 |
| | | TpCmale | 6.25386E-12 | 0.000 | 1.8606E-11 | 4.3601E-11 |
| | VehMale | VehFemale | 7.23743E-12 | 0.264 | -2.2615E-11 | 6.3103E-12 |
| | | CBDfemale | 7.23743E-12 | 0.010 | -3.3628E-11 | -4.7023E-12 |
| | | THCfemale | 6.89142E-12 | 0.141 | -2.4034E-11 | 3.5091E-12 |
| | | TpCfemale | 6.89142E-12 | 0.000 | -4.7340E-11 | -1.9797E-11 |
| | | CBDmale | 7.04730E-12 | 0.681 | -1.1177E-11 | 1.6989E-11 |
| | | THCmale | 7.04730E-12 | 0.033 | -2.9464E-11 | -1.2980E-12 |
| | | TpCmale | 6.89142E-12 | 0.722 | -1.6237E-11 | 1.1306E-11 |
| | CBDmale | VehFemale | 6.79503E-12 | 0.109 | -2.4637E-11 | 2.5201E-12 |
| | | CBDfemale | 6.79503E-12 | 0.002 | -3.5650E-11 | -8.4926E-12 |
| | | THCfemale | 6.42523E-12 | 0.045 | -2.6008E-11 | -3.2869E-13 |
| | | TpCfemale | 6.42523E-12 | 0.000 | -4.9315E-11 | -2.3635E-11 |
| | | VehMale | 7.04730E-12 | 0.681 | -1.6989E-11 | 1.1177E-11 |
| | | THCmale | 6.59215E-12 | 0.007 | -3.1460E-11 | -5.1137E-12 |
| | | TpCmale | 6.42523E-12 | 0.406 | -1.8211E-11 | 7.4682E-12 |
| | THCmale | VehFemale | 6.79503E-12 | 0.291 | -6.3504E-12 | 2.0807E-11 |
| | | CBDfemale | 6.79503E-12 | 0.580 | -1.7363E-11 | 9.7945E-12 |
| | | THCfemale | 6.42523E-12 | 0.429 | -7.7212E-12 | 1.7958E-11 |
| | | TpCfemale | 6.42523E-12 | 0.006 | -3.1028E-11 | -5.3482E-12 |
| | | VehMale | 7.04730E-12 | 0.033 | 1.2980E-12 | 2.9464E-11 |
| | | CBDmale | 6.59215E-12 | 0.007 | 5.1137E-12 | 3.1460E-11 |
| | | TpCmale | 6.42523E-12 | 0.049 | 7.5618E-14 | 2.5755E-11 |
| | TpCmale | VehFemale | 6.63322E-12 | 0.394 | -1.8943E-11 | 7.5684E-12 |
| | | CBDfemale | 6.63322E-12 | 0.014 | -2.9955E-11 | -3.4443E-12 |
| | | THCfemale | 6.25386E-12 | 0.217 | -2.0294E-11 | 4.7005E-12 |
| | | TpCfemale | 6.25386E-12 | 0.000 | -4.3601E-11 | -1.8606E-11 |
| | | VehMale | 6.89142E-12 | 0.722 | -1.1306E-11 | 1.6237E-11 |
| | | CBDmale | 6.42523E-12 | 0.406 | -7.4682E-12 | 1.8211E-11 |
| | | THCmale | 6.42523E-12 | 0.049 | -2.5755E-11 | -7.5618E-14 |
| *N*-docosahexaenoyl valine | VehFemale | CBDfemale | 3.90324E-12 | 0.040 | -1.6012E-11 | -4.0662E-13 |
| | | THCfemale | 3.71663E-12 | 0.017 | -1.6541E-11 | -1.6820E-12 |
| | | TpCfemale | 3.71663E-12 | 0.000 | -2.1642E-11 | -6.7829E-12 |
| | | VehMale | 4.03125E-12 | 0.444 | -1.1166E-11 | 4.9504E-12 |
| | | CBDmale | 3.80070E-12 | 0.938 | -7.3002E-12 | 7.8948E-12 |
| | | THCmale | 3.80070E-12 | 0.005 | -1.8767E-11 | -3.5722E-12 |
| | | TpCmale | 3.71663E-12 | 0.296 | -1.1349E-11 | 3.5103E-12 |
| | CBDfemale | VehFemale | 3.90324E-12 | 0.040 | 4.0662E-13 | 1.6012E-11 |
| | | THCfemale | 3.57738E-12 | 0.802 | -8.0534E-12 | 6.2487E-12 |
| | | TpCfemale | 3.57738E-12 | 0.098 | -1.3154E-11 | 1.1478E-12 |
| | | VehMale | 3.90324E-12 | 0.196 | -2.7014E-12 | 1.2904E-11 |
| | | CBDmale | 3.66465E-12 | 0.024 | 1.1808E-12 | 1.5832E-11 |
| | | THCmale | 3.66465E-12 | 0.422 | -1.0286E-11 | 4.3650E-12 |

| | | | | | |
|---|---|---|---|---|---|
| | TpCmale | 3.57738E-12 | 0.235 | -2.8611E-12 | 1.1441E-11 |
| THCfemale | VehFemale | 3.71663E-12 | 0.017 | 1.6820E-12 | 1.6541E-11 |
| | CBDfemale | 3.57738E-12 | 0.802 | -6.2487E-12 | 8.0534E-12 |
| | TpCfemale | 3.37279E-12 | 0.136 | -1.1843E-11 | 1.6412E-12 |
| | VehMale | 3.71663E-12 | 0.111 | -1.4260E-12 | 1.3433E-11 |
| | CBDmale | 3.46521E-12 | 0.009 | 2.4819E-12 | 1.6336E-11 |
| | THCmale | 3.46521E-12 | 0.555 | -8.9850E-12 | 4.8687E-12 |
| | TpCmale | 3.37279E-12 | 0.129 | -1.5498E-12 | 1.1934E-11 |
| TpCfemale | VehFemale | 3.71663E-12 | 0.000 | 6.7829E-12 | 2.1642E-11 |
| | CBDfemale | 3.57738E-12 | 0.098 | -1.1478E-12 | 1.3154E-11 |
| | THCfemale | 3.37279E-12 | 0.136 | -1.6412E-12 | 1.1843E-11 |
| | VehMale | 3.71663E-12 | 0.004 | 3.6749E-12 | 1.8534E-11 |
| | CBDmale | 3.46521E-12 | 0.000 | 7.5828E-12 | 2.1436E-11 |
| | THCmale | 3.46521E-12 | 0.383 | -3.8841E-12 | 9.9696E-12 |
| | TpCmale | 3.37279E-12 | 0.003 | 3.5511E-12 | 1.7035E-11 |
| VehMale | VehFemale | 4.03125E-12 | 0.444 | -4.9504E-12 | 1.1166E-11 |
| | CBDfemale | 3.90324E-12 | 0.196 | -1.2904E-11 | 2.7014E-12 |
| | THCfemale | 3.71663E-12 | 0.111 | -1.3433E-11 | 1.4260E-12 |
| | TpCfemale | 3.71663E-12 | 0.004 | -1.8534E-11 | -3.6749E-12 |
| | CBDmale | 3.80070E-12 | 0.374 | -4.1922E-12 | 1.1003E-11 |
| | THCmale | 3.80070E-12 | 0.038 | -1.5659E-11 | -4.6417E-13 |
| | TpCmale | 3.71663E-12 | 0.828 | -8.2406E-12 | 6.6183E-12 |
| CBDmale | VehFemale | 3.80070E-12 | 0.938 | -7.8948E-12 | 7.3002E-12 |
| | CBDfemale | 3.66465E-12 | 0.024 | -1.5832E-11 | -1.1808E-12 |
| | THCfemale | 3.46521E-12 | 0.009 | -1.6336E-11 | -2.4819E-12 |
| | TpCfemale | 3.46521E-12 | 0.000 | -2.1436E-11 | -7.5828E-12 |
| | VehMale | 3.80070E-12 | 0.374 | -1.1003E-11 | 4.1922E-12 |
| | THCmale | 3.55523E-12 | 0.002 | -1.8574E-11 | -4.3601E-12 |
| | TpCmale | 3.46521E-12 | 0.228 | -1.1143E-11 | 2.7104E-12 |
| THCmale | VehFemale | 3.80070E-12 | 0.005 | 3.5722E-12 | 1.8767E-11 |
| | CBDfemale | 3.66465E-12 | 0.422 | -4.3650E-12 | 1.0286E-11 |
| | THCfemale | 3.46521E-12 | 0.555 | -4.8687E-12 | 8.9850E-12 |
| | TpCfemale | 3.46521E-12 | 0.383 | -9.9696E-12 | 3.8841E-12 |
| | VehMale | 3.80070E-12 | 0.038 | 4.6417E-13 | 1.5659E-11 |
| | CBDmale | 3.55523E-12 | 0.002 | 4.3601E-12 | 1.8574E-11 |
| | TpCmale | 3.46521E-12 | 0.041 | 3.2363E-13 | 1.4177E-11 |
| TpCmale | VehFemale | 3.71663E-12 | 0.296 | -3.5103E-12 | 1.1349E-11 |
| | CBDfemale | 3.57738E-12 | 0.235 | -1.1441E-11 | 2.8611E-12 |
| | THCfemale | 3.37279E-12 | 0.129 | -1.1934E-11 | 1.5498E-12 |
| | TpCfemale | 3.37279E-12 | 0.003 | -1.7035E-11 | -3.5511E-12 |
| | VehMale | 3.71663E-12 | 0.828 | -6.6183E-12 | 8.2406E-12 |
| | CBDmale | 3.46521E-12 | 0.228 | -2.7104E-12 | 1.1143E-11 |
| | THCmale | 3.46521E-12 | 0.041 | -1.4177E-11 | -3.2363E-13 |

Supplemental Table 9.1 Descriptive statistics for *N*-acyl alanines (mol/L) in plasma collected from male and female adult mice 2h after receiving an i.p. injection of 10mg/kg CBD, THC, or THC+CBD

| | | N | Mean | Std. Deviation | Std. Error | 95% Confidence Interval for Mean | | Minimum | Maximum |
|---|---|---|---|---|---|---|---|---|---|
| | | | | | | Lower Bound | Upper Bound | | |
| *N*-palmitoyl alanine | Veh F | 7 | 3.2129E-08 | 3.23586E-09 | 1.22304E-09 | 2.9136E-08 | 3.5122E-08 | 2.89E-08 | 3.84E-08 |
| | CBD F | 8 | 3.0827E-08 | 4.22310E-09 | 1.49309E-09 | 2.7296E-08 | 3.4358E-08 | 2.52E-08 | 3.82E-08 |
| | THC F | 9 | 3.0438E-08 | 3.15107E-09 | 1.05036E-09 | 2.8016E-08 | 3.2861E-08 | 2.34E-08 | 3.35E-08 |
| | THC+CBD F | 10 | 3.2209E-08 | 2.24282E-09 | 7.09241E-10 | 3.0605E-08 | 3.3814E-08 | 2.85E-08 | 3.50E-08 |
| | Veh M | 7 | 3.2202E-08 | 1.98735E-09 | 7.51146E-10 | 3.0364E-08 | 3.4040E-08 | 2.98E-08 | 3.60E-08 |
| | CBD M | 10 | 3.2005E-08 | 3.41591E-09 | 1.08020E-09 | 2.9562E-08 | 3.4449E-08 | 2.54E-08 | 3.67E-08 |
| | THC M | 9 | 3.1304E-08 | 2.21984E-09 | 7.39948E-10 | 2.9598E-08 | 3.3010E-08 | 2.82E-08 | 3.46E-08 |
| | THC+CBD M | 10 | 3.2120E-08 | 2.59458E-09 | 8.20477E-10 | 3.0264E-08 | 3.3976E-08 | 2.84E-08 | 3.67E-08 |
| *N*-stearoyl alanine | Veh F | 6 | 2.0136E-08 | 5.81382E-09 | 2.37348E-09 | 1.4035E-08 | 2.6237E-08 | 1.38E-08 | 2.94E-08 |
| | CBD F | 9 | 2.0125E-08 | 9.90780E-09 | 3.30260E-09 | 1.2509E-08 | 2.7741E-08 | 1.01E-08 | 3.95E-08 |
| | THC F | 9 | 2.3799E-08 | 1.10447E-08 | 3.68157E-09 | 1.5309E-08 | 3.2288E-08 | 1.14E-08 | 3.70E-08 |
| | THC+CBD F | 10 | 2.0854E-08 | 1.02632E-08 | 3.24552E-09 | 1.3513E-08 | 2.8196E-08 | 9.95E-09 | 3.61E-08 |
| | Veh M | 7 | 1.6712E-08 | 3.06642E-09 | 1.15900E-09 | 1.3876E-08 | 1.9548E-08 | 1.35E-08 | 2.17E-08 |
| | CBD M | 11 | 2.0997E-08 | 8.92679E-09 | 2.69153E-09 | 1.5000E-08 | 2.6994E-08 | 1.13E-08 | 3.45E-08 |
| | THC M | 8 | 1.5198E-08 | 1.05758E-09 | 3.73912E-10 | 1.4314E-08 | 1.6082E-08 | 1.38E-08 | 1.70E-08 |
| | THC+CBD M | 9 | 1.5808E-08 | 2.28444E-09 | 7.61480E-10 | 1.4052E-08 | 1.7564E-08 | 1.32E-08 | 1.98E-08 |
| *N*-oleoyl alanine | Veh F | 7 | 2.9703E-08 | 3.86733E-09 | 1.46171E-09 | 2.6126E-08 | 3.3279E-08 | 2.57E-08 | 3.52E-08 |
| | CBD F | 9 | 3.1162E-08 | 7.51644E-09 | 2.50548E-09 | 2.5385E-08 | 3.6940E-08 | 2.26E-08 | 4.37E-08 |
| | THC F | 9 | 3.2326E-08 | 5.96729E-09 | 1.98910E-09 | 2.7739E-08 | 3.6913E-08 | 2.31E-08 | 4.39E-08 |
| | THC+CBD F | 11 | 2.9630E-08 | 4.68347E-09 | 1.41212E-09 | 2.6483E-08 | 3.2776E-08 | 2.29E-08 | 3.65E-08 |
| | Veh M | 8 | 3.4275E-08 | 5.53919E-09 | 1.95840E-09 | 2.9644E-08 | 3.8906E-08 | 2.61E-08 | 4.06E-08 |
| | CBD M | 10 | 3.2082E-08 | 5.86056E-09 | 1.85327E-09 | 2.7889E-08 | 3.6274E-08 | 2.68E-08 | 4.23E-08 |
| | THC M | 9 | 3.3135E-08 | 5.58597E-09 | 1.86199E-09 | 2.8841E-08 | 3.7428E-08 | 2.52E-08 | 4.36E-08 |
| | THC+CBD M | 9 | 3.0282E-08 | 3.22364E-09 | 1.07455E-09 | 2.7804E-08 | 3.2760E-08 | 2.46E-08 | 3.45E-08 |
| *N*-linoleoyl alanine | Veh F | 7 | 5.5483E-09 | 1.81061E-09 | 6.84347E-10 | 3.8737E-09 | 7.2228E-09 | 4.00E-09 | 9.22E-09 |
| | CBD F | 8 | 7.0179E-09 | 3.41371E-09 | 1.20693E-09 | 4.1639E-09 | 9.8718E-09 | 3.14E-09 | 1.30E-08 |
| | THC F | 8 | 6.3186E-09 | 3.02648E-09 | 1.07002E-09 | 3.7884E-09 | 8.8488E-09 | 2.94E-09 | 1.27E-08 |
| | THC+CBD F | 11 | 5.0250E-09 | 1.29621E-09 | 3.90822E-10 | 4.1542E-09 | 5.8958E-09 | 3.46E-09 | 6.67E-09 |
| | Veh M | 8 | 5.4262E-09 | 2.69105E-09 | 9.51430E-10 | 3.1765E-09 | 7.6760E-09 | 2.58E-09 | 9.91E-09 |
| | CBD M | 9 | 5.6421E-09 | 2.76581E-09 | 9.21938E-10 | 3.5161E-09 | 7.7681E-09 | 2.54E-09 | 9.67E-09 |
| | THC M | 9 | 5.2093E-09 | 1.30636E-09 | 4.35454E-10 | 4.2051E-09 | 6.2135E-09 | 2.68E-09 | 6.95E-09 |
| | THC+CBD M | 9 | 4.4501E-09 | 1.12696E-09 | 3.75655E-10 | 3.5839E-09 | 5.3164E-09 | 3.03E-09 | 6.37E-09 |
| *N*-arachidonoyl alanine | Veh F | BAL | | | | | | | |
| | CBD F | BDL | | | | | | | |
| | THC F | BDL | | | | | | | |
| | THC+CBD F | BAL | | | | | | | |
| | Veh M | BDL | | | | | | | |
| | CBD M | BDL | | | | | | | |
| | THC M | BDL | | | | | | | |
| | THC+CBD M | BDL | | | | | | | |
| *N*-docosahexaenoyl alanine | Veh F | BDL | | | | | | | |
| | CBD F | BDL | | | | | | | |
| | THC F | BDL | | | | | | | |
| | THC+CBD F | BDL | | | | | | | |
| | Veh M | BDL | | | | | | | |
| | CBD M | BAL | | | | | | | |
| | THC M | BDL | | | | | | | |
| | THC+CBD M | BDL | | | | | | | |

Supplemental Table 9.2 Descriptive statistics for *N*-acyl alanines (mol/g) in liver collected from male and female adult mice 2h after receiving an i.p. injection of 10mg/kg CBD, THC, or THC+CBD

| | | | | | | 95% Confidence Interval for Mean | | | |
|---|---|---|---|---|---|---|---|---|---|
| | | N | Mean | Std. Deviation | Std. Error | Lower Bound | Upper Bound | Minimum | Maximum |
| *N*-palmitoyl alanine | Veh F | 7 | 3.4475E-10 | 1.03479E-10 | 3.91113E-11 | 2.4905E-10 | 4.4045E-10 | 2.28E-10 | 5.1E-10 |
| | CBD F | 7 | 3.8856E-10 | 1.47639E-10 | 5.58023E-11 | 2.5201E-10 | 5.251E-10 | 2.1E-10 | 6.24E-10 |
| | THC F | 10 | 4.3261E-10 | 1.19646E-10 | 3.78354E-11 | 3.4702E-10 | 5.182E-10 | 2.71E-10 | 6.3E-10 |
| | THC+CBD F | 9 | 4.6444E-10 | 9.09783E-11 | 3.03261E-11 | 3.9451E-10 | 5.3438E-10 | 3.74E-10 | 6.74E-10 |
| | Veh M | 7 | 3.442E-10 | 7.98378E-11 | 3.01758E-11 | 2.7036E-10 | 4.1804E-10 | 2.28E-10 | 4.3E-10 |
| | CBD M | 10 | 3.0568E-10 | 6.67529E-11 | 2.11091E-11 | 2.5793E-10 | 3.5343E-10 | 2.33E-10 | 4.37E-10 |
| | THC M | 9 | 4.7123E-10 | 8.49188E-11 | 2.83063E-11 | 4.0596E-10 | 5.3651E-10 | 3.19E-10 | 5.78E-10 |
| | THC+CBD M | 10 | 3.2381E-10 | 8.29411E-11 | 2.62283E-11 | 2.6447E-10 | 3.8314E-10 | 1.94E-10 | 4.62E-10 |
| *N*-stearoyl alanine | Veh F | 7 | 5.5278E-11 | 2.51906E-11 | 9.52116E-12 | 3.1980E-11 | 7.8575E-11 | 1.84E-11 | 9.40E-11 |
| | CBD F | 8 | 9.5449E-11 | 5.59201E-11 | 1.97707E-11 | 4.8699E-11 | 1.4220E-10 | 2.90E-11 | 2.00E-10 |
| | THC F | 9 | 7.1623E-11 | 1.91315E-11 | 6.37716E-12 | 5.6918E-11 | 8.6329E-11 | 4.25E-11 | 1.05E-10 |
| | THC+CBD F | 9 | 6.9997E-11 | 2.92146E-11 | 9.73819E-12 | 4.7540E-11 | 9.2453E-11 | 2.47E-11 | 1.27E-10 |
| | Veh M | 7 | 5.7531E-11 | 1.69383E-11 | 6.40207E-12 | 4.1866E-11 | 7.3196E-11 | 4.12E-11 | 8.93E-11 |
| | CBD M | 9 | 4.8142E-11 | 2.79418E-11 | 9.31392E-12 | 2.6664E-11 | 6.9620E-11 | 2.73E-11 | 1.10E-10 |
| | THC M | 8 | 4.9729E-11 | 2.17440E-11 | 7.68768E-12 | 3.1551E-11 | 6.7908E-11 | 1.56E-11 | 7.45E-11 |
| | THC+CBD M | 9 | 5.0848E-11 | 1.83335E-11 | 6.11117E-12 | 3.6756E-11 | 6.4940E-11 | 3.00E-11 | 8.65E-11 |
| *N*-oleoyl alanine | Veh F | 7 | 1.9319E-10 | 6.35245E-11 | 2.40100E-11 | 1.3444E-10 | 2.5194E-10 | 1.20E-10 | 2.97E-10 |
| | CBD F | 7 | 2.2017E-10 | 7.90385E-11 | 2.98737E-11 | 1.4707E-10 | 2.9326E-10 | 1.28E-10 | 3.30E-10 |
| | THC F | 10 | 3.7050E-10 | 1.80657E-10 | 5.71288E-11 | 2.4126E-10 | 4.9973E-10 | 1.62E-10 | 7.20E-10 |
| | THC+CBD F | 10 | 3.1410E-10 | 9.38757E-11 | 2.96861E-11 | 2.4695E-10 | 3.8126E-10 | 1.78E-10 | 4.85E-10 |
| | Veh M | 7 | 1.8857E-10 | 4.81409E-11 | 1.81955E-11 | 1.4405E-10 | 2.3310E-10 | 1.32E-10 | 2.74E-10 |
| | CBD M | 10 | 1.7899E-10 | 4.28445E-11 | 1.35486E-11 | 1.4834E-10 | 2.0964E-10 | 9.93E-11 | 2.38E-10 |
| | THC M | 8 | 2.1799E-10 | 3.84079E-11 | 1.35792E-11 | 1.8588E-10 | 2.5010E-10 | 1.61E-10 | 2.69E-10 |
| | THC+CBD M | 11 | 2.5754E-10 | 8.85437E-11 | 2.66969E-11 | 1.9806E-10 | 3.1703E-10 | 1.05E-10 | 3.89E-10 |
| *N*-linoleoyl alanine | Veh F | 7 | 1.9272E-10 | 5.36512E-11 | 2.02783E-11 | 1.4310E-10 | 2.4234E-10 | 1.29E-10 | 2.77E-10 |
| | CBD F | 7 | 2.3105E-10 | 9.78619E-11 | 3.69883E-11 | 1.4054E-10 | 3.2155E-10 | 1.02E-10 | 3.67E-10 |
| | THC F | 10 | 2.7358E-10 | 7.42162E-11 | 2.34692E-11 | 2.2049E-10 | 3.2667E-10 | 1.72E-10 | 3.91E-10 |
| | THC+CBD F | 10 | 3.5452E-10 | 9.39568E-11 | 2.97117E-11 | 2.8730E-10 | 4.2173E-10 | 1.71E-10 | 5.06E-10 |
| | Veh M | 7 | 1.5425E-10 | 4.88630E-11 | 1.84685E-11 | 1.0906E-10 | 1.9944E-10 | 8.78E-11 | 2.22E-10 |
| | CBD M | 10 | 1.7138E-10 | 5.65910E-11 | 1.78956E-11 | 1.3090E-10 | 2.1186E-10 | 1.19E-10 | 2.66E-10 |
| | THC M | 8 | 3.2594E-10 | 2.85421E-11 | 1.00911E-11 | 3.0208E-10 | 3.4980E-10 | 2.81E-10 | 3.54E-10 |
| | THC+CBD M | 10 | 2.1867E-10 | 5.68564E-11 | 1.79796E-11 | 1.7800E-10 | 2.5934E-10 | 1.40E-10 | 3.25E-10 |
| *N*-arachidonoyl alanine | Veh F | 8 | 9.9540E-11 | 5.20317E-11 | 1.83960E-11 | 5.6041E-11 | 1.4304E-10 | 3.82E-11 | 1.96E-10 |
| | CBD F | 7 | 1.1301E-10 | 4.30704E-11 | 1.62791E-11 | 7.3176E-11 | 1.5284E-10 | 5.84E-11 | 1.60E-10 |
| | THC F | 10 | 1.4485E-10 | 4.24610E-11 | 1.34274E-11 | 1.1448E-10 | 1.7523E-10 | 8.38E-11 | 2.06E-10 |
| | THC+CBD F | 9 | 1.2963E-10 | 2.79469E-11 | 9.31564E-12 | 1.0815E-10 | 1.5111E-10 | 7.45E-11 | 1.66E-10 |
| | Veh M | 7 | 7.8762E-11 | 3.55208E-11 | 1.34256E-11 | 4.5911E-11 | 1.1161E-10 | 2.04E-11 | 1.27E-10 |
| | CBD M | 9 | 7.7231E-11 | 1.77087E-11 | 5.90288E-12 | 6.3619E-11 | 9.0843E-11 | 5.29E-11 | 1.07E-10 |
| | THC M | 8 | 1.4106E-10 | 1.73537E-11 | 6.13545E-12 | 1.2656E-10 | 1.5557E-10 | 1.06E-10 | 1.61E-10 |
| | THC+CBD M | 10 | 9.5942E-11 | 3.31502E-11 | 1.04830E-11 | 7.2227E-11 | 1.1966E-10 | 4.41E-11 | 1.46E-10 |
| *N*-docosahexaenoyl alanine | Veh F | 8 | 6.1512E-11 | 2.40980E-11 | 8.51994E-12 | 4.1365E-11 | 8.1658E-11 | 3.17E-11 | 9.48E-11 |
| | CBD F | 8 | 9.4546E-11 | 5.73820E-11 | 2.02876E-11 | 4.6573E-11 | 1.4252E-10 | 2.62E-11 | 1.97E-10 |
| | THC F | 10 | 1.0598E-10 | 4.37112E-11 | 1.38227E-11 | 7.4716E-11 | 1.3725E-10 | 5.93E-11 | 1.60E-10 |
| | THC+CBD F | 9 | 1.0075E-10 | 2.75117E-11 | 9.17057E-12 | 7.9599E-11 | 1.2189E-10 | 3.69E-11 | 1.32E-10 |
| | Veh M | 7 | 5.8666E-11 | 2.25374E-11 | 8.51832E-12 | 3.7822E-11 | 7.9510E-11 | 3.24E-11 | 9.30E-11 |
| | CBD M | 9 | 4.7607E-11 | 1.23659E-11 | 4.12198E-12 | 3.8101E-11 | 5.7112E-11 | 2.16E-11 | 6.24E-11 |
| | THC M | 9 | 1.0617E-10 | 4.08849E-11 | 1.36283E-11 | 7.4739E-11 | 1.3759E-10 | 2.51E-11 | 1.75E-10 |
| | THC+CBD M | 10 | 7.5087E-11 | 2.34727E-11 | 7.42271E-12 | 5.8296E-11 | 9.1879E-11 | 4.45E-11 | 1.31E-10 |

Supplemental Table 9.3 Independent t-test comparing male and female levels of *N*-acyl alanines measured in plasma collected from adult mice 2h after an i.p. injection of 10mg/kg CBD, THC, or THC+CBD

| | | | Levene's Test for Equality of Variances | | t-test for Equality of Means | | | | | | |
|---|---|---|---|---|---|---|---|---|---|---|---|
| | | | | | | | Significance | | | 95% Confidence Interval of the Difference | |
| | | | F | Sig. | t | df | One-Sided p | Two-Sided p | Std. Error Difference | Lower | Upper |
| Veh | *N*-palmitoyl alanine | Equal variances assumed | 1.444 | 0.253 | -0.051 | 12 | 0.480 | 0.960 | 1.43529E-09 | -3.20019E-09 | 3.05425E-09 |
| | | Equal variances not assumed | | | -0.051 | 9.963 | 0.480 | 0.960 | 1.43529E-09 | -3.27262E-09 | 3.12668E-09 |
| | *N*-stearoyl alanine | Equal variances assumed | 1.618 | 0.230 | 1.359 | 11 | 0.101 | 0.201 | 2.51853E-09 | -2.11931E-09 | 8.96718E-09 |
| | | Equal variances not assumed | | | 1.296 | 7.322 | 0.117 | 0.234 | 2.64134E-09 | -2.76662E-09 | 9.61449E-09 |
| | *N*-oleoyl alanine | Equal variances assumed | 1.333 | 0.269 | -1.825 | 13 | 0.045 | 0.091 | 2.50487E-09 | -9.98405E-09 | 8.38827E-10 |
| | | Equal variances not assumed | | | -1.871 | 12.460 | 0.042 | 0.085 | 2.44375E-09 | -9.87537E-09 | 7.30143E-10 |
| | *N*-linoleoyl alanine | Equal variances assumed | 2.265 | 0.156 | 0.101 | 13 | 0.460 | 0.921 | 1.20406E-09 | -2.47921E-09 | 2.72324E-09 |
| | | Equal variances not assumed | | | 0.104 | 12.282 | 0.459 | 0.919 | 1.17199E-09 | -2.42504E-09 | 2.66907E-09 |
| CBD | *N*-palmitoyl alanine | Equal variances assumed | 0.406 | 0.533 | -0.655 | 16 | 0.261 | 0.522 | 1.79788E-09 | -4.98949E-09 | 2.63319E-09 |
| | | Equal variances not assumed | | | -0.639 | 13.392 | 0.267 | 0.533 | 1.84287E-09 | -5.14762E-09 | 2.79132E-09 |
| | *N*-stearoyl alanine | Equal variances assumed | 0.016 | 0.899 | -0.207 | 18 | 0.419 | 0.838 | 4.21396E-09 | -9.72568E-09 | 7.98073E-09 |
| | | Equal variances not assumed | | | -0.205 | 16.377 | 0.420 | 0.840 | 4.26046E-09 | -9.88739E-09 | 8.14244E-09 |
| | *N*-oleoyl alanine | Equal variances assumed | 1.292 | 0.271 | -0.299 | 17 | 0.384 | 0.768 | 3.07432E-09 | -7.40578E-09 | 5.56672E-09 |
| | | Equal variances not assumed | | | -0.295 | 15.125 | 0.386 | 0.772 | 3.11642E-09 | -7.55725E-09 | 5.71819E-09 |

| | | | | | | | | | | | |
|---|---|---|---|---|---|---|---|---|---|---|---|
| | *N*-linoleoyl alanine | Equal variances assumed | 0.172 | 0.684 | 0.918 | 15 | 0.187 | 0.373 | 1.49911E-09 | -1.81952E-09 | 4.57104E-09 |
| | | Equal variances not assumed | | | 0.906 | 13.523 | 0.190 | 0.381 | 1.51877E-09 | -1.89247E-09 | 4.64399E-09 |
| THC | *N*-palmitoyl alanine | Equal variances assumed | 0.247 | 0.626 | -0.674 | 16 | 0.255 | 0.510 | 1.28483E-09 | -3.58934E-09 | 1.85807E-09 |
| | | Equal variances not assumed | | | -0.674 | 14.371 | 0.256 | 0.511 | 1.28483E-09 | -3.61464E-09 | 1.88338E-09 |
| | *N*-stearoyl alanine | Equal variances assumed | 97.503 | 0.000 | 2.186 | 15 | 0.023 | 0.045 | 3.93502E-09 | 2.13502E-10 | 1.69881E-08 |
| | | Equal variances not assumed | | | 2.324 | 8.165 | 0.024 | 0.048 | 3.70051E-09 | 9.73174E-11 | 1.71043E-08 |
| | *N*-oleoyl alanine | Equal variances assumed | 0.091 | 0.767 | -0.297 | 16 | 0.385 | 0.770 | 2.72461E-09 | -6.58475E-09 | 4.96709E-09 |
| | | Equal variances not assumed | | | -0.297 | 15.931 | 0.385 | 0.770 | 2.72461E-09 | -6.58679E-09 | 4.96913E-09 |
| | *N*-linoleoyl alanine | Equal variances assumed | 2.799 | 0.115 | 1.003 | 15 | 0.166 | 0.332 | 1.10641E-09 | -1.24893E-09 | 3.46760E-09 |
| | | Equal variances not assumed | | | 0.960 | 9.288 | 0.181 | 0.361 | 1.15523E-09 | -1.49170E-09 | 3.71036E-09 |
| THC+CBD | *N*-palmitoyl alanine | Equal variances assumed | 0.082 | 0.778 | 0.082 | 18 | 0.468 | 0.935 | 1.08453E-09 | -2.18911E-09 | 2.36791E-09 |
| | | Equal variances not assumed | | | 0.082 | 17.631 | 0.468 | 0.935 | 1.08453E-09 | -2.19254E-09 | 2.37134E-09 |
| | *N*-stearoyl alanine | Equal variances assumed | 20.404 | 0.000 | 1.439 | 17 | 0.084 | 0.168 | 3.50586E-09 | -2.35047E-09 | 1.24430E-08 |
| | | Equal variances not assumed | | | 1.514 | 9.984 | 0.081 | 0.161 | 3.33365E-09 | -2.38319E-09 | 1.24757E-08 |
| | *N*-oleoyl alanine | Equal variances assumed | 2.965 | 0.102 | -0.354 | 18 | 0.364 | 0.727 | 1.84252E-09 | -4.52377E-09 | 3.21820E-09 |
| | | Equal variances not assumed | | | -0.368 | 17.570 | 0.359 | 0.717 | 1.77446E-09 | -4.38735E-09 | 3.08178E-09 |
| | *N*-linoleoyl alanine | Equal variances assumed | 0.535 | 0.474 | 1.045 | 18 | 0.155 | 0.310 | 5.50094E-10 | -5.80862E-10 | 1.73055E-09 |
| | | Equal variances not assumed | | | 1.060 | 17.907 | 0.152 | 0.303 | 5.42087E-10 | -5.64463E-10 | 1.71415E-09 |

Supplemental Table 9.4 Independent t-test comparing male and female levels of *N*-acyl alanines measured in liver collected from adult mice 2h after an i.p. injection of 10mg/kg CBD, THC, or THC+CBD

| | | | Levene's Test for Equality of Variances | | t-test for Equality of Means | | | | | | | |
|---|---|---|---|---|---|---|---|---|---|---|---|---|
| | | | | | | | Significance | | | | 95% Confidence Interval of the Difference | |
| | | | F | Sig. | t | df | One-Sided p | Two-Sided p | Mean Difference | Std. Error Difference | Lower | Upper |
| Veh | *N*-palmitoyl alanine | Equal variances assumed | 0.603 | 0.452 | 0.011 | 12 | 0.496 | 0.991 | 5.47857E-13 | 4.93992E-11 | -1.07084E-10 | 1.08179E-10 |
| | | Equal variances not assumed | | | 0.011 | 11.274 | 0.496 | 0.991 | 5.47857E-13 | 4.93992E-11 | -1.07857E-10 | 1.08953E-10 |
| | *N*-stearoyl alanine | Equal variances assumed | 0.701 | 0.419 | -0.196 | 12 | 0.424 | 0.848 | -2.25313E-12 | 1.14734E-11 | -2.72515E-11 | 2.27453E-11 |
| | | Equal variances not assumed | | | -0.196 | 10.505 | 0.424 | 0.848 | -2.25313E-12 | 1.14734E-11 | -2.76520E-11 | 2.31457E-11 |
| | *N*-oleoyl alanine | Equal variances assumed | 1.568 | 0.234 | 0.153 | 12 | 0.440 | 0.881 | 4.61400E-12 | 3.01257E-11 | -6.10243E-11 | 7.02523E-11 |
| | | Equal variances not assumed | | | 0.153 | 11.182 | 0.441 | 0.881 | 4.61400E-12 | 3.01257E-11 | -6.15605E-11 | 7.07885E-11 |
| | *N*-linoleoyl alanine | Equal variances assumed | 0.001 | 0.981 | 1.403 | 12 | 0.093 | 0.186 | 3.84718E-11 | 2.74280E-11 | -2.12886E-11 | 9.82322E-11 |
| | | Equal variances not assumed | | | 1.403 | 11.897 | 0.093 | 0.186 | 3.84718E-11 | 2.74280E-11 | -2.13462E-11 | 9.82898E-11 |
| | *N*-arachidonoyl alanine | Equal variances assumed | 1.030 | 0.329 | 0.889 | 13 | 0.195 | 0.390 | 2.07783E-11 | 2.33765E-11 | -2.97234E-11 | 7.12801E-11 |
| | | Equal variances not assumed | | | 0.912 | 12.354 | 0.190 | 0.379 | 2.07783E-11 | 2.27741E-11 | -2.86850E-11 | 7.02416E-11 |
| | *N*-docosahexaenoyl alanine | Equal variances assumed | 0.075 | 0.789 | 0.235 | 13 | 0.409 | 0.818 | 2.84557E-12 | 1.21058E-11 | -2.33075E-11 | 2.89986E-11 |
| | | Equal variances not assumed | | | 0.236 | 12.923 | 0.408 | 0.817 | 2.84557E-12 | 1.20479E-11 | -2.31980E-11 | 2.88891E-11 |
| CBD | *N*-palmitoyl alanine | Equal variances assumed | 6.982 | 0.018 | 1.576 | 15 | 0.068 | 0.136 | 8.28741E-11 | 5.25998E-11 | -2.92398E-11 | 1.94988E-10 |
| | | Equal variances not assumed | | | 1.389 | 7.734 | 0.102 | 0.203 | 8.28741E-11 | 5.96615E-11 | -5.55320E-11 | 2.21280E-10 |
| | *N*-stearoyl alanine | Equal variances assumed | 3.354 | 0.087 | 2.248 | 15 | 0.020 | 0.040 | 4.73079E-11 | 2.10445E-11 | 2.45255E-12 | 9.21632E-11 |

| | | | | | | | | | | | | |
|---|---|---|---|---|---|---|---|---|---|---|---|---|
| | | Equal variances not assumed | | | 2.165 | 10.020 | 0.028 | 0.056 | 4.73079E-11 | 2.18548E-11 | -1.37448E-12 | 9.59902E-11 |
| | *N*-oleoyl alanine | Equal variances assumed | 3.261 | 0.091 | 1.392 | 15 | 0.092 | 0.184 | 4.11713E-11 | 2.95692E-11 | -2.18540E-11 | 1.04197E-10 |
| | | Equal variances not assumed | | | 1.255 | 8.483 | 0.121 | 0.243 | 4.11713E-11 | 3.28025E-11 | -3.37285E-11 | 1.16071E-10 |
| | *N*-linoleoyl alanine | Equal variances assumed | 4.101 | 0.061 | 1.596 | 15 | 0.066 | 0.131 | 5.96666E-11 | 3.73763E-11 | -1.99991E-11 | 1.39332E-10 |
| | | Equal variances not assumed | | | 1.452 | 8.816 | 0.091 | 0.181 | 5.96666E-11 | 4.10900E-11 | -3.35825E-11 | 1.52916E-10 |
| | *N*-arachidonoyl alanine | Equal variances assumed | 22.270 | 0.000 | 2.275 | 14 | 0.020 | 0.039 | 3.57788E-11 | 1.57297E-11 | 2.04203E-12 | 6.95155E-11 |
| | | Equal variances not assumed | | | 2.066 | 7.583 | 0.037 | 0.075 | 3.57788E-11 | 1.73163E-11 | -4.53786E-12 | 7.60954E-11 |
| | *N*-docosahexaenoyl alanine | Equal variances assumed | 16.275 | 0.001 | 2.401 | 15 | 0.015 | 0.030 | 4.69391E-11 | 1.95464E-11 | 5.27687E-12 | 8.86013E-11 |
| | | Equal variances not assumed | | | 2.267 | 7.579 | 0.027 | 0.055 | 4.69391E-11 | 2.07021E-11 | -1.26616E-12 | 9.51443E-11 |
| | *N*-palmitoyl alanine | Equal variances assumed | 1.817 | 0.195 | -0.803 | 17 | 0.217 | 0.433 | -3.86255E-11 | 4.81284E-11 | -1.40167E-10 | 6.29165E-11 |
| | | Equal variances not assumed | | | -0.817 | 16.189 | 0.213 | 0.426 | -3.86255E-11 | 4.72521E-11 | -1.38701E-10 | 6.14497E-11 |
| | *N*-stearoyl alanine | Equal variances assumed | 0.116 | 0.738 | 2.210 | 15 | 0.022 | 0.043 | 2.18942E-11 | 9.90891E-12 | 7.73818E-13 | 4.30145E-11 |
| | | Equal variances not assumed | | | 2.192 | 14.104 | 0.023 | 0.046 | 2.18942E-11 | 9.98842E-12 | 4.86004E-13 | 4.33023E-11 |
| THC | *N*-oleoyl alanine | Equal variances assumed | 10.826 | 0.005 | 2.332 | 16 | 0.017 | 0.033 | 1.52505E-10 | 6.53899E-11 | 1.38849E-11 | 2.91126E-10 |
| | | Equal variances not assumed | | | 2.597 | 10.005 | 0.013 | 0.027 | 1.52505E-10 | 5.87205E-11 | 2.16760E-11 | 2.83334E-10 |
| | *N*-linoleoyl alanine | Equal variances assumed | 4.855 | 0.043 | -1.878 | 16 | 0.039 | 0.079 | -5.23610E-11 | 2.78802E-11 | -1.11464E-10 | 6.74227E-12 |
| | | Equal variances not assumed | | | -2.050 | 12.104 | 0.031 | 0.063 | -5.23610E-11 | 2.55467E-11 | -1.07970E-10 | 3.24775E-12 |
| | *N*-arachidonoyl alanine | Equal variances assumed | 7.636 | 0.014 | 0.236 | 16 | 0.408 | 0.816 | 3.78950E-12 | 1.60570E-11 | -3.02499E-11 | 3.78289E-11 |
| | | Equal variances | | | 0.257 | 12.453 | 0.401 | 0.802 | 3.78950E-12 | 1.47627E-11 | -2.82465E-11 | 3.58255E-11 |

| | | | | | | | | | | | | |
|---|---|---|---|---|---|---|---|---|---|---|---|---|
| | | not assumed | | | | | | | | | | |
| | *N*-docosahexaenoyl alanine | Equal variances assumed | 1.304 | 0.269 | -0.009 | 17 | 0.496 | 0.993 | -1.81150E-13 | 1.94836E-11 | -4.12879E-11 | 4.09256E-11 |
| | | Equal variances not assumed | | | -0.009 | 16.966 | 0.496 | 0.993 | -1.81150E-13 | 1.94113E-11 | -4.11416E-11 | 4.07793E-11 |
| THC+CBD | *N*-palmitoyl alanine | Equal variances assumed | 0.000 | 0.991 | 3.526 | 17 | 0.001 | 0.003 | 1.40638E-10 | 3.98892E-11 | 5.64790E-11 | 2.24797E-10 |
| | | Equal variances not assumed | | | 3.508 | 16.325 | 0.001 | 0.003 | 1.40638E-10 | 4.00948E-11 | 5.57780E-11 | 2.25498E-10 |
| | *N*-stearoyl alanine | Equal variances assumed | 1.282 | 0.274 | 1.666 | 16 | 0.058 | 0.115 | 1.91487E-11 | 1.14969E-11 | -5.22359E-12 | 4.35211E-11 |
| | | Equal variances not assumed | | | 1.666 | 13.455 | 0.059 | 0.119 | 1.91487E-11 | 1.14969E-11 | -5.60368E-12 | 4.39012E-11 |
| | *N*-oleoyl alanine | Equal variances assumed | 0.050 | 0.826 | 1.421 | 19 | 0.086 | 0.172 | 5.65620E-11 | 3.98081E-11 | -2.67572E-11 | 1.39881E-10 |
| | | Equal variances not assumed | | | 1.417 | 18.534 | 0.087 | 0.173 | 5.65620E-11 | 3.99248E-11 | -2.71440E-11 | 1.40268E-10 |
| | *N*-linoleoyl alanine | Equal variances assumed | 0.969 | 0.338 | 3.912 | 18 | 0.001 | 0.001 | 1.35846E-10 | 3.47283E-11 | 6.28846E-11 | 2.08807E-10 |
| | | Equal variances not assumed | | | 3.912 | 14.812 | 0.001 | 0.001 | 1.35846E-10 | 3.47283E-11 | 6.17425E-11 | 2.09949E-10 |
| | *N*-arachidonoyl alanine | Equal variances assumed | 0.521 | 0.480 | 2.379 | 17 | 0.015 | 0.029 | 3.36854E-11 | 1.41568E-11 | 3.81718E-12 | 6.35536E-11 |
| | | Equal variances not assumed | | | 2.402 | 16.941 | 0.014 | 0.028 | 3.36854E-11 | 1.40241E-11 | 4.08939E-12 | 6.32814E-11 |
| | *N*-docosahexaenoyl alanine | Equal variances assumed | 0.012 | 0.913 | 2.194 | 17 | 0.021 | 0.042 | 2.56592E-11 | 1.16950E-11 | 9.84889E-13 | 5.03335E-11 |
| | | Equal variances not assumed | | | 2.175 | 15.864 | 0.023 | 0.045 | 2.56592E-11 | 1.17981E-11 | 6.30792E-13 | 5.06876E-11 |

Supplemental Table 9.5 One-way ANOVA for *N*-acyl alanines measured in plasma collected from adult mice 2h after an i.p. injection of 10mg/kg CBD, THC, or THC+CBD

| | | Sum of Squares | df | Mean Square | F | Sig. |
|---|---|---|---|---|---|---|
| *N*-palmitoyl alanine | Between Groups | 3.00E-17 | 7 | 4.29E-18 | 0.492 | 0.8369909 |
| | Within Groups | 5.41E-16 | 62 | 8.73E-18 | | |
| | Total | 5.71E-16 | 69 | | | |
| *N*-stearoyl alanine | Between Groups | 5.39E-16 | 7 | 7.70E-17 | 1.243 | 0.2939619 |
| | Within Groups | 3.78E-15 | 61 | 6.20E-17 | | |
| | Total | 4.32E-15 | 68 | | | |
| *N*-oleoyl alanine | Between Groups | 1.71E-16 | 7 | 2.44E-17 | 0.819 | 0.5746840 |
| | Within Groups | 1.90E-15 | 64 | 2.97E-17 | | |
| | Total | 2.07E-15 | 71 | | | |
| *N*-linoleoyl alanine | Between Groups | 3.71E-17 | 7 | 5.30E-18 | 1.017 | 0.4282239 |
| | Within Groups | 3.18E-16 | 61 | 5.21E-18 | | |
| | Total | 3.55E-16 | 68 | | | |

Supplemental Table 9.6 One-way ANOVA for *N*-acyl alanines measured in liver collected from adult mice 2h after an i.p. injection of 10mg/kg CBD, THC, or THC+CBD

| | | Sum of Squares | df | Mean Square | F | Sig. |
|---|---|---|---|---|---|---|
| *N*-palmitoyl alanine | Between Groups | 2.70E-19 | 7 | 3.85E-20 | 3.999 | 0.00117 |
| | Within Groups | 5.88E-19 | 61 | 9.64E-21 | | |
| | Total | 8.58E-19 | 68 | | | |
| *N*-stearoyl alanine | Between Groups | 1.49E-20 | 7 | 2.12E-21 | 2.491 | 0.02621 |
| | Within Groups | 4.94E-20 | 58 | 8.52E-22 | | |
| | Total | 6.43E-20 | 65 | | | |
| *N*-oleoyl alanine | Between Groups | 3.01E-19 | 7 | 4.30E-20 | 4.808 | 0.00023 |
| | Within Groups | 5.54E-19 | 62 | 8.93E-21 | | |
| | Total | 8.55E-19 | 69 | | | |
| *N*-linoleoyl alanine | Between Groups | 3.20E-19 | 7 | 4.57E-20 | 9.890 | 0.00000 |
| | Within Groups | 2.82E-19 | 61 | 4.62E-21 | | |
| | Total | 6.01E-19 | 68 | | | |
| *N*-arachidonoyl alanine | Between Groups | 4.27E-20 | 7 | 6.10E-21 | 4.904 | 0.00020 |
| | Within Groups | 7.46E-20 | 60 | 1.24E-21 | | |
| | Total | 1.17E-19 | 67 | | | |
| *N*-docosahexaenoyl alanine | Between Groups | 3.37E-20 | 7 | 4.82E-21 | 4.093 | 0.00095 |
| | Within Groups | 7.30E-20 | 62 | 1.18E-21 | | |
| | Total | 1.07E-19 | 69 | | | |

#### Supplemental Table 9.7 Fisher's LSD post-hoc test for plasma *N*-acyl alanine ANOVA

| Dependent Variable | | | Std. Error | Sig. | 95% Confidence Interval Lower Bound | 95% Confidence Interval Upper Bound |
|---|---|---|---|---|---|---|
| *N*-palmitoyl alanine | VehFemale | CBDfemale | 1.53E-09 | 0.3977 | -1.75E-09 | 4.36E-09 |
| | | THCfemale | 1.49E-09 | 0.2605 | -1.29E-09 | 4.67E-09 |
| | | TpCfemale | 1.46E-09 | 0.9562 | -2.99E-09 | 2.83E-09 |
| | | VehMale | 1.58E-09 | 0.9633 | -3.23E-09 | 3.08E-09 |
| | | CBDmale | 1.46E-09 | 0.9324 | -2.79E-09 | 3.03E-09 |
| | | THCmale | 1.49E-09 | 0.5814 | -2.15E-09 | 3.80E-09 |
| | | TpCmale | 1.46E-09 | 0.9950 | -2.90E-09 | 2.92E-09 |
| | CBDfemale | VehFemale | 1.53E-09 | 0.3977 | -4.36E-09 | 1.75E-09 |
| | | THCfemale | 1.44E-09 | 0.7876 | -2.48E-09 | 3.26E-09 |
| | | TpCfemale | 1.40E-09 | 0.3277 | -4.18E-09 | 1.42E-09 |
| | | VehMale | 1.53E-09 | 0.3719 | -4.43E-09 | 1.68E-09 |
| | | CBDmale | 1.40E-09 | 0.4037 | -3.98E-09 | 1.62E-09 |
| | | THCmale | 1.44E-09 | 0.7407 | -3.35E-09 | 2.39E-09 |
| | | TpCmale | 1.40E-09 | 0.3597 | -4.09E-09 | 1.51E-09 |
| | THCfemale | VehFemale | 1.49E-09 | 0.2605 | -4.67E-09 | 1.29E-09 |
| | | CBDfemale | 1.44E-09 | 0.7876 | -3.26E-09 | 2.48E-09 |
| | | TpCfemale | 1.36E-09 | 0.1968 | -4.48E-09 | 9.42E-10 |
| | | VehMale | 1.49E-09 | 0.2407 | -4.74E-09 | 1.21E-09 |
| | | CBDmale | 1.36E-09 | 0.2529 | -4.28E-09 | 1.15E-09 |
| | | THCmale | 1.39E-09 | 0.5365 | -3.65E-09 | 1.92E-09 |
| | | TpCmale | 1.36E-09 | 0.2201 | -4.39E-09 | 1.03E-09 |
| | TpCfemale | VehFemale | 1.46E-09 | 0.9562 | -2.83E-09 | 2.99E-09 |
| | | CBDfemale | 1.40E-09 | 0.3277 | -1.42E-09 | 4.18E-09 |
| | | THCfemale | 1.36E-09 | 0.1968 | -9.42E-10 | 4.48E-09 |
| | | VehMale | 1.46E-09 | 0.9960 | -2.90E-09 | 2.92E-09 |
| | | CBDmale | 1.32E-09 | 0.8776 | -2.44E-09 | 2.85E-09 |
| | | THCmale | 1.36E-09 | 0.5073 | -1.81E-09 | 3.62E-09 |
| | | TpCmale | 1.32E-09 | 0.9463 | -2.55E-09 | 2.73E-09 |
| | VehMale | VehFemale | 1.58E-09 | 0.9633 | -3.08E-09 | 3.23E-09 |
| | | CBDfemale | 1.53E-09 | 0.3719 | -1.68E-09 | 4.43E-09 |
| | | THCfemale | 1.49E-09 | 0.2407 | -1.21E-09 | 4.74E-09 |
| | | TpCfemale | 1.46E-09 | 0.9960 | -2.92E-09 | 2.90E-09 |
| | | CBDmale | 1.46E-09 | 0.8928 | -2.71E-09 | 3.11E-09 |
| | | THCmale | 1.49E-09 | 0.5486 | -2.08E-09 | 3.87E-09 |
| | | TpCmale | 1.46E-09 | 0.9552 | -2.83E-09 | 2.99E-09 |
| | CBDmale | VehFemale | 1.46E-09 | 0.9324 | -3.03E-09 | 2.79E-09 |
| | | CBDfemale | 1.40E-09 | 0.4037 | -1.62E-09 | 3.98E-09 |
| | | THCfemale | 1.36E-09 | 0.2529 | -1.15E-09 | 4.28E-09 |
| | | TpCfemale | 1.32E-09 | 0.8776 | -2.85E-09 | 2.44E-09 |
| | | VehMale | 1.46E-09 | 0.8928 | -3.11E-09 | 2.71E-09 |
| | | THCmale | 1.36E-09 | 0.6074 | -2.01E-09 | 3.41E-09 |
| | | TpCmale | 1.32E-09 | 0.9310 | -2.76E-09 | 2.53E-09 |
| | THCmale | VehFemale | 1.49E-09 | 0.5814 | -3.80E-09 | 2.15E-09 |
| | | CBDfemale | 1.44E-09 | 0.7407 | -2.39E-09 | 3.35E-09 |
| | | THCfemale | 1.39E-09 | 0.5365 | -1.92E-09 | 3.65E-09 |
| | | TpCfemale | 1.36E-09 | 0.5073 | -3.62E-09 | 1.81E-09 |
| | | VehMale | 1.49E-09 | 0.5486 | -3.87E-09 | 2.08E-09 |
| | | CBDmale | 1.36E-09 | 0.6074 | -3.41E-09 | 2.01E-09 |
| | | TpCmale | 1.36E-09 | 0.5500 | -3.53E-09 | 1.90E-09 |
| | TpCmale | VehFemale | 1.46E-09 | 0.9950 | -2.92E-09 | 2.90E-09 |
| | | CBDfemale | 1.40E-09 | 0.3597 | -1.51E-09 | 4.09E-09 |
| | | THCfemale | 1.36E-09 | 0.2201 | -1.03E-09 | 4.39E-09 |
| | | TpCfemale | 1.32E-09 | 0.9463 | -2.73E-09 | 2.55E-09 |
| | | VehMale | 1.46E-09 | 0.9552 | -2.99E-09 | 2.83E-09 |
| | | CBDmale | 1.32E-09 | 0.9310 | -2.53E-09 | 2.76E-09 |
| | | THCmale | 1.36E-09 | 0.5500 | -1.90E-09 | 3.53E-09 |
| *N*-stearoyl alanine | VehFemale | CBDfemale | 4.15E-09 | 0.9978 | -8.29E-09 | 8.31E-09 |

| | | | | | | |
|---|---|---|---|---|---|---|
| | | THCfemale | 4.15E-09 | 0.3809 | -1.20E-08 | 4.63E-09 |
| | | TpCfemale | 4.07E-09 | 0.8604 | -8.85E-09 | 7.41E-09 |
| | | VehMale | 4.38E-09 | 0.4374 | -5.33E-09 | 1.22E-08 |
| | | CBDmale | 4.00E-09 | 0.8301 | -8.85E-09 | 7.13E-09 |
| | | THCmale | 4.25E-09 | 0.2500 | -3.56E-09 | 1.34E-08 |
| | | TpCmale | 4.15E-09 | 0.3010 | -3.97E-09 | 1.26E-08 |
| | CBDfemale | VehFemale | 4.15E-09 | 0.9978 | -8.31E-09 | 8.29E-09 |
| | | THCfemale | 3.71E-09 | 0.3261 | -1.11E-08 | 3.75E-09 |
| | | TpCfemale | 3.62E-09 | 0.8408 | -7.96E-09 | 6.50E-09 |
| | | VehMale | 3.97E-09 | 0.3931 | -4.52E-09 | 1.13E-08 |
| | | CBDmale | 3.54E-09 | 0.8061 | -7.95E-09 | 6.20E-09 |
| | | THCmale | 3.83E-09 | 0.2027 | -2.72E-09 | 1.26E-08 |
| | | TpCmale | 3.71E-09 | 0.2493 | -3.10E-09 | 1.17E-08 |
| | THCfemale | VehFemale | 4.15E-09 | 0.3809 | -4.63E-09 | 1.20E-08 |
| | | CBDfemale | 3.71E-09 | 0.3261 | -3.75E-09 | 1.11E-08 |
| | | TpCfemale | 3.62E-09 | 0.4188 | -4.29E-09 | 1.02E-08 |
| | | VehMale | 3.97E-09 | 0.0791 | -8.47E-10 | 1.50E-08 |
| | | CBDmale | 3.54E-09 | 0.4316 | -4.27E-09 | 9.88E-09 |
| | | THCmale | 3.83E-09 | 0.0282 | 9.51E-10 | 1.63E-08 |
| | | TpCmale | 3.71E-09 | 0.0353 | 5.69E-10 | 1.54E-08 |
| | TpCfemale | VehFemale | 4.07E-09 | 0.8604 | -7.41E-09 | 8.85E-09 |
| | | CBDfemale | 3.62E-09 | 0.8408 | -6.50E-09 | 7.96E-09 |
| | | THCfemale | 3.62E-09 | 0.4188 | -1.02E-08 | 4.29E-09 |
| | | VehMale | 3.88E-09 | 0.2899 | -3.62E-09 | 1.19E-08 |
| | | CBDmale | 3.44E-09 | 0.9670 | -7.02E-09 | 6.74E-09 |
| | | THCmale | 3.73E-09 | 0.1350 | -1.81E-09 | 1.31E-08 |
| | | TpCmale | 3.62E-09 | 0.1681 | -2.19E-09 | 1.23E-08 |
| | VehMale | VehFemale | 4.38E-09 | 0.4374 | -1.22E-08 | 5.33E-09 |
| | | CBDfemale | 3.97E-09 | 0.3931 | -1.13E-08 | 4.52E-09 |
| | | THCfemale | 3.97E-09 | 0.0791 | -1.50E-08 | 8.47E-10 |
| | | TpCfemale | 3.88E-09 | 0.2899 | -1.19E-08 | 3.62E-09 |
| | | CBDmale | 3.81E-09 | 0.2647 | -1.19E-08 | 3.33E-09 |
| | | THCmale | 4.07E-09 | 0.7114 | -6.63E-09 | 9.66E-09 |
| | | TpCmale | 3.97E-09 | 0.8205 | -7.03E-09 | 8.84E-09 |
| | CBDmale | VehFemale | 4.00E-09 | 0.8301 | -7.13E-09 | 8.85E-09 |
| | | CBDfemale | 3.54E-09 | 0.8061 | -6.20E-09 | 7.95E-09 |
| | | THCfemale | 3.54E-09 | 0.4316 | -9.88E-09 | 4.27E-09 |
| | | TpCfemale | 3.44E-09 | 0.9670 | -6.74E-09 | 7.02E-09 |
| | | VehMale | 3.81E-09 | 0.2647 | -3.33E-09 | 1.19E-08 |
| | | THCmale | 3.66E-09 | 0.1181 | -1.52E-09 | 1.31E-08 |
| | | TpCmale | 3.54E-09 | 0.1477 | -1.89E-09 | 1.23E-08 |
| | THCmale | VehFemale | 4.25E-09 | 0.2500 | -1.34E-08 | 3.56E-09 |
| | | CBDfemale | 3.83E-09 | 0.2027 | -1.26E-08 | 2.72E-09 |
| | | THCfemale | 3.83E-09 | 0.0282 | -1.63E-08 | -9.51E-10 |
| | | TpCfemale | 3.73E-09 | 0.1350 | -1.31E-08 | 1.81E-09 |
| | | VehMale | 4.07E-09 | 0.7114 | -9.66E-09 | 6.63E-09 |
| | | CBDmale | 3.66E-09 | 0.1181 | -1.31E-08 | 1.52E-09 |
| | | TpCmale | 3.83E-09 | 0.8738 | -8.26E-09 | 7.04E-09 |
| | TpCmale | VehFemale | 4.15E-09 | 0.3010 | -1.26E-08 | 3.97E-09 |
| | | CBDfemale | 3.71E-09 | 0.2493 | -1.17E-08 | 3.10E-09 |
| | | THCfemale | 3.71E-09 | 0.0353 | -1.54E-08 | -5.69E-10 |
| | | TpCfemale | 3.62E-09 | 0.1681 | -1.23E-08 | 2.19E-09 |
| | | VehMale | 3.97E-09 | 0.8205 | -8.84E-09 | 7.03E-09 |
| | | CBDmale | 3.54E-09 | 0.1477 | -1.23E-08 | 1.89E-09 |
| | | THCmale | 3.83E-09 | 0.8738 | -7.04E-09 | 8.26E-09 |
| | VehFemale | CBDfemale | 2.75E-09 | 0.5971 | -6.95E-09 | 4.03E-09 |
| | | THCfemale | 2.75E-09 | 0.3433 | -8.11E-09 | 2.87E-09 |
| *N*-oleoyl alanine | | TpCfemale | 2.64E-09 | 0.9780 | -5.19E-09 | 5.34E-09 |
| | | VehMale | 2.82E-09 | 0.1101 | -1.02E-08 | 1.06E-09 |
| | | CBDmale | 2.69E-09 | 0.3792 | -7.75E-09 | 2.99E-09 |

| | | | | | | |
|---|---|---|---|---|---|---|
| | | THCmale | 2.75E-09 | 0.2162 | -8.92E-09 | 2.06E-09 |
| | | TpCmale | 2.75E-09 | 0.8335 | -6.07E-09 | 4.91E-09 |
| | CBDfemale | VehFemale | 2.75E-09 | 0.5971 | -4.03E-09 | 6.95E-09 |
| | | THCfemale | 2.57E-09 | 0.6523 | -6.30E-09 | 3.97E-09 |
| | | TpCfemale | 2.45E-09 | 0.5339 | -3.36E-09 | 6.43E-09 |
| | | VehMale | 2.65E-09 | 0.2443 | -8.41E-09 | 2.18E-09 |
| | | CBDmale | 2.51E-09 | 0.7148 | -5.92E-09 | 4.09E-09 |
| | | THCmale | 2.57E-09 | 0.4457 | -7.11E-09 | 3.16E-09 |
| | | TpCmale | 2.57E-09 | 0.7332 | -4.25E-09 | 6.01E-09 |
| | THCfemale | VehFemale | 2.75E-09 | 0.3433 | -2.87E-09 | 8.11E-09 |
| | | CBDfemale | 2.57E-09 | 0.6523 | -3.97E-09 | 6.30E-09 |
| | | TpCfemale | 2.45E-09 | 0.2754 | -2.20E-09 | 7.59E-09 |
| | | VehMale | 2.65E-09 | 0.4645 | -7.24E-09 | 3.34E-09 |
| | | CBDmale | 2.51E-09 | 0.9227 | -4.76E-09 | 5.25E-09 |
| | | THCmale | 2.57E-09 | 0.7540 | -5.94E-09 | 4.33E-09 |
| | | TpCmale | 2.57E-09 | 0.4295 | -3.09E-09 | 7.18E-09 |
| | TpCfemale | VehFemale | 2.64E-09 | 0.9780 | -5.34E-09 | 5.19E-09 |
| | | CBDfemale | 2.45E-09 | 0.5339 | -6.43E-09 | 3.36E-09 |
| | | THCfemale | 2.45E-09 | 0.2754 | -7.59E-09 | 2.20E-09 |
| | | VehMale | 2.53E-09 | 0.0714 | -9.71E-09 | 4.16E-10 |
| | | CBDmale | 2.38E-09 | 0.3072 | -7.21E-09 | 2.31E-09 |
| | | THCmale | 2.45E-09 | 0.1575 | -8.40E-09 | 1.39E-09 |
| | | TpCmale | 2.45E-09 | 0.7908 | -5.55E-09 | 4.24E-09 |
| | VehMale | VehFemale | 2.82E-09 | 0.1101 | -1.06E-09 | 1.02E-08 |
| | | CBDfemale | 2.65E-09 | 0.2443 | -2.18E-09 | 8.41E-09 |
| | | THCfemale | 2.65E-09 | 0.4645 | -3.34E-09 | 7.24E-09 |
| | | TpCfemale | 2.53E-09 | 0.0714 | -4.16E-10 | 9.71E-09 |
| | | CBDmale | 2.59E-09 | 0.3995 | -2.97E-09 | 7.36E-09 |
| | | THCmale | 2.65E-09 | 0.6683 | -4.15E-09 | 6.43E-09 |
| | | TpCmale | 2.65E-09 | 0.1367 | -1.30E-09 | 9.29E-09 |
| | CBDmale | VehFemale | 2.69E-09 | 0.3792 | -2.99E-09 | 7.75E-09 |
| | | CBDfemale | 2.51E-09 | 0.7148 | -4.09E-09 | 5.92E-09 |
| | | THCfemale | 2.51E-09 | 0.9227 | -5.25E-09 | 4.76E-09 |
| | | TpCfemale | 2.38E-09 | 0.3072 | -2.31E-09 | 7.21E-09 |
| | | VehMale | 2.59E-09 | 0.3995 | -7.36E-09 | 2.97E-09 |
| | | THCmale | 2.51E-09 | 0.6757 | -6.06E-09 | 3.95E-09 |
| | | TpCmale | 2.51E-09 | 0.4752 | -3.21E-09 | 6.80E-09 |
| | THCmale | VehFemale | 2.75E-09 | 0.2162 | -2.06E-09 | 8.92E-09 |
| | | CBDfemale | 2.57E-09 | 0.4457 | -3.16E-09 | 7.11E-09 |
| | | THCfemale | 2.57E-09 | 0.7540 | -4.33E-09 | 5.94E-09 |
| | | TpCfemale | 2.45E-09 | 0.1575 | -1.39E-09 | 8.40E-09 |
| | | VehMale | 2.65E-09 | 0.6683 | -6.43E-09 | 4.15E-09 |
| | | CBDmale | 2.51E-09 | 0.6757 | -3.95E-09 | 6.06E-09 |
| | | TpCmale | 2.57E-09 | 0.2713 | -2.28E-09 | 7.99E-09 |
| | TpCmale | VehFemale | 2.75E-09 | 0.8335 | -4.91E-09 | 6.07E-09 |
| | | CBDfemale | 2.57E-09 | 0.7332 | -6.01E-09 | 4.25E-09 |
| | | THCfemale | 2.57E-09 | 0.4295 | -7.18E-09 | 3.09E-09 |
| | | TpCfemale | 2.45E-09 | 0.7908 | -4.24E-09 | 5.55E-09 |
| | | VehMale | 2.65E-09 | 0.1367 | -9.29E-09 | 1.30E-09 |
| | | CBDmale | 2.51E-09 | 0.4752 | -6.80E-09 | 3.21E-09 |
| | | THCmale | 2.57E-09 | 0.2713 | -7.99E-09 | 2.28E-09 |
| | VehFemale | CBDfemale | 1.18E-09 | 0.2183 | -3.83E-09 | 8.93E-10 |
| | | THCfemale | 1.18E-09 | 0.5168 | -3.13E-09 | 1.59E-09 |
| | | TpCfemale | 1.10E-09 | 0.6371 | -1.68E-09 | 2.73E-09 |
| | | VehMale | 1.18E-09 | 0.9181 | -2.24E-09 | 2.48E-09 |
| *N*-linoleoyl alanine | | CBDmale | 1.15E-09 | 0.9352 | -2.39E-09 | 2.21E-09 |
| | | THCmale | 1.15E-09 | 0.7693 | -1.96E-09 | 2.64E-09 |
| | | TpCmale | 1.15E-09 | 0.3436 | -1.20E-09 | 3.40E-09 |
| | CBDfemale | VehFemale | 1.18E-09 | 0.2183 | -8.93E-10 | 3.83E-09 |
| | | THCfemale | 1.14E-09 | 0.5424 | -1.58E-09 | 2.98E-09 |

| | | | | | |
|---|---|---|---|---|---|
| | TpCfemale | 1.06E-09 | 0.0650 | -1.28E-10 | 4.11E-09 |
| | VehMale | 1.14E-09 | 0.1682 | -6.91E-10 | 3.87E-09 |
| | CBDmale | 1.11E-09 | 0.2196 | -8.42E-10 | 3.59E-09 |
| | THCmale | 1.11E-09 | 0.1081 | -4.09E-10 | 4.03E-09 |
| | TpCmale | 1.11E-09 | 0.0240 | 3.50E-10 | 4.79E-09 |
| THCfemale | VehFemale | 1.18E-09 | 0.5168 | -1.59E-09 | 3.13E-09 |
| | CBDfemale | 1.14E-09 | 0.5424 | -2.98E-09 | 1.58E-09 |
| | TpCfemale | 1.06E-09 | 0.2273 | -8.27E-10 | 3.41E-09 |
| | VehMale | 1.14E-09 | 0.4373 | -1.39E-09 | 3.17E-09 |
| | CBDmale | 1.11E-09 | 0.5442 | -1.54E-09 | 2.89E-09 |
| | THCmale | 1.11E-09 | 0.3212 | -1.11E-09 | 3.33E-09 |
| | TpCmale | 1.11E-09 | 0.0972 | -3.49E-10 | 4.09E-09 |
| TpCfemale | VehFemale | 1.10E-09 | 0.6371 | -2.73E-09 | 1.68E-09 |
| | CBDfemale | 1.06E-09 | 0.0650 | -4.11E-09 | 1.28E-10 |
| | THCfemale | 1.06E-09 | 0.2273 | -3.41E-09 | 8.27E-10 |
| | VehMale | 1.06E-09 | 0.7065 | -2.52E-09 | 1.72E-09 |
| | CBDmale | 1.03E-09 | 0.5497 | -2.67E-09 | 1.43E-09 |
| | THCmale | 1.03E-09 | 0.8580 | -2.24E-09 | 1.87E-09 |
| | TpCmale | 1.03E-09 | 0.5774 | -1.48E-09 | 2.63E-09 |
| VehMale | VehFemale | 1.18E-09 | 0.9181 | -2.48E-09 | 2.24E-09 |
| | CBDfemale | 1.14E-09 | 0.1682 | -3.87E-09 | 6.91E-10 |
| | THCfemale | 1.14E-09 | 0.4373 | -3.17E-09 | 1.39E-09 |
| | TpCfemale | 1.06E-09 | 0.7065 | -1.72E-09 | 2.52E-09 |
| | CBDmale | 1.11E-09 | 0.8463 | -2.43E-09 | 2.00E-09 |
| | THCmale | 1.11E-09 | 0.8456 | -2.00E-09 | 2.43E-09 |
| | TpCmale | 1.11E-09 | 0.3823 | -1.24E-09 | 3.19E-09 |
| CBDmale | VehFemale | 1.15E-09 | 0.9352 | -2.21E-09 | 2.39E-09 |
| | CBDfemale | 1.11E-09 | 0.2196 | -3.59E-09 | 8.42E-10 |
| | THCfemale | 1.11E-09 | 0.5442 | -2.89E-09 | 1.54E-09 |
| | TpCfemale | 1.03E-09 | 0.5497 | -1.43E-09 | 2.67E-09 |
| | VehMale | 1.11E-09 | 0.8463 | -2.00E-09 | 2.43E-09 |
| | THCmale | 1.08E-09 | 0.6889 | -1.72E-09 | 2.58E-09 |
| | TpCmale | 1.08E-09 | 0.2723 | -9.60E-10 | 3.34E-09 |
| THCmale | VehFemale | 1.15E-09 | 0.7693 | -2.64E-09 | 1.96E-09 |
| | CBDfemale | 1.11E-09 | 0.1081 | -4.03E-09 | 4.09E-10 |
| | THCfemale | 1.11E-09 | 0.3212 | -3.33E-09 | 1.11E-09 |
| | TpCfemale | 1.03E-09 | 0.8580 | -1.87E-09 | 2.24E-09 |
| | VehMale | 1.11E-09 | 0.8456 | -2.43E-09 | 2.00E-09 |
| | CBDmale | 1.08E-09 | 0.6889 | -2.58E-09 | 1.72E-09 |
| | TpCmale | 1.08E-09 | 0.4832 | -1.39E-09 | 2.91E-09 |
| TpCmale | VehFemale | 1.15E-09 | 0.3436 | -3.40E-09 | 1.20E-09 |
| | CBDfemale | 1.11E-09 | 0.0240 | -4.79E-09 | -3.50E-10 |
| | THCfemale | 1.11E-09 | 0.0972 | -4.09E-09 | 3.49E-10 |
| | TpCfemale | 1.03E-09 | 0.5774 | -2.63E-09 | 1.48E-09 |
| | VehMale | 1.11E-09 | 0.3823 | -3.19E-09 | 1.24E-09 |
| | CBDmale | 1.08E-09 | 0.2723 | -3.34E-09 | 9.60E-10 |
| | THCmale | 1.08E-09 | 0.4832 | -2.91E-09 | 1.39E-09 |

#### Supplemental Table 9.8 Fisher's LSD post-hoc test for liver *N*-acyl alanine ANOVA

| | (I) drugVsex | | Std. Error | Sig. | 95% Confidence Interval Lower Bound | Upper Bound |
|---|---|---|---|---|---|---|
| *N*-palmitoyl alanine | VehFemale | CBDfemale | 5.24811E-11 | 0.407 | -1.4875E-10 | 6.1134E-11 |
| | | THCfemale | 4.83851E-11 | 0.074 | -1.8461E-10 | 8.8942E-12 |
| | | TpCfemale | 4.94796E-11 | 0.019 | -2.1864E-10 | -2.0755E-11 |
| | | VehMale | 5.24811E-11 | 0.992 | -1.0439E-10 | 1.0549E-10 |
| | | CBDmale | 4.83851E-11 | 0.423 | -5.7686E-11 | 1.3582E-10 |
| | | THCmale | 4.94796E-11 | 0.013 | -2.2542E-10 | -2.7543E-11 |
| | | TpCmale | 4.83851E-11 | 0.667 | -7.5810E-11 | 1.1769E-10 |
| | CBDfemale | VehFemale | 5.24811E-11 | 0.407 | -6.1134E-11 | 1.4875E-10 |
| | | THCfemale | 4.83851E-11 | 0.366 | -1.4080E-10 | 5.2702E-11 |
| | | TpCfemale | 4.94796E-11 | 0.130 | -1.7483E-10 | 2.3053E-11 |
| | | VehMale | 5.24811E-11 | 0.401 | -6.0587E-11 | 1.4930E-10 |
| | | CBDmale | 4.83851E-11 | 0.092 | -1.3878E-11 | 1.7963E-10 |
| | | THCmale | 4.94796E-11 | 0.100 | -1.8162E-10 | 1.6265E-11 |
| | | TpCmale | 4.83851E-11 | 0.186 | -3.2002E-11 | 1.6150E-10 |
| | THCfemale | VehFemale | 4.83851E-11 | 0.074 | -8.8942E-12 | 1.8461E-10 |
| | | CBDfemale | 4.83851E-11 | 0.366 | -5.2702E-11 | 1.4080E-10 |
| | | TpCfemale | 4.51120E-11 | 0.483 | -1.2205E-10 | 5.8369E-11 |
| | | VehMale | 4.83851E-11 | 0.073 | -8.3463E-12 | 1.8516E-10 |
| | | CBDmale | 4.39088E-11 | 0.005 | 3.9123E-11 | 2.1473E-10 |
| | | THCmale | 4.51120E-11 | 0.395 | -1.2883E-10 | 5.1582E-11 |
| | | TpCmale | 4.39088E-11 | 0.016 | 2.0999E-11 | 1.9660E-10 |
| | TpCfemale | VehFemale | 4.94796E-11 | 0.019 | 2.0755E-11 | 2.1864E-10 |
| | | CBDfemale | 4.94796E-11 | 0.130 | -2.3053E-11 | 1.7483E-10 |
| | | THCfemale | 4.51120E-11 | 0.483 | -5.8369E-11 | 1.2205E-10 |
| | | VehMale | 4.94796E-11 | 0.018 | 2.1303E-11 | 2.1918E-10 |
| | | CBDmale | 4.51120E-11 | 0.001 | 6.8555E-11 | 2.4897E-10 |
| | | THCmale | 4.62839E-11 | 0.884 | -9.9338E-11 | 8.5763E-11 |
| | | TpCmale | 4.51120E-11 | 0.003 | 5.0431E-11 | 2.3084E-10 |
| | VehMale | VehFemale | 5.24811E-11 | 0.992 | -1.0549E-10 | 1.0439E-10 |
| | | CBDfemale | 5.24811E-11 | 0.401 | -1.4930E-10 | 6.0587E-11 |
| | | THCfemale | 4.83851E-11 | 0.073 | -1.8516E-10 | 8.3463E-12 |
| | | TpCfemale | 4.94796E-11 | 0.018 | -2.1918E-10 | -2.1303E-11 |
| | | CBDmale | 4.83851E-11 | 0.429 | -5.8234E-11 | 1.3527E-10 |
| | | THCmale | 4.94796E-11 | 0.013 | -2.2597E-10 | -2.8091E-11 |
| | | TpCmale | 4.83851E-11 | 0.675 | -7.6358E-11 | 1.1715E-10 |
| | CBDmale | VehFemale | 4.83851E-11 | 0.423 | -1.3582E-10 | 5.7686E-11 |
| | | CBDfemale | 4.83851E-11 | 0.092 | -1.7963E-10 | 1.3878E-11 |
| | | THCfemale | 4.39088E-11 | 0.005 | -2.1473E-10 | -3.9123E-11 |
| | | TpCfemale | 4.51120E-11 | 0.001 | -2.4897E-10 | -6.8555E-11 |
| | | VehMale | 4.83851E-11 | 0.429 | -1.3527E-10 | 5.8234E-11 |
| | | THCmale | 4.51120E-11 | 0.001 | -2.5576E-10 | -7.5343E-11 |
| | | TpCmale | 4.39088E-11 | 0.681 | -1.0593E-10 | 6.9677E-11 |
| | THCmale | VehFemale | 4.94796E-11 | 0.013 | 2.7543E-11 | 2.2542E-10 |
| | | CBDfemale | 4.94796E-11 | 0.100 | -1.6265E-11 | 1.8162E-10 |
| | | THCfemale | 4.51120E-11 | 0.395 | -5.1582E-11 | 1.2883E-10 |
| | | TpCfemale | 4.62839E-11 | 0.884 | -8.5763E-11 | 9.9338E-11 |
| | | VehMale | 4.94796E-11 | 0.013 | 2.8091E-11 | 2.2597E-10 |
| | | CBDmale | 4.51120E-11 | 0.001 | 7.5343E-11 | 2.5576E-10 |
| | | TpCmale | 4.51120E-11 | 0.002 | 5.7218E-11 | 2.3763E-10 |
| | TpCmale | VehFemale | 4.83851E-11 | 0.667 | -1.1769E-10 | 7.5810E-11 |
| | | CBDfemale | 4.83851E-11 | 0.186 | -1.6150E-10 | 3.2002E-11 |
| | | THCfemale | 4.39088E-11 | 0.016 | -1.9660E-10 | -2.0999E-11 |
| | | TpCfemale | 4.51120E-11 | 0.003 | -2.3084E-10 | -5.0431E-11 |
| | | VehMale | 4.83851E-11 | 0.675 | -1.1715E-10 | 7.6358E-11 |
| | | CBDmale | 4.39088E-11 | 0.681 | -6.9677E-11 | 1.0593E-10 |
| | | THCmale | 4.51120E-11 | 0.002 | -2.3763E-10 | -5.7218E-11 |
| *N*-stearoyl alanine | VehFemale | CBDfemale | 1.51072E-11 | 0.010 | -7.0412E-11 | -9.9313E-12 |

| | | | | | | |
|---|---|---|---|---|---|---|
| | | THCfemale | 1.47103E-11 | 0.271 | -4.5792E-11 | 1.3100E-11 |
| | | TpCfemale | 1.47103E-11 | 0.321 | -4.4165E-11 | 1.4727E-11 |
| | | VehMale | 1.56026E-11 | 0.886 | -3.3485E-11 | 2.8979E-11 |
| | | CBDmale | 1.47103E-11 | 0.629 | -2.2310E-11 | 3.6582E-11 |
| | | THCmale | 1.51072E-11 | 0.715 | -2.4692E-11 | 3.5789E-11 |
| | | TpCmale | 1.47103E-11 | 0.764 | -2.5016E-11 | 3.3876E-11 |
| | CBDfemale | VehFemale | 1.51072E-11 | 0.010 | 9.9313E-12 | 7.0412E-11 |
| | | THCfemale | 1.41837E-11 | 0.098 | -4.5658E-12 | 5.2218E-11 |
| | | TpCfemale | 1.41837E-11 | 0.078 | -2.9390E-12 | 5.3845E-11 |
| | | VehMale | 1.51072E-11 | 0.015 | 7.6782E-12 | 6.8159E-11 |
| | | CBDmale | 1.41837E-11 | 0.001 | 1.8916E-11 | 7.5700E-11 |
| | | THCmale | 1.45949E-11 | 0.003 | 1.6505E-11 | 7.4935E-11 |
| | | TpCmale | 1.41837E-11 | 0.003 | 1.6210E-11 | 7.2993E-11 |
| | THCfemale | VehFemale | 1.47103E-11 | 0.271 | -1.3100E-11 | 4.5792E-11 |
| | | CBDfemale | 1.41837E-11 | 0.098 | -5.2218E-11 | 4.5658E-12 |
| | | TpCfemale | 1.37602E-11 | 0.906 | -2.5917E-11 | 2.9171E-11 |
| | | VehMale | 1.47103E-11 | 0.342 | -1.5353E-11 | 4.3538E-11 |
| | | CBDmale | 1.37602E-11 | 0.093 | -4.0622E-12 | 5.1026E-11 |
| | | THCmale | 1.41837E-11 | 0.128 | -6.4976E-12 | 5.0286E-11 |
| | | TpCmale | 1.37602E-11 | 0.137 | -6.7685E-12 | 4.8320E-11 |
| | TpCfemale | VehFemale | 1.47103E-11 | 0.321 | -1.4727E-11 | 4.4165E-11 |
| | | CBDfemale | 1.41837E-11 | 0.078 | -5.3845E-11 | 2.9390E-12 |
| | | THCfemale | 1.37602E-11 | 0.906 | -2.9171E-11 | 2.5917E-11 |
| | | VehMale | 1.47103E-11 | 0.400 | -1.6980E-11 | 4.1912E-11 |
| | | CBDmale | 1.37602E-11 | 0.118 | -5.6890E-12 | 4.9399E-11 |
| | | THCmale | 1.41837E-11 | 0.158 | -8.1245E-12 | 4.8659E-11 |
| | | TpCmale | 1.37602E-11 | 0.169 | -8.3954E-12 | 4.6693E-11 |
| | VehMale | VehFemale | 1.56026E-11 | 0.886 | -2.8979E-11 | 3.3485E-11 |
| | | CBDfemale | 1.51072E-11 | 0.015 | -6.8159E-11 | -7.6782E-12 |
| | | THCfemale | 1.47103E-11 | 0.342 | -4.3538E-11 | 1.5353E-11 |
| | | TpCfemale | 1.47103E-11 | 0.400 | -4.1912E-11 | 1.6980E-11 |
| | | CBDmale | 1.47103E-11 | 0.526 | -2.0057E-11 | 3.8835E-11 |
| | | THCmale | 1.51072E-11 | 0.608 | -2.2439E-11 | 3.8042E-11 |
| | | TpCmale | 1.47103E-11 | 0.651 | -2.2763E-11 | 3.6129E-11 |
| | CBDmale | VehFemale | 1.47103E-11 | 0.629 | -3.6582E-11 | 2.2310E-11 |
| | | CBDfemale | 1.41837E-11 | 0.001 | -7.5700E-11 | -1.8916E-11 |
| | | THCfemale | 1.37602E-11 | 0.093 | -5.1026E-11 | 4.0622E-12 |
| | | TpCfemale | 1.37602E-11 | 0.118 | -4.9399E-11 | 5.6890E-12 |
| | | VehMale | 1.47103E-11 | 0.526 | -3.8835E-11 | 2.0057E-11 |
| | | THCmale | 1.41837E-11 | 0.911 | -2.9980E-11 | 2.6804E-11 |
| | | TpCmale | 1.37602E-11 | 0.845 | -3.0250E-11 | 2.4838E-11 |
| | THCmale | VehFemale | 1.51072E-11 | 0.715 | -3.5789E-11 | 2.4692E-11 |
| | | CBDfemale | 1.45949E-11 | 0.003 | -7.4935E-11 | -1.6505E-11 |
| | | THCfemale | 1.41837E-11 | 0.128 | -5.0286E-11 | 6.4976E-12 |
| | | TpCfemale | 1.41837E-11 | 0.158 | -4.8659E-11 | 8.1245E-12 |
| | | VehMale | 1.51072E-11 | 0.608 | -3.8042E-11 | 2.2439E-11 |
| | | CBDmale | 1.41837E-11 | 0.911 | -2.6804E-11 | 2.9980E-11 |
| | | TpCmale | 1.41837E-11 | 0.937 | -2.9510E-11 | 2.7273E-11 |
| | TpCmale | VehFemale | 1.47103E-11 | 0.764 | -3.3876E-11 | 2.5016E-11 |
| | | CBDfemale | 1.41837E-11 | 0.003 | -7.2993E-11 | -1.6210E-11 |
| | | THCfemale | 1.37602E-11 | 0.137 | -4.8320E-11 | 6.7685E-12 |
| | | TpCfemale | 1.37602E-11 | 0.169 | -4.6693E-11 | 8.3954E-12 |
| | | VehMale | 1.47103E-11 | 0.651 | -3.6129E-11 | 2.2763E-11 |
| | | CBDmale | 1.37602E-11 | 0.845 | -2.4838E-11 | 3.0250E-11 |
| | | THCmale | 1.41837E-11 | 0.937 | -2.7273E-11 | 2.9510E-11 |
| | VehFemale | CBDfemale | 5.05223E-11 | 0.595 | -1.2797E-10 | 7.4014E-11 |
| | | THCfemale | 4.65793E-11 | 0.000 | -2.7042E-10 | -8.4202E-11 |
| *N*-oleoyl alanine | | TpCfemale | 4.65793E-11 | 0.012 | -2.1403E-10 | -2.7806E-11 |
| | | VehMale | 5.05223E-11 | 0.928 | -9.6379E-11 | 1.0561E-10 |
| | | CBDmale | 4.65793E-11 | 0.762 | -7.8918E-11 | 1.0730E-10 |

| | | | | | | |
|---|---|---|---|---|---|---|
| | | THCmale | 4.89180E-11 | 0.614 | -1.2259E-10 | 7.2978E-11 |
| | | TpCmale | 4.56992E-11 | 0.164 | -1.5571E-10 | 2.6996E-11 |
| | CBDfemale | VehFemale | 5.05223E-11 | 0.595 | -7.4014E-11 | 1.2797E-10 |
| | | THCfemale | 4.65793E-11 | 0.002 | -2.4344E-10 | -5.7223E-11 |
| | | TpCfemale | 4.65793E-11 | 0.048 | -1.8705E-10 | -8.2781E-13 |
| | | VehMale | 5.05223E-11 | 0.534 | -6.9400E-11 | 1.3259E-10 |
| | | CBDmale | 4.65793E-11 | 0.380 | -5.1939E-11 | 1.3428E-10 |
| | | THCmale | 4.89180E-11 | 0.965 | -9.5615E-11 | 9.9957E-11 |
| | | TpCmale | 4.56992E-11 | 0.417 | -1.2873E-10 | 5.3975E-11 |
| | THCfemale | VehFemale | 4.65793E-11 | 0.000 | 8.4202E-11 | 2.7042E-10 |
| | | CBDfemale | 4.65793E-11 | 0.002 | 5.7223E-11 | 2.4344E-10 |
| | | TpCfemale | 4.22700E-11 | 0.187 | -2.8101E-11 | 1.4089E-10 |
| | | VehMale | 4.65793E-11 | 0.000 | 8.8816E-11 | 2.7504E-10 |
| | | CBDmale | 4.22700E-11 | 0.000 | 1.0701E-10 | 2.7600E-10 |
| | | THCmale | 4.48341E-11 | 0.001 | 6.2883E-11 | 2.4213E-10 |
| | | TpCmale | 4.12982E-11 | 0.008 | 3.0404E-11 | 1.9551E-10 |
| | TpCfemale | VehFemale | 4.65793E-11 | 0.012 | 2.7806E-11 | 2.1403E-10 |
| | | CBDfemale | 4.65793E-11 | 0.048 | 8.2781E-13 | 1.8705E-10 |
| | | THCfemale | 4.22700E-11 | 0.187 | -1.4089E-10 | 2.8101E-11 |
| | | VehMale | 4.65793E-11 | 0.009 | 3.2420E-11 | 2.1864E-10 |
| | | CBDmale | 4.22700E-11 | 0.002 | 5.0613E-11 | 2.1961E-10 |
| | | THCmale | 4.48341E-11 | 0.036 | 6.4876E-12 | 1.8573E-10 |
| | | TpCmale | 4.12982E-11 | 0.176 | -2.5992E-11 | 1.3912E-10 |
| | VehMale | VehFemale | 5.05223E-11 | 0.928 | -1.0561E-10 | 9.6379E-11 |
| | | CBDfemale | 5.05223E-11 | 0.534 | -1.3259E-10 | 6.9400E-11 |
| | | THCfemale | 4.65793E-11 | 0.000 | -2.7504E-10 | -8.8816E-11 |
| | | TpCfemale | 4.65793E-11 | 0.009 | -2.1864E-10 | -3.2420E-11 |
| | | CBDmale | 4.65793E-11 | 0.838 | -8.3532E-11 | 1.0269E-10 |
| | | THCmale | 4.89180E-11 | 0.550 | -1.2721E-10 | 6.8364E-11 |
| | | TpCmale | 4.56992E-11 | 0.136 | -1.6032E-10 | 2.2382E-11 |
| | CBDmale | VehFemale | 4.65793E-11 | 0.762 | -1.0730E-10 | 7.8918E-11 |
| | | CBDfemale | 4.65793E-11 | 0.380 | -1.3428E-10 | 5.1939E-11 |
| | | THCfemale | 4.22700E-11 | 0.000 | -2.7600E-10 | -1.0701E-10 |
| | | TpCfemale | 4.22700E-11 | 0.002 | -2.1961E-10 | -5.0613E-11 |
| | | VehMale | 4.65793E-11 | 0.838 | -1.0269E-10 | 8.3532E-11 |
| | | THCmale | 4.48341E-11 | 0.388 | -1.2862E-10 | 5.0622E-11 |
| | | TpCmale | 4.12982E-11 | 0.062 | -1.6110E-10 | 4.0061E-12 |
| | THCmale | VehFemale | 4.89180E-11 | 0.614 | -7.2978E-11 | 1.2259E-10 |
| | | CBDfemale | 4.89180E-11 | 0.965 | -9.9957E-11 | 9.5615E-11 |
| | | THCfemale | 4.48341E-11 | 0.001 | -2.4213E-10 | -6.2883E-11 |
| | | TpCfemale | 4.48341E-11 | 0.036 | -1.8573E-10 | -6.4876E-12 |
| | | VehMale | 4.89180E-11 | 0.550 | -6.8364E-11 | 1.2721E-10 |
| | | CBDmale | 4.48341E-11 | 0.388 | -5.0622E-11 | 1.2862E-10 |
| | | TpCmale | 4.39191E-11 | 0.371 | -1.2734E-10 | 4.8245E-11 |
| | TpCmale | VehFemale | 4.56992E-11 | 0.164 | -2.6996E-11 | 1.5571E-10 |
| | | CBDfemale | 4.56992E-11 | 0.417 | -5.3975E-11 | 1.2873E-10 |
| | | THCfemale | 4.12982E-11 | 0.008 | -1.9551E-10 | -3.0404E-11 |
| | | TpCfemale | 4.12982E-11 | 0.176 | -1.3912E-10 | 2.5992E-11 |
| | | VehMale | 4.56992E-11 | 0.136 | -2.2382E-11 | 1.6032E-10 |
| | | CBDmale | 4.12982E-11 | 0.062 | -4.0061E-12 | 1.6110E-10 |
| | | THCmale | 4.39191E-11 | 0.371 | -4.8245E-11 | 1.2734E-10 |
| *N*-linoleoyl alanine | VehFemale | CBDfemale | 3.63241E-11 | 0.296 | -1.1096E-10 | 3.4311E-11 |
| | | THCfemale | 3.34892E-11 | 0.019 | -1.4782E-10 | -1.3889E-11 |
| | | TpCfemale | 3.34892E-11 | 0.000 | -2.2876E-10 | -9.4827E-11 |
| | | VehMale | 3.63241E-11 | 0.294 | -3.4163E-11 | 1.1111E-10 |
| | | CBDmale | 3.34892E-11 | 0.526 | -4.5623E-11 | 8.8309E-11 |
| | | THCmale | 3.51707E-11 | 0.000 | -2.0354E-10 | -6.2887E-11 |
| | | TpCmale | 3.34892E-11 | 0.441 | -9.2912E-11 | 4.1019E-11 |
| | CBDfemale | VehFemale | 3.63241E-11 | 0.296 | -3.4311E-11 | 1.1096E-10 |
| | | THCfemale | 3.34892E-11 | 0.209 | -1.0950E-10 | 2.4435E-11 |

| | | | | | | |
|---|---|---|---|---|---|---|
| | | TpCfemale | 3.34892E-11 | 0.000 | -1.9043E-10 | -5.6503E-11 |
| | | VehMale | 3.63241E-11 | 0.039 | 4.1609E-12 | 1.4943E-10 |
| | | CBDmale | 3.34892E-11 | 0.080 | -7.2991E-12 | 1.2663E-10 |
| | | THCmale | 3.51707E-11 | 0.009 | -1.6522E-10 | -2.4564E-11 |
| | | TpCmale | 3.34892E-11 | 0.713 | -5.4588E-11 | 7.9343E-11 |
| | THCfemale | VehFemale | 3.34892E-11 | 0.019 | 1.3889E-11 | 1.4782E-10 |
| | | CBDfemale | 3.34892E-11 | 0.209 | -2.4435E-11 | 1.0950E-10 |
| | | TpCfemale | 3.03909E-11 | 0.010 | -1.4171E-10 | -2.0168E-11 |
| | | VehMale | 3.34892E-11 | 0.001 | 5.2360E-11 | 1.8629E-10 |
| | | CBDmale | 3.03909E-11 | 0.001 | 4.1427E-11 | 1.6297E-10 |
| | | THCmale | 3.22344E-11 | 0.109 | -1.1682E-10 | 1.2096E-11 |
| | | TpCmale | 3.03909E-11 | 0.076 | -5.8624E-12 | 1.1568E-10 |
| | TpCfemale | VehFemale | 3.34892E-11 | 0.000 | 9.4827E-11 | 2.2876E-10 |
| | | CBDfemale | 3.34892E-11 | 0.000 | 5.6503E-11 | 1.9043E-10 |
| | | THCfemale | 3.03909E-11 | 0.010 | 2.0168E-11 | 1.4171E-10 |
| | | VehMale | 3.34892E-11 | 0.000 | 1.3330E-10 | 2.6723E-10 |
| | | CBDmale | 3.03909E-11 | 0.000 | 1.2236E-10 | 2.4391E-10 |
| | | THCmale | 3.22344E-11 | 0.379 | -3.5880E-11 | 9.3034E-11 |
| | | TpCmale | 3.03909E-11 | 0.000 | 7.5076E-11 | 1.9662E-10 |
| | VehMale | VehFemale | 3.63241E-11 | 0.294 | -1.1111E-10 | 3.4163E-11 |
| | | CBDfemale | 3.63241E-11 | 0.039 | -1.4943E-10 | -4.1609E-12 |
| | | THCfemale | 3.34892E-11 | 0.001 | -1.8629E-10 | -5.2360E-11 |
| | | TpCfemale | 3.34892E-11 | 0.000 | -2.6723E-10 | -1.3330E-10 |
| | | CBDmale | 3.34892E-11 | 0.611 | -8.4094E-11 | 4.9837E-11 |
| | | THCmale | 3.51707E-11 | 0.000 | -2.4202E-10 | -1.0136E-10 |
| | | TpCmale | 3.34892E-11 | 0.059 | -1.3138E-10 | 2.5477E-12 |
| | CBDmale | VehFemale | 3.34892E-11 | 0.526 | -8.8309E-11 | 4.5623E-11 |
| | | CBDfemale | 3.34892E-11 | 0.080 | -1.2663E-10 | 7.2991E-12 |
| | | THCfemale | 3.03909E-11 | 0.001 | -1.6297E-10 | -4.1427E-11 |
| | | TpCfemale | 3.03909E-11 | 0.000 | -2.4391E-10 | -1.2236E-10 |
| | | VehMale | 3.34892E-11 | 0.611 | -4.9837E-11 | 8.4094E-11 |
| | | THCmale | 3.22344E-11 | 0.000 | -2.1902E-10 | -9.0102E-11 |
| | | TpCmale | 3.03909E-11 | 0.125 | -1.0806E-10 | 1.3481E-11 |
| | THCmale | VehFemale | 3.51707E-11 | 0.000 | 6.2887E-11 | 2.0354E-10 |
| | | CBDfemale | 3.51707E-11 | 0.009 | 2.4564E-11 | 1.6522E-10 |
| | | THCfemale | 3.22344E-11 | 0.109 | -1.2096E-11 | 1.1682E-10 |
| | | TpCfemale | 3.22344E-11 | 0.379 | -9.3034E-11 | 3.5880E-11 |
| | | VehMale | 3.51707E-11 | 0.000 | 1.0136E-10 | 2.4202E-10 |
| | | CBDmale | 3.22344E-11 | 0.000 | 9.0102E-11 | 2.1902E-10 |
| | | TpCmale | 3.22344E-11 | 0.001 | 4.2812E-11 | 1.7173E-10 |
| | TpCmale | VehFemale | 3.34892E-11 | 0.441 | -4.1019E-11 | 9.2912E-11 |
| | | CBDfemale | 3.34892E-11 | 0.713 | -7.9343E-11 | 5.4588E-11 |
| | | THCfemale | 3.03909E-11 | 0.076 | -1.1568E-10 | 5.8624E-12 |
| | | TpCfemale | 3.03909E-11 | 0.000 | -1.9662E-10 | -7.5076E-11 |
| | | VehMale | 3.34892E-11 | 0.059 | -2.5477E-12 | 1.3138E-10 |
| | | CBDmale | 3.03909E-11 | 0.125 | -1.3481E-11 | 1.0806E-10 |
| | | THCmale | 3.22344E-11 | 0.001 | -1.7173E-10 | -4.2812E-11 |
| | VehFemale | CBDfemale | 1.82534E-11 | 0.463 | -4.9982E-11 | 2.3043E-11 |
| | | THCfemale | 1.67295E-11 | 0.009 | -7.8777E-11 | -1.1849E-11 |
| | | TpCfemale | 1.71376E-11 | 0.084 | -6.4367E-11 | 4.1934E-12 |
| | | VehMale | 1.82534E-11 | 0.260 | -1.5734E-11 | 5.7291E-11 |
| | | CBDmale | 1.71376E-11 | 0.198 | -1.1971E-11 | 5.6589E-11 |
| | | THCmale | 1.76345E-11 | 0.022 | -7.6798E-11 | -6.2497E-12 |
| *N*-arachidonoyl alanine | | TpCmale | 1.67295E-11 | 0.830 | -2.9866E-11 | 3.7062E-11 |
| | CBDfemale | VehFemale | 1.82534E-11 | 0.463 | -2.3043E-11 | 4.9982E-11 |
| | | THCfemale | 1.73807E-11 | 0.072 | -6.6610E-11 | 2.9229E-12 |
| | | TpCfemale | 1.77739E-11 | 0.354 | -5.2170E-11 | 1.8936E-11 |
| | | VehMale | 1.88520E-11 | 0.074 | -3.4617E-12 | 7.1958E-11 |
| | | CBDmale | 1.77739E-11 | 0.049 | 2.2577E-13 | 7.1332E-11 |
| | | THCmale | 1.82534E-11 | 0.130 | -6.4566E-11 | 8.4580E-12 |

| | | | | | | |
|---|---|---|---|---|---|---|
| | | TpCmale | 1.73807E-11 | 0.330 | -1.7698E-11 | 5.1835E-11 |
| | THCfemale | VehFemale | 1.67295E-11 | 0.009 | 1.1849E-11 | 7.8777E-11 |
| | | CBDfemale | 1.73807E-11 | 0.072 | -2.9229E-12 | 6.6610E-11 |
| | | TpCfemale | 1.62049E-11 | 0.351 | -1.7188E-11 | 4.7641E-11 |
| | | VehMale | 1.73807E-11 | 0.000 | 3.1325E-11 | 1.0086E-10 |
| | | CBDmale | 1.62049E-11 | 0.000 | 3.5208E-11 | 1.0004E-10 |
| | | THCmale | 1.67295E-11 | 0.822 | -2.9675E-11 | 3.7254E-11 |
| | | TpCmale | 1.57727E-11 | 0.003 | 1.7362E-11 | 8.0462E-11 |
| | TpCfemale | VehFemale | 1.71376E-11 | 0.084 | -4.1934E-12 | 6.4367E-11 |
| | | CBDfemale | 1.77739E-11 | 0.354 | -1.8936E-11 | 5.2170E-11 |
| | | THCfemale | 1.62049E-11 | 0.351 | -4.7641E-11 | 1.7188E-11 |
| | | VehMale | 1.77739E-11 | 0.006 | 1.5312E-11 | 8.6418E-11 |
| | | CBDmale | 1.66259E-11 | 0.003 | 1.9139E-11 | 8.5653E-11 |
| | | THCmale | 1.71376E-11 | 0.507 | -4.5717E-11 | 2.2843E-11 |
| | | TpCmale | 1.62049E-11 | 0.042 | 1.2707E-12 | 6.6100E-11 |
| | VehMale | VehFemale | 1.82534E-11 | 0.260 | -5.7291E-11 | 1.5734E-11 |
| | | CBDfemale | 1.88520E-11 | 0.074 | -7.1958E-11 | 3.4617E-12 |
| | | THCfemale | 1.73807E-11 | 0.000 | -1.0086E-10 | -3.1325E-11 |
| | | TpCfemale | 1.77739E-11 | 0.006 | -8.6418E-11 | -1.5312E-11 |
| | | CBDmale | 1.77739E-11 | 0.932 | -3.4022E-11 | 3.7084E-11 |
| | | THCmale | 1.82534E-11 | 0.001 | -9.8814E-11 | -2.5790E-11 |
| | | TpCmale | 1.73807E-11 | 0.327 | -5.1946E-11 | 1.7587E-11 |
| | CBDmale | VehFemale | 1.71376E-11 | 0.198 | -5.6589E-11 | 1.1971E-11 |
| | | CBDfemale | 1.77739E-11 | 0.049 | -7.1332E-11 | -2.2577E-13 |
| | | THCfemale | 1.62049E-11 | 0.000 | -1.0004E-10 | -3.5208E-11 |
| | | TpCfemale | 1.66259E-11 | 0.003 | -8.5653E-11 | -1.9139E-11 |
| | | VehMale | 1.77739E-11 | 0.932 | -3.7084E-11 | 3.4022E-11 |
| | | THCmale | 1.71376E-11 | 0.000 | -9.8113E-11 | -2.9553E-11 |
| | | TpCmale | 1.62049E-11 | 0.253 | -5.1125E-11 | 1.3704E-11 |
| | THCmale | VehFemale | 1.76345E-11 | 0.022 | 6.2497E-12 | 7.6798E-11 |
| | | CBDfemale | 1.82534E-11 | 0.130 | -8.4580E-12 | 6.4566E-11 |
| | | THCfemale | 1.67295E-11 | 0.822 | -3.7254E-11 | 2.9675E-11 |
| | | TpCfemale | 1.71376E-11 | 0.507 | -2.2843E-11 | 4.5717E-11 |
| | | VehMale | 1.82534E-11 | 0.001 | 2.5790E-11 | 9.8814E-11 |
| | | CBDmale | 1.71376E-11 | 0.000 | 2.9553E-11 | 9.8113E-11 |
| | | TpCmale | 1.67295E-11 | 0.009 | 1.1658E-11 | 7.8586E-11 |
| | TpCmale | VehFemale | 1.67295E-11 | 0.830 | -3.7062E-11 | 2.9866E-11 |
| | | CBDfemale | 1.73807E-11 | 0.330 | -5.1835E-11 | 1.7698E-11 |
| | | THCfemale | 1.57727E-11 | 0.003 | -8.0462E-11 | -1.7362E-11 |
| | | TpCfemale | 1.62049E-11 | 0.042 | -6.6100E-11 | -1.2707E-12 |
| | | VehMale | 1.73807E-11 | 0.327 | -1.7587E-11 | 5.1946E-11 |
| | | CBDmale | 1.62049E-11 | 0.253 | -1.3704E-11 | 5.1125E-11 |
| | | THCmale | 1.67295E-11 | 0.009 | -7.8586E-11 | -1.1658E-11 |
| *N*-docosahexaenoyl alanine | VehFemale | CBDfemale | 1.71529E-11 | 0.059 | -6.7322E-11 | 1.2542E-12 |
| | | THCfemale | 1.62727E-11 | 0.008 | -7.7002E-11 | -1.1945E-11 |
| | | TpCfemale | 1.66696E-11 | 0.022 | -7.2557E-11 | -5.9127E-12 |
| | | VehMale | 1.77550E-11 | 0.873 | -3.2646E-11 | 3.8337E-11 |
| | | CBDmale | 1.66696E-11 | 0.407 | -1.9417E-11 | 4.7227E-11 |
| | | THCmale | 1.66696E-11 | 0.009 | -7.7977E-11 | -1.1332E-11 |
| | | TpCmale | 1.62727E-11 | 0.407 | -4.6104E-11 | 1.8953E-11 |
| | CBDfemale | VehFemale | 1.71529E-11 | 0.059 | -1.2542E-12 | 6.7322E-11 |
| | | THCfemale | 1.62727E-11 | 0.485 | -4.3968E-11 | 2.1089E-11 |
| | | TpCfemale | 1.66696E-11 | 0.711 | -3.9523E-11 | 2.7121E-11 |
| | | VehMale | 1.77550E-11 | 0.048 | 3.8792E-13 | 7.1371E-11 |
| | | CBDmale | 1.66696E-11 | 0.007 | 1.3617E-11 | 8.0261E-11 |
| | | THCmale | 1.66696E-11 | 0.488 | -4.4942E-11 | 2.1702E-11 |
| | | TpCmale | 1.62727E-11 | 0.236 | -1.3070E-11 | 5.1987E-11 |
| | THCfemale | VehFemale | 1.62727E-11 | 0.008 | 1.1945E-11 | 7.7002E-11 |
| | | CBDfemale | 1.62727E-11 | 0.485 | -2.1089E-11 | 4.3968E-11 |
| | | TpCfemale | 1.57625E-11 | 0.741 | -2.6270E-11 | 3.6747E-11 |

| | | | | | |
|---|---|---|---|---|---|
| | VehMale | 1.69061E-11 | 0.007 | 1.3524E-11 | 8.1114E-11 |
| | CBDmale | 1.57625E-11 | 0.000 | 2.6870E-11 | 8.9887E-11 |
| | THCmale | 1.57625E-11 | 0.991 | -3.1690E-11 | 3.1328E-11 |
| | TpCmale | 1.53420E-11 | 0.048 | 2.2928E-13 | 6.1566E-11 |
| TpCfemale | VehFemale | 1.66696E-11 | 0.022 | 5.9127E-12 | 7.2557E-11 |
| | CBDfemale | 1.66696E-11 | 0.711 | -2.7121E-11 | 3.9523E-11 |
| | THCfemale | 1.57625E-11 | 0.741 | -3.6747E-11 | 2.6270E-11 |
| | VehMale | 1.72885E-11 | 0.018 | 7.5211E-12 | 7.6640E-11 |
| | CBDmale | 1.61719E-11 | 0.002 | 2.0813E-11 | 8.5467E-11 |
| | THCmale | 1.61719E-11 | 0.739 | -3.7747E-11 | 2.6908E-11 |
| | TpCmale | 1.57625E-11 | 0.109 | -5.8495E-12 | 5.7168E-11 |
| VehMale | VehFemale | 1.77550E-11 | 0.873 | -3.8337E-11 | 3.2646E-11 |
| | CBDfemale | 1.77550E-11 | 0.048 | -7.1371E-11 | -3.8792E-13 |
| | THCfemale | 1.69061E-11 | 0.007 | -8.1114E-11 | -1.3524E-11 |
| | TpCfemale | 1.72885E-11 | 0.018 | -7.6640E-11 | -7.5211E-12 |
| | CBDmale | 1.72885E-11 | 0.525 | -2.3500E-11 | 4.5619E-11 |
| | THCmale | 1.72885E-11 | 0.008 | -8.2059E-11 | -1.2941E-11 |
| | TpCmale | 1.69061E-11 | 0.335 | -5.0216E-11 | 1.7374E-11 |
| CBDmale | VehFemale | 1.66696E-11 | 0.407 | -4.7227E-11 | 1.9417E-11 |
| | CBDfemale | 1.66696E-11 | 0.007 | -8.0261E-11 | -1.3617E-11 |
| | THCfemale | 1.57625E-11 | 0.000 | -8.9887E-11 | -2.6870E-11 |
| | TpCfemale | 1.61719E-11 | 0.002 | -8.5467E-11 | -2.0813E-11 |
| | VehMale | 1.72885E-11 | 0.525 | -4.5619E-11 | 2.3500E-11 |
| | THCmale | 1.61719E-11 | 0.001 | -9.0887E-11 | -2.6232E-11 |
| | TpCmale | 1.57625E-11 | 0.086 | -5.8989E-11 | 4.0280E-12 |
| THCmale | VehFemale | 1.66696E-11 | 0.009 | 1.1332E-11 | 7.7977E-11 |
| | CBDfemale | 1.66696E-11 | 0.488 | -2.1702E-11 | 4.4942E-11 |
| | THCfemale | 1.57625E-11 | 0.991 | -3.1328E-11 | 3.1690E-11 |
| | TpCfemale | 1.61719E-11 | 0.739 | -2.6908E-11 | 3.7747E-11 |
| | VehMale | 1.72885E-11 | 0.008 | 1.2941E-11 | 8.2059E-11 |
| | CBDmale | 1.61719E-11 | 0.001 | 2.6232E-11 | 9.0887E-11 |
| | TpCmale | 1.57625E-11 | 0.053 | -4.2995E-13 | 6.2587E-11 |
| TpCmale | VehFemale | 1.62727E-11 | 0.407 | -1.8953E-11 | 4.6104E-11 |
| | CBDfemale | 1.62727E-11 | 0.236 | -5.1987E-11 | 1.3070E-11 |
| | THCfemale | 1.53420E-11 | 0.048 | -6.1566E-11 | -2.2928E-13 |
| | TpCfemale | 1.57625E-11 | 0.109 | -5.7168E-11 | 5.8495E-12 |
| | VehMale | 1.69061E-11 | 0.335 | -1.7374E-11 | 5.0216E-11 |
| | CBDmale | 1.57625E-11 | 0.086 | -4.0280E-12 | 5.8989E-11 |
| | THCmale | 1.57625E-11 | 0.053 | -6.2587E-11 | 4.2995E-13 |

Supplemental Table 10.1 Descriptive statistics for *N*-acyl glycines (mol/L) in plasma collected from male and female adult mice 2h after receiving an i.p. injection of 10mg/kg CBD, THC, or THC+CBD

| | | | | | | 95% Confidence Interval for Mean | | | |
|---|---|---|---|---|---|---|---|---|---|
| | | N | Mean | Std. Deviation | Std. Error | Lower Bound | Upper Bound | Minimum | Maximum |
| *N*-palmitoyl glycine | Veh F | 7 | 5.7818E-09 | 1.65197E-09 | 6.24387E-10 | 4.2540E-09 | 7.3096E-09 | 2.85E-09 | 7.97E-09 |
| | CBD F | 8 | 4.4939E-09 | 1.92320E-09 | 6.79955E-10 | 2.8861E-09 | 6.1018E-09 | 2.45E-09 | 8.63E-09 |
| | THC F | 10 | 5.5683E-09 | 1.28502E-09 | 4.06359E-10 | 4.6491E-09 | 6.4876E-09 | 3.79E-09 | 7.89E-09 |
| | THC+CBD F | 11 | 6.4670E-09 | 1.90127E-09 | 5.73254E-10 | 5.1897E-09 | 7.7443E-09 | 2.92E-09 | 9.26E-09 |
| | Veh M | 8 | 4.7063E-09 | 8.42167E-10 | 2.97751E-10 | 4.0022E-09 | 5.4103E-09 | 3.33E-09 | 5.57E-09 |
| | CBD M | 11 | 6.2150E-09 | 2.34831E-09 | 7.08041E-10 | 4.6374E-09 | 7.7926E-09 | 2.34E-09 | 9.90E-09 |
| | THC M | 9 | 6.1828E-09 | 3.00835E-09 | 1.00278E-09 | 3.8703E-09 | 8.4952E-09 | 2.59E-09 | 1.14E-08 |
| | THC+CBD M | 9 | 5.0873E-09 | 1.48055E-09 | 4.93515E-10 | 3.9492E-09 | 6.2253E-09 | 2.67E-09 | 6.88E-09 |
| *N*-stearoyl glycine | Veh F | 6 | 3.0990E-09 | 1.48974E-09 | 6.08184E-10 | 1.5356E-09 | 4.6624E-09 | 1.25E-09 | 4.62E-09 |
| | CBD F | 8 | 2.6801E-09 | 1.92451E-09 | 6.80418E-10 | 1.0711E-09 | 4.2890E-09 | 1.27E-09 | 7.09E-09 |
| | THC F | 9 | 4.6190E-09 | 2.83887E-09 | 9.46291E-10 | 2.4368E-09 | 6.8011E-09 | 1.44E-09 | 9.96E-09 |
| | THC+CBD F | 10 | 3.0534E-09 | 1.31704E-09 | 4.16484E-10 | 2.1112E-09 | 3.9955E-09 | 9.94E-10 | 5.36E-09 |
| | Veh M | 6 | 2.6671E-09 | 7.95296E-10 | 3.24678E-10 | 1.8325E-09 | 3.5017E-09 | 1.81E-09 | 3.95E-09 |
| | CBD M | 11 | 3.8802E-09 | 1.00568E-09 | 3.03225E-10 | 3.2046E-09 | 4.5558E-09 | 2.78E-09 | 5.76E-09 |
| | THC M | 9 | 3.6384E-09 | 1.56134E-09 | 5.20447E-10 | 2.4383E-09 | 4.8386E-09 | 1.52E-09 | 6.28E-09 |
| | THC+CBD M | 10 | 3.3521E-09 | 1.33999E-09 | 4.23743E-10 | 2.3935E-09 | 4.3107E-09 | 1.04E-09 | 5.81E-09 |
| *N*-oleoyl glycine | Veh F | 7 | 2.8188E-09 | 7.93053E-10 | 2.99746E-10 | 2.0854E-09 | 3.5523E-09 | 1.79E-09 | 3.69E-09 |
| | CBD F | 7 | 2.5853E-09 | 8.49267E-10 | 3.20993E-10 | 1.7999E-09 | 3.3707E-09 | 1.80E-09 | 4.32E-09 |
| | THC F | 9 | 2.3329E-09 | 8.10631E-10 | 2.70210E-10 | 1.7098E-09 | 2.9560E-09 | 1.47E-09 | 3.68E-09 |
| | THC+CBD F | 11 | 3.0059E-09 | 1.12657E-09 | 3.39673E-10 | 2.2491E-09 | 3.7628E-09 | 1.42E-09 | 5.17E-09 |
| | Veh M | 7 | 2.6346E-09 | 6.64223E-10 | 2.51053E-10 | 2.0203E-09 | 3.2489E-09 | 1.90E-09 | 3.37E-09 |
| | CBD M | 10 | 3.5570E-09 | 2.18926E-09 | 6.92304E-10 | 1.9909E-09 | 5.1231E-09 | 9.03E-10 | 7.85E-09 |
| | THC M | 8 | 2.6565E-09 | 1.52525E-09 | 5.39258E-10 | 1.3813E-09 | 3.9316E-09 | 1.46E-09 | 5.01E-09 |
| | THC+CBD M | 9 | 3.2070E-09 | 1.15101E-09 | 3.83670E-10 | 2.3223E-09 | 4.0918E-09 | 1.58E-09 | 5.17E-09 |
| *N*-linoleoyl glycine | Veh F | 7 | 3.7055E-09 | 1.24400E-09 | 4.70189E-10 | 2.5550E-09 | 4.8560E-09 | 2.46E-09 | 5.39E-09 |
| | CBD F | 7 | 3.5523E-09 | 1.63270E-09 | 6.17103E-10 | 2.0423E-09 | 5.0623E-09 | 1.51E-09 | 5.93E-09 |
| | THC F | 9 | 2.3704E-09 | 7.83580E-10 | 2.61193E-10 | 1.7681E-09 | 2.9727E-09 | 1.10E-09 | 3.64E-09 |
| | THC+CBD F | 10 | 3.8174E-09 | 8.20213E-10 | 2.59374E-10 | 3.2306E-09 | 4.4041E-09 | 3.05E-09 | 5.39E-09 |
| | Veh M | 8 | 2.8640E-09 | 1.32460E-09 | 4.68317E-10 | 1.7566E-09 | 3.9714E-09 | 1.46E-09 | 5.15E-09 |
| | CBD M | 10 | 3.1907E-09 | 9.13762E-10 | 2.88957E-10 | 2.5370E-09 | 3.8443E-09 | 1.80E-09 | 4.51E-09 |
| | THC M | 9 | 4.4514E-09 | 2.89365E-09 | 9.64550E-10 | 2.2271E-09 | 6.6757E-09 | 9.82E-10 | 1.01E-08 |
| | THC+CBD M | 10 | 3.6373E-09 | 1.58390E-09 | 5.00874E-10 | 2.5042E-09 | 4.7703E-09 | 1.84E-09 | 6.13E-09 |
| *N*-arachidonoyl glycine | Veh F | BDL | | | | | | | |
| | CBD F | BDL | | | | | | | |
| | THC F | BDL | | | | | | | |
| | THC+CBD F | BDL | | | | | | | |
| | Veh M | BDL | | | | | | | |
| | CBD M | BDL | | | | | | | |
| | THC M | BAL | | | | | | | |
| | THC+CBD M | BDL | | | | | | | |
| | Veh F | BDL | | | | | | | |

| | | |
|---|---|---|
| *N*-docosahexaenoyl glycine | CBD F | BDL |
| | THC F | BDL |
| | THC+CBD F | BDL |
| | Veh M | BDL |
| | CBD M | BDL |
| | THC M | BDL |
| | THC+CBD M | BDL |

Supplemental Table 10.2 Descriptive statistics for *N*-acyl glycines (mol/g) in liver collected from male and female adult mice 2h after receiving an i.p. injection of 10mg/kg CBD, THC, or THC+CBD

| | | N | Mean | Std. Deviation | Std. Error | 95% Confidence Interval for Mean | | Minimum | Maximum |
|---|---|---|---|---|---|---|---|---|---|
| | | | | | | Lower Bound | Upper Bound | | |
| *N*-palmitoyl glycine | Veh F | 8 | 3.9916E-10 | 1.81586E-10 | 6.42005E-11 | 2.4735E-10 | 5.5097E-10 | 2.34E-10 | 7.39E-10 |
| | CBD F | 8 | 4.9495E-10 | 2.31654E-10 | 8.19019E-11 | 3.0128E-10 | 6.8861E-10 | 2.39E-10 | 8.27E-10 |
| | THC F | 10 | 4.8779E-10 | 1.23586E-10 | 3.90814E-11 | 3.9939E-10 | 5.7620E-10 | 3.32E-10 | 6.97E-10 |
| | THC+CBD F | 10 | 6.1717E-10 | 1.48038E-10 | 4.68137E-11 | 5.1127E-10 | 7.2307E-10 | 3.83E-10 | 9.00E-10 |
| | Veh M | 7 | 2.9587E-10 | 1.27494E-10 | 4.81883E-11 | 1.7796E-10 | 4.1378E-10 | 1.36E-10 | 4.70E-10 |
| | CBD M | 9 | 3.6153E-10 | 9.91350E-11 | 3.30450E-11 | 2.8533E-10 | 4.3773E-10 | 2.53E-10 | 4.90E-10 |
| | THC M | 9 | 5.9032E-10 | 1.54600E-10 | 5.15334E-11 | 4.7148E-10 | 7.0916E-10 | 3.66E-10 | 7.95E-10 |
| | THC+CBD M | 10 | 3.3228E-10 | 9.67588E-11 | 3.05978E-11 | 2.6306E-10 | 4.0150E-10 | 2.29E-10 | 5.31E-10 |
| *N*-stearoyl glycine | Veh F | 8 | 7.6721E-11 | 2.97130E-11 | 1.05051E-11 | 5.1880E-11 | 1.0156E-10 | 3.58E-11 | 1.02E-10 |
| | CBD F | 7 | 1.2156E-10 | 5.87324E-11 | 2.21988E-11 | 6.7239E-11 | 1.7588E-10 | 5.74E-11 | 2.03E-10 |
| | THC F | 9 | 1.3109E-10 | 4.69779E-11 | 1.56593E-11 | 9.4983E-11 | 1.6720E-10 | 8.82E-11 | 2.14E-10 |
| | THC+CBD F | 10 | 1.4133E-10 | 4.86368E-11 | 1.53803E-11 | 1.0654E-10 | 1.7612E-10 | 5.83E-11 | 2.07E-10 |
| | Veh M | 7 | 6.5120E-11 | 1.99122E-11 | 7.52610E-12 | 4.6705E-11 | 8.3536E-11 | 4.52E-11 | 1.03E-10 |
| | CBD M | 9 | 9.4338E-11 | 5.29104E-11 | 1.76368E-11 | 5.3668E-11 | 1.3501E-10 | 4.36E-11 | 2.12E-10 |
| | THC M | 9 | 1.1175E-10 | 3.82454E-11 | 1.27485E-11 | 8.2353E-11 | 1.4115E-10 | 4.28E-11 | 1.74E-10 |
| | THC+CBD M | 10 | 6.6350E-11 | 2.07089E-11 | 6.54873E-12 | 5.1536E-11 | 8.1165E-11 | 3.27E-11 | 9.76E-11 |
| *N*-oleoyl glycine | Veh F | 8 | 2.2407E-10 | 1.22054E-10 | 4.31528E-11 | 1.2203E-10 | 3.2611E-10 | 9.09E-11 | 4.44E-10 |
| | CBD F | 7 | 2.1150E-10 | 8.77884E-11 | 3.31809E-11 | 1.3031E-10 | 2.9269E-10 | 1.43E-10 | 3.60E-10 |
| | THC F | 10 | 4.3720E-10 | 1.47717E-10 | 4.67123E-11 | 3.3153E-10 | 5.4287E-10 | 2.34E-10 | 6.73E-10 |
| | THC+CBD F | 9 | 4.4920E-10 | 9.16581E-11 | 3.05527E-11 | 3.7875E-10 | 5.1966E-10 | 3.45E-10 | 6.62E-10 |
| | Veh M | 7 | 1.6672E-10 | 5.04240E-11 | 1.90585E-11 | 1.2009E-10 | 2.1336E-10 | 1.14E-10 | 2.57E-10 |
| | CBD M | 10 | 2.4564E-10 | 6.98420E-11 | 2.20860E-11 | 1.9568E-10 | 2.9561E-10 | 1.20E-10 | 3.63E-10 |
| | THC M | 8 | 3.3150E-10 | 8.20339E-11 | 2.90034E-11 | 2.6292E-10 | 4.0008E-10 | 1.96E-10 | 4.33E-10 |
| | THC+CBD M | 10 | 4.0360E-10 | 8.35112E-11 | 2.64086E-11 | 3.4386E-10 | 4.6334E-10 | 2.72E-10 | 5.35E-10 |
| *N*-linoleoyl glycine | Veh F | 8 | 3.1516E-10 | 1.47495E-10 | 5.21474E-11 | 1.9185E-10 | 4.3847E-10 | 1.49E-10 | 6.03E-10 |
| | CBD F | 8 | 4.3219E-10 | 2.56964E-10 | 9.08506E-11 | 2.1737E-10 | 6.4702E-10 | 1.54E-10 | 8.80E-10 |
| | THC F | 10 | 4.8339E-10 | 1.18105E-10 | 3.73482E-11 | 3.9890E-10 | 5.6788E-10 | 3.60E-10 | 6.89E-10 |
| | THC+CBD F | 9 | 7.3551E-10 | 1.92113E-10 | 6.40375E-11 | 5.8784E-10 | 8.8318E-10 | 4.17E-10 | 1.00E-09 |
| | Veh M | 7 | 2.2969E-10 | 1.13320E-10 | 4.28308E-11 | 1.2489E-10 | 3.3449E-10 | 1.12E-10 | 4.13E-10 |
| | CBD M | 9 | 3.4129E-10 | 4.86100E-11 | 1.62033E-11 | 3.0392E-10 | 3.7865E-10 | 2.68E-10 | 4.21E-10 |
| | THC M | 9 | 6.3291E-10 | 1.46782E-10 | 4.89272E-11 | 5.2008E-10 | 7.4573E-10 | 5.09E-10 | 8.69E-10 |
| | THC+CBD M | 11 | 7.1951E-10 | 3.52454E-10 | 1.06269E-10 | 4.8273E-10 | 9.5630E-10 | 3.06E-10 | 1.31E-09 |
| *N*-arachidonoyl glycine | Veh F | 8 | 5.0926E-10 | 2.87504E-10 | 1.01648E-10 | 2.6890E-10 | 7.4962E-10 | 2.36E-10 | 1.01E-09 |
| | CBD F | 7 | 6.6532E-10 | 2.52447E-10 | 9.54159E-11 | 4.3184E-10 | 8.9879E-10 | 3.98E-10 | 1.00E-09 |
| | THC F | 10 | 9.1193E-10 | 1.82479E-10 | 5.77050E-11 | 7.8139E-10 | 1.0425E-09 | 6.81E-10 | 1.15E-09 |
| | THC+CBD F | 9 | 1.0437E-09 | 1.95764E-10 | 6.52547E-11 | 8.9318E-10 | 1.1941E-09 | 8.31E-10 | 1.37E-09 |
| | Veh M | 7 | 4.8869E-10 | 2.75462E-10 | 1.04115E-10 | 2.3393E-10 | 7.4345E-10 | 1.49E-10 | 9.06E-10 |
| | CBD M | 10 | 6.6580E-10 | 1.88088E-10 | 5.94787E-11 | 5.3125E-10 | 8.0035E-10 | 4.12E-10 | 9.59E-10 |
| | THC M | 9 | 1.1159E-09 | 2.13962E-10 | 7.13205E-11 | 9.5143E-10 | 1.2804E-09 | 7.80E-10 | 1.37E-09 |
| | THC+CBD M | 11 | 9.6650E-10 | 3.40532E-10 | 1.02674E-10 | 7.3773E-10 | 1.1953E-09 | 5.46E-10 | 1.58E-09 |
| | Veh F | 8 | 7.5668E-11 | 3.53616E-11 | 1.25022E-11 | 4.6105E-11 | 1.0523E-10 | 3.94E-11 | 1.44E-10 |

| | | | | | | | | | |
|---|---|---|---|---|---|---|---|---|---|
| *N*-docosahexaenoyl glycine | CBD F | 8 | 1.2715E-10 | 8.04482E-11 | 2.84427E-11 | 5.9892E-11 | 1.9440E-10 | 4.93E-11 | 2.76E-10 |
| | THC F | 10 | 1.4666E-10 | 4.79560E-11 | 1.51650E-11 | 1.1236E-10 | 1.8097E-10 | 6.93E-11 | 2.25E-10 |
| | THC+CBD F | 9 | 1.7942E-10 | 4.18222E-11 | 1.39407E-11 | 1.4727E-10 | 2.1157E-10 | 1.04E-10 | 2.25E-10 |
| | Veh M | 7 | 6.9034E-11 | 2.97456E-11 | 1.12428E-11 | 4.1524E-11 | 9.6544E-11 | 3.48E-11 | 1.22E-10 |
| | CBD M | 10 | 7.9901E-11 | 2.70016E-11 | 8.53866E-12 | 6.0585E-11 | 9.9217E-11 | 5.00E-11 | 1.31E-10 |
| | THC M | 9 | 1.8204E-10 | 4.69519E-11 | 1.56506E-11 | 1.4595E-10 | 2.1813E-10 | 1.11E-10 | 2.69E-10 |
| | THC+CBD M | 11 | 1.5702E-10 | 8.14854E-11 | 2.45688E-11 | 1.0228E-10 | 2.1176E-10 | 5.72E-11 | 3.09E-10 |

Supplemental Table 10.3 Independent t-test comparing male and female levels of *N*-acyl glycines measured in plasma collected from adult mice 2h after an i.p. injection of 10mg/kg CBD, THC, or THC+CBD

| | | | Levene's Test for Equality of Variances | | t-test for Equality of Means | | | | | | |
|---|---|---|---|---|---|---|---|---|---|---|---|
| | | | | | | | Significance | | | 95% Confidence Interval of the Difference | |
| | | | F | Sig. | t | df | One-Sided p | Two-Sided p | Std. Error Difference | Lower | Upper |
| Veh | *N*-palmitoyl glycine | Equal variances assumed | 2.114 | 0.170 | 1.622 | 13 | 0.064 | 0.129 | 6.63078E-10 | -3.56961E-10 | 2.50803E-09 |
| | | Equal variances not assumed | | | 1.555 | 8.655 | 0.078 | 0.156 | 6.91747E-10 | -4.98855E-10 | 2.64992E-09 |
| | *N*-stearoyl glycine | Equal variances assumed | 3.353 | 0.097 | 0.626 | 10 | 0.273 | 0.545 | 6.89423E-10 | -1.10424E-09 | 1.96801E-09 |
| | | Equal variances not assumed | | | 0.626 | 7.636 | 0.275 | 0.549 | 6.89423E-10 | -1.17122E-09 | 2.03499E-09 |
| | *N*-oleoyl glycine | Equal variances assumed | 0.677 | 0.427 | 0.471 | 12 | 0.323 | 0.646 | 3.90992E-10 | -6.67628E-10 | 1.03617E-09 |
| | | Equal variances not assumed | | | 0.471 | 11.642 | 0.323 | 0.646 | 3.90992E-10 | -6.70545E-10 | 1.03909E-09 |
| | *N*-linoleoyl glycine | Equal variances assumed | 0.003 | 0.955 | 1.262 | 13 | 0.114 | 0.229 | 6.66618E-10 | -5.98597E-10 | 2.28168E-09 |
| | | Equal variances not assumed | | | 1.268 | 12.915 | 0.114 | 0.227 | 6.63625E-10 | -5.93092E-10 | 2.27618E-09 |
| CBD | *N*-palmitoyl glycine | Equal variances assumed | 0.523 | 0.479 | -1.696 | 17 | 0.054 | 0.108 | 1.01450E-09 | -3.86150E-09 | 4.19313E-10 |
| | | Equal variances not assumed | | | -1.753 | 16.681 | 0.049 | 0.098 | 9.81663E-10 | -3.79524E-09 | 3.53053E-10 |
| | *N*-stearoyl glycine | Equal variances assumed | 1.455 | 0.244 | -1.774 | 17 | 0.047 | 0.094 | 6.76557E-10 | -2.62752E-09 | 2.27305E-10 |
| | | Equal variances not assumed | | | -1.611 | 9.786 | 0.069 | 0.139 | 7.44925E-10 | -2.86483E-09 | 4.64617E-10 |
| | *N*-oleoyl glycine | Equal variances assumed | 4.351 | 0.054 | -1.108 | 15 | 0.143 | 0.285 | 8.76613E-10 | -2.84012E-09 | 8.96792E-10 |
| | | Equal variances not assumed | | | -1.273 | 12.424 | 0.113 | 0.226 | 7.63100E-10 | -2.62804E-09 | 6.84711E-10 |
| | *N*-linoleoyl glycine | Equal variances assumed | 3.654 | 0.075 | 0.586 | 15 | 0.283 | 0.567 | 6.16944E-10 | -9.53390E-10 | 1.67658E-09 |

| | | | | | | | | | | | |
|---|---|---|---|---|---|---|---|---|---|---|---|
| | | Equal variances not assumed | | | 0.531 | 8.643 | 0.305 | 0.609 | 6.81404E-10 | -1.18962E-09 | 1.91281E-09 |
| THC | *N*-palmitoyl glycine | Equal variances assumed | 9.736 | 0.006 | -0.590 | 17 | 0.281 | 0.563 | 1.04099E-09 | -2.81075E-09 | 1.58184E-09 |
| | | Equal variances not assumed | | | -0.568 | 10.589 | 0.291 | 0.582 | 1.08199E-09 | -3.00721E-09 | 1.77831E-09 |
| | *N*-stearoyl glycine | Equal variances assumed | 2.521 | 0.132 | 0.908 | 16 | 0.189 | 0.377 | 1.07997E-09 | -1.30887E-09 | 3.26999E-09 |
| | | Equal variances not assumed | | | 0.908 | 12.434 | 0.191 | 0.381 | 1.07997E-09 | -1.36341E-09 | 3.32453E-09 |
| | *N*-oleoyl glycine | Equal variances assumed | 7.729 | 0.014 | -0.556 | 15 | 0.293 | 0.587 | 5.82308E-10 | -1.56470E-09 | 9.17619E-10 |
| | | Equal variances not assumed | | | -0.536 | 10.384 | 0.301 | 0.603 | 6.03169E-10 | -1.66079E-09 | 1.01370E-09 |
| | *N*-linoleoyl glycine | Equal variances assumed | 6.834 | 0.019 | -2.083 | 16 | 0.027 | 0.054 | 9.99289E-10 | -4.19943E-09 | 3.73685E-11 |
| | | Equal variances not assumed | | | -2.083 | 9.167 | 0.033 | 0.066 | 9.99289E-10 | -4.33532E-09 | 1.73258E-10 |
| THC+CBD | *N*-palmitoyl glycine | Equal variances assumed | 0.065 | 0.802 | 1.777 | 18 | 0.046 | 0.092 | 7.76220E-10 | -2.51081E-10 | 3.01048E-09 |
| | | Equal variances not assumed | | | 1.824 | 17.974 | 0.042 | 0.085 | 7.56424E-10 | -2.09654E-10 | 2.96905E-09 |
| | *N*-stearoyl glycine | Equal variances assumed | 0.055 | 0.817 | -0.503 | 18 | 0.311 | 0.621 | 5.94153E-10 | -1.54699E-09 | 9.49545E-10 |
| | | Equal variances not assumed | | | -0.503 | 17.995 | 0.311 | 0.621 | 5.94153E-10 | -1.54702E-09 | 9.49572E-10 |
| | *N*-oleoyl glycine | Equal variances assumed | 0.058 | 0.812 | -0.393 | 18 | 0.349 | 0.699 | 5.11266E-10 | -1.27525E-09 | 8.73013E-10 |
| | | Equal variances not assumed | | | -0.392 | 17.067 | 0.350 | 0.700 | 5.12426E-10 | -1.28192E-09 | 8.79681E-10 |
| | *N*-linoleoyl glycine | Equal variances assumed | 7.442 | 0.014 | 0.319 | 18 | 0.377 | 0.753 | 5.64048E-10 | -1.00495E-09 | 1.36510E-09 |
| | | Equal variances not assumed | | | 0.319 | 13.503 | 0.377 | 0.754 | 5.64048E-10 | -1.03388E-09 | 1.39403E-09 |

Supplemental Table 10.4 Independent t-test comparing male and female levels of *N*-acyl glycines measured in liver collected from adult mice 2h after an i.p. injection of 10mg/kg CBD, THC, or THC+CBD

| | | Levene's Test for Equality of Variances | | t-test for Equality of Means | | | | | | | |
|---|---|---|---|---|---|---|---|---|---|---|---|
| | | | | | | Significance | | | | 95% Confidence Interval of the Difference | |
| | | F | Sig. | t | df | One-Sided p | Two-Sided p | Mean Difference | Std. Error Difference | Lower | Upper |
| *N*-palmitoyl glycine | Equal variances assumed | 1.239 | 0.286 | 1.256 | 13 | 0.116 | 0.231 | 1.03286E-10 | 8.22517E-11 | -7.44077E-11 | 2.80980E-10 |
| | Equal variances not assumed | | | 1.287 | 12.486 | 0.111 | 0.222 | 1.03286E-10 | 8.02734E-11 | -7.08628E-11 | 2.77435E-10 |
| *N*-stearoyl glycine | Equal variances assumed | 4.651 | 0.050 | 0.874 | 13 | 0.199 | 0.398 | 1.16003E-11 | 1.32798E-11 | -1.70890E-11 | 4.02896E-11 |
| | Equal variances not assumed | | | 0.898 | 12.261 | 0.193 | 0.387 | 1.16003E-11 | 1.29229E-11 | -1.64898E-11 | 3.96904E-11 |
| *N*-oleoyl glycine | Equal variances assumed | 4.016 | 0.066 | 1.156 | 13 | 0.134 | 0.269 | 5.73509E-11 | 4.96284E-11 | -4.98647E-11 | 1.64567E-10 |
| | Equal variances not assumed | | | 1.216 | 9.572 | 0.127 | 0.253 | 5.73509E-11 | 4.71740E-11 | -4.83995E-11 | 1.63101E-10 |
| *N*-linoleoyl glycine | Equal variances assumed | 0.240 | 0.632 | 1.243 | 13 | 0.118 | 0.236 | 8.54744E-11 | 6.87404E-11 | -6.30303E-11 | 2.33979E-10 |
| | Equal variances not assumed | | | 1.267 | 12.822 | 0.114 | 0.228 | 8.54744E-11 | 6.74821E-11 | -6.05176E-11 | 2.31466E-10 |
| *N*-arachidonoyl glycine | Equal variances assumed | 0.058 | 0.813 | 0.141 | 13 | 0.445 | 0.890 | 2.05647E-11 | 1.45954E-10 | -2.94750E-10 | 3.35879E-10 |
| | Equal variances not assumed | | | 0.141 | 12.868 | 0.445 | 0.890 | 2.05647E-11 | 1.45507E-10 | -2.94111E-10 | 3.35240E-10 |
| *N*-docosahexaenoyl glycine | Equal variances assumed | 0.384 | 0.546 | 0.390 | 13 | 0.352 | 0.703 | 6.63379E-12 | 1.70217E-11 | -3.01393E-11 | 4.34068E-11 |
| | Equal variances not assumed | | | 0.395 | 12.989 | 0.350 | 0.700 | 6.63379E-12 | 1.68138E-11 | -2.96934E-11 | 4.29610E-11 |
| *N*-palmitoyl glycine | Equal variances assumed | 19.083 | 0.001 | 1.578 | 15 | 0.068 | 0.135 | 1.33416E-10 | 8.45605E-11 | -4.68207E-11 | 3.13652E-10 |
| | Equal variances not assumed | | | 1.511 | 9.250 | 0.082 | 0.164 | 1.33416E-10 | 8.83170E-11 | -6.55510E-11 | 3.32383E-10 |
| *N*-stearoyl glycine | Equal variances assumed | 0.664 | 0.429 | 0.974 | 14 | 0.173 | 0.347 | 2.72189E-11 | 2.79595E-11 | -3.27482E-11 | 8.71861E-11 |

| | | | | | | | | | | | |
|---|---|---|---|---|---|---|---|---|---|---|---|
| | Equal variances not assumed | | | 0.960 | 12.292 | 0.178 | 0.356 | 2.72189E-11 | 2.83521E-11 | -3.43926E-11 | 8.88304E-11 |
| *N*-oleoyl glycine | Equal variances assumed | 1.160 | 0.299 | -0.894 | 15 | 0.193 | 0.386 | -3.41432E-11 | 3.82026E-11 | -1.15570E-10 | 4.72838E-11 |
| | Equal variances not assumed | | | -0.857 | 11.049 | 0.205 | 0.410 | -3.41432E-11 | 3.98593E-11 | -1.21826E-10 | 5.35395E-11 |
| *N*-linoleoyl glycine | Equal variances assumed | 12.475 | 0.003 | 1.045 | 15 | 0.156 | 0.313 | 9.09055E-11 | 8.70239E-11 | -9.45816E-11 | 2.76393E-10 |
| | Equal variances not assumed | | | 0.985 | 7.446 | 0.178 | 0.356 | 9.09055E-11 | 9.22843E-11 | -1.24691E-10 | 3.06502E-10 |
| *N*-arachidonoyl glycine | Equal variances assumed | 1.163 | 0.298 | -0.005 | 15 | 0.498 | 0.996 | -4.85786E-13 | 1.06517E-10 | -2.27521E-10 | 2.26549E-10 |
| | Equal variances not assumed | | | -0.004 | 10.511 | 0.498 | 0.997 | -4.85786E-13 | 1.12436E-10 | -2.49369E-10 | 2.48397E-10 |
| *N*-docosahexaenoyl glycine | Equal variances assumed | 10.124 | 0.006 | 1.749 | 16 | 0.050 | 0.099 | 4.72475E-11 | 2.70065E-11 | -1.00038E-11 | 1.04499E-10 |
| | Equal variances not assumed | | | 1.591 | 8.266 | 0.075 | 0.149 | 4.72475E-11 | 2.96968E-11 | -2.08512E-11 | 1.15346E-10 |
| *N*-palmitoyl glycine | Equal variances assumed | 0.569 | 0.461 | -1.605 | 17 | 0.063 | 0.127 | -1.02526E-10 | 6.38870E-11 | -2.37316E-10 | 3.22638E-11 |
| | Equal variances not assumed | | | -1.585 | 15.338 | 0.067 | 0.133 | -1.02526E-10 | 6.46765E-11 | -2.40116E-10 | 3.50643E-11 |
| *N*-stearoyl glycine | Equal variances assumed | 1.485 | 0.241 | 0.958 | 16 | 0.176 | 0.352 | 1.93421E-11 | 2.01925E-11 | -2.34641E-11 | 6.21483E-11 |
| | Equal variances not assumed | | | 0.958 | 15.368 | 0.176 | 0.353 | 1.93421E-11 | 2.01925E-11 | -2.36076E-11 | 6.22918E-11 |
| *N*-oleoyl glycine | Equal variances assumed | 2.637 | 0.124 | 1.806 | 16 | 0.045 | 0.090 | 1.05705E-10 | 5.85157E-11 | -1.83428E-11 | 2.29753E-10 |
| | Equal variances not assumed | | | 1.922 | 14.505 | 0.037 | 0.074 | 1.05705E-10 | 5.49839E-11 | -1.18398E-11 | 2.23250E-10 |
| *N*-linoleoyl glycine | Equal variances assumed | 0.969 | 0.339 | -2.458 | 17 | 0.012 | 0.025 | -1.49516E-10 | 6.08227E-11 | -2.77841E-10 | -2.11917E-11 |
| | Equal variances not assumed | | | -2.429 | 15.393 | 0.014 | 0.028 | -1.49516E-10 | 6.15529E-11 | -2.80422E-10 | -1.86108E-11 |
| *N*-arachidonoyl glycine | Equal variances assumed | 0.003 | 0.954 | -2.243 | 17 | 0.019 | 0.039 | -2.03972E-10 | 9.09376E-11 | -3.95833E-10 | -1.21104E-11 |
| | Equal variances | | | -2.223 | 15.861 | 0.021 | 0.041 | -2.03972E-10 | 9.17414E-11 | -3.98594E-10 | -9.34997E-12 |

| | | | | | | | | | | | |
|---|---|---|---|---|---|---|---|---|---|---|---|
| | not assumed | | | | | | | | | | |
| *N*-docosahexaenoyl glycine | Equal variances assumed | 0.000 | 0.985 | -1.622 | 17 | 0.062 | 0.123 | -3.53813E-11 | 2.18184E-11 | -8.14140E-11 | 1.06515E-11 |
| | Equal variances not assumed | | | -1.624 | 16.862 | 0.062 | 0.123 | -3.53813E-11 | 2.17927E-11 | -8.13884E-11 | 1.06259E-11 |
| *N*-palmitoyl glycine | Equal variances assumed | 1.410 | 0.251 | 5.094 | 18 | 0.000 | 0.000 | 2.84894E-10 | 5.59263E-11 | 1.67398E-10 | 4.02391E-10 |
| | Equal variances not assumed | | | 5.094 | 15.503 | 0.000 | 0.000 | 2.84894E-10 | 5.59263E-11 | 1.66026E-10 | 4.03762E-10 |
| *N*-stearoyl glycine | Equal variances assumed | 8.387 | 0.010 | 4.485 | 18 | 0.000 | 0.000 | 7.49780E-11 | 1.67165E-11 | 3.98580E-11 | 1.10098E-10 |
| | Equal variances not assumed | | | 4.485 | 12.159 | 0.000 | 0.001 | 7.49780E-11 | 1.67165E-11 | 3.86089E-11 | 1.11347E-10 |
| *N*-oleoyl glycine | Equal variances assumed | 0.136 | 0.717 | 1.135 | 17 | 0.136 | 0.272 | 4.55992E-11 | 4.01757E-11 | -3.91642E-11 | 1.30363E-10 |
| | Equal variances not assumed | | | 1.129 | 16.321 | 0.138 | 0.275 | 4.55992E-11 | 4.03842E-11 | -3.98746E-11 | 1.31073E-10 |
| *N*-linoleoyl glycine | Equal variances assumed | 4.608 | 0.046 | 0.122 | 18 | 0.452 | 0.904 | 1.59972E-11 | 1.31362E-10 | -2.59983E-10 | 2.91978E-10 |
| | Equal variances not assumed | | | 0.129 | 15.952 | 0.450 | 0.899 | 1.59972E-11 | 1.24072E-10 | -2.47089E-10 | 2.79083E-10 |
| *N*-arachidonoyl glycine | Equal variances assumed | 2.161 | 0.159 | 0.601 | 18 | 0.278 | 0.555 | 7.71578E-11 | 1.28280E-10 | -1.92348E-10 | 3.46664E-10 |
| | Equal variances not assumed | | | 0.634 | 16.371 | 0.267 | 0.535 | 7.71578E-11 | 1.21656E-10 | -1.80267E-10 | 3.34582E-10 |
| *N*-docosahexaenoyl glycine | Equal variances assumed | 7.035 | 0.016 | 0.746 | 18 | 0.233 | 0.465 | 2.24009E-11 | 3.00377E-11 | -4.07060E-11 | 8.55077E-11 |
| | Equal variances not assumed | | | 0.793 | 15.471 | 0.220 | 0.440 | 2.24009E-11 | 2.82483E-11 | -3.76498E-11 | 8.24515E-11 |

Supplemental Table 10.5 One-way ANOVA for *N*-acyl glycines measured in plasma collected from adult mice 2h after an i.p. injection of 10mg/kg CBD, THC, or THC+CBD

| | | Sum of Squares | df | Mean Square | F | Sig. |
|---|---|---|---|---|---|---|
| *N*-palmitoyl glycine | Between Groups | 3.42E-17 | 7 | 4.89E-18 | 1.306 | 0.2617487 |
| | Within Groups | 2.43E-16 | 65 | 3.74E-18 | | |
| | Total | 2.78E-16 | 72 | | | |
| *N*-stearoyl glycine | Between Groups | 2.55E-17 | 7 | 3.64E-18 | 1.337 | 0.2488780 |
| | Within Groups | 1.66E-16 | 61 | 2.72E-18 | | |
| | Total | 1.92E-16 | 68 | | | |
| *N*-oleoyl glycine | Between Groups | 9.87E-18 | 7 | 1.41E-18 | 0.857 | 0.5455419 |
| | Within Groups | 9.87E-17 | 60 | 1.65E-18 | | |
| | Total | 1.09E-16 | 67 | | | |
| *N*-linoleoyl glycine | Between Groups | 2.52E-17 | 7 | 3.60E-18 | 1.531 | 0.1736287 |
| | Within Groups | 1.46E-16 | 62 | 2.35E-18 | | |
| | Total | 1.71E-16 | 69 | | | |

Supplemental Table 10.6 One-way ANOVA for *N*-acyl glycines measured in liver collected from adult mice 2h after an i.p. injection of 10mg/kg CBD, THC, or THC+CBD

| | | Sum of Squares | df | Mean Square | F | Sig. |
|---|---|---|---|---|---|---|
| *N*-palmitoyl glycine | Between Groups | 8.82E-19 | 7 | 1.26E-19 | 5.702 | 0.00004 |
| | Within Groups | 1.39E-18 | 63 | 2.21E-20 | | |
| | Total | 2.28E-18 | 70 | | | |
| *N*-stearoyl glycine | Between Groups | 5.45E-20 | 7 | 7.78E-21 | 4.473 | 0.00045 |
| | Within Groups | 1.06E-19 | 61 | 1.74E-21 | | |
| | Total | 1.61E-19 | 68 | | | |
| *N*-oleoyl glycine | Between Groups | 7.34E-19 | 7 | 1.05E-19 | 10.972 | 0.00000 |
| | Within Groups | 5.83E-19 | 61 | 9.56E-21 | | |
| | Total | 1.32E-18 | 68 | | | |
| *N*-linoleoyl glycine | Between Groups | 2.24E-18 | 7 | 3.20E-19 | 7.914 | 0.00000 |
| | Within Groups | 2.55E-18 | 63 | 4.04E-20 | | |
| | Total | 4.78E-18 | 70 | | | |
| *N*-arachidonoyl glycine | Between Groups | 3.50E-18 | 7 | 5.01E-19 | 8.156 | 0.00000 |
| | Within Groups | 3.87E-18 | 63 | 6.14E-20 | | |
| | Total | 7.37E-18 | 70 | | | |
| *N*-docosahexaenoyl glycine | Between Groups | 1.32E-19 | 7 | 1.89E-20 | 6.534 | 0.00001 |
| | Within Groups | 1.85E-19 | 64 | 2.89E-21 | | |
| | Total | 3.17E-19 | 71 | | | |

#### Supplemental Table 10.7 Fisher's LSD post-hoc test for plasma *N*-acyl glycines ANOVA

| Dependent Variable | | | Std. Error | Sig. | 95% Confidence Interval Lower Bound | 95% Confidence Interval Upper Bound |
|---|---|---|---|---|---|---|
| *N*-palmitoyl glycine | VehFemale | CBDfemale | 1.00E-09 | 0.2030 | -7.12E-10 | 3.29E-09 |
| | | THCfemale | 9.53E-10 | 0.8236 | -1.69E-09 | 2.12E-09 |
| | | TpCfemale | 9.35E-10 | 0.4665 | -2.55E-09 | 1.18E-09 |
| | | VehMale | 1.00E-09 | 0.2868 | -9.24E-10 | 3.08E-09 |
| | | CBDmale | 9.35E-10 | 0.6448 | -2.30E-09 | 1.44E-09 |
| | | THCmale | 9.75E-10 | 0.6822 | -2.35E-09 | 1.55E-09 |
| | | TpCmale | 9.75E-10 | 0.4788 | -1.25E-09 | 2.64E-09 |
| | CBDfemale | VehFemale | 1.00E-09 | 0.2030 | -3.29E-09 | 7.12E-10 |
| | | THCfemale | 9.18E-10 | 0.2460 | -2.91E-09 | 7.58E-10 |
| | | TpCfemale | 8.99E-10 | 0.0318 | -3.77E-09 | -1.78E-10 |
| | | VehMale | 9.67E-10 | 0.8270 | -2.14E-09 | 1.72E-09 |
| | | CBDmale | 8.99E-10 | 0.0600 | -3.52E-09 | 7.44E-11 |
| | | THCmale | 9.40E-10 | 0.0771 | -3.57E-09 | 1.89E-10 |
| | | TpCmale | 9.40E-10 | 0.5302 | -2.47E-09 | 1.28E-09 |
| | THCfemale | VehFemale | 9.53E-10 | 0.8236 | -2.12E-09 | 1.69E-09 |
| | | CBDfemale | 9.18E-10 | 0.2460 | -7.58E-10 | 2.91E-09 |
| | | TpCfemale | 8.45E-10 | 0.2917 | -2.59E-09 | 7.90E-10 |
| | | VehMale | 9.18E-10 | 0.3510 | -9.71E-10 | 2.69E-09 |
| | | CBDmale | 8.45E-10 | 0.4470 | -2.34E-09 | 1.04E-09 |
| | | THCmale | 8.89E-10 | 0.4919 | -2.39E-09 | 1.16E-09 |
| | | TpCmale | 8.89E-10 | 0.5903 | -1.29E-09 | 2.26E-09 |
| | TpCfemale | VehFemale | 9.35E-10 | 0.4665 | -1.18E-09 | 2.55E-09 |
| | | CBDfemale | 8.99E-10 | 0.0318 | 1.78E-10 | 3.77E-09 |
| | | THCfemale | 8.45E-10 | 0.2917 | -7.90E-10 | 2.59E-09 |
| | | VehMale | 8.99E-10 | 0.0545 | -3.48E-11 | 3.56E-09 |
| | | CBDmale | 8.25E-10 | 0.7610 | -1.40E-09 | 1.90E-09 |
| | | THCmale | 8.70E-10 | 0.7449 | -1.45E-09 | 2.02E-09 |
| | | TpCmale | 8.70E-10 | 0.1175 | -3.57E-10 | 3.12E-09 |
| | VehMale | VehFemale | 1.00E-09 | 0.2868 | -3.08E-09 | 9.24E-10 |
| | | CBDfemale | 9.67E-10 | 0.8270 | -1.72E-09 | 2.14E-09 |
| | | THCfemale | 9.18E-10 | 0.3510 | -2.69E-09 | 9.71E-10 |
| | | TpCfemale | 8.99E-10 | 0.0545 | -3.56E-09 | 3.48E-11 |
| | | CBDmale | 8.99E-10 | 0.0981 | -3.30E-09 | 2.87E-10 |
| | | THCmale | 9.40E-10 | 0.1211 | -3.35E-09 | 4.01E-10 |
| | | TpCmale | 9.40E-10 | 0.6866 | -2.26E-09 | 1.50E-09 |
| | CBDmale | VehFemale | 9.35E-10 | 0.6448 | -1.44E-09 | 2.30E-09 |
| | | CBDfemale | 8.99E-10 | 0.0600 | -7.44E-11 | 3.52E-09 |
| | | THCfemale | 8.45E-10 | 0.4470 | -1.04E-09 | 2.34E-09 |
| | | TpCfemale | 8.25E-10 | 0.7610 | -1.90E-09 | 1.40E-09 |
| | | VehMale | 8.99E-10 | 0.0981 | -2.87E-10 | 3.30E-09 |
| | | THCmale | 8.70E-10 | 0.9705 | -1.70E-09 | 1.77E-09 |
| | | TpCmale | 8.70E-10 | 0.1993 | -6.09E-10 | 2.86E-09 |
| | THCmale | VehFemale | 9.75E-10 | 0.6822 | -1.55E-09 | 2.35E-09 |
| | | CBDfemale | 9.40E-10 | 0.0771 | -1.89E-10 | 3.57E-09 |
| | | THCfemale | 8.89E-10 | 0.4919 | -1.16E-09 | 2.39E-09 |
| | | TpCfemale | 8.70E-10 | 0.7449 | -2.02E-09 | 1.45E-09 |
| | | VehMale | 9.40E-10 | 0.1211 | -4.01E-10 | 3.35E-09 |
| | | CBDmale | 8.70E-10 | 0.9705 | -1.77E-09 | 1.70E-09 |
| | | TpCmale | 9.12E-10 | 0.2341 | -7.26E-10 | 2.92E-09 |
| | TpCmale | VehFemale | 9.75E-10 | 0.4788 | -2.64E-09 | 1.25E-09 |
| | | CBDfemale | 9.40E-10 | 0.5302 | -1.28E-09 | 2.47E-09 |
| | | THCfemale | 8.89E-10 | 0.5903 | -2.26E-09 | 1.29E-09 |
| | | TpCfemale | 8.70E-10 | 0.1175 | -3.12E-09 | 3.57E-10 |
| | | VehMale | 9.40E-10 | 0.6866 | -1.50E-09 | 2.26E-09 |
| | | CBDmale | 8.70E-10 | 0.1993 | -2.86E-09 | 6.09E-10 |
| | | THCmale | 9.12E-10 | 0.2341 | -2.92E-09 | 7.26E-10 |
| *N*-stearoyl glycine | VehFemale | CBDfemale | 8.91E-10 | 0.6399 | -1.36E-09 | 2.20E-09 |

| | | | | | | |
|---|---|---|---|---|---|---|
| | | THCfemale | 8.70E-10 | 0.0855 | -3.26E-09 | 2.19E-10 |
| | | TpCfemale | 8.52E-10 | 0.9575 | -1.66E-09 | 1.75E-09 |
| | | VehMale | 9.53E-10 | 0.6519 | -1.47E-09 | 2.34E-09 |
| | | CBDmale | 8.37E-10 | 0.3545 | -2.46E-09 | 8.93E-10 |
| | | THCmale | 8.70E-10 | 0.5373 | -2.28E-09 | 1.20E-09 |
| | | TpCmale | 8.52E-10 | 0.7674 | -1.96E-09 | 1.45E-09 |
| | CBDfemale | VehFemale | 8.91E-10 | 0.6399 | -2.20E-09 | 1.36E-09 |
| | | THCfemale | 8.02E-10 | 0.0186 | -3.54E-09 | -3.36E-10 |
| | | TpCfemale | 7.83E-10 | 0.6351 | -1.94E-09 | 1.19E-09 |
| | | VehMale | 8.91E-10 | 0.9884 | -1.77E-09 | 1.79E-09 |
| | | CBDmale | 7.67E-10 | 0.1227 | -2.73E-09 | 3.33E-10 |
| | | THCmale | 8.02E-10 | 0.2366 | -2.56E-09 | 6.45E-10 |
| | | TpCmale | 7.83E-10 | 0.3939 | -2.24E-09 | 8.93E-10 |
| | THCfemale | VehFemale | 8.70E-10 | 0.0855 | -2.19E-10 | 3.26E-09 |
| | | CBDfemale | 8.02E-10 | 0.0186 | 3.36E-10 | 3.54E-09 |
| | | TpCfemale | 7.58E-10 | 0.0432 | 4.98E-11 | 3.08E-09 |
| | | VehMale | 8.70E-10 | 0.0284 | 2.13E-10 | 3.69E-09 |
| | | CBDmale | 7.42E-10 | 0.3230 | -7.44E-10 | 2.22E-09 |
| | | THCmale | 7.78E-10 | 0.2122 | -5.75E-10 | 2.54E-09 |
| | | TpCmale | 7.58E-10 | 0.0998 | -2.49E-10 | 2.78E-09 |
| | TpCfemale | VehFemale | 8.52E-10 | 0.9575 | -1.75E-09 | 1.66E-09 |
| | | CBDfemale | 7.83E-10 | 0.6351 | -1.19E-09 | 1.94E-09 |
| | | THCfemale | 7.58E-10 | 0.0432 | -3.08E-09 | -4.98E-11 |
| | | VehMale | 8.52E-10 | 0.6519 | -1.32E-09 | 2.09E-09 |
| | | CBDmale | 7.21E-10 | 0.2559 | -2.27E-09 | 6.15E-10 |
| | | THCmale | 7.58E-10 | 0.4432 | -2.10E-09 | 9.31E-10 |
| | | TpCmale | 7.38E-10 | 0.6870 | -1.77E-09 | 1.18E-09 |
| | VehMale | VehFemale | 9.53E-10 | 0.6519 | -2.34E-09 | 1.47E-09 |
| | | CBDfemale | 8.91E-10 | 0.9884 | -1.79E-09 | 1.77E-09 |
| | | THCfemale | 8.70E-10 | 0.0284 | -3.69E-09 | -2.13E-10 |
| | | TpCfemale | 8.52E-10 | 0.6519 | -2.09E-09 | 1.32E-09 |
| | | CBDmale | 8.37E-10 | 0.1525 | -2.89E-09 | 4.61E-10 |
| | | THCmale | 8.70E-10 | 0.2684 | -2.71E-09 | 7.67E-10 |
| | | TpCmale | 8.52E-10 | 0.4245 | -2.39E-09 | 1.02E-09 |
| | CBDmale | VehFemale | 8.37E-10 | 0.3545 | -8.93E-10 | 2.46E-09 |
| | | CBDfemale | 7.67E-10 | 0.1227 | -3.33E-10 | 2.73E-09 |
| | | THCfemale | 7.42E-10 | 0.3230 | -2.22E-09 | 7.44E-10 |
| | | TpCfemale | 7.21E-10 | 0.2559 | -6.15E-10 | 2.27E-09 |
| | | VehMale | 8.37E-10 | 0.1525 | -4.61E-10 | 2.89E-09 |
| | | THCmale | 7.42E-10 | 0.7455 | -1.24E-09 | 1.72E-09 |
| | | TpCmale | 7.21E-10 | 0.4666 | -9.13E-10 | 1.97E-09 |
| | THCmale | VehFemale | 8.70E-10 | 0.5373 | -1.20E-09 | 2.28E-09 |
| | | CBDfemale | 8.02E-10 | 0.2366 | -6.45E-10 | 2.56E-09 |
| | | THCfemale | 7.78E-10 | 0.2122 | -2.54E-09 | 5.75E-10 |
| | | TpCfemale | 7.58E-10 | 0.4432 | -9.31E-10 | 2.10E-09 |
| | | VehMale | 8.70E-10 | 0.2684 | -7.67E-10 | 2.71E-09 |
| | | CBDmale | 7.42E-10 | 0.7455 | -1.72E-09 | 1.24E-09 |
| | | TpCmale | 7.58E-10 | 0.7070 | -1.23E-09 | 1.80E-09 |
| | TpCmale | VehFemale | 8.52E-10 | 0.7674 | -1.45E-09 | 1.96E-09 |
| | | CBDfemale | 7.83E-10 | 0.3939 | -8.93E-10 | 2.24E-09 |
| | | THCfemale | 7.58E-10 | 0.0998 | -2.78E-09 | 2.49E-10 |
| | | TpCfemale | 7.38E-10 | 0.6870 | -1.18E-09 | 1.77E-09 |
| | | VehMale | 8.52E-10 | 0.4245 | -1.02E-09 | 2.39E-09 |
| | | CBDmale | 7.21E-10 | 0.4666 | -1.97E-09 | 9.13E-10 |
| | | THCmale | 7.58E-10 | 0.7070 | -1.80E-09 | 1.23E-09 |
| *N*-oleoyl glycine | VehFemale | CBDfemale | 6.86E-10 | 0.7346 | -1.14E-09 | 1.60E-09 |
| | | THCfemale | 6.46E-10 | 0.4552 | -8.07E-10 | 1.78E-09 |
| | | TpCfemale | 6.20E-10 | 0.7639 | -1.43E-09 | 1.05E-09 |
| | | VehMale | 6.86E-10 | 0.7890 | -1.19E-09 | 1.56E-09 |
| | | CBDmale | 6.32E-10 | 0.2475 | -2.00E-09 | 5.26E-10 |

| | | | | | | |
|---|---|---|---|---|---|---|
| | | THCmale | 6.64E-10 | 0.8076 | -1.17E-09 | 1.49E-09 |
| | | TpCmale | 6.46E-10 | 0.5504 | -1.68E-09 | 9.05E-10 |
| | CBDfemale | VehFemale | 6.86E-10 | 0.7346 | -1.60E-09 | 1.14E-09 |
| | | THCfemale | 6.46E-10 | 0.6976 | -1.04E-09 | 1.55E-09 |
| | | TpCfemale | 6.20E-10 | 0.5002 | -1.66E-09 | 8.20E-10 |
| | | VehMale | 6.86E-10 | 0.9430 | -1.42E-09 | 1.32E-09 |
| | | CBDmale | 6.32E-10 | 0.1295 | -2.24E-09 | 2.93E-10 |
| | | THCmale | 6.64E-10 | 0.9150 | -1.40E-09 | 1.26E-09 |
| | | TpCmale | 6.46E-10 | 0.3400 | -1.91E-09 | 6.71E-10 |
| | THCfemale | VehFemale | 6.46E-10 | 0.4552 | -1.78E-09 | 8.07E-10 |
| | | CBDfemale | 6.46E-10 | 0.6976 | -1.55E-09 | 1.04E-09 |
| | | TpCfemale | 5.77E-10 | 0.2477 | -1.83E-09 | 4.80E-10 |
| | | VehMale | 6.46E-10 | 0.6424 | -1.59E-09 | 9.91E-10 |
| | | CBDmale | 5.89E-10 | 0.0421 | -2.40E-09 | -4.52E-11 |
| | | THCmale | 6.23E-10 | 0.6056 | -1.57E-09 | 9.23E-10 |
| | | TpCmale | 6.05E-10 | 0.1535 | -2.08E-09 | 3.35E-10 |
| | TpCfemale | VehFemale | 6.20E-10 | 0.7639 | -1.05E-09 | 1.43E-09 |
| | | CBDfemale | 6.20E-10 | 0.5002 | -8.20E-10 | 1.66E-09 |
| | | THCfemale | 5.77E-10 | 0.2477 | -4.80E-10 | 1.83E-09 |
| | | VehMale | 6.20E-10 | 0.5516 | -8.69E-10 | 1.61E-09 |
| | | CBDmale | 5.60E-10 | 0.3294 | -1.67E-09 | 5.70E-10 |
| | | THCmale | 5.96E-10 | 0.5598 | -8.43E-10 | 1.54E-09 |
| | | TpCmale | 5.77E-10 | 0.7284 | -1.35E-09 | 9.52E-10 |
| | VehMale | VehFemale | 6.86E-10 | 0.7890 | -1.56E-09 | 1.19E-09 |
| | | CBDfemale | 6.86E-10 | 0.9430 | -1.32E-09 | 1.42E-09 |
| | | THCfemale | 6.46E-10 | 0.6424 | -9.91E-10 | 1.59E-09 |
| | | TpCfemale | 6.20E-10 | 0.5516 | -1.61E-09 | 8.69E-10 |
| | | CBDmale | 6.32E-10 | 0.1497 | -2.19E-09 | 3.42E-10 |
| | | THCmale | 6.64E-10 | 0.9738 | -1.35E-09 | 1.31E-09 |
| | | TpCmale | 6.46E-10 | 0.3794 | -1.87E-09 | 7.21E-10 |
| | CBDmale | VehFemale | 6.32E-10 | 0.2475 | -5.26E-10 | 2.00E-09 |
| | | CBDfemale | 6.32E-10 | 0.1295 | -2.93E-10 | 2.24E-09 |
| | | THCfemale | 5.89E-10 | 0.0421 | 4.52E-11 | 2.40E-09 |
| | | TpCfemale | 5.60E-10 | 0.3294 | -5.70E-10 | 1.67E-09 |
| | | VehMale | 6.32E-10 | 0.1497 | -3.42E-10 | 2.19E-09 |
| | | THCmale | 6.08E-10 | 0.1441 | -3.17E-10 | 2.12E-09 |
| | | TpCmale | 5.89E-10 | 0.5549 | -8.29E-10 | 1.53E-09 |
| | THCmale | VehFemale | 6.64E-10 | 0.8076 | -1.49E-09 | 1.17E-09 |
| | | CBDfemale | 6.64E-10 | 0.9150 | -1.26E-09 | 1.40E-09 |
| | | THCfemale | 6.23E-10 | 0.6056 | -9.23E-10 | 1.57E-09 |
| | | TpCfemale | 5.96E-10 | 0.5598 | -1.54E-09 | 8.43E-10 |
| | | VehMale | 6.64E-10 | 0.9738 | -1.31E-09 | 1.35E-09 |
| | | CBDmale | 6.08E-10 | 0.1441 | -2.12E-09 | 3.17E-10 |
| | | TpCmale | 6.23E-10 | 0.3806 | -1.80E-09 | 6.96E-10 |
| | TpCmale | VehFemale | 6.46E-10 | 0.5504 | -9.05E-10 | 1.68E-09 |
| | | CBDfemale | 6.46E-10 | 0.3400 | -6.71E-10 | 1.91E-09 |
| | | THCfemale | 6.05E-10 | 0.1535 | -3.35E-10 | 2.08E-09 |
| | | TpCfemale | 5.77E-10 | 0.7284 | -9.52E-10 | 1.35E-09 |
| | | VehMale | 6.46E-10 | 0.3794 | -7.21E-10 | 1.87E-09 |
| | | CBDmale | 5.89E-10 | 0.5549 | -1.53E-09 | 8.29E-10 |
| | | THCmale | 6.23E-10 | 0.3806 | -6.96E-10 | 1.80E-09 |
| *N*-linoleoyl glycine | VehFemale | CBDfemale | 8.19E-10 | 0.8522 | -1.48E-09 | 1.79E-09 |
| | | THCfemale | 7.72E-10 | 0.0888 | -2.09E-10 | 2.88E-09 |
| | | TpCfemale | 7.55E-10 | 0.8828 | -1.62E-09 | 1.40E-09 |
| | | VehMale | 7.93E-10 | 0.2928 | -7.44E-10 | 2.43E-09 |
| | | CBDmale | 7.55E-10 | 0.4979 | -9.95E-10 | 2.02E-09 |
| | | THCmale | 7.72E-10 | 0.3379 | -2.29E-09 | 7.98E-10 |
| | | TpCmale | 7.55E-10 | 0.9283 | -1.44E-09 | 1.58E-09 |
| | CBDfemale | VehFemale | 8.19E-10 | 0.8522 | -1.79E-09 | 1.48E-09 |
| | | THCfemale | 7.72E-10 | 0.1310 | -3.62E-10 | 2.73E-09 |

| | | | | | |
|---|---|---|---|---|---|
| | TpCfemale | 7.55E-10 | 0.7268 | -1.77E-09 | 1.24E-09 |
| | VehMale | 7.93E-10 | 0.3888 | -8.97E-10 | 2.27E-09 |
| | CBDmale | 7.55E-10 | 0.6338 | -1.15E-09 | 1.87E-09 |
| | THCmale | 7.72E-10 | 0.2488 | -2.44E-09 | 6.45E-10 |
| | TpCmale | 7.55E-10 | 0.9107 | -1.59E-09 | 1.42E-09 |
| THCfemale | VehFemale | 7.72E-10 | 0.0888 | -2.88E-09 | 2.09E-10 |
| | CBDfemale | 7.72E-10 | 0.1310 | -2.73E-09 | 3.62E-10 |
| | TpCfemale | 7.04E-10 | 0.0441 | -2.85E-09 | -3.95E-11 |
| | VehMale | 7.45E-10 | 0.5099 | -1.98E-09 | 9.95E-10 |
| | CBDmale | 7.04E-10 | 0.2485 | -2.23E-09 | 5.87E-10 |
| | THCmale | 7.22E-10 | 0.0054 | -3.53E-09 | -6.37E-10 |
| | TpCmale | 7.04E-10 | 0.0768 | -2.67E-09 | 1.41E-10 |
| TpCfemale | VehFemale | 7.55E-10 | 0.8828 | -1.40E-09 | 1.62E-09 |
| | CBDfemale | 7.55E-10 | 0.7268 | -1.24E-09 | 1.77E-09 |
| | THCfemale | 7.04E-10 | 0.0441 | 3.95E-11 | 2.85E-09 |
| | VehMale | 7.27E-10 | 0.1945 | -5.00E-10 | 2.41E-09 |
| | CBDmale | 6.85E-10 | 0.3640 | -7.43E-10 | 2.00E-09 |
| | THCmale | 7.04E-10 | 0.3714 | -2.04E-09 | 7.73E-10 |
| | TpCmale | 6.85E-10 | 0.7936 | -1.19E-09 | 1.55E-09 |
| VehMale | VehFemale | 7.93E-10 | 0.2928 | -2.43E-09 | 7.44E-10 |
| | CBDfemale | 7.93E-10 | 0.3888 | -2.27E-09 | 8.97E-10 |
| | THCfemale | 7.45E-10 | 0.5099 | -9.95E-10 | 1.98E-09 |
| | TpCfemale | 7.27E-10 | 0.1945 | -2.41E-09 | 5.00E-10 |
| | CBDmale | 7.27E-10 | 0.6547 | -1.78E-09 | 1.13E-09 |
| | THCmale | 7.45E-10 | 0.0370 | -3.08E-09 | -9.89E-11 |
| | TpCmale | 7.27E-10 | 0.2915 | -2.23E-09 | 6.80E-10 |
| CBDmale | VehFemale | 7.55E-10 | 0.4979 | -2.02E-09 | 9.95E-10 |
| | CBDfemale | 7.55E-10 | 0.6338 | -1.87E-09 | 1.15E-09 |
| | THCfemale | 7.04E-10 | 0.2485 | -5.87E-10 | 2.23E-09 |
| | TpCfemale | 6.85E-10 | 0.3640 | -2.00E-09 | 7.43E-10 |
| | VehMale | 7.27E-10 | 0.6547 | -1.13E-09 | 1.78E-09 |
| | THCmale | 7.04E-10 | 0.0783 | -2.67E-09 | 1.47E-10 |
| | TpCmale | 6.85E-10 | 0.5170 | -1.82E-09 | 9.23E-10 |
| THCmale | VehFemale | 7.72E-10 | 0.3379 | -7.98E-10 | 2.29E-09 |
| | CBDfemale | 7.72E-10 | 0.2488 | -6.45E-10 | 2.44E-09 |
| | THCfemale | 7.22E-10 | 0.0054 | 6.37E-10 | 3.53E-09 |
| | TpCfemale | 7.04E-10 | 0.3714 | -7.73E-10 | 2.04E-09 |
| | VehMale | 7.45E-10 | 0.0370 | 9.89E-11 | 3.08E-09 |
| | CBDmale | 7.04E-10 | 0.0783 | -1.47E-10 | 2.67E-09 |
| | TpCmale | 7.04E-10 | 0.2520 | -5.93E-10 | 2.22E-09 |
| TpCmale | VehFemale | 7.55E-10 | 0.9283 | -1.58E-09 | 1.44E-09 |
| | CBDfemale | 7.55E-10 | 0.9107 | -1.42E-09 | 1.59E-09 |
| | THCfemale | 7.04E-10 | 0.0768 | -1.41E-10 | 2.67E-09 |
| | TpCfemale | 6.85E-10 | 0.7936 | -1.55E-09 | 1.19E-09 |
| | VehMale | 7.27E-10 | 0.2915 | -6.80E-10 | 2.23E-09 |
| | CBDmale | 6.85E-10 | 0.5170 | -9.23E-10 | 1.82E-09 |
| | THCmale | 7.04E-10 | 0.2520 | -2.22E-09 | 5.93E-10 |

#### Supplemental Table 10.8 Fisher's LSD post-hoc test for liver *N*-acyl glycines ANOVA

| Dependent Variable | | | Std. Error | Sig. | 95% Confidence Interval | |
|---|---|---|---|---|---|---|
| | | | | | Lower Bound | Upper Bound |
| *N*-palmitoyl glycine | VehFemale | CBDfemale | 7.43431E-11 | 0.202 | -2.4435E-10 | 5.2774E-11 |
| | | THCfemale | 7.05281E-11 | 0.213 | -2.2958E-10 | 5.2303E-11 |
| | | TpCfemale | 7.05281E-11 | 0.003 | -3.5895E-10 | -7.7076E-11 |
| | | VehMale | 7.69525E-11 | 0.184 | -5.0491E-11 | 2.5706E-10 |
| | | CBDmale | 7.22485E-11 | 0.604 | -1.0675E-10 | 1.8200E-10 |
| | | THCmale | 7.22485E-11 | 0.010 | -3.3554E-10 | -4.6785E-11 |
| | | TpCmale | 7.05281E-11 | 0.347 | -7.4060E-11 | 2.0782E-10 |
| | CBDfemale | VehFemale | 7.43431E-11 | 0.202 | -5.2774E-11 | 2.4435E-10 |
| | | THCfemale | 7.05281E-11 | 0.920 | -1.3379E-10 | 1.4809E-10 |
| | | TpCfemale | 7.05281E-11 | 0.088 | -2.6317E-10 | 1.8713E-11 |
| | | VehMale | 7.69525E-11 | 0.012 | 4.5298E-11 | 3.5285E-10 |
| | | CBDmale | 7.22485E-11 | 0.069 | -1.0961E-11 | 2.7779E-10 |
| | | THCmale | 7.22485E-11 | 0.192 | -2.3975E-10 | 4.9004E-11 |
| | | TpCmale | 7.05281E-11 | 0.024 | 2.1729E-11 | 3.0361E-10 |
| | THCfemale | VehFemale | 7.05281E-11 | 0.213 | -5.2303E-11 | 2.2958E-10 |
| | | CBDfemale | 7.05281E-11 | 0.920 | -1.4809E-10 | 1.3379E-10 |
| | | TpCfemale | 6.64945E-11 | 0.056 | -2.6226E-10 | 3.4997E-12 |
| | | VehMale | 7.32734E-11 | 0.011 | 4.5497E-11 | 3.3835E-10 |
| | | CBDmale | 6.83166E-11 | 0.069 | -1.0257E-11 | 2.6278E-10 |
| | | THCmale | 6.83166E-11 | 0.138 | -2.3905E-10 | 3.3994E-11 |
| | | TpCmale | 6.64945E-11 | 0.023 | 2.2637E-11 | 2.8839E-10 |
| | TpCfemale | VehFemale | 7.05281E-11 | 0.003 | 7.7076E-11 | 3.5895E-10 |
| | | CBDfemale | 7.05281E-11 | 0.088 | -1.8713E-11 | 2.6317E-10 |
| | | THCfemale | 6.64945E-11 | 0.056 | -3.4997E-12 | 2.6226E-10 |
| | | VehMale | 7.32734E-11 | 0.000 | 1.7488E-10 | 4.6773E-10 |
| | | CBDmale | 6.83166E-11 | 0.000 | 1.1912E-10 | 3.9216E-10 |
| | | THCmale | 6.83166E-11 | 0.696 | -1.0967E-10 | 1.6337E-10 |
| | | TpCmale | 6.64945E-11 | 0.000 | 1.5202E-10 | 4.1777E-10 |
| | VehMale | VehFemale | 7.69525E-11 | 0.184 | -2.5706E-10 | 5.0491E-11 |
| | | CBDfemale | 7.69525E-11 | 0.012 | -3.5285E-10 | -4.5298E-11 |
| | | THCfemale | 7.32734E-11 | 0.011 | -3.3835E-10 | -4.5497E-11 |
| | | TpCfemale | 7.32734E-11 | 0.000 | -4.6773E-10 | -1.7488E-10 |
| | | CBDmale | 7.49308E-11 | 0.384 | -2.1540E-10 | 8.4078E-11 |
| | | THCmale | 7.49308E-11 | 0.000 | -4.4419E-10 | -1.4471E-10 |
| | | TpCmale | 7.32734E-11 | 0.621 | -1.8283E-10 | 1.1002E-10 |
| | CBDmale | VehFemale | 7.22485E-11 | 0.604 | -1.8200E-10 | 1.0675E-10 |
| | | CBDfemale | 7.22485E-11 | 0.069 | -2.7779E-10 | 1.0961E-11 |
| | | THCfemale | 6.83166E-11 | 0.069 | -2.6278E-10 | 1.0257E-11 |
| | | TpCfemale | 6.83166E-11 | 0.000 | -3.9216E-10 | -1.1912E-10 |
| | | VehMale | 7.49308E-11 | 0.384 | -8.4078E-11 | 2.1540E-10 |
| | | THCmale | 7.00914E-11 | 0.002 | -3.6886E-10 | -8.8722E-11 |
| | | TpCmale | 6.83166E-11 | 0.670 | -1.0727E-10 | 1.6577E-10 |
| | THCmale | VehFemale | 7.22485E-11 | 0.010 | 4.6785E-11 | 3.3554E-10 |
| | | CBDfemale | 7.22485E-11 | 0.192 | -4.9004E-11 | 2.3975E-10 |
| | | THCfemale | 6.83166E-11 | 0.138 | -3.3994E-11 | 2.3905E-10 |
| | | TpCfemale | 6.83166E-11 | 0.696 | -1.6337E-10 | 1.0967E-10 |
| | | VehMale | 7.49308E-11 | 0.000 | 1.4471E-10 | 4.4419E-10 |
| | | CBDmale | 7.00914E-11 | 0.002 | 8.8722E-11 | 3.6886E-10 |
| | | TpCmale | 6.83166E-11 | 0.000 | 1.2152E-10 | 3.9456E-10 |
| | TpCmale | VehFemale | 7.05281E-11 | 0.347 | -2.0782E-10 | 7.4060E-11 |
| | | CBDfemale | 7.05281E-11 | 0.024 | -3.0361E-10 | -2.1729E-11 |
| | | THCfemale | 6.64945E-11 | 0.023 | -2.8839E-10 | -2.2637E-11 |
| | | TpCfemale | 6.64945E-11 | 0.000 | -4.1777E-10 | -1.5202E-10 |
| | | VehMale | 7.32734E-11 | 0.621 | -1.1002E-10 | 1.8283E-10 |
| | | CBDmale | 6.83166E-11 | 0.670 | -1.6577E-10 | 1.0727E-10 |
| | | THCmale | 6.83166E-11 | 0.000 | -3.9456E-10 | -1.2152E-10 |
| *N*-stearoyl glycine | VehFemale | CBDfemale | 2.15906E-11 | 0.042 | -8.8010E-11 | -1.6637E-12 |

| | | | | | | |
|---|---|---|---|---|---|---|
| | | THCfemale | 2.02708E-11 | 0.009 | -9.4907E-11 | -1.3839E-11 |
| | | TpCfemale | 1.97881E-11 | 0.002 | -1.0418E-10 | -2.5039E-11 |
| | | VehMale | 2.15906E-11 | 0.593 | -3.1573E-11 | 5.4773E-11 |
| | | CBDmale | 2.02708E-11 | 0.388 | -5.8152E-11 | 2.2916E-11 |
| | | THCmale | 2.02708E-11 | 0.089 | -7.5565E-11 | 5.5032E-12 |
| | | TpCmale | 1.97881E-11 | 0.602 | -2.9199E-11 | 4.9939E-11 |
| | CBDfemale | VehFemale | 2.15906E-11 | 0.042 | 1.6637E-12 | 8.8010E-11 |
| | | THCfemale | 2.10234E-11 | 0.652 | -5.1575E-11 | 3.2503E-11 |
| | | TpCfemale | 2.05584E-11 | 0.340 | -6.0880E-11 | 2.1338E-11 |
| | | VehMale | 2.22987E-11 | 0.014 | 1.1848E-11 | 1.0103E-10 |
| | | CBDmale | 2.10234E-11 | 0.200 | -1.4820E-11 | 6.9258E-11 |
| | | THCmale | 2.10234E-11 | 0.643 | -3.2233E-11 | 5.1845E-11 |
| | | TpCmale | 2.05584E-11 | 0.009 | 1.4098E-11 | 9.6316E-11 |
| | THCfemale | VehFemale | 2.02708E-11 | 0.009 | 1.3839E-11 | 9.4907E-11 |
| | | CBDfemale | 2.10234E-11 | 0.652 | -3.2503E-11 | 5.1575E-11 |
| | | TpCfemale | 1.91676E-11 | 0.595 | -4.8563E-11 | 2.8093E-11 |
| | | VehMale | 2.10234E-11 | 0.003 | 2.3934E-11 | 1.0801E-10 |
| | | CBDmale | 1.96656E-11 | 0.066 | -2.5687E-12 | 7.6079E-11 |
| | | THCmale | 1.96656E-11 | 0.329 | -1.9982E-11 | 5.8666E-11 |
| | | TpCmale | 1.91676E-11 | 0.001 | 2.6415E-11 | 1.0307E-10 |
| | TpCfemale | VehFemale | 1.97881E-11 | 0.002 | 2.5039E-11 | 1.0418E-10 |
| | | CBDfemale | 2.05584E-11 | 0.340 | -2.1338E-11 | 6.0880E-11 |
| | | THCfemale | 1.91676E-11 | 0.595 | -2.8093E-11 | 4.8563E-11 |
| | | VehMale | 2.05584E-11 | 0.000 | 3.5099E-11 | 1.1732E-10 |
| | | CBDmale | 1.91676E-11 | 0.017 | 8.6621E-12 | 8.5318E-11 |
| | | THCmale | 1.91676E-11 | 0.128 | -8.7509E-12 | 6.7905E-11 |
| | | TpCmale | 1.86564E-11 | 0.000 | 3.7672E-11 | 1.1228E-10 |
| | VehMale | VehFemale | 2.15906E-11 | 0.593 | -5.4773E-11 | 3.1573E-11 |
| | | CBDfemale | 2.22987E-11 | 0.014 | -1.0103E-10 | -1.1848E-11 |
| | | THCfemale | 2.10234E-11 | 0.003 | -1.0801E-10 | -2.3934E-11 |
| | | TpCfemale | 2.05584E-11 | 0.000 | -1.1732E-10 | -3.5099E-11 |
| | | CBDmale | 2.10234E-11 | 0.170 | -7.1257E-11 | 1.2821E-11 |
| | | THCmale | 2.10234E-11 | 0.030 | -8.8670E-11 | -4.5922E-12 |
| | | TpCmale | 2.05584E-11 | 0.952 | -4.2339E-11 | 3.9879E-11 |
| | CBDmale | VehFemale | 2.02708E-11 | 0.388 | -2.2916E-11 | 5.8152E-11 |
| | | CBDfemale | 2.10234E-11 | 0.200 | -6.9258E-11 | 1.4820E-11 |
| | | THCfemale | 1.96656E-11 | 0.066 | -7.6079E-11 | 2.5687E-12 |
| | | TpCfemale | 1.91676E-11 | 0.017 | -8.5318E-11 | -8.6621E-12 |
| | | VehMale | 2.10234E-11 | 0.170 | -1.2821E-11 | 7.1257E-11 |
| | | THCmale | 1.96656E-11 | 0.379 | -5.6737E-11 | 2.1911E-11 |
| | | TpCmale | 1.91676E-11 | 0.149 | -1.0340E-11 | 6.6316E-11 |
| | THCmale | VehFemale | 2.02708E-11 | 0.089 | -5.5032E-12 | 7.5565E-11 |
| | | CBDfemale | 2.10234E-11 | 0.643 | -5.1845E-11 | 3.2233E-11 |
| | | THCfemale | 1.96656E-11 | 0.329 | -5.8666E-11 | 1.9982E-11 |
| | | TpCfemale | 1.91676E-11 | 0.128 | -6.7905E-11 | 8.7509E-12 |
| | | VehMale | 2.10234E-11 | 0.030 | 4.5922E-12 | 8.8670E-11 |
| | | CBDmale | 1.96656E-11 | 0.379 | -2.1911E-11 | 5.6737E-11 |
| | | TpCmale | 1.91676E-11 | 0.021 | 7.0728E-12 | 8.3729E-11 |
| | TpCmale | VehFemale | 1.97881E-11 | 0.602 | -4.9939E-11 | 2.9199E-11 |
| | | CBDfemale | 2.05584E-11 | 0.009 | -9.6316E-11 | -1.4098E-11 |
| | | THCfemale | 1.91676E-11 | 0.001 | -1.0307E-10 | -2.6415E-11 |
| | | TpCfemale | 1.86564E-11 | 0.000 | -1.1228E-10 | -3.7672E-11 |
| | | VehMale | 2.05584E-11 | 0.952 | -3.9879E-11 | 4.2339E-11 |
| | | CBDmale | 1.91676E-11 | 0.149 | -6.6316E-11 | 1.0340E-11 |
| | | THCmale | 1.91676E-11 | 0.021 | -8.3729E-11 | -7.0728E-12 |
| *N*-oleoyl glycine | VehFemale | CBDfemale | 5.06029E-11 | 0.805 | -8.8614E-11 | 1.1376E-10 |
| | | THCfemale | 4.63783E-11 | 0.000 | -3.0587E-10 | -1.2039E-10 |
| | | TpCfemale | 4.75096E-11 | 0.000 | -3.2013E-10 | -1.3013E-10 |
| | | VehMale | 5.06029E-11 | 0.262 | -4.3836E-11 | 1.5854E-10 |
| | | CBDmale | 4.63783E-11 | 0.644 | -1.1431E-10 | 7.1168E-11 |

| | | | | | | |
|---|---|---|---|---|---|---|
| | | THCmale | 4.88870E-11 | 0.032 | -2.0518E-10 | -9.6694E-12 |
| | | TpCmale | 4.63783E-11 | 0.000 | -2.7227E-10 | -8.6790E-11 |
| | CBDfemale | VehFemale | 5.06029E-11 | 0.805 | -1.1376E-10 | 8.8614E-11 |
| | | THCfemale | 4.81836E-11 | 0.000 | -3.2205E-10 | -1.2935E-10 |
| | | TpCfemale | 4.92735E-11 | 0.000 | -3.3623E-10 | -1.3917E-10 |
| | | VehMale | 5.22624E-11 | 0.395 | -5.9727E-11 | 1.4928E-10 |
| | | CBDmale | 4.81836E-11 | 0.481 | -1.3049E-10 | 6.2206E-11 |
| | | THCmale | 5.06029E-11 | 0.021 | -2.2118E-10 | -1.8811E-11 |
| | | TpCmale | 4.81836E-11 | 0.000 | -2.8845E-10 | -9.5752E-11 |
| | THCfemale | VehFemale | 4.63783E-11 | 0.000 | 1.2039E-10 | 3.0587E-10 |
| | | CBDfemale | 4.81836E-11 | 0.000 | 1.2935E-10 | 3.2205E-10 |
| | | TpCfemale | 4.49241E-11 | 0.790 | -1.0183E-10 | 7.7833E-11 |
| | | VehMale | 4.81836E-11 | 0.000 | 1.7413E-10 | 3.6683E-10 |
| | | CBDmale | 4.37259E-11 | 0.000 | 1.0412E-10 | 2.7899E-10 |
| | | THCmale | 4.63783E-11 | 0.026 | 1.2966E-11 | 1.9844E-10 |
| | | TpCmale | 4.37259E-11 | 0.445 | -5.3834E-11 | 1.2104E-10 |
| | TpCfemale | VehFemale | 4.75096E-11 | 0.000 | 1.3013E-10 | 3.2013E-10 |
| | | CBDfemale | 4.92735E-11 | 0.000 | 1.3917E-10 | 3.3623E-10 |
| | | THCfemale | 4.49241E-11 | 0.790 | -7.7833E-11 | 1.0183E-10 |
| | | VehMale | 4.92735E-11 | 0.000 | 1.8395E-10 | 3.8101E-10 |
| | | CBDmale | 4.49241E-11 | 0.000 | 1.1373E-10 | 2.9339E-10 |
| | | THCmale | 4.75096E-11 | 0.016 | 2.2702E-11 | 2.1270E-10 |
| | | TpCmale | 4.49241E-11 | 0.314 | -4.4232E-11 | 1.3543E-10 |
| | VehMale | VehFemale | 5.06029E-11 | 0.262 | -1.5854E-10 | 4.3836E-11 |
| | | CBDfemale | 5.22624E-11 | 0.395 | -1.4928E-10 | 5.9727E-11 |
| | | THCfemale | 4.81836E-11 | 0.000 | -3.6683E-10 | -1.7413E-10 |
| | | TpCfemale | 4.92735E-11 | 0.000 | -3.8101E-10 | -1.8395E-10 |
| | | CBDmale | 4.81836E-11 | 0.107 | -1.7527E-10 | 1.7427E-11 |
| | | THCmale | 5.06029E-11 | 0.002 | -2.6596E-10 | -6.3589E-11 |
| | | TpCmale | 4.81836E-11 | 0.000 | -3.3323E-10 | -1.4053E-10 |
| | CBDmale | VehFemale | 4.63783E-11 | 0.644 | -7.1168E-11 | 1.1431E-10 |
| | | CBDfemale | 4.81836E-11 | 0.481 | -6.2206E-11 | 1.3049E-10 |
| | | THCfemale | 4.37259E-11 | 0.000 | -2.7899E-10 | -1.0412E-10 |
| | | TpCfemale | 4.49241E-11 | 0.000 | -2.9339E-10 | -1.1373E-10 |
| | | VehMale | 4.81836E-11 | 0.107 | -1.7427E-11 | 1.7527E-10 |
| | | THCmale | 4.63783E-11 | 0.069 | -1.7859E-10 | 6.8849E-12 |
| | | TpCmale | 4.37259E-11 | 0.001 | -2.4539E-10 | -7.0523E-11 |
| | THCmale | VehFemale | 4.88870E-11 | 0.032 | 9.6694E-12 | 2.0518E-10 |
| | | CBDfemale | 5.06029E-11 | 0.021 | 1.8811E-11 | 2.2118E-10 |
| | | THCfemale | 4.63783E-11 | 0.026 | -1.9844E-10 | -1.2966E-11 |
| | | TpCfemale | 4.75096E-11 | 0.016 | -2.1270E-10 | -2.2702E-11 |
| | | VehMale | 5.06029E-11 | 0.002 | 6.3589E-11 | 2.6596E-10 |
| | | CBDmale | 4.63783E-11 | 0.069 | -6.8849E-12 | 1.7859E-10 |
| | | TpCmale | 4.63783E-11 | 0.125 | -1.6484E-10 | 2.0635E-11 |
| | TpCmale | VehFemale | 4.63783E-11 | 0.000 | 8.6790E-11 | 2.7227E-10 |
| | | CBDfemale | 4.81836E-11 | 0.000 | 9.5752E-11 | 2.8845E-10 |
| | | THCfemale | 4.37259E-11 | 0.445 | -1.2104E-10 | 5.3834E-11 |
| | | TpCfemale | 4.49241E-11 | 0.314 | -1.3543E-10 | 4.4232E-11 |
| | | VehMale | 4.81836E-11 | 0.000 | 1.4053E-10 | 3.3323E-10 |
| | | CBDmale | 4.37259E-11 | 0.001 | 7.0523E-11 | 2.4539E-10 |
| | | THCmale | 4.63783E-11 | 0.125 | -2.0635E-11 | 1.6484E-10 |
| *N*-linoleoyl glycine | VehFemale | CBDfemale | 1.00512E-10 | 0.249 | -3.1789E-10 | 8.3827E-11 |
| | | THCfemale | 9.53536E-11 | 0.083 | -3.5877E-10 | 2.2324E-11 |
| | | TpCfemale | 9.76797E-11 | 0.000 | -6.1555E-10 | -2.2515E-10 |
| | | VehMale | 1.04039E-10 | 0.414 | -1.2243E-10 | 2.9338E-10 |
| | | CBDmale | 9.76797E-11 | 0.790 | -2.2132E-10 | 1.6907E-10 |
| | | THCmale | 9.76797E-11 | 0.002 | -5.1294E-10 | -1.2254E-10 |
| | | TpCmale | 9.34074E-11 | 0.000 | -5.9101E-10 | -2.1769E-10 |
| | CBDfemale | VehFemale | 1.00512E-10 | 0.249 | -8.3827E-11 | 3.1789E-10 |
| | | THCfemale | 9.53536E-11 | 0.593 | -2.4174E-10 | 1.3935E-10 |

| | | | | | | |
|---|---|---|---|---|---|---|
| | | TpCfemale | 9.76797E-11 | 0.003 | -4.9852E-10 | -1.0812E-10 |
| | | VehMale | 1.04039E-10 | 0.056 | -5.4020E-12 | 4.1041E-10 |
| | | CBDmale | 9.76797E-11 | 0.356 | -1.0429E-10 | 2.8610E-10 |
| | | THCmale | 9.76797E-11 | 0.044 | -3.9591E-10 | -5.5149E-12 |
| | | TpCmale | 9.34074E-11 | 0.003 | -4.7398E-10 | -1.0066E-10 |
| | THCfemale | VehFemale | 9.53536E-11 | 0.083 | -2.2324E-11 | 3.5877E-10 |
| | | CBDfemale | 9.53536E-11 | 0.593 | -1.3935E-10 | 2.4174E-10 |
| | | TpCfemale | 9.23637E-11 | 0.008 | -4.3670E-10 | -6.7548E-11 |
| | | VehMale | 9.90653E-11 | 0.013 | 5.5734E-11 | 4.5167E-10 |
| | | CBDmale | 9.23637E-11 | 0.129 | -4.2473E-11 | 3.2668E-10 |
| | | THCmale | 9.23637E-11 | 0.110 | -3.3409E-10 | 3.5058E-11 |
| | | TpCmale | 8.78333E-11 | 0.009 | -4.1165E-10 | -6.0605E-11 |
| | TpCfemale | VehFemale | 9.76797E-11 | 0.000 | 2.2515E-10 | 6.1555E-10 |
| | | CBDfemale | 9.76797E-11 | 0.003 | 1.0812E-10 | 4.9852E-10 |
| | | THCfemale | 9.23637E-11 | 0.008 | 6.7548E-11 | 4.3670E-10 |
| | | VehMale | 1.01306E-10 | 0.000 | 3.0338E-10 | 7.0827E-10 |
| | | CBDmale | 9.47632E-11 | 0.000 | 2.0485E-10 | 5.8359E-10 |
| | | THCmale | 9.47632E-11 | 0.283 | -8.6763E-11 | 2.9198E-10 |
| | | TpCmale | 9.03532E-11 | 0.860 | -1.6456E-10 | 1.9655E-10 |
| | VehMale | VehFemale | 1.04039E-10 | 0.414 | -2.9338E-10 | 1.2243E-10 |
| | | CBDfemale | 1.04039E-10 | 0.056 | -4.1041E-10 | 5.4020E-12 |
| | | THCfemale | 9.90653E-11 | 0.013 | -4.5167E-10 | -5.5734E-11 |
| | | TpCfemale | 1.01306E-10 | 0.000 | -7.0827E-10 | -3.0338E-10 |
| | | CBDmale | 1.01306E-10 | 0.275 | -3.1404E-10 | 9.0846E-11 |
| | | THCmale | 1.01306E-10 | 0.000 | -6.0566E-10 | -2.0077E-10 |
| | | TpCmale | 9.71934E-11 | 0.000 | -6.8405E-10 | -2.9560E-10 |
| | CBDmale | VehFemale | 9.76797E-11 | 0.790 | -1.6907E-10 | 2.2132E-10 |
| | | CBDfemale | 9.76797E-11 | 0.356 | -2.8610E-10 | 1.0429E-10 |
| | | THCfemale | 9.23637E-11 | 0.129 | -3.2668E-10 | 4.2473E-11 |
| | | TpCfemale | 9.47632E-11 | 0.000 | -5.8359E-10 | -2.0485E-10 |
| | | VehMale | 1.01306E-10 | 0.275 | -9.0846E-11 | 3.1404E-10 |
| | | THCmale | 9.47632E-11 | 0.003 | -4.8099E-10 | -1.0225E-10 |
| | | TpCmale | 9.03532E-11 | 0.000 | -5.5878E-10 | -1.9767E-10 |
| | THCmale | VehFemale | 9.76797E-11 | 0.002 | 1.2254E-10 | 5.1294E-10 |
| | | CBDfemale | 9.76797E-11 | 0.044 | 5.5149E-12 | 3.9591E-10 |
| | | THCfemale | 9.23637E-11 | 0.110 | -3.5058E-11 | 3.3409E-10 |
| | | TpCfemale | 9.47632E-11 | 0.283 | -2.9198E-10 | 8.6763E-11 |
| | | VehMale | 1.01306E-10 | 0.000 | 2.0077E-10 | 6.0566E-10 |
| | | CBDmale | 9.47632E-11 | 0.003 | 1.0225E-10 | 4.8099E-10 |
| | | TpCmale | 9.03532E-11 | 0.341 | -2.6717E-10 | 9.3947E-11 |
| | TpCmale | VehFemale | 9.34074E-11 | 0.000 | 2.1769E-10 | 5.9101E-10 |
| | | CBDfemale | 9.34074E-11 | 0.003 | 1.0066E-10 | 4.7398E-10 |
| | | THCfemale | 8.78333E-11 | 0.009 | 6.0605E-11 | 4.1165E-10 |
| | | TpCfemale | 9.03532E-11 | 0.860 | -1.9655E-10 | 1.6456E-10 |
| | | VehMale | 9.71934E-11 | 0.000 | 2.9560E-10 | 6.8405E-10 |
| | | CBDmale | 9.03532E-11 | 0.000 | 1.9767E-10 | 5.5878E-10 |
| | | THCmale | 9.03532E-11 | 0.341 | -9.3947E-11 | 2.6717E-10 |
| *N*-arachidonoyl glycine | VehFemale | CBDfemale | 1.28220E-10 | 0.228 | -4.1229E-10 | 1.0017E-10 |
| | | THCfemale | 1.17516E-10 | 0.001 | -6.3751E-10 | -1.6783E-10 |
| | | TpCfemale | 1.20382E-10 | 0.000 | -7.7497E-10 | -2.9384E-10 |
| | | VehMale | 1.28220E-10 | 0.873 | -2.3566E-10 | 2.7679E-10 |
| | | CBDmale | 1.17516E-10 | 0.188 | -3.9138E-10 | 7.8289E-11 |
| | | THCmale | 1.20382E-10 | 0.000 | -8.4721E-10 | -3.6608E-10 |
| | | TpCmale | 1.15117E-10 | 0.000 | -6.8729E-10 | -2.2720E-10 |
| | CBDfemale | VehFemale | 1.28220E-10 | 0.228 | -1.0017E-10 | 4.1229E-10 |
| | | THCfemale | 1.22090E-10 | 0.048 | -4.9059E-10 | -2.6309E-12 |
| | | TpCfemale | 1.24852E-10 | 0.004 | -6.2784E-10 | -1.2884E-10 |
| | | VehMale | 1.32425E-10 | 0.187 | -8.8005E-11 | 4.4126E-10 |
| | | CBDmale | 1.22090E-10 | 0.997 | -2.4446E-10 | 2.4349E-10 |
| | | THCmale | 1.24852E-10 | 0.001 | -7.0008E-10 | -2.0108E-10 |

| | | | | | | |
|---|---|---|---|---|---|---|
| | | TpCmale | 1.19783E-10 | 0.014 | -5.4055E-10 | -6.1815E-11 |
| | THCfemale | VehFemale | 1.17516E-10 | 0.001 | 1.6783E-10 | 6.3751E-10 |
| | | CBDfemale | 1.22090E-10 | 0.048 | 2.6309E-12 | 4.9059E-10 |
| | | TpCfemale | 1.13831E-10 | 0.252 | -3.5920E-10 | 9.5741E-11 |
| | | VehMale | 1.22090E-10 | 0.001 | 1.7926E-10 | 6.6721E-10 |
| | | CBDmale | 1.10795E-10 | 0.030 | 2.4717E-11 | 4.6753E-10 |
| | | THCmale | 1.13831E-10 | 0.078 | -4.3144E-10 | 2.3501E-11 |
| | | TpCmale | 1.08248E-10 | 0.616 | -2.7089E-10 | 1.6174E-10 |
| | TpCfemale | VehFemale | 1.20382E-10 | 0.000 | 2.9384E-10 | 7.7497E-10 |
| | | CBDfemale | 1.24852E-10 | 0.004 | 1.2884E-10 | 6.2784E-10 |
| | | THCfemale | 1.13831E-10 | 0.252 | -9.5741E-11 | 3.5920E-10 |
| | | VehMale | 1.24852E-10 | 0.000 | 3.0547E-10 | 8.0446E-10 |
| | | CBDmale | 1.13831E-10 | 0.002 | 1.5038E-10 | 6.0533E-10 |
| | | THCmale | 1.16788E-10 | 0.538 | -3.0562E-10 | 1.6114E-10 |
| | | TpCmale | 1.11353E-10 | 0.491 | -1.4536E-10 | 2.9968E-10 |
| | VehMale | VehFemale | 1.28220E-10 | 0.873 | -2.7679E-10 | 2.3566E-10 |
| | | CBDfemale | 1.32425E-10 | 0.187 | -4.4126E-10 | 8.8005E-11 |
| | | THCfemale | 1.22090E-10 | 0.001 | -6.6721E-10 | -1.7926E-10 |
| | | TpCfemale | 1.24852E-10 | 0.000 | -8.0446E-10 | -3.0547E-10 |
| | | CBDmale | 1.22090E-10 | 0.152 | -4.2109E-10 | 6.6866E-11 |
| | | THCmale | 1.24852E-10 | 0.000 | -8.7670E-10 | -3.7771E-10 |
| | | TpCmale | 1.19783E-10 | 0.000 | -7.1718E-10 | -2.3844E-10 |
| | CBDmale | VehFemale | 1.17516E-10 | 0.188 | -7.8289E-11 | 3.9138E-10 |
| | | CBDfemale | 1.22090E-10 | 0.997 | -2.4349E-10 | 2.4446E-10 |
| | | THCfemale | 1.10795E-10 | 0.030 | -4.6753E-10 | -2.4717E-11 |
| | | TpCfemale | 1.13831E-10 | 0.002 | -6.0533E-10 | -1.5038E-10 |
| | | VehMale | 1.22090E-10 | 0.152 | -6.6866E-11 | 4.2109E-10 |
| | | THCmale | 1.13831E-10 | 0.000 | -6.7757E-10 | -2.2262E-10 |
| | | TpCmale | 1.08248E-10 | 0.007 | -5.1701E-10 | -8.4381E-11 |
| | THCmale | VehFemale | 1.20382E-10 | 0.000 | 3.6608E-10 | 8.4721E-10 |
| | | CBDfemale | 1.24852E-10 | 0.001 | 2.0108E-10 | 7.0008E-10 |
| | | THCfemale | 1.13831E-10 | 0.078 | -2.3501E-11 | 4.3144E-10 |
| | | TpCfemale | 1.16788E-10 | 0.538 | -1.6114E-10 | 3.0562E-10 |
| | | VehMale | 1.24852E-10 | 0.000 | 3.7771E-10 | 8.7670E-10 |
| | | CBDmale | 1.13831E-10 | 0.000 | 2.2262E-10 | 6.7757E-10 |
| | | TpCmale | 1.11353E-10 | 0.185 | -7.3123E-11 | 3.7192E-10 |
| | TpCmale | VehFemale | 1.15117E-10 | 0.000 | 2.2720E-10 | 6.8729E-10 |
| | | CBDfemale | 1.19783E-10 | 0.014 | 6.1815E-11 | 5.4055E-10 |
| | | THCfemale | 1.08248E-10 | 0.616 | -1.6174E-10 | 2.7089E-10 |
| | | TpCfemale | 1.11353E-10 | 0.491 | -2.9968E-10 | 1.4536E-10 |
| | | VehMale | 1.19783E-10 | 0.000 | 2.3844E-10 | 7.1718E-10 |
| | | CBDmale | 1.08248E-10 | 0.007 | 8.4381E-11 | 5.1701E-10 |
| | | THCmale | 1.11353E-10 | 0.185 | -3.7192E-10 | 7.3123E-11 |
| *N*-docosahexaenoyl glycine | VehFemale | CBDfemale | 2.68570E-11 | 0.060 | -1.0513E-10 | 2.1724E-12 |
| | | THCfemale | 2.54788E-11 | 0.007 | -1.2189E-10 | -2.0094E-11 |
| | | TpCfemale | 2.61003E-11 | 0.000 | -1.5589E-10 | -5.1610E-11 |
| | | VehMale | 2.77996E-11 | 0.812 | -4.8902E-11 | 6.2170E-11 |
| | | CBDmale | 2.54788E-11 | 0.869 | -5.5133E-11 | 4.6667E-11 |
| | | THCmale | 2.61003E-11 | 0.000 | -1.5852E-10 | -5.4234E-11 |
| | | TpCmale | 2.49588E-11 | 0.002 | -1.3121E-10 | -3.1489E-11 |
| | CBDfemale | VehFemale | 2.68570E-11 | 0.060 | -2.1724E-12 | 1.0513E-10 |
| | | THCfemale | 2.54788E-11 | 0.447 | -7.0413E-11 | 3.1386E-11 |
| | | TpCfemale | 2.61003E-11 | 0.049 | -1.0441E-10 | -1.2901E-13 |
| | | VehMale | 2.77996E-11 | 0.041 | 2.5783E-12 | 1.1365E-10 |
| | | CBDmale | 2.54788E-11 | 0.068 | -3.6522E-12 | 9.8147E-11 |
| | | THCmale | 2.61003E-11 | 0.039 | -1.0704E-10 | -2.7535E-12 |
| | | TpCmale | 2.49588E-11 | 0.236 | -7.9730E-11 | 1.9991E-11 |
| | THCfemale | VehFemale | 2.54788E-11 | 0.007 | 2.0094E-11 | 1.2189E-10 |
| | | CBDfemale | 2.54788E-11 | 0.447 | -3.1386E-11 | 7.0413E-11 |
| | | TpCfemale | 2.46799E-11 | 0.189 | -8.2061E-11 | 1.6547E-11 |

| | | | | | | |
|---|---|---|---|---|---|---|
| | | VehMale | 2.64705E-11 | 0.005 | 2.4747E-11 | 1.3051E-10 |
| | | CBDmale | 2.40216E-11 | 0.007 | 1.8772E-11 | 1.1475E-10 |
| | | THCmale | 2.46799E-11 | 0.157 | -8.4685E-11 | 1.3922E-11 |
| | | TpCmale | 2.34693E-11 | 0.661 | -5.7241E-11 | 3.6529E-11 |
| | TpCfemale | VehFemale | 2.61003E-11 | 0.000 | 5.1610E-11 | 1.5589E-10 |
| | | CBDfemale | 2.61003E-11 | 0.049 | 1.2901E-13 | 1.0441E-10 |
| | | THCfemale | 2.46799E-11 | 0.189 | -1.6547E-11 | 8.2061E-11 |
| | | VehMale | 2.70693E-11 | 0.000 | 5.6308E-11 | 1.6446E-10 |
| | | CBDmale | 2.46799E-11 | 0.000 | 5.0214E-11 | 1.4882E-10 |
| | | THCmale | 2.53210E-11 | 0.918 | -5.3209E-11 | 4.7960E-11 |
| | | TpCmale | 2.41426E-11 | 0.357 | -2.5830E-11 | 7.0631E-11 |
| | VehMale | VehFemale | 2.77996E-11 | 0.812 | -6.2170E-11 | 4.8902E-11 |
| | | CBDfemale | 2.77996E-11 | 0.041 | -1.1365E-10 | -2.5783E-12 |
| | | THCfemale | 2.64705E-11 | 0.005 | -1.3051E-10 | -2.4747E-11 |
| | | TpCfemale | 2.70693E-11 | 0.000 | -1.6446E-10 | -5.6308E-11 |
| | | CBDmale | 2.64705E-11 | 0.683 | -6.3748E-11 | 4.2014E-11 |
| | | THCmale | 2.70693E-11 | 0.000 | -1.6709E-10 | -5.8932E-11 |
| | | TpCmale | 2.59704E-11 | 0.001 | -1.3987E-10 | -3.6102E-11 |
| | CBDmale | VehFemale | 2.54788E-11 | 0.869 | -4.6667E-11 | 5.5133E-11 |
| | | CBDfemale | 2.54788E-11 | 0.068 | -9.8147E-11 | 3.6522E-12 |
| | | THCfemale | 2.40216E-11 | 0.007 | -1.1475E-10 | -1.8772E-11 |
| | | TpCfemale | 2.46799E-11 | 0.000 | -1.4882E-10 | -5.0214E-11 |
| | | VehMale | 2.64705E-11 | 0.683 | -4.2014E-11 | 6.3748E-11 |
| | | THCmale | 2.46799E-11 | 0.000 | -1.5145E-10 | -5.2839E-11 |
| | | TpCmale | 2.34693E-11 | 0.002 | -1.2400E-10 | -3.0232E-11 |
| | THCmale | VehFemale | 2.61003E-11 | 0.000 | 5.4234E-11 | 1.5852E-10 |
| | | CBDfemale | 2.61003E-11 | 0.039 | 2.7535E-12 | 1.0704E-10 |
| | | THCfemale | 2.46799E-11 | 0.157 | -1.3922E-11 | 8.4685E-11 |
| | | TpCfemale | 2.53210E-11 | 0.918 | -4.7960E-11 | 5.3209E-11 |
| | | VehMale | 2.70693E-11 | 0.000 | 5.8932E-11 | 1.6709E-10 |
| | | CBDmale | 2.46799E-11 | 0.000 | 5.2839E-11 | 1.5145E-10 |
| | | TpCmale | 2.41426E-11 | 0.304 | -2.3205E-11 | 7.3256E-11 |
| | TpCmale | VehFemale | 2.49588E-11 | 0.002 | 3.1489E-11 | 1.3121E-10 |
| | | CBDfemale | 2.49588E-11 | 0.236 | -1.9991E-11 | 7.9730E-11 |
| | | THCfemale | 2.34693E-11 | 0.661 | -3.6529E-11 | 5.7241E-11 |
| | | TpCfemale | 2.41426E-11 | 0.357 | -7.0631E-11 | 2.5830E-11 |
| | | VehMale | 2.59704E-11 | 0.001 | 3.6102E-11 | 1.3987E-10 |
| | | CBDmale | 2.34693E-11 | 0.002 | 3.0232E-11 | 1.2400E-10 |
| | | THCmale | 2.41426E-11 | 0.304 | -7.3256E-11 | 2.3205E-11 |

Supplemental Table 11.1 Descriptive statistics for *N*-acyl leucines (mol/L) in plasma collected from male and female adult mice 2h after receiving an i.p. injection of 10mg/kg CBD, THC, or THC+CBD

| | | N | Mean | Std. Deviation | Std. Error | 95% Confidence Interval for Mean | | Minimum | Maximum |
|---|---|---|---|---|---|---|---|---|---|
| | | | | | | Lower Bound | Upper Bound | | |
| *N*-palmitoyl leucine | Veh F | 7 | 4.3113E-09 | 2.26131E-09 | 8.54697E-10 | 2.2199E-09 | 6.4026E-09 | 1.62E-09 | 8.90E-09 |
| | CBD F | 8 | 4.1172E-09 | 2.76021E-09 | 9.75881E-10 | 1.8096E-09 | 6.4248E-09 | 1.72E-09 | 8.86E-09 |
| | THC F | 9 | 4.2310E-09 | 1.90182E-09 | 6.33942E-10 | 2.7691E-09 | 5.6929E-09 | 1.93E-09 | 6.86E-09 |
| | THC+CBD F | 10 | 5.4789E-09 | 2.10326E-09 | 6.65108E-10 | 3.9743E-09 | 6.9835E-09 | 3.19E-09 | 9.32E-09 |
| | Veh M | 8 | 4.8820E-09 | 3.59471E-09 | 1.27092E-09 | 1.8768E-09 | 7.8873E-09 | 1.20E-09 | 9.71E-09 |
| | CBD M | 11 | 4.7829E-09 | 3.16689E-09 | 9.54854E-10 | 2.6553E-09 | 6.9104E-09 | 1.18E-09 | 8.94E-09 |
| | THC M | 9 | 7.1151E-09 | 4.63953E-09 | 1.54651E-09 | 3.5488E-09 | 1.0681E-08 | 2.21E-09 | 1.40E-08 |
| | THC+CBD M | 10 | 8.0013E-09 | 5.49677E-09 | 1.73823E-09 | 4.0691E-09 | 1.1933E-08 | 1.48E-09 | 1.85E-08 |
| *N*-stearoyl leucine | Veh F | 7 | 1.1560E-09 | 8.55795E-10 | 3.23460E-10 | 3.6452E-10 | 1.9475E-09 | 4.70E-10 | 2.89E-09 |
| | CBD F | 9 | 1.5601E-09 | 1.35868E-09 | 4.52894E-10 | 5.1572E-10 | 2.6045E-09 | 4.06E-10 | 4.24E-09 |
| | THC F | 9 | 1.3122E-09 | 7.61964E-10 | 2.53988E-10 | 7.2649E-10 | 1.8979E-09 | 4.83E-10 | 2.55E-09 |
| | THC+CBD F | 10 | 1.7225E-09 | 8.99396E-10 | 2.84414E-10 | 1.0791E-09 | 2.3659E-09 | 6.00E-10 | 3.01E-09 |
| | Veh M | 8 | 1.6042E-09 | 1.40228E-09 | 4.95781E-10 | 4.3191E-10 | 2.7766E-09 | 3.09E-10 | 3.55E-09 |
| | CBD M | 11 | 1.4145E-09 | 1.00786E-09 | 3.03882E-10 | 7.3740E-10 | 2.0916E-09 | 3.69E-10 | 2.71E-09 |
| | THC M | 9 | 2.2327E-09 | 1.62411E-09 | 5.41369E-10 | 9.8432E-10 | 3.4811E-09 | 5.87E-10 | 4.51E-09 |
| | THC+CBD M | 10 | 2.4571E-09 | 1.83609E-09 | 5.80623E-10 | 1.1436E-09 | 3.7705E-09 | 3.28E-10 | 5.07E-09 |
| *N*-oleoyl leucine | Veh F | 7 | 2.6641E-09 | 1.70464E-09 | 6.44292E-10 | 1.0876E-09 | 4.2406E-09 | 9.60E-10 | 5.95E-09 |
| | CBD F | 8 | 2.8210E-09 | 2.20169E-09 | 7.78415E-10 | 9.8038E-10 | 4.6617E-09 | 1.07E-09 | 6.37E-09 |
| | THC F | 9 | 2.8189E-09 | 1.27883E-09 | 4.26275E-10 | 1.8359E-09 | 3.8019E-09 | 1.55E-09 | 5.02E-09 |
| | THC+CBD F | 10 | 3.2874E-09 | 1.48812E-09 | 4.70586E-10 | 2.2228E-09 | 4.3519E-09 | 1.61E-09 | 5.75E-09 |
| | Veh M | 8 | 3.8017E-09 | 3.49819E-09 | 1.23680E-09 | 8.7716E-10 | 6.7263E-09 | 3.50E-10 | 9.18E-09 |
| | CBD M | 11 | 3.1584E-09 | 2.36505E-09 | 7.13088E-10 | 1.5695E-09 | 4.7472E-09 | 6.50E-10 | 6.58E-09 |
| | THC M | 9 | 4.1113E-09 | 2.85960E-09 | 9.53201E-10 | 1.9132E-09 | 6.3093E-09 | 9.60E-10 | 8.53E-09 |
| | THC+CBD M | 10 | 5.9750E-09 | 4.66882E-09 | 1.47641E-09 | 2.6351E-09 | 9.3148E-09 | 8.35E-10 | 1.32E-08 |
| *N*-linoleoyl leucine | Veh F | 7 | 3.1564E-09 | 1.66809E-09 | 6.30480E-10 | 1.6136E-09 | 4.6991E-09 | 1.35E-09 | 6.09E-09 |
| | CBD F | 8 | 2.7444E-09 | 1.92982E-09 | 6.82295E-10 | 1.1310E-09 | 4.3578E-09 | 1.11E-09 | 6.73E-09 |
| | THC F | 10 | 2.7093E-09 | 1.61476E-09 | 5.10633E-10 | 1.5542E-09 | 3.8645E-09 | 1.10E-09 | 5.58E-09 |
| | THC+CBD F | 10 | 3.9374E-09 | 2.04714E-09 | 6.47364E-10 | 2.4730E-09 | 5.4019E-09 | 1.44E-09 | 7.47E-09 |
| | Veh M | 8 | 3.1426E-09 | 2.47845E-09 | 8.76266E-10 | 1.0705E-09 | 5.2146E-09 | 9.97E-10 | 6.54E-09 |
| | CBD M | 11 | 3.2785E-09 | 2.33696E-09 | 7.04621E-10 | 1.7085E-09 | 4.8485E-09 | 8.42E-10 | 7.70E-09 |
| | THC M | 9 | 4.4198E-09 | 2.96668E-09 | 9.88893E-10 | 2.1394E-09 | 6.7001E-09 | 1.16E-09 | 8.86E-09 |
| | THC+CBD M | 10 | 4.5747E-09 | 3.26582E-09 | 1.03274E-09 | 2.2385E-09 | 6.9109E-09 | 8.94E-10 | 1.06E-08 |
| *N*-docosahexaenoyl leucine | Veh F | 7 | 8.7192E-10 | 4.50022E-10 | 1.70092E-10 | 4.5572E-10 | 1.2881E-09 | 4.94E-10 | 1.78E-09 |
| | CBD F | 7 | 8.2886E-10 | 4.36374E-10 | 1.64934E-10 | 4.2529E-10 | 1.2324E-09 | 3.14E-10 | 1.43E-09 |
| | THC F | 9 | 1.4308E-09 | 8.37962E-10 | 2.79321E-10 | 7.8671E-10 | 2.0749E-09 | 3.75E-10 | 3.08E-09 |
| | THC+CBD F | 10 | 1.8799E-09 | 1.23852E-09 | 3.91655E-10 | 9.9396E-10 | 2.7659E-09 | 4.06E-10 | 3.36E-09 |
| | Veh M | 7 | 1.3094E-09 | 1.04722E-09 | 3.95812E-10 | 3.4091E-10 | 2.2779E-09 | 3.46E-10 | 2.96E-09 |
| | CBD M | 8 | 1.4072E-09 | 9.56365E-10 | 3.38126E-10 | 6.0771E-10 | 2.2068E-09 | 3.42E-10 | 2.81E-09 |
| | THC M | 9 | 1.7792E-09 | 1.39237E-09 | 4.64122E-10 | 7.0893E-10 | 2.8495E-09 | 4.67E-10 | 4.47E-09 |
| | THC+CBD M | 9 | 1.3672E-09 | 8.40122E-10 | 2.80041E-10 | 7.2147E-10 | 2.0130E-09 | 3.78E-10 | 2.46E-09 |

Supplemental Table 11.2 Descriptive statistics for *N*-acyl leucines (mol/g) in liver collected from male and female adult mice 2h after receiving an i.p. injection of 10mg/kg CBD, THC, or THC+CBD

| | | N | Mean | Std. Deviation | Std. Error | 95% Confidence Interval for Mean | | Minimum | Maximum |
|---|---|---|---|---|---|---|---|---|---|
| | | | | | | Lower Bound | Upper Bound | | |
| *N*-palmitoyl leucine | Veh F | 8 | 2.2231E-11 | 1.14580E-11 | 4.05102E-12 | 1.2651E-11 | 3.1810E-11 | 8.34E-12 | 3.99E-11 |
| | CBD F | 8 | 3.1644E-11 | 2.21559E-11 | 7.83329E-12 | 1.3121E-11 | 5.0166E-11 | 9.79E-12 | 6.98E-11 |
| | THC F | 10 | 2.5876E-11 | 9.31691E-12 | 2.94626E-12 | 1.9211E-11 | 3.2541E-11 | 1.35E-11 | 3.83E-11 |
| | THC+CBD F | 10 | 3.6621E-11 | 9.57477E-12 | 3.02781E-12 | 2.9771E-11 | 4.3470E-11 | 2.01E-11 | 5.02E-11 |
| | Veh M | 7 | 1.8968E-11 | 8.76815E-12 | 3.31405E-12 | 1.0859E-11 | 2.7077E-11 | 9.01E-12 | 3.44E-11 |
| | CBD M | 10 | 1.3450E-11 | 5.55550E-12 | 1.75680E-12 | 9.4759E-12 | 1.7424E-11 | 8.32E-12 | 2.41E-11 |
| | THC M | 9 | 3.2995E-11 | 1.53116E-11 | 5.10387E-12 | 2.1226E-11 | 4.4765E-11 | 1.55E-11 | 6.31E-11 |
| | THC+CBD M | 10 | 1.3080E-11 | 6.07929E-12 | 1.92244E-12 | 8.7308E-12 | 1.7429E-11 | 5.07E-12 | 2.56E-11 |
| *N*-stearoyl leucine | Veh F | 8 | 1.3818E-12 | 7.36644E-13 | 2.60443E-13 | 7.6594E-13 | 1.9976E-12 | 4.02E-13 | 2.75E-12 |
| | CBD F | 7 | 1.5425E-12 | 9.06144E-13 | 3.42490E-13 | 7.0449E-13 | 2.3806E-12 | 5.42E-13 | 3.22E-12 |
| | THC F | 10 | 1.5053E-12 | 6.35446E-13 | 2.00946E-13 | 1.0507E-12 | 1.9599E-12 | 6.39E-13 | 2.23E-12 |
| | THC+CBD F | 10 | 2.9610E-12 | 1.20364E-12 | 3.80623E-13 | 2.0999E-12 | 3.8220E-12 | 8.52E-13 | 4.49E-12 |
| | Veh M | 7 | 1.8354E-12 | 7.30183E-13 | 2.75983E-13 | 1.1601E-12 | 2.5108E-12 | 9.92E-13 | 2.77E-12 |
| | CBD M | 9 | 8.2481E-13 | 4.41912E-13 | 1.47304E-13 | 4.8513E-13 | 1.1645E-12 | 3.09E-13 | 1.57E-12 |
| | THC M | 8 | 1.6621E-12 | 9.28040E-13 | 3.28112E-13 | 8.8624E-13 | 2.4380E-12 | 3.26E-13 | 3.14E-12 |
| | THC+CBD M | 10 | 1.3417E-12 | 9.95096E-13 | 3.14677E-13 | 6.2987E-13 | 2.0536E-12 | 3.11E-13 | 3.70E-12 |
| *N*-oleoyl leucine | Veh F | 7 | 9.6788E-12 | 3.78357E-12 | 1.43005E-12 | 6.1796E-12 | 1.3178E-11 | 5.59E-12 | 1.48E-11 |
| | CBD F | 7 | 1.1937E-11 | 4.95869E-12 | 1.87421E-12 | 7.3511E-12 | 1.6523E-11 | 6.98E-12 | 2.09E-11 |
| | THC F | 10 | 1.8665E-11 | 9.39872E-12 | 2.97214E-12 | 1.1942E-11 | 2.5389E-11 | 8.58E-12 | 3.71E-11 |
| | THC+CBD F | 10 | 2.3830E-11 | 6.89305E-12 | 2.17977E-12 | 1.8899E-11 | 2.8761E-11 | 1.23E-11 | 3.24E-11 |
| | Veh M | 7 | 1.0163E-11 | 2.51565E-12 | 9.50825E-13 | 7.8363E-12 | 1.2489E-11 | 6.44E-12 | 1.28E-11 |
| | CBD M | 9 | 7.0013E-12 | 2.13142E-12 | 7.10475E-13 | 5.3629E-12 | 8.6396E-12 | 3.69E-12 | 1.06E-11 |
| | THC M | 8 | 1.3666E-11 | 4.62603E-12 | 1.63555E-12 | 9.7990E-12 | 1.7534E-11 | 8.14E-12 | 2.13E-11 |
| | THC+CBD M | 10 | 9.8180E-12 | 3.97840E-12 | 1.25808E-12 | 6.9720E-12 | 1.2664E-11 | 4.49E-12 | 1.79E-11 |
| *N*-linoleoyl leucine | Veh F | 8 | 1.0060E-11 | 5.52173E-12 | 1.95223E-12 | 5.4432E-12 | 1.4676E-11 | 3.05E-12 | 1.81E-11 |
| | CBD F | 8 | 1.6010E-11 | 1.49419E-11 | 5.28276E-12 | 3.5185E-12 | 2.8502E-11 | 3.28E-12 | 4.50E-11 |
| | THC F | 10 | 1.1590E-11 | 5.52303E-12 | 1.74654E-12 | 7.6391E-12 | 1.5541E-11 | 4.77E-12 | 2.22E-11 |
| | THC+CBD F | 10 | 2.1057E-11 | 6.74990E-12 | 2.13451E-12 | 1.6228E-11 | 2.5885E-11 | 1.10E-11 | 3.40E-11 |
| | Veh M | 7 | 5.9777E-12 | 3.43370E-12 | 1.29782E-12 | 2.8021E-12 | 9.1534E-12 | 1.79E-12 | 1.15E-11 |
| | CBD M | 10 | 6.0264E-12 | 3.26208E-12 | 1.03156E-12 | 3.6928E-12 | 8.3599E-12 | 1.82E-12 | 1.23E-11 |
| | THC M | 9 | 1.4883E-11 | 6.84857E-12 | 2.28286E-12 | 9.6188E-12 | 2.0147E-11 | 6.98E-12 | 2.64E-11 |
| | THC+CBD M | 10 | 8.4216E-12 | 4.00926E-12 | 1.26784E-12 | 5.5535E-12 | 1.1290E-11 | 3.29E-12 | 1.60E-11 |
| *N*-docosahexaenoyl leucine | Veh F | 8 | 1.1481E-11 | 5.27938E-12 | 1.86654E-12 | 7.0671E-12 | 1.5894E-11 | 4.28E-12 | 1.78E-11 |
| | CBD F | 8 | 1.7559E-11 | 1.55418E-11 | 5.49484E-12 | 4.5659E-12 | 3.0552E-11 | 2.89E-12 | 4.19E-11 |
| | THC F | 10 | 1.7081E-11 | 7.68272E-12 | 2.42949E-12 | 1.1585E-11 | 2.2577E-11 | 6.64E-12 | 2.64E-11 |
| | THC+CBD F | 10 | 2.4089E-11 | 1.04517E-11 | 3.30511E-12 | 1.6613E-11 | 3.1566E-11 | 1.04E-11 | 4.03E-11 |
| | Veh M | 7 | 9.7168E-12 | 5.25438E-12 | 1.98597E-12 | 4.8573E-12 | 1.4576E-11 | 2.82E-12 | 1.77E-11 |
| | CBD M | 9 | 5.8108E-12 | 2.06486E-12 | 6.88286E-13 | 4.2236E-12 | 7.3979E-12 | 3.28E-12 | 9.69E-12 |
| | THC M | 9 | 1.7924E-11 | 9.51748E-12 | 3.17249E-12 | 1.0609E-11 | 2.5240E-11 | 6.38E-12 | 3.30E-11 |
| | THC+CBD M | 10 | 1.0641E-11 | 7.38143E-12 | 2.33421E-12 | 5.3602E-12 | 1.5921E-11 | 2.86E-12 | 2.53E-11 |

Supplemental Table 11.3 Independent t-test comparing male and female levels of *N*-acyl leucines measured in plasma collected from adult mice 2h after an i.p. injection of 10mg/kg CBD, THC, or THC+CBD

| | | | Levene's Test for Equality of Variances | | t-test for Equality of Means | | | | | | |
|---|---|---|---|---|---|---|---|---|---|---|---|
| | | | | | | | Significance | | | 95% Confidence Interval of the Difference | |
| | | | F | Sig. | t | df | One-Sided p | Two-Sided p | Std. Error Difference | Lower | Upper |
| Veh | *N*-palmitoyl leucine | Equal variances assumed | 4.250 | 0.060 | -0.361 | 13 | 0.362 | 0.724 | 1.57984E-09 | -3.98381E-09 | 2.84228E-09 |
| | | Equal variances not assumed | | | -0.373 | 11.919 | 0.358 | 0.716 | 1.53158E-09 | -3.91031E-09 | 2.76878E-09 |
| | *N*-stearoyl leucine | Equal variances assumed | 5.811 | 0.031 | -0.733 | 13 | 0.238 | 0.477 | 6.11683E-10 | -1.76971E-09 | 8.73208E-10 |
| | | Equal variances not assumed | | | -0.757 | 11.745 | 0.232 | 0.464 | 5.91967E-10 | -1.74115E-09 | 8.44647E-10 |
| | *N*-oleoyl leucine | Equal variances assumed | 5.066 | 0.042 | -0.781 | 13 | 0.225 | 0.449 | 1.45747E-09 | -4.28629E-09 | 2.01107E-09 |
| | | Equal variances not assumed | | | -0.816 | 10.420 | 0.216 | 0.433 | 1.39455E-09 | -4.22799E-09 | 1.95277E-09 |
| | *N*-linoleoyl leucine | Equal variances assumed | 4.245 | 0.060 | 0.012 | 13 | 0.495 | 0.990 | 1.10904E-09 | -2.38213E-09 | 2.40973E-09 |
| | | Equal variances not assumed | | | 0.013 | 12.283 | 0.495 | 0.990 | 1.07951E-09 | -2.33226E-09 | 2.35986E-09 |
| | *N*-docosahexaenoyl leucine | Equal variances assumed | 11.457 | 0.005 | -1.016 | 12 | 0.165 | 0.330 | 4.30812E-10 | -1.37617E-09 | 5.01147E-10 |
| | | Equal variances not assumed | | | -1.016 | 8.143 | 0.170 | 0.339 | 4.30812E-10 | -1.42794E-09 | 5.52915E-10 |
| CBD | *N*-palmitoyl leucine | Equal variances assumed | 1.288 | 0.272 | -0.477 | 17 | 0.320 | 0.640 | 1.39682E-09 | -3.61273E-09 | 2.28132E-09 |
| | | Equal variances not assumed | | | -0.488 | 16.337 | 0.316 | 0.632 | 1.36532E-09 | -3.55520E-09 | 2.22379E-09 |
| | *N*-stearoyl leucine | Equal variances assumed | 1.053 | 0.318 | 0.275 | 18 | 0.393 | 0.786 | 5.28917E-10 | -9.65607E-10 | 1.25682E-09 |
| | | Equal variances not assumed | | | 0.267 | 14.477 | 0.397 | 0.793 | 5.45396E-10 | -1.02054E-09 | 1.31176E-09 |
| | *N*-oleoyl leucine | Equal variances assumed | 0.433 | 0.519 | -0.316 | 17 | 0.378 | 0.756 | 1.06834E-09 | -2.59133E-09 | 1.91667E-09 |

| | | | | | | | | | | |
|---|---|---|---|---|---|---|---|---|---|---|
| | | Equal variances not assumed | | | -0.320 | 15.860 | 0.377 | 0.753 | 1.05566E-09 | -2.57684E-09 | 1.90218E-09 |
| | *N*-linoleoyl leucine | Equal variances assumed | 1.268 | 0.276 | -0.528 | 17 | 0.302 | 0.605 | 1.01229E-09 | -2.66986E-09 | 1.60162E-09 |
| | | Equal variances not assumed | | | -0.545 | 16.642 | 0.297 | 0.593 | 9.80825E-10 | -2.60687E-09 | 1.53863E-09 |
| | *N*-docosahexaenoyl leucine | Equal variances assumed | 10.779 | 0.006 | -1.467 | 13 | 0.083 | 0.166 | 3.94284E-10 | -1.43018E-09 | 2.73414E-10 |
| | | Equal variances not assumed | | | -1.537 | 10.063 | 0.078 | 0.155 | 3.76208E-10 | -1.41592E-09 | 2.59152E-10 |
| THC | *N*-palmitoyl leucine | Equal variances assumed | 9.863 | 0.006 | -1.726 | 16 | 0.052 | 0.104 | 1.67140E-09 | -6.42727E-09 | 6.59141E-10 |
| | | Equal variances not assumed | | | -1.726 | 10.615 | 0.057 | 0.113 | 1.67140E-09 | -6.57914E-09 | 8.11016E-10 |
| | *N*-stearoyl leucine | Equal variances assumed | 13.074 | 0.002 | -1.539 | 16 | 0.072 | 0.143 | 5.97989E-10 | -2.18821E-09 | 3.47147E-10 |
| | | Equal variances not assumed | | | -1.539 | 11.359 | 0.076 | 0.151 | 5.97989E-10 | -2.23164E-09 | 3.90574E-10 |
| | *N*-oleoyl leucine | Equal variances assumed | 13.805 | 0.002 | -1.238 | 16 | 0.117 | 0.234 | 1.04418E-09 | -3.50587E-09 | 9.21231E-10 |
| | | Equal variances not assumed | | | -1.238 | 11.077 | 0.121 | 0.241 | 1.04418E-09 | -3.58859E-09 | 1.00395E-09 |
| | *N*-linoleoyl leucine | Equal variances assumed | 8.751 | 0.009 | -1.584 | 17 | 0.066 | 0.132 | 1.07972E-09 | -3.98843E-09 | 5.67574E-10 |
| | | Equal variances not assumed | | | -1.537 | 12.072 | 0.075 | 0.150 | 1.11295E-09 | -4.13373E-09 | 7.12875E-10 |
| | *N*-docosahexaenoyl leucine | Equal variances assumed | 2.446 | 0.137 | -0.643 | 16 | 0.265 | 0.529 | 5.41691E-10 | -1.49671E-09 | 7.99959E-10 |
| | | Equal variances not assumed | | | -0.643 | 13.123 | 0.266 | 0.531 | 5.41691E-10 | -1.51751E-09 | 8.20763E-10 |
| THC+CBD | *N*-palmitoyl leucine | Equal variances assumed | 7.981 | 0.011 | -1.355 | 18 | 0.096 | 0.192 | 1.86113E-09 | -6.43244E-09 | 1.38775E-09 |
| | | Equal variances not assumed | | | -1.355 | 11.580 | 0.101 | 0.201 | 1.86113E-09 | -6.59378E-09 | 1.54908E-09 |
| | *N*-stearoyl leucine | Equal variances assumed | 8.741 | 0.008 | -1.136 | 18 | 0.135 | 0.271 | 6.46540E-10 | -2.09294E-09 | 6.23721E-10 |

| | | | | | | | | | | |
|---|---|---|---|---|---|---|---|---|---|---|
| | Equal variances not assumed | | | -1.136 | 13.084 | 0.138 | 0.276 | 6.46540E-10 | -2.13047E-09 | 6.61246E-10 |
| *N*-oleoyl leucine | Equal variances assumed | 16.453 | 0.001 | -1.734 | 18 | 0.050 | 0.100 | 1.54959E-09 | -5.94318E-09 | 5.67971E-10 |
| | Equal variances not assumed | | | -1.734 | 10.810 | 0.056 | 0.111 | 1.54959E-09 | -6.10556E-09 | 7.30357E-10 |
| *N*-linoleoyl leucine | Equal variances assumed | 2.157 | 0.159 | -0.523 | 18 | 0.304 | 0.607 | 1.21887E-09 | -3.19804E-09 | 1.92345E-09 |
| | Equal variances not assumed | | | -0.523 | 15.127 | 0.304 | 0.609 | 1.21887E-09 | -3.23335E-09 | 1.95877E-09 |
| *N*-docosahexaenoyl leucine | Equal variances assumed | 5.392 | 0.033 | 1.043 | 17 | 0.156 | 0.311 | 4.91487E-10 | -5.24245E-10 | 1.54965E-09 |
| | Equal variances not assumed | | | 1.065 | 15.884 | 0.151 | 0.303 | 4.81473E-10 | -5.08582E-10 | 1.53398E-09 |

Supplemental Table 11.4 Independent t-test comparing male and female levels of *N*-acyl leucines measured in liver collected from adult mice 2h after an i.p. injection of 10mg/kg CBD, THC, or THC+CBD

| | | | Levene's Test for Equality of Variances | | t-test for Equality of Means | | | | | | | |
|---|---|---|---|---|---|---|---|---|---|---|---|---|
| | | | | | | | Significance | | | | 95% Confidence Interval of the Difference | |
| | | | F | Sig. | t | df | One-Sided p | Two-Sided p | Mean Difference | Std. Error Difference | Lower | Upper |
| Veh | *N*-palmitoyl leucine | Equal variances assumed | 1.020 | 0.331 | 0.612 | 13 | 0.276 | 0.551 | 3.26281E-12 | 5.33291E-12 | -8.25825E-12 | 1.47839E-11 |
| | | Equal variances not assumed | | | 0.623 | 12.811 | 0.272 | 0.544 | 3.26281E-12 | 5.23390E-12 | -8.06135E-12 | 1.45870E-11 |
| | *N*-stearoyl leucine | Equal variances assumed | 0.166 | 0.690 | -1.195 | 13 | 0.127 | 0.254 | -4.53651E-13 | 3.79710E-13 | -1.27396E-12 | 3.66662E-13 |
| | | Equal variances not assumed | | | -1.195 | 12.767 | 0.127 | 0.254 | -4.53651E-13 | 3.79470E-13 | -1.27497E-12 | 3.67669E-13 |
| | *N*-oleoyl leucine | Equal variances assumed | 2.941 | 0.112 | -0.282 | 12 | 0.391 | 0.783 | -4.84030E-13 | 1.71730E-12 | -4.22571E-12 | 3.25765E-12 |
| | | Equal variances not assumed | | | -0.282 | 10.438 | 0.392 | 0.784 | -4.84030E-13 | 1.71730E-12 | -4.28878E-12 | 3.32072E-12 |
| | *N*-linoleoyl leucine | Equal variances assumed | 1.726 | 0.212 | 1.687 | 13 | 0.058 | 0.115 | 4.08181E-12 | 2.41974E-12 | -1.14571E-12 | 9.30933E-12 |
| | | Equal variances not assumed | | | 1.741 | 11.853 | 0.054 | 0.108 | 4.08181E-12 | 2.34425E-12 | -1.03289E-12 | 9.19651E-12 |
| | *N*-docosahexaenoyl leucine | Equal variances assumed | 0.000 | 0.993 | 0.647 | 13 | 0.264 | 0.529 | 1.76395E-12 | 2.72637E-12 | -4.12603E-12 | 7.65392E-12 |
| | | Equal variances not assumed | | | 0.647 | 12.753 | 0.264 | 0.529 | 1.76395E-12 | 2.72545E-12 | -4.13565E-12 | 7.66355E-12 |
| CBD | *N*-palmitoyl leucine | Equal variances assumed | 14.636 | 0.001 | 2.517 | 16 | 0.011 | 0.023 | 1.81936E-11 | 7.22687E-12 | 2.87329E-12 | 3.35138E-11 |
| | | Equal variances not assumed | | | 2.266 | 7.707 | 0.027 | 0.054 | 1.81936E-11 | 8.02788E-12 | -4.42113E-13 | 3.68292E-11 |
| | *N*-stearoyl leucine | Equal variances assumed | 3.678 | 0.076 | 2.092 | 14 | 0.028 | 0.055 | 7.17720E-13 | 3.43092E-13 | -1.81383E-14 | 1.45358E-12 |
| | | Equal variances not assumed | | | 1.925 | 8.214 | 0.045 | 0.089 | 7.17720E-13 | 3.72824E-13 | -1.38126E-13 | 1.57357E-12 |
| | *N*-oleoyl leucine | Equal variances assumed | 6.806 | 0.021 | 2.703 | 14 | 0.009 | 0.017 | 4.93587E-12 | 1.82636E-12 | 1.01871E-12 | 8.85303E-12 |

| | | | | | | | | | | | | |
|---|---|---|---|---|---|---|---|---|---|---|---|---|
| | | Equal variances not assumed | | | 2.463 | 7.729 | 0.020 | 0.040 | 4.93587E-12 | 2.00435E-12 | 2.85427E-13 | 9.58632E-12 |
| | *N*-linoleoyl leucine | Equal variances assumed | 11.405 | 0.004 | 2.067 | 16 | 0.028 | 0.055 | 9.98391E-12 | 4.82949E-12 | -2.54143E-13 | 2.02220E-11 |
| | | Equal variances not assumed | | | 1.855 | 7.535 | 0.052 | 0.103 | 9.98391E-12 | 5.38253E-12 | -2.56268E-12 | 2.25305E-11 |
| | *N*-docosahexaenoyl leucine | Equal variances assumed | 11.950 | 0.004 | 2.255 | 15 | 0.020 | 0.040 | 1.17484E-11 | 5.21073E-12 | 6.42017E-13 | 2.28548E-11 |
| | | Equal variances not assumed | | | 2.122 | 7.220 | 0.035 | 0.070 | 1.17484E-11 | 5.53778E-12 | -1.26595E-12 | 2.47628E-11 |
| | *N*-palmitoyl leucine | Equal variances assumed | 1.573 | 0.227 | -1.239 | 17 | 0.116 | 0.232 | -7.11918E-12 | 5.74395E-12 | -1.92379E-11 | 4.99950E-12 |
| | | Equal variances not assumed | | | -1.208 | 12.943 | 0.124 | 0.249 | -7.11918E-12 | 5.89321E-12 | -1.98564E-11 | 5.61808E-12 |
| | *N*-stearoyl leucine | Equal variances assumed | 0.834 | 0.375 | -0.425 | 16 | 0.338 | 0.676 | -1.56797E-13 | 3.68626E-13 | -9.38249E-13 | 6.24655E-13 |
| | | Equal variances not assumed | | | -0.408 | 11.930 | 0.345 | 0.691 | -1.56797E-13 | 3.84755E-13 | -9.95649E-13 | 6.82055E-13 |
| | *N*-oleoyl leucine | Equal variances assumed | 4.431 | 0.051 | 1.371 | 16 | 0.095 | 0.189 | 4.99902E-12 | 3.64508E-12 | -2.72820E-12 | 1.27262E-11 |
| THC | | Equal variances not assumed | | | 1.474 | 13.665 | 0.082 | 0.163 | 4.99902E-12 | 3.39243E-12 | -2.29379E-12 | 1.22918E-11 |
| | *N*-linoleoyl leucine | Equal variances assumed | 1.089 | 0.311 | -1.159 | 17 | 0.131 | 0.262 | -3.29302E-12 | 2.84058E-12 | -9.28612E-12 | 2.70008E-12 |
| | | Equal variances not assumed | | | -1.146 | 15.412 | 0.135 | 0.269 | -3.29302E-12 | 2.87434E-12 | -9.40528E-12 | 2.81924E-12 |
| | *N*-docosahexaenoyl leucine | Equal variances assumed | 0.330 | 0.573 | -0.214 | 17 | 0.417 | 0.833 | -8.43609E-13 | 3.94916E-12 | -9.17561E-12 | 7.48839E-12 |
| | | Equal variances not assumed | | | -0.211 | 15.420 | 0.418 | 0.836 | -8.43609E-13 | 3.99589E-12 | -9.34047E-12 | 7.65325E-12 |
| | *N*-palmitoyl leucine | Equal variances assumed | 2.244 | 0.151 | 6.564 | 18 | 0.000 | 0.000 | 2.35410E-11 | 3.58656E-12 | 1.60059E-11 | 3.10760E-11 |
| THC+CBD | | Equal variances not assumed | | | 6.564 | 15.242 | 0.000 | 0.000 | 2.35410E-11 | 3.58656E-12 | 1.59070E-11 | 3.11750E-11 |
| | *N*-stearoyl leucine | Equal variances assumed | 0.866 | 0.364 | 3.279 | 18 | 0.002 | 0.004 | 1.61924E-12 | 4.93858E-13 | 5.81683E-13 | 2.65680E-12 |
| | | Equal variances | | | 3.279 | 17.386 | 0.002 | 0.004 | 1.61924E-12 | 4.93858E-13 | 5.79049E-13 | 2.65943E-12 |

| | | | | | | | | | | | |
|---|---|---|---|---|---|---|---|---|---|---|---|
| | not assumed | | | | | | | | | | |
| *N*-oleoyl leucine | Equal variances assumed | 3.955 | 0.062 | 5.568 | 18 | 0.000 | 0.000 | 1.40122E-11 | 2.51678E-12 | 8.72464E-12 | 1.92997E-11 |
| | Equal variances not assumed | | | 5.568 | 14.397 | 0.000 | 0.000 | 1.40122E-11 | 2.51678E-12 | 8.62818E-12 | 1.93962E-11 |
| *N*-linoleoyl leucine | Equal variances assumed | 0.721 | 0.407 | 5.089 | 18 | 0.000 | 0.000 | 1.26352E-11 | 2.48265E-12 | 7.41939E-12 | 1.78511E-11 |
| | Equal variances not assumed | | | 5.089 | 14.648 | 0.000 | 0.000 | 1.26352E-11 | 2.48265E-12 | 7.33249E-12 | 1.79380E-11 |
| *N*-docosahexaenoyl leucine | Equal variances assumed | 1.966 | 0.178 | 3.324 | 18 | 0.002 | 0.004 | 1.34489E-11 | 4.04627E-12 | 4.94797E-12 | 2.19498E-11 |
| | Equal variances not assumed | | | 3.324 | 16.189 | 0.002 | 0.004 | 1.34489E-11 | 4.04627E-12 | 4.87931E-12 | 2.20184E-11 |

Supplemental Table 11.5 One-way ANOVA for *N*-acyl leucines measured in plasma collected from adult mice 2h after an i.p. injection of 10mg/kg CBD, THC, or THC+CBD

| | | Sum of Squares | df | Mean Square | F | Sig. |
|---|---|---|---|---|---|---|
| *N*-palmitoyl leucine | Between Groups | 1.34E-16 | 7 | 1.92E-17 | 1.558 | 0.1643922 |
| | Within Groups | 7.88E-16 | 64 | 1.23E-17 | | |
| | Total | 9.22E-16 | 71 | | | |
| *N*-stearoyl leucine | Between Groups | 1.29E-17 | 7 | 1.84E-18 | 1.122 | 0.3603952 |
| | Within Groups | 1.06E-16 | 65 | 1.64E-18 | | |
| | Total | 1.19E-16 | 72 | | | |
| *N*-oleoyl leucine | Between Groups | 7.86E-17 | 7 | 1.12E-17 | 1.474 | 0.1924588 |
| | Within Groups | 4.88E-16 | 64 | 7.62E-18 | | |
| | Total | 5.66E-16 | 71 | | | |
| *N*-linoleoyl leucine | Between Groups | 3.42E-17 | 7 | 4.89E-18 | 0.864 | 0.5395065 |
| | Within Groups | 3.68E-16 | 65 | 5.66E-18 | | |
| | Total | 4.02E-16 | 72 | | | |
| *N*-docosahexaenoyl leucine | Between Groups | 7.91E-18 | 7 | 1.13E-18 | 1.172 | 0.3329729 |
| | Within Groups | 5.59E-17 | 58 | 9.64E-19 | | |
| | Total | 6.38E-17 | 65 | | | |

Supplemental Table 11.6 One-way ANOVA for *N*-acyl leucines measured in liver collected from adult mice 2h after an i.p. injection of 10mg/kg CBD, THC, or THC+CBD

| | | Sum of Squares | df | Mean Square | F | Sig. |
|---|---|---|---|---|---|---|
| *N*-palmitoyl leucine | Between Groups | 5.32E-21 | 7 | 7.61E-22 | 5.464 | 0.00006 |
| | Within Groups | 8.91E-21 | 64 | 1.39E-22 | | |
| | Total | 1.42E-20 | 71 | | | |
| *N*-stearoyl leucine | Between Groups | 2.54E-23 | 7 | 3.62E-24 | 4.902 | 0.00020 |
| | Within Groups | 4.51E-23 | 61 | 7.39E-25 | | |
| | Total | 7.05E-23 | 68 | | | |
| *N*-oleoyl leucine | Between Groups | 2.05E-21 | 7 | 2.92E-22 | 9.627 | 0.00000 |
| | Within Groups | 1.82E-21 | 60 | 3.04E-23 | | |
| | Total | 3.87E-21 | 67 | | | |
| *N*-linoleoyl leucine | Between Groups | 1.79E-21 | 7 | 2.56E-22 | 5.209 | 0.00010 |
| | Within Groups | 3.15E-21 | 64 | 4.92E-23 | | |
| | Total | 4.94E-21 | 71 | | | |
| *N*-docosahexaenoyl leucine | Between Groups | 2.23E-21 | 7 | 3.18E-22 | 4.164 | 0.00080 |
| | Within Groups | 4.82E-21 | 63 | 7.64E-23 | | |
| | Total | 7.04E-21 | 70 | | | |

#### Supplemental Table 11.7 Fisher's LSD post-hoc test for plasma *N*-acyl leucine ANOVA

| Dependent Variable | | | Std. Error | Sig. | 95% Confidence Interval Lower Bound | Upper Bound |
|---|---|---|---|---|---|---|
| *N*-palmitoyl leucine | VehFemale | CBDfemale | 1.82E-09 | 0.9152 | -3.43E-09 | 3.82E-09 |
| | | THCfemale | 1.77E-09 | 0.9639 | -3.45E-09 | 3.61E-09 |
| | | TpCfemale | 1.73E-09 | 0.5018 | -4.62E-09 | 2.29E-09 |
| | | VehMale | 1.82E-09 | 0.7543 | -4.20E-09 | 3.06E-09 |
| | | CBDmale | 1.70E-09 | 0.7819 | -3.86E-09 | 2.92E-09 |
| | | THCmale | 1.77E-09 | 0.1177 | -6.34E-09 | 7.28E-10 |
| | | TpCmale | 1.73E-09 | 0.0366 | -7.14E-09 | -2.36E-10 |
| | CBDfemale | VehFemale | 1.82E-09 | 0.9152 | -3.82E-09 | 3.43E-09 |
| | | THCfemale | 1.70E-09 | 0.9470 | -3.52E-09 | 3.29E-09 |
| | | TpCfemale | 1.66E-09 | 0.4162 | -4.69E-09 | 1.96E-09 |
| | | VehMale | 1.75E-09 | 0.6643 | -4.27E-09 | 2.74E-09 |
| | | CBDmale | 1.63E-09 | 0.6844 | -3.92E-09 | 2.59E-09 |
| | | THCmale | 1.70E-09 | 0.0834 | -6.40E-09 | 4.07E-10 |
| | | TpCmale | 1.66E-09 | 0.0227 | -7.21E-09 | -5.60E-10 |
| | THCfemale | VehFemale | 1.77E-09 | 0.9639 | -3.61E-09 | 3.45E-09 |
| | | CBDfemale | 1.70E-09 | 0.9470 | -3.29E-09 | 3.52E-09 |
| | | TpCfemale | 1.61E-09 | 0.4417 | -4.47E-09 | 1.97E-09 |
| | | VehMale | 1.70E-09 | 0.7038 | -4.06E-09 | 2.75E-09 |
| | | CBDmale | 1.58E-09 | 0.7275 | -3.70E-09 | 2.60E-09 |
| | | THCmale | 1.65E-09 | 0.0860 | -6.19E-09 | 4.20E-10 |
| | | TpCmale | 1.61E-09 | 0.0225 | -6.99E-09 | -5.50E-10 |
| | TpCfemale | VehFemale | 1.73E-09 | 0.5018 | -2.29E-09 | 4.62E-09 |
| | | CBDfemale | 1.66E-09 | 0.4162 | -1.96E-09 | 4.69E-09 |
| | | THCfemale | 1.61E-09 | 0.4417 | -1.97E-09 | 4.47E-09 |
| | | VehMale | 1.66E-09 | 0.7210 | -2.73E-09 | 3.92E-09 |
| | | CBDmale | 1.53E-09 | 0.6513 | -2.37E-09 | 3.76E-09 |
| | | THCmale | 1.61E-09 | 0.3139 | -4.86E-09 | 1.58E-09 |
| | | TpCmale | 1.57E-09 | 0.1128 | -5.66E-09 | 6.12E-10 |
| | VehMale | VehFemale | 1.82E-09 | 0.7543 | -3.06E-09 | 4.20E-09 |
| | | CBDfemale | 1.75E-09 | 0.6643 | -2.74E-09 | 4.27E-09 |
| | | THCfemale | 1.70E-09 | 0.7038 | -2.75E-09 | 4.06E-09 |
| | | TpCfemale | 1.66E-09 | 0.7210 | -3.92E-09 | 2.73E-09 |
| | | CBDmale | 1.63E-09 | 0.9517 | -3.16E-09 | 3.36E-09 |
| | | THCmale | 1.70E-09 | 0.1949 | -5.64E-09 | 1.17E-09 |
| | | TpCmale | 1.66E-09 | 0.0654 | -6.44E-09 | 2.05E-10 |
| | CBDmale | VehFemale | 1.70E-09 | 0.7819 | -2.92E-09 | 3.86E-09 |
| | | CBDfemale | 1.63E-09 | 0.6844 | -2.59E-09 | 3.92E-09 |
| | | THCfemale | 1.58E-09 | 0.7275 | -2.60E-09 | 3.70E-09 |
| | | TpCfemale | 1.53E-09 | 0.6513 | -3.76E-09 | 2.37E-09 |
| | | VehMale | 1.63E-09 | 0.9517 | -3.36E-09 | 3.16E-09 |
| | | THCmale | 1.58E-09 | 0.1440 | -5.48E-09 | 8.18E-10 |
| | | TpCmale | 1.53E-09 | 0.0397 | -6.28E-09 | -1.56E-10 |
| | THCmale | VehFemale | 1.77E-09 | 0.1177 | -7.28E-10 | 6.34E-09 |
| | | CBDfemale | 1.70E-09 | 0.0834 | -4.07E-10 | 6.40E-09 |
| | | THCfemale | 1.65E-09 | 0.0860 | -4.20E-10 | 6.19E-09 |
| | | TpCfemale | 1.61E-09 | 0.3139 | -1.58E-09 | 4.86E-09 |
| | | VehMale | 1.70E-09 | 0.1949 | -1.17E-09 | 5.64E-09 |
| | | CBDmale | 1.58E-09 | 0.1440 | -8.18E-10 | 5.48E-09 |
| | | TpCmale | 1.61E-09 | 0.5844 | -4.11E-09 | 2.33E-09 |
| | TpCmale | VehFemale | 1.73E-09 | 0.0366 | 2.36E-10 | 7.14E-09 |
| | | CBDfemale | 1.66E-09 | 0.0227 | 5.60E-10 | 7.21E-09 |
| | | THCfemale | 1.61E-09 | 0.0225 | 5.50E-10 | 6.99E-09 |
| | | TpCfemale | 1.57E-09 | 0.1128 | -6.12E-10 | 5.66E-09 |
| | | VehMale | 1.66E-09 | 0.0654 | -2.05E-10 | 6.44E-09 |
| | | CBDmale | 1.53E-09 | 0.0397 | 1.56E-10 | 6.28E-09 |
| | | THCmale | 1.61E-09 | 0.5844 | -2.33E-09 | 4.11E-09 |
| *N*-stearoyl leucine | VehFemale | CBDfemale | 6.45E-10 | 0.5331 | -1.69E-09 | 8.84E-10 |

| | | | | | | |
|---|---|---|---|---|---|---|
| | | THCfemale | 6.45E-10 | 0.8094 | -1.44E-09 | 1.13E-09 |
| | | TpCfemale | 6.31E-10 | 0.3724 | -1.83E-09 | 6.93E-10 |
| | | VehMale | 6.62E-10 | 0.5009 | -1.77E-09 | 8.75E-10 |
| | | CBDmale | 6.19E-10 | 0.6775 | -1.49E-09 | 9.77E-10 |
| | | THCmale | 6.45E-10 | 0.0998 | -2.36E-09 | 2.11E-10 |
| | | TpCmale | 6.31E-10 | 0.0431 | -2.56E-09 | -4.16E-11 |
| | CBDfemale | VehFemale | 6.45E-10 | 0.5331 | -8.84E-10 | 1.69E-09 |
| | | THCfemale | 6.03E-10 | 0.6825 | -9.57E-10 | 1.45E-09 |
| | | TpCfemale | 5.88E-10 | 0.7833 | -1.34E-09 | 1.01E-09 |
| | | VehMale | 6.22E-10 | 0.9436 | -1.29E-09 | 1.20E-09 |
| | | CBDmale | 5.75E-10 | 0.8010 | -1.00E-09 | 1.29E-09 |
| | | THCmale | 6.03E-10 | 0.2690 | -1.88E-09 | 5.32E-10 |
| | | TpCmale | 5.88E-10 | 0.1320 | -2.07E-09 | 2.77E-10 |
| | THCfemale | VehFemale | 6.45E-10 | 0.8094 | -1.13E-09 | 1.44E-09 |
| | | CBDfemale | 6.03E-10 | 0.6825 | -1.45E-09 | 9.57E-10 |
| | | TpCfemale | 5.88E-10 | 0.4878 | -1.58E-09 | 7.64E-10 |
| | | VehMale | 6.22E-10 | 0.6402 | -1.53E-09 | 9.50E-10 |
| | | CBDmale | 5.75E-10 | 0.8594 | -1.25E-09 | 1.05E-09 |
| | | THCmale | 6.03E-10 | 0.1319 | -2.13E-09 | 2.84E-10 |
| | | TpCmale | 5.88E-10 | 0.0558 | -2.32E-09 | 2.94E-11 |
| | TpCfemale | VehFemale | 6.31E-10 | 0.3724 | -6.93E-10 | 1.83E-09 |
| | | CBDfemale | 5.88E-10 | 0.7833 | -1.01E-09 | 1.34E-09 |
| | | THCfemale | 5.88E-10 | 0.4878 | -7.64E-10 | 1.58E-09 |
| | | VehMale | 6.07E-10 | 0.8462 | -1.09E-09 | 1.33E-09 |
| | | CBDmale | 5.59E-10 | 0.5837 | -8.09E-10 | 1.42E-09 |
| | | THCmale | 5.88E-10 | 0.3887 | -1.68E-09 | 6.64E-10 |
| | | TpCmale | 5.72E-10 | 0.2038 | -1.88E-09 | 4.08E-10 |
| | VehMale | VehFemale | 6.62E-10 | 0.5009 | -8.75E-10 | 1.77E-09 |
| | | CBDfemale | 6.22E-10 | 0.9436 | -1.20E-09 | 1.29E-09 |
| | | THCfemale | 6.22E-10 | 0.6402 | -9.50E-10 | 1.53E-09 |
| | | TpCfemale | 6.07E-10 | 0.8462 | -1.33E-09 | 1.09E-09 |
| | | CBDmale | 5.95E-10 | 0.7507 | -9.98E-10 | 1.38E-09 |
| | | THCmale | 6.22E-10 | 0.3159 | -1.87E-09 | 6.13E-10 |
| | | TpCmale | 6.07E-10 | 0.1648 | -2.07E-09 | 3.59E-10 |
| | CBDmale | VehFemale | 6.19E-10 | 0.6775 | -9.77E-10 | 1.49E-09 |
| | | CBDfemale | 5.75E-10 | 0.8010 | -1.29E-09 | 1.00E-09 |
| | | THCfemale | 5.75E-10 | 0.8594 | -1.05E-09 | 1.25E-09 |
| | | TpCfemale | 5.59E-10 | 0.5837 | -1.42E-09 | 8.09E-10 |
| | | VehMale | 5.95E-10 | 0.7507 | -1.38E-09 | 9.98E-10 |
| | | THCmale | 5.75E-10 | 0.1597 | -1.97E-09 | 3.31E-10 |
| | | TpCmale | 5.59E-10 | 0.0668 | -2.16E-09 | 7.41E-11 |
| | THCmale | VehFemale | 6.45E-10 | 0.0998 | -2.11E-10 | 2.36E-09 |
| | | CBDfemale | 6.03E-10 | 0.2690 | -5.32E-10 | 1.88E-09 |
| | | THCfemale | 6.03E-10 | 0.1319 | -2.84E-10 | 2.13E-09 |
| | | TpCfemale | 5.88E-10 | 0.3887 | -6.64E-10 | 1.68E-09 |
| | | VehMale | 6.22E-10 | 0.3159 | -6.13E-10 | 1.87E-09 |
| | | CBDmale | 5.75E-10 | 0.1597 | -3.31E-10 | 1.97E-09 |
| | | TpCmale | 5.88E-10 | 0.7040 | -1.40E-09 | 9.50E-10 |
| | TpCmale | VehFemale | 6.31E-10 | 0.0431 | 4.16E-11 | 2.56E-09 |
| | | CBDfemale | 5.88E-10 | 0.1320 | -2.77E-10 | 2.07E-09 |
| | | THCfemale | 5.88E-10 | 0.0558 | -2.94E-11 | 2.32E-09 |
| | | TpCfemale | 5.72E-10 | 0.2038 | -4.08E-10 | 1.88E-09 |
| | | VehMale | 6.07E-10 | 0.1648 | -3.59E-10 | 2.07E-09 |
| | | CBDmale | 5.59E-10 | 0.0668 | -7.41E-11 | 2.16E-09 |
| | | THCmale | 5.88E-10 | 0.7040 | -9.50E-10 | 1.40E-09 |
| *N*-oleoy leucine | VehFemale | CBDfemale | 1.43E-09 | 0.9129 | -3.01E-09 | 2.70E-09 |
| | | THCfemale | 1.39E-09 | 0.9117 | -2.93E-09 | 2.62E-09 |
| | | TpCfemale | 1.36E-09 | 0.6484 | -3.34E-09 | 2.09E-09 |
| | | VehMale | 1.43E-09 | 0.4288 | -3.99E-09 | 1.72E-09 |
| | | CBDmale | 1.33E-09 | 0.7123 | -3.16E-09 | 2.17E-09 |

| | | | | | | |
|---|---|---|---|---|---|---|
| | | THCmale | 1.39E-09 | 0.3021 | -4.23E-09 | 1.33E-09 |
| | | TpCmale | 1.36E-09 | 0.0177 | -6.03E-09 | -5.94E-10 |
| | CBDfemale | VehFemale | 1.43E-09 | 0.9129 | -2.70E-09 | 3.01E-09 |
| | | THCfemale | 1.34E-09 | 0.9988 | -2.68E-09 | 2.68E-09 |
| | | TpCfemale | 1.31E-09 | 0.7229 | -3.08E-09 | 2.15E-09 |
| | | VehMale | 1.38E-09 | 0.4799 | -3.74E-09 | 1.78E-09 |
| | | CBDmale | 1.28E-09 | 0.7934 | -2.90E-09 | 2.22E-09 |
| | | THCmale | 1.34E-09 | 0.3397 | -3.97E-09 | 1.39E-09 |
| | | TpCmale | 1.31E-09 | 0.0189 | -5.77E-09 | -5.38E-10 |
| | THCfemale | VehFemale | 1.39E-09 | 0.9117 | -2.62E-09 | 2.93E-09 |
| | | CBDfemale | 1.34E-09 | 0.9988 | -2.68E-09 | 2.68E-09 |
| | | TpCfemale | 1.27E-09 | 0.7131 | -3.00E-09 | 2.07E-09 |
| | | VehMale | 1.34E-09 | 0.4664 | -3.66E-09 | 1.70E-09 |
| | | CBDmale | 1.24E-09 | 0.7853 | -2.82E-09 | 2.14E-09 |
| | | THCmale | 1.30E-09 | 0.3243 | -3.89E-09 | 1.31E-09 |
| | | TpCmale | 1.27E-09 | 0.0154 | -5.69E-09 | -6.23E-10 |
| | TpCfemale | VehFemale | 1.36E-09 | 0.6484 | -2.09E-09 | 3.34E-09 |
| | | CBDfemale | 1.31E-09 | 0.7229 | -2.15E-09 | 3.08E-09 |
| | | THCfemale | 1.27E-09 | 0.7131 | -2.07E-09 | 3.00E-09 |
| | | VehMale | 1.31E-09 | 0.6957 | -3.13E-09 | 2.10E-09 |
| | | CBDmale | 1.21E-09 | 0.9152 | -2.28E-09 | 2.54E-09 |
| | | THCmale | 1.27E-09 | 0.5182 | -3.36E-09 | 1.71E-09 |
| | | TpCmale | 1.23E-09 | 0.0332 | -5.15E-09 | -2.22E-10 |
| | VehMale | VehFemale | 1.43E-09 | 0.4288 | -1.72E-09 | 3.99E-09 |
| | | CBDfemale | 1.38E-09 | 0.4799 | -1.78E-09 | 3.74E-09 |
| | | THCfemale | 1.34E-09 | 0.4664 | -1.70E-09 | 3.66E-09 |
| | | TpCfemale | 1.31E-09 | 0.6957 | -2.10E-09 | 3.13E-09 |
| | | CBDmale | 1.28E-09 | 0.6176 | -1.92E-09 | 3.21E-09 |
| | | THCmale | 1.34E-09 | 0.8182 | -2.99E-09 | 2.37E-09 |
| | | TpCmale | 1.31E-09 | 0.1018 | -4.79E-09 | 4.42E-10 |
| | CBDmale | VehFemale | 1.33E-09 | 0.7123 | -2.17E-09 | 3.16E-09 |
| | | CBDfemale | 1.28E-09 | 0.7934 | -2.22E-09 | 2.90E-09 |
| | | THCfemale | 1.24E-09 | 0.7853 | -2.14E-09 | 2.82E-09 |
| | | TpCfemale | 1.21E-09 | 0.9152 | -2.54E-09 | 2.28E-09 |
| | | VehMale | 1.28E-09 | 0.6176 | -3.21E-09 | 1.92E-09 |
| | | THCmale | 1.24E-09 | 0.4453 | -3.43E-09 | 1.53E-09 |
| | | TpCmale | 1.21E-09 | 0.0227 | -5.23E-09 | -4.07E-10 |
| | THCmale | VehFemale | 1.39E-09 | 0.3021 | -1.33E-09 | 4.23E-09 |
| | | CBDfemale | 1.34E-09 | 0.3397 | -1.39E-09 | 3.97E-09 |
| | | THCfemale | 1.30E-09 | 0.3243 | -1.31E-09 | 3.89E-09 |
| | | TpCfemale | 1.27E-09 | 0.5182 | -1.71E-09 | 3.36E-09 |
| | | VehMale | 1.34E-09 | 0.8182 | -2.37E-09 | 2.99E-09 |
| | | CBDmale | 1.24E-09 | 0.4453 | -1.53E-09 | 3.43E-09 |
| | | TpCmale | 1.27E-09 | 0.1466 | -4.40E-09 | 6.70E-10 |
| | TpCmale | VehFemale | 1.36E-09 | 0.0177 | 5.94E-10 | 6.03E-09 |
| | | CBDfemale | 1.31E-09 | 0.0189 | 5.38E-10 | 5.77E-09 |
| | | THCfemale | 1.27E-09 | 0.0154 | 6.23E-10 | 5.69E-09 |
| | | TpCfemale | 1.23E-09 | 0.0332 | 2.22E-10 | 5.15E-09 |
| | | VehMale | 1.31E-09 | 0.1018 | -4.42E-10 | 4.79E-09 |
| | | CBDmale | 1.21E-09 | 0.0227 | 4.07E-10 | 5.23E-09 |
| | | THCmale | 1.27E-09 | 0.1466 | -6.70E-10 | 4.40E-09 |
| *N*-linoleoyl leucine | VehFemale | CBDfemale | 1.23E-09 | 0.7391 | -2.05E-09 | 2.87E-09 |
| | | THCfemale | 1.17E-09 | 0.7043 | -1.89E-09 | 2.79E-09 |
| | | TpCfemale | 1.17E-09 | 0.5077 | -3.12E-09 | 1.56E-09 |
| | | VehMale | 1.23E-09 | 0.9911 | -2.45E-09 | 2.47E-09 |
| | | CBDmale | 1.15E-09 | 0.9157 | -2.42E-09 | 2.18E-09 |
| | | THCmale | 1.20E-09 | 0.2959 | -3.66E-09 | 1.13E-09 |
| | | TpCmale | 1.17E-09 | 0.2308 | -3.76E-09 | 9.23E-10 |
| | CBDfemale | VehFemale | 1.23E-09 | 0.7391 | -2.87E-09 | 2.05E-09 |
| | | THCfemale | 1.13E-09 | 0.9753 | -2.22E-09 | 2.29E-09 |

| | | | | | | |
|---|---|---|---|---|---|---|
| | | TpCfemale | 1.13E-09 | 0.2944 | -3.45E-09 | 1.06E-09 |
| | | VehMale | 1.19E-09 | 0.7389 | -2.77E-09 | 1.98E-09 |
| | | CBDmale | 1.11E-09 | 0.6306 | -2.74E-09 | 1.67E-09 |
| | | THCmale | 1.16E-09 | 0.1521 | -3.98E-09 | 6.34E-10 |
| | | TpCmale | 1.13E-09 | 0.1097 | -4.08E-09 | 4.24E-10 |
| | THCfemale | VehFemale | 1.17E-09 | 0.7043 | -2.79E-09 | 1.89E-09 |
| | | CBDfemale | 1.13E-09 | 0.9753 | -2.29E-09 | 2.22E-09 |
| | | TpCfemale | 1.06E-09 | 0.2527 | -3.35E-09 | 8.97E-10 |
| | | VehMale | 1.13E-09 | 0.7023 | -2.69E-09 | 1.82E-09 |
| | | CBDmale | 1.04E-09 | 0.5859 | -2.65E-09 | 1.51E-09 |
| | | THCmale | 1.09E-09 | 0.1225 | -3.89E-09 | 4.73E-10 |
| | | TpCmale | 1.06E-09 | 0.0843 | -3.99E-09 | 2.60E-10 |
| | TpCfemale | VehFemale | 1.17E-09 | 0.5077 | -1.56E-09 | 3.12E-09 |
| | | CBDfemale | 1.13E-09 | 0.2944 | -1.06E-09 | 3.45E-09 |
| | | THCfemale | 1.06E-09 | 0.2527 | -8.97E-10 | 3.35E-09 |
| | | VehMale | 1.13E-09 | 0.4838 | -1.46E-09 | 3.05E-09 |
| | | CBDmale | 1.04E-09 | 0.5284 | -1.42E-09 | 2.74E-09 |
| | | THCmale | 1.09E-09 | 0.6605 | -2.67E-09 | 1.70E-09 |
| | | TpCmale | 1.06E-09 | 0.5513 | -2.76E-09 | 1.49E-09 |
| | VehMale | VehFemale | 1.23E-09 | 0.9911 | -2.47E-09 | 2.45E-09 |
| | | CBDfemale | 1.19E-09 | 0.7389 | -1.98E-09 | 2.77E-09 |
| | | THCfemale | 1.13E-09 | 0.7023 | -1.82E-09 | 2.69E-09 |
| | | TpCfemale | 1.13E-09 | 0.4838 | -3.05E-09 | 1.46E-09 |
| | | CBDmale | 1.11E-09 | 0.9025 | -2.34E-09 | 2.07E-09 |
| | | THCmale | 1.16E-09 | 0.2734 | -3.59E-09 | 1.03E-09 |
| | | TpCmale | 1.13E-09 | 0.2090 | -3.69E-09 | 8.22E-10 |
| | CBDmale | VehFemale | 1.15E-09 | 0.9157 | -2.18E-09 | 2.42E-09 |
| | | CBDfemale | 1.11E-09 | 0.6306 | -1.67E-09 | 2.74E-09 |
| | | THCfemale | 1.04E-09 | 0.5859 | -1.51E-09 | 2.65E-09 |
| | | TpCfemale | 1.04E-09 | 0.5284 | -2.74E-09 | 1.42E-09 |
| | | VehMale | 1.11E-09 | 0.9025 | -2.07E-09 | 2.34E-09 |
| | | THCmale | 1.07E-09 | 0.2898 | -3.28E-09 | 9.95E-10 |
| | | TpCmale | 1.04E-09 | 0.2169 | -3.37E-09 | 7.80E-10 |
| | THCmale | VehFemale | 1.20E-09 | 0.2959 | -1.13E-09 | 3.66E-09 |
| | | CBDfemale | 1.16E-09 | 0.1521 | -6.34E-10 | 3.98E-09 |
| | | THCfemale | 1.09E-09 | 0.1225 | -4.73E-10 | 3.89E-09 |
| | | TpCfemale | 1.09E-09 | 0.6605 | -1.70E-09 | 2.67E-09 |
| | | VehMale | 1.16E-09 | 0.2734 | -1.03E-09 | 3.59E-09 |
| | | CBDmale | 1.07E-09 | 0.2898 | -9.95E-10 | 3.28E-09 |
| | | TpCmale | 1.09E-09 | 0.8877 | -2.34E-09 | 2.03E-09 |
| | TpCmale | VehFemale | 1.17E-09 | 0.2308 | -9.23E-10 | 3.76E-09 |
| | | CBDfemale | 1.13E-09 | 0.1097 | -4.24E-10 | 4.08E-09 |
| | | THCfemale | 1.06E-09 | 0.0843 | -2.60E-10 | 3.99E-09 |
| | | TpCfemale | 1.06E-09 | 0.5513 | -1.49E-09 | 2.76E-09 |
| | | VehMale | 1.13E-09 | 0.2090 | -8.22E-10 | 3.69E-09 |
| | | CBDmale | 1.04E-09 | 0.2169 | -7.80E-10 | 3.37E-09 |
| | | THCmale | 1.09E-09 | 0.8877 | -2.03E-09 | 2.34E-09 |
| *N*-docosahexaenoyl leucine | VehFemale | CBDfemale | 5.25E-10 | 0.9349 | -1.01E-09 | 1.09E-09 |
| | | THCfemale | 4.95E-10 | 0.2633 | -1.55E-09 | 4.32E-10 |
| | | TpCfemale | 4.84E-10 | 0.0417 | -1.98E-09 | -3.94E-11 |
| | | VehMale | 5.25E-10 | 0.4079 | -1.49E-09 | 6.13E-10 |
| | | CBDmale | 5.08E-10 | 0.2965 | -1.55E-09 | 4.82E-10 |
| | | THCmale | 4.95E-10 | 0.0719 | -1.90E-09 | 8.32E-11 |
| | | TpCmale | 4.95E-10 | 0.3210 | -1.49E-09 | 4.95E-10 |
| | CBDfemale | VehFemale | 5.25E-10 | 0.9349 | -1.09E-09 | 1.01E-09 |
| | | THCfemale | 4.95E-10 | 0.2287 | -1.59E-09 | 3.89E-10 |
| | | TpCfemale | 4.84E-10 | 0.0339 | -2.02E-09 | -8.25E-11 |
| | | VehMale | 5.25E-10 | 0.3636 | -1.53E-09 | 5.70E-10 |
| | | CBDmale | 5.08E-10 | 0.2597 | -1.60E-09 | 4.39E-10 |
| | | THCmale | 4.95E-10 | 0.0597 | -1.94E-09 | 4.02E-11 |

| | | | | | |
|---|---|---|---|---|---|
| | TpCmale | 4.95E-10 | 0.2811 | -1.53E-09 | 4.52E-10 |
| THCfemale | VehFemale | 4.95E-10 | 0.2633 | -4.32E-10 | 1.55E-09 |
| | CBDfemale | 4.95E-10 | 0.2287 | -3.89E-10 | 1.59E-09 |
| | TpCfemale | 4.51E-10 | 0.3236 | -1.35E-09 | 4.54E-10 |
| | VehMale | 4.95E-10 | 0.8071 | -8.69E-10 | 1.11E-09 |
| | CBDmale | 4.77E-10 | 0.9608 | -9.31E-10 | 9.79E-10 |
| | THCmale | 4.63E-10 | 0.4547 | -1.27E-09 | 5.78E-10 |
| | TpCmale | 4.63E-10 | 0.8912 | -8.63E-10 | 9.90E-10 |
| TpCfemale | VehFemale | 4.84E-10 | 0.0417 | 3.94E-11 | 1.98E-09 |
| | CBDfemale | 4.84E-10 | 0.0339 | 8.25E-11 | 2.02E-09 |
| | THCfemale | 4.51E-10 | 0.3236 | -4.54E-10 | 1.35E-09 |
| | VehMale | 4.84E-10 | 0.2432 | -3.98E-10 | 1.54E-09 |
| | CBDmale | 4.66E-10 | 0.3144 | -4.60E-10 | 1.41E-09 |
| | THCmale | 4.51E-10 | 0.8241 | -8.02E-10 | 1.00E-09 |
| | TpCmale | 4.51E-10 | 0.2604 | -3.90E-10 | 1.42E-09 |
| VehMale | VehFemale | 5.25E-10 | 0.4079 | -6.13E-10 | 1.49E-09 |
| | CBDfemale | 5.25E-10 | 0.3636 | -5.70E-10 | 1.53E-09 |
| | THCfemale | 4.95E-10 | 0.8071 | -1.11E-09 | 8.69E-10 |
| | TpCfemale | 4.84E-10 | 0.2432 | -1.54E-09 | 3.98E-10 |
| | CBDmale | 5.08E-10 | 0.8480 | -1.12E-09 | 9.19E-10 |
| | THCmale | 4.95E-10 | 0.3464 | -1.46E-09 | 5.21E-10 |
| | TpCmale | 4.95E-10 | 0.9074 | -1.05E-09 | 9.33E-10 |
| CBDmale | VehFemale | 5.08E-10 | 0.2965 | -4.82E-10 | 1.55E-09 |
| | CBDfemale | 5.08E-10 | 0.2597 | -4.39E-10 | 1.60E-09 |
| | THCfemale | 4.77E-10 | 0.9608 | -9.79E-10 | 9.31E-10 |
| | TpCfemale | 4.66E-10 | 0.3144 | -1.41E-09 | 4.60E-10 |
| | VehMale | 5.08E-10 | 0.8480 | -9.19E-10 | 1.12E-09 |
| | THCmale | 4.77E-10 | 0.4388 | -1.33E-09 | 5.83E-10 |
| | TpCmale | 4.77E-10 | 0.9335 | -9.15E-10 | 9.95E-10 |
| THCmale | VehFemale | 4.95E-10 | 0.0719 | -8.32E-11 | 1.90E-09 |
| | CBDfemale | 4.95E-10 | 0.0597 | -4.02E-11 | 1.94E-09 |
| | THCfemale | 4.63E-10 | 0.4547 | -5.78E-10 | 1.27E-09 |
| | TpCfemale | 4.51E-10 | 0.8241 | -1.00E-09 | 8.02E-10 |
| | VehMale | 4.95E-10 | 0.3464 | -5.21E-10 | 1.46E-09 |
| | CBDmale | 4.77E-10 | 0.4388 | -5.83E-10 | 1.33E-09 |
| | TpCmale | 4.63E-10 | 0.3771 | -5.15E-10 | 1.34E-09 |
| TpCmale | VehFemale | 4.95E-10 | 0.3210 | -4.95E-10 | 1.49E-09 |
| | CBDfemale | 4.95E-10 | 0.2811 | -4.52E-10 | 1.53E-09 |
| | THCfemale | 4.63E-10 | 0.8912 | -9.90E-10 | 8.63E-10 |
| | TpCfemale | 4.51E-10 | 0.2604 | -1.42E-09 | 3.90E-10 |
| | VehMale | 4.95E-10 | 0.9074 | -9.33E-10 | 1.05E-09 |
| | CBDmale | 4.77E-10 | 0.9335 | -9.95E-10 | 9.15E-10 |
| | THCmale | 4.63E-10 | 0.3771 | -1.34E-09 | 5.15E-10 |

#### Supplemental Table 11.8 Fisher's LSD post-hoc test for liver *N*-acyl leucine ANOVA

| Dependent Variable | | | Std. Error | Sig. | 95% Confidence Interval Lower Bound | Upper Bound |
|---|---|---|---|---|---|---|
| *N*-palmitoyl leucine | VehFemale | CBDfemale | 5.89914E-12 | 0.115 | -2.1198E-11 | 2.3719E-12 |
| | | THCfemale | 5.59641E-12 | 0.517 | -1.4826E-11 | 7.5344E-12 |
| | | TpCfemale | 5.59641E-12 | 0.012 | -2.5570E-11 | -3.2099E-12 |
| | | VehMale | 6.10619E-12 | 0.595 | -8.9357E-12 | 1.5461E-11 |
| | | CBDmale | 5.59641E-12 | 0.122 | -2.3995E-12 | 1.9961E-11 |
| | | THCmale | 5.73293E-12 | 0.065 | -2.2218E-11 | 6.8800E-13 |
| | | TpCmale | 5.59641E-12 | 0.107 | -2.0292E-12 | 2.0331E-11 |
| | CBDfemale | VehFemale | 5.89914E-12 | 0.115 | -2.3719E-12 | 2.1198E-11 |
| | | THCfemale | 5.59641E-12 | 0.307 | -5.4128E-12 | 1.6947E-11 |
| | | TpCfemale | 5.59641E-12 | 0.377 | -1.6157E-11 | 6.2031E-12 |
| | | VehMale | 6.10619E-12 | 0.042 | 4.7727E-13 | 2.4874E-11 |
| | | CBDmale | 5.59641E-12 | 0.002 | 7.0134E-12 | 2.9374E-11 |
| | | THCmale | 5.73293E-12 | 0.814 | -1.2805E-11 | 1.0101E-11 |
| | | TpCmale | 5.59641E-12 | 0.002 | 7.3838E-12 | 2.9744E-11 |
| | THCfemale | VehFemale | 5.59641E-12 | 0.517 | -7.5344E-12 | 1.4826E-11 |
| | | CBDfemale | 5.59641E-12 | 0.307 | -1.6947E-11 | 5.4128E-12 |
| | | TpCfemale | 5.27635E-12 | 0.046 | -2.1285E-11 | -2.0362E-13 |
| | | VehMale | 5.81425E-12 | 0.239 | -4.7068E-12 | 1.8524E-11 |
| | | CBDmale | 5.27635E-12 | 0.022 | 1.8855E-12 | 2.2967E-11 |
| | | THCmale | 5.42093E-12 | 0.194 | -1.7949E-11 | 3.7104E-12 |
| | | TpCmale | 5.27635E-12 | 0.018 | 2.2559E-12 | 2.3337E-11 |
| | TpCfemale | VehFemale | 5.59641E-12 | 0.012 | 3.2099E-12 | 2.5570E-11 |
| | | CBDfemale | 5.59641E-12 | 0.377 | -6.2031E-12 | 1.6157E-11 |
| | | THCfemale | 5.27635E-12 | 0.046 | 2.0362E-13 | 2.1285E-11 |
| | | VehMale | 5.81425E-12 | 0.003 | 6.0375E-12 | 2.9268E-11 |
| | | CBDmale | 5.27635E-12 | 0.000 | 1.2630E-11 | 3.3711E-11 |
| | | THCmale | 5.42093E-12 | 0.506 | -7.2044E-12 | 1.4455E-11 |
| | | TpCmale | 5.27635E-12 | 0.000 | 1.3000E-11 | 3.4082E-11 |
| | VehMale | VehFemale | 6.10619E-12 | 0.595 | -1.5461E-11 | 8.9357E-12 |
| | | CBDfemale | 6.10619E-12 | 0.042 | -2.4874E-11 | -4.7727E-13 |
| | | THCfemale | 5.81425E-12 | 0.239 | -1.8524E-11 | 4.7068E-12 |
| | | TpCfemale | 5.81425E-12 | 0.003 | -2.9268E-11 | -6.0375E-12 |
| | | CBDmale | 5.81425E-12 | 0.346 | -6.0975E-12 | 1.7133E-11 |
| | | THCmale | 5.94577E-12 | 0.021 | -2.5906E-11 | -2.1496E-12 |
| | | TpCmale | 5.81425E-12 | 0.315 | -5.7272E-12 | 1.7503E-11 |
| | CBDmale | VehFemale | 5.59641E-12 | 0.122 | -1.9961E-11 | 2.3995E-12 |
| | | CBDfemale | 5.59641E-12 | 0.002 | -2.9374E-11 | -7.0134E-12 |
| | | THCfemale | 5.27635E-12 | 0.022 | -2.2967E-11 | -1.8855E-12 |
| | | TpCfemale | 5.27635E-12 | 0.000 | -3.3711E-11 | -1.2630E-11 |
| | | VehMale | 5.81425E-12 | 0.346 | -1.7133E-11 | 6.0975E-12 |
| | | THCmale | 5.42093E-12 | 0.001 | -3.0375E-11 | -8.7159E-12 |
| | | TpCmale | 5.27635E-12 | 0.944 | -1.0170E-11 | 1.0911E-11 |
| | THCmale | VehFemale | 5.73293E-12 | 0.065 | -6.8800E-13 | 2.2218E-11 |
| | | CBDfemale | 5.73293E-12 | 0.814 | -1.0101E-11 | 1.2805E-11 |
| | | THCfemale | 5.42093E-12 | 0.194 | -3.7104E-12 | 1.7949E-11 |
| | | TpCfemale | 5.42093E-12 | 0.506 | -1.4455E-11 | 7.2044E-12 |
| | | VehMale | 5.94577E-12 | 0.021 | 2.1496E-12 | 2.5906E-11 |
| | | CBDmale | 5.42093E-12 | 0.001 | 8.7159E-12 | 3.0375E-11 |
| | | TpCmale | 5.42093E-12 | 0.000 | 9.0862E-12 | 3.0745E-11 |
| | TpCmale | VehFemale | 5.59641E-12 | 0.107 | -2.0331E-11 | 2.0292E-12 |
| | | CBDfemale | 5.59641E-12 | 0.002 | -2.9744E-11 | -7.3838E-12 |
| | | THCfemale | 5.27635E-12 | 0.018 | -2.3337E-11 | -2.2559E-12 |
| | | TpCfemale | 5.27635E-12 | 0.000 | -3.4082E-11 | -1.3000E-11 |
| | | VehMale | 5.81425E-12 | 0.315 | -1.7503E-11 | 5.7272E-12 |
| | | CBDmale | 5.27635E-12 | 0.944 | -1.0911E-11 | 1.0170E-11 |
| | | THCmale | 5.42093E-12 | 0.000 | -3.0745E-11 | -9.0862E-12 |
| *N*-stearoyl leucine | VehFemale | CBDfemale | 4.45015E-13 | 0.719 | -1.0506E-12 | 7.2912E-13 |

| | | | | | | |
|---|---|---|---|---|---|---|
| | | THCfemale | 4.07863E-13 | 0.763 | -9.3909E-13 | 6.9206E-13 |
| | | TpCfemale | 4.07863E-13 | 0.000 | -2.3947E-12 | -7.6360E-13 |
| | | VehMale | 4.45015E-13 | 0.312 | -1.3435E-12 | 4.3621E-13 |
| | | CBDmale | 4.17812E-13 | 0.187 | -2.7848E-13 | 1.3924E-12 |
| | | THCmale | 4.29925E-13 | 0.517 | -1.1400E-12 | 5.7938E-13 |
| | | TpCmale | 4.07863E-13 | 0.922 | -7.7550E-13 | 8.5564E-13 |
| | CBDfemale | VehFemale | 4.45015E-13 | 0.719 | -7.2912E-13 | 1.0506E-12 |
| | | THCfemale | 4.23739E-13 | 0.930 | -8.1010E-13 | 8.8454E-13 |
| | | TpCfemale | 4.23739E-13 | 0.001 | -2.2658E-12 | -5.7111E-13 |
| | | VehMale | 4.59609E-13 | 0.526 | -1.2120E-12 | 6.2613E-13 |
| | | CBDmale | 4.33324E-13 | 0.103 | -1.4876E-13 | 1.5842E-12 |
| | | THCmale | 4.45015E-13 | 0.789 | -1.0094E-12 | 7.7029E-13 |
| | | TpCmale | 4.23739E-13 | 0.637 | -6.4651E-13 | 1.0481E-12 |
| | THCfemale | VehFemale | 4.07863E-13 | 0.763 | -6.9206E-13 | 9.3909E-13 |
| | | CBDfemale | 4.23739E-13 | 0.930 | -8.8454E-13 | 8.1010E-13 |
| | | TpCfemale | 3.84537E-13 | 0.000 | -2.2246E-12 | -6.8673E-13 |
| | | VehMale | 4.23739E-13 | 0.439 | -1.1775E-12 | 5.1718E-13 |
| | | CBDmale | 3.95074E-13 | 0.090 | -1.0950E-13 | 1.4705E-12 |
| | | THCmale | 4.07863E-13 | 0.702 | -9.7237E-13 | 6.5878E-13 |
| | | TpCmale | 3.84537E-13 | 0.672 | -6.0534E-13 | 9.3251E-13 |
| | TpCfemale | VehFemale | 4.07863E-13 | 0.000 | 7.6360E-13 | 2.3947E-12 |
| | | CBDfemale | 4.23739E-13 | 0.001 | 5.7111E-13 | 2.2658E-12 |
| | | THCfemale | 3.84537E-13 | 0.000 | 6.8673E-13 | 2.2246E-12 |
| | | VehMale | 4.23739E-13 | 0.010 | 2.7820E-13 | 1.9728E-12 |
| | | CBDmale | 3.95074E-13 | 0.000 | 1.3462E-12 | 2.9262E-12 |
| | | THCmale | 4.07863E-13 | 0.002 | 4.8329E-13 | 2.1144E-12 |
| | | TpCmale | 3.84537E-13 | 0.000 | 8.5031E-13 | 2.3882E-12 |
| | VehMale | VehFemale | 4.45015E-13 | 0.312 | -4.3621E-13 | 1.3435E-12 |
| | | CBDfemale | 4.59609E-13 | 0.526 | -6.2613E-13 | 1.2120E-12 |
| | | THCfemale | 4.23739E-13 | 0.439 | -5.1718E-13 | 1.1775E-12 |
| | | TpCfemale | 4.23739E-13 | 0.010 | -1.9728E-12 | -2.7820E-13 |
| | | CBDmale | 4.33324E-13 | 0.023 | 1.4415E-13 | 1.8771E-12 |
| | | THCmale | 4.45015E-13 | 0.698 | -7.1652E-13 | 1.0632E-12 |
| | | TpCmale | 4.23739E-13 | 0.248 | -3.5360E-13 | 1.3410E-12 |
| | CBDmale | VehFemale | 4.17812E-13 | 0.187 | -1.3924E-12 | 2.7848E-13 |
| | | CBDfemale | 4.33324E-13 | 0.103 | -1.5842E-12 | 1.4876E-13 |
| | | THCfemale | 3.95074E-13 | 0.090 | -1.4705E-12 | 1.0950E-13 |
| | | TpCfemale | 3.95074E-13 | 0.000 | -2.9262E-12 | -1.3462E-12 |
| | | VehMale | 4.33324E-13 | 0.023 | -1.8771E-12 | -1.4415E-13 |
| | | THCmale | 4.17812E-13 | 0.050 | -1.6728E-12 | -1.8270E-15 |
| | | TpCmale | 3.95074E-13 | 0.196 | -1.3069E-12 | 2.7309E-13 |
| | THCmale | VehFemale | 4.29925E-13 | 0.517 | -5.7938E-13 | 1.1400E-12 |
| | | CBDfemale | 4.45015E-13 | 0.789 | -7.7029E-13 | 1.0094E-12 |
| | | THCfemale | 4.07863E-13 | 0.702 | -6.5878E-13 | 9.7237E-13 |
| | | TpCfemale | 4.07863E-13 | 0.002 | -2.1144E-12 | -4.8329E-13 |
| | | VehMale | 4.45015E-13 | 0.698 | -1.0632E-12 | 7.1652E-13 |
| | | CBDmale | 4.17812E-13 | 0.050 | 1.8270E-15 | 1.6728E-12 |
| | | TpCmale | 4.07863E-13 | 0.435 | -4.9519E-13 | 1.1360E-12 |
| | TpCmale | VehFemale | 4.07863E-13 | 0.922 | -8.5564E-13 | 7.7550E-13 |
| | | CBDfemale | 4.23739E-13 | 0.637 | -1.0481E-12 | 6.4651E-13 |
| | | THCfemale | 3.84537E-13 | 0.672 | -9.3251E-13 | 6.0534E-13 |
| | | TpCfemale | 3.84537E-13 | 0.000 | -2.3882E-12 | -8.5031E-13 |
| | | VehMale | 4.23739E-13 | 0.248 | -1.3410E-12 | 3.5360E-13 |
| | | CBDmale | 3.95074E-13 | 0.196 | -2.7309E-13 | 1.3069E-12 |
| | | THCmale | 4.07863E-13 | 0.435 | -1.1360E-12 | 4.9519E-13 |
| | VehFemale | CBDfemale | 2.94605E-12 | 0.446 | -8.1513E-12 | 3.6347E-12 |
| | | THCfemale | 2.71613E-12 | 0.002 | -1.4420E-11 | -3.5536E-12 |
| *N*-oleoy leucine | | TpCfemale | 2.71613E-12 | 0.000 | -1.9584E-11 | -8.7183E-12 |
| | | VehMale | 2.94605E-12 | 0.870 | -6.3770E-12 | 5.4090E-12 |
| | | CBDmale | 2.77757E-12 | 0.339 | -2.8784E-12 | 8.2335E-12 |

| | | | | | | |
|---|---|---|---|---|---|---|
| | | THCmale | 2.85250E-12 | 0.167 | -9.6935E-12 | 1.7182E-12 |
| | | TpCmale | 2.71613E-12 | 0.959 | -5.5722E-12 | 5.2939E-12 |
| | CBDfemale | VehFemale | 2.94605E-12 | 0.446 | -3.6347E-12 | 8.1513E-12 |
| | | THCfemale | 2.71613E-12 | 0.016 | -1.2161E-11 | -1.2953E-12 |
| | | TpCfemale | 2.71613E-12 | 0.000 | -1.7326E-11 | -6.4600E-12 |
| | | VehMale | 2.94605E-12 | 0.549 | -4.1187E-12 | 7.6673E-12 |
| | | CBDmale | 2.77757E-12 | 0.081 | -6.2009E-13 | 1.0492E-11 |
| | | THCmale | 2.85250E-12 | 0.547 | -7.4352E-12 | 3.9766E-12 |
| | | TpCmale | 2.71613E-12 | 0.438 | -3.3139E-12 | 7.5522E-12 |
| | THCfemale | VehFemale | 2.71613E-12 | 0.002 | 3.5536E-12 | 1.4420E-11 |
| | | CBDfemale | 2.71613E-12 | 0.016 | 1.2953E-12 | 1.2161E-11 |
| | | TpCfemale | 2.46485E-12 | 0.040 | -1.0095E-11 | -2.3429E-13 |
| | | VehMale | 2.71613E-12 | 0.003 | 3.0696E-12 | 1.3936E-11 |
| | | CBDmale | 2.53239E-12 | 0.000 | 6.5987E-12 | 1.6730E-11 |
| | | THCmale | 2.61436E-12 | 0.061 | -2.3048E-13 | 1.0229E-11 |
| | | TpCmale | 2.46485E-12 | 0.001 | 3.9171E-12 | 1.3778E-11 |
| | TpCfemale | VehFemale | 2.71613E-12 | 0.000 | 8.7183E-12 | 1.9584E-11 |
| | | CBDfemale | 2.71613E-12 | 0.000 | 6.4600E-12 | 1.7326E-11 |
| | | THCfemale | 2.46485E-12 | 0.040 | 2.3429E-13 | 1.0095E-11 |
| | | VehMale | 2.71613E-12 | 0.000 | 8.2343E-12 | 1.9100E-11 |
| | | CBDmale | 2.53239E-12 | 0.000 | 1.1763E-11 | 2.1894E-11 |
| | | THCmale | 2.61436E-12 | 0.000 | 4.9342E-12 | 1.5393E-11 |
| | | TpCmale | 2.46485E-12 | 0.000 | 9.0818E-12 | 1.8943E-11 |
| | VehMale | VehFemale | 2.94605E-12 | 0.870 | -5.4090E-12 | 6.3770E-12 |
| | | CBDfemale | 2.94605E-12 | 0.549 | -7.6673E-12 | 4.1187E-12 |
| | | THCfemale | 2.71613E-12 | 0.003 | -1.3936E-11 | -3.0696E-12 |
| | | TpCfemale | 2.71613E-12 | 0.000 | -1.9100E-11 | -8.2343E-12 |
| | | CBDmale | 2.77757E-12 | 0.260 | -2.3944E-12 | 8.7175E-12 |
| | | THCmale | 2.85250E-12 | 0.224 | -9.2095E-12 | 2.2023E-12 |
| | | TpCmale | 2.71613E-12 | 0.899 | -5.0882E-12 | 5.7779E-12 |
| | CBDmale | VehFemale | 2.77757E-12 | 0.339 | -8.2335E-12 | 2.8784E-12 |
| | | CBDfemale | 2.77757E-12 | 0.081 | -1.0492E-11 | 6.2009E-13 |
| | | THCfemale | 2.53239E-12 | 0.000 | -1.6730E-11 | -6.5987E-12 |
| | | TpCfemale | 2.53239E-12 | 0.000 | -2.1894E-11 | -1.1763E-11 |
| | | VehMale | 2.77757E-12 | 0.260 | -8.7175E-12 | 2.3944E-12 |
| | | THCmale | 2.67814E-12 | 0.016 | -1.2022E-11 | -1.3081E-12 |
| | | TpCmale | 2.53239E-12 | 0.270 | -7.8823E-12 | 2.2488E-12 |
| | THCmale | VehFemale | 2.85250E-12 | 0.167 | -1.7182E-12 | 9.6935E-12 |
| | | CBDfemale | 2.85250E-12 | 0.547 | -3.9766E-12 | 7.4352E-12 |
| | | THCfemale | 2.61436E-12 | 0.061 | -1.0229E-11 | 2.3048E-13 |
| | | TpCfemale | 2.61436E-12 | 0.000 | -1.5393E-11 | -4.9342E-12 |
| | | VehMale | 2.85250E-12 | 0.224 | -2.2023E-12 | 9.2095E-12 |
| | | CBDmale | 2.67814E-12 | 0.016 | 1.3081E-12 | 1.2022E-11 |
| | | TpCmale | 2.61436E-12 | 0.146 | -1.3811E-12 | 9.0780E-12 |
| | TpCmale | VehFemale | 2.71613E-12 | 0.959 | -5.2939E-12 | 5.5722E-12 |
| | | CBDfemale | 2.71613E-12 | 0.438 | -7.5522E-12 | 3.3139E-12 |
| | | THCfemale | 2.46485E-12 | 0.001 | -1.3778E-11 | -3.9171E-12 |
| | | TpCfemale | 2.46485E-12 | 0.000 | -1.8943E-11 | -9.0818E-12 |
| | | VehMale | 2.71613E-12 | 0.899 | -5.7779E-12 | 5.0882E-12 |
| | | CBDmale | 2.53239E-12 | 0.270 | -2.2488E-12 | 7.8823E-12 |
| | | THCmale | 2.61436E-12 | 0.146 | -9.0780E-12 | 1.3811E-12 |
| *N*-linoleoyl leucine | VehFemale | CBDfemale | 3.50626E-12 | 0.095 | -1.2955E-11 | 1.0538E-12 |
| | | THCfemale | 3.32633E-12 | 0.647 | -8.1756E-12 | 5.1146E-12 |
| | | TpCfemale | 3.32633E-12 | 0.002 | -1.7642E-11 | -4.3522E-12 |
| | | VehMale | 3.62933E-12 | 0.265 | -3.1686E-12 | 1.1332E-11 |
| | | CBDmale | 3.32633E-12 | 0.230 | -2.6119E-12 | 1.0678E-11 |
| | | THCmale | 3.40748E-12 | 0.162 | -1.1631E-11 | 1.9837E-12 |
| | | TpCmale | 3.32633E-12 | 0.624 | -5.0072E-12 | 8.2831E-12 |
| | CBDfemale | VehFemale | 3.50626E-12 | 0.095 | -1.0538E-12 | 1.2955E-11 |
| | | THCfemale | 3.32633E-12 | 0.189 | -2.2249E-12 | 1.1065E-11 |

| | | | | | | |
|---|---|---|---|---|---|---|
| | | TpCfemale | 3.32633E-12 | 0.134 | -1.1692E-11 | 1.5986E-12 |
| | | VehMale | 3.62933E-12 | 0.007 | 2.7821E-12 | 1.7283E-11 |
| | | CBDmale | 3.32633E-12 | 0.004 | 3.3388E-12 | 1.6629E-11 |
| | | THCmale | 3.40748E-12 | 0.742 | -5.6800E-12 | 7.9344E-12 |
| | | TpCmale | 3.32633E-12 | 0.026 | 9.4356E-13 | 1.4234E-11 |
| | THCfemale | VehFemale | 3.32633E-12 | 0.647 | -5.1146E-12 | 8.1756E-12 |
| | | CBDfemale | 3.32633E-12 | 0.189 | -1.1065E-11 | 2.2249E-12 |
| | | TpCfemale | 3.13610E-12 | 0.004 | -1.5732E-11 | -3.2017E-12 |
| | | VehMale | 3.45581E-12 | 0.109 | -1.2914E-12 | 1.2516E-11 |
| | | CBDmale | 3.13610E-12 | 0.081 | -7.0137E-13 | 1.1829E-11 |
| | | THCmale | 3.22203E-12 | 0.311 | -9.7298E-12 | 3.1437E-12 |
| | | TpCmale | 3.13610E-12 | 0.316 | -3.0966E-12 | 9.4335E-12 |
| | TpCfemale | VehFemale | 3.32633E-12 | 0.002 | 4.3522E-12 | 1.7642E-11 |
| | | CBDfemale | 3.32633E-12 | 0.134 | -1.5986E-12 | 1.1692E-11 |
| | | THCfemale | 3.13610E-12 | 0.004 | 3.2017E-12 | 1.5732E-11 |
| | | VehMale | 3.45581E-12 | 0.000 | 8.1753E-12 | 2.1983E-11 |
| | | CBDmale | 3.13610E-12 | 0.000 | 8.7654E-12 | 2.1296E-11 |
| | | THCmale | 3.22203E-12 | 0.060 | -2.6301E-13 | 1.2611E-11 |
| | | TpCmale | 3.13610E-12 | 0.000 | 6.3702E-12 | 1.8900E-11 |
| | VehMale | VehFemale | 3.62933E-12 | 0.265 | -1.1332E-11 | 3.1686E-12 |
| | | CBDfemale | 3.62933E-12 | 0.007 | -1.7283E-11 | -2.7821E-12 |
| | | THCfemale | 3.45581E-12 | 0.109 | -1.2516E-11 | 1.2914E-12 |
| | | TpCfemale | 3.45581E-12 | 0.000 | -2.1983E-11 | -8.1753E-12 |
| | | CBDmale | 3.45581E-12 | 0.989 | -6.9524E-12 | 6.8551E-12 |
| | | THCmale | 3.53398E-12 | 0.014 | -1.5965E-11 | -1.8454E-12 |
| | | TpCmale | 3.45581E-12 | 0.482 | -9.3477E-12 | 4.4599E-12 |
| | CBDmale | VehFemale | 3.32633E-12 | 0.230 | -1.0678E-11 | 2.6119E-12 |
| | | CBDfemale | 3.32633E-12 | 0.004 | -1.6629E-11 | -3.3388E-12 |
| | | THCfemale | 3.13610E-12 | 0.081 | -1.1829E-11 | 7.0137E-13 |
| | | TpCfemale | 3.13610E-12 | 0.000 | -2.1296E-11 | -8.7654E-12 |
| | | VehMale | 3.45581E-12 | 0.989 | -6.8551E-12 | 6.9524E-12 |
| | | THCmale | 3.22203E-12 | 0.008 | -1.5293E-11 | -2.4200E-12 |
| | | TpCmale | 3.13610E-12 | 0.448 | -8.6603E-12 | 3.8698E-12 |
| | THCmale | VehFemale | 3.40748E-12 | 0.162 | -1.9837E-12 | 1.1631E-11 |
| | | CBDfemale | 3.40748E-12 | 0.742 | -7.9344E-12 | 5.6800E-12 |
| | | THCfemale | 3.22203E-12 | 0.311 | -3.1437E-12 | 9.7298E-12 |
| | | TpCfemale | 3.22203E-12 | 0.060 | -1.2611E-11 | 2.6301E-13 |
| | | VehMale | 3.53398E-12 | 0.014 | 1.8454E-12 | 1.5965E-11 |
| | | CBDmale | 3.22203E-12 | 0.008 | 2.4200E-12 | 1.5293E-11 |
| | | TpCmale | 3.22203E-12 | 0.049 | 2.4735E-14 | 1.2898E-11 |
| | TpCmale | VehFemale | 3.32633E-12 | 0.624 | -8.2831E-12 | 5.0072E-12 |
| | | CBDfemale | 3.32633E-12 | 0.026 | -1.4234E-11 | -9.4356E-13 |
| | | THCfemale | 3.13610E-12 | 0.316 | -9.4335E-12 | 3.0966E-12 |
| | | TpCfemale | 3.13610E-12 | 0.000 | -1.8900E-11 | -6.3702E-12 |
| | | VehMale | 3.45581E-12 | 0.482 | -4.4599E-12 | 9.3477E-12 |
| | | CBDmale | 3.13610E-12 | 0.448 | -3.8698E-12 | 8.6603E-12 |
| | | THCmale | 3.22203E-12 | 0.049 | -1.2898E-11 | -2.4735E-14 |
| *N*-docosahexaenoyl leucine | VehFemale | CBDfemale | 4.37120E-12 | 0.169 | -1.4814E-11 | 2.6567E-12 |
| | | THCfemale | 4.14689E-12 | 0.182 | -1.3887E-11 | 2.6869E-12 |
| | | TpCfemale | 4.14689E-12 | 0.003 | -2.0896E-11 | -4.3218E-12 |
| | | VehMale | 4.52463E-12 | 0.698 | -7.2778E-12 | 1.0806E-11 |
| | | CBDmale | 4.24805E-12 | 0.187 | -2.8191E-12 | 1.4159E-11 |
| | | THCmale | 4.24805E-12 | 0.134 | -1.4933E-11 | 2.0455E-12 |
| | | TpCmale | 4.14689E-12 | 0.840 | -7.4467E-12 | 9.1271E-12 |
| | CBDfemale | VehFemale | 4.37120E-12 | 0.169 | -2.6567E-12 | 1.4814E-11 |
| | | THCfemale | 4.14689E-12 | 0.909 | -7.8084E-12 | 8.7654E-12 |
| | | TpCfemale | 4.14689E-12 | 0.120 | -1.4817E-11 | 1.7567E-12 |
| | | VehMale | 4.52463E-12 | 0.088 | -1.1994E-12 | 1.6884E-11 |
| | | CBDmale | 4.24805E-12 | 0.007 | 3.2594E-12 | 2.0237E-11 |
| | | THCmale | 4.24805E-12 | 0.932 | -8.8542E-12 | 8.1239E-12 |

| | | | | | |
|---|---|---|---|---|---|
| | TpCmale | 4.14689E-12 | 0.100 | -1.3682E-12 | 1.5206E-11 |
| THCfemale | VehFemale | 4.14689E-12 | 0.182 | -2.6869E-12 | 1.3887E-11 |
| | CBDfemale | 4.14689E-12 | 0.909 | -8.7654E-12 | 7.8084E-12 |
| | TpCfemale | 3.90972E-12 | 0.078 | -1.4822E-11 | 8.0425E-13 |
| | VehMale | 4.30831E-12 | 0.092 | -1.2456E-12 | 1.5973E-11 |
| | CBDmale | 4.01686E-12 | 0.007 | 3.2429E-12 | 1.9297E-11 |
| | THCmale | 4.01686E-12 | 0.834 | -8.8707E-12 | 7.1834E-12 |
| | TpCmale | 3.90972E-12 | 0.104 | -1.3728E-12 | 1.4253E-11 |
| TpCfemale | VehFemale | 4.14689E-12 | 0.003 | 4.3218E-12 | 2.0896E-11 |
| | CBDfemale | 4.14689E-12 | 0.120 | -1.7567E-12 | 1.4817E-11 |
| | THCfemale | 3.90972E-12 | 0.078 | -8.0425E-13 | 1.4822E-11 |
| | VehMale | 4.30831E-12 | 0.001 | 5.7631E-12 | 2.2982E-11 |
| | CBDmale | 4.01686E-12 | 0.000 | 1.0252E-11 | 2.6306E-11 |
| | THCmale | 4.01686E-12 | 0.130 | -1.8620E-12 | 1.4192E-11 |
| | TpCmale | 3.90972E-12 | 0.001 | 5.6359E-12 | 2.1262E-11 |
| VehMale | VehFemale | 4.52463E-12 | 0.698 | -1.0806E-11 | 7.2778E-12 |
| | CBDfemale | 4.52463E-12 | 0.088 | -1.6884E-11 | 1.1994E-12 |
| | THCfemale | 4.30831E-12 | 0.092 | -1.5973E-11 | 1.2456E-12 |
| | TpCfemale | 4.30831E-12 | 0.001 | -2.2982E-11 | -5.7631E-12 |
| | CBDmale | 4.40576E-12 | 0.379 | -4.8982E-12 | 1.2710E-11 |
| | THCmale | 4.40576E-12 | 0.067 | -1.7012E-11 | 5.9670E-13 |
| | TpCmale | 4.30831E-12 | 0.831 | -9.5332E-12 | 7.6857E-12 |
| CBDmale | VehFemale | 4.24805E-12 | 0.187 | -1.4159E-11 | 2.8191E-12 |
| | CBDfemale | 4.24805E-12 | 0.007 | -2.0237E-11 | -3.2594E-12 |
| | THCfemale | 4.01686E-12 | 0.007 | -1.9297E-11 | -3.2429E-12 |
| | TpCfemale | 4.01686E-12 | 0.000 | -2.6306E-11 | -1.0252E-11 |
| | VehMale | 4.40576E-12 | 0.379 | -1.2710E-11 | 4.8982E-12 |
| | THCmale | 4.12121E-12 | 0.005 | -2.0349E-11 | -3.8780E-12 |
| | TpCmale | 4.01686E-12 | 0.234 | -1.2857E-11 | 3.1973E-12 |
| THCmale | VehFemale | 4.24805E-12 | 0.134 | -2.0455E-12 | 1.4933E-11 |
| | CBDfemale | 4.24805E-12 | 0.932 | -8.1239E-12 | 8.8542E-12 |
| | THCfemale | 4.01686E-12 | 0.834 | -7.1834E-12 | 8.8707E-12 |
| | TpCfemale | 4.01686E-12 | 0.130 | -1.4192E-11 | 1.8620E-12 |
| | VehMale | 4.40576E-12 | 0.067 | -5.9670E-13 | 1.7012E-11 |
| | CBDmale | 4.12121E-12 | 0.005 | 3.8780E-12 | 2.0349E-11 |
| | TpCmale | 4.01686E-12 | 0.075 | -7.4329E-13 | 1.5311E-11 |
| TpCmale | VehFemale | 4.14689E-12 | 0.840 | -9.1271E-12 | 7.4467E-12 |
| | CBDfemale | 4.14689E-12 | 0.100 | -1.5206E-11 | 1.3682E-12 |
| | THCfemale | 3.90972E-12 | 0.104 | -1.4253E-11 | 1.3728E-12 |
| | TpCfemale | 3.90972E-12 | 0.001 | -2.1262E-11 | -5.6359E-12 |
| | VehMale | 4.30831E-12 | 0.831 | -7.6857E-12 | 9.5332E-12 |
| | CBDmale | 4.01686E-12 | 0.234 | -3.1973E-12 | 1.2857E-11 |
| | THCmale | 4.01686E-12 | 0.075 | -1.5311E-11 | 7.4329E-13 |

Supplemental Table 12.1 Descriptive statistics for *N*-acyl prolines (mol/L) in plasma collected from male and female adult mice 2h after receiving an i.p. injection of 10mg/kg CBD, THC, or THC+CBD

| | | N | Mean | Std. Deviation | Std. Error | 95% Confidence Interval for Mean | | Minimum | Maximum |
|---|---|---|---|---|---|---|---|---|---|
| | | | | | | Lower Bound | Upper Bound | | |
| *N*-palmitoyl proline | Veh F | BAL | | | | | | | |
| | CBD F | BAL | | | | | | | |
| | THC F | BAL | | | | | | | |
| | THC+CBD F | 4 | 2.7072E-09 | 1.35663E-09 | 6.78316E-10 | 5.4851E-10 | 4.8659E-09 | 8.67E-10 | 4.05E-09 |
| | Veh M | BAL | | | | | | | |
| | CBD M | 8 | 1.9845E-09 | 5.20301E-10 | 1.83954E-10 | 1.5495E-09 | 2.4195E-09 | 1.26E-09 | 2.96E-09 |
| | THC M | 4 | 1.8455E-09 | 3.48592E-10 | 1.74296E-10 | 1.2908E-09 | 2.4002E-09 | 1.40E-09 | 2.22E-09 |
| | THC+CBD M | BAL | | | | | | | |
| *N*-stearoyl proline | Veh F | BAL | | | | | | | |
| | CBD F | BAL | | | | | | | |
| | THC F | BAL | | | | | | | |
| | THC+CBD F | BDL | | | | | | | |
| | Veh M | BAL | | | | | | | |
| | CBD M | BAL | | | | | | | |
| | THC M | BAL | | | | | | | |
| | THC+CBD M | BAL | | | | | | | |
| *N*-oleoyl proline | Veh F | BDL | | | | | | | |
| | CBD F | BDL | | | | | | | |
| | THC F | BDL | | | | | | | |
| | THC+CBD F | BDL | | | | | | | |
| | Veh M | BDL | | | | | | | |
| | CBD M | BDL | | | | | | | |
| | THC M | BAL | | | | | | | |
| | THC+CBD M | BDL | | | | | | | |
| *N*-linoleoyl proline | Veh F | BDL | | | | | | | |
| | CBD F | BAL | | | | | | | |
| | THC F | BDL | | | | | | | |
| | THC+CBD F | BDL | | | | | | | |
| | Veh M | BDL | | | | | | | |
| | CBD M | BAL | | | | | | | |
| | THC M | BDL | | | | | | | |
| | THC+CBD M | BDL | | | | | | | |
| *N*-arachidonoyl proline | Veh F | BAL | | | | | | | |
| | CBD F | BDL | | | | | | | |
| | THC F | BDL | | | | | | | |
| | THC+CBD F | BDL | | | | | | | |
| | Veh M | BDL | | | | | | | |
| | CBD M | BDL | | | | | | | |
| | THC M | BDL | | | | | | | |
| | THC+CBD M | BDL | | | | | | | |
| | Veh F | BDL | | | | | | | |

| | | |
|---|---|---|
| *N*-docosahexaenoyl proline | CBD F | BAL |
| | THC F | BDL |
| | THC+CBD F | BDL |
| | Veh M | BDL |
| | CBD M | BDL |
| | THC M | BDL |
| | THC+CBD M | BAL |

Supplemental Table 12.2 Descriptive statistics for *N*-acyl prolines (mol/L) in liver collected from male and female adult mice 2h after receiving an i.p. injection of 10mg/kg CBD, THC, or THC+CBD

| | | N | Mean | Std. Deviation | Std. Error | 95% Confidence Interval for Mean | | Minimum | Maximum |
|---|---|---|---|---|---|---|---|---|---|
| | | | | | | Lower Bound | Upper Bound | | |
| *N*-palmitoyl proline | Veh F | 8 | 6.4666E-12 | 3.84600E-12 | 1.35977E-12 | 3.2513E-12 | 9.6819E-12 | 2.37E-12 | 1.29E-11 |
| | CBD F | 7 | 7.1635E-12 | 2.74715E-12 | 1.03832E-12 | 4.6229E-12 | 9.7042E-12 | 4.67E-12 | 1.12E-11 |
| | THC F | 7 | 3.6143E-12 | 1.40127E-12 | 5.29632E-13 | 2.3183E-12 | 4.9103E-12 | 2.03E-12 | 5.48E-12 |
| | THC+CBD F | 7 | 3.1470E-12 | 7.87197E-13 | 2.97532E-13 | 2.4190E-12 | 3.8751E-12 | 1.60E-12 | 3.76E-12 |
| | Veh M | 5 | 2.7857E-12 | 6.40408E-13 | 2.86399E-13 | 1.9905E-12 | 3.5808E-12 | 2.11E-12 | 3.69E-12 |
| | CBD M | 7 | 4.6709E-12 | 2.90833E-12 | 1.09925E-12 | 1.9811E-12 | 7.3606E-12 | 1.75E-12 | 9.96E-12 |
| | THC M | 7 | 5.8458E-12 | 2.69377E-12 | 1.01815E-12 | 3.3545E-12 | 8.3371E-12 | 3.04E-12 | 1.04E-11 |
| | THC+CBD M | 11 | 5.7903E-12 | 3.03013E-12 | 9.13620E-13 | 3.7547E-12 | 7.8260E-12 | 2.52E-12 | 1.13E-11 |
| *N*-stearoyl proline | Veh F | BAL | | | | | | | |
| | CBD F | BAL | | | | | | | |
| | THC F | 4 | 1.7167E-12 | 7.20216E-13 | 3.60108E-13 | 5.7069E-13 | 2.8627E-12 | 9.90E-13 | 2.71E-12 |
| | THC+CBD F | BAL | | | | | | | |
| | Veh M | BAL | | | | | | | |
| | CBD M | BAL | | | | | | | |
| | THC M | BAL | | | | | | | |
| | THC+CBD M | BAL | | | | | | | |
| *N*-oleoyl proline | Veh F | 4 | 3.6600E-12 | 1.20852E-12 | 6.04260E-13 | 1.7370E-12 | 5.5831E-12 | 2.44E-12 | 5.28E-12 |
| | CBD F | 4 | 5.7299E-12 | 1.67913E-12 | 8.39564E-13 | 3.0580E-12 | 8.4017E-12 | 3.34E-12 | 7.08E-12 |
| | THC F | BAL | | | | | | | |
| | THC+CBD F | 7 | 4.4027E-12 | 1.84280E-12 | 6.96514E-13 | 2.6984E-12 | 6.1070E-12 | 1.82E-12 | 7.26E-12 |
| | Veh M | BAL | | | | | | | |
| | CBD M | BAL | | | | | | | |
| | THC M | 4 | 5.7728E-12 | 3.81627E-12 | 1.90813E-12 | -2.9971E-13 | 1.1845E-11 | 2.44E-12 | 1.12E-11 |
| | THC+CBD M | BAL | | | | | | | |
| *N*-linoleoyl proline | Veh F | BAL | | | | | | | |
| | CBD F | BAL | | | | | | | |
| | THC F | BDL | | | | | | | |
| | THC+CBD F | BAL | 1.4952E-12 | 3.82934E-13 | 2.70775E-13 | -1.9453E-12 | 4.9358E-12 | 1.22E-12 | 1.77E-12 |
| | Veh M | 5 | 1.4922E-12 | 1.03316E-12 | 4.62041E-13 | 2.0938E-13 | 2.7750E-12 | 4.38E-13 | 2.59E-12 |
| | CBD M | 4 | 1.5128E-12 | 7.04387E-13 | 3.52193E-13 | 3.9201E-13 | 2.6337E-12 | 7.16E-13 | 2.43E-12 |
| | THC M | BDL | | | | | | | |
| | THC+CBD M | BAL | | | | | | | |
| *N*-arachidonoyl proline | Veh F | BAL | | | | | | | |
| | CBD F | BAL | | | | | | | |
| | THC F | BDL | | | | | | | |
| | THC+CBD F | BAL | | | | | | | |
| | Veh M | BDL | | | | | | | |
| | CBD M | BDL | | | | | | | |
| | THC M | BDL | | | | | | | |
| | THC+CBD M | BDL | | | | | | | |
| | Veh F | BDL | | | | | | | |

| | | |
|---|---|---|
| *N*-docosahexaenoyl proline | CBD F | BAL |
| | THC F | BDL |
| | THC+CBD F | BDL |
| | Veh M | BDL |
| | CBD M | BAL |
| | THC M | BAL |
| | THC+CBD M | BDL |

Supplemental Table 12.3 Independent t-test comparing male and female levels of *N*-acyl prolines measured in liver collected from adult mice 2h after an i.p. injection of 10mg/kg CBD, THC, or THC+CBD

| | | | Levene's Test for Equality of Variances | | t-test for Equality of Means | | | | | | | |
|---|---|---|---|---|---|---|---|---|---|---|---|---|
| | | | | | | | Significance | | | | 95% Confidence Interval of the Difference | |
| | | | F | Sig. | t | df | One-Sided p | Two-Sided p | Mean Difference | Std. Error Difference | Lower | Upper |
| Veh | *N*-palmitoyl proline | Equal variances assumed | 10.073 | 0.009 | 2.088 | 11 | 0.030 | 0.061 | 3.68092E-12 | 1.76286E-12 | -1.99095E-13 | 7.56094E-12 |
| | | Equal variances not assumed | | | 2.649 | 7.609 | 0.015 | 0.031 | 3.68092E-12 | 1.38960E-12 | 4.47585E-13 | 6.91426E-12 |
| CBD | *N*-palmitoyl proline | Equal variances assumed | 0.008 | 0.931 | 1.648 | 12 | 0.063 | 0.125 | 2.49269E-12 | 1.51210E-12 | -8.01900E-13 | 5.78729E-12 |
| | | Equal variances not assumed | | | 1.648 | 11.961 | 0.063 | 0.125 | 2.49269E-12 | 1.51210E-12 | -8.03086E-13 | 5.78847E-12 |
| THC | *N*-palmitoyl proline | Equal variances assumed | 2.909 | 0.114 | -1.944 | 12 | 0.038 | 0.076 | -2.23152E-12 | 1.14767E-12 | -4.73207E-12 | 2.69032E-13 |
| | | Equal variances not assumed | | | -1.944 | 9.026 | 0.042 | 0.084 | -2.23152E-12 | 1.14767E-12 | -4.82659E-12 | 3.63559E-13 |
| THC+CBD | *N*-palmitoyl proline | Equal variances assumed | 12.550 | 0.003 | -2.237 | 16 | 0.020 | 0.040 | -2.64331E-12 | 1.18144E-12 | -5.14785E-12 | -1.38760E-13 |
| | | Equal variances not assumed | | | -2.751 | 12.008 | 0.009 | 0.018 | -2.64331E-12 | 9.60847E-13 | -4.73665E-12 | -5.49965E-13 |

Supplemental Table 12.4 One-way ANOVA for *N*-acyl prolines measured in plasma collected from adult mice 2h after an i.p. injection of 10mg/kg CBD, THC, or THC+CBD

| | | Sum of Squares | df | Mean Square | F | Sig. |
|---|---|---|---|---|---|---|
| *N*-palmitoyl proline | Between Groups | 4.78E-18 | 7 | 6.82E-19 | 0.705 | 0.6680156 |
| | Within Groups | 2.13E-17 | 22 | 9.68E-19 | | |
| | Total | 2.61E-17 | 29 | | | |

Supplemental Table 12.5 One-way ANOVA for *N*-acyl prolines measured in liver collected from adult mice 2h after an i.p. injection of 10mg/kg CBD, THC, or THC+CBD

| | | Sum of Squares | df | Mean Square | F | Sig. |
|---|---|---|---|---|---|---|
| *N*-palmitoyl proline | Between Groups | 1.24E-22 | 7 | 1.77E-23 | 2.568 | 0.02400 |
| | Within Groups | 3.52E-22 | 51 | 6.90E-24 | | |
| | Total | 4.76E-22 | 58 | | | |
| *N*-oleoyl proline | Between Groups | 8.94E-24 | 3 | 2.98E-24 | 1.199 | 0.34614 |
| | Within Groups | 3.48E-23 | 14 | 2.49E-24 | | |
| | Total | 4.37E-23 | 17 | | | |
| *N*-linoleoyl proline | Between Groups | 9.46E-28 | 1 | 9.46E-28 | 0.001 | 0.97389 |
| | Within Groups | 5.76E-24 | 7 | 8.23E-25 | | |
| | Total | 5.76E-24 | 8 | | | |

#### Supplemental Table 12.6 Fisher's LSD post-hoc test for plasma *N*-acyl proline ANOVA

| Dependent Variable | | | Std. Error | Sig. | 95% Confidence Interval Lower Bound | 95% Confidence Interval Upper Bound |
|---|---|---|---|---|---|---|
| *N*-palmitoyl proline | TpCfemale | CBDmale | 6.02E-10 | 0.2430 | -5.27E-10 | 1.97E-09 |
| | | THCmale | 6.96E-10 | 0.2285 | -5.81E-10 | 2.30E-09 |
| | CBDmale | TpCfemale | 6.02E-10 | 0.2430 | -1.97E-09 | 5.27E-10 |
| | | THCmale | 6.02E-10 | 0.8196 | -1.11E-09 | 1.39E-09 |
| | THCmale | TpCfemale | 6.96E-10 | 0.2285 | -2.30E-09 | 5.81E-10 |
| | | CBDmale | 6.02E-10 | 0.8196 | -1.39E-09 | 1.11E-09 |

#### Supplemental Table 12.7 Fisher's LSD post-hoc test for liver *N*-acyl proline ANOVA

| Dependent Variable | | | Std. Error | Sig. | 95% Confidence Interval Lower Bound | 95% Confidence Interval Upper Bound |
|---|---|---|---|---|---|---|
| *N*-palmitoyl proline | VehFemale | CBDfemale | 1.35982E-12 | 0.610 | -3.4269E-12 | 2.0330E-12 |
| | | THCfemale | 1.35982E-12 | 0.041 | 1.2235E-13 | 5.5822E-12 |
| | | TpCfemale | 1.35982E-12 | 0.018 | 5.8962E-13 | 6.0495E-12 |
| | | VehMale | 1.49786E-12 | 0.017 | 6.7385E-13 | 6.6880E-12 |
| | | CBDmale | 1.35982E-12 | 0.193 | -9.3420E-13 | 4.5257E-12 |
| | | THCmale | 1.35982E-12 | 0.650 | -2.1092E-12 | 3.3507E-12 |
| | | TpCmale | 1.22085E-12 | 0.582 | -1.7747E-12 | 3.1272E-12 |
| | CBDfemale | VehFemale | 1.35982E-12 | 0.610 | -2.0330E-12 | 3.4269E-12 |
| | | THCfemale | 1.40441E-12 | 0.015 | 7.2978E-13 | 6.3687E-12 |
| | | TpCfemale | 1.40441E-12 | 0.006 | 1.1970E-12 | 6.8360E-12 |
| | | VehMale | 1.53846E-12 | 0.006 | 1.2893E-12 | 7.4665E-12 |
| | | CBDmale | 1.40441E-12 | 0.082 | -3.2678E-13 | 5.3122E-12 |
| | | THCmale | 1.40441E-12 | 0.353 | -1.5017E-12 | 4.1372E-12 |
| | | TpCmale | 1.27034E-12 | 0.285 | -1.1771E-12 | 3.9235E-12 |
| | THCfemale | VehFemale | 1.35982E-12 | 0.041 | -5.5822E-12 | -1.2235E-13 |
| | | CBDfemale | 1.40441E-12 | 0.015 | -6.3687E-12 | -7.2978E-13 |
| | | TpCfemale | 1.40441E-12 | 0.741 | -2.3522E-12 | 3.2867E-12 |
| | | VehMale | 1.53846E-12 | 0.593 | -2.2600E-12 | 3.9172E-12 |
| | | CBDmale | 1.40441E-12 | 0.455 | -3.8760E-12 | 1.7629E-12 |
| | | THCmale | 1.40441E-12 | 0.118 | -5.0510E-12 | 5.8796E-13 |
| | | TpCmale | 1.27034E-12 | 0.093 | -4.7264E-12 | 3.7427E-13 |
| | TpCfemale | VehFemale | 1.35982E-12 | 0.018 | -6.0495E-12 | -5.8962E-13 |
| | | CBDfemale | 1.40441E-12 | 0.006 | -6.8360E-12 | -1.1970E-12 |
| | | THCfemale | 1.40441E-12 | 0.741 | -3.2867E-12 | 2.3522E-12 |
| | | VehMale | 1.53846E-12 | 0.815 | -2.7272E-12 | 3.4499E-12 |
| | | CBDmale | 1.40441E-12 | 0.283 | -4.3433E-12 | 1.2957E-12 |
| | | THCmale | 1.40441E-12 | 0.060 | -5.5183E-12 | 1.2069E-13 |
| | | TpCmale | 1.27034E-12 | 0.042 | -5.1936E-12 | -9.2995E-14 |
| | VehMale | VehFemale | 1.49786E-12 | 0.017 | -6.6880E-12 | -6.7385E-13 |
| | | CBDfemale | 1.53846E-12 | 0.006 | -7.4665E-12 | -1.2893E-12 |
| | | THCfemale | 1.53846E-12 | 0.593 | -3.9172E-12 | 2.2600E-12 |
| | | TpCfemale | 1.53846E-12 | 0.815 | -3.4499E-12 | 2.7272E-12 |
| | | CBDmale | 1.53846E-12 | 0.226 | -4.9738E-12 | 1.2034E-12 |
| | | THCmale | 1.53846E-12 | 0.052 | -6.1487E-12 | 2.8438E-14 |
| | | TpCmale | 1.41712E-12 | 0.039 | -5.8497E-12 | -1.5968E-13 |
| | CBDmale | VehFemale | 1.35982E-12 | 0.193 | -4.5257E-12 | 9.3420E-13 |
| | | CBDfemale | 1.40441E-12 | 0.082 | -5.3122E-12 | 3.2678E-13 |
| | | THCfemale | 1.40441E-12 | 0.455 | -1.7629E-12 | 3.8760E-12 |
| | | TpCfemale | 1.40441E-12 | 0.283 | -1.2957E-12 | 4.3433E-12 |
| | | VehMale | 1.53846E-12 | 0.226 | -1.2034E-12 | 4.9738E-12 |
| | | THCmale | 1.40441E-12 | 0.407 | -3.9944E-12 | 1.6445E-12 |
| | | TpCmale | 1.27034E-12 | 0.382 | -3.6698E-12 | 1.4308E-12 |
| | THCmale | VehFemale | 1.35982E-12 | 0.650 | -3.3507E-12 | 2.1092E-12 |
| | | CBDfemale | 1.40441E-12 | 0.353 | -4.1372E-12 | 1.5017E-12 |
| | | THCfemale | 1.40441E-12 | 0.118 | -5.8796E-13 | 5.0510E-12 |
| | | TpCfemale | 1.40441E-12 | 0.060 | -1.2069E-13 | 5.5183E-12 |
| | | VehMale | 1.53846E-12 | 0.052 | -2.8438E-14 | 6.1487E-12 |
| | | CBDmale | 1.40441E-12 | 0.407 | -1.6445E-12 | 3.9944E-12 |
| | | TpCmale | 1.27034E-12 | 0.965 | -2.4948E-12 | 2.6058E-12 |
| | TpCmale | VehFemale | 1.22085E-12 | 0.582 | -3.1272E-12 | 1.7747E-12 |
| | | CBDfemale | 1.27034E-12 | 0.285 | -3.9235E-12 | 1.1771E-12 |
| | | THCfemale | 1.27034E-12 | 0.093 | -3.7427E-13 | 4.7264E-12 |
| | | TpCfemale | 1.27034E-12 | 0.042 | 9.2995E-14 | 5.1936E-12 |
| | | VehMale | 1.41712E-12 | 0.039 | 1.5968E-13 | 5.8497E-12 |
| | | CBDmale | 1.27034E-12 | 0.382 | -1.4308E-12 | 3.6698E-12 |
| | | THCmale | 1.27034E-12 | 0.965 | -2.6058E-12 | 2.4948E-12 |
| *N*-oleoyl proline | VehFemale | CBDfemale | 1.11475E-12 | 0.085 | -4.4607E-12 | 3.2110E-13 |

| | | | | | |
|---|---|---|---|---|---|
| | THCfemale | 1.20407E-12 | 0.543 | -3.3336E-12 | 1.8313E-12 |
| | TpCfemale | 9.88120E-13 | 0.465 | -2.8619E-12 | 1.3767E-12 |
| CBDfemale | F Veh | 1.11475E-12 | 0.085 | -3.2110E-13 | 4.4607E-12 |
| | THCfemale | 1.20407E-12 | 0.292 | -1.2638E-12 | 3.9011E-12 |
| | TpCfemale | 9.88120E-13 | 0.201 | -7.9214E-13 | 3.4465E-12 |
| THCfemale | F Veh | 1.20407E-12 | 0.543 | -1.8313E-12 | 3.3336E-12 |
| | CBDfemale | 1.20407E-12 | 0.292 | -3.9011E-12 | 1.2638E-12 |
| | TpCfemale | 1.08788E-12 | 0.994 | -2.3247E-12 | 2.3418E-12 |
| TpCfemale | F Veh | 9.88120E-13 | 0.465 | -1.3767E-12 | 2.8619E-12 |
| | CBDfemale | 9.88120E-13 | 0.201 | -3.4465E-12 | 7.9214E-13 |
| | THCfemale | 1.08788E-12 | 0.994 | -2.3418E-12 | 2.3247E-12 |

Supplemental Table 13.1 Descriptive statistics for *N*-acyl serines (mol/L) in plasma collected from male and female adult mice 2h after receiving an i.p. injection of 10mg/kg CBD, THC, or THC+CBD

| | | N | Mean | Std. Deviation | Std. Error | 95% Confidence Interval for Mean | | Minimum | Maximum |
|---|---|---|---|---|---|---|---|---|---|
| | | | | | | Lower Bound | Upper Bound | | |
| *N*-palmitoyl serine | Veh F | 7 | 2.9460E-09 | 1.56919E-09 | 5.93097E-10 | 1.4947E-09 | 4.3973E-09 | 3.98E-10 | 4.71E-09 |
| | CBD F | 6 | 2.5694E-09 | 1.21630E-09 | 4.96554E-10 | 1.2930E-09 | 3.8458E-09 | 9.85E-10 | 4.30E-09 |
| | THC F | 5 | 2.0059E-09 | 1.68796E-09 | 7.54877E-10 | -9.0000E-11 | 4.1018E-09 | 2.72E-10 | 3.91E-09 |
| | THC+CBD F | 9 | 1.5124E-09 | 9.30684E-10 | 3.10228E-10 | 7.9703E-10 | 2.2278E-09 | 4.75E-10 | 3.05E-09 |
| | Veh M | 5 | 2.5581E-09 | 1.55415E-09 | 6.95037E-10 | 6.2838E-10 | 4.4878E-09 | 4.02E-10 | 4.36E-09 |
| | CBD M | 4 | 3.9464E-09 | 1.55741E-09 | 7.78705E-10 | 1.4682E-09 | 6.4246E-09 | 1.89E-09 | 5.67E-09 |
| | THC M | 4 | 2.1715E-09 | 1.25129E-09 | 6.25644E-10 | 1.8038E-10 | 4.1625E-09 | 3.72E-10 | 3.19E-09 |
| | THC+CBD M | 4 | 3.0971E-09 | 5.39584E-10 | 2.69792E-10 | 2.2385E-09 | 3.9557E-09 | 2.52E-09 | 3.74E-09 |
| *N*-stearoyl serine | Veh F | 5 | 3.1379E-09 | 2.73239E-09 | 1.22196E-09 | -2.5478E-10 | 6.5306E-09 | 1.18E-09 | 7.89E-09 |
| | CBD F | BAL | | | | | | | |
| | THC F | 4 | 4.2209E-09 | 1.29214E-09 | 6.46070E-10 | 2.1649E-09 | 6.2770E-09 | 2.62E-09 | 5.61E-09 |
| | THC+CBD F | 4 | 3.5948E-09 | 2.72635E-09 | 1.36317E-09 | -7.4340E-10 | 7.9331E-09 | 5.86E-10 | 7.06E-09 |
| | Veh M | BAL | | | | | | | |
| | CBD M | BAL | | | | | | | |
| | THC M | BAL | | | | | | | |
| | THC+CBD M | 4 | 1.4786E-09 | 2.51699E-10 | 1.25850E-10 | 1.0781E-09 | 1.8791E-09 | 1.25E-09 | 1.84E-09 |
| *N*-oleoyl serine | Veh F | BDL | | | | | | | |
| | CBD F | BAL | | | | | | | |
| | THC F | BAL | | | | | | | |
| | THC+CBD F | BDL | | | | | | | |
| | Veh M | BAL | | | | | | | |
| | CBD M | BAL | | | | | | | |
| | THC M | BAL | | | | | | | |
| | THC+CBD M | BAL | | | | | | | |
| *N*-linoleoyl serine | Veh F | BDL | | | | | | | |
| | CBD F | BAL | | | | | | | |
| | THC F | BAL | | | | | | | |
| | THC+CBD F | BDL | | | | | | | |
| | Veh M | BDL | | | | | | | |
| | CBD M | BDL | | | | | | | |
| | THC M | BAL | | | | | | | |
| | THC+CBD M | BDL | | | | | | | |
| *N*-arachidonoyl serine | Veh F | BDL | | | | | | | |
| | CBD F | BDL | | | | | | | |
| | THC F | BDL | | | | | | | |
| | THC+CBD F | BDL | | | | | | | |
| | Veh M | BDL | | | | | | | |
| | CBD M | BAL | | | | | | | |
| | THC M | BAL | | | | | | | |
| | THC+CBD M | BDL | | | | | | | |
| | Veh F | BDL | | | | | | | |

| | | |
|---|---|---|
| *N*-docosahexaenoyl serine | CBD F | BDL |
| | THC F | BDL |
| | THC+CBD F | BDL |
| | Veh M | BDL |
| | CBD M | BDL |
| | THC M | BDL |
| | THC+CBD M | BDL |

Supplemental Table 13.2 Descriptive statistics for *N*-acyl serines (mol/g) in liver collected from male and female adult mice 2h after receiving an i.p. injection of 10mg/kg CBD, THC, or THC+CBD

| | | N | Mean | Std. Deviation | Std. Error | 95% Confidence Interval for Mean | | Minimum | Maximum |
|---|---|---|---|---|---|---|---|---|---|
| | | | | | | Lower Bound | Upper Bound | | |
| *N*-palmitoyl serine | Veh F | 7 | 2.9460E-09 | 1.56919E-09 | 5.93097E-10 | 1.4947E-09 | 4.3973E-09 | 3.98E-10 | 4.71E-09 |
| | CBD F | 6 | 2.5694E-09 | 1.21630E-09 | 4.96554E-10 | 1.2930E-09 | 3.8458E-09 | 9.85E-10 | 4.30E-09 |
| | THC F | 5 | 2.0059E-09 | 1.68796E-09 | 7.54877E-10 | -9.0000E-11 | 4.1018E-09 | 2.72E-10 | 3.91E-09 |
| | THC+CBD F | 9 | 1.5124E-09 | 9.30684E-10 | 3.10228E-10 | 7.9703E-10 | 2.2278E-09 | 4.75E-10 | 3.05E-09 |
| | Veh M | 5 | 2.5581E-09 | 1.55415E-09 | 6.95037E-10 | 6.2838E-10 | 4.4878E-09 | 4.02E-10 | 4.36E-09 |
| | CBD M | 4 | 3.9464E-09 | 1.55741E-09 | 7.78705E-10 | 1.4682E-09 | 6.4246E-09 | 1.89E-09 | 5.67E-09 |
| | THC M | 4 | 2.1715E-09 | 1.25129E-09 | 6.25644E-10 | 1.8038E-10 | 4.1625E-09 | 3.72E-10 | 3.19E-09 |
| | THC+CBD M | 4 | 3.0971E-09 | 5.39584E-10 | 2.69792E-10 | 2.2385E-09 | 3.9557E-09 | 2.52E-09 | 3.74E-09 |
| *N*-stearoyl serine | Veh F | 5 | 3.1379E-09 | 2.73239E-09 | 1.22196E-09 | -2.5478E-10 | 6.5306E-09 | 1.18E-09 | 7.89E-09 |
| | CBD F | BAL | | | | | | | |
| | THC F | 4 | 4.2209E-09 | 1.29214E-09 | 6.46070E-10 | 2.1649E-09 | 6.2770E-09 | 2.62E-09 | 5.61E-09 |
| | THC+CBD F | 4 | 3.5948E-09 | 2.72635E-09 | 1.36317E-09 | -7.4340E-10 | 7.9331E-09 | 5.86E-10 | 7.06E-09 |
| | Veh M | BAL | | | | | | | |
| | CBD M | BAL | | | | | | | |
| | THC M | BAL | | | | | | | |
| | THC+CBD M | 4 | 1.4786E-09 | 2.51699E-10 | 1.25850E-10 | 1.0781E-09 | 1.8791E-09 | 1.25E-09 | 1.84E-09 |
| *N*-oleoyl serine | Veh F | BDL | | | | | | | |
| | CBD F | BAL | | | | | | | |
| | THC F | BAL | | | | | | | |
| | THC+CBD F | BDL | | | | | | | |
| | Veh M | BAL | | | | | | | |
| | CBD M | BAL | | | | | | | |
| | THC M | BAL | | | | | | | |
| | THC+CBD M | BAL | | | | | | | |
| *N*-linoleoyl serine | Veh F | BDL | | | | | | | |
| | CBD F | BAL | | | | | | | |
| | THC F | BAL | | | | | | | |
| | THC+CBD F | BDL | | | | | | | |
| | Veh M | BDL | | | | | | | |
| | CBD M | BDL | | | | | | | |
| | THC M | BAL | | | | | | | |
| | THC+CBD M | BDL | | | | | | | |
| *N*-arachidonoyl serine | Veh F | BDL | | | | | | | |
| | CBD F | BDL | | | | | | | |
| | THC F | BDL | | | | | | | |
| | THC+CBD F | BDL | | | | | | | |
| | Veh M | BDL | | | | | | | |
| | CBD M | BAL | | | | | | | |
| | THC M | BAL | | | | | | | |
| | THC+CBD M | BDL | | | | | | | |
| | Veh F | BDL | | | | | | | |

| | | |
|---|---|---|
| *N*-docosahexaenoyl serine | CBD F | BDL |
| | THC F | BDL |
| | THC+CBD F | BDL |
| | Veh M | BDL |
| | CBD M | BDL |
| | THC M | BDL |
| | THC+CBD M | BDL |

Supplemental Table 13.3 Independent t-test comparing male and female levels of *N*-acyl serines measured in plasma collected from adult mice 2h after an i.p. injection of 10mg/kg CBD, THC, or THC+CBD

| | | | Levene's Test for Equality of Variances | | t-test for Equality of Means | | | | | | |
|---|---|---|---|---|---|---|---|---|---|---|---|
| | | | | | | | Significance | | | 95% Confidence Interval of the Difference | |
| | | | F | Sig. | t | df | One-Sided p | Two-Sided p | Std. Error Difference | Lower | Upper |
| Veh | *N*-palmitoyl serine | Equal variances assumed | 0.001 | 0.970 | 0.424 | 10 | 0.340 | 0.681 | 9.15310E-10 | -1.65155E-09 | 2.42732E-09 |
| | | Equal variances not assumed | | | 0.425 | 8.826 | 0.341 | 0.681 | 9.13696E-10 | -1.68525E-09 | 2.46103E-09 |
| CBD | *N*-palmitoyl serine | Equal variances assumed | 0.016 | 0.902 | -1.575 | 8 | 0.077 | 0.154 | 8.74213E-10 | -3.39291E-09 | 6.38965E-10 |
| | | Equal variances not assumed | | | -1.491 | 5.400 | 0.096 | 0.192 | 9.23552E-10 | -3.69911E-09 | 9.45158E-10 |
| THC | *N*-palmitoyl serine | Equal variances assumed | 0.898 | 0.375 | -0.163 | 7 | 0.438 | 0.875 | 1.01716E-09 | -2.57078E-09 | 2.23962E-09 |
| | | Equal variances not assumed | | | -0.169 | 6.987 | 0.435 | 0.871 | 9.80444E-10 | -2.48484E-09 | 2.15368E-09 |
| THC+CBD | *N*-palmitoyl serine | Equal variances assumed | 1.466 | 0.251 | -3.131 | 11 | 0.005 | 0.010 | 5.06116E-10 | -2.69860E-09 | -4.70692E-10 |
| | | Equal variances not assumed | | | -3.854 | 9.772 | 0.002 | 0.003 | 4.11131E-10 | -2.50361E-09 | -6.65680E-10 |
| | *N*-stearoyl serine | Equal variances assumed | 6.087 | 0.049 | 1.546 | 6 | 0.087 | 0.173 | 1.36897E-09 | -1.23353E-09 | 5.46597E-09 |
| | | Equal variances not assumed | | | 1.546 | 3.051 | 0.109 | 0.218 | 1.36897E-09 | -2.19943E-09 | 6.43188E-09 |

Supplemental Table 13.4 Independent t-test comparing male and female levels of *N*-acyl serines measured in liver collected from adult mice 2h after an i.p. injection of 10mg/kg CBD, THC, or THC+CBD

| | | | Levene's Test for Equality of Variances | | t-test for Equality of Means | | | | | | | |
|---|---|---|---|---|---|---|---|---|---|---|---|---|
| | | | | | | | Significance | | | | 95% Confidence Interval of the Difference | |
| | | | F | Sig. | t | df | One-Sided p | Two-Sided p | Mean Difference | Std. Error Difference | Lower | Upper |
| Veh | *N*-palmitoyl serine | Equal variances assumed | 0.357 | 0.562 | 0.543 | 11 | 0.299 | 0.598 | 9.83000E-12 | 1.81113E-11 | -3.00327E-11 | 4.96927E-11 |
| | | Equal variances not assumed | | | 0.563 | 10.240 | 0.293 | 0.586 | 9.83000E-12 | 1.74751E-11 | -2.89836E-11 | 4.86436E-11 |
| | *N*-stearoyl serine | Equal variances assumed | 8.170 | 0.016 | 0.519 | 11 | 0.307 | 0.614 | 4.37470E-12 | 8.42952E-12 | -1.41785E-11 | 2.29279E-11 |
| | | Equal variances not assumed | | | 0.560 | 6.629 | 0.297 | 0.594 | 4.37470E-12 | 7.81023E-12 | -1.43054E-11 | 2.30548E-11 |
| | *N*-oleoyl serine | Equal variances assumed | 6.783 | 0.023 | 2.258 | 12 | 0.022 | 0.043 | 2.23932E-11 | 9.91715E-12 | 7.85608E-13 | 4.40009E-11 |
| | | Equal variances not assumed | | | 2.586 | 8.156 | 0.016 | 0.032 | 2.23932E-11 | 8.65908E-12 | 2.49164E-12 | 4.22948E-11 |
| | *N*-linoleoyl serine | Equal variances assumed | 0.534 | 0.479 | 1.456 | 12 | 0.086 | 0.171 | 2.95718E-11 | 2.03094E-11 | -1.46785E-11 | 7.38221E-11 |
| | | Equal variances not assumed | | | 1.456 | 11.949 | 0.086 | 0.171 | 2.95718E-11 | 2.03094E-11 | -1.46996E-11 | 7.38432E-11 |
| | *N*-arachidonoyl serine | Equal variances assumed | 0.472 | 0.505 | 0.919 | 12 | 0.188 | 0.376 | 1.45346E-12 | 1.58183E-12 | -1.99306E-12 | 4.89997E-12 |
| | | Equal variances not assumed | | | 0.919 | 9.616 | 0.190 | 0.381 | 1.45346E-12 | 1.58183E-12 | -2.09024E-12 | 4.99715E-12 |
| | *N*-docosahexaenoyl serine | Equal variances assumed | 2.793 | 0.123 | 0.526 | 11 | 0.305 | 0.609 | 4.87444E-12 | 9.26980E-12 | -1.55282E-11 | 2.52771E-11 |
| | | Equal variances not assumed | | | 0.582 | 10.930 | 0.286 | 0.572 | 4.87444E-12 | 8.36945E-12 | -1.35611E-11 | 2.33100E-11 |
| CBD | *N*-palmitoyl serine | Equal variances assumed | 1.148 | 0.301 | 1.079 | 15 | 0.149 | 0.298 | 2.40378E-11 | 2.22772E-11 | -2.34449E-11 | 7.15204E-11 |
| | | Equal variances not assumed | | | 1.189 | 14.543 | 0.127 | 0.254 | 2.40378E-11 | 2.02205E-11 | -1.91795E-11 | 6.72551E-11 |
| | *N*-stearoyl serine | Equal variances assumed | 1.104 | 0.312 | 0.540 | 13 | 0.299 | 0.598 | 7.99680E-12 | 1.48005E-11 | -2.39777E-11 | 3.99713E-11 |

| | | | | | | | | | | | | |
|---|---|---|---|---|---|---|---|---|---|---|---|---|
| | | Equal variances not assumed | | | 0.516 | 8.367 | 0.309 | 0.619 | 7.99680E-12 | 1.54837E-11 | -2.74380E-11 | 4.34316E-11 |
| | *N*-oleoyl serine | Equal variances assumed | 16.191 | 0.001 | 1.430 | 15 | 0.087 | 0.173 | 1.28286E-11 | 8.97251E-12 | -6.29584E-12 | 3.19531E-11 |
| | | Equal variances not assumed | | | 1.223 | 6.852 | 0.131 | 0.262 | 1.28286E-11 | 1.04936E-11 | -1.20941E-11 | 3.77513E-11 |
| | *N*-linoleoyl serine | Equal variances assumed | 18.331 | 0.001 | 1.561 | 15 | 0.070 | 0.139 | 5.91740E-11 | 3.79144E-11 | -2.16388E-11 | 1.39987E-10 |
| | | Equal variances not assumed | | | 1.312 | 6.433 | 0.117 | 0.234 | 5.91740E-11 | 4.50943E-11 | -4.93936E-11 | 1.67741E-10 |
| | *N*-arachidonoyl serine | Equal variances assumed | 8.564 | 0.012 | 1.942 | 13 | 0.037 | 0.074 | 4.44225E-12 | 2.28735E-12 | -4.99274E-13 | 9.38377E-12 |
| | | Equal variances not assumed | | | 1.661 | 6.137 | 0.073 | 0.147 | 4.44225E-12 | 2.67424E-12 | -2.06612E-12 | 1.09506E-11 |
| | *N*-docosahexaenoyl serine | Equal variances assumed | 19.183 | 0.001 | 2.610 | 13 | 0.011 | 0.022 | 4.60096E-11 | 1.76269E-11 | 7.92908E-12 | 8.40902E-11 |
| | | Equal variances not assumed | | | 2.117 | 5.191 | 0.043 | 0.086 | 4.60096E-11 | 2.17340E-11 | -9.24743E-12 | 1.01267E-10 |
| | *N*-palmitoyl serine | Equal variances assumed | 1.219 | 0.285 | -0.653 | 17 | 0.261 | 0.522 | -1.78195E-11 | 2.72869E-11 | -7.53898E-11 | 3.97509E-11 |
| | | Equal variances not assumed | | | -0.643 | 14.716 | 0.265 | 0.530 | -1.78195E-11 | 2.77271E-11 | -7.70180E-11 | 4.13791E-11 |
| | *N*-stearoyl serine | Equal variances assumed | 3.372 | 0.086 | 1.233 | 15 | 0.118 | 0.237 | 1.62861E-11 | 1.32097E-11 | -1.18697E-11 | 4.44418E-11 |
| | | Equal variances not assumed | | | 1.164 | 7.528 | 0.140 | 0.280 | 1.62861E-11 | 1.39973E-11 | -1.63475E-11 | 4.89196E-11 |
| THC | *N*-oleoyl serine | Equal variances assumed | 1.017 | 0.328 | 2.047 | 16 | 0.029 | 0.057 | 3.52080E-11 | 1.72003E-11 | -1.25502E-12 | 7.16709E-11 |
| | | Equal variances not assumed | | | 2.117 | 15.948 | 0.025 | 0.050 | 3.52080E-11 | 1.66284E-11 | -5.19582E-14 | 7.04679E-11 |
| | *N*-linoleoyl serine | Equal variances assumed | 0.071 | 0.794 | -0.366 | 16 | 0.360 | 0.719 | -1.44560E-11 | 3.95054E-11 | -9.82038E-11 | 6.92918E-11 |
| | | Equal variances not assumed | | | -0.366 | 15.939 | 0.360 | 0.719 | -1.44560E-11 | 3.95054E-11 | -9.82297E-11 | 6.93177E-11 |
| | *N*-arachidonoyl serine | Equal variances assumed | 0.000 | 0.985 | 0.412 | 16 | 0.343 | 0.685 | 1.20664E-12 | 2.92551E-12 | -4.99517E-12 | 7.40844E-12 |
| | | Equal variances | | | 0.412 | 15.828 | 0.343 | 0.686 | 1.20664E-12 | 2.92551E-12 | -5.00064E-12 | 7.41391E-12 |

| | | | | | | | | | | | |
|---|---|---|---|---|---|---|---|---|---|---|---|
| | | not assumed | | | | | | | | | |
| | *N*-docosahexaenoyl serine | Equal variances assumed | 3.729 | 0.071 | -1.089 | 16 | 0.146 | 0.292 | -1.51803E-11 | 1.39404E-11 | -4.47327E-11 | 1.43720E-11 |
| | | Equal variances not assumed | | | -1.089 | 13.603 | 0.148 | 0.295 | -1.51803E-11 | 1.39404E-11 | -4.51617E-11 | 1.48010E-11 |
| | *N*-palmitoyl serine | Equal variances assumed | 0.193 | 0.665 | 2.350 | 19 | 0.015 | 0.030 | 6.86257E-11 | 2.92065E-11 | 7.49585E-12 | 1.29756E-10 |
| | | Equal variances not assumed | | | 2.316 | 16.524 | 0.017 | 0.034 | 6.86257E-11 | 2.96359E-11 | 5.96200E-12 | 1.31289E-10 |
| | *N*-stearoyl serine | Equal variances assumed | 5.943 | 0.026 | 1.485 | 17 | 0.078 | 0.156 | 7.89333E-12 | 5.31572E-12 | -3.32186E-12 | 1.91085E-11 |
| | | Equal variances not assumed | | | 1.443 | 12.329 | 0.087 | 0.174 | 7.89333E-12 | 5.47175E-12 | -3.99338E-12 | 1.97800E-11 |
| | *N*-oleoyl serine | Equal variances assumed | 1.904 | 0.186 | 3.990 | 17 | 0.000 | 0.001 | 4.93012E-11 | 1.23570E-11 | 2.32303E-11 | 7.53721E-11 |
| | | Equal variances not assumed | | | 4.084 | 15.469 | 0.000 | 0.001 | 4.93012E-11 | 1.20732E-11 | 2.36355E-11 | 7.49668E-11 |
| THC+CBD | *N*-linoleoyl serine | Equal variances assumed | 0.790 | 0.387 | 2.423 | 17 | 0.013 | 0.027 | 1.07140E-10 | 4.42181E-11 | 1.38483E-11 | 2.00432E-10 |
| | | Equal variances not assumed | | | 2.464 | 16.397 | 0.013 | 0.025 | 1.07140E-10 | 4.34882E-11 | 1.51306E-11 | 1.99150E-10 |
| | *N*-arachidonoyl serine | Equal variances assumed | 3.813 | 0.068 | 2.597 | 17 | 0.009 | 0.019 | 8.40892E-12 | 3.23757E-12 | 1.57825E-12 | 1.52396E-11 |
| | | Equal variances not assumed | | | 2.502 | 10.824 | 0.015 | 0.030 | 8.40892E-12 | 3.36049E-12 | 9.97831E-13 | 1.58200E-11 |
| | *N*-docosahexaenoyl serine | Equal variances assumed | 3.927 | 0.063 | 1.956 | 18 | 0.033 | 0.066 | 4.16917E-11 | 2.13199E-11 | -3.09980E-12 | 8.64833E-11 |
| | | Equal variances not assumed | | | 1.956 | 12.498 | 0.037 | 0.073 | 4.16917E-11 | 2.13199E-11 | -4.55599E-12 | 8.79395E-11 |

Supplemental Table 13.5 One-way ANOVA for *N*-acyl serines measured in plasma collected from adult mice 2h after an i.p. injection of 10mg/kg CBD, THC, or THC+CBD

| | | Sum of Squares | df | Mean Square | F | Sig. |
|---|---|---|---|---|---|---|
| *N*-palmitoyl serine | Between Groups | 2.17E-17 | 7 | 3.09E-18 | 1.767 | 0.1244593 |
| | Within Groups | 6.30E-17 | 36 | 1.75E-18 | | |
| | Total | 8.47E-17 | 43 | | | |
| *N*-stearoyl serine | Between Groups | 0.000 | 6 | 0.000 | 0.929 | 0.4992819 |
| | Within Groups | 0.000 | 17 | 0.000 | | |
| | Total | 0.000 | 23 | | | |

Supplemental Table 13.6 One-way ANOVA for *N*-acyl serines measured in liver collected from adult mice 2h after an i.p. injection of 10mg/kg CBD, THC, or THC+CBD

| | | Sum of Squares | df | Mean Square | F | Sig. |
|---|---|---|---|---|---|---|
| *N*-palmitoyl serine | Between Groups | 6.89E-20 | 7 | 9.84E-21 | 3.261 | 0.00518 |
| | Within Groups | 1.87E-19 | 62 | 3.02E-21 | | |
| | Total | 2.56E-19 | 69 | | | |
| *N*-stearoyl serine | Between Groups | 2.14E-21 | 7 | 3.06E-22 | 0.646 | 0.71622 |
| | Within Groups | 2.65E-20 | 56 | 4.74E-22 | | |
| | Total | 2.87E-20 | 63 | | | |
| *N*-oleoyl serine | Between Groups | 4.30E-20 | 7 | 6.14E-21 | 8.694 | 0.00000 |
| | Within Groups | 4.24E-20 | 60 | 7.06E-22 | | |
| | Total | 8.53E-20 | 67 | | | |
| *N*-linoleoyl serine | Between Groups | 1.84E-19 | 7 | 2.62E-20 | 4.188 | 0.00082 |
| | Within Groups | 3.76E-19 | 60 | 6.27E-21 | | |
| | Total | 5.60E-19 | 67 | | | |
| *N*-arachidonoyl serine | Between Groups | 9.11E-22 | 7 | 1.30E-22 | 4.171 | 0.00089 |
| | Within Groups | 1.81E-21 | 58 | 3.12E-23 | | |
| | Total | 2.72E-21 | 65 | | | |
| *N*-docosahexaenoyl serine | Between Groups | 2.40E-20 | 7 | 3.43E-21 | 2.753 | 0.01539 |
| | Within Groups | 7.24E-20 | 58 | 1.25E-21 | | |
| | Total | 9.64E-20 | 65 | | | |

#### Supplemental Table 13.7 Fisher's LSD post-hoc test for plasma *N*-acyl serine ANOVA

| Dependent Variable | | | Std. Error | Sig. | 95% Confidence Interval Lower Bound | 95% Confidence Interval Upper Bound |
|---|---|---|---|---|---|---|
| *N*-palmitoyl serine | VehFemale | CBDfemale | 7.36E-10 | 6.12E-01 | -1.12E-09 | 1.87E-09 |
| | | THCfemale | 7.75E-10 | 2.33E-01 | -6.31E-10 | 2.51E-09 |
| | | TpCfemale | 6.67E-10 | 3.83E-02 | 8.15E-11 | 2.79E-09 |
| | | VehMale | 7.75E-10 | 6.20E-01 | -1.18E-09 | 1.96E-09 |
| | | CBDmale | 8.29E-10 | 2.36E-01 | -2.68E-09 | 6.81E-10 |
| | | THCmale | 8.29E-10 | 3.56E-01 | -9.07E-10 | 2.46E-09 |
| | | TpCmale | 8.29E-10 | 8.56E-01 | -1.83E-09 | 1.53E-09 |
| | CBDfemale | VehFemale | 7.36E-10 | 6.12E-01 | -1.87E-09 | 1.12E-09 |
| | | THCfemale | 8.01E-10 | 4.86E-01 | -1.06E-09 | 2.19E-09 |
| | | TpCfemale | 6.97E-10 | 1.38E-01 | -3.57E-10 | 2.47E-09 |
| | | VehMale | 8.01E-10 | 9.89E-01 | -1.61E-09 | 1.64E-09 |
| | | CBDmale | 8.54E-10 | 1.16E-01 | -3.11E-09 | 3.55E-10 |
| | | THCmale | 8.54E-10 | 6.44E-01 | -1.33E-09 | 2.13E-09 |
| | | TpCmale | 8.54E-10 | 5.41E-01 | -2.26E-09 | 1.20E-09 |
| | THCfemale | VehFemale | 7.75E-10 | 2.33E-01 | -2.51E-09 | 6.31E-10 |
| | | CBDfemale | 8.01E-10 | 4.86E-01 | -2.19E-09 | 1.06E-09 |
| | | TpCfemale | 7.38E-10 | 5.08E-01 | -1.00E-09 | 1.99E-09 |
| | | VehMale | 8.37E-10 | 5.13E-01 | -2.25E-09 | 1.14E-09 |
| | | CBDmale | 8.87E-10 | 3.54E-02 | -3.74E-09 | -1.41E-10 |
| | | THCmale | 8.87E-10 | 8.53E-01 | -1.97E-09 | 1.63E-09 |
| | | TpCmale | 8.87E-10 | 2.27E-01 | -2.89E-09 | 7.09E-10 |
| | TpCfemale | VehFemale | 6.67E-10 | 3.83E-02 | -2.79E-09 | -8.15E-11 |
| | | CBDfemale | 6.97E-10 | 1.38E-01 | -2.47E-09 | 3.57E-10 |
| | | THCfemale | 7.38E-10 | 5.08E-01 | -1.99E-09 | 1.00E-09 |
| | | VehMale | 7.38E-10 | 1.65E-01 | -2.54E-09 | 4.51E-10 |
| | | CBDmale | 7.95E-10 | 4.15E-03 | -4.05E-09 | -8.22E-10 |
| | | THCmale | 7.95E-10 | 4.13E-01 | -2.27E-09 | 9.53E-10 |
| | | TpCmale | 7.95E-10 | 5.38E-02 | -3.20E-09 | 2.77E-11 |
| | VehMale | VehFemale | 7.75E-10 | 6.20E-01 | -1.96E-09 | 1.18E-09 |
| | | CBDfemale | 8.01E-10 | 9.89E-01 | -1.64E-09 | 1.61E-09 |
| | | THCfemale | 8.37E-10 | 5.13E-01 | -1.14E-09 | 2.25E-09 |
| | | TpCfemale | 7.38E-10 | 1.65E-01 | -4.51E-10 | 2.54E-09 |
| | | CBDmale | 8.87E-10 | 1.26E-01 | -3.19E-09 | 4.12E-10 |
| | | THCmale | 8.87E-10 | 6.66E-01 | -1.41E-09 | 2.19E-09 |
| | | TpCmale | 8.87E-10 | 5.47E-01 | -2.34E-09 | 1.26E-09 |
| | CBDmale | VehFemale | 8.29E-10 | 2.36E-01 | -6.81E-10 | 2.68E-09 |
| | | CBDfemale | 8.54E-10 | 1.16E-01 | -3.55E-10 | 3.11E-09 |
| | | THCfemale | 8.87E-10 | 3.54E-02 | 1.41E-10 | 3.74E-09 |
| | | TpCfemale | 7.95E-10 | 4.15E-03 | 8.22E-10 | 4.05E-09 |
| | | VehMale | 8.87E-10 | 1.26E-01 | -4.12E-10 | 3.19E-09 |
| | | THCmale | 9.35E-10 | 6.58E-02 | -1.22E-10 | 3.67E-09 |
| | | TpCmale | 9.35E-10 | 3.70E-01 | -1.05E-09 | 2.75E-09 |
| | THCmale | VehFemale | 8.29E-10 | 3.56E-01 | -2.46E-09 | 9.07E-10 |
| | | CBDfemale | 8.54E-10 | 6.44E-01 | -2.13E-09 | 1.33E-09 |
| | | THCfemale | 8.87E-10 | 8.53E-01 | -1.63E-09 | 1.97E-09 |
| | | TpCfemale | 7.95E-10 | 4.13E-01 | -9.53E-10 | 2.27E-09 |
| | | VehMale | 8.87E-10 | 6.66E-01 | -2.19E-09 | 1.41E-09 |
| | | CBDmale | 9.35E-10 | 6.58E-02 | -3.67E-09 | 1.22E-10 |
| | | TpCmale | 9.35E-10 | 3.29E-01 | -2.82E-09 | 9.72E-10 |
| | TpCmale | VehFemale | 8.29E-10 | 8.56E-01 | -1.53E-09 | 1.83E-09 |
| | | CBDfemale | 8.54E-10 | 5.41E-01 | -1.20E-09 | 2.26E-09 |
| | | THCfemale | 8.87E-10 | 2.27E-01 | -7.09E-10 | 2.89E-09 |
| | | TpCfemale | 7.95E-10 | 5.38E-02 | -2.77E-11 | 3.20E-09 |
| | | VehMale | 8.87E-10 | 5.47E-01 | -1.26E-09 | 2.34E-09 |
| | | CBDmale | 9.35E-10 | 3.70E-01 | -2.75E-09 | 1.05E-09 |
| | | THCmale | 9.35E-10 | 3.29E-01 | -9.72E-10 | 2.82E-09 |
| *N*-stearoyl serine | VehFemale | THCfemale | 1.31E-09 | 4.19E-01 | -3.8408E-09 | 1.6747E-09 |

| | | | | | |
|---|---|---|---|---|---|
| | TpCfemale | 1.31E-09 | 7.31E-01 | -3.2147E-09 | 2.3009E-09 |
| | TpCmale | 1.31E-09 | 2.21E-01 | -1.0984E-09 | 4.4171E-09 |
| THCfemale | VehFemale | 1.31E-09 | 4.19E-01 | -1.6747E-09 | 3.8408E-09 |
| | TpCfemale | 1.38E-09 | 6.55E-01 | -2.2808E-09 | 3.5331E-09 |
| | TpCmale | 1.38E-09 | 6.29E-02 | -1.6460E-10 | 5.6493E-09 |
| TpCfemale | VehFemale | 1.31E-09 | 7.31E-01 | -2.3009E-09 | 3.2147E-09 |
| | THCfemale | 1.38E-09 | 6.55E-01 | -3.5331E-09 | 2.2808E-09 |
| | TpCmale | 1.38E-09 | 1.43E-01 | -7.9072E-10 | 5.0232E-09 |
| TpCmale | VehFemale | 1.31E-09 | 2.21E-01 | -4.4171E-09 | 1.0984E-09 |
| | THCfemale | 1.38E-09 | 6.29E-02 | -5.6493E-09 | 1.6460E-10 |
| | TpCfemale | 1.38E-09 | 1.43E-01 | -5.0232E-09 | 7.9072E-10 |

#### Supplemental Table 13.8 Fisher's LSD post-hoc test for liver *N*-acyl serine ANOVA

| Dependent Variable | | | Std. Error | Sig. | 95% Confidence Interval Lower Bound | Upper Bound |
|---|---|---|---|---|---|---|
| *N*-palmitoyl serine | VehFemale | CBDfemale | 2.93684E-11 | 0.128 | -1.0406E-10 | 1.3355E-11 |
| | | THCfemale | 2.70763E-11 | 0.050 | -1.0836E-10 | -1.0798E-13 |
| | | TpCfemale | 2.70763E-11 | 0.002 | -1.4241E-10 | -3.4161E-11 |
| | | VehMale | 3.05676E-11 | 0.749 | -5.1274E-11 | 7.0934E-11 |
| | | CBDmale | 2.70763E-11 | 0.434 | -7.5439E-11 | 3.2811E-11 |
| | | THCmale | 2.76888E-11 | 0.012 | -1.2740E-10 | -1.6703E-11 |
| | | TpCmale | 2.65647E-11 | 0.462 | -7.2762E-11 | 3.3442E-11 |
| | CBDfemale | VehFemale | 2.93684E-11 | 0.128 | -1.3355E-11 | 1.0406E-10 |
| | | THCfemale | 2.70763E-11 | 0.744 | -6.3006E-11 | 4.5244E-11 |
| | | TpCfemale | 2.70763E-11 | 0.118 | -9.7058E-11 | 1.1191E-11 |
| | | VehMale | 3.05676E-11 | 0.076 | -5.9217E-12 | 1.1629E-10 |
| | | CBDmale | 2.70763E-11 | 0.378 | -3.0087E-11 | 7.8163E-11 |
| | | THCmale | 2.76888E-11 | 0.339 | -8.2049E-11 | 2.8649E-11 |
| | | TpCmale | 2.65647E-11 | 0.337 | -2.7410E-11 | 7.8794E-11 |
| | THCfemale | VehFemale | 2.70763E-11 | 0.050 | 1.0798E-13 | 1.0836E-10 |
| | | CBDfemale | 2.70763E-11 | 0.744 | -4.5244E-11 | 6.3006E-11 |
| | | TpCfemale | 2.45714E-11 | 0.171 | -8.3170E-11 | 1.5065E-11 |
| | | VehMale | 2.83726E-11 | 0.027 | 7.3468E-12 | 1.2078E-10 |
| | | CBDmale | 2.45714E-11 | 0.185 | -1.6199E-11 | 8.2036E-11 |
| | | THCmale | 2.52447E-11 | 0.483 | -6.8283E-11 | 3.2644E-11 |
| | | TpCmale | 2.40064E-11 | 0.155 | -1.3415E-11 | 8.2561E-11 |
| | TpCfemale | VehFemale | 2.70763E-11 | 0.002 | 3.4161E-11 | 1.4241E-10 |
| | | CBDfemale | 2.70763E-11 | 0.118 | -1.1191E-11 | 9.7058E-11 |
| | | THCfemale | 2.45714E-11 | 0.171 | -1.5065E-11 | 8.3170E-11 |
| | | VehMale | 2.83726E-11 | 0.001 | 4.1400E-11 | 1.5483E-10 |
| | | CBDmale | 2.45714E-11 | 0.008 | 1.7854E-11 | 1.1609E-10 |
| | | THCmale | 2.52447E-11 | 0.523 | -3.4230E-11 | 6.6697E-11 |
| | | TpCmale | 2.40064E-11 | 0.006 | 2.0638E-11 | 1.1661E-10 |
| | VehMale | VehFemale | 3.05676E-11 | 0.749 | -7.0934E-11 | 5.1274E-11 |
| | | CBDfemale | 3.05676E-11 | 0.076 | -1.1629E-10 | 5.9217E-12 |
| | | THCfemale | 2.83726E-11 | 0.027 | -1.2078E-10 | -7.3468E-12 |
| | | TpCfemale | 2.83726E-11 | 0.001 | -1.5483E-10 | -4.1400E-11 |
| | | CBDmale | 2.83726E-11 | 0.277 | -8.7860E-11 | 2.5572E-11 |
| | | THCmale | 2.89576E-11 | 0.006 | -1.3977E-10 | -2.3997E-11 |
| | | TpCmale | 2.78847E-11 | 0.294 | -8.5231E-11 | 2.6251E-11 |
| | CBDmale | VehFemale | 2.70763E-11 | 0.434 | -3.2811E-11 | 7.5439E-11 |
| | | CBDfemale | 2.70763E-11 | 0.378 | -7.8163E-11 | 3.0087E-11 |
| | | THCfemale | 2.45714E-11 | 0.185 | -8.2036E-11 | 1.6199E-11 |
| | | TpCfemale | 2.45714E-11 | 0.008 | -1.1609E-10 | -1.7854E-11 |
| | | VehMale | 2.83726E-11 | 0.277 | -2.5572E-11 | 8.7860E-11 |
| | | THCmale | 2.52447E-11 | 0.049 | -1.0120E-10 | -2.7458E-13 |
| | | TpCmale | 2.40064E-11 | 0.945 | -4.6334E-11 | 4.9643E-11 |
| | THCmale | VehFemale | 2.76888E-11 | 0.012 | 1.6703E-11 | 1.2740E-10 |
| | | CBDfemale | 2.76888E-11 | 0.339 | -2.8649E-11 | 8.2049E-11 |
| | | THCfemale | 2.52447E-11 | 0.483 | -3.2644E-11 | 6.8283E-11 |
| | | TpCfemale | 2.52447E-11 | 0.523 | -6.6697E-11 | 3.4230E-11 |
| | | VehMale | 2.89576E-11 | 0.006 | 2.3997E-11 | 1.3977E-10 |
| | | CBDmale | 2.52447E-11 | 0.049 | 2.7458E-13 | 1.0120E-10 |
| | | TpCmale | 2.46951E-11 | 0.038 | 3.0274E-12 | 1.0176E-10 |
| | TpCmale | VehFemale | 2.65647E-11 | 0.462 | -3.3442E-11 | 7.2762E-11 |
| | | CBDfemale | 2.65647E-11 | 0.337 | -7.8794E-11 | 2.7410E-11 |
| | | THCfemale | 2.40064E-11 | 0.155 | -8.2561E-11 | 1.3415E-11 |
| | | TpCfemale | 2.40064E-11 | 0.006 | -1.1661E-10 | -2.0638E-11 |
| | | VehMale | 2.78847E-11 | 0.294 | -2.6251E-11 | 8.5231E-11 |
| | | CBDmale | 2.40064E-11 | 0.945 | -4.9643E-11 | 4.6334E-11 |
| | | THCmale | 2.46951E-11 | 0.038 | -1.0176E-10 | -3.0274E-12 |
| *N*-stearoyl serine | VehFemale | CBDfemale | 1.16316E-11 | 0.683 | -2.8081E-11 | 1.8521E-11 |

| | | | | | | |
|---|---|---|---|---|---|---|
| | | THCfemale | 1.12623E-11 | 0.429 | -3.1528E-11 | 1.3594E-11 |
| | | TpCfemale | 1.09664E-11 | 0.902 | -2.0608E-11 | 2.3329E-11 |
| | | VehMale | 1.21066E-11 | 0.719 | -1.9878E-11 | 2.8627E-11 |
| | | CBDmale | 1.12623E-11 | 0.776 | -1.9344E-11 | 2.5778E-11 |
| | | THCmale | 1.09664E-11 | 0.507 | -1.4649E-11 | 2.9287E-11 |
| | | TpCmale | 1.07238E-11 | 0.392 | -1.2228E-11 | 3.0736E-11 |
| | CBDfemale | VehFemale | 1.16316E-11 | 0.683 | -1.8521E-11 | 2.8081E-11 |
| | | THCfemale | 1.12623E-11 | 0.711 | -2.6748E-11 | 1.8374E-11 |
| | | TpCfemale | 1.09664E-11 | 0.578 | -1.5828E-11 | 2.8109E-11 |
| | | VehMale | 1.21066E-11 | 0.453 | -1.5097E-11 | 3.3407E-11 |
| | | CBDmale | 1.12623E-11 | 0.481 | -1.4564E-11 | 3.0558E-11 |
| | | THCmale | 1.09664E-11 | 0.275 | -9.8690E-12 | 3.4068E-11 |
| | | TpCmale | 1.07238E-11 | 0.196 | -7.4483E-12 | 3.5517E-11 |
| | THCfemale | VehFemale | 1.12623E-11 | 0.429 | -1.3594E-11 | 3.1528E-11 |
| | | CBDfemale | 1.12623E-11 | 0.711 | -1.8374E-11 | 2.6748E-11 |
| | | TpCfemale | 1.05738E-11 | 0.333 | -1.0854E-11 | 3.1509E-11 |
| | | VehMale | 1.17522E-11 | 0.261 | -1.0201E-11 | 3.6884E-11 |
| | | CBDmale | 1.08804E-11 | 0.268 | -9.6125E-12 | 3.3980E-11 |
| | | THCmale | 1.05738E-11 | 0.129 | -4.8959E-12 | 3.7468E-11 |
| | | TpCmale | 1.03220E-11 | 0.083 | -2.4567E-12 | 3.8898E-11 |
| | TpCfemale | VehFemale | 1.09664E-11 | 0.902 | -2.3329E-11 | 2.0608E-11 |
| | | CBDfemale | 1.09664E-11 | 0.578 | -2.8109E-11 | 1.5828E-11 |
| | | THCfemale | 1.05738E-11 | 0.333 | -3.1509E-11 | 1.0854E-11 |
| | | VehMale | 1.14689E-11 | 0.794 | -1.9961E-11 | 2.5989E-11 |
| | | CBDmale | 1.05738E-11 | 0.861 | -1.9326E-11 | 2.3038E-11 |
| | | THCmale | 1.02581E-11 | 0.564 | -1.4591E-11 | 2.6508E-11 |
| | | TpCmale | 9.99838E-12 | 0.433 | -1.2136E-11 | 2.7922E-11 |
| | VehMale | VehFemale | 1.21066E-11 | 0.719 | -2.8627E-11 | 1.9878E-11 |
| | | CBDfemale | 1.21066E-11 | 0.453 | -3.3407E-11 | 1.5097E-11 |
| | | THCfemale | 1.17522E-11 | 0.261 | -3.6884E-11 | 1.0201E-11 |
| | | TpCfemale | 1.14689E-11 | 0.794 | -2.5989E-11 | 1.9961E-11 |
| | | CBDmale | 1.17522E-11 | 0.922 | -2.4701E-11 | 2.2384E-11 |
| | | THCmale | 1.14689E-11 | 0.798 | -2.0031E-11 | 2.5919E-11 |
| | | TpCmale | 1.12372E-11 | 0.666 | -1.7632E-11 | 2.7390E-11 |
| | CBDmale | VehFemale | 1.12623E-11 | 0.776 | -2.5778E-11 | 1.9344E-11 |
| | | CBDfemale | 1.12623E-11 | 0.481 | -3.0558E-11 | 1.4564E-11 |
| | | THCfemale | 1.08804E-11 | 0.268 | -3.3980E-11 | 9.6125E-12 |
| | | TpCfemale | 1.05738E-11 | 0.861 | -2.3038E-11 | 1.9326E-11 |
| | | VehMale | 1.17522E-11 | 0.922 | -2.2384E-11 | 2.4701E-11 |
| | | THCmale | 1.05738E-11 | 0.699 | -1.7079E-11 | 2.5284E-11 |
| | | TpCmale | 1.03220E-11 | 0.561 | -1.4640E-11 | 2.6715E-11 |
| | THCmale | VehFemale | 1.09664E-11 | 0.507 | -2.9287E-11 | 1.4649E-11 |
| | | CBDfemale | 1.09664E-11 | 0.275 | -3.4068E-11 | 9.8690E-12 |
| | | THCfemale | 1.05738E-11 | 0.129 | -3.7468E-11 | 4.8959E-12 |
| | | TpCfemale | 1.02581E-11 | 0.564 | -2.6508E-11 | 1.4591E-11 |
| | | VehMale | 1.14689E-11 | 0.798 | -2.5919E-11 | 2.0031E-11 |
| | | CBDmale | 1.05738E-11 | 0.699 | -2.5284E-11 | 1.7079E-11 |
| | | TpCmale | 9.99838E-12 | 0.847 | -1.8094E-11 | 2.1964E-11 |
| | TpCmale | VehFemale | 1.07238E-11 | 0.392 | -3.0736E-11 | 1.2228E-11 |
| | | CBDfemale | 1.07238E-11 | 0.196 | -3.5517E-11 | 7.4483E-12 |
| | | THCfemale | 1.03220E-11 | 0.083 | -3.8898E-11 | 2.4567E-12 |
| | | TpCfemale | 9.99838E-12 | 0.433 | -2.7922E-11 | 1.2136E-11 |
| | | VehMale | 1.12372E-11 | 0.666 | -2.7390E-11 | 1.7632E-11 |
| | | CBDmale | 1.03220E-11 | 0.561 | -2.6715E-11 | 1.4640E-11 |
| | | THCmale | 9.99838E-12 | 0.847 | -2.1964E-11 | 1.8094E-11 |
| *N*-oleoyl serine | VehFemale | CBDfemale | 1.37505E-11 | 0.665 | -3.3488E-11 | 2.1522E-11 |
| | | THCfemale | 1.26025E-11 | 0.000 | -7.8027E-11 | -2.7609E-11 |
| | | TpCfemale | 1.29099E-11 | 0.000 | -7.5949E-11 | -2.4302E-11 |
| | | VehMale | 1.43486E-11 | 0.124 | -6.3082E-12 | 5.1095E-11 |
| | | CBDmale | 1.26025E-11 | 0.589 | -1.8363E-11 | 3.2054E-11 |

| | | | | | | |
|---|---|---|---|---|---|---|
| | | THCmale | 1.32842E-11 | 0.190 | -4.4183E-11 | 8.9621E-12 |
| | | TpCmale | 1.26025E-11 | 0.948 | -2.6033E-11 | 2.4385E-11 |
| | CBDfemale | VehFemale | 1.37505E-11 | 0.665 | -2.1522E-11 | 3.3488E-11 |
| | | THCfemale | 1.30931E-11 | 0.001 | -7.3025E-11 | -2.0645E-11 |
| | | TpCfemale | 1.33892E-11 | 0.002 | -7.0925E-11 | -1.7360E-11 |
| | | VehMale | 1.47813E-11 | 0.060 | -1.1905E-12 | 5.7944E-11 |
| | | CBDmale | 1.30931E-11 | 0.331 | -1.3361E-11 | 3.9019E-11 |
| | | THCmale | 1.37505E-11 | 0.401 | -3.9132E-11 | 1.5878E-11 |
| | | TpCmale | 1.30931E-11 | 0.695 | -2.1031E-11 | 3.1349E-11 |
| | THCfemale | VehFemale | 1.26025E-11 | 0.000 | 2.7609E-11 | 7.8027E-11 |
| | | CBDfemale | 1.30931E-11 | 0.001 | 2.0645E-11 | 7.3025E-11 |
| | | TpCfemale | 1.22074E-11 | 0.826 | -2.1726E-11 | 2.7111E-11 |
| | | VehMale | 1.37199E-11 | 0.000 | 4.7768E-11 | 1.0266E-10 |
| | | CBDmale | 1.18818E-11 | 0.000 | 3.5896E-11 | 8.3431E-11 |
| | | THCmale | 1.26025E-11 | 0.007 | 9.9992E-12 | 6.0417E-11 |
| | | TpCmale | 1.18818E-11 | 0.000 | 2.8227E-11 | 7.5761E-11 |
| | TpCfemale | VehFemale | 1.29099E-11 | 0.000 | 2.4302E-11 | 7.5949E-11 |
| | | CBDfemale | 1.33892E-11 | 0.002 | 1.7360E-11 | 7.0925E-11 |
| | | THCfemale | 1.22074E-11 | 0.826 | -2.7111E-11 | 2.1726E-11 |
| | | VehMale | 1.40028E-11 | 0.000 | 4.4509E-11 | 1.0053E-10 |
| | | CBDmale | 1.22074E-11 | 0.000 | 3.2552E-11 | 8.1389E-11 |
| | | THCmale | 1.29099E-11 | 0.014 | 6.6914E-12 | 5.8339E-11 |
| | | TpCmale | 1.22074E-11 | 0.000 | 2.4883E-11 | 7.3720E-11 |
| | VehMale | VehFemale | 1.43486E-11 | 0.124 | -5.1095E-11 | 6.3082E-12 |
| | | CBDfemale | 1.47813E-11 | 0.060 | -5.7944E-11 | 1.1905E-12 |
| | | THCfemale | 1.37199E-11 | 0.000 | -1.0266E-10 | -4.7768E-11 |
| | | TpCfemale | 1.40028E-11 | 0.000 | -1.0053E-10 | -4.4509E-11 |
| | | CBDmale | 1.37199E-11 | 0.262 | -4.2992E-11 | 1.1896E-11 |
| | | THCmale | 1.43486E-11 | 0.007 | -6.8705E-11 | -1.1302E-11 |
| | | TpCmale | 1.37199E-11 | 0.096 | -5.0661E-11 | 4.2264E-12 |
| | CBDmale | VehFemale | 1.26025E-11 | 0.589 | -3.2054E-11 | 1.8363E-11 |
| | | CBDfemale | 1.30931E-11 | 0.331 | -3.9019E-11 | 1.3361E-11 |
| | | THCfemale | 1.18818E-11 | 0.000 | -8.3431E-11 | -3.5896E-11 |
| | | TpCfemale | 1.22074E-11 | 0.000 | -8.1389E-11 | -3.2552E-11 |
| | | VehMale | 1.37199E-11 | 0.262 | -1.1896E-11 | 4.2992E-11 |
| | | THCmale | 1.26025E-11 | 0.057 | -4.9664E-11 | 7.5321E-13 |
| | | TpCmale | 1.18818E-11 | 0.521 | -3.1437E-11 | 1.6097E-11 |
| | THCmale | VehFemale | 1.32842E-11 | 0.190 | -8.9621E-12 | 4.4183E-11 |
| | | CBDfemale | 1.37505E-11 | 0.401 | -1.5878E-11 | 3.9132E-11 |
| | | THCfemale | 1.26025E-11 | 0.007 | -6.0417E-11 | -9.9992E-12 |
| | | TpCfemale | 1.29099E-11 | 0.014 | -5.8339E-11 | -6.6914E-12 |
| | | VehMale | 1.43486E-11 | 0.007 | 1.1302E-11 | 6.8705E-11 |
| | | CBDmale | 1.26025E-11 | 0.057 | -7.5321E-13 | 4.9664E-11 |
| | | TpCmale | 1.26025E-11 | 0.188 | -8.4228E-12 | 4.1995E-11 |
| | TpCmale | VehFemale | 1.26025E-11 | 0.948 | -2.4385E-11 | 2.6033E-11 |
| | | CBDfemale | 1.30931E-11 | 0.695 | -3.1349E-11 | 2.1031E-11 |
| | | THCfemale | 1.18818E-11 | 0.000 | -7.5761E-11 | -2.8227E-11 |
| | | TpCfemale | 1.22074E-11 | 0.000 | -7.3720E-11 | -2.4883E-11 |
| | | VehMale | 1.37199E-11 | 0.096 | -4.2264E-12 | 5.0661E-11 |
| | | CBDmale | 1.18818E-11 | 0.521 | -1.6097E-11 | 3.1437E-11 |
| | | THCmale | 1.26025E-11 | 0.188 | -4.1995E-11 | 8.4228E-12 |
| *N*-linoleoyl serine | VehFemale | CBDfemale | 4.23100E-11 | 0.372 | -1.2270E-10 | 4.6563E-11 |
| | | THCfemale | 3.98902E-11 | 0.164 | -1.3604E-10 | 2.3545E-11 |
| | | TpCfemale | 3.98902E-11 | 0.001 | -2.1988E-10 | -6.0290E-11 |
| | | VehMale | 4.23100E-11 | 0.487 | -5.5061E-11 | 1.1420E-10 |
| | | CBDmale | 3.90078E-11 | 0.590 | -5.6923E-11 | 9.9132E-11 |
| | | THCmale | 3.98902E-11 | 0.081 | -1.5050E-10 | 9.0893E-12 |
| | | TpCmale | 3.90078E-11 | 0.402 | -1.1097E-10 | 4.5085E-11 |
| | CBDfemale | VehFemale | 4.23100E-11 | 0.372 | -4.6563E-11 | 1.2270E-10 |
| | | THCfemale | 3.98902E-11 | 0.650 | -9.7970E-11 | 6.1614E-11 |

| | | | | | | |
|---|---|---|---|---|---|---|
| | | TpCfemale | 3.98902E-11 | 0.013 | -1.8181E-10 | -2.2221E-11 |
| | | VehMale | 4.23100E-11 | 0.115 | -1.6992E-11 | 1.5227E-10 |
| | | CBDmale | 3.90078E-11 | 0.135 | -1.8853E-11 | 1.3720E-10 |
| | | THCmale | 3.98902E-11 | 0.417 | -1.1243E-10 | 4.7159E-11 |
| | | TpCmale | 3.90078E-11 | 0.896 | -7.2901E-11 | 8.3154E-11 |
| | THCfemale | VehFemale | 3.98902E-11 | 0.164 | -2.3545E-11 | 1.3604E-10 |
| | | CBDfemale | 3.98902E-11 | 0.650 | -6.1614E-11 | 9.7970E-11 |
| | | TpCfemale | 3.73139E-11 | 0.028 | -1.5847E-10 | -9.1970E-12 |
| | | VehMale | 3.98902E-11 | 0.035 | 6.0265E-12 | 1.6561E-10 |
| | | CBDmale | 3.63691E-11 | 0.038 | 4.6028E-12 | 1.5010E-10 |
| | | THCmale | 3.73139E-11 | 0.700 | -8.9095E-11 | 6.0183E-11 |
| | | TpCmale | 3.63691E-11 | 0.524 | -4.9445E-11 | 9.6053E-11 |
| | TpCfemale | VehFemale | 3.98902E-11 | 0.001 | 6.0290E-11 | 2.1988E-10 |
| | | CBDfemale | 3.98902E-11 | 0.013 | 2.2221E-11 | 1.8181E-10 |
| | | THCfemale | 3.73139E-11 | 0.028 | 9.1970E-12 | 1.5847E-10 |
| | | VehMale | 3.98902E-11 | 0.000 | 8.9862E-11 | 2.4945E-10 |
| | | CBDmale | 3.63691E-11 | 0.000 | 8.8439E-11 | 2.3394E-10 |
| | | THCmale | 3.73139E-11 | 0.068 | -5.2590E-12 | 1.4402E-10 |
| | | TpCmale | 3.63691E-11 | 0.005 | 3.4391E-11 | 1.7989E-10 |
| | VehMale | VehFemale | 4.23100E-11 | 0.487 | -1.1420E-10 | 5.5061E-11 |
| | | CBDfemale | 4.23100E-11 | 0.115 | -1.5227E-10 | 1.6992E-11 |
| | | THCfemale | 3.98902E-11 | 0.035 | -1.6561E-10 | -6.0265E-12 |
| | | TpCfemale | 3.98902E-11 | 0.000 | -2.4945E-10 | -8.9862E-11 |
| | | CBDmale | 3.90078E-11 | 0.829 | -8.6494E-11 | 6.9560E-11 |
| | | THCmale | 3.98902E-11 | 0.015 | -1.8007E-10 | -2.0482E-11 |
| | | TpCmale | 3.90078E-11 | 0.114 | -1.4054E-10 | 1.5513E-11 |
| | CBDmale | VehFemale | 3.90078E-11 | 0.590 | -9.9132E-11 | 5.6923E-11 |
| | | CBDfemale | 3.90078E-11 | 0.135 | -1.3720E-10 | 1.8853E-11 |
| | | THCfemale | 3.63691E-11 | 0.038 | -1.5010E-10 | -4.6028E-12 |
| | | TpCfemale | 3.63691E-11 | 0.000 | -2.3394E-10 | -8.8439E-11 |
| | | VehMale | 3.90078E-11 | 0.829 | -6.9560E-11 | 8.6494E-11 |
| | | THCmale | 3.63691E-11 | 0.014 | -1.6456E-10 | -1.9059E-11 |
| | | TpCmale | 3.53990E-11 | 0.132 | -1.2486E-10 | 1.6761E-11 |
| | THCmale | VehFemale | 3.98902E-11 | 0.081 | -9.0893E-12 | 1.5050E-10 |
| | | CBDfemale | 3.98902E-11 | 0.417 | -4.7159E-11 | 1.1243E-10 |
| | | THCfemale | 3.73139E-11 | 0.700 | -6.0183E-11 | 8.9095E-11 |
| | | TpCfemale | 3.73139E-11 | 0.068 | -1.4402E-10 | 5.2590E-12 |
| | | VehMale | 3.98902E-11 | 0.015 | 2.0482E-11 | 1.8007E-10 |
| | | CBDmale | 3.63691E-11 | 0.014 | 1.9059E-11 | 1.6456E-10 |
| | | TpCmale | 3.63691E-11 | 0.303 | -3.4989E-11 | 1.1051E-10 |
| | TpCmale | VehFemale | 3.90078E-11 | 0.402 | -4.5085E-11 | 1.1097E-10 |
| | | CBDfemale | 3.90078E-11 | 0.896 | -8.3154E-11 | 7.2901E-11 |
| | | THCfemale | 3.63691E-11 | 0.524 | -9.6053E-11 | 4.9445E-11 |
| | | TpCfemale | 3.63691E-11 | 0.005 | -1.7989E-10 | -3.4391E-11 |
| | | VehMale | 3.90078E-11 | 0.114 | -1.5513E-11 | 1.4054E-10 |
| | | CBDmale | 3.53990E-11 | 0.132 | -1.6761E-11 | 1.2486E-10 |
| | | THCmale | 3.63691E-11 | 0.303 | -1.1051E-10 | 3.4989E-11 |
| *N*-arachidonoyl serine | VehFemale | CBDfemale | 3.10814E-12 | 0.061 | -1.2153E-11 | 2.9029E-13 |
| | | THCfemale | 2.81542E-12 | 0.019 | -1.2458E-11 | -1.1866E-12 |
| | | TpCfemale | 2.81542E-12 | 0.000 | -1.6137E-11 | -4.8658E-12 |
| | | VehMale | 2.98620E-12 | 0.628 | -4.5241E-12 | 7.4310E-12 |
| | | CBDmale | 2.81542E-12 | 0.599 | -7.1248E-12 | 4.1466E-12 |
| | | THCmale | 2.81542E-12 | 0.051 | -1.1251E-11 | 2.0025E-14 |
| | | TpCmale | 2.75314E-12 | 0.450 | -7.6036E-12 | 3.4184E-12 |
| | CBDfemale | VehFemale | 3.10814E-12 | 0.061 | -2.9029E-13 | 1.2153E-11 |
| | | THCfemale | 2.94444E-12 | 0.763 | -6.7849E-12 | 5.0030E-12 |
| | | TpCfemale | 2.94444E-12 | 0.126 | -1.0464E-11 | 1.3238E-12 |
| | | VehMale | 3.10814E-12 | 0.021 | 1.1632E-12 | 1.3606E-11 |
| | | CBDmale | 2.94444E-12 | 0.137 | -1.4517E-12 | 1.0336E-11 |
| | | THCmale | 2.94444E-12 | 0.915 | -5.5783E-12 | 6.2096E-12 |

| | | | | | | |
|---|---|---|---|---|---|---|
| | | TpCmale | 2.88495E-12 | 0.189 | -1.9361E-12 | 9.6136E-12 |
| | THCfemale | VehFemale | 2.81542E-12 | 0.019 | 1.1866E-12 | 1.2458E-11 |
| | | CBDfemale | 2.94444E-12 | 0.763 | -5.0030E-12 | 6.7849E-12 |
| | | TpCfemale | 2.63358E-12 | 0.168 | -8.9509E-12 | 1.5925E-12 |
| | | VehMale | 2.81542E-12 | 0.005 | 2.6401E-12 | 1.3911E-11 |
| | | CBDmale | 2.63358E-12 | 0.047 | 6.1516E-14 | 1.0605E-11 |
| | | THCmale | 2.63358E-12 | 0.649 | -4.0651E-12 | 6.4783E-12 |
| | | TpCmale | 2.56690E-12 | 0.071 | -4.0850E-13 | 9.8679E-12 |
| | TpCfemale | VehFemale | 2.81542E-12 | 0.000 | 4.8658E-12 | 1.6137E-11 |
| | | CBDfemale | 2.94444E-12 | 0.126 | -1.3238E-12 | 1.0464E-11 |
| | | THCfemale | 2.63358E-12 | 0.168 | -1.5925E-12 | 8.9509E-12 |
| | | VehMale | 2.81542E-12 | 0.000 | 6.3193E-12 | 1.7591E-11 |
| | | CBDmale | 2.63358E-12 | 0.001 | 3.7407E-12 | 1.4284E-11 |
| | | THCmale | 2.63358E-12 | 0.069 | -3.8584E-13 | 1.0158E-11 |
| | | TpCmale | 2.56690E-12 | 0.002 | 3.2707E-12 | 1.3547E-11 |
| | VehMale | VehFemale | 2.98620E-12 | 0.628 | -7.4310E-12 | 4.5241E-12 |
| | | CBDfemale | 3.10814E-12 | 0.021 | -1.3606E-11 | -1.1632E-12 |
| | | THCfemale | 2.81542E-12 | 0.005 | -1.3911E-11 | -2.6401E-12 |
| | | TpCfemale | 2.81542E-12 | 0.000 | -1.7591E-11 | -6.3193E-12 |
| | | CBDmale | 2.81542E-12 | 0.300 | -8.5782E-12 | 2.6931E-12 |
| | | THCmale | 2.81542E-12 | 0.015 | -1.2705E-11 | -1.4334E-12 |
| | | TpCmale | 2.75314E-12 | 0.203 | -9.0571E-12 | 1.9650E-12 |
| | CBDmale | VehFemale | 2.81542E-12 | 0.599 | -4.1466E-12 | 7.1248E-12 |
| | | CBDfemale | 2.94444E-12 | 0.137 | -1.0336E-11 | 1.4517E-12 |
| | | THCfemale | 2.63358E-12 | 0.047 | -1.0605E-11 | -6.1516E-14 |
| | | TpCfemale | 2.63358E-12 | 0.001 | -1.4284E-11 | -3.7407E-12 |
| | | VehMale | 2.81542E-12 | 0.300 | -2.6931E-12 | 8.5782E-12 |
| | | THCmale | 2.63358E-12 | 0.123 | -9.3983E-12 | 1.1451E-12 |
| | | TpCmale | 2.56690E-12 | 0.815 | -5.7417E-12 | 4.5347E-12 |
| | THCmale | VehFemale | 2.81542E-12 | 0.051 | -2.0025E-14 | 1.1251E-11 |
| | | CBDfemale | 2.94444E-12 | 0.915 | -6.2096E-12 | 5.5783E-12 |
| | | THCfemale | 2.63358E-12 | 0.649 | -6.4783E-12 | 4.0651E-12 |
| | | TpCfemale | 2.63358E-12 | 0.069 | -1.0158E-11 | 3.8584E-13 |
| | | VehMale | 2.81542E-12 | 0.015 | 1.4334E-12 | 1.2705E-11 |
| | | CBDmale | 2.63358E-12 | 0.123 | -1.1451E-12 | 9.3983E-12 |
| | | TpCmale | 2.56690E-12 | 0.175 | -1.6151E-12 | 8.6613E-12 |
| | TpCmale | VehFemale | 2.75314E-12 | 0.450 | -3.4184E-12 | 7.6036E-12 |
| | | CBDfemale | 2.88495E-12 | 0.189 | -9.6136E-12 | 1.9361E-12 |
| | | THCfemale | 2.56690E-12 | 0.071 | -9.8679E-12 | 4.0850E-13 |
| | | TpCfemale | 2.56690E-12 | 0.002 | -1.3547E-11 | -3.2707E-12 |
| | | VehMale | 2.75314E-12 | 0.203 | -1.9650E-12 | 9.0571E-12 |
| | | CBDmale | 2.56690E-12 | 0.815 | -4.5347E-12 | 5.7417E-12 |
| | | THCmale | 2.56690E-12 | 0.175 | -8.6613E-12 | 1.6151E-12 |
| *N*-docosahexaenoyl serine | VehFemale | CBDfemale | 1.90743E-11 | 0.075 | -7.2734E-11 | 3.6286E-12 |
| | | THCfemale | 1.71618E-11 | 0.642 | -4.2386E-11 | 2.6321E-11 |
| | | TpCfemale | 1.67532E-11 | 0.008 | -7.9769E-11 | -1.2699E-11 |
| | | VehMale | 2.01348E-11 | 0.810 | -3.5430E-11 | 4.5179E-11 |
| | | CBDmale | 1.71618E-11 | 0.507 | -2.2896E-11 | 4.5810E-11 |
| | | THCmale | 1.71618E-11 | 0.181 | -5.7566E-11 | 1.1140E-11 |
| | | TpCmale | 1.67532E-11 | 0.787 | -3.8077E-11 | 2.8993E-11 |
| | CBDfemale | VehFemale | 1.90743E-11 | 0.075 | -3.6286E-12 | 7.2734E-11 |
| | | THCfemale | 1.86146E-11 | 0.160 | -1.0741E-11 | 6.3782E-11 |
| | | TpCfemale | 1.82385E-11 | 0.524 | -4.8189E-11 | 2.4828E-11 |
| | | VehMale | 2.13866E-11 | 0.070 | -3.3827E-12 | 8.2237E-11 |
| | | CBDmale | 1.86146E-11 | 0.016 | 8.7484E-12 | 8.3271E-11 |
| | | THCmale | 1.86146E-11 | 0.545 | -2.5921E-11 | 4.8601E-11 |
| | | TpCmale | 1.82385E-11 | 0.105 | -6.4975E-12 | 6.6519E-11 |
| | THCfemale | VehFemale | 1.71618E-11 | 0.642 | -2.6321E-11 | 4.2386E-11 |
| | | CBDfemale | 1.86146E-11 | 0.160 | -6.3782E-11 | 1.0741E-11 |
| | | TpCfemale | 1.62279E-11 | 0.022 | -7.0685E-11 | -5.7176E-12 |

| | | | | | | |
|---|---|---|---|---|---|---|
| | | VehMale | 1.96999E-11 | 0.515 | -2.6527E-11 | 5.2340E-11 |
| | | CBDmale | 1.66494E-11 | 0.247 | -1.3838E-11 | 5.2817E-11 |
| | | THCmale | 1.66494E-11 | 0.366 | -4.8508E-11 | 1.8147E-11 |
| | | TpCmale | 1.62279E-11 | 0.830 | -2.8993E-11 | 3.5974E-11 |
| | TpCfemale | VehFemale | 1.67532E-11 | 0.008 | 1.2699E-11 | 7.9769E-11 |
| | | CBDfemale | 1.82385E-11 | 0.524 | -2.4828E-11 | 4.8189E-11 |
| | | THCfemale | 1.62279E-11 | 0.022 | 5.7176E-12 | 7.0685E-11 |
| | | VehMale | 1.93449E-11 | 0.011 | 1.2385E-11 | 8.9831E-11 |
| | | CBDmale | 1.62279E-11 | 0.001 | 2.5207E-11 | 9.0174E-11 |
| | | THCmale | 1.62279E-11 | 0.161 | -9.4627E-12 | 5.5505E-11 |
| | | TpCmale | 1.57950E-11 | 0.011 | 1.0075E-11 | 7.3309E-11 |
| | VehMale | VehFemale | 2.01348E-11 | 0.810 | -4.5179E-11 | 3.5430E-11 |
| | | CBDfemale | 2.13866E-11 | 0.070 | -8.2237E-11 | 3.3827E-12 |
| | | THCfemale | 1.96999E-11 | 0.515 | -5.2340E-11 | 2.6527E-11 |
| | | TpCfemale | 1.93449E-11 | 0.011 | -8.9831E-11 | -1.2385E-11 |
| | | CBDmale | 1.96999E-11 | 0.739 | -3.2851E-11 | 4.6016E-11 |
| | | THCmale | 1.96999E-11 | 0.159 | -6.7521E-11 | 1.1346E-11 |
| | | TpCmale | 1.93449E-11 | 0.628 | -4.8139E-11 | 2.9307E-11 |
| | CBDmale | VehFemale | 1.71618E-11 | 0.507 | -4.5810E-11 | 2.2896E-11 |
| | | CBDfemale | 1.86146E-11 | 0.016 | -8.3271E-11 | -8.7484E-12 |
| | | THCfemale | 1.66494E-11 | 0.247 | -5.2817E-11 | 1.3838E-11 |
| | | TpCfemale | 1.62279E-11 | 0.001 | -9.0174E-11 | -2.5207E-11 |
| | | VehMale | 1.96999E-11 | 0.739 | -4.6016E-11 | 3.2851E-11 |
| | | THCmale | 1.66494E-11 | 0.042 | -6.7997E-11 | -1.3421E-12 |
| | | TpCmale | 1.62279E-11 | 0.328 | -4.8482E-11 | 1.6485E-11 |
| | THCmale | VehFemale | 1.71618E-11 | 0.181 | -1.1140E-11 | 5.7566E-11 |
| | | CBDfemale | 1.86146E-11 | 0.545 | -4.8601E-11 | 2.5921E-11 |
| | | THCfemale | 1.66494E-11 | 0.366 | -1.8147E-11 | 4.8508E-11 |
| | | TpCfemale | 1.62279E-11 | 0.161 | -5.5505E-11 | 9.4627E-12 |
| | | VehMale | 1.96999E-11 | 0.159 | -1.1346E-11 | 6.7521E-11 |
| | | CBDmale | 1.66494E-11 | 0.042 | 1.3421E-12 | 6.7997E-11 |
| | | TpCmale | 1.62279E-11 | 0.255 | -1.3813E-11 | 5.1154E-11 |
| | TpCmale | VehFemale | 1.67532E-11 | 0.787 | -2.8993E-11 | 3.8077E-11 |
| | | CBDfemale | 1.82385E-11 | 0.105 | -6.6519E-11 | 6.4975E-12 |
| | | THCfemale | 1.62279E-11 | 0.830 | -3.5974E-11 | 2.8993E-11 |
| | | TpCfemale | 1.57950E-11 | 0.011 | -7.3309E-11 | -1.0075E-11 |
| | | VehMale | 1.93449E-11 | 0.628 | -2.9307E-11 | 4.8139E-11 |
| | | CBDmale | 1.62279E-11 | 0.328 | -1.6485E-11 | 4.8482E-11 |
| | | THCmale | 1.62279E-11 | 0.255 | -5.1154E-11 | 1.3813E-11 |

Supplemental Table 14.1 Descriptive statistics for *N*-acyl taurines (mol/L) in plasma collected from male and female adult mice 2h after receiving an i.p. injection of 10mg/kg CBD, THC, or THC+CBD

| | | N | Mean | Std. Deviation | Std. Error | 95% Confidence Interval for Mean | | Minimum | Maximum |
|---|---|---|---|---|---|---|---|---|---|
| | | | | | | Lower Bound | Upper Bound | | |
| *N*-palmitoyl taurine | Veh F | BAL | | | | | | | |
| | CBD F | BAL | | | | | | | |
| | THC F | BAL | | | | | | | |
| | THC+CBD F | BAL | | | | | | | |
| | Veh M | BDL | | | | | | | |
| | CBD M | BAL | | | | | | | |
| | THC M | BAL | | | | | | | |
| | THC+CBD M | BAL | | | | | | | |
| *N*-stearoyl taurine | Veh F | BDL | | | | | | | |
| | CBD F | BAL | | | | | | | |
| | THC F | BAL | | | | | | | |
| | THC+CBD F | BAL | | | | | | | |
| | Veh M | BAL | | | | | | | |
| | CBD M | BAL | | | | | | | |
| | THC M | BAL | | | | | | | |
| | THC+CBD M | BAL | | | | | | | |
| *N*-oleoyl taurine | Veh F | BAL | | | | | | | |
| | CBD F | 5 | 6.3550E-08 | 2.42924E-08 | 1.08639E-08 | 3.3387E-08 | 9.3713E-08 | 3.79E-08 | 9.95E-08 |
| | THC F | 5 | 5.9617E-08 | 8.08006E-09 | 3.61351E-09 | 4.9584E-08 | 6.9650E-08 | 5.03E-08 | 7.08E-08 |
| | THC+CBD F | 7 | 7.0024E-08 | 1.35739E-08 | 5.13045E-09 | 5.7470E-08 | 8.2577E-08 | 5.47E-08 | 9.67E-08 |
| | Veh M | BAL | | | | | | | |
| | CBD M | BAL | | | | | | | |
| | THC M | BAL | | | | | | | |
| | THC+CBD M | 4 | 6.1558E-08 | 6.12583E-09 | 3.06291E-09 | 5.1811E-08 | 7.1306E-08 | 5.45E-08 | 6.92E-08 |
| *N*-arachidonoyl taurine | Veh F | BDL | | | | | | | |
| | CBD F | BAL | | | | | | | |
| | THC F | BDL | | | | | | | |
| | THC+CBD F | BDL | | | | | | | |
| | Veh M | BDL | | | | | | | |
| | CBD M | BDL | | | | | | | |
| | THC M | BDL | | | | | | | |
| | THC+CBD M | BDL | | | | | | | |

Supplemental Table 14.2 Descriptive statistics for *N*-acyl taurines (mol/g) in liver collected from male and female adult mice 2h after receiving an i.p. injection of 10mg/kg CBD, THC, or THC+CBD

| | | N | Mean | Std. Deviation | Std. Error | 95% Confidence Interval for Mean | | Minimum | Maximum |
|---|---|---|---|---|---|---|---|---|---|
| | | | | | | Lower Bound | Upper Bound | | |
| *N*-palmitoyl taurine | Veh F | 8 | 2.7084E-09 | 7.63639E-10 | 2.69987E-10 | 2.0700E-09 | 3.3468E-09 | 1.89E-09 | 4.11E-09 |
| | CBD F | 8 | 2.3274E-09 | 1.23597E-09 | 4.36980E-10 | 1.2941E-09 | 3.3607E-09 | 8.89E-10 | 4.62E-09 |
| | THC F | 9 | 3.8833E-09 | 5.65830E-10 | 1.88610E-10 | 3.4484E-09 | 4.3182E-09 | 3.03E-09 | 4.55E-09 |
| | THC+CBD F | 9 | 3.0574E-09 | 1.07088E-09 | 3.56960E-10 | 2.2342E-09 | 3.8805E-09 | 1.84E-09 | 5.14E-09 |
| | Veh M | 7 | 1.9477E-09 | 3.59517E-10 | 1.35885E-10 | 1.6152E-09 | 2.2802E-09 | 1.60E-09 | 2.66E-09 |
| | CBD M | 10 | 2.5793E-09 | 1.55174E-09 | 4.90703E-10 | 1.4693E-09 | 3.6894E-09 | 8.36E-10 | 5.64E-09 |
| | THC M | 9 | 3.4015E-09 | 1.10477E-09 | 3.68257E-10 | 2.5523E-09 | 4.2507E-09 | 1.32E-09 | 4.88E-09 |
| | THC+CBD M | 11 | 3.4783E-09 | 1.05484E-09 | 3.18047E-10 | 2.7696E-09 | 4.1869E-09 | 1.63E-09 | 5.25E-09 |
| *N*-stearoyl taurine | Veh F | 6 | 1.3419E-10 | 7.83565E-11 | 3.19889E-11 | 5.1961E-11 | 2.1642E-10 | 7.48E-11 | 2.50E-10 |
| | CBD F | 4 | 1.3715E-10 | 1.73455E-11 | 8.67274E-12 | 1.0955E-10 | 1.6475E-10 | 1.14E-10 | 1.53E-10 |
| | THC F | 10 | 2.3307E-10 | 6.14218E-11 | 1.94233E-11 | 1.8913E-10 | 2.7700E-10 | 1.50E-10 | 3.36E-10 |
| | THC+CBD F | 9 | 2.7434E-10 | 1.66873E-10 | 5.56245E-11 | 1.4607E-10 | 4.0261E-10 | 1.01E-10 | 5.43E-10 |
| | Veh M | 4 | 7.9442E-11 | 3.59962E-11 | 1.79981E-11 | 2.2164E-11 | 1.3672E-10 | 4.40E-11 | 1.19E-10 |
| | CBD M | 5 | 9.9301E-11 | 6.29729E-11 | 2.81623E-11 | 2.1110E-11 | 1.7749E-10 | 4.07E-11 | 1.79E-10 |
| | THC M | 4 | 1.0304E-10 | 8.03157E-11 | 4.01578E-11 | -2.4761E-11 | 2.3084E-10 | 4.80E-11 | 2.20E-10 |
| | THC+CBD M | 8 | 1.0054E-10 | 5.55859E-11 | 1.96526E-11 | 5.4070E-11 | 1.4701E-10 | 4.27E-11 | 2.13E-10 |
| *N*-oleoyl taurine | Veh F | 8 | 1.2799E-09 | 4.90364E-10 | 1.73370E-10 | 8.6993E-10 | 1.6898E-09 | 7.91E-10 | 2.11E-09 |
| | CBD F | 8 | 9.4773E-10 | 5.06881E-10 | 1.79209E-10 | 5.2397E-10 | 1.3715E-09 | 2.63E-10 | 1.88E-09 |
| | THC F | 10 | 2.9022E-09 | 1.27958E-09 | 4.04640E-10 | 1.9869E-09 | 3.8176E-09 | 1.41E-09 | 4.81E-09 |
| | THC+CBD F | 9 | 2.0561E-09 | 7.06864E-10 | 2.35621E-10 | 1.5127E-09 | 2.5994E-09 | 1.00E-09 | 3.44E-09 |
| | Veh M | 7 | 1.1007E-09 | 6.80259E-10 | 2.57114E-10 | 4.7161E-10 | 1.7299E-09 | 4.23E-10 | 2.27E-09 |
| | CBD M | 10 | 9.5912E-10 | 3.63590E-10 | 1.14977E-10 | 6.9903E-10 | 1.2192E-09 | 3.39E-10 | 1.32E-09 |
| | THC M | 8 | 1.1466E-09 | 4.67337E-10 | 1.65229E-10 | 7.5588E-10 | 1.5373E-09 | 3.72E-10 | 1.83E-09 |
| | THC+CBD M | 10 | 2.6043E-09 | 7.83272E-10 | 2.47692E-10 | 2.0440E-09 | 3.1646E-09 | 1.75E-09 | 4.36E-09 |
| *N*-arachidonoyl taurine | Veh F | 8 | 2.1342E-09 | 9.50735E-10 | 3.36135E-10 | 1.3394E-09 | 2.9290E-09 | 1.19E-09 | 3.94E-09 |
| | CBD F | 8 | 2.0647E-09 | 1.40095E-09 | 4.95310E-10 | 8.9344E-10 | 3.2359E-09 | 4.61E-10 | 3.92E-09 |
| | THC F | 10 | 5.1709E-09 | 1.71197E-09 | 5.41373E-10 | 3.9463E-09 | 6.3956E-09 | 2.61E-09 | 8.40E-09 |
| | THC+CBD F | 10 | 5.7855E-09 | 2.75117E-09 | 8.69995E-10 | 3.8175E-09 | 7.7536E-09 | 1.68E-09 | 1.09E-08 |
| | Veh M | 7 | 1.1352E-09 | 1.99729E-10 | 7.54904E-11 | 9.5044E-10 | 1.3199E-09 | 8.26E-10 | 1.43E-09 |
| | CBD M | 9 | 2.2082E-09 | 1.28277E-09 | 4.27591E-10 | 1.2221E-09 | 3.1942E-09 | 9.04E-10 | 4.98E-09 |
| | THC M | 9 | 4.5639E-09 | 1.53985E-09 | 5.13284E-10 | 3.3802E-09 | 5.7475E-09 | 1.50E-09 | 6.45E-09 |
| | THC+CBD M | 11 | 5.3254E-09 | 2.23852E-09 | 6.74939E-10 | 3.8215E-09 | 6.8293E-09 | 2.35E-09 | 9.35E-09 |

Supplemental Table 14.3 Independent t-test comparing male and female levels of *N*-acyl taurines measured in plasma collected from adult mice 2h after an i.p. injection of 10mg/kg CBD, THC, or THC+CBD

| | | | Levene's Test for Equality of Variances | | t-test for Equality of Means | | | | | | |
|---|---|---|---|---|---|---|---|---|---|---|---|
| | | | | | | | Significance | | | 95% Confidence Interval of the Difference | |
| | | | F | Sig. | t | df | One-Sided p | Two-Sided p | Std. Error Difference | Lower | Upper |
| THC+CBD | *N*-oleoyl taurine | Equal variances assumed | 1.228 | 0.296 | 1.161 | 9 | 0.138 | 0.276 | 7.29179E-09 | -8.02952E-09 | 2.49608E-08 |
| | | Equal variances not assumed | | | 1.417 | 8.803 | 0.095 | 0.191 | 5.97520E-09 | -5.09749E-09 | 2.20288E-08 |

#### Supplemental Table 14.4 Independent t-test comparing male and female levels of *N*-acyl taurines measured in liver collected from adult mice 2h after an i.p. injection of 10mg/kg CBD, THC, or THC+CBD

| | | | Levene's Test for Equality of Variances | | t-test for Equality of Means | | | | | | | |
|---|---|---|---|---|---|---|---|---|---|---|---|---|
| | | | | | | | Significance | | | | 95% Confidence Interval of the Difference | |
| | | | F | Sig. | t | df | One-Sided p | Two-Sided p | Mean Difference | Std. Error Difference | Lower | Upper |
| Veh | *N*-palmitoyl taurine | Equal variances assumed | 2.033 | 0.178 | 2.404 | 13 | 0.016 | 0.032 | 7.60665E-10 | 3.16364E-10 | 7.72011E-11 | 1.44413E-09 |
| | | Equal variances not assumed | | | 2.517 | 10.230 | 0.015 | 0.030 | 7.60665E-10 | 3.02254E-10 | 8.92443E-11 | 1.43209E-09 |
| | *N*-stearoyl taurine | Equal variances assumed | 5.867 | 0.042 | 1.290 | 8 | 0.117 | 0.233 | 5.47492E-11 | 4.24423E-11 | -4.31229E-11 | 1.52621E-10 |
| | | Equal variances not assumed | | | 1.492 | 7.426 | 0.089 | 0.177 | 5.47492E-11 | 3.67045E-11 | -3.10433E-11 | 1.40542E-10 |
| | *N*-oleoyl taurine | Equal variances assumed | 0.914 | 0.356 | 0.591 | 13 | 0.282 | 0.565 | 1.79143E-10 | 3.03133E-10 | -4.75737E-10 | 8.34022E-10 |
| | | Equal variances not assumed | | | 0.578 | 10.785 | 0.288 | 0.575 | 1.79143E-10 | 3.10104E-10 | -5.05053E-10 | 8.63339E-10 |
| | *N*-arachidonoyl taurine | Equal variances assumed | 8.602 | 0.012 | 2.716 | 13 | 0.009 | 0.018 | 9.99030E-10 | 3.67833E-10 | 2.04374E-10 | 1.79369E-09 |
| | | Equal variances not assumed | | | 2.900 | 7.701 | 0.010 | 0.021 | 9.99030E-10 | 3.44508E-10 | 1.99193E-10 | 1.79887E-09 |
| CBD | *N*-palmitoyl taurine | Equal variances assumed | 0.202 | 0.659 | -0.373 | 16 | 0.357 | 0.714 | -2.51896E-10 | 6.74629E-10 | -1.68204E-09 | 1.17825E-09 |
| | | Equal variances | | | -0.383 | 15.999 | 0.353 | 0.706 | -2.51896E-10 | 6.57070E-10 | -1.64483E-09 | 1.14104E-09 |

| | | | | | | | | | | | | |
|---|---|---|---|---|---|---|---|---|---|---|---|---|
| | | not assumed | | | | | | | | | | |
| | *N*-stearoyl taurine | Equal variances assumed | 16.443 | 0.005 | 1.153 | 7 | 0.143 | 0.287 | 3.78477E-11 | 3.28290E-11 | -3.97806E-11 | 1.15476E-10 |
| | | Equal variances not assumed | | | 1.284 | 4.738 | 0.129 | 0.258 | 3.78477E-11 | 2.94675E-11 | -3.91791E-11 | 1.14874E-10 |
| | *N*-oleoyl taurine | Equal variances assumed | 0.405 | 0.533 | -0.056 | 16 | 0.478 | 0.956 | -1.13928E-11 | 2.04994E-10 | -4.45961E-10 | 4.23176E-10 |
| | | Equal variances not assumed | | | -0.054 | 12.325 | 0.479 | 0.958 | -1.13928E-11 | 2.12922E-10 | -4.73958E-10 | 4.51173E-10 |
| | *N*-arachidonoyl taurine | Equal variances assumed | 0.820 | 0.380 | -0.221 | 15 | 0.414 | 0.828 | -1.43503E-10 | 6.50744E-10 | -1.53053E-09 | 1.24352E-09 |
| | | Equal variances not assumed | | | -0.219 | 14.348 | 0.415 | 0.829 | -1.43503E-10 | 6.54344E-10 | -1.54374E-09 | 1.25673E-09 |
| | *N*-palmitoyl taurine | Equal variances assumed | 2.968 | 0.104 | 1.164 | 16 | 0.131 | 0.261 | 4.81784E-10 | 4.13747E-10 | -3.95320E-10 | 1.35889E-09 |
| | | Equal variances not assumed | | | 1.164 | 11.927 | 0.134 | 0.267 | 4.81784E-10 | 4.13747E-10 | -4.20306E-10 | 1.38388E-09 |
| THC | *N*-stearoyl taurine | Equal variances assumed | 0.147 | 0.708 | 3.298 | 12 | 0.003 | 0.006 | 1.30026E-10 | 3.94303E-11 | 4.41147E-11 | 2.15937E-10 |
| | | Equal variances not assumed | | | 2.915 | 4.486 | 0.019 | 0.038 | 1.30026E-10 | 4.46085E-11 | 1.12893E-11 | 2.48763E-10 |
| | *N*-oleoyl taurine | Equal variances assumed | 8.759 | 0.009 | 3.671 | 16 | 0.001 | 0.002 | 1.75563E-09 | 4.78251E-10 | 7.41784E-10 | 2.76948E-09 |

| | | | | | | | | | | | |
|---|---|---|---|---|---|---|---|---|---|---|---|
| | | Equal variances not assumed | | | 4.017 | 11.829 | 0.001 | 0.002 | 1.75563E-09 | 4.37074E-10 | 8.01796E-10 | 2.70947E-09 |
| | *N*-arachidonoyl taurine | Equal variances assumed | 0.111 | 0.743 | 0.809 | 17 | 0.215 | 0.430 | 6.07080E-10 | 7.50420E-10 | -9.76168E-10 | 2.19033E-09 |
| | | Equal variances not assumed | | | 0.814 | 16.999 | 0.214 | 0.427 | 6.07080E-10 | 7.46020E-10 | -9.66888E-10 | 2.18105E-09 |
| | *N*-palmitoyl taurine | Equal variances assumed | 0.002 | 0.969 | -0.882 | 18 | 0.195 | 0.390 | -4.20907E-10 | 4.77333E-10 | -1.42375E-09 | 5.81933E-10 |
| | | Equal variances not assumed | | | -0.880 | 17.115 | 0.195 | 0.391 | -4.20907E-10 | 4.78094E-10 | -1.42908E-09 | 5.87269E-10 |
| | *N*-stearoyl taurine | Equal variances assumed | 8.881 | 0.009 | 2.802 | 15 | 0.007 | 0.013 | 1.73794E-10 | 6.20248E-11 | 4.15916E-11 | 3.05997E-10 |
| | | Equal variances not assumed | | | 2.946 | 9.945 | 0.007 | 0.015 | 1.73794E-10 | 5.89941E-11 | 4.22482E-11 | 3.05340E-10 |
| THC+CBD | *N*-oleoyl taurine | Equal variances assumed | 0.066 | 0.800 | -1.595 | 17 | 0.065 | 0.129 | -5.48252E-10 | 3.43815E-10 | -1.27364E-09 | 1.77133E-10 |
| | | Equal variances not assumed | | | -1.604 | 16.999 | 0.064 | 0.127 | -5.48252E-10 | 3.41861E-10 | -1.26952E-09 | 1.73016E-10 |
| | *N*-arachidonoyl taurine | Equal variances assumed | 0.435 | 0.517 | 0.422 | 19 | 0.339 | 0.678 | 4.60131E-10 | 1.08993E-09 | -1.82113E-09 | 2.74139E-09 |
| | | Equal variances not assumed | | | 0.418 | 17.416 | 0.341 | 0.681 | 4.60131E-10 | 1.10111E-09 | -1.85878E-09 | 2.77904E-09 |

Supplemental Table 14.5 One-way ANOVA for *N*-acyl taurines measured in plasma collected from adult mice 2h after an i.p. injection of 10mg/kg CBD, THC, or THC+CBD

| | | Sum of Squares | df | Mean Square | F | Sig. |
|---|---|---|---|---|---|---|
| *N*-oleoyl taurine | Between Groups | 6.32E-16 | 7 | 9.02E-17 | 0.374 | 0.9082361 |
| | Within Groups | 5.55E-15 | 23 | 2.41E-16 | | |
| | Total | 6.18E-15 | 30 | | | |

Supplemental Table 14.6 One-way ANOVA for *N*-acyl taurines measured in liver collected from adult mice 2h after an i.p. injection of 10mg/kg CBD, THC, or THC+CBD

| | | Sum of Squares | df | Mean Square | F | Sig. |
|---|---|---|---|---|---|---|
| *N*-palmitoyl taurine | Between Groups | 2.48E-17 | 7 | 3.54E-18 | 3.194 | 0.00587 |
| | Within Groups | 6.98E-17 | 63 | 1.11E-18 | | |
| | Total | 9.46E-17 | 70 | | | |
| *N*-stearoyl taurine | Between Groups | 2.62E-19 | 7 | 3.74E-20 | 4.505 | 0.00081 |
| | Within Groups | 3.49E-19 | 42 | 8.31E-21 | | |
| | Total | 6.11E-19 | 49 | | | |
| *N*-oleoyl taurine | Between Groups | 4.01E-17 | 7 | 5.74E-18 | 10.700 | 0.00000 |
| | Within Groups | 3.32E-17 | 62 | 5.36E-19 | | |
| | Total | 7.34E-17 | 69 | | | |
| *N*-arachidonoyl taurine | Between Groups | 2.08E-16 | 7 | 2.97E-17 | 9.639 | 0.00000 |
| | Within Groups | 1.97E-16 | 64 | 3.08E-18 | | |
| | Total | 4.05E-16 | 71 | | | |

#### Supplemental Table 14.7 Fisher's LSD post-hoc test for plasma *N*-acyl taurine ANOVA

| Dependent Variable | | | Std. Error | Sig. | 95% Confidence Interval Lower Bound | Upper Bound |
|---|---|---|---|---|---|---|
| *N*-oleoyl taurine | CBDfemale | THCfemale | 9.83E-09 | 0.6926 | -1.64E-08 | 2.43E-08 |
| | | TpCfemale | 9.10E-09 | 0.4839 | -2.53E-08 | 1.23E-08 |
| | | TpCmale | 1.04E-08 | 0.8501 | -1.96E-08 | 2.36E-08 |
| | THCfemale | CBDfemale | 9.83E-09 | 0.6926 | -2.43E-08 | 1.64E-08 |
| | | TpCfemale | 9.10E-09 | 0.2645 | -2.92E-08 | 8.41E-09 |
| | | TpCmale | 1.04E-08 | 0.8539 | -2.35E-08 | 1.96E-08 |
| | TpCfemale | CBDfemale | 9.10E-09 | 0.4839 | -1.23E-08 | 2.53E-08 |
| | | THCfemale | 9.10E-09 | 0.2645 | -8.41E-09 | 2.92E-08 |
| | | TpCmale | 9.74E-09 | 0.3937 | -1.17E-08 | 2.86E-08 |
| | TpCmale | CBDfemale | 1.04E-08 | 0.8501 | -2.36E-08 | 1.96E-08 |
| | | THCfemale | 1.04E-08 | 0.8539 | -1.96E-08 | 2.35E-08 |
| | | TpCfemale | 9.74E-09 | 0.3937 | -2.86E-08 | 1.17E-08 |

#### Supplemental Table 14.8 Fisher's LSD post-hoc test for liver *N*-acyl taurine ANOVA

| Dependent Variable | | | Std. Error | Sig. | 95% Confidence Interval Lower Bound | Upper Bound |
|---|---|---|---|---|---|---|
| | VehFemale | CBDfemale | 5.26476E-10 | 0.472 | -6.7111E-10 | 1.4330E-09 |
| | | THCfemale | 5.11642E-10 | 0.025 | -2.1974E-09 | -1.5250E-10 |
| | | TpCfemale | 5.11642E-10 | 0.498 | -1.3714E-09 | 6.7343E-10 |
| | | VehMale | 5.44954E-10 | 0.168 | -3.2834E-10 | 1.8497E-09 |
| | | CBDmale | 4.99459E-10 | 0.797 | -8.6902E-10 | 1.1272E-09 |
| | | THCmale | 5.11642E-10 | 0.180 | -1.7156E-09 | 3.2929E-10 |
| | | TpCmale | 4.89265E-10 | 0.121 | -1.7476E-09 | 2.0780E-10 |
| | CBDfemale | VehFemale | 5.26476E-10 | 0.472 | -1.4330E-09 | 6.7111E-10 |
| | | THCfemale | 5.11642E-10 | 0.003 | -2.5783E-09 | -5.3346E-10 |
| | | TpCfemale | 5.11642E-10 | 0.159 | -1.7524E-09 | 2.9246E-10 |
| | | VehMale | 5.44954E-10 | 0.489 | -7.0931E-10 | 1.4687E-09 |
| | | CBDmale | 4.99459E-10 | 0.616 | -1.2500E-09 | 7.4619E-10 |
| | | THCmale | 5.11642E-10 | 0.040 | -2.0966E-09 | -5.1680E-11 |
| | | TpCmale | 4.89265E-10 | 0.022 | -2.1286E-09 | -1.7316E-10 |
| | THCfemale | VehFemale | 5.11642E-10 | 0.025 | 1.5250E-10 | 2.1974E-09 |
| | | CBDfemale | 5.11642E-10 | 0.003 | 5.3346E-10 | 2.5783E-09 |
| | | TpCfemale | 4.96366E-10 | 0.101 | -1.6598E-10 | 1.8178E-09 |
| | | VehMale | 5.30638E-10 | 0.001 | 8.7520E-10 | 2.9960E-09 |
| | | CBDmale | 4.83798E-10 | 0.009 | 3.3721E-10 | 2.2708E-09 |
| | | THCmale | 4.96366E-10 | 0.335 | -5.1012E-10 | 1.4737E-09 |
| | | TpCmale | 4.73266E-10 | 0.395 | -5.4073E-10 | 1.3508E-09 |
| | TpCfemale | VehFemale | 5.11642E-10 | 0.498 | -6.7343E-10 | 1.3714E-09 |
| | | CBDfemale | 5.11642E-10 | 0.159 | -2.9246E-10 | 1.7524E-09 |
| | | THCfemale | 4.96366E-10 | 0.101 | -1.8178E-09 | 1.6598E-10 |
| | | VehMale | 5.30638E-10 | 0.041 | 4.9276E-11 | 2.1701E-09 |
| | | CBDmale | 4.83798E-10 | 0.327 | -4.8872E-10 | 1.4449E-09 |
| | | THCmale | 4.96366E-10 | 0.491 | -1.3361E-09 | 6.4777E-10 |
| *N*-palmitoyl taurine | | TpCmale | 4.73266E-10 | 0.377 | -1.3667E-09 | 5.2484E-10 |
| | VehMale | VehFemale | 5.44954E-10 | 0.168 | -1.8497E-09 | 3.2834E-10 |
| | | CBDfemale | 5.44954E-10 | 0.489 | -1.4687E-09 | 7.0931E-10 |
| | | THCfemale | 5.30638E-10 | 0.001 | -2.9960E-09 | -8.7520E-10 |
| | | TpCfemale | 5.30638E-10 | 0.041 | -2.1701E-09 | -4.9276E-11 |
| | | CBDmale | 5.18900E-10 | 0.228 | -1.6685E-09 | 4.0535E-10 |
| | | THCmale | 5.30638E-10 | 0.008 | -2.5142E-09 | -3.9342E-10 |
| | | TpCmale | 5.09095E-10 | 0.004 | -2.5479E-09 | -5.1323E-10 |
| | CBDmale | VehFemale | 4.99459E-10 | 0.797 | -1.1272E-09 | 8.6902E-10 |
| | | CBDfemale | 4.99459E-10 | 0.616 | -7.4619E-10 | 1.2500E-09 |
| | | THCfemale | 4.83798E-10 | 0.009 | -2.2708E-09 | -3.3721E-10 |
| | | TpCfemale | 4.83798E-10 | 0.327 | -1.4449E-09 | 4.8872E-10 |
| | | VehMale | 5.18900E-10 | 0.228 | -4.0535E-10 | 1.6685E-09 |
| | | THCmale | 4.83798E-10 | 0.094 | -1.7890E-09 | 1.4457E-10 |
| | | TpCmale | 4.60068E-10 | 0.055 | -1.8184E-09 | 2.0388E-11 |
| | THCmale | VehFemale | 5.11642E-10 | 0.180 | -3.2929E-10 | 1.7156E-09 |
| | | CBDfemale | 5.11642E-10 | 0.040 | 5.1680E-11 | 2.0966E-09 |
| | | THCfemale | 4.96366E-10 | 0.335 | -1.4737E-09 | 5.1012E-10 |
| | | TpCfemale | 4.96366E-10 | 0.491 | -6.4777E-10 | 1.3361E-09 |
| | | VehMale | 5.30638E-10 | 0.008 | 3.9342E-10 | 2.5142E-09 |
| | | CBDmale | 4.83798E-10 | 0.094 | -1.4457E-10 | 1.7890E-09 |
| | | TpCmale | 4.73266E-10 | 0.872 | -1.0225E-09 | 8.6898E-10 |
| | TpCmale | VehFemale | 4.89265E-10 | 0.121 | -2.0780E-10 | 1.7476E-09 |
| | | CBDfemale | 4.89265E-10 | 0.022 | 1.7316E-10 | 2.1286E-09 |
| | | THCfemale | 4.73266E-10 | 0.395 | -1.3508E-09 | 5.4073E-10 |
| | | TpCfemale | 4.73266E-10 | 0.377 | -5.2484E-10 | 1.3667E-09 |
| | | VehMale | 5.09095E-10 | 0.004 | 5.1323E-10 | 2.5479E-09 |
| | | CBDmale | 4.60068E-10 | 0.055 | -2.0388E-11 | 1.8184E-09 |

| | | | | | | |
|---|---|---|---|---|---|---|
| | | THCmale | 4.73266E-10 | 0.872 | -8.6898E-10 | 1.0225E-09 |
| | VehFemale | CBDfemale | 5.88463E-11 | 0.960 | -1.2171E-10 | 1.1580E-10 |
| | | THCfemale | 4.70770E-11 | 0.042 | -1.9388E-10 | -3.8689E-12 |
| | | TpCfemale | 4.80478E-11 | 0.006 | -2.3711E-10 | -4.3180E-11 |
| | | VehMale | 5.88463E-11 | 0.357 | -6.4007E-11 | 1.7351E-10 |
| | | CBDmale | 5.52027E-11 | 0.531 | -7.6514E-11 | 1.4629E-10 |
| | | THCmale | 5.88463E-11 | 0.599 | -8.7605E-11 | 1.4991E-10 |
| | | TpCmale | 4.92343E-11 | 0.498 | -6.5709E-11 | 1.3301E-10 |
| | CBDfemale | VehFemale | 5.88463E-11 | 0.960 | -1.1580E-10 | 1.2171E-10 |
| | | THCfemale | 5.39335E-11 | 0.083 | -2.0476E-10 | 1.2926E-11 |
| | | TpCfemale | 5.47829E-11 | 0.016 | -2.4774E-10 | -2.6630E-11 |
| | | VehMale | 6.44629E-11 | 0.376 | -7.2384E-11 | 1.8780E-10 |
| | | CBDmale | 6.11549E-11 | 0.539 | -8.5568E-11 | 1.6126E-10 |
| | | THCmale | 6.44629E-11 | 0.599 | -9.5982E-11 | 1.6420E-10 |
| | | TpCmale | 5.58265E-11 | 0.516 | -7.6054E-11 | 1.4927E-10 |
| | THCfemale | VehFemale | 4.70770E-11 | 0.042 | 3.8689E-12 | 1.9388E-10 |
| | | CBDfemale | 5.39335E-11 | 0.083 | -1.2926E-11 | 2.0476E-10 |
| | | TpCfemale | 4.18871E-11 | 0.330 | -1.2580E-10 | 4.3261E-11 |
| | | VehMale | 5.39335E-11 | 0.007 | 4.4781E-11 | 2.6247E-10 |
| | | CBDmale | 4.99327E-11 | 0.011 | 3.2996E-11 | 2.3453E-10 |
| | | THCmale | 5.39335E-11 | 0.020 | 2.1184E-11 | 2.3887E-10 |
| | | TpCmale | 4.32430E-11 | 0.004 | 4.5256E-11 | 2.1979E-10 |
| | TpCfemale | VehFemale | 4.80478E-11 | 0.006 | 4.3180E-11 | 2.3711E-10 |
| | | CBDfemale | 5.47829E-11 | 0.016 | 2.6630E-11 | 2.4774E-10 |
| | | THCfemale | 4.18871E-11 | 0.330 | -4.3261E-11 | 1.2580E-10 |
| | | VehMale | 5.47829E-11 | 0.001 | 8.4337E-11 | 3.0545E-10 |
| | | CBDmale | 5.08490E-11 | 0.001 | 7.2417E-11 | 2.7765E-10 |
| | | THCmale | 5.47829E-11 | 0.003 | 6.0740E-11 | 2.8185E-10 |
| | | TpCmale | 4.42979E-11 | 0.000 | 8.4398E-11 | 2.6319E-10 |
| *N*-stearoyl taurine | VehMale | VehFemale | 5.88463E-11 | 0.357 | -1.7351E-10 | 6.4007E-11 |
| | | CBDfemale | 6.44629E-11 | 0.376 | -1.8780E-10 | 7.2384E-11 |
| | | THCfemale | 5.39335E-11 | 0.007 | -2.6247E-10 | -4.4781E-11 |
| | | TpCfemale | 5.47829E-11 | 0.001 | -3.0545E-10 | -8.4337E-11 |
| | | CBDmale | 6.11549E-11 | 0.747 | -1.4328E-10 | 1.0356E-10 |
| | | THCmale | 6.44629E-11 | 0.716 | -1.5369E-10 | 1.0649E-10 |
| | | TpCmale | 5.58265E-11 | 0.707 | -1.3376E-10 | 9.1563E-11 |
| | CBDmale | VehFemale | 5.52027E-11 | 0.531 | -1.4629E-10 | 7.6514E-11 |
| | | CBDfemale | 6.11549E-11 | 0.539 | -1.6126E-10 | 8.5568E-11 |
| | | THCfemale | 4.99327E-11 | 0.011 | -2.3453E-10 | -3.2996E-11 |
| | | TpCfemale | 5.08490E-11 | 0.001 | -2.7765E-10 | -7.2417E-11 |
| | | VehMale | 6.11549E-11 | 0.747 | -1.0356E-10 | 1.4328E-10 |
| | | THCmale | 6.11549E-11 | 0.952 | -1.2715E-10 | 1.1968E-10 |
| | | TpCmale | 5.19716E-11 | 0.981 | -1.0612E-10 | 1.0364E-10 |
| | THCmale | VehFemale | 5.88463E-11 | 0.599 | -1.4991E-10 | 8.7605E-11 |
| | | CBDfemale | 6.44629E-11 | 0.599 | -1.6420E-10 | 9.5982E-11 |
| | | THCfemale | 5.39335E-11 | 0.020 | -2.3887E-10 | -2.1184E-11 |
| | | TpCfemale | 5.47829E-11 | 0.003 | -2.8185E-10 | -6.0740E-11 |
| | | VehMale | 6.44629E-11 | 0.716 | -1.0649E-10 | 1.5369E-10 |
| | | CBDmale | 6.11549E-11 | 0.952 | -1.1968E-10 | 1.2715E-10 |
| | | TpCmale | 5.58265E-11 | 0.965 | -1.1016E-10 | 1.1516E-10 |
| | TpCmale | VehFemale | 4.92343E-11 | 0.498 | -1.3301E-10 | 6.5709E-11 |
| | | CBDfemale | 5.58265E-11 | 0.516 | -1.4927E-10 | 7.6054E-11 |
| | | THCfemale | 4.32430E-11 | 0.004 | -2.1979E-10 | -4.5256E-11 |
| | | TpCfemale | 4.42979E-11 | 0.000 | -2.6319E-10 | -8.4398E-11 |
| | | VehMale | 5.58265E-11 | 0.707 | -9.1563E-11 | 1.3376E-10 |
| | | CBDmale | 5.19716E-11 | 0.981 | -1.0364E-10 | 1.0612E-10 |
| | | THCmale | 5.58265E-11 | 0.965 | -1.1516E-10 | 1.1016E-10 |
| *N*-oleoyl taurine | VehFemale | CBDfemale | 3.66059E-10 | 0.368 | -3.9958E-10 | 1.0639E-09 |
| | | THCfemale | 3.47274E-10 | 0.000 | -2.3165E-09 | -9.2814E-10 |

| | | | | | | |
|---|---|---|---|---|---|---|
| | | TpCfemale | 3.55745E-10 | 0.033 | -1.4873E-09 | -6.5066E-11 |
| | | VehMale | 3.78907E-10 | 0.638 | -5.7828E-10 | 9.3657E-10 |
| | | CBDmale | 3.47274E-10 | 0.359 | -3.7343E-10 | 1.0150E-09 |
| | | THCmale | 3.66059E-10 | 0.717 | -5.9844E-10 | 8.6504E-10 |
| | | TpCmale | 3.47274E-10 | 0.000 | -2.0186E-09 | -6.3025E-10 |
| | CBDfemale | VehFemale | 3.66059E-10 | 0.368 | -1.0639E-09 | 3.9958E-10 |
| | | THCfemale | 3.47274E-10 | 0.000 | -2.6487E-09 | -1.2603E-09 |
| | | TpCfemale | 3.55745E-10 | 0.003 | -1.8195E-09 | -3.9722E-10 |
| | | VehMale | 3.78907E-10 | 0.688 | -9.1044E-10 | 6.0441E-10 |
| | | CBDmale | 3.47274E-10 | 0.974 | -7.0558E-10 | 6.8280E-10 |
| | | THCmale | 3.66059E-10 | 0.589 | -9.3059E-10 | 5.3289E-10 |
| | | TpCmale | 3.47274E-10 | 0.000 | -2.3508E-09 | -9.6241E-10 |
| | THCfemale | VehFemale | 3.47274E-10 | 0.000 | 9.2814E-10 | 2.3165E-09 |
| | | CBDfemale | 3.47274E-10 | 0.000 | 1.2603E-09 | 2.6487E-09 |
| | | TpCfemale | 3.36385E-10 | 0.014 | 1.7372E-10 | 1.5186E-09 |
| | | VehMale | 3.60791E-10 | 0.000 | 1.0803E-09 | 2.5227E-09 |
| | | CBDmale | 3.27413E-10 | 0.000 | 1.2886E-09 | 2.5976E-09 |
| | | THCmale | 3.47274E-10 | 0.000 | 1.0614E-09 | 2.4498E-09 |
| | | TpCmale | 3.27413E-10 | 0.366 | -3.5660E-10 | 9.5238E-10 |
| | TpCfemale | VehFemale | 3.55745E-10 | 0.033 | 6.5066E-11 | 1.4873E-09 |
| | | CBDfemale | 3.55745E-10 | 0.003 | 3.9722E-10 | 1.8195E-09 |
| | | THCfemale | 3.36385E-10 | 0.014 | -1.5186E-09 | -1.7372E-10 |
| | | VehMale | 3.68952E-10 | 0.012 | 2.1781E-10 | 1.6929E-09 |
| | | CBDmale | 3.36385E-10 | 0.002 | 4.2453E-10 | 1.7694E-09 |
| | | THCmale | 3.55745E-10 | 0.013 | 1.9837E-10 | 1.6206E-09 |
| | | TpCmale | 3.36385E-10 | 0.108 | -1.2207E-09 | 1.2417E-10 |
| | VehMale | VehFemale | 3.78907E-10 | 0.638 | -9.3657E-10 | 5.7828E-10 |
| | | CBDfemale | 3.78907E-10 | 0.688 | -6.0441E-10 | 9.1044E-10 |
| | | THCfemale | 3.60791E-10 | 0.000 | -2.5227E-09 | -1.0803E-09 |
| | | TpCfemale | 3.68952E-10 | 0.012 | -1.6929E-09 | -2.1781E-10 |
| | | CBDmale | 3.60791E-10 | 0.696 | -5.7959E-10 | 8.6283E-10 |
| | | THCmale | 3.78907E-10 | 0.904 | -8.0326E-10 | 7.1158E-10 |
| | | TpCmale | 3.60791E-10 | 0.000 | -2.2248E-09 | -7.8237E-10 |
| | CBDmale | VehFemale | 3.47274E-10 | 0.359 | -1.0150E-09 | 3.7343E-10 |
| | | CBDfemale | 3.47274E-10 | 0.974 | -6.8280E-10 | 7.0558E-10 |
| | | THCfemale | 3.27413E-10 | 0.000 | -2.5976E-09 | -1.2886E-09 |
| | | TpCfemale | 3.36385E-10 | 0.002 | -1.7694E-09 | -4.2453E-10 |
| | | VehMale | 3.60791E-10 | 0.696 | -8.6283E-10 | 5.7959E-10 |
| | | THCmale | 3.47274E-10 | 0.591 | -8.8165E-10 | 5.0673E-10 |
| | | TpCmale | 3.27413E-10 | 0.000 | -2.2997E-09 | -9.9072E-10 |
| | THCmale | VehFemale | 3.66059E-10 | 0.717 | -8.6504E-10 | 5.9844E-10 |
| | | CBDfemale | 3.66059E-10 | 0.589 | -5.3289E-10 | 9.3059E-10 |
| | | THCfemale | 3.47274E-10 | 0.000 | -2.4498E-09 | -1.0614E-09 |
| | | TpCfemale | 3.55745E-10 | 0.013 | -1.6206E-09 | -1.9837E-10 |
| | | VehMale | 3.78907E-10 | 0.904 | -7.1158E-10 | 8.0326E-10 |
| | | CBDmale | 3.47274E-10 | 0.591 | -5.0673E-10 | 8.8165E-10 |
| | | TpCmale | 3.47274E-10 | 0.000 | -2.1519E-09 | -7.6355E-10 |
| | TpCmale | VehFemale | 3.47274E-10 | 0.000 | 6.3025E-10 | 2.0186E-09 |
| | | CBDfemale | 3.47274E-10 | 0.000 | 9.6241E-10 | 2.3508E-09 |
| | | THCfemale | 3.27413E-10 | 0.366 | -9.5238E-10 | 3.5660E-10 |
| | | TpCfemale | 3.36385E-10 | 0.108 | -1.2417E-10 | 1.2207E-09 |
| | | VehMale | 3.60791E-10 | 0.000 | 7.8237E-10 | 2.2248E-09 |
| | | CBDmale | 3.27413E-10 | 0.000 | 9.9072E-10 | 2.2997E-09 |
| | | THCmale | 3.47274E-10 | 0.000 | 7.6355E-10 | 2.1519E-09 |
| *N*-arachidonoyl taurine | VehFemale | CBDfemale | 8.77332E-10 | 0.937 | -1.6831E-09 | 1.8222E-09 |
| | | THCfemale | 8.32310E-10 | 0.001 | -4.6995E-09 | -1.3740E-09 |
| | | TpCfemale | 8.32310E-10 | 0.000 | -5.3141E-09 | -1.9886E-09 |
| | | VehMale | 9.08125E-10 | 0.275 | -8.1516E-10 | 2.8132E-09 |
| | | CBDmale | 8.52613E-10 | 0.931 | -1.7773E-09 | 1.6293E-09 |

| | | | | | |
|---|---|---|---|---|---|
| | THCmale | 8.52613E-10 | 0.006 | -4.1330E-09 | -7.2637E-10 |
| | TpCmale | 8.15322E-10 | 0.000 | -4.8200E-09 | -1.5624E-09 |
| CBDfemale | VehFemale | 8.77332E-10 | 0.937 | -1.8222E-09 | 1.6831E-09 |
| | THCfemale | 8.32310E-10 | 0.000 | -4.7690E-09 | -1.4435E-09 |
| | TpCfemale | 8.32310E-10 | 0.000 | -5.3836E-09 | -2.0581E-09 |
| | VehMale | 9.08125E-10 | 0.310 | -8.8469E-10 | 2.7437E-09 |
| | CBDmale | 8.52613E-10 | 0.867 | -1.8468E-09 | 1.5598E-09 |
| | THCmale | 8.52613E-10 | 0.005 | -4.2025E-09 | -7.9590E-10 |
| | TpCmale | 8.15322E-10 | 0.000 | -4.8895E-09 | -1.6319E-09 |
| THCfemale | VehFemale | 8.32310E-10 | 0.001 | 1.3740E-09 | 4.6995E-09 |
| | CBDfemale | 8.32310E-10 | 0.000 | 1.4435E-09 | 4.7690E-09 |
| | TpCfemale | 7.84709E-10 | 0.436 | -2.1822E-09 | 9.5304E-10 |
| | VehMale | 8.64708E-10 | 0.000 | 2.3083E-09 | 5.7632E-09 |
| | CBDmale | 8.06212E-10 | 0.000 | 1.3522E-09 | 4.5734E-09 |
| | THCmale | 8.06212E-10 | 0.454 | -1.0035E-09 | 2.2177E-09 |
| | TpCmale | 7.66668E-10 | 0.841 | -1.6861E-09 | 1.3771E-09 |
| TpCfemale | VehFemale | 8.32310E-10 | 0.000 | 1.9886E-09 | 5.3141E-09 |
| | CBDfemale | 8.32310E-10 | 0.000 | 2.0581E-09 | 5.3836E-09 |
| | THCfemale | 7.84709E-10 | 0.436 | -9.5304E-10 | 2.1822E-09 |
| | VehMale | 8.64708E-10 | 0.000 | 2.9229E-09 | 6.3778E-09 |
| | CBDmale | 8.06212E-10 | 0.000 | 1.9668E-09 | 5.1880E-09 |
| | THCmale | 8.06212E-10 | 0.135 | -3.8892E-10 | 2.8323E-09 |
| | TpCmale | 7.66668E-10 | 0.551 | -1.0715E-09 | 1.9917E-09 |
| VehMale | VehFemale | 9.08125E-10 | 0.275 | -2.8132E-09 | 8.1516E-10 |
| | CBDfemale | 9.08125E-10 | 0.310 | -2.7437E-09 | 8.8469E-10 |
| | THCfemale | 8.64708E-10 | 0.000 | -5.7632E-09 | -2.3083E-09 |
| | TpCfemale | 8.64708E-10 | 0.000 | -6.3778E-09 | -2.9229E-09 |
| | CBDmale | 8.84267E-10 | 0.229 | -2.8395E-09 | 6.9352E-10 |
| | THCmale | 8.84267E-10 | 0.000 | -5.1952E-09 | -1.6622E-09 |
| | TpCmale | 8.48369E-10 | 0.000 | -5.8851E-09 | -2.4954E-09 |
| CBDmale | VehFemale | 8.52613E-10 | 0.931 | -1.6293E-09 | 1.7773E-09 |
| | CBDfemale | 8.52613E-10 | 0.867 | -1.5598E-09 | 1.8468E-09 |
| | THCfemale | 8.06212E-10 | 0.000 | -4.5734E-09 | -1.3522E-09 |
| | TpCfemale | 8.06212E-10 | 0.000 | -5.1880E-09 | -1.9668E-09 |
| | VehMale | 8.84267E-10 | 0.229 | -6.9352E-10 | 2.8395E-09 |
| | THCmale | 8.27156E-10 | 0.006 | -4.0081E-09 | -7.0326E-10 |
| | TpCmale | 7.88663E-10 | 0.000 | -4.6928E-09 | -1.5417E-09 |
| THCmale | VehFemale | 8.52613E-10 | 0.006 | 7.2637E-10 | 4.1330E-09 |
| | CBDfemale | 8.52613E-10 | 0.005 | 7.9590E-10 | 4.2025E-09 |
| | THCfemale | 8.06212E-10 | 0.454 | -2.2177E-09 | 1.0035E-09 |
| | TpCfemale | 8.06212E-10 | 0.135 | -2.8323E-09 | 3.8892E-10 |
| | VehMale | 8.84267E-10 | 0.000 | 1.6622E-09 | 5.1952E-09 |
| | CBDmale | 8.27156E-10 | 0.006 | 7.0326E-10 | 4.0081E-09 |
| | TpCmale | 7.88663E-10 | 0.338 | -2.3371E-09 | 8.1399E-10 |
| TpCmale | VehFemale | 8.15322E-10 | 0.000 | 1.5624E-09 | 4.8200E-09 |
| | CBDfemale | 8.15322E-10 | 0.000 | 1.6319E-09 | 4.8895E-09 |
| | THCfemale | 7.66668E-10 | 0.841 | -1.3771E-09 | 1.6861E-09 |
| | TpCfemale | 7.66668E-10 | 0.551 | -1.9917E-09 | 1.0715E-09 |
| | VehMale | 8.48369E-10 | 0.000 | 2.4954E-09 | 5.8851E-09 |
| | CBDmale | 7.88663E-10 | 0.000 | 1.5417E-09 | 4.6928E-09 |
| | THCmale | 7.88663E-10 | 0.338 | -8.1399E-10 | 2.3371E-09 |

Supplemental Table 15.1 Descriptive statistics for *N*-acyl GABAs (mol/L) in liver collected from male and female adult mice 2h after receiving an i.p. injection of 10mg/kg CBD, THC, or THC+CBD

| | | | | | | 95% Confidence Interval for Mean | | | |
|---|---|---|---|---|---|---|---|---|---|
| | | N | Mean | Std. Deviation | Std. Error | Lower Bound | Upper Bound | Minimum | Maximum |
| *N*-palmitoyl GABA | Veh F | 7 | 7.3104E-12 | 2.77857E-12 | 1.05020E-12 | 4.7406E-12 | 9.8801E-12 | 4.40E-12 | 1.25E-11 |
| | CBD F | 7 | 1.5110E-11 | 1.15979E-11 | 4.38358E-12 | 4.3841E-12 | 2.5837E-11 | 4.02E-12 | 3.35E-11 |
| | THC F | 9 | 9.7434E-12 | 4.62865E-12 | 1.54288E-12 | 6.1855E-12 | 1.3301E-11 | 2.35E-12 | 1.83E-11 |
| | THC+CBD F | 10 | 1.5452E-11 | 4.89721E-12 | 1.54863E-12 | 1.1949E-11 | 1.8955E-11 | 5.94E-12 | 2.03E-11 |
| | Veh M | 4 | 5.3495E-12 | 2.35379E-12 | 1.17690E-12 | 1.6041E-12 | 9.0949E-12 | 3.61E-12 | 8.83E-12 |
| | CBD M | 6 | 4.4265E-12 | 7.63619E-13 | 3.11746E-13 | 3.6251E-12 | 5.2279E-12 | 3.28E-12 | 5.30E-12 |
| | THC M | 9 | 1.5164E-11 | 7.11370E-12 | 2.37123E-12 | 9.6955E-12 | 2.0632E-11 | 5.49E-12 | 2.62E-11 |
| | THC+CBD M | 7 | 5.2784E-12 | 3.02697E-12 | 1.14409E-12 | 2.4790E-12 | 8.0779E-12 | 1.73E-12 | 9.36E-12 |
| *N*-stearoyl GABA | Veh F | 5 | 8.9903E-13 | 4.28788E-13 | 1.91760E-13 | 3.6662E-13 | 1.4314E-12 | 3.04E-13 | 1.51E-12 |
| | CBD F | 5 | 1.8272E-12 | 1.25847E-12 | 5.62805E-13 | 2.6462E-13 | 3.3898E-12 | 5.64E-13 | 3.86E-12 |
| | THC F | 5 | 1.6092E-12 | 1.34663E-12 | 6.02230E-13 | -6.2815E-14 | 3.2813E-12 | 3.48E-13 | 3.87E-12 |
| | THC+CBD F | 7 | 1.0818E-12 | 7.23537E-13 | 2.73471E-13 | 4.1262E-13 | 1.7509E-12 | 3.40E-13 | 2.46E-12 |
| | Veh M | 6 | 1.9498E-12 | 5.52356E-13 | 2.25498E-13 | 1.3702E-12 | 2.5295E-12 | 1.30E-12 | 2.73E-12 |
| | CBD M | 6 | 2.8189E-12 | 2.17049E-12 | 8.86100E-13 | 5.4109E-13 | 5.0967E-12 | 1.10E-12 | 6.15E-12 |
| | THC M | 7 | 1.9824E-12 | 8.39844E-13 | 3.17431E-13 | 1.2057E-12 | 2.7591E-12 | 1.14E-12 | 3.02E-12 |
| | THC+CBD M | 8 | 1.4456E-12 | 7.57606E-13 | 2.67854E-13 | 8.1221E-13 | 2.0790E-12 | 4.99E-13 | 2.96E-12 |
| *N*-oleoyl GABA | Veh F | 7 | 4.5984E-12 | 2.37178E-12 | 8.96450E-13 | 2.4048E-12 | 6.7919E-12 | 2.28E-12 | 9.58E-12 |
| | CBD F | 6 | 8.6198E-12 | 7.36769E-12 | 3.00785E-12 | 8.8785E-13 | 1.6352E-11 | 2.57E-12 | 2.00E-11 |
| | THC F | 10 | 8.6759E-12 | 5.47622E-12 | 1.73173E-12 | 4.7585E-12 | 1.2593E-11 | 2.01E-12 | 1.95E-11 |
| | THC+CBD F | 10 | 9.8337E-12 | 4.25230E-12 | 1.34470E-12 | 6.7918E-12 | 1.2876E-11 | 4.05E-12 | 1.77E-11 |
| | Veh M | 5 | 2.7652E-12 | 1.71767E-12 | 7.68165E-13 | 6.3240E-13 | 4.8979E-12 | 1.40E-12 | 5.59E-12 |
| | CBD M | 6 | 2.1884E-12 | 9.40157E-13 | 3.83817E-13 | 1.2018E-12 | 3.1750E-12 | 8.14E-13 | 3.52E-12 |
| | THC M | 8 | 5.5177E-12 | 3.08109E-12 | 1.08933E-12 | 2.9419E-12 | 8.0936E-12 | 2.65E-12 | 1.10E-11 |
| | THC+CBD M | 9 | 4.1133E-12 | 2.04191E-12 | 6.80636E-13 | 2.5438E-12 | 5.6829E-12 | 2.14E-12 | 7.87E-12 |
| *N*-linoleoyl GABA | Veh F | 8 | 1.9530E-10 | 1.31464E-10 | 4.64795E-11 | 8.5391E-11 | 3.0520E-10 | 6.83E-11 | 4.58E-10 |
| | CBD F | 5 | 6.9258E-10 | 4.92185E-10 | 2.20112E-10 | 8.1454E-11 | 1.3037E-09 | 5.78E-11 | 1.33E-09 |
| | THC F | 8 | 2.1678E-10 | 1.42391E-10 | 5.03428E-11 | 9.7742E-11 | 3.3583E-10 | 6.48E-11 | 4.58E-10 |
| | THC+CBD F | 10 | 5.2963E-10 | 2.63632E-10 | 8.33677E-11 | 3.4104E-10 | 7.1823E-10 | 1.13E-10 | 1.03E-09 |
| | Veh M | BAL | | | | | | | |
| | CBD M | 7 | 7.3175E-11 | 3.55761E-11 | 1.34465E-11 | 4.0272E-11 | 1.0608E-10 | 3.24E-11 | 1.30E-10 |
| | THC M | 9 | 4.0748E-10 | 2.28248E-10 | 7.60826E-11 | 2.3203E-10 | 5.8293E-10 | 7.78E-11 | 6.84E-10 |
| | THC+CBD M | 8 | 2.6677E-10 | 2.40695E-10 | 8.50985E-11 | 6.5547E-11 | 4.6800E-10 | 2.66E-11 | 7.37E-10 |
| *N*-arachidonoyl GABA | Veh F | 5 | 1.7797E-12 | 4.52972E-13 | 2.02575E-13 | 1.2172E-12 | 2.3421E-12 | 1.20E-12 | 2.39E-12 |
| | CBD F | 5 | 2.5928E-12 | 1.11787E-12 | 4.99925E-13 | 1.2048E-12 | 3.9809E-12 | 8.87E-13 | 3.80E-12 |
| | THC F | 5 | 2.4940E-12 | 3.93705E-13 | 1.76070E-13 | 2.0051E-12 | 2.9828E-12 | 2.14E-12 | 2.93E-12 |
| | THC+CBD F | 9 | 3.2983E-12 | 1.40887E-12 | 4.69624E-13 | 2.2154E-12 | 4.3813E-12 | 8.52E-13 | 5.47E-12 |
| | Veh M | BDL | | | | | | | |
| | CBD M | BAL | | | | | | | |
| | THC M | 7 | 3.3371E-12 | 2.33385E-12 | 8.82111E-13 | 1.1787E-12 | 5.4956E-12 | 1.21E-12 | 7.41E-12 |
| | THC+CBD M | 5 | 2.0313E-12 | 1.24829E-12 | 5.58252E-13 | 4.8133E-13 | 3.5812E-12 | 9.86E-13 | 4.09E-12 |
| *N*-docosahexaenoyl GABA | Veh F | 5 | 1.2485E-12 | 2.42350E-13 | 1.08382E-13 | 9.4755E-13 | 1.5494E-12 | 8.54E-13 | 1.46E-12 |

| | | | | | | | | | |
|---|---|---|---|---|---|---|---|---|---|
| | CBD F | 5 | 1.7282E-12 | 1.10261E-12 | 4.93102E-13 | 3.5911E-13 | 3.0973E-12 | 5.70E-13 | 3.26E-12 |
| | THC F | 8 | 2.8141E-12 | 1.65932E-12 | 5.86659E-13 | 1.4269E-12 | 4.2013E-12 | 8.07E-13 | 6.11E-12 |
| | THC+CBD F | 10 | 2.5385E-12 | 1.17087E-12 | 3.70261E-13 | 1.7009E-12 | 3.3761E-12 | 7.45E-13 | 4.73E-12 |
| | Veh M | BAL | | | | | | 7.80E-13 | 7.80E-13 |
| | CBD M | 4 | 1.1765E-12 | 5.92231E-13 | 2.96116E-13 | 2.3414E-13 | 2.1189E-12 | 4.67E-13 | 1.92E-12 |
| | THC M | 9 | 2.4914E-12 | 1.46287E-12 | 4.87624E-13 | 1.3669E-12 | 3.6158E-12 | 1.00E-12 | 5.21E-12 |
| | THC+CBD M | 9 | 1.5541E-12 | 1.05448E-12 | 3.51494E-13 | 7.4351E-13 | 2.3646E-12 | 4.44E-13 | 3.22E-12 |

Supplemental Table 15.2 Independent t-test comparing male and female levels of *N*-acyl GABAs measured in liver collected from adult mice 2h after an i.p. injection of 10mg/kg CBD, THC, or THC+CBD

| | | | Levene's Test for Equality of Variances | | t-test for Equality of Means | | | | | | | |
|---|---|---|---|---|---|---|---|---|---|---|---|---|
| | | | | | | | Significance | | | | 95% Confidence Interval of the Difference | |
| | | | F | Sig. | t | df | One-Sided p | Two-Sided p | Mean Difference | Std. Error Difference | Lower | Upper |
| Veh | *N*-palmitoyl GABA | Equal variances assumed | 0.054 | 0.822 | 1.183 | 9 | 0.134 | 0.267 | 1.96087E-12 | 1.65757E-12 | -1.78881E-12 | 5.71056E-12 |
| | | Equal variances not assumed | | | 1.243 | 7.350 | 0.126 | 0.252 | 1.96087E-12 | 1.57734E-12 | -1.73326E-12 | 5.65501E-12 |
| | *N*-stearoyl GABA | Equal variances assumed | 1.375 | 0.271 | -3.462 | 9 | 0.004 | 0.007 | -1.05081E-12 | 3.03499E-13 | -1.73737E-12 | -3.64248E-13 |
| | | Equal variances not assumed | | | -3.550 | 8.978 | 0.003 | 0.006 | -1.05081E-12 | 2.96009E-13 | -1.72068E-12 | -3.80938E-13 |
| | *N*-oleoyl GABA | Equal variances assumed | 0.062 | 0.808 | 1.467 | 10 | 0.087 | 0.173 | 1.83320E-12 | 1.24974E-12 | -9.51384E-13 | 4.61779E-12 |
| | | Equal variances not assumed | | | 1.553 | 9.977 | 0.076 | 0.152 | 1.83320E-12 | 1.18055E-12 | -7.98041E-13 | 4.46445E-12 |
| | *N*-linoleoyl GABA | Equal variances assumed | 2.420 | 0.158 | 0.624 | 8 | 0.275 | 0.550 | 6.07006E-11 | 9.72503E-11 | -1.63559E-10 | 2.84960E-10 |
| | | Equal variances not assumed | | | 1.294 | 7.239 | 0.118 | 0.235 | 6.07006E-11 | 4.68986E-11 | -4.94582E-11 | 1.70859E-10 |
| | *N*-docosahexaenoyl GABA | Equal variances assumed | | | 1.765 | 4 | 0.076 | 0.152 | 4.68674E-13 | 2.65481E-13 | -2.68420E-13 | 1.20577E-12 |
| | | Equal variances not | | | | | | | 4.68674E-13 | | | |

| | | | | | | | | | | | | |
|---|---|---|---|---|---|---|---|---|---|---|---|---|
| | | [continued] assumed | | | | | | | | | | |
| CBD | *N*-palmitoyl GABA | Equal variances assumed | 12.304 | 0.005 | 2.238 | 11 | 0.023 | 0.047 | 1.06838E-11 | 4.77406E-12 | 1.76182E-13 | 2.11914E-11 |
| | | Equal variances not assumed | | | 2.431 | 6.061 | 0.025 | 0.051 | 1.06838E-11 | 4.39465E-12 | -4.34857E-14 | 2.14111E-11 |
| | *N*-stearoyl GABA | Equal variances assumed | 2.467 | 0.151 | -0.899 | 9 | 0.196 | 0.392 | -9.91674E-13 | 1.10352E-12 | -3.48800E-12 | 1.50465E-12 |
| | | Equal variances not assumed | | | -0.945 | 8.183 | 0.186 | 0.372 | -9.91674E-13 | 1.04972E-12 | -3.40295E-12 | 1.41960E-12 |
| | *N*-oleoyl GABA | Equal variances assumed | 19.397 | 0.001 | 2.121 | 10 | 0.030 | 0.060 | 6.43137E-12 | 3.03224E-12 | -3.24870E-13 | 1.31876E-11 |
| | | Equal variances not assumed | | | 2.121 | 5.163 | 0.043 | 0.086 | 6.43137E-12 | 3.03224E-12 | -1.28989E-12 | 1.41526E-11 |
| | *N*-linoleoyl GABA | Equal variances assumed | 13.920 | 0.004 | 3.385 | 10 | 0.003 | 0.007 | 6.19408E-10 | 1.82983E-10 | 2.11697E-10 | 1.02712E-09 |
| | | Equal variances not assumed | | | 2.809 | 4.030 | 0.024 | 0.048 | 6.19408E-10 | 2.20522E-10 | 8.92595E-12 | 1.22989E-09 |
| | *N*-docosahexaenoyl GABA | Equal variances assumed | 2.061 | 0.194 | 0.895 | 7 | 0.200 | 0.401 | 5.51665E-13 | 6.16655E-13 | -9.06493E-13 | 2.00982E-12 |
| | | Equal variances not assumed | | | 0.959 | 6.311 | 0.186 | 0.373 | 5.51665E-13 | 5.75182E-13 | -8.39119E-13 | 1.94245E-12 |
| THC | *N*-palmitoyl GABA | Equal variances assumed | 2.809 | 0.113 | -1.916 | 16 | 0.037 | 0.073 | -5.42013E-12 | 2.82900E-12 | -1.14173E-11 | 5.77084E-13 |
| | | Equal varian[continued] | | | -1.916 | 13.744 | 0.038 | 0.076 | -5.42013E-12 | 2.82900E-12 | -1.14983E-11 | 6.58081E-13 |

| | | | | | | | | | | | | |
|---|---|---|---|---|---|---|---|---|---|---|---|---|
| | | ces not assumed | | | | | | | | | | |
| | *N*-stearoyl GABA | Equal variances assumed | 0.218 | 0.650 | -0.595 | 10 | 0.283 | 0.565 | -3.73134E-13 | 6.27530E-13 | -1.77136E-12 | 1.02509E-12 |
| | | Equal variances not assumed | | | -0.548 | 6.212 | 0.301 | 0.603 | -3.73134E-13 | 6.80767E-13 | -2.02524E-12 | 1.27898E-12 |
| | *N*-oleoyl GABA | Equal variances assumed | 3.171 | 0.094 | 1.452 | 16 | 0.083 | 0.166 | 3.15821E-12 | 2.17485E-12 | -1.45225E-12 | 7.76868E-12 |
| | | Equal variances not assumed | | | 1.544 | 14.594 | 0.072 | 0.144 | 3.15821E-12 | 2.04586E-12 | -1.21303E-12 | 7.52945E-12 |
| | *N*-linoleoyl GABA | Equal variances assumed | 4.661 | 0.047 | -2.033 | 15 | 0.030 | 0.060 | -1.90696E-10 | 9.37784E-11 | -3.90580E-10 | 9.18751E-12 |
| | | Equal variances not assumed | | | -2.090 | 13.567 | 0.028 | 0.056 | -1.90696E-10 | 9.12302E-11 | -3.86954E-10 | 5.56095E-12 |
| | *N*-arachidonoyl GABA | Equal variances assumed | 7.317 | 0.022 | -0.789 | 10 | 0.224 | 0.448 | -8.43174E-13 | 1.06853E-12 | -3.22400E-12 | 1.53765E-12 |
| | | Equal variances not assumed | | | -0.937 | 6.472 | 0.191 | 0.382 | -8.43174E-13 | 8.99511E-13 | -3.00585E-12 | 1.31950E-12 |
| | *N*-docosahexaenoyl GABA | Equal variances assumed | 0.002 | 0.962 | 0.426 | 15 | 0.338 | 0.676 | 3.22747E-13 | 7.56875E-13 | -1.29049E-12 | 1.93599E-12 |
| | | Equal variances not assumed | | | 0.423 | 14.117 | 0.339 | 0.679 | 3.22747E-13 | 7.62854E-13 | -1.31214E-12 | 1.95763E-12 |
| THC+CBD | *N*-palmitoyl GABA | Equal variances assumed | 1.891 | 0.189 | 4.859 | 15 | 0.000 | 0.000 | 1.01737E-11 | 2.09397E-12 | 5.71048E-12 | 1.46368E-11 |

| | | | | | | | | | | | |
|---|---|---|---|---|---|---|---|---|---|---|---|
| | Equal variances not assumed | | | 5.284 | 14.864 | 0.000 | 0.000 | 1.01737E-11 | 1.92541E-12 | 6.06647E-12 | 1.42809E-11 |
| *N*-stearoyl GABA | Equal variances assumed | 0.004 | 0.950 | -0.947 | 13 | 0.180 | 0.361 | -3.63801E-13 | 3.84061E-13 | -1.19351E-12 | 4.65912E-13 |
| | Equal variances not assumed | | | -0.950 | 12.876 | 0.180 | 0.359 | -3.63801E-13 | 3.82795E-13 | -1.19159E-12 | 4.63985E-13 |
| *N*-oleoyl GABA | Equal variances assumed | 5.287 | 0.034 | 3.666 | 17 | 0.001 | 0.002 | 5.72035E-12 | 1.56050E-12 | 2.42799E-12 | 9.01271E-12 |
| | Equal variances not assumed | | | 3.795 | 13.226 | 0.001 | 0.002 | 5.72035E-12 | 1.50714E-12 | 2.47001E-12 | 8.97069E-12 |
| *N*-linoleoyl GABA | Equal variances assumed | 0.150 | 0.704 | 2.183 | 16 | 0.022 | 0.044 | 2.62861E-10 | 1.20413E-10 | 7.59799E-12 | 5.18125E-10 |
| | Equal variances not assumed | | | 2.207 | 15.663 | 0.021 | 0.043 | 2.62861E-10 | 1.19130E-10 | 9.87484E-12 | 5.15848E-10 |
| *N*-arachidonoyl GABA | Equal variances assumed | 0.005 | 0.946 | 1.673 | 12 | 0.060 | 0.120 | 1.26705E-12 | 7.57153E-13 | -3.82648E-13 | 2.91674E-12 |
| | Equal variances not assumed | | | 1.737 | 9.329 | 0.058 | 0.115 | 1.26705E-12 | 7.29515E-13 | -3.74408E-13 | 2.90850E-12 |
| *N*-docosahexaenoyl GABA | Equal variances assumed | 0.036 | 0.852 | 1.917 | 17 | 0.036 | 0.072 | 9.84420E-13 | 5.13506E-13 | -9.89824E-14 | 2.06782E-12 |
| | Equal variances not assumed | | | 1.928 | 16.999 | 0.035 | 0.071 | 9.84420E-13 | 5.10530E-13 | -9.27085E-14 | 2.06155E-12 |

Supplemental Table 15.3 One-way ANOVA for *N*-acyl GABAs measured in liver collected from adult mice 2h after an i.p. injection of 10mg/kg CBD, THC, or THC+CBD

| | | Sum of Squares | df | Mean Square | F | Sig. |
|---|---|---|---|---|---|---|
| *N*-palmitoyl GABA | Between Groups | 1.18E-21 | 7 | 1.69E-22 | 5.017 | 0.00022 |
| | Within Groups | 1.72E-21 | 51 | 3.37E-23 | | |
| | Total | 2.90E-21 | 58 | | | |
| *N*-stearoyl GABA | Between Groups | 1.50E-23 | 7 | 2.14E-24 | 1.726 | 0.12978 |
| | Within Groups | 5.08E-23 | 41 | 1.24E-24 | | |
| | Total | 6.58E-23 | 48 | | | |
| *N*-oleoyl GABA | Between Groups | 4.45E-22 | 7 | 6.36E-23 | 3.945 | 0.00156 |
| | Within Groups | 8.54E-22 | 53 | 1.61E-23 | | |
| | Total | 1.30E-21 | 60 | | | |
| *N*-linoleoyl GABA | Between Groups | 1.85E-18 | 6 | 3.08E-19 | 5.509 | 0.00021 |
| | Within Groups | 2.69E-18 | 48 | 5.60E-20 | | |
| | Total | 4.54E-18 | 54 | | | |
| *N*-arachidonoyl GABA | Between Groups | 1.78E-23 | 6 | 2.97E-24 | 1.500 | 0.21078 |
| | Within Groups | 6.13E-23 | 31 | 1.98E-24 | | |
| | Total | 7.91E-23 | 37 | | | |
| *N*-docosahexaenoyl GABA | Between Groups | 1.78E-23 | 6 | 2.96E-24 | 1.996 | 0.08711 |
| | Within Groups | 6.38E-23 | 43 | 1.48E-24 | | |
| | Total | 8.15E-23 | 49 | | | |

#### Supplemental Table 15.4 Fisher's LSD post-hoc test for liver *N*-acyl GABAs ANOVA

| Dependent Variable | | | Std. Error | Sig. | 95% Confidence Interval Lower Bound | 95% Confidence Interval Upper Bound |
|---|---|---|---|---|---|---|
| *N*-palmitoyl GABA | VehFemale | CBDfemale | 3.10414E-12 | 0.015 | -1.4032E-11 | -1.5681E-12 |
| | | THCfemale | 2.92662E-12 | 0.410 | -8.3085E-12 | 3.4424E-12 |
| | | TpCfemale | 2.86188E-12 | 0.006 | -1.3887E-11 | -2.3963E-12 |
| | | VehMale | 3.63993E-12 | 0.592 | -5.3466E-12 | 9.2683E-12 |
| | | CBDmale | 3.23090E-12 | 0.376 | -3.6024E-12 | 9.3702E-12 |
| | | THCmale | 2.92662E-12 | 0.010 | -1.3729E-11 | -1.9778E-12 |
| | | TpCmale | 3.10414E-12 | 0.516 | -4.1999E-12 | 8.2638E-12 |
| | CBDfemale | VehFemale | 3.10414E-12 | 0.015 | 1.5681E-12 | 1.4032E-11 |
| | | THCfemale | 2.92662E-12 | 0.073 | -5.0854E-13 | 1.1242E-11 |
| | | TpCfemale | 2.86188E-12 | 0.905 | -6.0873E-12 | 5.4037E-12 |
| | | VehMale | 3.63993E-12 | 0.010 | 2.4534E-12 | 1.7068E-11 |
| | | CBDmale | 3.23090E-12 | 0.002 | 4.1975E-12 | 1.7170E-11 |
| | | THCmale | 2.92662E-12 | 0.986 | -5.9287E-12 | 5.8222E-12 |
| | | TpCmale | 3.10414E-12 | 0.003 | 3.6000E-12 | 1.6064E-11 |
| | THCfemale | VehFemale | 2.92662E-12 | 0.410 | -3.4424E-12 | 8.3085E-12 |
| | | CBDfemale | 2.92662E-12 | 0.073 | -1.1242E-11 | 5.0854E-13 |
| | | TpCfemale | 2.66828E-12 | 0.037 | -1.1065E-11 | -3.5188E-13 |
| | | VehMale | 3.48977E-12 | 0.214 | -2.6121E-12 | 1.1400E-11 |
| | | CBDmale | 3.06073E-12 | 0.088 | -8.2774E-13 | 1.1462E-11 |
| | | THCmale | 2.73760E-12 | 0.053 | -1.0916E-11 | 7.5831E-14 |
| | | TpCmale | 2.92662E-12 | 0.133 | -1.4104E-12 | 1.0340E-11 |
| | TpCfemale | VehFemale | 2.86188E-12 | 0.006 | 2.3963E-12 | 1.3887E-11 |
| | | CBDfemale | 2.86188E-12 | 0.905 | -5.4037E-12 | 6.0873E-12 |
| | | THCfemale | 2.66828E-12 | 0.037 | 3.5188E-13 | 1.1065E-11 |
| | | VehMale | 3.43566E-12 | 0.005 | 3.2052E-12 | 1.7000E-11 |
| | | CBDmale | 2.99889E-12 | 0.001 | 5.0051E-12 | 1.7046E-11 |
| | | THCmale | 2.66828E-12 | 0.914 | -5.0682E-12 | 5.6454E-12 |
| | | TpCmale | 2.86188E-12 | 0.001 | 4.4282E-12 | 1.5919E-11 |
| | VehMale | VehFemale | 3.63993E-12 | 0.592 | -9.2683E-12 | 5.3466E-12 |
| | | CBDfemale | 3.63993E-12 | 0.010 | -1.7068E-11 | -2.4534E-12 |
| | | THCfemale | 3.48977E-12 | 0.214 | -1.1400E-11 | 2.6121E-12 |
| | | TpCfemale | 3.43566E-12 | 0.005 | -1.7000E-11 | -3.2052E-12 |
| | | CBDmale | 3.74861E-12 | 0.806 | -6.6027E-12 | 8.4486E-12 |
| | | THCmale | 3.48977E-12 | 0.007 | -1.6820E-11 | -2.8081E-12 |
| | | TpCmale | 3.63993E-12 | 0.985 | -7.2364E-12 | 7.3785E-12 |
| | CBDmale | VehFemale | 3.23090E-12 | 0.376 | -9.3702E-12 | 3.6024E-12 |
| | | CBDfemale | 3.23090E-12 | 0.002 | -1.7170E-11 | -4.1975E-12 |
| | | THCfemale | 3.06073E-12 | 0.088 | -1.1462E-11 | 8.2774E-13 |
| | | TpCfemale | 2.99889E-12 | 0.001 | -1.7046E-11 | -5.0051E-12 |
| | | VehMale | 3.74861E-12 | 0.806 | -8.4486E-12 | 6.6027E-12 |
| | | THCmale | 3.06073E-12 | 0.001 | -1.6882E-11 | -4.5924E-12 |
| | | TpCmale | 3.23090E-12 | 0.793 | -7.3382E-12 | 5.6344E-12 |
| | THCmale | VehFemale | 2.92662E-12 | 0.010 | 1.9778E-12 | 1.3729E-11 |
| | | CBDfemale | 2.92662E-12 | 0.986 | -5.8222E-12 | 5.9287E-12 |
| | | THCfemale | 2.73760E-12 | 0.053 | -7.5831E-14 | 1.0916E-11 |
| | | TpCfemale | 2.66828E-12 | 0.914 | -5.6454E-12 | 5.0682E-12 |
| | | VehMale | 3.48977E-12 | 0.007 | 2.8081E-12 | 1.6820E-11 |
| | | CBDmale | 3.06073E-12 | 0.001 | 4.5924E-12 | 1.6882E-11 |
| | | TpCmale | 2.92662E-12 | 0.001 | 4.0097E-12 | 1.5761E-11 |
| | TpCmale | VehFemale | 3.10414E-12 | 0.516 | -8.2638E-12 | 4.1999E-12 |
| | | CBDfemale | 3.10414E-12 | 0.003 | -1.6064E-11 | -3.6000E-12 |
| | | THCfemale | 2.92662E-12 | 0.133 | -1.0340E-11 | 1.4104E-12 |
| | | TpCfemale | 2.86188E-12 | 0.001 | -1.5919E-11 | -4.4282E-12 |
| | | VehMale | 3.63993E-12 | 0.985 | -7.3785E-12 | 7.2364E-12 |
| | | CBDmale | 3.23090E-12 | 0.793 | -5.6344E-12 | 7.3382E-12 |

| | | | | | | |
|---|---|---|---|---|---|---|
| | | THCmale | 2.92662E-12 | 0.001 | -1.5761E-11 | -4.0097E-12 |
| *N*-stearoyl GABA | VehFemale | CBDfemale | 7.03965E-13 | 0.195 | -2.3499E-12 | 4.9351E-13 |
| | | THCfemale | 7.03965E-13 | 0.319 | -2.1319E-12 | 7.1148E-13 |
| | | TpCfemale | 6.51745E-13 | 0.781 | -1.4990E-12 | 1.1335E-12 |
| | | VehMale | 6.73995E-13 | 0.127 | -2.4120E-12 | 3.1035E-13 |
| | | CBDmale | 6.73995E-13 | 0.007 | -3.2810E-12 | -5.5869E-13 |
| | | THCmale | 6.51745E-13 | 0.104 | -2.3996E-12 | 2.3288E-13 |
| | | TpCmale | 6.34545E-13 | 0.394 | -1.8280E-12 | 7.3494E-13 |
| | CBDfemale | VehFemale | 7.03965E-13 | 0.195 | -4.9351E-13 | 2.3499E-12 |
| | | THCfemale | 7.03965E-13 | 0.758 | -1.2037E-12 | 1.6397E-12 |
| | | TpCfemale | 6.51745E-13 | 0.259 | -5.7079E-13 | 2.0617E-12 |
| | | VehMale | 6.73995E-13 | 0.857 | -1.4838E-12 | 1.2385E-12 |
| | | CBDmale | 6.73995E-13 | 0.149 | -2.3528E-12 | 3.6949E-13 |
| | | THCmale | 6.51745E-13 | 0.813 | -1.4714E-12 | 1.1611E-12 |
| | | TpCmale | 6.34545E-13 | 0.551 | -8.9986E-13 | 1.6631E-12 |
| | THCfemale | VehFemale | 7.03965E-13 | 0.319 | -7.1148E-13 | 2.1319E-12 |
| | | CBDfemale | 7.03965E-13 | 0.758 | -1.6397E-12 | 1.2037E-12 |
| | | TpCfemale | 6.51745E-13 | 0.423 | -7.8876E-13 | 1.8437E-12 |
| | | VehMale | 6.73995E-13 | 0.616 | -1.7018E-12 | 1.0206E-12 |
| | | CBDmale | 6.73995E-13 | 0.080 | -2.5708E-12 | 1.5152E-13 |
| | | THCmale | 6.51745E-13 | 0.570 | -1.6894E-12 | 9.4309E-13 |
| | | TpCmale | 6.34545E-13 | 0.798 | -1.1178E-12 | 1.4452E-12 |
| | TpCfemale | VehFemale | 6.51745E-13 | 0.781 | -1.1335E-12 | 1.4990E-12 |
| | | CBDfemale | 6.51745E-13 | 0.259 | -2.0617E-12 | 5.7079E-13 |
| | | THCfemale | 6.51745E-13 | 0.423 | -1.8437E-12 | 7.8876E-13 |
| | | VehMale | 6.19253E-13 | 0.169 | -2.1187E-12 | 3.8254E-13 |
| | | CBDmale | 6.19253E-13 | 0.008 | -2.9877E-12 | -4.8650E-13 |
| | | THCmale | 5.94959E-13 | 0.138 | -2.1021E-12 | 3.0095E-13 |
| | | TpCmale | 5.76066E-13 | 0.531 | -1.5272E-12 | 7.9959E-13 |
| | VehMale | VehFemale | 6.73995E-13 | 0.127 | -3.1035E-13 | 2.4120E-12 |
| | | CBDfemale | 6.73995E-13 | 0.857 | -1.2385E-12 | 1.4838E-12 |
| | | THCfemale | 6.73995E-13 | 0.616 | -1.0206E-12 | 1.7018E-12 |
| | | TpCfemale | 6.19253E-13 | 0.169 | -3.8254E-13 | 2.1187E-12 |
| | | CBDmale | 6.42629E-13 | 0.184 | -2.1669E-12 | 4.2877E-13 |
| | | THCmale | 6.19253E-13 | 0.958 | -1.2831E-12 | 1.2181E-12 |
| | | TpCmale | 6.01124E-13 | 0.406 | -7.0973E-13 | 1.7183E-12 |
| | CBDmale | VehFemale | 6.73995E-13 | 0.007 | 5.5869E-13 | 3.2810E-12 |
| | | CBDfemale | 6.73995E-13 | 0.149 | -3.6949E-13 | 2.3528E-12 |
| | | THCfemale | 6.73995E-13 | 0.080 | -1.5152E-13 | 2.5708E-12 |
| | | TpCfemale | 6.19253E-13 | 0.008 | 4.8650E-13 | 2.9877E-12 |
| | | VehMale | 6.42629E-13 | 0.184 | -4.2877E-13 | 2.1669E-12 |
| | | THCmale | 6.19253E-13 | 0.184 | -4.1410E-13 | 2.0871E-12 |
| | | TpCmale | 6.01124E-13 | 0.028 | 1.5931E-13 | 2.5873E-12 |
| | THCmale | VehFemale | 6.51745E-13 | 0.104 | -2.3288E-13 | 2.3996E-12 |
| | | CBDfemale | 6.51745E-13 | 0.813 | -1.1611E-12 | 1.4714E-12 |
| | | THCfemale | 6.51745E-13 | 0.570 | -9.4309E-13 | 1.6894E-12 |
| | | TpCfemale | 5.94959E-13 | 0.138 | -3.0095E-13 | 2.1021E-12 |
| | | VehMale | 6.19253E-13 | 0.958 | -1.2181E-12 | 1.2831E-12 |
| | | CBDmale | 6.19253E-13 | 0.184 | -2.0871E-12 | 4.1410E-13 |
| | | TpCmale | 5.76066E-13 | 0.357 | -6.2660E-13 | 1.7002E-12 |
| | TpCmale | VehFemale | 6.34545E-13 | 0.394 | -7.3494E-13 | 1.8280E-12 |
| | | CBDfemale | 6.34545E-13 | 0.551 | -1.6631E-12 | 8.9986E-13 |
| | | THCfemale | 6.34545E-13 | 0.798 | -1.4452E-12 | 1.1178E-12 |
| | | TpCfemale | 5.76066E-13 | 0.531 | -7.9959E-13 | 1.5272E-12 |
| | | VehMale | 6.01124E-13 | 0.406 | -1.7183E-12 | 7.0973E-13 |
| | | CBDmale | 6.01124E-13 | 0.028 | -2.5873E-12 | -1.5931E-13 |
| | | THCmale | 5.76066E-13 | 0.357 | -1.7002E-12 | 6.2660E-13 |
| *N*-oleoyl GABA | VehFemale | CBDfemale | 2.23304E-12 | 0.077 | -8.5003E-12 | 4.5750E-13 |
| | | THCfemale | 1.97799E-12 | 0.044 | -8.0449E-12 | -1.1020E-13 |

| | | | | | | |
|---|---|---|---|---|---|---|
| | | TpCfemale | 1.97799E-12 | 0.011 | -9.2027E-12 | -1.2680E-12 |
| | | VehMale | 2.35020E-12 | 0.439 | -2.8807E-12 | 6.5471E-12 |
| | | CBDmale | 2.23304E-12 | 0.285 | -2.0689E-12 | 6.8889E-12 |
| | | THCmale | 2.07730E-12 | 0.660 | -5.0859E-12 | 3.2472E-12 |
| | | TpCmale | 2.02273E-12 | 0.811 | -3.5721E-12 | 4.5421E-12 |
| | CBDfemale | VehFemale | 2.23304E-12 | 0.077 | -4.5750E-13 | 8.5003E-12 |
| | | THCfemale | 2.07268E-12 | 0.978 | -4.2134E-12 | 4.1011E-12 |
| | | TpCfemale | 2.07268E-12 | 0.561 | -5.3712E-12 | 2.9434E-12 |
| | | VehMale | 2.43044E-12 | 0.020 | 9.7976E-13 | 1.0729E-11 |
| | | CBDmale | 2.31733E-12 | 0.008 | 1.7834E-12 | 1.1079E-11 |
| | | THCmale | 2.16766E-12 | 0.158 | -1.2457E-12 | 7.4498E-12 |
| | | TpCmale | 2.11542E-12 | 0.038 | 2.6342E-13 | 8.7494E-12 |
| | THCfemale | VehFemale | 1.97799E-12 | 0.044 | 1.1020E-13 | 8.0449E-12 |
| | | CBDfemale | 2.07268E-12 | 0.978 | -4.1011E-12 | 4.2134E-12 |
| | | TpCfemale | 1.79500E-12 | 0.522 | -4.7581E-12 | 2.4425E-12 |
| | | VehMale | 2.19841E-12 | 0.010 | 1.5013E-12 | 1.0320E-11 |
| | | CBDmale | 2.07268E-12 | 0.003 | 2.3302E-12 | 1.0645E-11 |
| | | THCmale | 1.90388E-12 | 0.103 | -6.6049E-13 | 6.9769E-12 |
| | | TpCmale | 1.84418E-12 | 0.017 | 8.6361E-13 | 8.2615E-12 |
| | TpCfemale | VehFemale | 1.97799E-12 | 0.011 | 1.2680E-12 | 9.2027E-12 |
| | | CBDfemale | 2.07268E-12 | 0.561 | -2.9434E-12 | 5.3712E-12 |
| | | THCfemale | 1.79500E-12 | 0.522 | -2.4425E-12 | 4.7581E-12 |
| | | VehMale | 2.19841E-12 | 0.002 | 2.6591E-12 | 1.1478E-11 |
| | | CBDmale | 2.07268E-12 | 0.001 | 3.4880E-12 | 1.1803E-11 |
| | | THCmale | 1.90388E-12 | 0.027 | 4.9728E-13 | 8.1347E-12 |
| | | TpCmale | 1.84418E-12 | 0.003 | 2.0214E-12 | 9.4193E-12 |
| | VehMale | VehFemale | 2.35020E-12 | 0.439 | -6.5471E-12 | 2.8807E-12 |
| | | CBDfemale | 2.43044E-12 | 0.020 | -1.0729E-11 | -9.7976E-13 |
| | | THCfemale | 2.19841E-12 | 0.010 | -1.0320E-11 | -1.5013E-12 |
| | | TpCfemale | 2.19841E-12 | 0.002 | -1.1478E-11 | -2.6591E-12 |
| | | CBDmale | 2.43044E-12 | 0.813 | -4.2981E-12 | 5.4516E-12 |
| | | THCmale | 2.28818E-12 | 0.234 | -7.3420E-12 | 1.8370E-12 |
| | | TpCmale | 2.23875E-12 | 0.550 | -5.8385E-12 | 3.1422E-12 |
| | CBDmale | VehFemale | 2.23304E-12 | 0.285 | -6.8889E-12 | 2.0689E-12 |
| | | CBDfemale | 2.31733E-12 | 0.008 | -1.1079E-11 | -1.7834E-12 |
| | | THCfemale | 2.07268E-12 | 0.003 | -1.0645E-11 | -2.3302E-12 |
| | | TpCfemale | 2.07268E-12 | 0.001 | -1.1803E-11 | -3.4880E-12 |
| | | VehMale | 2.43044E-12 | 0.813 | -5.4516E-12 | 4.2981E-12 |
| | | THCmale | 2.16766E-12 | 0.131 | -7.6771E-12 | 1.0185E-12 |
| | | TpCmale | 2.11542E-12 | 0.367 | -6.1679E-12 | 2.3181E-12 |
| | THCmale | VehFemale | 2.07730E-12 | 0.660 | -3.2472E-12 | 5.0859E-12 |
| | | CBDfemale | 2.16766E-12 | 0.158 | -7.4498E-12 | 1.2457E-12 |
| | | THCfemale | 1.90388E-12 | 0.103 | -6.9769E-12 | 6.6049E-13 |
| | | TpCfemale | 1.90388E-12 | 0.027 | -8.1347E-12 | -4.9728E-13 |
| | | VehMale | 2.28818E-12 | 0.234 | -1.8370E-12 | 7.3420E-12 |
| | | CBDmale | 2.16766E-12 | 0.131 | -1.0185E-12 | 7.6771E-12 |
| | | TpCmale | 1.95032E-12 | 0.475 | -2.5075E-12 | 5.3162E-12 |
| | TpCmale | VehFemale | 2.02273E-12 | 0.811 | -4.5421E-12 | 3.5721E-12 |
| | | CBDfemale | 2.11542E-12 | 0.038 | -8.7494E-12 | -2.6342E-13 |
| | | THCfemale | 1.84418E-12 | 0.017 | -8.2615E-12 | -8.6361E-13 |
| | | TpCfemale | 1.84418E-12 | 0.003 | -9.4193E-12 | -2.0214E-12 |
| | | VehMale | 2.23875E-12 | 0.550 | -3.1422E-12 | 5.8385E-12 |
| | | CBDmale | 2.11542E-12 | 0.367 | -2.3181E-12 | 6.1679E-12 |
| | | THCmale | 1.95032E-12 | 0.475 | -5.3162E-12 | 2.5075E-12 |
| *N*-linoleoyl GABA | VehFemale | CBDfemale | 1.34890E-10 | 0.001 | -7.6850E-10 | -2.2607E-10 |
| | | THCfemale | 1.18307E-10 | 0.857 | -2.5936E-10 | 2.1639E-10 |
| | | TpCfemale | 1.12235E-10 | 0.005 | -5.6000E-10 | -1.0867E-10 |
| | | CBDmale | 1.22459E-10 | 0.324 | -1.2410E-10 | 3.6834E-10 |
| | | THCmale | 1.14973E-10 | 0.071 | -4.4335E-10 | 1.8987E-11 |

| | | | | | | |
|---|---|---|---|---|---|---|
| | | TpCmale | 1.18307E-10 | 0.549 | -3.0935E-10 | 1.6640E-10 |
| | CBDfemale | VehFemale | 1.34890E-10 | 0.001 | 2.2607E-10 | 7.6850E-10 |
| | | THCfemale | 1.34890E-10 | 0.001 | 2.0458E-10 | 7.4701E-10 |
| | | TpCfemale | 1.29598E-10 | 0.215 | -9.7627E-11 | 4.2352E-10 |
| | | CBDmale | 1.38546E-10 | 0.000 | 3.4084E-10 | 8.9797E-10 |
| | | THCmale | 1.31976E-10 | 0.036 | 1.9746E-11 | 5.5046E-10 |
| | | TpCmale | 1.34890E-10 | 0.003 | 1.5459E-10 | 6.9702E-10 |
| | THCfemale | VehFemale | 1.18307E-10 | 0.857 | -2.1639E-10 | 2.5936E-10 |
| | | CBDfemale | 1.34890E-10 | 0.001 | -7.4701E-10 | -2.0458E-10 |
| | | TpCfemale | 1.12235E-10 | 0.008 | -5.3852E-10 | -8.7186E-11 |
| | | CBDmale | 1.22459E-10 | 0.247 | -1.0261E-10 | 3.8983E-10 |
| | | THCmale | 1.14973E-10 | 0.104 | -4.2187E-10 | 4.0473E-11 |
| | | TpCmale | 1.18307E-10 | 0.675 | -2.8786E-10 | 1.8788E-10 |
| | TpCfemale | VehFemale | 1.12235E-10 | 0.005 | 1.0867E-10 | 5.6000E-10 |
| | | CBDfemale | 1.29598E-10 | 0.215 | -4.2352E-10 | 9.7627E-11 |
| | | THCfemale | 1.12235E-10 | 0.008 | 8.7186E-11 | 5.3852E-10 |
| | | CBDmale | 1.16604E-10 | 0.000 | 2.2201E-10 | 6.9091E-10 |
| | | THCmale | 1.08716E-10 | 0.267 | -9.6434E-11 | 3.4074E-10 |
| | | TpCmale | 1.12235E-10 | 0.023 | 3.7197E-11 | 4.8853E-10 |
| | CBDmale | VehFemale | 1.22459E-10 | 0.324 | -3.6834E-10 | 1.2410E-10 |
| | | CBDfemale | 1.38546E-10 | 0.000 | -8.9797E-10 | -3.4084E-10 |
| | | THCfemale | 1.22459E-10 | 0.247 | -3.8983E-10 | 1.0261E-10 |
| | | TpCfemale | 1.16604E-10 | 0.000 | -6.9091E-10 | -2.2201E-10 |
| | | THCmale | 1.19242E-10 | 0.007 | -5.7406E-10 | -9.4554E-11 |
| | | TpCmale | 1.22459E-10 | 0.120 | -4.3982E-10 | 5.2622E-11 |
| | THCmale | VehFemale | 1.14973E-10 | 0.071 | -1.8987E-11 | 4.4335E-10 |
| | | CBDfemale | 1.31976E-10 | 0.036 | -5.5046E-10 | -1.9746E-11 |
| | | THCfemale | 1.14973E-10 | 0.104 | -4.0473E-11 | 4.2187E-10 |
| | | TpCfemale | 1.08716E-10 | 0.267 | -3.4074E-10 | 9.6434E-11 |
| | | CBDmale | 1.19242E-10 | 0.007 | 9.4554E-11 | 5.7406E-10 |
| | | TpCmale | 1.14973E-10 | 0.227 | -9.0462E-11 | 3.7188E-10 |
| | TpCmale | VehFemale | 1.18307E-10 | 0.549 | -1.6640E-10 | 3.0935E-10 |
| | | CBDfemale | 1.34890E-10 | 0.003 | -6.9702E-10 | -1.5459E-10 |
| | | THCfemale | 1.18307E-10 | 0.675 | -1.8788E-10 | 2.8786E-10 |
| | | TpCfemale | 1.12235E-10 | 0.023 | -4.8853E-10 | -3.7197E-11 |
| | | CBDmale | 1.22459E-10 | 0.120 | -5.2622E-11 | 4.3982E-10 |
| | | THCmale | 1.14973E-10 | 0.227 | -3.7188E-10 | 9.0462E-11 |
| *N*-arachidonoyl GABA | VehFemale | CBDfemale | 8.89209E-13 | 0.368 | -2.6267E-12 | 1.0004E-12 |
| | | THCfemale | 8.89209E-13 | 0.428 | -2.5278E-12 | 1.0993E-12 |
| | | TpCfemale | 7.84209E-13 | 0.062 | -3.1181E-12 | 8.0743E-14 |
| | | THCmale | 8.23248E-13 | 0.068 | -3.2365E-12 | 1.2156E-13 |
| | | TpCmale | 8.89209E-13 | 0.779 | -2.0652E-12 | 1.5619E-12 |
| | CBDfemale | VehFemale | 8.89209E-13 | 0.368 | -1.0004E-12 | 2.6267E-12 |
| | | THCfemale | 8.89209E-13 | 0.912 | -1.7147E-12 | 1.9124E-12 |
| | | TpCfemale | 7.84209E-13 | 0.375 | -2.3049E-12 | 8.9391E-13 |
| | | THCmale | 8.23248E-13 | 0.373 | -2.4233E-12 | 9.3473E-13 |
| | | TpCmale | 8.89209E-13 | 0.532 | -1.2520E-12 | 2.3751E-12 |
| | THCfemale | VehFemale | 8.89209E-13 | 0.428 | -1.0993E-12 | 2.5278E-12 |
| | | CBDfemale | 8.89209E-13 | 0.912 | -1.9124E-12 | 1.7147E-12 |
| | | TpCfemale | 7.84209E-13 | 0.313 | -2.4038E-12 | 7.9504E-13 |
| | | THCmale | 8.23248E-13 | 0.314 | -2.5222E-12 | 8.3585E-13 |
| | | TpCmale | 8.89209E-13 | 0.607 | -1.3509E-12 | 2.2762E-12 |
| | TpCfemale | VehFemale | 7.84209E-13 | 0.062 | -8.0743E-14 | 3.1181E-12 |
| | | CBDfemale | 7.84209E-13 | 0.375 | -8.9391E-13 | 2.3049E-12 |
| | | THCfemale | 7.84209E-13 | 0.313 | -7.9504E-13 | 2.4038E-12 |
| | | THCmale | 7.08539E-13 | 0.957 | -1.4839E-12 | 1.4063E-12 |
| | | TpCmale | 7.84209E-13 | 0.116 | -3.3236E-13 | 2.8665E-12 |
| | THCmale | VehFemale | 8.23248E-13 | 0.068 | -1.2156E-13 | 3.2365E-12 |
| | | CBDfemale | 8.23248E-13 | 0.373 | -9.3473E-13 | 2.4233E-12 |

| | | | | | | |
|---|---|---|---|---|---|---|
| | | THCfemale | 8.23248E-13 | 0.314 | -8.3585E-13 | 2.5222E-12 |
| | | TpCfemale | 7.08539E-13 | 0.957 | -1.4063E-12 | 1.4839E-12 |
| | | TpCmale | 8.23248E-13 | 0.123 | -3.7317E-13 | 2.9849E-12 |
| | TpCmale | VehFemale | 8.89209E-13 | 0.779 | -1.5619E-12 | 2.0652E-12 |
| | | CBDfemale | 8.89209E-13 | 0.532 | -2.3751E-12 | 1.2520E-12 |
| | | THCfemale | 8.89209E-13 | 0.607 | -2.2762E-12 | 1.3509E-12 |
| | | TpCfemale | 7.84209E-13 | 0.116 | -2.8665E-12 | 3.3236E-13 |
| | | THCmale | 8.23248E-13 | 0.123 | -2.9849E-12 | 3.7317E-13 |
| *N*-docosahexaenoyl GABA | VehFemale | CBDfemale | 7.70245E-13 | 0.537 | -2.0331E-12 | 1.0736E-12 |
| | | THCfemale | 6.94290E-13 | 0.029 | -2.9658E-12 | -1.6546E-13 |
| | | TpCfemale | 6.67052E-13 | 0.060 | -2.6353E-12 | 5.5227E-14 |
| | | CBDmale | 8.16969E-13 | 0.930 | -1.5756E-12 | 1.7195E-12 |
| | | THCmale | 6.79293E-13 | 0.074 | -2.6128E-12 | 1.2704E-13 |
| | | TpCmale | 6.79293E-13 | 0.655 | -1.6755E-12 | 1.0643E-12 |
| | CBDfemale | VehFemale | 7.70245E-13 | 0.537 | -1.0736E-12 | 2.0331E-12 |
| | | THCfemale | 6.94290E-13 | 0.125 | -2.4861E-12 | 3.1425E-13 |
| | | TpCfemale | 6.67052E-13 | 0.231 | -2.1555E-12 | 5.3494E-13 |
| | | CBDmale | 8.16969E-13 | 0.503 | -1.0959E-12 | 2.1992E-12 |
| | | THCmale | 6.79293E-13 | 0.267 | -2.1331E-12 | 6.0675E-13 |
| | | TpCmale | 6.79293E-13 | 0.799 | -1.1958E-12 | 1.5440E-12 |
| | THCfemale | VehFemale | 6.94290E-13 | 0.029 | 1.6546E-13 | 2.9658E-12 |
| | | CBDfemale | 6.94290E-13 | 0.125 | -3.1425E-13 | 2.4861E-12 |
| | | TpCfemale | 5.77684E-13 | 0.636 | -8.8939E-13 | 1.4406E-12 |
| | | CBDmale | 7.45787E-13 | 0.034 | 1.3357E-13 | 3.1416E-12 |
| | | THCmale | 5.91776E-13 | 0.588 | -8.7068E-13 | 1.5162E-12 |
| | | TpCmale | 5.91776E-13 | 0.039 | 6.6611E-14 | 2.4535E-12 |
| | TpCfemale | VehFemale | 6.67052E-13 | 0.060 | -5.5227E-14 | 2.6353E-12 |
| | | CBDfemale | 6.67052E-13 | 0.231 | -5.3494E-13 | 2.1555E-12 |
| | | THCfemale | 5.77684E-13 | 0.636 | -1.4406E-12 | 8.8939E-13 |
| | | CBDmale | 7.20499E-13 | 0.065 | -9.1058E-14 | 2.8150E-12 |
| | | THCmale | 5.59570E-13 | 0.933 | -1.0814E-12 | 1.1756E-12 |
| | | TpCmale | 5.59570E-13 | 0.086 | -1.4406E-13 | 2.1129E-12 |
| | CBDmale | VehFemale | 8.16969E-13 | 0.930 | -1.7195E-12 | 1.5756E-12 |
| | | CBDfemale | 8.16969E-13 | 0.503 | -2.1992E-12 | 1.0959E-12 |
| | | THCfemale | 7.45787E-13 | 0.034 | -3.1416E-12 | -1.3357E-13 |
| | | TpCfemale | 7.20499E-13 | 0.065 | -2.8150E-12 | 9.1058E-14 |
| | | THCmale | 7.31846E-13 | 0.079 | -2.7907E-12 | 1.6107E-13 |
| | | TpCmale | 7.31846E-13 | 0.609 | -1.8535E-12 | 1.0984E-12 |
| | THCmale | VehFemale | 6.79293E-13 | 0.074 | -1.2704E-13 | 2.6128E-12 |
| | | CBDfemale | 6.79293E-13 | 0.267 | -6.0675E-13 | 2.1331E-12 |
| | | THCfemale | 5.91776E-13 | 0.588 | -1.5162E-12 | 8.7068E-13 |
| | | TpCfemale | 5.59570E-13 | 0.933 | -1.1756E-12 | 1.0814E-12 |
| | | CBDmale | 7.31846E-13 | 0.079 | -1.6107E-13 | 2.7907E-12 |
| | | TpCmale | 5.74107E-13 | 0.110 | -2.2050E-13 | 2.0951E-12 |
| | TpCmale | VehFemale | 6.79293E-13 | 0.655 | -1.0643E-12 | 1.6755E-12 |
| | | CBDfemale | 6.79293E-13 | 0.799 | -1.5440E-12 | 1.1958E-12 |
| | | THCfemale | 5.91776E-13 | 0.039 | -2.4535E-12 | -6.6611E-14 |
| | | TpCfemale | 5.59570E-13 | 0.086 | -2.1129E-12 | 1.4406E-13 |
| | | CBDmale | 7.31846E-13 | 0.609 | -1.0984E-12 | 1.8535E-12 |
| | | THCmale | 5.74107E-13 | 0.110 | -2.0951E-12 | 2.2050E-13 |

#### Supplemental Table 16.1 Descriptive statistics for *N*-acyl methionines (mol/g) in liver collected from male and female adult mice 2h after receiving an i.p. injection of 10mg/kg CBD, THC, or THC+CBD

| N | Mean | Std. Deviation | Std. Error | 95% Confidence Interval for Mean | Minimum | Maximum |
|---|---|---|---|---|---|---|

| | | | | | | Lower Bound | Upper Bound | | |
|---|---|---|---|---|---|---|---|---|---|
| *N*-palmitoyl methionine | Veh F | 8 | 5.3819E-11 | 2.62546E-11 | 9.28242E-12 | 3.1870E-11 | 7.5769E-11 | 2.19E-11 | 8.44E-11 |
| | CBD F | 7 | 5.8965E-11 | 3.47520E-11 | 1.31350E-11 | 2.6825E-11 | 9.1105E-11 | 1.33E-11 | 1.01E-10 |
| | THC F | 10 | 6.9220E-11 | 2.86804E-11 | 9.06954E-12 | 4.8703E-11 | 8.9737E-11 | 4.20E-11 | 1.26E-10 |
| | THC+CBD F | 9 | 8.9649E-11 | 2.57316E-11 | 8.57719E-12 | 6.9870E-11 | 1.0943E-10 | 3.50E-11 | 1.13E-10 |
| | Veh M | 7 | 3.4639E-11 | 1.28599E-11 | 4.86058E-12 | 2.2745E-11 | 4.6532E-11 | 1.78E-11 | 5.05E-11 |
| | CBD M | 10 | 3.0435E-11 | 1.00245E-11 | 3.17002E-12 | 2.3264E-11 | 3.7606E-11 | 1.31E-11 | 4.40E-11 |
| | THC M | 9 | 7.0110E-11 | 2.96303E-11 | 9.87678E-12 | 4.7334E-11 | 9.2886E-11 | 3.15E-11 | 1.14E-10 |
| | THC+CBD M | 10 | 2.8815E-11 | 1.70628E-11 | 5.39572E-12 | 1.6609E-11 | 4.1020E-11 | 1.27E-11 | 6.78E-11 |
| *N*-stearoyl methionine | Veh F | 8 | 3.8949E-12 | 2.26492E-12 | 8.00771E-13 | 2.0014E-12 | 5.7884E-12 | 1.43E-12 | 7.73E-12 |
| | CBD F | 6 | 8.9513E-12 | 5.78811E-12 | 2.36299E-12 | 2.8770E-12 | 1.5026E-11 | 4.24E-12 | 1.75E-11 |
| | THC F | 10 | 4.7194E-12 | 2.61916E-12 | 8.28251E-13 | 2.8458E-12 | 6.5930E-12 | 2.15E-12 | 9.90E-12 |
| | THC+CBD F | 10 | 7.5215E-12 | 2.63814E-12 | 8.34252E-13 | 5.6343E-12 | 9.4087E-12 | 3.55E-12 | 1.20E-11 |
| | Veh M | 7 | 3.4329E-12 | 1.88853E-12 | 7.13799E-13 | 1.6863E-12 | 5.1795E-12 | 1.32E-12 | 6.78E-12 |
| | CBD M | 4 | 2.7720E-12 | 7.92047E-13 | 3.96023E-13 | 1.5116E-12 | 4.0323E-12 | 1.60E-12 | 3.27E-12 |
| | THC M | 7 | 4.9284E-12 | 1.85440E-12 | 7.00899E-13 | 3.2133E-12 | 6.6434E-12 | 2.04E-12 | 7.44E-12 |
| | THC+CBD M | 8 | 2.4388E-12 | 1.51548E-12 | 5.35803E-13 | 1.1718E-12 | 3.7058E-12 | 1.23E-12 | 5.40E-12 |
| *N*-oleoyl methionine | Veh F | 8 | 5.5329E-11 | 2.88185E-11 | 1.01889E-11 | 3.1236E-11 | 7.9422E-11 | 2.99E-11 | 1.04E-10 |
| | CBD F | 7 | 5.4200E-11 | 1.92155E-11 | 7.26279E-12 | 3.6429E-11 | 7.1972E-11 | 1.88E-11 | 7.53E-11 |
| | THC F | 10 | 9.6074E-11 | 4.07651E-11 | 1.28911E-11 | 6.6913E-11 | 1.2524E-10 | 4.55E-11 | 1.75E-10 |
| | THC+CBD F | 9 | 1.0590E-10 | 2.06322E-11 | 6.87739E-12 | 9.0042E-11 | 1.2176E-10 | 6.16E-11 | 1.30E-10 |
| | Veh M | 7 | 3.7227E-11 | 1.13381E-11 | 4.28541E-12 | 2.6741E-11 | 4.7713E-11 | 2.42E-11 | 5.24E-11 |
| | CBD M | 10 | 3.8907E-11 | 1.30860E-11 | 4.13815E-12 | 2.9546E-11 | 4.8269E-11 | 2.21E-11 | 6.14E-11 |
| | THC M | 8 | 6.0247E-11 | 2.25059E-11 | 7.95705E-12 | 4.1432E-11 | 7.9063E-11 | 3.23E-11 | 9.78E-11 |
| | THC+CBD M | 10 | 4.3078E-11 | 1.93921E-11 | 6.13234E-12 | 2.9205E-11 | 5.6950E-11 | 2.04E-11 | 9.06E-11 |
| *N*-linoleoyl methionine | Veh F | 8 | 3.8966E-11 | 2.02536E-11 | 7.16074E-12 | 2.2033E-11 | 5.5898E-11 | 1.08E-11 | 7.30E-11 |
| | CBD F | 7 | 5.3406E-11 | 4.37111E-11 | 1.65212E-11 | 1.2980E-11 | 9.3832E-11 | 2.42E-12 | 9.71E-11 |
| | THC F | 9 | 5.0141E-11 | 2.03080E-11 | 6.76933E-12 | 3.4531E-11 | 6.5751E-11 | 2.44E-11 | 7.89E-11 |
| | THC+CBD F | 9 | 1.0216E-10 | 3.44266E-11 | 1.14755E-11 | 7.5696E-11 | 1.2862E-10 | 4.74E-11 | 1.44E-10 |
| | Veh M | 7 | 1.9781E-11 | 1.07574E-11 | 4.06590E-12 | 9.8317E-12 | 2.9730E-11 | 6.63E-12 | 3.70E-11 |
| | CBD M | 9 | 2.0955E-11 | 5.26971E-12 | 1.75657E-12 | 1.6905E-11 | 2.5006E-11 | 1.43E-11 | 3.05E-11 |
| | THC M | 9 | 5.5632E-11 | 2.77896E-11 | 9.26321E-12 | 3.4271E-11 | 7.6993E-11 | 1.86E-11 | 9.63E-11 |
| | THC+CBD M | 10 | 2.5354E-11 | 1.73987E-11 | 5.50194E-12 | 1.2907E-11 | 3.7800E-11 | 1.15E-11 | 6.19E-11 |
| *N*-arachidonoyl methionine | Veh F | 8 | 3.8280E-11 | 2.34639E-11 | 8.29573E-12 | 1.8664E-11 | 5.7897E-11 | 9.66E-12 | 6.98E-11 |
| | CBD F | 7 | 4.8857E-11 | 3.04386E-11 | 1.15047E-11 | 2.0706E-11 | 7.7008E-11 | 4.61E-12 | 9.13E-11 |
| | THC F | 10 | 4.9524E-11 | 1.80298E-11 | 5.70151E-12 | 3.6626E-11 | 6.2422E-11 | 2.98E-11 | 7.88E-11 |
| | THC+CBD F | 9 | 6.3954E-11 | 1.63050E-11 | 5.43499E-12 | 5.1420E-11 | 7.6487E-11 | 3.46E-11 | 9.31E-11 |
| | Veh M | 7 | 2.8493E-11 | 1.62683E-11 | 6.14883E-12 | 1.3447E-11 | 4.3539E-11 | 5.77E-12 | 5.28E-11 |
| | CBD M | 10 | 2.5233E-11 | 1.23595E-11 | 3.90841E-12 | 1.6391E-11 | 3.4074E-11 | 1.32E-11 | 4.72E-11 |
| | THC M | 9 | 4.7462E-11 | 2.15348E-11 | 7.17827E-12 | 3.0909E-11 | 6.4015E-11 | 1.26E-11 | 7.63E-11 |
| | THC+CBD M | 11 | 2.8569E-11 | 2.27673E-11 | 6.86461E-12 | 1.3274E-11 | 4.3864E-11 | 7.23E-12 | 7.08E-11 |
| *N*-docosahexaenoyl methionine | Veh F | 8 | 1.9372E-11 | 1.36547E-11 | 4.82765E-12 | 7.9569E-12 | 3.0788E-11 | 4.41E-12 | 3.84E-11 |
| | CBD F | 7 | 2.5188E-11 | 2.22901E-11 | 8.42485E-12 | 4.5732E-12 | 4.5803E-11 | 2.29E-12 | 6.37E-11 |
| | THC F | 9 | 2.7145E-11 | 1.12860E-11 | 3.76201E-12 | 1.8469E-11 | 3.5820E-11 | 8.26E-12 | 4.15E-11 |
| | THC+CBD F | 9 | 4.3194E-11 | 1.70195E-11 | 5.67317E-12 | 3.0112E-11 | 5.6276E-11 | 2.00E-11 | 6.93E-11 |
| | Veh M | 7 | 9.9292E-12 | 6.33402E-12 | 2.39404E-12 | 4.0713E-12 | 1.5787E-11 | 3.56E-12 | 2.23E-11 |
| | CBD M | 9 | 1.0216E-11 | 5.00926E-12 | 1.66975E-12 | 6.3653E-12 | 1.4066E-11 | 3.06E-12 | 1.72E-11 |

| | | | | | | | | | |
|---|---|---|---|---|---|---|---|---|---|
| | THC M | 9 | 3.1757E-11 | 1.57360E-11 | 5.24535E-12 | 1.9661E-11 | 4.3853E-11 | 7.95E-12 | 5.86E-11 |
| | THC+CBD M | 11 | 2.3511E-11 | 2.49284E-11 | 7.51619E-12 | 6.7641E-12 | 4.0258E-11 | 2.54E-12 | 7.15E-11 |

Supplemental Table 16.2 Independent t-test comparing male and female levels of *N*-acyl methionines measured in liver collected from adult mice 2h after an i.p. injection of 10mg/kg CBD, THC, or THC+CBD

| | | | Levene's Test for Equality of Variances | | t-test for Equality of Means | | | | | | | |
|---|---|---|---|---|---|---|---|---|---|---|---|---|
| | | | | | | | Significance | | | | 95% Confidence Interval of the Difference | |
| | | | F | Sig. | t | df | One-Sided p | Two-Sided p | Mean Difference | Std. Error Difference | Lower | Upper |
| Veh | *N*-palmitoyl methionine | Equal variances assumed | 10.309 | 0.007 | 1.752 | 13 | 0.052 | 0.103 | 1.91805E-11 | 1.09482E-11 | -4.47173E-12 | 4.28328E-11 |
| | | Equal variances not assumed | | | 1.831 | 10.449 | 0.048 | 0.096 | 1.91805E-11 | 1.04780E-11 | -4.03079E-12 | 4.23918E-11 |
| | *N*-stearoyl methionine | Equal variances assumed | 0.224 | 0.644 | 0.425 | 13 | 0.339 | 0.678 | 4.62023E-13 | 1.08665E-12 | -1.88554E-12 | 2.80959E-12 |
| | | Equal variances not assumed | | | 0.431 | 12.982 | 0.337 | 0.674 | 4.62023E-13 | 1.07273E-12 | -1.85580E-12 | 2.77984E-12 |
| | *N*-oleoyl methionine | Equal variances assumed | 5.530 | 0.035 | 1.554 | 13 | 0.072 | 0.144 | 1.81022E-11 | 1.16481E-11 | -7.06191E-12 | 4.32663E-11 |
| | | Equal variances not assumed | | | 1.638 | 9.354 | 0.067 | 0.135 | 1.81022E-11 | 1.10534E-11 | -6.75882E-12 | 4.29632E-11 |
| | *N*-linoleoyl methionine | Equal variances assumed | 2.477 | 0.140 | 2.238 | 13 | 0.022 | 0.043 | 1.91851E-11 | 8.57153E-12 | 6.67469E-13 | 3.77028E-11 |
| | | Equal variances not assumed | | | 2.330 | 10.917 | 0.020 | 0.040 | 1.91851E-11 | 8.23455E-12 | 1.04424E-12 | 3.73260E-11 |
| | *N*-arachidonoyl methionine | Equal variances assumed | 2.276 | 0.155 | 0.924 | 13 | 0.186 | 0.372 | 9.78714E-12 | 1.05889E-11 | -1.30888E-11 | 3.26631E-11 |
| | | Equal variances not assumed | | | 0.948 | 12.428 | 0.181 | 0.361 | 9.78714E-12 | 1.03261E-11 | -1.26258E-11 | 3.22001E-11 |
| | *N*-docosahexaenoyl methionine | Equal variances assumed | 5.678 | 0.033 | 1.673 | 13 | 0.059 | 0.118 | 9.44320E-12 | 5.64373E-12 | -2.74933E-12 | 2.16357E-11 |
| | | Equal variances not assumed | | | 1.752 | 10.150 | 0.055 | 0.110 | 9.44320E-12 | 5.38866E-12 | -2.53946E-12 | 2.14259E-11 |
| CBD | *N*-palmitoyl methionine | Equal variances assumed | 21.660 | 0.000 | 2.484 | 15 | 0.013 | 0.025 | 2.85301E-11 | 1.14875E-11 | 4.04504E-12 | 5.30151E-11 |
| | | Equal variances not assumed | | | 2.111 | 6.704 | 0.037 | 0.074 | 2.85301E-11 | 1.35121E-11 | -3.70917E-12 | 6.07693E-11 |
| | *N*-stearoyl methionine | Equal variances assumed | 13.690 | 0.006 | 2.080 | 8 | 0.036 | 0.071 | 6.17931E-12 | 2.97028E-12 | -6.70167E-13 | 1.30288E-11 |

| | | | | | | | | | | | | |
|---|---|---|---|---|---|---|---|---|---|---|---|---|
| | | Equal variances not assumed | | | 2.579 | 5.278 | 0.024 | 0.047 | 6.17931E-12 | 2.39594E-12 | 1.16612E-13 | 1.22420E-11 |
| | *N*-oleoyl methionine | Equal variances assumed | 1.048 | 0.322 | 1.961 | 15 | 0.034 | 0.069 | 1.52930E-11 | 7.79880E-12 | -1.32974E-12 | 3.19157E-11 |
| | | Equal variances not assumed | | | 1.830 | 9.837 | 0.049 | 0.098 | 1.52930E-11 | 8.35897E-12 | -3.37388E-12 | 3.39599E-11 |
| | *N*-linoleoyl methionine | Equal variances assumed | 94.098 | 0.000 | 2.229 | 14 | 0.021 | 0.043 | 3.24506E-11 | 1.45600E-11 | 1.22253E-12 | 6.36786E-11 |
| | | Equal variances not assumed | | | 1.953 | 6.136 | 0.049 | 0.098 | 3.24506E-11 | 1.66143E-11 | -7.98612E-12 | 7.28872E-11 |
| | *N*-arachidonoyl methionine | Equal variances assumed | 9.506 | 0.008 | 2.230 | 15 | 0.021 | 0.041 | 2.36246E-11 | 1.05954E-11 | 1.04103E-12 | 4.62082E-11 |
| | | Equal variances not assumed | | | 1.944 | 7.399 | 0.045 | 0.091 | 2.36246E-11 | 1.21505E-11 | -4.79549E-12 | 5.20447E-11 |
| | *N*-docosahexaenoyl methionine | Equal variances assumed | 11.420 | 0.004 | 1.971 | 14 | 0.034 | 0.069 | 1.49723E-11 | 7.59738E-12 | -1.32248E-12 | 3.12670E-11 |
| | | Equal variances not assumed | | | 1.743 | 6.473 | 0.064 | 0.128 | 1.49723E-11 | 8.58873E-12 | -5.67678E-12 | 3.56213E-11 |
| | *N*-palmitoyl methionine | Equal variances assumed | 0.227 | 0.639 | -0.066 | 17 | 0.474 | 0.948 | -8.89790E-13 | 1.33849E-11 | -2.91295E-11 | 2.73499E-11 |
| | | Equal variances not assumed | | | -0.066 | 16.654 | 0.474 | 0.948 | -8.89790E-13 | 1.34092E-11 | -2.92256E-11 | 2.74460E-11 |
| | *N*-stearoyl methionine | Equal variances assumed | 1.524 | 0.236 | -0.181 | 15 | 0.429 | 0.859 | -2.08986E-13 | 1.15484E-12 | -2.67047E-12 | 2.25250E-12 |
| | | Equal variances not assumed | | | -0.193 | 14.981 | 0.425 | 0.850 | -2.08986E-13 | 1.08502E-12 | -2.52189E-12 | 2.10392E-12 |
| THC | *N*-oleoyl methionine | Equal variances assumed | 2.620 | 0.125 | 2.221 | 16 | 0.021 | 0.041 | 3.58272E-11 | 1.61301E-11 | 1.63282E-12 | 7.00215E-11 |
| | | Equal variances not assumed | | | 2.365 | 14.465 | 0.016 | 0.032 | 3.58272E-11 | 1.51491E-11 | 3.43334E-12 | 6.82210E-11 |
| | *N*-linoleoyl methionine | Equal variances assumed | 1.607 | 0.223 | -0.479 | 16 | 0.319 | 0.639 | -5.49073E-12 | 1.14731E-11 | -2.98125E-11 | 1.88311E-11 |
| | | Equal variances not assumed | | | -0.479 | 14.648 | 0.320 | 0.639 | -5.49073E-12 | 1.14731E-11 | -2.99962E-11 | 1.90147E-11 |
| | *N*-arachidonoyl methionine | Equal variances assumed | 0.843 | 0.372 | 0.227 | 17 | 0.412 | 0.823 | 2.06174E-12 | 9.07762E-12 | -1.70904E-11 | 2.12139E-11 |
| | | Equal variances | | | 0.225 | 15.717 | 0.412 | 0.825 | 2.06174E-12 | 9.16705E-12 | -1.74000E-11 | 2.15234E-11 |

| | | | | | | | | | | | |
|---|---|---|---|---|---|---|---|---|---|---|---|
| | | not assumed | | | | | | | | | |
| | *N*-docosahexaenoyl methionine | Equal variances assumed | 0.649 | 0.432 | -0.715 | 16 | 0.243 | 0.485 | -4.61231E-12 | 6.45495E-12 | -1.82962E-11 | 9.07157E-12 |
| | | Equal variances not assumed | | | -0.715 | 14.508 | 0.243 | 0.486 | -4.61231E-12 | 6.45495E-12 | -1.84114E-11 | 9.18682E-12 |
| THC+CBD | *N*-palmitoyl methionine | Equal variances assumed | 1.464 | 0.243 | 6.135 | 17 | 0.000 | 0.000 | 6.08348E-11 | 9.91552E-12 | 3.99149E-11 | 8.17547E-11 |
| | | Equal variances not assumed | | | 6.004 | 13.680 | 0.000 | 0.000 | 6.08348E-11 | 1.01332E-11 | 3.90535E-11 | 8.26161E-11 |
| | *N*-stearoyl methionine | Equal variances assumed | 2.447 | 0.137 | 4.831 | 16 | 0.000 | 0.000 | 5.08273E-12 | 1.05210E-12 | 2.85237E-12 | 7.31309E-12 |
| | | Equal variances not assumed | | | 5.126 | 14.733 | 0.000 | 0.000 | 5.08273E-12 | 9.91494E-13 | 2.96607E-12 | 7.19939E-12 |
| | *N*-oleoyl methionine | Equal variances assumed | 0.133 | 0.720 | 6.842 | 17 | 0.000 | 0.000 | 6.28240E-11 | 9.18260E-12 | 4.34504E-11 | 8.21976E-11 |
| | | Equal variances not assumed | | | 6.818 | 16.504 | 0.000 | 0.000 | 6.28240E-11 | 9.21434E-12 | 4.33389E-11 | 8.23091E-11 |
| | *N*-linoleoyl methionine | Equal variances assumed | 11.562 | 0.003 | 6.238 | 17 | 0.000 | 0.000 | 7.68052E-11 | 1.23117E-11 | 5.08298E-11 | 1.02781E-10 |
| | | Equal variances not assumed | | | 6.035 | 11.558 | 0.000 | 0.000 | 7.68052E-11 | 1.27263E-11 | 4.89588E-11 | 1.04652E-10 |
| | *N*-arachidonoyl methionine | Equal variances assumed | 1.804 | 0.196 | 3.906 | 18 | 0.001 | 0.001 | 3.53846E-11 | 9.05795E-12 | 1.63545E-11 | 5.44146E-11 |
| | | Equal variances not assumed | | | 4.041 | 17.749 | 0.000 | 0.001 | 3.53846E-11 | 8.75568E-12 | 1.69709E-11 | 5.37983E-11 |
| | *N*-docosahexaenoyl methionine | Equal variances assumed | 2.164 | 0.159 | 2.011 | 18 | 0.030 | 0.059 | 1.96829E-11 | 9.78532E-12 | -8.75341E-13 | 4.02411E-11 |
| | | Equal variances not assumed | | | 2.090 | 17.528 | 0.026 | 0.051 | 1.96829E-11 | 9.41690E-12 | -1.39530E-13 | 3.95052E-11 |

Supplemental Table 16.3 One-way ANOVA for *N*-acyl methionines measured in liver collected from adult mice 2h after an i.p. injection of 10mg/kg CBD, THC, or THC+CBD

| | | Sum of Squares | df | Mean Square | F | Sig. |
|---|---|---|---|---|---|---|
| *N*-palmitoyl methionine | Between Groups | 3.08E-20 | 7 | 4.40E-21 | 7.506 | 0.00000 |
| | Within Groups | 3.63E-20 | 62 | 5.86E-22 | | |
| | Total | 6.71E-20 | 69 | | | |
| *N*-stearoyl methionine | Between Groups | 2.57E-22 | 7 | 3.67E-23 | 4.926 | 0.00025 |
| | Within Groups | 3.88E-22 | 52 | 7.46E-24 | | |
| | Total | 6.45E-22 | 59 | | | |
| *N*-oleoyl methionine | Between Groups | 4.30E-20 | 7 | 6.14E-21 | 10.508 | 0.00000 |
| | Within Groups | 3.56E-20 | 61 | 5.84E-22 | | |
| | Total | 7.86E-20 | 68 | | | |
| *N*-linoleoyl methionine | Between Groups | 4.49E-20 | 7 | 6.41E-21 | 10.413 | 0.00000 |
| | Within Groups | 3.69E-20 | 60 | 6.16E-22 | | |
| | Total | 8.18E-20 | 67 | | | |
| *N*-arachidonoyl methionine | Between Groups | 1.16E-20 | 7 | 1.66E-21 | 3.972 | 0.00118 |
| | Within Groups | 2.63E-20 | 63 | 4.18E-22 | | |
| | Total | 3.79E-20 | 70 | | | |
| *N*-docosahexaenoyl methionine | Between Groups | 7.22E-21 | 7 | 1.03E-21 | 3.871 | 0.00151 |
| | Within Groups | 1.63E-20 | 61 | 2.67E-22 | | |
| | Total | 2.35E-20 | 68 | | | |

#### Supplemental Table 16.4 Fisher's LSD post-hoc test for liver *N*-acyl methionines ANOVA

| Dependent Variable | | | Std. Error | Sig. | 95% Confidence Interval Lower Bound | 95% Confidence Interval Upper Bound |
|---|---|---|---|---|---|---|
| | VehFemale | CBDfemale | 1.25251E-11 | 0.683 | -3.0183E-11 | 1.9892E-11 |
| | | THCfemale | 1.14794E-11 | 0.185 | -3.8348E-11 | 7.5462E-12 |
| | | TpCfemale | 1.17595E-11 | 0.003 | -5.9337E-11 | -1.2323E-11 |
| | | VehMale | 1.25251E-11 | 0.131 | -5.8567E-12 | 4.4218E-11 |
| | | CBDmale | 1.14794E-11 | 0.046 | 4.3749E-13 | 4.6332E-11 |
| | | THCmale | 1.17595E-11 | 0.171 | -3.9797E-11 | 7.2162E-12 |
| | | TpCmale | 1.14794E-11 | 0.033 | 2.0579E-12 | 4.7952E-11 |
| | CBDfemale | VehFemale | 1.25251E-11 | 0.683 | -1.9892E-11 | 3.0183E-11 |
| | | THCfemale | 1.19263E-11 | 0.393 | -3.4096E-11 | 1.3585E-11 |
| | | TpCfemale | 1.21960E-11 | 0.014 | -5.5064E-11 | -6.3048E-12 |
| | | VehMale | 1.29358E-11 | 0.065 | -1.5323E-12 | 5.0184E-11 |
| | | CBDmale | 1.19263E-11 | 0.020 | 4.6898E-12 | 5.2370E-11 |
| | | THCmale | 1.21960E-11 | 0.364 | -3.5525E-11 | 1.3234E-11 |
| | | TpCmale | 1.19263E-11 | 0.014 | 6.3102E-12 | 5.3991E-11 |
| | THCfemale | VehFemale | 1.14794E-11 | 0.185 | -7.5462E-12 | 3.8348E-11 |
| | | CBDfemale | 1.19263E-11 | 0.393 | -1.3585E-11 | 3.4096E-11 |
| | | TpCfemale | 1.11195E-11 | 0.071 | -4.2657E-11 | 1.7984E-12 |
| | | VehMale | 1.19263E-11 | 0.005 | 1.0741E-11 | 5.8422E-11 |
| | | CBDmale | 1.08229E-11 | 0.001 | 1.7151E-11 | 6.0420E-11 |
| | | THCmale | 1.11195E-11 | 0.936 | -2.3117E-11 | 2.1338E-11 |
| | | TpCmale | 1.08229E-11 | 0.000 | 1.8771E-11 | 6.2040E-11 |
| | TpCfemale | VehFemale | 1.17595E-11 | 0.003 | 1.2323E-11 | 5.9337E-11 |
| | | CBDfemale | 1.21960E-11 | 0.014 | 6.3048E-12 | 5.5064E-11 |
| | | THCfemale | 1.11195E-11 | 0.071 | -1.7984E-12 | 4.2657E-11 |
| | | VehMale | 1.21960E-11 | 0.000 | 3.0631E-11 | 7.9390E-11 |
| | | CBDmale | 1.11195E-11 | 0.000 | 3.6987E-11 | 8.1442E-11 |
| | | THCmale | 1.14083E-11 | 0.092 | -3.2657E-12 | 4.2344E-11 |
| | | TpCmale | 1.11195E-11 | 0.000 | 3.8607E-11 | 8.3062E-11 |
| *N*-palmitoyl methionine | VehMale | VehFemale | 1.25251E-11 | 0.131 | -4.4218E-11 | 5.8567E-12 |
| | | CBDfemale | 1.29358E-11 | 0.065 | -5.0184E-11 | 1.5323E-12 |
| | | THCfemale | 1.19263E-11 | 0.005 | -5.8422E-11 | -1.0741E-11 |
| | | TpCfemale | 1.21960E-11 | 0.000 | -7.9390E-11 | -3.0631E-11 |
| | | CBDmale | 1.19263E-11 | 0.726 | -1.9636E-11 | 2.8044E-11 |
| | | THCmale | 1.21960E-11 | 0.005 | -5.9851E-11 | -1.1092E-11 |
| | | TpCmale | 1.19263E-11 | 0.627 | -1.8016E-11 | 2.9665E-11 |
| | CBDmale | VehFemale | 1.14794E-11 | 0.046 | -4.6332E-11 | -4.3749E-13 |
| | | CBDfemale | 1.19263E-11 | 0.020 | -5.2370E-11 | -4.6898E-12 |
| | | THCfemale | 1.08229E-11 | 0.001 | -6.0420E-11 | -1.7151E-11 |
| | | TpCfemale | 1.11195E-11 | 0.000 | -8.1442E-11 | -3.6987E-11 |
| | | VehMale | 1.19263E-11 | 0.726 | -2.8044E-11 | 1.9636E-11 |
| | | THCmale | 1.11195E-11 | 0.001 | -6.1903E-11 | -1.7448E-11 |
| | | TpCmale | 1.08229E-11 | 0.881 | -2.0014E-11 | 2.3255E-11 |
| | THCmale | VehFemale | 1.17595E-11 | 0.171 | -7.2162E-12 | 3.9797E-11 |
| | | CBDfemale | 1.21960E-11 | 0.364 | -1.3234E-11 | 3.5525E-11 |
| | | THCfemale | 1.11195E-11 | 0.936 | -2.1338E-11 | 2.3117E-11 |
| | | TpCfemale | 1.14083E-11 | 0.092 | -4.2344E-11 | 3.2657E-12 |
| | | VehMale | 1.21960E-11 | 0.005 | 1.1092E-11 | 5.9851E-11 |
| | | CBDmale | 1.11195E-11 | 0.001 | 1.7448E-11 | 6.1903E-11 |
| | | TpCmale | 1.11195E-11 | 0.000 | 1.9068E-11 | 6.3523E-11 |
| | TpCmale | VehFemale | 1.14794E-11 | 0.033 | -4.7952E-11 | -2.0579E-12 |
| | | CBDfemale | 1.19263E-11 | 0.014 | -5.3991E-11 | -6.3102E-12 |
| | | THCfemale | 1.08229E-11 | 0.000 | -6.2040E-11 | -1.8771E-11 |
| | | TpCfemale | 1.11195E-11 | 0.000 | -8.3062E-11 | -3.8607E-11 |
| | | VehMale | 1.19263E-11 | 0.627 | -2.9665E-11 | 1.8016E-11 |
| | | CBDmale | 1.08229E-11 | 0.881 | -2.3255E-11 | 2.0014E-11 |
| | | THCmale | 1.11195E-11 | 0.000 | -6.3523E-11 | -1.9068E-11 |
| *N*-stearoyl methionine | VehFemale | CBDfemale | 1.47482E-12 | 0.001 | -8.0158E-12 | -2.0969E-12 |

| | | | | | | |
|---|---|---|---|---|---|---|
| | | THCfemale | 1.29535E-12 | 0.527 | -3.4238E-12 | 1.7748E-12 |
| | | TpCfemale | 1.29535E-12 | 0.007 | -6.2259E-12 | -1.0273E-12 |
| | | VehMale | 1.41334E-12 | 0.745 | -2.3741E-12 | 3.2981E-12 |
| | | CBDmale | 1.67229E-12 | 0.505 | -2.2328E-12 | 4.4786E-12 |
| | | THCmale | 1.41334E-12 | 0.468 | -3.8696E-12 | 1.8026E-12 |
| | | TpCmale | 1.36542E-12 | 0.291 | -1.2838E-12 | 4.1960E-12 |
| | CBDfemale | VehFemale | 1.47482E-12 | 0.001 | 2.0969E-12 | 8.0158E-12 |
| | | THCfemale | 1.41020E-12 | 0.004 | 1.4021E-12 | 7.0617E-12 |
| | | TpCfemale | 1.41020E-12 | 0.315 | -1.4000E-12 | 4.2595E-12 |
| | | VehMale | 1.51930E-12 | 0.001 | 2.4697E-12 | 8.5671E-12 |
| | | CBDmale | 1.76275E-12 | 0.001 | 2.6421E-12 | 9.7165E-12 |
| | | THCmale | 1.51930E-12 | 0.011 | 9.7420E-13 | 7.0716E-12 |
| | | TpCmale | 1.47482E-12 | 0.000 | 3.5530E-12 | 9.4719E-12 |
| | THCfemale | VehFemale | 1.29535E-12 | 0.527 | -1.7748E-12 | 3.4238E-12 |
| | | CBDfemale | 1.41020E-12 | 0.004 | -7.0617E-12 | -1.4021E-12 |
| | | TpCfemale | 1.22127E-12 | 0.026 | -5.2528E-12 | -3.5147E-13 |
| | | VehMale | 1.34577E-12 | 0.344 | -1.4140E-12 | 3.9870E-12 |
| | | CBDmale | 1.61559E-12 | 0.234 | -1.2945E-12 | 5.1893E-12 |
| | | THCmale | 1.34577E-12 | 0.877 | -2.9095E-12 | 2.4915E-12 |
| | | TpCmale | 1.29535E-12 | 0.084 | -3.1870E-13 | 4.8799E-12 |
| | TpCfemale | VehFemale | 1.29535E-12 | 0.007 | 1.0273E-12 | 6.2259E-12 |
| | | CBDfemale | 1.41020E-12 | 0.315 | -4.2595E-12 | 1.4000E-12 |
| | | THCfemale | 1.22127E-12 | 0.026 | 3.5147E-13 | 5.2528E-12 |
| | | VehMale | 1.34577E-12 | 0.004 | 1.3882E-12 | 6.7891E-12 |
| | | CBDmale | 1.61559E-12 | 0.005 | 1.5076E-12 | 7.9915E-12 |
| | | THCmale | 1.34577E-12 | 0.059 | -1.0736E-13 | 5.2936E-12 |
| | | TpCmale | 1.29535E-12 | 0.000 | 2.4834E-12 | 7.6820E-12 |
| | VehMale | VehFemale | 1.41334E-12 | 0.745 | -3.2981E-12 | 2.3741E-12 |
| | | CBDfemale | 1.51930E-12 | 0.001 | -8.5671E-12 | -2.4697E-12 |
| | | THCfemale | 1.34577E-12 | 0.344 | -3.9870E-12 | 1.4140E-12 |
| | | TpCfemale | 1.34577E-12 | 0.004 | -6.7891E-12 | -1.3882E-12 |
| | | CBDmale | 1.71164E-12 | 0.701 | -2.7738E-12 | 4.0956E-12 |
| | | THCmale | 1.45970E-12 | 0.310 | -4.4246E-12 | 1.4336E-12 |
| | | TpCmale | 1.41334E-12 | 0.485 | -1.8420E-12 | 3.8302E-12 |
| | CBDmale | VehFemale | 1.67229E-12 | 0.505 | -4.4786E-12 | 2.2328E-12 |
| | | CBDfemale | 1.76275E-12 | 0.001 | -9.7165E-12 | -2.6421E-12 |
| | | THCfemale | 1.61559E-12 | 0.234 | -5.1893E-12 | 1.2945E-12 |
| | | TpCfemale | 1.61559E-12 | 0.005 | -7.9915E-12 | -1.5076E-12 |
| | | VehMale | 1.71164E-12 | 0.701 | -4.0956E-12 | 2.7738E-12 |
| | | THCmale | 1.71164E-12 | 0.213 | -5.5911E-12 | 1.2782E-12 |
| | | TpCmale | 1.67229E-12 | 0.843 | -3.0225E-12 | 3.6889E-12 |
| | THCmale | VehFemale | 1.41334E-12 | 0.468 | -1.8026E-12 | 3.8696E-12 |
| | | CBDfemale | 1.51930E-12 | 0.011 | -7.0716E-12 | -9.7420E-13 |
| | | THCfemale | 1.34577E-12 | 0.877 | -2.4915E-12 | 2.9095E-12 |
| | | TpCfemale | 1.34577E-12 | 0.059 | -5.2936E-12 | 1.0736E-13 |
| | | VehMale | 1.45970E-12 | 0.310 | -1.4336E-12 | 4.4246E-12 |
| | | CBDmale | 1.71164E-12 | 0.213 | -1.2782E-12 | 5.5911E-12 |
| | | TpCmale | 1.41334E-12 | 0.084 | -3.4649E-13 | 5.3257E-12 |
| | TpCmale | VehFemale | 1.36542E-12 | 0.291 | -4.1960E-12 | 1.2838E-12 |
| | | CBDfemale | 1.47482E-12 | 0.000 | -9.4719E-12 | -3.5530E-12 |
| | | THCfemale | 1.29535E-12 | 0.084 | -4.8799E-12 | 3.1870E-13 |
| | | TpCfemale | 1.29535E-12 | 0.000 | -7.6820E-12 | -2.4834E-12 |
| | | VehMale | 1.41334E-12 | 0.485 | -3.8302E-12 | 1.8420E-12 |
| | | CBDmale | 1.67229E-12 | 0.843 | -3.6889E-12 | 3.0225E-12 |
| | | THCmale | 1.41334E-12 | 0.084 | -5.3257E-12 | 3.4649E-13 |
| *N*-oleoyl methionine | VehFemale | CBDfemale | 1.25088E-11 | 0.928 | -2.3885E-11 | 2.6141E-11 |
| | | THCfemale | 1.14645E-11 | 0.001 | -6.3670E-11 | -1.7821E-11 |
| | | TpCfemale | 1.17441E-11 | 0.000 | -7.4057E-11 | -2.7089E-11 |
| | | VehMale | 1.25088E-11 | 0.153 | -6.9106E-12 | 4.3115E-11 |
| | | CBDmale | 1.14645E-11 | 0.157 | -6.5033E-12 | 3.9346E-11 |

| | | | | | | |
|---|---|---|---|---|---|---|
| | | THCmale | 1.20846E-11 | 0.685 | -2.9083E-11 | 1.9246E-11 |
| | | TpCmale | 1.14645E-11 | 0.289 | -1.0674E-11 | 3.5176E-11 |
| | CBDfemale | VehFemale | 1.25088E-11 | 0.928 | -2.6141E-11 | 2.3885E-11 |
| | | THCfemale | 1.19107E-11 | 0.001 | -6.5691E-11 | -1.8057E-11 |
| | | TpCfemale | 1.21801E-11 | 0.000 | -7.6057E-11 | -2.7346E-11 |
| | | VehMale | 1.29190E-11 | 0.194 | -8.8592E-12 | 4.2807E-11 |
| | | CBDmale | 1.19107E-11 | 0.204 | -8.5240E-12 | 3.9110E-11 |
| | | THCmale | 1.25088E-11 | 0.631 | -3.1060E-11 | 1.8966E-11 |
| | | TpCmale | 1.19107E-11 | 0.354 | -1.2694E-11 | 3.4940E-11 |
| | THCfemale | VehFemale | 1.14645E-11 | 0.001 | 1.7821E-11 | 6.3670E-11 |
| | | CBDfemale | 1.19107E-11 | 0.001 | 1.8057E-11 | 6.5691E-11 |
| | | TpCfemale | 1.11050E-11 | 0.380 | -3.2033E-11 | 1.2378E-11 |
| | | VehMale | 1.19107E-11 | 0.000 | 3.5031E-11 | 8.2665E-11 |
| | | CBDmale | 1.08088E-11 | 0.000 | 3.5553E-11 | 7.8780E-11 |
| | | THCmale | 1.14645E-11 | 0.003 | 1.2903E-11 | 5.8752E-11 |
| | | TpCmale | 1.08088E-11 | 0.000 | 3.1383E-11 | 7.4610E-11 |
| | TpCfemale | VehFemale | 1.17441E-11 | 0.000 | 2.7089E-11 | 7.4057E-11 |
| | | CBDfemale | 1.21801E-11 | 0.000 | 2.7346E-11 | 7.6057E-11 |
| | | THCfemale | 1.11050E-11 | 0.380 | -1.2378E-11 | 3.2033E-11 |
| | | VehMale | 1.21801E-11 | 0.000 | 4.4319E-11 | 9.3031E-11 |
| | | CBDmale | 1.11050E-11 | 0.000 | 4.4788E-11 | 8.9200E-11 |
| | | THCmale | 1.17441E-11 | 0.000 | 2.2171E-11 | 6.9138E-11 |
| | | TpCmale | 1.11050E-11 | 0.000 | 4.0618E-11 | 8.5030E-11 |
| | VehMale | VehFemale | 1.25088E-11 | 0.153 | -4.3115E-11 | 6.9106E-12 |
| | | CBDfemale | 1.29190E-11 | 0.194 | -4.2807E-11 | 8.8592E-12 |
| | | THCfemale | 1.19107E-11 | 0.000 | -8.2665E-11 | -3.5031E-11 |
| | | TpCfemale | 1.21801E-11 | 0.000 | -9.3031E-11 | -4.4319E-11 |
| | | CBDmale | 1.19107E-11 | 0.888 | -2.5498E-11 | 2.2136E-11 |
| | | THCmale | 1.25088E-11 | 0.071 | -4.8033E-11 | 1.9922E-12 |
| | | TpCmale | 1.19107E-11 | 0.625 | -2.9668E-11 | 1.7966E-11 |
| | CBDmale | VehFemale | 1.14645E-11 | 0.157 | -3.9346E-11 | 6.5033E-12 |
| | | CBDfemale | 1.19107E-11 | 0.204 | -3.9110E-11 | 8.5240E-12 |
| | | THCfemale | 1.08088E-11 | 0.000 | -7.8780E-11 | -3.5553E-11 |
| | | TpCfemale | 1.11050E-11 | 0.000 | -8.9200E-11 | -4.4788E-11 |
| | | VehMale | 1.19107E-11 | 0.888 | -2.2136E-11 | 2.5498E-11 |
| | | THCmale | 1.14645E-11 | 0.068 | -4.4264E-11 | 1.5849E-12 |
| | | TpCmale | 1.08088E-11 | 0.701 | -2.5784E-11 | 1.7443E-11 |
| | THCmale | VehFemale | 1.20846E-11 | 0.685 | -1.9246E-11 | 2.9083E-11 |
| | | CBDfemale | 1.25088E-11 | 0.631 | -1.8966E-11 | 3.1060E-11 |
| | | THCfemale | 1.14645E-11 | 0.003 | -5.8752E-11 | -1.2903E-11 |
| | | TpCfemale | 1.17441E-11 | 0.000 | -6.9138E-11 | -2.2171E-11 |
| | | VehMale | 1.25088E-11 | 0.071 | -1.9922E-12 | 4.8033E-11 |
| | | CBDmale | 1.14645E-11 | 0.068 | -1.5849E-12 | 4.4264E-11 |
| | | TpCmale | 1.14645E-11 | 0.139 | -5.7551E-12 | 4.0094E-11 |
| | TpCmale | VehFemale | 1.14645E-11 | 0.289 | -3.5176E-11 | 1.0674E-11 |
| | | CBDfemale | 1.19107E-11 | 0.354 | -3.4940E-11 | 1.2694E-11 |
| | | THCfemale | 1.08088E-11 | 0.000 | -7.4610E-11 | -3.1383E-11 |
| | | TpCfemale | 1.11050E-11 | 0.000 | -8.5030E-11 | -4.0618E-11 |
| | | VehMale | 1.19107E-11 | 0.625 | -1.7966E-11 | 2.9668E-11 |
| | | CBDmale | 1.08088E-11 | 0.701 | -1.7443E-11 | 2.5784E-11 |
| | | THCmale | 1.14645E-11 | 0.139 | -4.0094E-11 | 5.7551E-12 |
| *N*-linoleoyl methionine | VehFemale | CBDfemale | 1.28409E-11 | 0.265 | -4.0126E-11 | 1.1245E-11 |
| | | THCfemale | 1.20560E-11 | 0.358 | -3.5291E-11 | 1.2941E-11 |
| | | TpCfemale | 1.20560E-11 | 0.000 | -8.7309E-11 | -3.9078E-11 |
| | | VehMale | 1.28409E-11 | 0.140 | -6.5006E-12 | 4.4871E-11 |
| | | CBDmale | 1.20560E-11 | 0.140 | -6.1053E-12 | 4.2126E-11 |
| | | THCmale | 1.20560E-11 | 0.172 | -4.0781E-11 | 7.4498E-12 |
| | | TpCmale | 1.17689E-11 | 0.252 | -9.9293E-12 | 3.7153E-11 |
| | CBDfemale | VehFemale | 1.28409E-11 | 0.265 | -1.1245E-11 | 4.0126E-11 |
| | | THCfemale | 1.25036E-11 | 0.795 | -2.1746E-11 | 2.8276E-11 |

| | | | | | | |
|---|---|---|---|---|---|---|
| | | TpCfemale | 1.25036E-11 | 0.000 | -7.3764E-11 | -2.3742E-11 |
| | | VehMale | 1.32621E-11 | 0.014 | 7.0973E-12 | 6.0153E-11 |
| | | CBDmale | 1.25036E-11 | 0.012 | 7.4397E-12 | 5.7461E-11 |
| | | THCmale | 1.25036E-11 | 0.859 | -2.7236E-11 | 2.2785E-11 |
| | | TpCmale | 1.22270E-11 | 0.025 | 3.5947E-12 | 5.2510E-11 |
| | THCfemale | VehFemale | 1.20560E-11 | 0.358 | -1.2941E-11 | 3.5291E-11 |
| | | CBDfemale | 1.25036E-11 | 0.795 | -2.8276E-11 | 2.1746E-11 |
| | | TpCfemale | 1.16960E-11 | 0.000 | -7.5414E-11 | -2.8622E-11 |
| | | VehMale | 1.25036E-11 | 0.018 | 5.3493E-12 | 5.5371E-11 |
| | | CBDmale | 1.16960E-11 | 0.015 | 5.7898E-12 | 5.2581E-11 |
| | | THCmale | 1.16960E-11 | 0.640 | -2.8886E-11 | 1.7905E-11 |
| | | TpCmale | 1.13999E-11 | 0.034 | 1.9840E-12 | 4.7590E-11 |
| | TpCfemale | VehFemale | 1.20560E-11 | 0.000 | 3.9078E-11 | 8.7309E-11 |
| | | CBDfemale | 1.25036E-11 | 0.000 | 2.3742E-11 | 7.3764E-11 |
| | | THCfemale | 1.16960E-11 | 0.000 | 2.8622E-11 | 7.5414E-11 |
| | | VehMale | 1.25036E-11 | 0.000 | 5.7367E-11 | 1.0739E-10 |
| | | CBDmale | 1.16960E-11 | 0.000 | 5.7808E-11 | 1.0460E-10 |
| | | THCmale | 1.16960E-11 | 0.000 | 2.3132E-11 | 6.9923E-11 |
| | | TpCmale | 1.13999E-11 | 0.000 | 5.4002E-11 | 9.9608E-11 |
| | VehMale | VehFemale | 1.28409E-11 | 0.140 | -4.4871E-11 | 6.5006E-12 |
| | | CBDfemale | 1.32621E-11 | 0.014 | -6.0153E-11 | -7.0973E-12 |
| | | THCfemale | 1.25036E-11 | 0.018 | -5.5371E-11 | -5.3493E-12 |
| | | TpCfemale | 1.25036E-11 | 0.000 | -1.0739E-10 | -5.7367E-11 |
| | | CBDmale | 1.25036E-11 | 0.925 | -2.6186E-11 | 2.3836E-11 |
| | | THCmale | 1.25036E-11 | 0.006 | -6.0862E-11 | -1.0840E-11 |
| | | TpCmale | 1.22270E-11 | 0.650 | -3.0031E-11 | 1.8885E-11 |
| | CBDmale | VehFemale | 1.20560E-11 | 0.140 | -4.2126E-11 | 6.1053E-12 |
| | | CBDfemale | 1.25036E-11 | 0.012 | -5.7461E-11 | -7.4397E-12 |
| | | THCfemale | 1.16960E-11 | 0.015 | -5.2581E-11 | -5.7898E-12 |
| | | TpCfemale | 1.16960E-11 | 0.000 | -1.0460E-10 | -5.7808E-11 |
| | | VehMale | 1.25036E-11 | 0.925 | -2.3836E-11 | 2.6186E-11 |
| | | THCmale | 1.16960E-11 | 0.004 | -5.8072E-11 | -1.1281E-11 |
| | | TpCmale | 1.13999E-11 | 0.701 | -2.7201E-11 | 1.8405E-11 |
| | THCmale | VehFemale | 1.20560E-11 | 0.172 | -7.4498E-12 | 4.0781E-11 |
| | | CBDfemale | 1.25036E-11 | 0.859 | -2.2785E-11 | 2.7236E-11 |
| | | THCfemale | 1.16960E-11 | 0.640 | -1.7905E-11 | 2.8886E-11 |
| | | TpCfemale | 1.16960E-11 | 0.000 | -6.9923E-11 | -2.3132E-11 |
| | | VehMale | 1.25036E-11 | 0.006 | 1.0840E-11 | 6.0862E-11 |
| | | CBDmale | 1.16960E-11 | 0.004 | 1.1281E-11 | 5.8072E-11 |
| | | TpCmale | 1.13999E-11 | 0.010 | 7.4747E-12 | 5.3081E-11 |
| | TpCmale | VehFemale | 1.17689E-11 | 0.252 | -3.7153E-11 | 9.9293E-12 |
| | | CBDfemale | 1.22270E-11 | 0.025 | -5.2510E-11 | -3.5947E-12 |
| | | THCfemale | 1.13999E-11 | 0.034 | -4.7590E-11 | -1.9840E-12 |
| | | TpCfemale | 1.13999E-11 | 0.000 | -9.9608E-11 | -5.4002E-11 |
| | | VehMale | 1.22270E-11 | 0.650 | -1.8885E-11 | 3.0031E-11 |
| | | CBDmale | 1.13999E-11 | 0.701 | -1.8405E-11 | 2.7201E-11 |
| | | THCmale | 1.13999E-11 | 0.010 | -5.3081E-11 | -7.4747E-12 |
| *N*-arachidonoyl methionine | VehFemale | CBDfemale | 1.05788E-11 | 0.321 | -3.1717E-11 | 1.0563E-11 |
| | | THCfemale | 9.69567E-12 | 0.251 | -3.0619E-11 | 8.1314E-12 |
| | | TpCfemale | 9.93218E-12 | 0.012 | -4.5521E-11 | -5.8254E-12 |
| | | VehMale | 1.05788E-11 | 0.358 | -1.1353E-11 | 3.0927E-11 |
| | | CBDmale | 9.69567E-12 | 0.183 | -6.3276E-12 | 3.2423E-11 |
| | | THCmale | 9.93218E-12 | 0.359 | -2.9030E-11 | 1.0666E-11 |
| | | TpCmale | 9.49777E-12 | 0.310 | -9.2685E-12 | 2.8691E-11 |
| | CBDfemale | VehFemale | 1.05788E-11 | 0.321 | -1.0563E-11 | 3.1717E-11 |
| | | THCfemale | 1.00731E-11 | 0.947 | -2.0796E-11 | 1.9463E-11 |
| | | TpCfemale | 1.03009E-11 | 0.148 | -3.5681E-11 | 5.4884E-12 |
| | | VehMale | 1.09258E-11 | 0.067 | -1.4693E-12 | 4.2198E-11 |
| | | CBDmale | 1.00731E-11 | 0.022 | 3.4952E-12 | 4.3754E-11 |
| | | THCmale | 1.03009E-11 | 0.893 | -1.9190E-11 | 2.1980E-11 |

| | | | | | | |
|---|---|---|---|---|---|---|
| | | TpCmale | 9.88274E-12 | 0.044 | 5.3911E-13 | 4.0037E-11 |
| | THCfemale | VehFemale | 9.69567E-12 | 0.251 | -8.1314E-12 | 3.0619E-11 |
| | | CBDfemale | 1.00731E-11 | 0.947 | -1.9463E-11 | 2.0796E-11 |
| | | TpCfemale | 9.39165E-12 | 0.129 | -3.3197E-11 | 4.3382E-12 |
| | | VehMale | 1.00731E-11 | 0.041 | 9.0152E-13 | 4.1160E-11 |
| | | CBDmale | 9.14116E-12 | 0.010 | 6.0243E-12 | 4.2559E-11 |
| | | THCmale | 9.39165E-12 | 0.827 | -1.6706E-11 | 2.0829E-11 |
| | | TpCmale | 8.93099E-12 | 0.022 | 3.1079E-12 | 3.8802E-11 |
| | TpCfemale | VehFemale | 9.93218E-12 | 0.012 | 5.8254E-12 | 4.5521E-11 |
| | | CBDfemale | 1.03009E-11 | 0.148 | -5.4884E-12 | 3.5681E-11 |
| | | THCfemale | 9.39165E-12 | 0.129 | -4.3382E-12 | 3.3197E-11 |
| | | VehMale | 1.03009E-11 | 0.001 | 1.4876E-11 | 5.6045E-11 |
| | | CBDmale | 9.39165E-12 | 0.000 | 1.9953E-11 | 5.7489E-11 |
| | | THCmale | 9.63563E-12 | 0.092 | -2.7640E-12 | 3.5747E-11 |
| | | TpCmale | 9.18721E-12 | 0.000 | 1.7025E-11 | 5.3744E-11 |
| | VehMale | VehFemale | 1.05788E-11 | 0.358 | -3.0927E-11 | 1.1353E-11 |
| | | CBDfemale | 1.09258E-11 | 0.067 | -4.2198E-11 | 1.4693E-12 |
| | | THCfemale | 1.00731E-11 | 0.041 | -4.1160E-11 | -9.0152E-13 |
| | | TpCfemale | 1.03009E-11 | 0.001 | -5.6045E-11 | -1.4876E-11 |
| | | CBDmale | 1.00731E-11 | 0.747 | -1.6869E-11 | 2.3390E-11 |
| | | THCmale | 1.03009E-11 | 0.070 | -3.9554E-11 | 1.6155E-12 |
| | | TpCmale | 9.88274E-12 | 0.994 | -1.9825E-11 | 1.9673E-11 |
| | CBDmale | VehFemale | 9.69567E-12 | 0.183 | -3.2423E-11 | 6.3276E-12 |
| | | CBDfemale | 1.00731E-11 | 0.022 | -4.3754E-11 | -3.4952E-12 |
| | | THCfemale | 9.14116E-12 | 0.010 | -4.2559E-11 | -6.0243E-12 |
| | | TpCfemale | 9.39165E-12 | 0.000 | -5.7489E-11 | -1.9953E-11 |
| | | VehMale | 1.00731E-11 | 0.747 | -2.3390E-11 | 1.6869E-11 |
| | | THCmale | 9.39165E-12 | 0.021 | -4.0997E-11 | -3.4620E-12 |
| | | TpCmale | 8.93099E-12 | 0.710 | -2.1184E-11 | 1.4511E-11 |
| | THCmale | VehFemale | 9.93218E-12 | 0.359 | -1.0666E-11 | 2.9030E-11 |
| | | CBDfemale | 1.03009E-11 | 0.893 | -2.1980E-11 | 1.9190E-11 |
| | | THCfemale | 9.39165E-12 | 0.827 | -2.0829E-11 | 1.6706E-11 |
| | | TpCfemale | 9.63563E-12 | 0.092 | -3.5747E-11 | 2.7640E-12 |
| | | VehMale | 1.03009E-11 | 0.070 | -1.6155E-12 | 3.9554E-11 |
| | | CBDmale | 9.39165E-12 | 0.021 | 3.4620E-12 | 4.0997E-11 |
| | | TpCmale | 9.18721E-12 | 0.044 | 5.3413E-13 | 3.7252E-11 |
| | TpCmale | VehFemale | 9.49777E-12 | 0.310 | -2.8691E-11 | 9.2685E-12 |
| | | CBDfemale | 9.88274E-12 | 0.044 | -4.0037E-11 | -5.3911E-13 |
| | | THCfemale | 8.93099E-12 | 0.022 | -3.8802E-11 | -3.1079E-12 |
| | | TpCfemale | 9.18721E-12 | 0.000 | -5.3744E-11 | -1.7025E-11 |
| | | VehMale | 9.88274E-12 | 0.994 | -1.9673E-11 | 1.9825E-11 |
| | | CBDmale | 8.93099E-12 | 0.710 | -1.4511E-11 | 2.1184E-11 |
| | | THCmale | 9.18721E-12 | 0.044 | -3.7252E-11 | -5.3413E-13 |
| *N*-docosahexaenoyl methionine | VehFemale | CBDfemale | 8.44961E-12 | 0.494 | -2.2712E-11 | 1.1080E-11 |
| | | THCfemale | 7.93310E-12 | 0.331 | -2.3635E-11 | 8.0911E-12 |
| | | TpCfemale | 7.93310E-12 | 0.004 | -3.9685E-11 | -7.9584E-12 |
| | | VehMale | 8.44961E-12 | 0.268 | -7.4528E-12 | 2.6339E-11 |
| | | CBDmale | 7.93310E-12 | 0.253 | -6.7066E-12 | 2.5020E-11 |
| | | THCmale | 7.93310E-12 | 0.124 | -2.8248E-11 | 3.4788E-12 |
| | | TpCmale | 7.58613E-12 | 0.587 | -1.9308E-11 | 1.1031E-11 |
| | CBDfemale | VehFemale | 8.44961E-12 | 0.494 | -1.1080E-11 | 2.2712E-11 |
| | | THCfemale | 8.22763E-12 | 0.813 | -1.8409E-11 | 1.4496E-11 |
| | | TpCfemale | 8.22763E-12 | 0.032 | -3.4458E-11 | -1.5539E-12 |
| | | VehMale | 8.72672E-12 | 0.085 | -2.1913E-12 | 3.2709E-11 |
| | | CBDmale | 8.22763E-12 | 0.074 | -1.4799E-12 | 3.1424E-11 |
| | | THCmale | 8.22763E-12 | 0.428 | -2.3021E-11 | 9.8833E-12 |
| | | TpCmale | 7.89361E-12 | 0.832 | -1.4107E-11 | 1.7461E-11 |
| | THCfemale | VehFemale | 7.93310E-12 | 0.331 | -8.0911E-12 | 2.3635E-11 |
| | | CBDfemale | 8.22763E-12 | 0.813 | -1.4496E-11 | 1.8409E-11 |
| | | TpCfemale | 7.69624E-12 | 0.041 | -3.1439E-11 | -6.5997E-13 |

| | | | | | |
|---|---|---|---|---|---|
| | VehMale | 8.22763E-12 | 0.041 | 7.6315E-13 | 3.3667E-11 |
| | CBDmale | 7.69624E-12 | 0.032 | 1.5392E-12 | 3.2318E-11 |
| | THCmale | 7.69624E-12 | 0.551 | -2.0002E-11 | 1.0777E-11 |
| | TpCmale | 7.33808E-12 | 0.622 | -1.1040E-11 | 1.8307E-11 |
| TpCfemale | VehFemale | 7.93310E-12 | 0.004 | 7.9584E-12 | 3.9685E-11 |
| | CBDfemale | 8.22763E-12 | 0.032 | 1.5539E-12 | 3.4458E-11 |
| | THCfemale | 7.69624E-12 | 0.041 | 6.5997E-13 | 3.1439E-11 |
| | VehMale | 8.22763E-12 | 0.000 | 1.6813E-11 | 4.9717E-11 |
| | CBDmale | 7.69624E-12 | 0.000 | 1.7589E-11 | 4.8368E-11 |
| | THCmale | 7.69624E-12 | 0.142 | -3.9523E-12 | 2.6827E-11 |
| | TpCmale | 7.33808E-12 | 0.009 | 5.0095E-12 | 3.4356E-11 |
| VehMale | VehFemale | 8.44961E-12 | 0.268 | -2.6339E-11 | 7.4528E-12 |
| | CBDfemale | 8.72672E-12 | 0.085 | -3.2709E-11 | 2.1913E-12 |
| | THCfemale | 8.22763E-12 | 0.041 | -3.3667E-11 | -7.6315E-13 |
| | TpCfemale | 8.22763E-12 | 0.000 | -4.9717E-11 | -1.6813E-11 |
| | CBDmale | 8.22763E-12 | 0.972 | -1.6739E-11 | 1.6166E-11 |
| | THCmale | 8.22763E-12 | 0.010 | -3.8280E-11 | -5.3755E-12 |
| | TpCmale | 7.89361E-12 | 0.090 | -2.9366E-11 | 2.2022E-12 |
| CBDmale | VehFemale | 7.93310E-12 | 0.253 | -2.5020E-11 | 6.7066E-12 |
| | CBDfemale | 8.22763E-12 | 0.074 | -3.1424E-11 | 1.4799E-12 |
| | THCfemale | 7.69624E-12 | 0.032 | -3.2318E-11 | -1.5392E-12 |
| | TpCfemale | 7.69624E-12 | 0.000 | -4.8368E-11 | -1.7589E-11 |
| | VehMale | 8.22763E-12 | 0.972 | -1.6166E-11 | 1.6739E-11 |
| | THCmale | 7.69624E-12 | 0.007 | -3.6931E-11 | -6.1515E-12 |
| | TpCmale | 7.33808E-12 | 0.075 | -2.7969E-11 | 1.3779E-12 |
| THCmale | VehFemale | 7.93310E-12 | 0.124 | -3.4788E-12 | 2.8248E-11 |
| | CBDfemale | 8.22763E-12 | 0.428 | -9.8833E-12 | 2.3021E-11 |
| | THCfemale | 7.69624E-12 | 0.551 | -1.0777E-11 | 2.0002E-11 |
| | TpCfemale | 7.69624E-12 | 0.142 | -2.6827E-11 | 3.9523E-12 |
| | VehMale | 8.22763E-12 | 0.010 | 5.3755E-12 | 3.8280E-11 |
| | CBDmale | 7.69624E-12 | 0.007 | 6.1515E-12 | 3.6931E-11 |
| | TpCmale | 7.33808E-12 | 0.266 | -6.4278E-12 | 2.2919E-11 |
| TpCmale | VehFemale | 7.58613E-12 | 0.587 | -1.1031E-11 | 1.9308E-11 |
| | CBDfemale | 7.89361E-12 | 0.832 | -1.7461E-11 | 1.4107E-11 |
| | THCfemale | 7.33808E-12 | 0.622 | -1.8307E-11 | 1.1040E-11 |
| | TpCfemale | 7.33808E-12 | 0.009 | -3.4356E-11 | -5.0095E-12 |
| | VehMale | 7.89361E-12 | 0.090 | -2.2022E-12 | 2.9366E-11 |
| | CBDmale | 7.33808E-12 | 0.075 | -1.3779E-12 | 2.7969E-11 |
| | THCmale | 7.33808E-12 | 0.266 | -2.2919E-11 | 6.4278E-12 |

Supplemental Table 17.1 Descriptive statistics for *N*-acyl tryptophans (mol/L) in liver collected from male and female adult mice 2h after receiving an i.p. injection of 10mg/kg CBD, THC, or THC+CBD

| | | N | Mean | Std. Deviation | Std. Error | 95% Confidence Interval for Mean | | Minimum | Maximum |
|---|---|---|---|---|---|---|---|---|---|
| | | | | | | Lower Bound | Upper Bound | | |
| *N*-palmitoyl tryptophan | Veh F | BAL | | | | | | | |
| | CBD F | BAL | | | | | | | |
| | THC F | 4 | 2.1327E-12 | 9.10766E-14 | 4.55383E-14 | 1.9878E-12 | 2.2776E-12 | 2.05E-12 | 2.26E-12 |
| | THC+CBD F | BAL | | | | | | | |
| | Veh M | BAL | | | | | | | |
| | CBD M | BAL | | | | | | | |
| | THC M | BAL | | | | | | | |
| | THC+CBD M | BAL | | | | | | | |
| *N*-stearoyl tryptophan | Veh F | 4 | 4.4552E-12 | 1.34156E-12 | 6.70780E-13 | 2.3205E-12 | 6.5899E-12 | 3.32E-12 | 6.37E-12 |
| | CBD F | BDL | | | | | | | |
| | THC F | BDL | | | | | | | |
| | THC+CBD F | BAL | | | | | | | |
| | Veh M | BAL | | | | | | | |
| | CBD M | BAL | | | | | | | |
| | THC M | BAL | | | | | | | |
| | THC+CBD M | BAL | | | | | | | |
| *N*-oleoyl tryptophan | Veh F | BDL | | | | | | | |
| | CBD F | BAL | | | | | | | |
| | THC F | BDL | | | | | | | |
| | THC+CBD F | BAL | | | | | | | |
| | Veh M | BDL | | | | | | | |
| | CBD M | BDL | | | | | | | |
| | THC M | BDL | | | | | | | |
| | THC+CBD M | BDL | | | | | | | |
| *N*-linoleoyl tryptophan | Veh F | BAL | | | | | | | |
| | CBD F | 4 | 3.6253E-10 | 1.72652E-10 | 8.63261E-11 | 8.7802E-11 | 6.3726E-10 | 2.00E-10 | 5.98E-10 |
| | THC F | 4 | 2.9478E-10 | 1.24240E-10 | 6.21202E-11 | 9.7082E-11 | 4.9247E-10 | 1.68E-10 | 4.66E-10 |
| | THC+CBD F | 5 | 3.4607E-10 | 1.84679E-10 | 8.25909E-11 | 1.1676E-10 | 5.7538E-10 | 1.70E-10 | 6.37E-10 |
| | Veh M | BDL | | | | | | | |
| | CBD M | 5 | 2.4587E-10 | 6.09889E-11 | 2.72751E-11 | 1.7015E-10 | 3.2160E-10 | 1.83E-10 | 3.13E-10 |
| | THC M | BAL | | | | | | | |
| | THC+CBD M | 4 | 1.8537E-10 | 8.35193E-11 | 4.17597E-11 | 5.2473E-11 | 3.1827E-10 | 9.77E-11 | 2.87E-10 |
| *N*-arachidonoyl tryptophan | Veh F | BDL | | | | | | | |
| | CBD F | BDL | | | | | | | |

| | | |
|---|---|---|
| | THC F | BDL |
| | THC+CBD F | BDL |
| | Veh M | BDL |
| | CBD M | BDL |
| | THC M | BDL |
| | THC+CBD M | BDL |
| *N*-docosahexaenoyl tryptophan | Veh F | BDL |
| | CBD F | BDL |
| | THC F | BDL |
| | THC+CBD F | BAL |
| | Veh M | BDL |
| | CBD M | BDL |
| | THC M | BDL |
| | THC+CBD M | BDL |

Supplemental Table 17.2 Independent t-test comparing male and female levels of *N*-acyl tryptophans measured in liver collected from adult mice 2h after an i.p. injection of 10mg/kg CBD, THC, or THC+CBD

| | | | Levene's Test for Equality of Variances | | t-test for Equality of Means | | | | | | | |
|---|---|---|---|---|---|---|---|---|---|---|---|---|
| | | | | | | | Significance | | | | 95% Confidence Interval of the Difference | |
| | | | F | Sig. | t | df | One-Sided p | Two-Sided p | Mean Difference | Std. Error Difference | Lower | Upper |
| CBD | *N*-linoleoyl tryptophan | Equal variances assumed | 2.738 | 0.142 | 1.425 | 7 | 0.099 | 0.197 | 1.16656E-10 | 8.18860E-11 | -7.69734E-11 | 3.10286E-10 |
| | | Equal variances not assumed | | | 1.289 | 3.602 | 0.137 | 0.274 | 1.16656E-10 | 9.05325E-11 | -1.46046E-10 | 3.79358E-10 |
| THC+CBD | *N*-linoleoyl tryptophan | Equal variances assumed | 2.455 | 0.161 | 1.598 | 7 | 0.077 | 0.154 | 1.60699E-10 | 1.00576E-10 | -7.71245E-11 | 3.98523E-10 |
| | | Equal variances not assumed | | | 1.736 | 5.801 | 0.067 | 0.135 | 1.60699E-10 | 9.25480E-11 | -6.76530E-11 | 3.89051E-10 |

Supplemental Table 17.3 One-way ANOVA for *N*-acyl tryptophans measured in liver collected from adult mice 2h after an i.p. injection of 10mg/kg CBD, THC, or THC+CBD

| | | Sum of Squares | df | Mean Square | F | Sig. |
|---|---|---|---|---|---|---|
| *N*-linoleoyl tryptophan | Between Groups | 9.03E-20 | 4 | 2.26E-20 | 1.246 | 0.32922 |
| | Within Groups | 3.08E-19 | 17 | 1.81E-20 | | |
| | Total | 3.98E-19 | 21 | | | |

#### Supplemental Table 17.4 Fisher's LSD post-hoc test for liver *N*-acyl tryptophans ANOVA

| | (I) drugVsexLtyrosine | | Std. Error | Sig. | 95% Confidence Interval Lower Bound | Upper Bound |
|---|---|---|---|---|---|---|
| *N*-linoleoyl tryptophan | CBDFemale | THCfemale | 9.51722E-11 | 0.486 | -1.3304E-10 | 2.6855E-10 |
| | | TpCfemale | 9.02883E-11 | 0.857 | -1.7403E-10 | 2.0695E-10 |
| | | CBDmale | 9.02883E-11 | 0.214 | -7.3835E-11 | 3.0715E-10 |
| | | TpCmale | 9.51722E-11 | 0.080 | -2.3636E-11 | 3.7796E-10 |
| | THCFemale | CBDfemale | 9.51722E-11 | 0.486 | -2.6855E-10 | 1.3304E-10 |
| | | TpCfemale | 9.02883E-11 | 0.577 | -2.4179E-10 | 1.3920E-10 |
| | | CBDmale | 9.02883E-11 | 0.595 | -1.4159E-10 | 2.3939E-10 |
| | | TpCmale | 9.51722E-11 | 0.266 | -9.1391E-11 | 3.1020E-10 |
| | TpCFemale | CBDfemale | 9.02883E-11 | 0.857 | -2.0695E-10 | 1.7403E-10 |
| | | THCfemale | 9.02883E-11 | 0.577 | -1.3920E-10 | 2.4179E-10 |
| | | CBDmale | 8.51246E-11 | 0.255 | -7.9401E-11 | 2.7979E-10 |
| | | TpCmale | 9.02883E-11 | 0.093 | -2.9793E-11 | 3.5119E-10 |
| | CBDMale | CBDfemale | 9.02883E-11 | 0.214 | -3.0715E-10 | 7.3835E-11 |
| | | THCfemale | 9.02883E-11 | 0.595 | -2.3939E-10 | 1.4159E-10 |
| | | TpCfemale | 8.51246E-11 | 0.255 | -2.7979E-10 | 7.9401E-11 |
| | | TpCmale | 9.02883E-11 | 0.512 | -1.2999E-10 | 2.5100E-10 |
| | TpCMale | CBDfemale | 9.51722E-11 | 0.080 | -3.7796E-10 | 2.3636E-11 |
| | | THCfemale | 9.51722E-11 | 0.266 | -3.1020E-10 | 9.1391E-11 |
| | | TpCfemale | 9.02883E-11 | 0.093 | -3.5119E-10 | 2.9793E-11 |
| | | CBDmale | 9.02883E-11 | 0.512 | -2.5100E-10 | 1.2999E-10 |

Supplemental Table 18.1 Descriptive statistics for prostaglandins (mol/g) in liver collected from male and female adult mice 2h after receiving an i.p. injection of 10mg/kg CBD, THC, or THC+CBD

| | | N | Mean | Std. Deviation | Std. Error | 95% Confidence Interval for Mean | | Minimum | Maximum |
|---|---|---|---|---|---|---|---|---|---|
| | | | | | | Lower Bound | Upper Bound | | |
| $PGE_2$ | Veh F | 8 | 1.7492E-10 | 7.96230E-11 | 2.81510E-11 | 1.0835E-10 | 2.4149E-10 | 6.01E-11 | 2.71E-10 |
| | CBD F | 8 | 3.3165E-10 | 1.16305E-10 | 4.11201E-11 | 2.3441E-10 | 4.2888E-10 | 1.63E-10 | 5.29E-10 |
| | THC F | 10 | 4.4081E-10 | 1.91061E-10 | 6.04189E-11 | 3.0413E-10 | 5.7749E-10 | 2.30E-10 | 7.81E-10 |
| | THC+CBD F | 9 | 3.4855E-10 | 9.30058E-11 | 3.10019E-11 | 2.7706E-10 | 4.2004E-10 | 2.17E-10 | 5.02E-10 |
| | Veh M | 7 | 1.6524E-10 | 9.90316E-11 | 3.74304E-11 | 7.3653E-11 | 2.5683E-10 | 6.60E-11 | 2.85E-10 |
| | CBD M | 10 | 2.4956E-10 | 9.76879E-11 | 3.08916E-11 | 1.7968E-10 | 3.1944E-10 | 1.03E-10 | 4.27E-10 |
| | THC M | 9 | 3.3768E-10 | 1.12449E-10 | 3.74831E-11 | 2.5125E-10 | 4.2412E-10 | 1.67E-10 | 4.66E-10 |
| | THC+CBD M | 10 | 3.0450E-10 | 9.54596E-11 | 3.01870E-11 | 2.3621E-10 | 3.7279E-10 | 1.53E-10 | 4.64E-10 |
| PGF2α | Veh F | 8 | 8.0552E-10 | 3.93691E-10 | 1.39191E-10 | 4.7638E-10 | 1.1347E-09 | 1.42E-10 | 1.34E-09 |
| | CBD F | 8 | 1.3923E-09 | 3.17575E-10 | 1.12280E-10 | 1.1268E-09 | 1.6578E-09 | 8.62E-10 | 1.82E-09 |
| | THC F | 10 | 1.3899E-09 | 5.72419E-10 | 1.81015E-10 | 9.8043E-10 | 1.7994E-09 | 7.50E-10 | 2.24E-09 |
| | THC+CBD F | 9 | 1.0035E-09 | 3.27778E-10 | 1.09259E-10 | 7.5156E-10 | 1.2555E-09 | 6.94E-10 | 1.52E-09 |
| | Veh M | 6 | 1.0702E-09 | 6.39857E-10 | 2.61220E-10 | 3.9872E-10 | 1.7417E-09 | 5.70E-10 | 1.89E-09 |
| | CBD M | 10 | 1.1575E-09 | 5.21393E-10 | 1.64879E-10 | 7.8455E-10 | 1.5305E-09 | 5.33E-10 | 2.08E-09 |
| | THC M | 8 | 1.3480E-09 | 1.73797E-10 | 6.14466E-11 | 1.2028E-09 | 1.4933E-09 | 1.12E-09 | 1.57E-09 |
| | THC+CBD M | 10 | 8.9989E-10 | 3.59749E-10 | 1.13763E-10 | 6.4254E-10 | 1.1572E-09 | 3.69E-10 | 1.44E-09 |
| 6-keto-$PGF_{1\alpha}$ | Veh F | 5 | 1.2296E-09 | 5.32582E-10 | 2.38178E-10 | 5.6828E-10 | 1.8909E-09 | 8.11E-10 | 2.10E-09 |
| | CBD F | 5 | 1.8042E-09 | 1.08753E-09 | 4.86359E-10 | 4.5389E-10 | 3.1546E-09 | 6.73E-10 | 3.23E-09 |
| | THC F | 8 | 3.0712E-09 | 2.49235E-09 | 8.81177E-10 | 9.8759E-10 | 5.1549E-09 | 7.54E-10 | 8.06E-09 |
| | THC+CBD F | 7 | 2.0596E-09 | 9.03011E-10 | 3.41306E-10 | 1.2245E-09 | 2.8948E-09 | 9.01E-10 | 2.96E-09 |
| | Veh M | BDL | | | | | | | |
| | CBD M | BDL | | | | | | | |
| | THC M | BAL | 1.2946E-09 | | | | | 1.29E-09 | 1.29E-09 |
| | THC+CBD M | 6 | 3.3578E-09 | 3.75908E-09 | 1.53464E-09 | -5.8712E-10 | 7.3027E-09 | 3.80E-10 | 8.39E-09 |

Supplemental Table 18.2 Independent t-test comparing male and female levels of prostaglandins measured in liver collected from adult mice 2h after an i.p. injection of 10mg/kg CBD, THC, or THC+CBD

| | | | Levene's Test for Equality of Variances | | t-test for Equality of Means | | | | | | | |
|---|---|---|---|---|---|---|---|---|---|---|---|---|
| | | | | | | | Significance | | | | 95% Confidence Interval of the Difference | |
| | | | F | Sig. | t | df | One-Sided p | Two-Sided p | Mean Difference | Std. Error Difference | Lower | Upper |
| Veh | $PGE_2$ | Equal variances assumed | 2.225 | 0.160 | 0.210 | 13 | 0.419 | 0.837 | 9.67694E-12 | 4.61176E-11 | -8.99541E-11 | 1.09308E-10 |
| | | Equal variances not assumed | | | 0.207 | 11.542 | 0.420 | 0.840 | 9.67694E-12 | 4.68350E-11 | -9.28186E-11 | 1.12172E-10 |
| | PGF2α | Equal variances assumed | 3.685 | 0.079 | -0.959 | 12 | 0.178 | 0.356 | -2.64691E-10 | 2.75909E-10 | -8.65845E-10 | 3.36463E-10 |
| | | Equal variances not assumed | | | -0.894 | 7.794 | 0.199 | 0.398 | -2.64691E-10 | 2.95990E-10 | -9.50405E-10 | 4.21023E-10 |
| CBD | $PGE_2$ | Equal variances assumed | 0.127 | 0.726 | 1.629 | 16 | 0.061 | 0.123 | 8.20856E-11 | 5.03918E-11 | -2.47401E-11 | 1.88911E-10 |
| | | Equal variances not assumed | | | 1.596 | 13.730 | 0.067 | 0.133 | 8.20856E-11 | 5.14311E-11 | -2.84272E-11 | 1.92598E-10 |
| | PGF2α | Equal variances assumed | 1.929 | 0.184 | 1.115 | 16 | 0.141 | 0.281 | 2.34793E-10 | 2.10556E-10 | -2.11566E-10 | 6.81152E-10 |
| | | Equal variances not assumed | | | 1.177 | 15.106 | 0.129 | 0.257 | 2.34793E-10 | 1.99479E-10 | -1.90126E-10 | 6.59712E-10 |
| THC | $PGE_2$ | Equal variances assumed | 3.737 | 0.070 | 1.412 | 17 | 0.088 | 0.176 | 1.03126E-10 | 7.30488E-11 | -5.09939E-11 | 2.57245E-10 |
| | | Equal variances not assumed | | | 1.450 | 14.795 | 0.084 | 0.168 | 1.03126E-10 | 7.11015E-11 | -4.86064E-11 | 2.54858E-10 |
| | PGF2α | Equal variances assumed | 21.870 | 0.000 | 0.199 | 16 | 0.423 | 0.845 | 4.18632E-11 | 2.10816E-10 | -4.05046E-10 | 4.88773E-10 |
| | | Equal variances not assumed | | | 0.219 | 11.006 | 0.415 | 0.831 | 4.18632E-11 | 1.91160E-10 | -3.78850E-10 | 4.62576E-10 |
| | 6-keto-PGF1alpha | Equal variances assumed | | | 0.672 | 7 | 0.262 | 0.523 | 1.77665E-09 | 2.64353E-09 | -4.47430E-09 | 8.02761E-09 |
| | | Equal variances not assumed | | | | | | | 1.77665E-09 | | | |

| | | | | | | | | | | | |
|---|---|---|---|---|---|---|---|---|---|---|---|
| | $PGE_2$ | Equal variances assumed | 0.008 | 0.930 | 1.016 | 17 | 0.162 | 0.324 | 4.40468E-11 | 4.33338E-11 | -4.73794E-11 | 1.35473E-10 |
| | | Equal variances not assumed | | | 1.018 | 16.876 | 0.162 | 0.323 | 4.40468E-11 | 4.32709E-11 | -4.72979E-11 | 1.35392E-10 |
| | PGF2α | Equal variances assumed | 0.066 | 0.800 | 0.654 | 17 | 0.261 | 0.522 | 1.03621E-10 | 1.58550E-10 | -2.30890E-10 | 4.38133E-10 |
| THC+CBD | | Equal variances not assumed | | | 0.657 | 16.994 | 0.260 | 0.520 | 1.03621E-10 | 1.57732E-10 | -2.29173E-10 | 4.36416E-10 |
| | 6-keto-PGF1alpha | Equal variances assumed | 21.671 | 0.001 | -0.890 | 11 | 0.196 | 0.392 | -1.29814E-09 | 1.45799E-09 | -4.50717E-09 | 1.91088E-09 |
| | | Equal variances not assumed | | | -0.826 | 5.496 | 0.222 | 0.443 | -1.29814E-09 | 1.57213E-09 | -5.23223E-09 | 2.63595E-09 |

Supplemental Table 18.3 One-way ANOVA for prostaglandins measured in liver collected from adult mice 2h after an i.p. injection of 10mg/kg CBD, THC, or THC+CBD

| | | Sum of Squares | df | Mean Square | F | Sig. |
|---|---|---|---|---|---|---|
| $PGE_2$ | Between Groups | 0.000 | 7 | 0.000 | 5.393 | 0.00007 |
| | Within Groups | 0.000 | 63 | 0.000 | | |
| | Total | 0.000 | 70 | | | |
| PGF2α | Between Groups | 0.000 | 7 | 0.000 | 2.393 | 0.03132 |
| | Within Groups | 0.000 | 61 | 0.000 | | |
| | Total | 0.000 | 68 | | | |
| 6-keto-PGF1α | Between Groups | 0.000 | 3 | 0.000 | 1.507 | 0.24192 |
| | Within Groups | 0.000 | 21 | 0.000 | | |
| | Total | 0.000 | 24 | | | |

#### Supplemental Table 18.4 Fisher's LSD post-hoc test for liver prostaglandins ANOVA

| Dependent Variable | | | Std. Error | Sig. | 95% Confidence Interval Lower Bound | 95% Confidence Interval Upper Bound |
|---|---|---|---|---|---|---|
| PGE2 | VehFemale | CBDfemale | 5.86E-11 | 9.50E-03 | -2.7378E-10 | -3.9667E-11 |
| | | THCfemale | 5.56E-11 | 1.07E-05 | -3.7694E-10 | -1.5484E-10 |
| | | TpCfemale | 5.69E-11 | 3.35E-03 | -2.8739E-10 | -5.9866E-11 |
| | | VehMale | 6.06E-11 | 8.74E-01 | -1.1149E-10 | 1.3084E-10 |
| | | CBDmale | 5.56E-11 | 1.84E-01 | -1.8569E-10 | 3.6411E-11 |
| | | THCmale | 5.69E-11 | 5.75E-03 | -2.7652E-10 | -4.9002E-11 |
| | | TpCmale | 5.56E-11 | 2.29E-02 | -2.4063E-10 | -1.8528E-11 |
| | CBDfemale | VehFemale | 5.86E-11 | 9.50E-03 | 3.9667E-11 | 2.7378E-10 |
| | | THCfemale | 5.56E-11 | 5.39E-02 | -2.2021E-10 | 1.8890E-12 |
| | | TpCfemale | 5.69E-11 | 7.68E-01 | -1.3066E-10 | 9.6860E-11 |
| | | VehMale | 6.06E-11 | 7.89E-03 | 4.5235E-11 | 2.8757E-10 |
| | | CBDmale | 5.56E-11 | 1.45E-01 | -2.8966E-11 | 1.9314E-10 |
| | | THCmale | 5.69E-11 | 9.16E-01 | -1.1980E-10 | 1.0772E-10 |
| | | TpCmale | 5.56E-11 | 6.27E-01 | -8.3906E-11 | 1.3820E-10 |
| | THCfemale | VehFemale | 5.56E-11 | 1.07E-05 | 1.5484E-10 | 3.7694E-10 |
| | | CBDfemale | 5.56E-11 | 5.39E-02 | -1.8890E-12 | 2.2021E-10 |
| | | TpCfemale | 5.38E-11 | 9.15E-02 | -1.5308E-11 | 1.9983E-10 |
| | | VehMale | 5.77E-11 | 1.12E-05 | 1.6019E-10 | 3.9094E-10 |
| | | CBDmale | 5.24E-11 | 5.34E-04 | 8.6548E-11 | 2.9595E-10 |
| | | THCmale | 5.38E-11 | 5.99E-02 | -4.4440E-12 | 2.1070E-10 |
| | | TpCmale | 5.24E-11 | 1.16E-02 | 3.1608E-11 | 2.4101E-10 |
| | TpCfemale | VehFemale | 5.69E-11 | 3.35E-03 | 5.9866E-11 | 2.8739E-10 |
| | | CBDfemale | 5.69E-11 | 7.68E-01 | -9.6860E-11 | 1.3066E-10 |
| | | THCfemale | 5.38E-11 | 9.15E-02 | -1.9983E-10 | 1.5308E-11 |
| | | VehMale | 5.90E-11 | 2.85E-03 | 6.5320E-11 | 3.0129E-10 |
| | | CBDmale | 5.38E-11 | 7.06E-02 | -8.5831E-12 | 2.0656E-10 |
| | | THCmale | 5.52E-11 | 8.45E-01 | -9.9500E-11 | 1.2123E-10 |
| | | TpCmale | 5.38E-11 | 4.16E-01 | -6.3523E-11 | 1.5162E-10 |
| | VehMale | VehFemale | 6.06E-11 | 8.74E-01 | -1.3084E-10 | 1.1149E-10 |
| | | CBDfemale | 6.06E-11 | 7.89E-03 | -2.8757E-10 | -4.5235E-11 |
| | | THCfemale | 5.77E-11 | 1.12E-05 | -3.9094E-10 | -1.6019E-10 |
| | | TpCfemale | 5.90E-11 | 2.85E-03 | -3.0129E-10 | -6.5320E-11 |
| | | CBDmale | 5.77E-11 | 1.49E-01 | -1.9969E-10 | 3.1057E-11 |
| | | THCmale | 5.90E-11 | 4.84E-03 | -2.9042E-10 | -5.4456E-11 |
| | | TpCmale | 5.77E-11 | 1.88E-02 | -2.5463E-10 | -2.3883E-11 |
| | CBDmale | VehFemale | 5.56E-11 | 1.84E-01 | -3.6411E-11 | 1.8569E-10 |
| | | CBDfemale | 5.56E-11 | 1.45E-01 | -1.9314E-10 | 2.8966E-11 |
| | | THCfemale | 5.24E-11 | 5.34E-04 | -2.9595E-10 | -8.6548E-11 |
| | | TpCfemale | 5.38E-11 | 7.06E-02 | -2.0656E-10 | 8.5831E-12 |
| | | VehMale | 5.77E-11 | 1.49E-01 | -3.1057E-11 | 1.9969E-10 |
| | | THCmale | 5.38E-11 | 1.07E-01 | -1.9569E-10 | 1.9447E-11 |
| | | TpCmale | 5.24E-11 | 2.98E-01 | -1.5964E-10 | 4.9761E-11 |
| | THCmale | VehFemale | 5.69E-11 | 5.75E-03 | 4.9002E-11 | 2.7652E-10 |
| | | CBDfemale | 5.69E-11 | 9.16E-01 | -1.0772E-10 | 1.1980E-10 |
| | | THCfemale | 5.38E-11 | 5.99E-02 | -2.1070E-10 | 4.4440E-12 |
| | | TpCfemale | 5.52E-11 | 8.45E-01 | -1.2123E-10 | 9.9500E-11 |
| | | VehMale | 5.90E-11 | 4.84E-03 | 5.4456E-11 | 2.9042E-10 |
| | | CBDmale | 5.38E-11 | 1.07E-01 | -1.9447E-11 | 1.9569E-10 |
| | | TpCmale | 5.38E-11 | 5.40E-01 | -7.4387E-11 | 1.4075E-10 |
| | TpCmale | VehFemale | 5.56E-11 | 2.29E-02 | 1.8528E-11 | 2.4063E-10 |
| | | CBDfemale | 5.56E-11 | 6.27E-01 | -1.3820E-10 | 8.3906E-11 |
| | | THCfemale | 5.24E-11 | 1.16E-02 | -2.4101E-10 | -3.1608E-11 |
| | | TpCfemale | 5.38E-11 | 4.16E-01 | -1.5162E-10 | 6.3523E-11 |
| | | VehMale | 5.77E-11 | 1.88E-02 | 2.3883E-11 | 2.5463E-10 |
| | | CBDmale | 5.24E-11 | 2.98E-01 | -4.9761E-11 | 1.5964E-10 |

| | | | | | | |
|---|---|---|---|---|---|---|
| | | THCmale | 5.38E-11 | 5.40E-01 | -1.4075E-10 | 7.4387E-11 |
| PGF2α | VehFemale | CBDfemale | 2.17E-10 | 8.80E-03 | -1.0203E-09 | -1.5328E-10 |
| | | THCfemale | 2.06E-10 | 6.10E-03 | -9.9568E-10 | -1.7311E-10 |
| | | TpCfemale | 2.11E-10 | 3.51E-01 | -6.1931E-10 | 2.2332E-10 |
| | | VehMale | 2.34E-10 | 2.63E-01 | -7.3296E-10 | 2.0358E-10 |
| | | CBDmale | 2.06E-10 | 9.21E-02 | -7.6331E-10 | 5.9267E-11 |
| | | THCmale | 2.17E-10 | 1.50E-02 | -9.7607E-10 | -1.0900E-10 |
| | | TpCmale | 2.06E-10 | 6.48E-01 | -5.0566E-10 | 3.1691E-10 |
| | CBDfemale | VehFemale | 2.17E-10 | 8.80E-03 | 1.5328E-10 | 1.0203E-09 |
| | | THCfemale | 2.06E-10 | 9.91E-01 | -4.0887E-10 | 4.1370E-10 |
| | | TpCfemale | 2.11E-10 | 6.98E-02 | -3.2503E-11 | 8.1014E-10 |
| | | VehMale | 2.34E-10 | 1.74E-01 | -1.4615E-10 | 7.9039E-10 |
| | | CBDmale | 2.06E-10 | 2.58E-01 | -1.7649E-10 | 6.4608E-10 |
| | | THCmale | 2.17E-10 | 8.39E-01 | -3.8925E-10 | 4.7781E-10 |
| | | TpCmale | 2.06E-10 | 1.97E-02 | 8.1152E-11 | 9.0372E-10 |
| | THCfemale | VehFemale | 2.06E-10 | 6.10E-03 | 1.7311E-10 | 9.9568E-10 |
| | | CBDfemale | 2.06E-10 | 9.91E-01 | -4.1370E-10 | 4.0887E-10 |
| | | TpCfemale | 1.99E-10 | 5.71E-02 | -1.1990E-11 | 7.8479E-10 |
| | | VehMale | 2.24E-10 | 1.58E-01 | -1.2805E-10 | 7.6746E-10 |
| | | CBDmale | 1.94E-10 | 2.35E-01 | -1.5539E-10 | 6.2014E-10 |
| | | THCmale | 2.06E-10 | 8.39E-01 | -3.6942E-10 | 4.5315E-10 |
| | | TpCmale | 1.94E-10 | 1.41E-02 | 1.0226E-10 | 8.7779E-10 |
| | TpCfemale | VehFemale | 2.11E-10 | 3.51E-01 | -2.2332E-10 | 6.1931E-10 |
| | | CBDfemale | 2.11E-10 | 6.98E-02 | -8.1014E-10 | 3.2503E-11 |
| | | THCfemale | 1.99E-10 | 5.71E-02 | -7.8479E-10 | 1.1990E-11 |
| | | VehMale | 2.29E-10 | 7.71E-01 | -5.2368E-10 | 3.9029E-10 |
| | | CBDmale | 1.99E-10 | 4.42E-01 | -5.5241E-10 | 2.4437E-10 |
| | | THCmale | 2.11E-10 | 1.07E-01 | -7.6586E-10 | 7.6782E-11 |
| | | TpCmale | 1.99E-10 | 6.05E-01 | -2.9477E-10 | 5.0201E-10 |
| | VehMale | VehFemale | 2.34E-10 | 2.63E-01 | -2.0358E-10 | 7.3296E-10 |
| | | CBDfemale | 2.34E-10 | 1.74E-01 | -7.9039E-10 | 1.4615E-10 |
| | | THCfemale | 2.24E-10 | 1.58E-01 | -7.6746E-10 | 1.2805E-10 |
| | | TpCfemale | 2.29E-10 | 7.71E-01 | -3.9029E-10 | 5.2368E-10 |
| | | CBDmale | 2.24E-10 | 6.98E-01 | -5.3508E-10 | 3.6042E-10 |
| | | THCmale | 2.34E-10 | 2.40E-01 | -7.4611E-10 | 1.9043E-10 |
| | | TpCmale | 2.24E-10 | 4.50E-01 | -2.7744E-10 | 6.1807E-10 |
| | CBDmale | VehFemale | 2.06E-10 | 9.21E-02 | -5.9267E-11 | 7.6331E-10 |
| | | CBDfemale | 2.06E-10 | 2.58E-01 | -6.4608E-10 | 1.7649E-10 |
| | | THCfemale | 1.94E-10 | 2.35E-01 | -6.2014E-10 | 1.5539E-10 |
| | | TpCfemale | 1.99E-10 | 4.42E-01 | -2.4437E-10 | 5.5241E-10 |
| | | VehMale | 2.24E-10 | 6.98E-01 | -3.6042E-10 | 5.3508E-10 |
| | | THCmale | 2.06E-10 | 3.58E-01 | -6.0180E-10 | 2.2077E-10 |
| | | TpCmale | 1.94E-10 | 1.89E-01 | -1.3012E-10 | 6.4541E-10 |
| | THCmale | VehFemale | 2.17E-10 | 1.50E-02 | 1.0900E-10 | 9.7607E-10 |
| | | CBDfemale | 2.17E-10 | 8.39E-01 | -4.7781E-10 | 3.8925E-10 |
| | | THCfemale | 2.06E-10 | 8.39E-01 | -4.5315E-10 | 3.6942E-10 |
| | | TpCfemale | 2.11E-10 | 1.07E-01 | -7.6782E-11 | 7.6586E-10 |
| | | VehMale | 2.34E-10 | 2.40E-01 | -1.9043E-10 | 7.4611E-10 |
| | | CBDmale | 2.06E-10 | 3.58E-01 | -2.2077E-10 | 6.0180E-10 |
| | | TpCmale | 2.06E-10 | 3.32E-02 | 3.6872E-11 | 8.5944E-10 |
| | TpCmale | VehFemale | 2.06E-10 | 6.48E-01 | -3.1691E-10 | 5.0566E-10 |
| | | CBDfemale | 2.06E-10 | 1.97E-02 | -9.0372E-10 | -8.1152E-11 |
| | | THCfemale | 1.94E-10 | 1.41E-02 | -8.7779E-10 | -1.0226E-10 |
| | | TpCfemale | 1.99E-10 | 6.05E-01 | -5.0201E-10 | 2.9477E-10 |
| | | VehMale | 2.24E-10 | 4.50E-01 | -6.1807E-10 | 2.7744E-10 |
| | | CBDmale | 1.94E-10 | 1.89E-01 | -6.4541E-10 | 1.3012E-10 |
| | | THCmale | 2.06E-10 | 3.32E-02 | -8.5944E-10 | -3.6872E-11 |
| 6-keto-PGF1α | VehFemale | CBDfemale | 1.02E-09 | 5.78E-01 | -2.6885E-09 | 1.5391E-09 |
| | | THCfemale | 9.16E-10 | 5.74E-02 | -3.7470E-09 | 6.3683E-11 |

| | | | | | |
|---|---|---|---|---|---|
| | TpCfemale | 9.41E-10 | 3.88E-01 | -2.7871E-09 | 1.1269E-09 |
| CBDfemale | VehFemale | 1.02E-09 | 5.78E-01 | -1.5391E-09 | 2.6885E-09 |
| | THCfemale | 9.16E-10 | 1.81E-01 | -3.1724E-09 | 6.3835E-10 |
| | TpCfemale | 9.41E-10 | 7.89E-01 | -2.2124E-09 | 1.7016E-09 |
| THCfemale | VehFemale | 9.16E-10 | 5.74E-02 | -6.3683E-11 | 3.7470E-09 |
| | CBDfemale | 9.16E-10 | 1.81E-01 | -6.3835E-10 | 3.1724E-09 |
| | TpCfemale | 8.32E-10 | 2.37E-01 | -7.1816E-10 | 2.7414E-09 |
| TpCfemale | VehFemale | 9.41E-10 | 3.88E-01 | -1.1269E-09 | 2.7871E-09 |
| | CBDfemale | 9.41E-10 | 7.89E-01 | -1.7016E-09 | 2.2124E-09 |
| | THCfemale | 8.32E-10 | 2.37E-01 | -2.7414E-09 | 7.1816E-10 |
